# Unravelling neuronal and glial differences in ceramide composition, synthesis, and sensitivity to toxicity

**DOI:** 10.1101/2023.12.06.569570

**Authors:** John J. McInnis, Disha Sood, Lilu Guo, Michael R. Dufault, Mariana Garcia, Rachel Passaro, Grace Gao, Bailin Zhang, James C. Dodge

## Abstract

Ceramides are lipids that play vital roles in complex lipid synthesis, membrane function, and cell signaling. Disrupted ceramide homeostasis is implicated in cell-death and several neurologic diseases. Ceramides are often analyzed in tissue, but this approach fails to resolve cell-type differences in ceramide homeostasis that are likely essential to understanding cell and non-cell autonomous contributions to neurodegeneration. We show that human iPSC-derived neurons and glia differ in their rate of ceramide synthesis, ceramide isoform composition, and responses to altered ceramide levels. RNA-sequencing of cells treated to increase or decrease ceramides revealed connections to inflammation, ER stress, and apoptosis. Moreover, introducing labeled sphinganine showed that glia readily synthesize ceramide de novo and that neurons are relatively more sensitive to ceramide toxicity. Our findings provide a framework for understanding neurologic diseases with sphingolipid alternations and insights in to designing therapeutics that target ceramide for treating them.

## Introduction

Ceramides are sphingolipids that perform both structural and signaling functions within the cell. Structurally, ceramides can regulate membrane fluidity and curvature under normal conditions and contribute to membrane pore formation during cellular stress (Kaltenegger et al., 2021; Pinto et al., 2011). Ceramide is generated through two different pathways: *de novo* synthesis, and through its salvage following the hydrolysis of complex lipids such as sphingomyelin, glucosylceramide, and galatosylceramide (Kitatani et al., 2008)(Figure 1a). The *de novo* synthesis pathway is initiated by the rate limiting enzyme complex serine palmitoyltransferase (SPT), followed by the synthesis of dihydroceramide by six ceramide synthases (CerS1-6). Dihydroceramide is then desaturated by DEGS to form ceramide (Merrill, 1983; Mizutani et al., 2005; Pant et al., 2019; Pewzner-Jung et al., 2006; Planas-Serra et al., 2023; Ruangsiriluk et al., 2012; Ternes et al., 2002; Venkataraman et al., 2002). Each CerS preferentially generates ceramides with specific fatty acid chain lengths, conferring different properties to each ceramide species.

**Figure 1.**
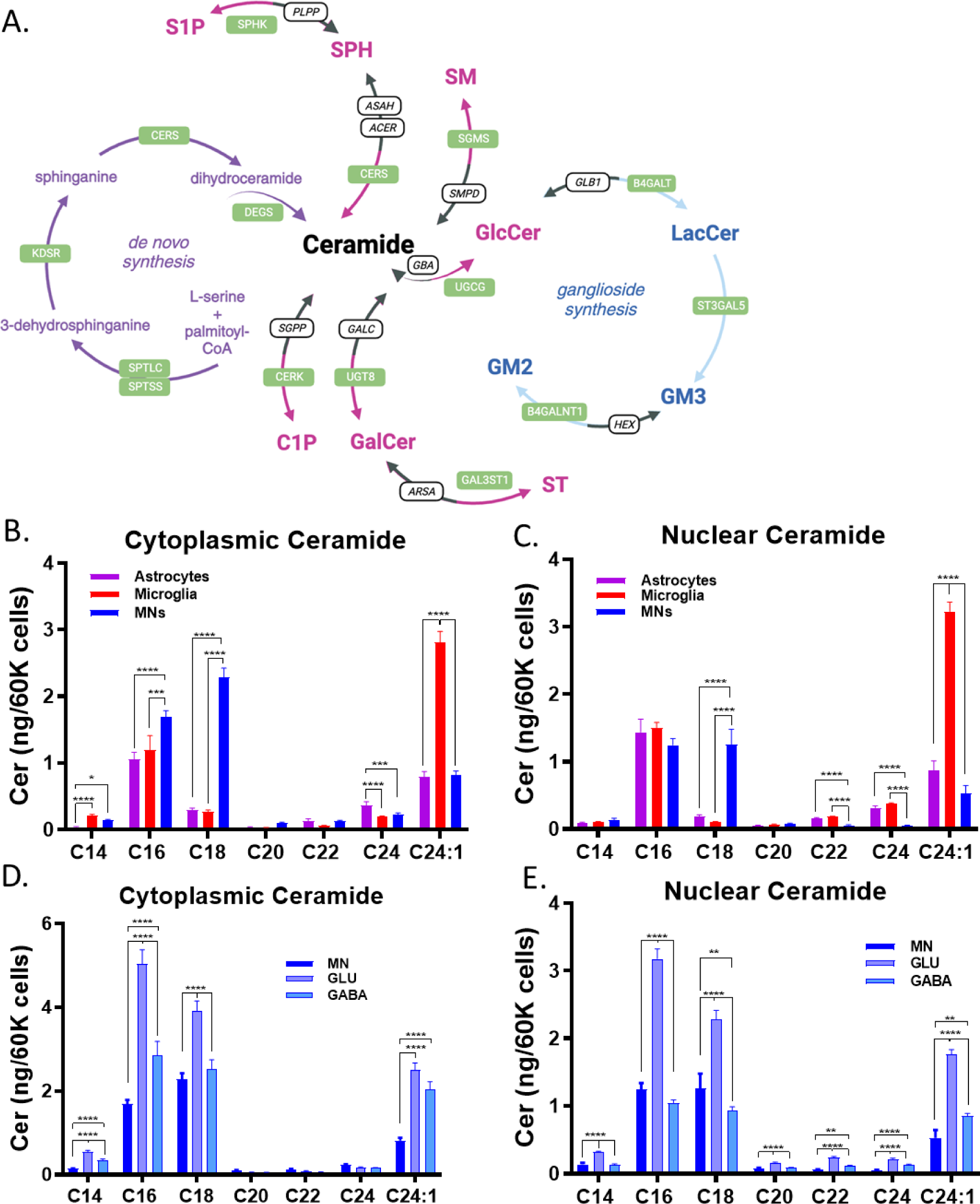
Human iPSC-derived neurons and glia differ in ceramide profiles. **A)** Diagram illustrating the different pathways involved in ceramide homeostasis. Graphs of both cytoplasmic **(B)** and nuclear **(C)** profiles of different ceramide chain lengths (*N* = 8 wells per condition). Motor neurons (MNs) had significantly higher levels of cytosolic and nuclear C18-ceramide (\*\*\*\**p<*0.01). Graphs comparing the cytoplasmic **(D)** and nuclear **(E)** profiles of different neuronal subtypes. Statistical comparisons made by one way ANOVA within each ceramide chain length (\**p*<0.05, \*\**p*<0.01, \*\*\**p*<0.001, \*\*\*\**p*<0.0001; *n* = 8 replicate wells).

Disrupted ceramide homeostasis (i.e., ceramide cacostasis) is implicated in the pathogenesis of both rare and common diseases of the CNS. Farber disease (FD) is a rare lysosomal storage disease caused by a complete deficiency in acid ceramidase (ASAH1), an enzyme that breaks down ceramide into sphingosine. In FD patients and FD models, the resultant accumulation of ceramide leads to inflamed joints, infiltration of macrophages into multiple tissues, and neurological dysfunction (Alayoubi et al., 2013). Mutations that result in partial loss of ASAH1 activity are associated with non-5q spinal muscular atrophy, and morpholino knockdown of ASAH1 in zebrafish leads to a loss of motor neuron axons, further bolstering evidence that ceramide accumulation triggers neurodegeneration (Zhou et al., 2012). Interestingly, SPTLC1 (an SPT complex member) gain of function mutations that increase ceramide synthesis also lead to motor neuron disease in children (Johnson et al., 2021; Kölbel et al., 2022; Mohassel et al., 2021). Moreover, mutations in SPTLC that result in deoxysphingolipid accumulation lead to Hereditary Sensory and Autonomic Neuropathy Type I (HSAN-I), a disease with sensory loss and distal muscle weakness (Dawkins et al., 2001; Kölbel et al., 2022; Penno et al., 2010). On the other hand, loss of function of DEGS1 disrupts ceramide de novo synthesis and leads to hypomyelinating leukodystrophy type 18 (HDL18) (Pant et al., 2019; Planas-Serra et al., 2023), illustrating the importance of precise ceramide synthesis regulation for brain development and function.

Studies indicate that ceramide cacostasis also contributes to the pathogenesis of more common neurodegenerative diseases including Parkinson’s disease (PD) and Alzheimer Disease (AD). For example, ceramide is increased in PD patient plasma but decreased in cortical areas (Abbott et al., 2014; Guedes et al., 2017; Mielke et al., 2013). Knockout of PD risk genes in both flies (*iPLA2-VIA* knockout) and mice (*Lrrk2* knockout) resulted in increased ceramides, and overexpression of cellular alpha-synuclein (a pathological hallmark of PD) caused ceramide accumulation (Ferrazza et al., 2016; Lin et al., 2018). Multiple studies point to increased ceramide levels in AD patients (Cutler et al., 2004; Han et al., 2011; Kim et al., 2017). Ceramides are elevated in serum exosomes of 5xFAD model mice, toxic amyloid treatment increases astrocyte ceramide levels, and ceramide containing exosomes activate cell death pathways (Elsherbini et al., 2020; Wang et al., 2012). Inhibition of ceramide synthesis is beneficial in mouse models of AD (TgCRND8), reducing both cortical ceramide and hyperphosphorylated tau levels (Geekiyanage et al., 2013).

While ceramide dysregulation contributes to multiple CNS diseases, it is unclear how ceramide levels are regulated in different CNS cell-types (Johnson et al., 2021; Mohassel et al., 2021). Here, we utilized motor and cortical neurons, astrocytes, and microglia derived from human induced pluripotent stem cells (iPSCs), to understand cellular differences in ceramide homeostasis. Each cell type had a unique ceramide composition, expression profile of ceramide synthases and ceramidases, and ceramide synthase activity. Specifically, glia tended to have higher levels of ceramide synthesis related genes, higher de novo synthesis activity, and increased sensitivity to CerS inhibition. By RNA sequencing, we additionally show that glial, but not neuronal, RNA expression profiles are altered by CerS inhibition.

Notably, the ceramide profile of neurons appeared uniquely stable relative to microglia and astrocytes. Glia, and particularly astrocytes, have been implicated in the release of toxic lipid species, with neurons and oligodendrocytes often disproportionately affected by the toxicity. We therefore sought to understand how each cell might respond to ceramide precursors. Despite a very stable ceramide profile, neurons were relatively more susceptible to sphinganine toxicity than astrocytes. Surprisingly, microglia showed similar levels of sensitivity to sphinganine toxicity to neurons. These results were recapitulated to a certain extent by inhibition of glucosylceramide synthase inhibitor that triggered ceramide accumulation, as well as treatment of neurons with sphinganine in a liposomal formulation. This baseline understanding of how different cell types synthesize ceramide and respond to alterations in ceramide synthesis, may provide insights into the mechanisms behind multiple neurodegenerative diseases and illuminate new avenues for therapeutic intervention.

## Results

### Neurons and glia have distinct ceramide profiles

Previous studies have established that mouse CNS cell-types differ in their general lipidomic profiles (Fitzner et al., 2020). However, the regulation of ceramide homeostasis in these cell types is not understood. Specifically, it is not known whether they share common ceramide species or whether ceramide generation occurs through distinct pathways (Figure 1A). Furthermore, although sphingolipids play important roles in nuclear signaling and metabolism (Lucki and Sewer, 2012), the subcellular composition of ceramides in CNS cell-types has not been investigated. Therefore, we utilized liquid chromatography/mass spectrometry (LC/MS) to profile nuclear and cytosolic fraction ceramides in human iPSC-derived cells (Figure 1B-E). This allowed us to establish whether there are basal differences in human neuronal and glial ceramide profiles. Cytosolic C16- and C18-ceramide were highest in motor neurons (MNs), whereas astrocytes had the highest C24-ceramide and microglia the highest C24:1 (Figure 1B). Astrocytes and microglia had significantly higher levels of nuclear fraction long chain ceramides (C22, C24, and C24:1) than MNs (Figure 1C). Consistent with the cytosolic fraction, MNs had significantly higher levels of nuclear C18 than glial cells (Figure 1C). *In vivo* studies have shown that CerS1 is neuronally localized and generates C18-ceramide (Ginkel et al., 2012). This illustrates that the differing ceramide profiles in our iPSC-derived neuronal cell types may match *in vivo* profiles.

### Ceramide profiles in different neuronal subtypes also vary

As neurons are often uniquely affected in diseases with sphingolipid dysregulation, we next determined the relative levels of ceramides in glutamatergic and GABAergic iPSC derived cortical neurons. Consistent with MNs, C18-ceramide levels were high in cortical neurons relative to the levels seen in glia (Figure 1D,E). Glutamatergic neurons had the highest levels of C16- and C18-ceramide in both nuclear and cytosolic fractions (Figure 1D) and had the highest levels of all ceramide chain lengths in the nuclear fraction (Figure 1E). MNs had uniquely low levels of C24:1 in both fractions and C14 in the cytosolic fraction (Figure 1D,E). This establishes that, while neurons have some commonalities in their ceramide profiles, the neuronal-subtype ceramide signatures are significantly different in ways that could be relevant to normal physiology and disease.

### Bulk RNA-Seq characterization of ceramide pathway members within MNs and glia

Our lipidomics data establish that neurons and glia have distinct ceramide profiles. To identify the underlying genetic expression driving our lipidomic profile differences, we performed RNA sequencing on iPSC-derived astrocytes, MNs, and microglia. Due to differences in cellular viability, RNA libraries from neurons and glia were prepared differently. We are not comparing cellular expression directly by RNA-sequencing, we only assessed the relative expression levels within each cell-type. Regardless, differences between astrocytes and microglia, which were processed similarly, are likely physiologically relevant. These data suggest that both glia and MNs depend on *SPTLC1*, *CERS2*, *DEGS1*, and *SPTSSA*, for ceramide production and that MNs might be distinctly dependent on *SPTSSB* and *DEGS2* (Figure 2A). MNs, astrocytes, and microglia have relatively lower *CERS3* expression relative to other ceramide synthesis pathway members (Figure 2A). Microglia had uniquely low *CerS1* expression as compared to other CerS. ORMDLs, negative regulators of SPTLC (Cai et al., 2016) were expressed in all cell types with *ORMDL1* and *ORMDL3* levels being relatively higher than *ORMDL2* in astrocytes and MNs (Figure 2A). Expression of enzymes comprising the sphingolipid anteome, metabolic pathways (e.g., NADPH, L-serine and Acyl-CoA) that directly converge on ceramide synthesis (Santos et al., 2022), showed similar patterns of expression across all cell types with a few exceptions (Supplemental Figure 1C). For example, enzymes mediating L-serine synthesis including *PHGDH, PSAT1, PSPH* were amongst the most highly expressed anteome members within astrocytes, which was not the case in microglia. *ACSL1* and *ACSL5,* members of the acyl-CoA synthase family, had low expression relative to other ACSLs in MNs and astrocytes but expressed at levels comparable to other ACSLs in microglia.

**Figure 2.**
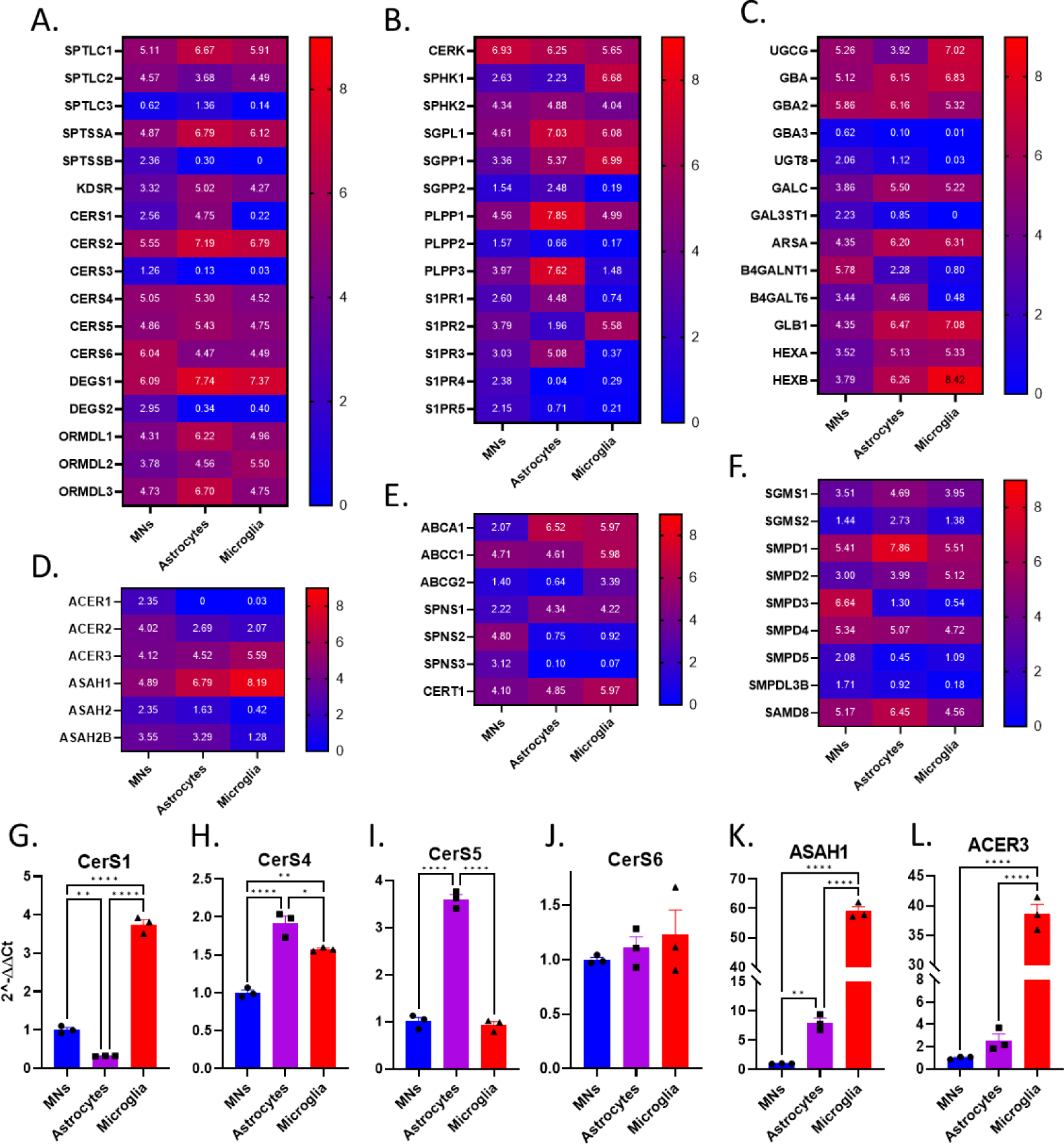
RNA-seq and qPCR characterization of ceramide homeostatic pathways in iPSC-derived motor neurons, astrocytes, and microglia. **A-F)** Heatmaps illustrating the expression levels of genes involved in ceramide synthesis **(A)**, ceramide/sphingosine phosphorylation and dephosphorylation **(B)**, glycosphingolipid utilization and generation of ceramide **(C)**, ceramide hydrolysis **(D)**, ceramide transport **(E)**, and sphingomyelin homeostasis, in iPSC-derived cells analyzed by bulk RNA-Seq (*n* = 3 replicate wells for astrocytes and motor neurons (MNs), *n* = 4 microglia replicate wells). **G-L)** iPSC-derived cells were lysed and analyzed by Taqman qPCR for the expression of genes involved in ceramide synthesis and ceramidases. All gene expression was normalized to GAPDH controls. (One-way ANOVA, Tukey’s multiple comparisons test, \**p*<0.05, \*\**p*<0.01, \*\*\*\**p*<0.0001; *n* = 3 replicate wells, 3 qPCR reactions per well).

We also found cell type differences in the expression of enzymes regulating the generation of ceramide via its salvage from the dephosphorylation or hydrolysis of more complex lipids, the use of ceramide as a substate in lipid synthesis, ceramide hydrolysis, and ceramide transport (Figure 2B-F). *PLPP1*-*3* are enzymes that dephosphorylate ceramide-1P to ceramide. All cells had relatively low *PLPP2* expression and microglia had low *PLPP3* as compared to *PLPP1* (Figure 2B). Lysosomal enzymes (i.e., *GALC, ARSA, GLB1, HEXB* and *SMPD1*) that generate ceramide through lipid hydrolysis had high levels of expression in glia relative to other pathway members (Figure 2C, F). *SMPD3*, a non-lysosomal hydrolytic enzyme of sphingomyelin, was the most highly expressed SMPD gene in MNs, while being amongst the lowest in glia (Figure 2F). Similarly, enzymes regulating ceramide hydrolysis also varied by cell type with *ASAH1* being enriched in glia and *ACER1* in MNs (Figure 2D). The expression pattern of lipid synthases that use ceramide to make more complex lipids were comparable across cell types except for *UGCG*, which was enriched in microglia relative to other synthases (Figure 2C). Differences in the expression of sphingolipid transport enzymes were also noted for glia and MNs, with *ABCA1* and *SPNS1* enriched in glia and elevated levels of *SPNS2* and *SPNS3* in MNs (Figure 2E). Finally, some differences were found in the expression of enzymes that control the phosphorylation status of sphingosine, the hydrolytic product of ceramide, and its receptors. For example, *SPHK1* levels in microglia were high compared to expression levels of similar enzymes, which was not the case astrocytes (Figure 2B). In MNs *S1PR1, SP1R2, S1PR3, S1PR4* and *S1PR5* were detected at comparable expression levels. By contrast, levels of *S1PR2* and *S1PR3* were elevated relative to other S1PRs in microglia and astrocytes respectively (Figure 2B).

Because we were not able to directly compare gene expression of different cell-types by RNA-sequencing, we performed qPCRs on ceramide synthesis pathway members to perform more direct comparisons of CerS. The SPT complex is an essential rate limiting step in ceramide synthesis, so we also assessed the expression levels of complex members *SPTLC1, SPTLC2, SPTSSA,* and *SPTSSB.* Microglia had higher expression of multiple synthesis genes (*CERS1, SPTLC1, SPTLC2, SPTSSA*) as compared to astrocytes and MNs (Figures 2G, S1A, and S1B). Microglia also expressed ceramidases (*ASAH1* and *ACER3*) at significantly higher levels (Figure 2K and 2L). Notably, *SPTSSA* was highest in microglia, as this preferentially uses shorter chain acyl-CoAs to produce long chain bases, whereas MNs were highest for *SPTSSB* (Figure S2B). *SPTSSB* prefers longer chain acyl-CoAs such as C18 to produce long chain bases (Han et al., 2009). By qPCR, MNs also expressed significantly higher levels of CerS1 than astrocytes (Figure 2G). CerS1 generates C18-fatty acid chain length and is known to be enriched in neurons *in vivo*. Astrocytes were highest for expression of CerS4 and CerS5 (Figure 2H).

### Neurons and glia differ in ceramide synthesis enzyme activities

RNA expression levels do not always correlate well with protein expression or protein function in the CNS (Taski et al., Nat Communications 2022), as might be indicated by our RNA-seq, qPCR, and lipidomics results. For example, CerC18 was highest in MNs relative to glia, *CERS1* expression measured by RNA-seq was low in MNs relative to other ceramide synthesis pathway members, and *CERS1* expression measured by qPCR was higher than astrocytes but lower than microglia (Figures 1 and 2). To clarify the synthesis activity of each cell type, we characterized the functional activity of the CerS enzymes (Figure 3A) by utilizing a ceramide synthesis activity assay. Briefly, palmitoyl-CoA and NBD-sphinganine were incubated at 37-degrees with astroglial, microglial, and MN lysates, followed by isolation and fluorescence reads of NBD-ceramide produced from NBD-sphinganine by CerS. This allows the use of different palmitoyl-CoA chain length precursors to assay CerS5/6 (16:0 palmitoyl-CoA), CerS1/4 (18:0 palmitoyl-CoA), or CerS2 (24:1 palmitoyl-CoA) dependent ceramide production (Tidhar et al., 2015). Microglia had significantly higher levels of CerS1/4 (Figure 3B), CerS2 (Figure 3C), and CerS5/6 activity (Figure 3D), than did astrocytes and MNs. Astrocytes had significantly higher CerS5/6 activity than MNs (Figure 3D), whereas MNs had significantly higher CerS1/4 activity than astrocytes (Figure 3B). This is consistent with our data showing high CerC18 levels in MNs (Figure 1), and *in vivo* reports suggesting that neurons have uniquely high CerS1 expression and CerC18 levels (Ginkel et al., 2012). We take this as additional confirmation that our iPSC derived cells maintain features of *in vivo* cell types. Based on comparing the different activity assays, this suggests that astrocytes might be uniquely dependent on CerS5/6 activity for ceramide synthesis, while neurons uniquely dependent on CerS1. This pattern does not necessarily follow from the RNA-seq, qPCR analysis, and lipidomics. All microglial CerS activity was higher than the other cell types, which was not predicted from RNA measures. Additionally, the unique dependence of astrocytes on CerS5/6 does not match the relative levels of astroglial CerC14 and CerC16 compared to neurons and microglia. These discrepancies between RNA levels and CerS activity, indicate that functional outputs for ceramide synthesis are needed. The lack of alignment between lipid levels and CerS activity is also likely due to cell type dependent differences in the generation of ceramide from alternative pathways.

**Figure 3.**
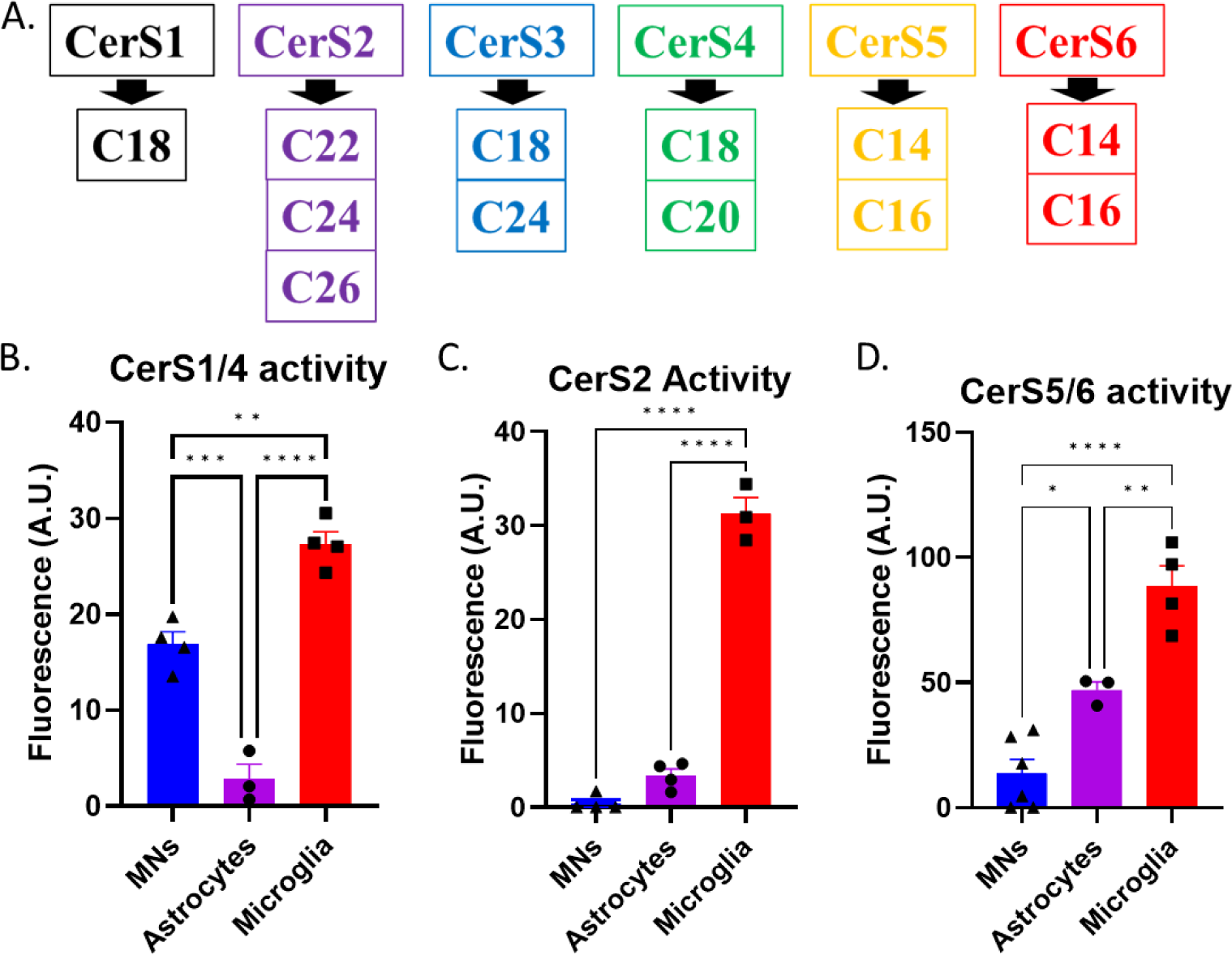
Assay of CerS1/4, Cers2 and Cers5/6 enzymatic activities in motor neuron and glial cell lysates. **A)** Schematic of the different fatty acid carbon chain lengths produced by different CerS enzymes. **B)** Assay of CerS1/4 (18:0 palmitoyl-CoA and NBD-sphinganine added to lysate) activity in astrocyte, microglia, and motor neuron (MN) cell lysates. MNs had significantly higher CerS1/4 than did astrocytes (One-way ANOVA, Tukey’s multiple comparisons test, \*\*\**P*<0.001). **C)** Assay of CerS2 (24:1 palmitoyl-CoA and NBD-sphinganine added to lysate) activity in astrocyte, microglial, and MN cell lysates. **D)** Assay of CerS5/6 (16:0 palmitoyl-CoA and NBD-sphinganine added to lysate) activity in astrocyte, microglial, and MN cell lysates. **B, C, D)** Microglia had the highest CerS activity for all ceramide synthases measured (One-way ANOVA, Tukey’s multiple comparisons test, \*\*\*\**p*<0.0001, \*\*\**p*<0.001, \*\**p*<0.01). **D)** Astrocytes had significantly higher CerS5/6 activity than did MNs (One-way ANOVA, Tukey’s multiple comparisons test, \**p*<0.05). All synthesis activity was normalized to cell number (activity per 200k cells) by taking nuclear cell counts and subtracting dead cells using nuclear cell-death stains from representative culture wells (activity assay is *n* = 3-6 wells of plated cells as shown in graphs, with *n* = 3 CerS activity reactions performed per well).

### Pharmacological inhibition of CerS has cell-type specific effects on ceramide profiles

While basal ceramide levels and RNA expression data are informative, it is important to determine if CerS activity drives basal ceramide levels in differentiated neurons and glia. By treating with the pan-CerS inhibitor fumonisin-B1 (FB1), we isolated the portion of the ceramide profile driven by *de novo* CerS synthesis, versus lipids that are generated through alternative pathways (Figure 1A). This might clarify differences observed in CerS-activity, lipidomics, and RNA expression. A variety of ceramide chain lengths were reduced after 24 hours of FB1 treatment of glial cells (Figures 4A, 4B, 4C, S2C, and S2E). Notably, the magnitude of reduction in astrocyte ceramides was larger than changes in microglial ceramide in response to FB1 inhibition (Figures 4A, 4B, 4C, S2C, and S2E). This could indicate differences in the levels of ceramide synthesis between astrocytes and microglia, or a greater compensatory change in microglia to keep ceramide levels stable via other ceramide generating pathways. Microglia did have less significant ceramide changes in the cytoplasmic fraction (Figure 4C), indicating that cellular compartments such as lysosomes might give microglia unique capacities to compensate for changes in lipid synthesis. Interestingly, some of the largest changes in astrocyte ceramides were in CerC14 and CerC16, consistent with the high levels of CerS5/6 activity in these cells (Figures 3D, 4B, and 4E). Microglia had some of the largest changes in longer chain ceramides, consistent with their high level of CerS2 activity.

**Figure 4.**
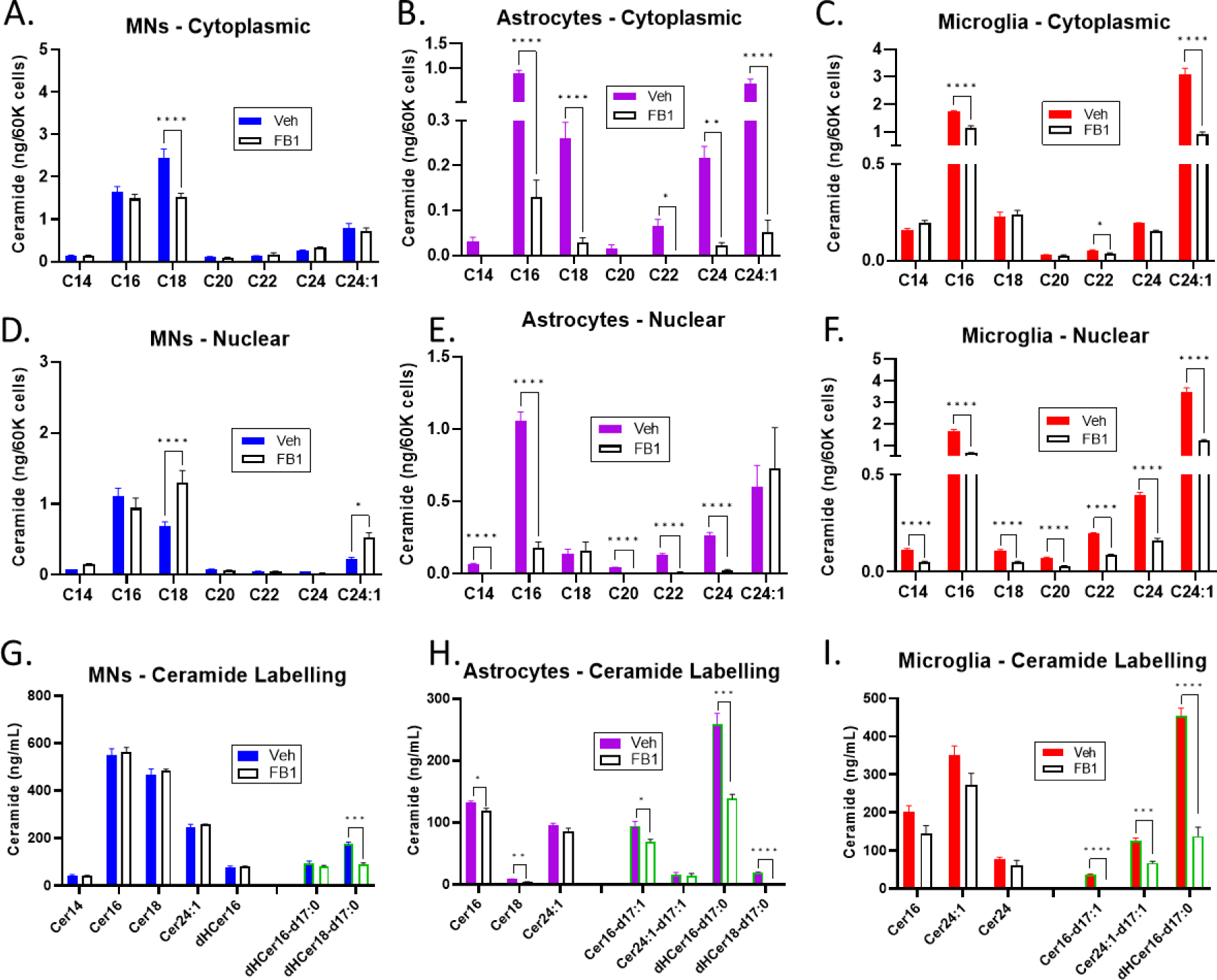
Ceramide profiles of iPSC derived neural cell types treated with FB1 and/or d17:0-sphinganine. **A-F)** Motor neuron (MN**) (A and D)** astrocyte **(B and E)** and microglial **(C and F)** cytoplasmic and nuclear ceramide profiles after 72 hours of CerS inhibition (3 μM FB1 treatment). **G-I)** By introducing a traceable sphinganine (d17:0) precursor, in the presence or absence of 3 μM FB1, we were able to track the active CerS-dependent ceramide synthesis in each cell type over a 2-hour period. A significant percentage of astroglial **(H)** and microglial **(I)** ceramides were labelled (bars with green outlines) after the two hours. Notably, MNs **(G)** had a lower percent of each ceramide labelled. Further labelling data in Figures S3 and S4. Each ceramide chain length vehicle vs FB1 treatment was analyzed by Student’s T-test (\*\*\*\**P*<0.0001, \*\*\**P*<0.001, \**P*<0.05; *n* = 4 replicates).

In MNs, there were minimal changes in ceramide in response to FB1 inhibition – even after 72 hours of treatment. MNs showed some small changes in CerC18 and CerC24:1 (Figures 4A and S2A). It is possible that FB1 targets the CerS1 activity in MNs to slightly alter lipid levels and/or distribution, or that there could be compensatory changes in other regulators of ceramide. By contrast, in response to FB1 treatment, glutamatergic and GABAergic cortical neurons both showed more robust significant reductions in ceramides (Figure S5). This may suggest that MNs have some distinct properties in maintaining ceramide stability.

### Labelling with d17:0-sphinganine confirms glial and neuronal ceramide synthesis rates

Perturbation of CerS and measures of overall ceramide levels may fail to adequately detect small changes in synthesis or the overall *de novo* synthesis capacity of each cell type. For a more sensitive measure of acute ceramide synthesis by CerS, we introduced a sphinganine precursor with an unnatural 17-carbon chain length (d17:0-sphinganine). This label is then incorporated into ceramide produced through CerS/*de novo* ceramide synthesis pathway. We used whole-cell lysate instead of fractionated samples to increase the sensitivity for detecting both labeled and unlabeled products. Across cell types, label incorporation was higher in dihydro-ceramides (dhCer) than in ceramides, consistent with the newly synthesized lipids incorporating the label, but labelled dihydroceramide not yet converted to ceramide by DEGS activity (Figures 4G, 4H, and 4I). This, along with the lower label incorporation in FB1 treated cells, is indicative that the d17:0-sphinganine label is labelling ceramide generated through *de novo* synthesis as expected. There was a greater reduction in labelled ceramides with FB1 treatment than in unlabeled ceramides, consistent with labelled ceramides acting as a metric of acute ceramide synthesis (Figures 4G, 4H, and 4I).

Labelled ceramide was a more significant proportion of total ceramide in glia than MNs after two hours of incubation (Figures 4G, 4H, 4I, and S4). This is consistent with our observation that MNs have significantly lower activity from multiple CerS than glial cells and show a minimal change in ceramide following FB1 treatment. This may also suggest that the high level of MN CerS1/4 activity in our enzyme assay may not directly parallel CerS1 kinetics in MNs *in vitro*, or account for ceramide incorporation into more complex lipids or its breakdown. Notably, newly synthesized lipids tracked cellular CerS activity in most cases. MNs had high CerS1/4 activity and low activity for other CerS (Figure 3B), and correspondingly the limited newly synthesized ceramides were mostly dhCerC18 (Figures 4G, S3E, and S3F) which results from CerS1 activity. Consistent with these findings CerS5/6 (Figure 3D) was maximal in astrocytes, which correspondingly had higher amounts of labelled dHCer16 and Cer16 (Figure 4H). Microglia were high for activity of all CerS enzymes (Figure 3) and had high levels of newly synthesized Cer16 and Cer24:
(Figure 4I). The main inconsistency between CerS-activity and ceramide labelling is the high level of microglial CerS1/4 (Figure 3B) activity but no detectable levels of newly synthesized Cer18 (Figure 4I). Future experiments will need to determine the relative contribution of CerS1 and CerS4 in regulating Cer18 levels in microglia. If CerS1 plays a more limited role than Cers4, which is suggested by our RNA-seq findings, this would be consistent with the essential role for CerS4 in regulating immune cells and the context specificity of the ceramide chain lengths generated by CerS4 (El-Hindi et al., 2022). This discrepancy could also be due to microglia having an increased capacity for hydrolyzing CerC18 or incorporating it into more complex sphingolipids and glycosphingolipids compared to other cell types.

### Inhibition of glucosylceramide synthesis alters ceramide profiles in neurons and glia

The *de novo* ceramide synthesis pathway is highly dependent on CerS generation of dHCer and its subsequent conversion to ceramide by DEGS. FB1 was a logical tool to reduce ceramide synthesis and the addition of sphinganine allowed us to track *de novo* synthesis in these cell types. To induce ceramide accumulation in cells without the addition of an exogenous lipid, we treated cells with a glucosylceramide synthase inhibitor (GCSi; 667161) to prevent ceramide from being used to synthesize glucosylceramide (Figure 1A). Given the differences in glial and MN dependence on *de novo* ceramide synthesis, we predicted that cellular differences in ceramide accumulation would likely occur after GCSi treatment. In astrocytes, GCSi treatment significantly increased CerC14 and CerC16 (Figures 5B and 5E), consistent with high CerS5/6 activity and the reduction in the same ceramides with FB1 treatment. Microglia also had small accumulations in CerC14 and CerC16 in response to GCSi (Figures 5C and 5F). The most dramatic increase in MNs after GCSi treatment was C18-ceramide in the cytoplasmic fraction (Figures 5A and 5D), the expected chain length based on MN CerS1 activity and which ceramides were reduced after FB1 treatment. Ceramide changes in response to GCSi differed between nuclear and cytosolic fractions, potentially due to differences in glucosylceramide intracellular flux amongst different cellular compartments. We note that microglia did not significantly accumulate CerC24 or Cer24:1(although there was a trend) in response to GCSi treatment as we would have expected from CerS-activity, ceramide labelling, and FB1 treatment. This observation also supports the idea that microglia are more readily adaptive to elevated ceramide levels (e.g., by increasing hydrolysis or increasing complex lipid synthesis) compared to astrocytes and MNs.

**Figure 5.**
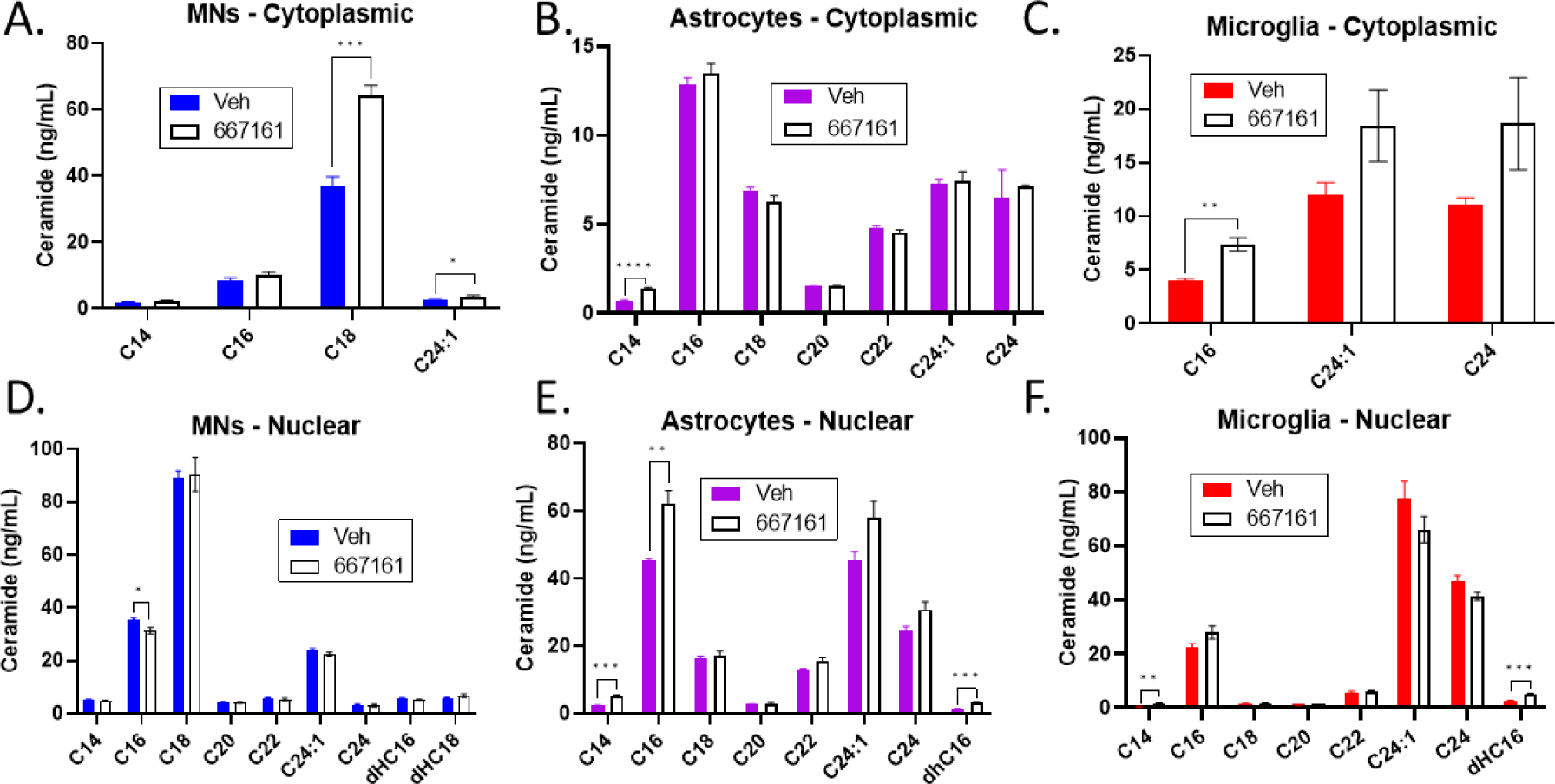
Ceramide Profiles of iPSC derived neural cell types treated with GCSi. **A-F)** To alter ceramide production through inhibition of downstream glucosylceramide production, MNs **(A and D)**, astrocytes **(B and E)** and microglia **(C and F)** were treated with a GCSi (Genz-667161) and ceramides were measured in nuclear and cytoplasmic fractions. Each ceramide chain length vehicle vs GCSi treatment was analyzed by Student’s T-test, (\*\*\*\**P*<0.0001, \*\*\**P*<0.001, \**P*<0.05; *n* = 4 wells).

### Bulk RNAseq of cells treated with FB1 to reduce ceramide synthesis and GCSi to induce ceramide accumulation

While we observed different gene expression profiles across cell-types by qPCR and RNA-seq, this did not determine which of their cellular functions depend on ceramide synthesis. Ceramide labelling, FB1 treatment, and GCSi treatment all revealed cell-type differences in ceramide homeostasis. To assess how ceramide accumulation influences cellular function we treated each cell type with the pan-CerS inhibitor FB1 or GCSi (Genz-667161), then performed RNA-seq. As in previous experiments, cells were cultured for 1-week post thaw then treated with 3 μM FB1 or Genz-667161 for 72 hours before lysis. After CerS inhibition, we observed many differentially expressed genes (DEGs) in both astrocytes (1820 genes) and microglia (1734 genes) (Figures 6B, 6C, and 6D). Notably, there were no DEGs in MNs after FB1 treatment, consistent with our data showing minimal reliance on ceramide synthesis in MNs relative to glia (Figure 6A). GCSi induced significant DEGs in all three cell types (astrocytes 409 DEGs, Microglia 920 DEGs, and MNs 576 DEGs) (Figures 6F, 6G, 6H, and 6I).

**Figure 6.**
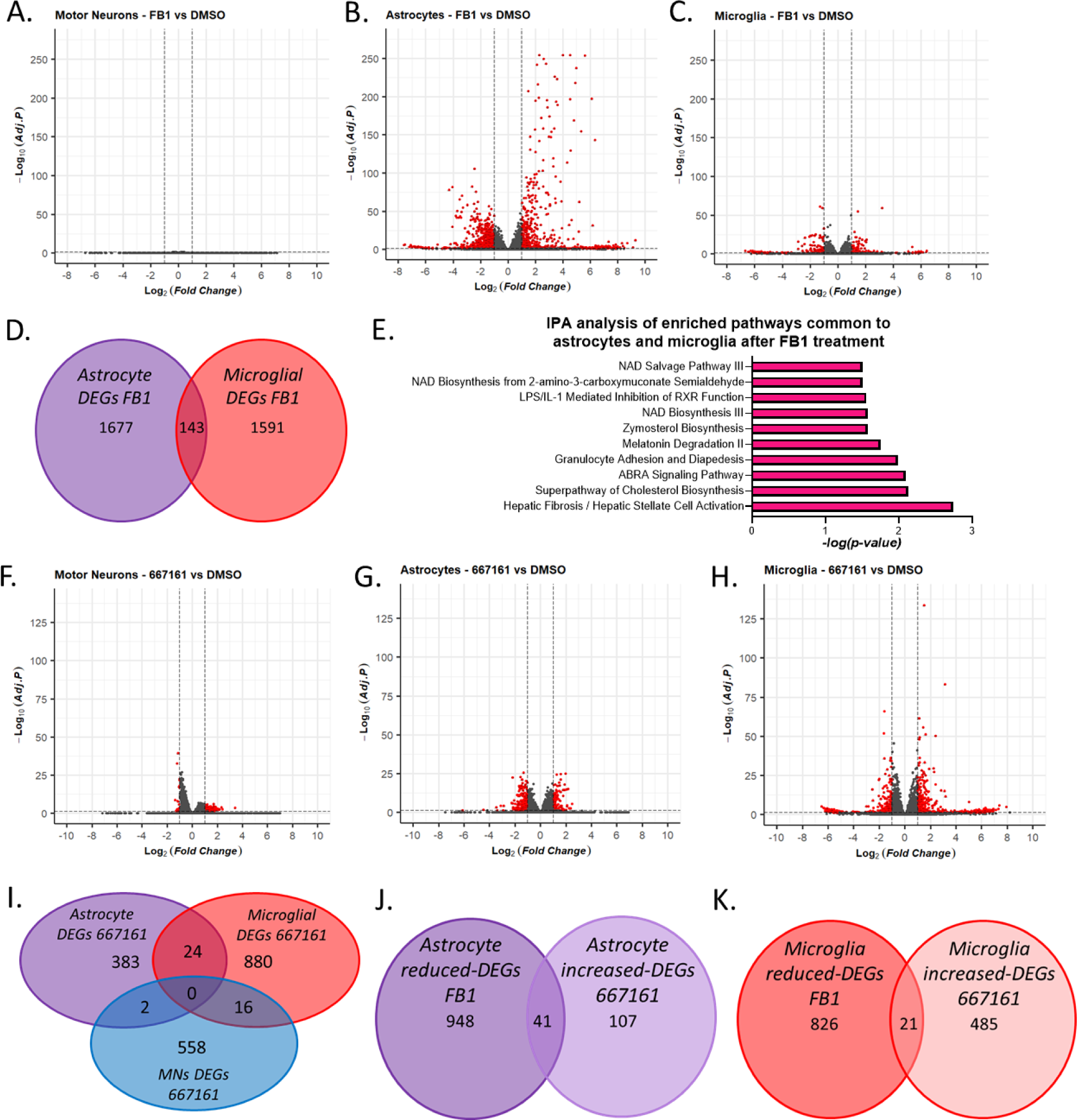
Bulk RNAseq of astrocytes and motor neurons treated with FB1 and GCSi. **A-E)** Motor neurons (MNs), astrocytes, and microglia were cultured for 7-days, then treated with 3 μM FB1 for 72-hours, as with previous protocols. We then isolated RNA and samples were sent for bulk sequencing. **A-C)** Volcano plots of RNA-seq data in MNs, astrocytes, and microglia after FB1 treatment. **B)** Astrocytes showed 1,820 significant differentially expressed genes (sDEGs) in response to FB1 treatment. **C)** Microglia showed 1,734 sDEGs in response to FB1 treatment. **A)** MNs had no sDEGs. **D)** In response to FB1 treatment, astrocytes and microglia had 143 common sDEGs. **E)** The top 10 common IPA pathway changes in both astrocytes and microglia after FB1 treatment graphed by -log(p-value). **F-H)** Volcano plots of RNA expression in astrocytes, microglia, and MNs after 72 hours 3 μM GCSi treatment. **I)** Venn diagram of common gene expression changes after GCSi in MNs, astrocytes, and microglia. **J and K)** Venn diagrams of genes that both reduce expression after FB1 treatment and increase expression after GCSi treatment in astrocytes or microglia.

Ceramide is implicated in many cellular functions, so we first looked at Ingenuity Pathway Analysis (IPA) to determine if ceramides had different functions across cell types. Previous studies showed that ceramide accumulation might be a hallmark of glial reactivity (de Wit et al., 2019). Our RNA-seq data indicate that ceramide levels may act as a key regulator of reactivity and inflammation. Within the top 10 differentially regulated IPA pathways in response to FB1, astrocytes had multiple pathways involved in inflammatory regulation (*Hepatic Fibrosis*, *Agranulocyte Adhesion and Diapedesis*, *Pulmonary Fibrosis Idiopathic Signaling Pathway*, *Neuroinflammation Signaling Pathway*, *Osteoarthritis Pathway*, and *Ferroptosis Signaling Pathway*) (Supplemental excel Compiled-PathwayAnalsyis: astrocyte FB1 treatment pathways). Microglial inflammatory pathways were similarly impacted by FB1 treatment (*Granulocyte Adhesion and Diapedesis*, *Agranulocyte Adhesion and Diapedesis*, *Hepatic Fibrosis*, *Role of IL-17A in Psoriasis*, *Airway Pathology in Chronic Obstructive Pulmonary*) (Supplemental excel Compiled-PathwayAnalsyis: microglia FB1 treatment pathways). Additionally, consistent with their role in brain lipid regulation, FB1 significantly altered the *Superpathway of Cholesterol Biosynthesis* in astrocytes. Of interest, astrocytes also had pathway changes that could relate to axon guidance (*Axonal Guidance Signaling*). Common pathway changes in response to FB1 treatment in both astrocytes and microglia showed themes in lipid synthesis, NAD regulation, and inflammation (Figure 6E).

GCSi treatment altered many inflammatory pathways in astrocytes, similar to FB1 treatment (*Acute Phase Response Signaling*, *HMGB1 Signaling, TNFR2 Signaling, Atherosclerosis Signaling, Agranulocyte Adhesion and Diapedesis, Airway Pathology in Chronic Obstructive Pulmonary Disease*, *NOD1/2 Signaling Pathway*) (Supplemental excel Compiled-PathwayAnalsyis: astrocyte pathways in GCSi treatment). Astrocytes also showed changes in lipid/cholesterol regulation after GCSi treatment (*Hepatic Cholestasis*, *LXR/RXR Activation*). GCSi induced a variety of inflammatory changes in microglia as well, many involved in inflammatory cell migration and infiltration (*Granulocyte Adhesion and Diapedesis*, *Agranulocyte Adhesion and Diapedesis*, *Hepatic Fibrosis / Hepatic Stellate Cell Activation*) (Supplemental excel Compiled-PathwayAnalsyis: microglia pathways in GCSi treatment).

While MNs did not respond to FB1, there were a variety of pathway changes in response to GCSi. The changes centered around amino acid degradation (*Glutamate Degradation II*, *L-cysteine Degradation I*, and *Aspartate Degradation II*), lipid/cholesterol regulation (*LXR/RXR Activation* and *FXR/RXR Activation*), and inflammatory responses (*Osteoarthritis Pathway* and *Role of Cytokines in Mediating Communication*). Consistent pathway changes across these cell-types indicate that ceramides may have key roles in regulating inflammation and cholesterol homeostasis.

To assess which pathways were commonly altered in response to ceramide changes, we analyzed the pathways that changed the most in response to FB1 and GCSi across all three cell types. Of the 20 most highly affected pathways, many were involved in inflammatory functions (*Pathogen Induced Cytokine Storm Signaling Pathway*, *IL-33 Signaling Pathway*, *IL-17 Signaling*, *Neuroinflammation Signaling Pathway*, *Macrophage Classical Activation Signaling Pathway*, and *HMGB1 Signaling*). Many of these pathways were modified in opposing directions in response to FB1 and GCSi in astrocytes but changed in the same direction in microglia (Figure S7). Notably, IL-33 is known to be an astrocyte derived cytokine capable of regulating microglial synaptic pruning, and IL-17 similarly regulates synaptic activity (Luo et al., 2019; Vainchtein et al., 2018).

We then assessed changes in response to FB1 and GCSi at the individual gene level. Astrocytes had a small number of genes (6 DEGs) that were both up in response to FB1 treatment and down in response to GCSi (Supplemental document 1). These related to mitochondrial function (*PDK4*), fatty acid metabolism (*PDK4* and *ANGPTL4*), water balance (*AQP4*), and ER stress/apoptosis (*KLHDC7B*). Alternatively, 41-DEGs decreased in response to FB1 and increased in response to GCSi (Figure 6J and supplemental document 2). These related to ER stress/apoptosis (*BIRC3*, *PLN*), cholesterol regulation (*SREBF1*), inflammatory regulation (*CCL2*, *CTGF*, *TAGLN*), angiogenesis (*MYBL1*, *THSB1*), antioxidant activity (*GSTA1*) and cellular adhesion/cytoskeletal dynamics/cellular motility (*MPP4*, *ACTA2*, *RSPH4A*, *THBS1*, *DYNLT5*, *MMP10*, *STXBP6*). This left 106 DEGs that changed in the same direction in astrocytes in response to both FB1 and GCSi treatment (supplemental document 3).

Microglia had 21 DEGs that were down in response to FB1 treatment and up in response to GCSi (Supplemental document 4). These DEGs were involved in cellular adhesion/cytoskeletal dynamics/cellular motility (*MYH16*, *MMP21*, *CLDN5*), inflammatory regulation (*NTSR1*, *HMGA2*, *HTRA3*), depalmitoylation (*NOTUM*), blood coagulation (*F13A1*), and sulfatation (*SULT1B1*, *ARSI*) (Figure 6K). There were zero microglial genes that were up in response to FB1 treatment and down in response to GCSi treatment. This left 257 genes that changed in the same direction in response to both FB1 and GCSi treatment (Supplemental document5).

Of interest were common and unique gene changes between cell types. In response to FB1 treatment, there were 143 DEGs in common between astrocytes and microglia (Supplemental document 6). Of the 143 shared DEGs, 38 genes increased expression. These related to ER stress/apoptosis (*STC2*, *CHAC1*), inflammatory regulation (*AREG*, *BEX2*, *TREML3P*, *FOXD1*), glycolysis (*GCKR*), antioxidant activity (*CHAC1*), NAD metabolism/mitochondrial function (*ALDH1L2*, *NMNAT2*), and cellular adhesion/cytoskeletal dynamics/cellular motility (*PCDH19*, *PAPPA*). There were 41 downregulated genes in astrocytes and microglia in response to FB1 treatment. These included genes involved in cholesterol and fatty acid regulation (*MSMO1*, *HMGCS1*, *CYP2U1-AS1*), inflammatory regulation (*RNASE1*, *CXCL14*, *TGFB2-AS1*, *PTGES3L*), oxidative stress (*MAOB*), axon and synaptic development (*CBLN2*, *TMEM108*), sulfatation (*ARSI*), and cellular adhesion/cytoskeletal dynamics/cellular motility (*DNAH6*, *MYH2*, *ADAM19*, *ADAMTS12*, *CLDN5*, *TPM2*).

In response to GCSi treatment, there were no common DEGs amongst all three cell-types. However, GCSi treatment induced 24 common DEGs between astrocytes and microglia, astrocytes and MNs shared 2 DEGs, and microglia and MNs had 16 common DEGs (Supplemental document 7). The 16 common DEGs between microglia and MNs had functions in inflammatory regulation (*IL18RAP*, *GIMAP4*, *CD1B*, *TAFA3*) and ubiquitin degradation of proteins (*ASB15*). CD1B is of particular interest as it functions to present lipid and glycolipid self-antigens (Shahine, 2018). The 24 common DEGs between microglia and astrocytes had functions in extracellular matrix and blood clotting regulation (*TFIP2*, *SERPINE1*, *PCED1B*, *CLDN5*, *SPON1*), inflammatory regulation (*IL11*, *CLCF1*), and apoptosis (*DTHD1*). It is notable that both *IL11* and *CLCF1* had increased expression in response to GCSi and both promote oligodendrocyte survival (Gurfein et al., 2009; Ji-Wei et al., 2022; Zhang et al., 2006). It is possible that lipid stress induced by GCSi treatment is detected by glial cells as a sign of dysmyelination or demyelination. In response to GCSi astrocytes and MNs had only 2 DEGs, *PRRG* and *KCNE1B*.

### Ceramide precursors are differentially toxic between glia and neurons

We showed that ceramide synthesis is significantly different between iPSC-derived cell types. Additionally, glia and neurons incorporated sphinganine into ceramide chain lengths differently. In our RNA-seq data, CerS inhibition and GCSi induced gene changes related to ER stress, ferroptosis, and apoptosis. Sphingolipids are dysregulated in a variety of neurological diseases, and we showed that sphinganine can be turned into ceramide when applied to glia and, to a lesser extent, neurons. We therefore treated cells to alter ceramide and/or sphinganine levels, and analyzed the resulting toxicity by Cell-Titer Glo (CTG), DAPI/cytotox staining, and neurite analysis.

We first sought to determine the toxic doses of GCSi (Genz-667161), as these may directly relate to the ceramide accumulation seen in Figure 5. Despite having large changes in nuclear ceramides with GCSi treatment (Figure 5E), astrocytes were more resistant to GCSi toxicity than MNs and microglia (Figure 7A-C). We did see small decreases in astroglial viability at 72 hours (Fig 7B), but both microglia and MNs showed more significant toxicity (Figure 7A and 7C).

**Figure 7.**
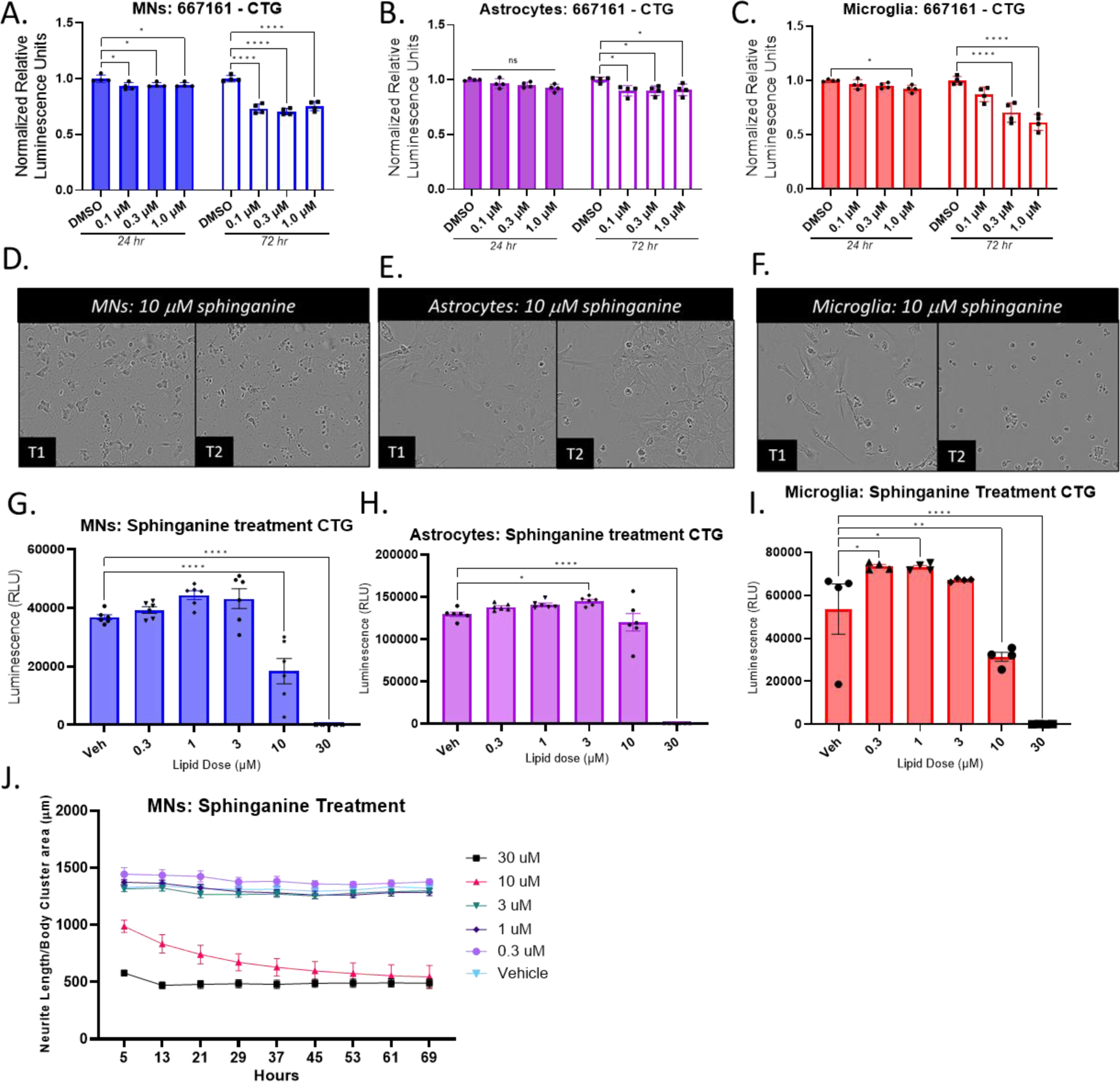
Introduction of a ceramide precursor and inhibition of downstream glucosylceramide synthesis are disproportionately toxic to motor neurons and microglia. **A-C)** Astrocytes, microglia, and motor neurons (MNs) were cultured for 7-days, then treated with increasing doses of GCSi. **D, E, F)** Sphinganine was dosed into cells at increasing concentrations and Incucyte live imaging and analysis was performed. Representative images show MN cell-death and disintegrating neurites, microglia showed signs of cell death, while astrocytes remained largely resilient to 10 μM sphinganine treatment. **G, H, I)** Cell titer glo (CTG) analysis of MN cell viability was consistent with neurite findings that toxicity starts at 10 μM doses with similar levels of toxicity in microglia, but resistance to 10 μM dose toxicity in astrocytes. **J)** Live imaging of MNs and analysis of neurites by phase microscopy revealed acute toxicity at 10 μM and higher. Data analyze by one-way ANOVA with Bonferroni’s multiple comparisons test, \*\*\*\**p*<0.0001, \*\*\**p*<0.001, \*\**p*<0.01, \**P*<0.05 (*n* = 4 or 6 technical replicates as depicted in graphs).

To alter ceramide levels by a different mechanism, we treated cells with increasing sphinganine concentrations to determine the effects on the viability of different cell-types. Both sphinganine (Figures 7D, 7G, 7J, S8A, S8B, and S8C) and sphingosine (Figures S8D-G) were significantly toxic to neurons at 10 μM doses, as assessed by CTG, DAPI/cytotox staining, and neurite analysis. High doses were acutely toxic and 10 μM sphinganine treatment showed more acute toxicity than 10 μM sphingosine.

Sphinganine was toxic to astrocytes as well, but only starting at the 30 μM dose (Figures 7E, 7H, S9A, S9C, and S9D). This illustrates a potential resistance of astroglial cells to this sphingolipid driven toxicity. We therefore treated microglia to see if other glia had similar resistance to toxicity. Microglia had higher mRNA expression of ceramidases, so we anticipated that microglia might have heightened resistance to sphinganine toxicity. In contrast to our expectations, and similar to MNs, microglia underwent cell death starting at 10 μM doses (Figures 7F, 7I, S9E, S9F, and S9G). We also observed significant acute morphological changes in microglial cells in response to sub-toxic doses (3 μM) of sphinganine (supplemental video 2). These morphological changes normalized over time, though there were signs of intracellular inclusion (possibly lipid droplets). It’s known that lipid accumulation can alter microglial activity (Victor et al., 2022), but future studies will need to ascertain whether sphinganine similarly alters their activity.

### Liposomal delivery significantly alters sphinganine delivery to neurons

Bulk application of free lipids is not the most physiological delivery and therefore may not recapitulate *in vivo* lipid-related toxicity. There is evidence that astrocytes can release lipoparticles and/or exosomes containing toxic lipids including ceramides (Guttenplan et al., 2021; Wang et al., 2012). Therefore, we repeated our sphingolipid treatment assay with delivery of sphinganine, but in a lipoparticle/liposomal format. NBD-sphinganine was mixed with POPC in a 5:95 ratio and delivered to cells as liposomes or liposomes bound to APOE. We compared this delivery to liposomes lacking sphinganine (with and without APOE) as well as delivery of free NBD-sphinganine at doses equivalent to that of NBD-sphinganine delivered by liposomes. NBD (green) allowed us to track the rate of incorporation and localization of the NBD-sphinganine within MNs, though we can’t rule out that the sphinganine could be incorporated into other lipids and/or metabolized in some way.

We observe that treatment of MNs with NBD-sphinganine (NBD-SA) in both a free and liposomal (LP) format led to NBD localization in MNs, with no green signal detected in POPC treated neurons (Figures 8B-E and S10A). A large portion of NBD signal in free NBD-sphinganine treatment localized to cell bodies and dead cell clumps. It is difficult to remove these clumps without negatively impacting live MN health. We can’t exclude that a large portion of free NBD-sphinganine signal in our analyses could localize to dead cells. Notably, we saw many neurites that were NBD^+^ in the liposomal treated cells (NBD-SA+POPC+APOE LP; Figure 8B, 8C, and 8E).

**Figure 8.**
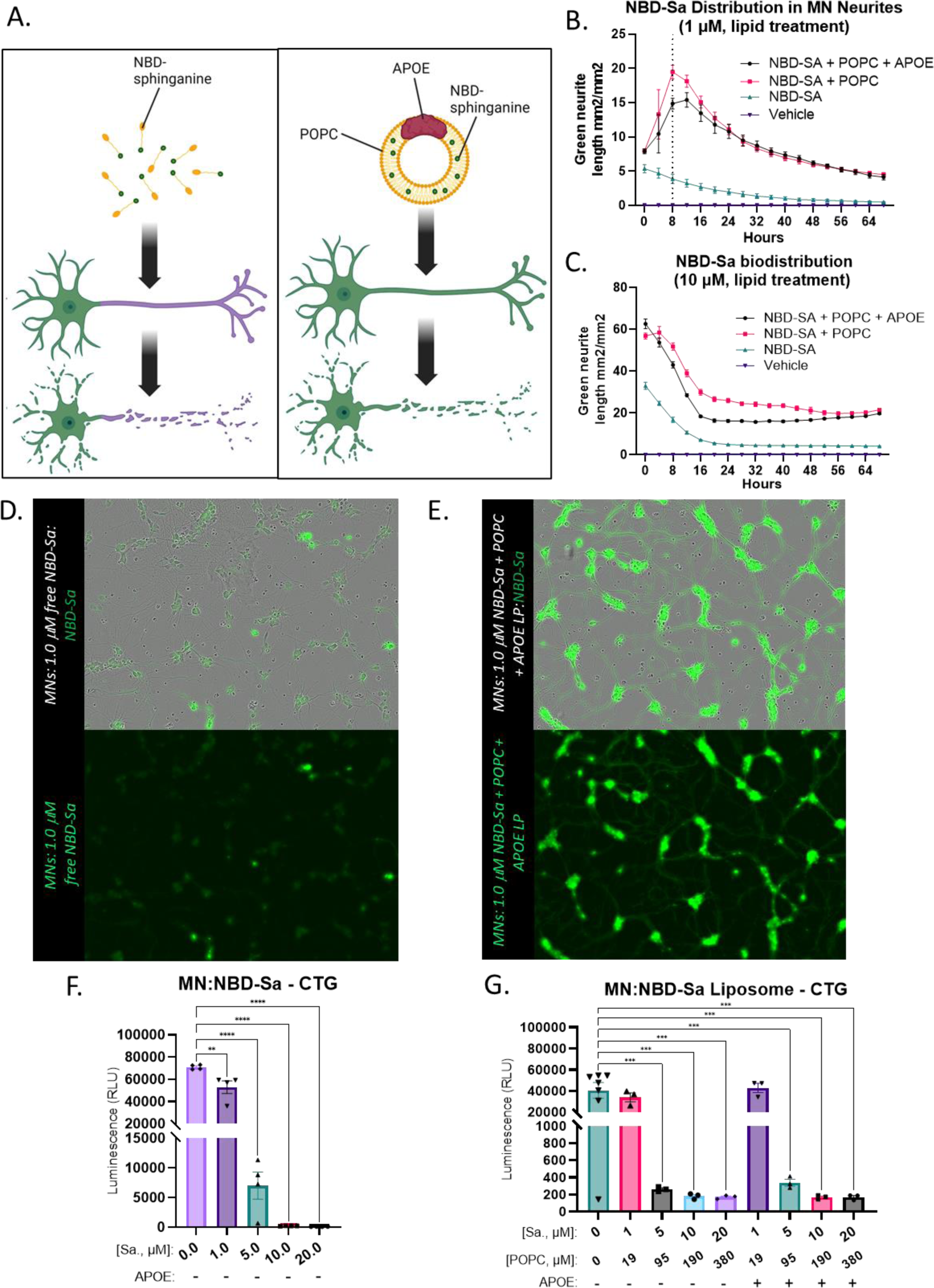
Liposomal delivery of sphinganine to motor neurons. **A)** Diagrams illustrating treatment of motor neurons (MNs) with liposomes with NBD-labelled (green) sphinganine and liposomes lacking sphinganine. These were delivered either bound or not bound to APOE. These treatments were compared to delivery of free NBD-sphinganine. **B and C)** To quantitatively assess if liposomal delivery increased NBD-label incorporation into neurites, we performed an analysis for green labelled neurites. Graphs illustrate the delivery of 1.0 μM and 10.0 μM of NBD-sphinganine in both free and liposomal forms, with increased NBD^+^ neurite area in liposomal delivery regardless of APOE status. Dotted line in **(B)** is where images were taken for **D and E)** 8 hrs post-treatment, showing significant NBD labelling in neurites when NBD-sphinganine was delivered in liposomes **(E)** but did not see the same extent of labelling in the free NBD-sphinganine **(D)** delivery condition. No green labelling was observed in neurons treated with liposomes lacking NBD-sphinganine (Figure S10). We noted **F and G)** Cell titer glo (CTG) analysis showed that NBD-sphinganine had a similar toxicity curve when delivered freely or delivered in liposomes. Notably, liposomes without NBD-sphinganine showed minimal toxicity (Figure S10), showing that APOE and the carrier lipid POPC (19x the concentration of NBD-sphinganine in the liposome) are not contributing significantly to lipid toxicity. Diagrams generated in BioRender.

After 72 hours of treatment, CTG was performed for cell viability. POPC and APOE were kept in consistent proportion to the sphinganine. NBD-sphinganine was similarly toxic when delivered freely or in liposomal formulations, regardless of whether the liposomes had APOE bound or not (Figures 8F, 8G, S10C, and S10D). We note that POPC delivered in the same doses, but lacking sphinganine, did not show significant toxicity (Figures S10A and S10B). As POPC was a significantly higher proportion of the liposome than sphinganine, this indicates that bulk lipid application alone does not drive toxicity.

Based on the finding that neurites were NBD^+^, we utilized Incucyte neurite analysis for NBD^+^ (green) neurites to assess lipid incorporation. At multiple doses, liposomal delivery of NBD-sphinganine induced significantly more NBD incorporation into neurites than free NBD-sphinganine delivery (Figures 8B-E, S10A and S10E). APOE did not significantly alter this labelling. This might indicate better incorporation of the lipid into the whole cell or that delivery of the lipid through a different pathway might alter neurite incorporation.

We additionally analyzed neurites by phase microscopy in cells treated with liposomal NBD-sphinganine, as a measure of neuronal health. Consistent with our previous free sphinganine treatment experiment (Figures 7 and S8), there was acute neurite loss in response to sphinganine treatment then a plateau, as dead neurites were not retracted and were still picked up by the analysis software. Surprisingly, the analysis detected more neurite loss in neurons treated with NBD-sphinganine liposomes bound with APOE (Figures S10C and S10D). In both instances, the neurites appeared dead but without APOE the neurite debris stayed in place, whereas there might have been more neurite retraction with APOE. This was a subtle effect picked up by coarse phase analysis, so future studies will need to better characterize the effect and if APOE alters neurodegeneration mechanisms.

## Methods

### iPSC derived cell culture protocols

6-well, 12-well, or 24-well plates were coated using 2.5 mL PDL (1 mg/mL, Sigma P7886) in 47.5 mL borate buffer (100 mM Boric Acid, 75 mM Sodium Chloride, pH 8.4, Boston Bioproducts Inc. C-8852S) overnight at room temperature. Plates were then washed with water three times and allowed to dry at room temperature before plating. PDL coated plates were stored at 4-degrees for a maximum of one month before plating.

BrainXell differentiated microglia were thawed by water-bath and diluted 5 mL in microglial medium, centrifuged at 300 x g for 5 minutes, resuspended in 1 mL microglial media, and counted by hemocytometer or Countess. Cells were plated at specified densities for each assay in microglial media (DMEM/F-12, Gibco 1330-032; N2, Gibco 17502048; B27, Gibco #17504044; NEAA, Gibco #11140-050; Chemically Defined Lipid Mix, Gibco #11905-031; Glutamax, Gibco #35050-061; AA2P solution, Sigma #A8960; 2-Mercaptoethanol, Sigma #M3148; Pen/Strep, Gibco #15140-122; 20 ng/mL M-CSF, Peprotech #300-25; 100 ng/mL IL-34, Peprotech #200-34; 2 ng/mL TGF-β1 Peprotech #100-21C) and only half media changes were ever performed.

Each vial of BrainXell differentiated motor neurons was thawed by water-bath, diluted to 5 mLs with media, then counted by hemocytometer or Countess. Cells were not centrifuged and pipetting up and down was limited. Cells were then plated on day-0 at specified densities in motor neuron seeding media (DMEM/F12, Gibco 1330-032; Neurobasal Medium, Life Technologies #21103-049; B27 supplement, Life Technologies #21103-049; Gibco #17504044; N2 supplement, Gibco 17502048; GlutaMAX, Gibco #35050-061; BrainXell Motor neuron seeding supplement) on PDL coated plates. On day-1, a full media change was performed using the day-0 media, with the addition of growth factors (10 ng/mL BDNF, Peprotech #450-02; 10 ng/mL GDNF, Peprotech #450-10; 1 ng/mL TGF-β1, Peprotech #100-21C; 1:10 of Geltrex diluted 1:10 in DMEM/F-12, Life Technologies #A1413201). On day-4, half media was changed with day-0 media containing BrainXell day-4 supplement instead of seeding supplement and with BDNF, GDNF, and TGF-β1. Day-7 and on, the same media as day-4 was used, but without the BrainXell supplement.

BrainXell astrocyte vials were thawed and diluted with media to 5 mLs and counted by hemocytometer or Countess. Cells were then plated on day-0 at specified densities on PDL coated plates in astrocyte seeding medium (DMEM/F12, Gibco 1330-032; Neurobasal Medium, Life Technologies #21103-049; N2 supplement, Gibco 17502048; GlutaMAX, Gibco #35050-061; BrainXell Astrocyte Supplement; Fetal Bovine Serum, ThermoFisher A3840001). On day-1, media was changed in full (DMEM/F12, Gibco 1330-032; Neurobasal Medium, Life Technologies #21103-049; N2 supplement, Gibco 17502048; GlutaMAX, Gibco #35050-061; BrainXell Astrocyte Supplement), thereafter astrocytes were fed with the same media by half media changes every 3-4 days.

### RNA sequencing

After 72 hours of cell treatment with 3 μM FB1 or Genz-667161, RNA was isolated by Qiagen RNeasy Mini Kit (Qiagen cat# 74104) and sent to Azenta for RNA sequencing. RNA quality control was performed at Azenta using Qubit system. RIN values lower than 6.0 were considered suboptimal. As neuronal samples often had RIN values of lower than 6.0 because of difficulties removing dead cells without compromising healthy neurons, rRNA depletion library preparation was used for neurons followed by Illumina 2 x 150 bp sequencing, ∼30 M reads/sample. Astrocyte and microglial RNA was prepped using Poly-A library preparation by Illumina 2 x 150 bp sequencing, ∼30 M reads/sample.

### RNA sequencing data analysis

RNASeq pre-processing to raw counts was performed by Azenta as previously described. Analysis for differentially expressed genes (DEGs) was performed using the DESeq2 (Love et al., 2014) implemented within Array Studio (Version 11, Omicsoft Corporation, Research Triangle Park, NC, USA). Analyses was restricted to intra cell type comparisons. Each condition was contrasted with the DMSO control. Genes were considered statistically significantly altered if the absolute value of fold change |FC| >2 and adj.pValue < 0.05. Volcano plots were generated using the “EnhancedVolcano” package (Blighe et al. 2023), within the R/RStudio environment (version 3.6.3/version 1.1.463). Data were analyzed through the use of IPA (QIAGEN Inc., https://www.qiagenbioinformatics.com/products/ingenuitypathway-analysis). Data is uploaded to database at Synapse ID# syn53113026.

### Lipidomics Cell preparation

Cells for lipidomics other than labelling were plated at 60k/well in a 24-well dish. For all other experiments astrocytes (20k/96-well, 240k/24-well, 600k/6-well), microglia (40k/96-well, 240k/24-well, 500k/12-well), and motor neurons (40k/96-well, 240k/24-well, and 500k/12-well) were cultured according to BrainXell cell culture protocols outlined above. Cells were then lysed, scraped, and processed for lipidomics with a protocol adapted from Mitchell et al. (2015). Briefly, cells were lysed and scraped in lipidomics lysis buffer (10 mM HEPES, 10 mM NaCl, 1 mM KH2PO4, 5 mM NaHCO3, 5 mM EDTA, 1 mM CaCl2, and 0.5 mM MgCl2), lysed using bead tissue lyser or by 15 x shearing through a syringe, and 25ul of 20% sucrose was added per 500 μl of lysate. Lysate was then centrifuged at 6,300 x g at 4-degrees for 14 minutes. Supernatant formed the cytosolic fraction. The pellet was washed, centrifuged, then the nuclear pellet was resuspended and stored at −80 degrees Celsius until analysis.

### Ceramide and dHCer profiling

Cytoplasm or nuclear fraction (50 μL) was extracted with 1mL of extraction solution containing internal standard (methanol/acetonitrile/water, 80/15/5 %, v/v/v containing 10ng/mL d35Cer16 as IS). The mixture was vortexed for 15 minutes in multi-tube vortex and spined down for 8 minutes at 8400g. The supernatant (200 μL) was carefully transferred to a MS vial for analysis. Standard curves (0.03-1000 ng/mL) were prepared in the same way as samples with Ceramide standards: Ceramide_C14, Ceramide_C16, Ceramide_C18, Ceramide_C24, Ceramide_C24:1, and dH-Ceramide_C16.

LC-MS/MS analysis was conducted on a Waters Acquity UPLC system coupled with a Sciex QTRAP6500 mass spectrometer using multiple-reaction monitoring (MRM). Ceramide profile is separated by a Waters Acquity UPLC BEH C8 (100×2.1mm, 1.7um particles, cat#186002878) at a flow rate of 0.75 mL/min. The column is heated at 60C during the run. The mobile phase consists of A) 0.2% Formic Acid; 5 mM Ammonium formate in DI water and B) 0.2% Formic Acid; 5 mM Ammonium formate in (50:50) Methanol:Acetonitrile. The injection volume was 5 μL and the total runtime was 6 min. The step gradient was as follows: 0–0.2 min, 85 % solvent B; 0.2–1.8 min, 85 to 98 % solvent B; 1.81-3.4 min, 100% solvent B, 3.4–3.5 min, 100 to 85 % solvent B; 3.5–6 min 85 % solvent B.

The ESI+ source temperature was 450 °C; the ESI needle was 4,500 V; the declustering potential was 60 V; the entrance potential was 10 V; and the collision cell exit potential was 10 V. The collision and curtain gas were set at medium and 15, respectively. GS1 and GS2 were set at 80 and 20. The collision energy was 34 eV for Ceramide and 40 eV for dHCer. For MRM, the dwell time was set at 50 ms for each of the signal from transitions.

Data were acquired and analyzed by MultiQuant 3.0 (AB Sciex). Calibration curves were constructed by plotting the corresponding peak area ratios of analyte/internal standard versus the corresponding analyte concentrations using 1/x weighing linear regression analysis.

### qPCR analysis

Cells were washed two times with half media removal and addition of PBS, then complete volume was removed, and direct Cells-to-Ct lysis buffer (Invitrogen, A25602) was added, and lysates frozen at −80 degrees. Quantitative PCRs were performed according to kit instructions using ThermoFisher taqman gene expression assays with GAPDH probes as loading controls on a QuantStudio 7 Flex Real-Time PCR System. Data was analyzed using -deltadeltaCt relative to MN expression levels and alternatively -deltaCt to relative gene expression.

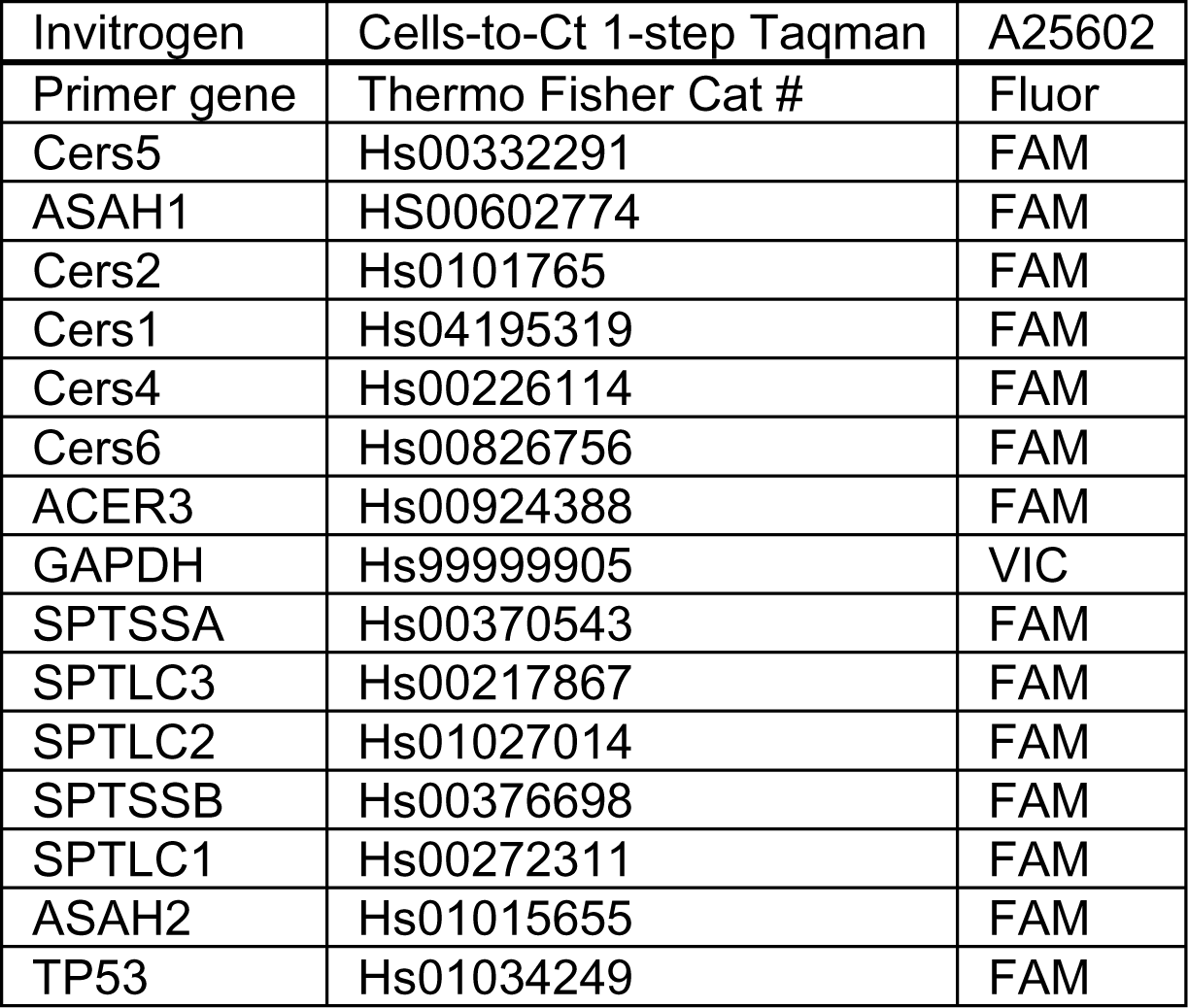

### Ceramide synthesis assays

Cells were lysed and scraped on ice using HEPES buffer (20 mM HEPES, 25 mM KCl, 250 mM Sucrose, 2 mM MgCl2) made according to kit (640011, 640012, 640014, Avanti Polar Lipids Inc.) instructions (Tidhar et al., 2015). Lysates, NBD-sphinganine (810206P, Avanti Polar Lipids Inc.), and Acyl-CoA (18:0, 870718P, Avanti Polar Lipids Inc.; 16:0, 870716P, Avanti Polar Lipids Inc.; 24:1, 870725P, Avanti Polar Lipids Inc.) for reaction were water-bath sonicated prior to the reaction. All reagents were kept warm at 37-degrees prior to the start of the reaction. Reactions were performed on a thermal cycler for temperature consistency and reactions were neutralized using methanol containing 1% formic acid. Lipid column (8E-S001-BGB, Phenomenex) washes and elutions were performed according to manufacturer instructions (640011, 640012, 640014, Avanti Polar Lipids Inc.) using a vacuum manifold (AH0-8950, Phenomenex) and NBD-ceramide levels were read by FlexStation 3 microplate reader.

### d17:0-sphinganine labelling and ceramide measures

Stock d17:0-sphinganine (860654P, Avanti Polar Lipids Inc.) was warm water-bath sonicated at 37-degrees Celsius for 15-20 minutes prior to addition to cells at 1 μM for 2 hours or 24 hours. For inhibitor conditions, FB1 (Sigma-Aldrich, F1147) was added at 3 μM and pre-incubated for 1-hour prior to d17:0-sphinganine addition.

### Preparation of NBD-sphinganine/POPC liposomes

We made a mixture of POPC (16:0-18:1 PC; Avanti 850457) and NBD-sphinganine (Avanti 810206P) with 95% POPC and 5% of NBD-sphinganine in 100% ethanol. Lipids were then added to PBS at a 1:9 ratio of ethanol/lipid solution to PBS. Components were vortexed to mix components, then incubated at 37-degrees to form mixture. Resultant mixtures were freeze/thawed 3-times, rapidly on dry ice then room temperature to increase the uniformity of the liposomes in the mixture. We then used the lipid solution to resuspend APOE (abcam ab280330) at a ratio of 3 μg of APOE for every 10 μg of lipid in the NBD-sphinganine:POPC liposomes (equal volume of POPC liposomes were added to APOE for controls). All dosing was then performed based on the molarity of NBD-sphinganine added to cells, or equal volumes of similarly prepared control solutions.

### Reagent List

**Reagent List:**
|  |  |  |
| --- | --- | --- |
| Cells-to-Ct 1-step Taqman | A25602 |  |
| Cells-to-Ct 1-step Taqman | A25602 | Invitrogen |
| Cers5 FAM taqman primer | Hs00332291 | ThermoFisher |
| ASAH1 FAM taqman primer | HS00602774 | FAM |
| Cers2 FAM taqman primer | Hs0101765 | FAM |
| Cers1 FAM taqman primer | Hs04195319 | FAM |
| Cers4 FAM taqman primer | Hs00226114 | FAM |
| Cers6 FAM taqman primer | Hs00826756 | FAM |
| ACER3 FAM taqman primer | Hs00924388 | FAM |
| GAPDH VIC taqman primer | Hs99999905 | VIC |
| SPTSSA taqman primer | Hs00370543 | FAM |
| SPTLC3 taqman primer | Hs00217867 | FAM |
| SPTLC2 taqman primer | Hs01027014 | FAM |
| SPTSSB taqman primer | Hs00376698 | FAM |
| SPTLC1 taqman primer | Hs00272311 | FAM |
| ASAH2 taqman primer | Hs01015655 | FAM |
| TP53 taqman primer | Hs01034249 | FAM |
| Strata C18-E 96/well lipid binding column | 8E-S001-BGB | Phenomenex |
| Vacuum manifold | AH0-8950 | Phenomenex |
| Fumonisin B1 | F1147 | Sigma-Aldrich |
| NBD-sphinganine | 810206P | Avanti |
| Acyl-CoA 18:0 | 870718P | Avanti |
| Acyl-CoA 16:0 | 870716P | Avanti |
| Acyl-CoA 24:1 | 870725P | Avanti |
| Methanol | 34860 | SIGMA |
| Formic Acid | FX0440 | EMD Millipore |
| HPLC grade water | W5-4 | Fishersci |
| Ammonium Acetate | A1542 | SIGMA |
| MicroAMP Opical 96-well Plate | N8010560 | Life Technologies |
| Celigo |  | Nexcelom |
| Plate reader | FlexStation3 | Molecular Devices |
| Incucyte SX5 |  | Sartorius |
| borate buffer (100 mM Boric Acid, 75 mM Sodium Chloride, pH 8.4) | C-8852S | Boston Bioproducts Inc. |
| PDL (1 mg/mL) | P7886 | SIGMA |
| Hu APOE | ab280330 | Abcam |
| POPC | 850457 | Avanti |
| DMEM/F-12 | 1330-032 | Gibco |
| N-2 supplement | 17502048 | Gibco |
| B-27 supplement | 17504044 | Gibco |
| NEAA | 11140-050 | Gibco |
| Chemically Defined Lipid Mix | 11905-031 | Gibco |
| Glutamax | 35050-061 | Gibco |
| AA2P | A8960 | Sigma |
| 2-Mercaptoethanol | M3148 | Sigma |
| Pen/Strep | 15140-122 | Gibco |
| M-CSF | 300-25 | Peprotech |
| IL-34 | 200-34 | Peprotech |
| TGF-b1 | 100-21C | Peprotech |
| Neurobasal Medium | 21103-049 | Life Technologies |
| GlutaMAX | 35050-061 | Gibco |
| BDNF | 450-02 | Peprtech |
| GDNF | 450-10 | Peprtech |
| Geltrex | A1413201 | Life Technologies |
| Fetal Bovine Serum | A3840001 | ThermoFisher |
| Qiagen RNeasy Mini Kit | 74104 | Qiagen |
| UPLC BEH C8 | 186002878 | Waters Acquity |
| CerS activity kits | 640011,<br>640012,<br>640014 | Avanti Polar Lipids |
| d17:0-sphinganine | 860654P | Avanti Polar Lipids |

## Discussion

Diseases of the CNS are often characterized by glial reactivity and the selective vulnerability of neuronal and oligodendroglial cells to cell-death. The nature of the cellular dysfunction and death, as well as the vulnerable cell types, underly each disease’s symptomology. Ceramide is a sphingolipid essential to normal cellular functions but is also involved in cell-death pathways. Despite the variable dysfunctions and cell types affected in Alzheimer’s disease, multiple sclerosis, ALS, and Farber disease, CNS ceramide accumulation has been observed in each of these diseases (Han et al., 2002). Ceramide also accumulates during normal process of aging and is associated with cognitive decline. Cellular stress and disease can induce ceramide accumulation in mitochondrial membranes, of interest since *de novo* ceramide biosynthesis occurs at the ER-mitochondria associate membranes (MAMs) (Planas-Serra et al, 2023). At high concentrations, ceramides form pores in mitochondrial membranes, releasing cytochrome C, and contributing to the initiation of cell death. This and other important functions of ceramide in ER function, membrane trafficking and rafting, as well as metabolism, likely explain why disrupted ceramide homeostasis is associated with a variety of CNS diseases.

### Basal regulation of ceramide levels

Despite the known role of ceramide in neurodegeneration and the possibility that astroglial ceramides are markers of neuroinflammation, how different cell types of the CNS regulate ceramide has not been fully elucidated (de Wit et al., 2019; Dodge, 2017). Efforts have been made to characterize mouse cell-type specific lipid regulation, but this did not focus on ceramides (Fitzner et al., 2020). Our understanding of ceramide alterations in disease largely derives from lipidomics analysis of whole tissue lysates and CSF, or transcriptomic observations (Barrier et al., 2010; Fernández-Beltrán et al., 2021; Henriques et al., 2017; Pousinis et al., 2020). Additionally, some essential signaling mechanisms related to ceramide have been determined in cancer cells (Fekry et al., 2018).

We utilized healthy human iPSC-derived cells to determine how ceramides are regulated differently across CNS cell-types. There is evidence from other works that disease relevant changes in lipid profiles can be found in iPSC-derived cells (Lieberman et al., 2022; Tomasello et al., 2022). We found that neurons had higher levels of C18-ceramide than other cells, consistent with *in vivo* studies (Ginkel et al., 2012). In many cases, our RNA data did not match well with activity and lipidomics data. It is well known that RNA signatures don’t always match protein expression (Tasaki et al., 2022). CerS expression may not directly correlate with CerS activity as multiple proteins are known to modulate CerS activity, some through post-translational modifications (Sassa et al., 2016; Tomasello et al., 2022). Future studies will need to determine if the discrepancies identified here are due to translational regulation, post-translational modifications, or similar.

### Assessing ceramide production across cell-types

While we could not address how cell-type specific CerS post-translational modifications altered cellular regulation of ceramides, we determined the functional lipid output of CerS in each cell-type. Upon addition of sphinganine, a precursor for ceramide synthesis, it was notable that each cell-type largely turned the sphinganine precursor into ceramide chain lengths predicted from our CerS activity assays. Astroglial CerS5/6 generating Cer16, microglia CerS2/5/6 generating Cer16 and Cer24:1, and motor neurons (MNs) CerS1/4 generating Cer18 and to a lesser extent Cer16. It’s notable that MNs treated with precursor were able to generate detectable labelled C18-ceramide, but it was a significantly smaller percentage of the overall ceramide profile than the labelled ceramides in the glial cells. Additionally, MN ceramide profile was minimally altered by FB1 treatment (pan-CerS inhibitor), which seems to indicate that MN do not rely largely on de novo ceramide synthesis to function.

Sphinganine addition allowed us to increase ceramide precursor. An alternative way to determine relative ceramide production, is to block complex glycosphingolipid production downstream of ceramide. In our experiments we used a GCSi, which blocks the generation of glucosylceramide from ceramide and also affects disease progression in preclinical models of ALS, PD and AD (Sardi et al., 2017). Notably, after GCSi treatment iPSC-derived neurons and glia accumulated similar ceramide species that occurred in response after the addition of the sphinganine precursor. This provided further confirmation that homeostatic regulation of certain ceramide species differs amongst astrocytes, microglia and neurons. Additionally, GCSi induction of ceramide may explain exacerbation of disease phenotypes for some neurodegenerative diseases (Dodge et al., 2015).

### Identification of pathways dependent on ceramide levels via RNA-sequencing

Determining the ceramides that each cell type produces in response to increased precursor and inhibited downstream synthesis was key to understanding cellular metabolic differences. We wanted to further understand how altered ceramide levels influence the functions of different cell-types by analyzing changes in RNA expression after FB1 and GCSi treatment. Surprisingly, but consistent with our other data, MNs had no statistically significant gene changes in response to FB1 treatment. Astrocytes and microglia both had >1.5k differentially expressed genes (DEGs) in response to FB1. This seems to indicate that ceramide synthesis occurs largely in glial cells over these time scales, and that MNs do not synthesize large amounts of ceramide *de novo*, nor show significant responses to inhibition of ceramide synthesis. In contrast to this finding, all three cell-types had many significant DEGs after GCSi treatment. This possibly indicates that while MNs do not rely heavily on ceramide synthesis, they do respond to small increases in ceramide accumulation and/or inhibition of glucosylceramide synthesis.

We additionally assessed pathways impacted by FB1 and GCSi treatment, and commonalities between cell-type response to FB1 and GCSi. Ceramide alterations resulted in changes to both lipid homeostasis and inflammatory activity. This could underlie some of the lipid, cholesterol, and inflammatory changes characteristic of neurodegenerative diseases. We note that these changes were accompanied by specific responses such as increased pro-myelination and oligodendrocyte survival signals (*IL11* and *CLCF1*) after GCSi treatment and decreased lipid self-antigen presentation (*CD1B*) (Gurfein et al., 2009; Ji-Wei et al., 2022; Shahine, 2018; Zhang et al., 2006). This could indicate that ceramide accumulation acts as a signal of myelin dysfunction leading to an adaptive response. This could potentially extend to signals regulating axon growth and synapses as well, because FB1 treatment of astrocytes and microglia decreased expression of genes involved in axon growth and synaptic development (*CBLN2* and *TMEM108*), and we saw alterations in axon guidance pathways in astrocytes after FB1 treatment. This latter finding also likely highlights that the homeostatic regulation of ceramide within an optimal range is critical for the normal function of the CNS – too little or too much ceramide likely leads to a cacostatic state. Indeed, LOF mutations in DEGS1 provoke lower ceramide and increased DhCer levels, leading to childhood onset leukodystrophy (Pant et al., 2019).

Notably, some of the changes in response to FB1 and GCSi treatment mirrored changes found in neurodegenerative diseases with altered ceramide homeostasis. CCL2, a cytokine that is elevated in mouse models of Farber disease (Dworski et al., 2017), decreased after FB1 treatment and increased after GSCi. Similar changes were observed for the biomarker *MMP10*, which is predictive of AD symptom progression (Martino Adami et al., 2022), and *SREBF1* which has variants that are associated with PD risk (Ivatt et al., 2014). IL-17 signaling was similarly changed by altered ceramides and is known to be dysregulated in Alzheimer’s disease, Parkinson’s disease, and multiple sclerosis (Brigas et al., 2021; Chen et al., 2020; Gautam et al., 2023).

There were also indications that ceramides are essential to the functions of a mitochondrial, ER stress, lipid dysregulation, and apoptosis axis (*STC2*, *CHAC1, ALDH1L2*, *NMNAT2, PDK4, KLHDC7B, BIRC3*, *PLN, SREBF1*). Ceramide is synthesized in the ER, at the mitochondria associated membranes (Planas-Serra et al, 2023), regulates mitochondrial function and dynamics, can induce mitochondrial pores in programmed cell death, and are essential to brain development as well as long term neuronal and glial health. Thus, ceramide dysregulation likely plays an important role in the ER and mitochondrial dysfunction present in many neurodegenerative diseases. Conversely, dysregulation of ceramides could also induce adaptive responses through increased expression of genes that promote regenerative mechanisms such as remyelination. Our data indicate that ceramides are functionally connected to metabolic, lipid and inflammatory pathways relevant for neurodegenerative diseases. It will need to be determined if manipulating ceramides in preclinical neurodegenerative models can alter these biomarkers and inflammatory factors in ways that modify disease progression.

### Understanding differences in cell-type sensitivity to sphinganine/ceramide toxicity

Given our RNA expression findings, we had significant questions surrounding how ceramide dysregulation would alter cell survival. In certain disease contexts, astrocytes can take on a reactive state that is toxic to neurons and oligodendrocytes (Liddelow et al., 2017). The precise identity and source of the toxicity remains to be determined and is likely context dependent, but there is evidence that the ceramide synthesis pathway could be involved (Arredondo et al., 2022; Chung et al., 2023; Guttenplan et al., 2021; Victor et al., 2022; Wieder et al., 2023). This is consistent with evidence that glial cells can produce saturated lipids (sphingolipids?) that are toxic to neurons and that astrocytes may even release ceramide extracellularly in response to disease stimuli, triggering neuronal cell-death (Guttenplan et al., 2021; Wang et al., 2012).

Based on our findings of ceramide generation from precursors, we treated each cell type with escalating doses of sphinganine to see if the differences in ceramide handling between cell-types also altered vulnerability to toxicity. Astrocytes synthesized the largest proportion of ceramides using labelled sphinganine precursor, so it was notable that astrocytes also showed the most resistance to sphinganine toxicity. Microglia and neurons both had decreased viability starting at 10 μM sphinganine. Of interest, our live imaging revealed that sub-toxic doses of sphinganine led to acute morphological changes in microglia that then normalized with time (Supplemental Video 2). Microglia may respond to toxic doses of lipid with morphologic changes and resolve the stress in some cases (Figure S4E), and in other cases the toxic dose of lipid may trigger cell-death (Figure 7I). Future studies will need to determine whether morphological changes represent an acute inflammatory response to sphinganine and whether similar acute responses occur in neurons and astrocytes at sub-toxic doses. Notably, a similar morphological alteration is seen by others in response to oleic acid and that this added microglial lipid burden may contribute to neuronal dysfunction (Victor et al., 2022).

While addition of sphinganine is a method of increasing ceramide and assessing toxicity, we are adding significant excess lipid. We therefore used a well characterized inhibitor of glucosylceramide synthase to see if ceramide would accumulate and if there was associated toxicity. With this inhibitor treatment, we did find accumulation of certain ceramides. Notably, the ceramide chain lengths that accumulated in each cell type tended to be those generated by each cell type’s most active CerS. We also found a significant impact on cell viability with GCSi treatment in all three cell types. Paralleling our findings with sphinganine treatment, astrocyte viability was not impacted to the same extent as microglia and MNs with GCSi treatment. RNA sequencing of cells treated with GCSi revealed a similar number of DEGs in both MNs and astrocytes, but the extent of the changes was larger in astrocytes.

Given the central role of ceramides in the synthesis of more complex lipids and the known function of astrocytes in providing lipids to neurons, we sought to deliver sphinganine to neurons in a more physiological manner in order to address how this would alter trafficking and toxicity (Guttenplan et al., 2021; Qi et al., 2021). It is known that lipids are often transferred in lipoparticle/APOE mediated form. Additionally, ceramides can be found in blood derived high density lipoprotein (HDL), indicating that ceramides can be packaged into such apolipoprotein particles (Argraves et al., 2011). We therefore adapted our sphinganine delivery into a liposomal formulation as a rough mimic of HDL delivery. Our liposomal preparations had minimal impact on cellular viability when just the carrier lipid (POPC) and APOE were delivered. As POPC made up ∼95% of the liposomes, this indicated that simple bulk lipid addition to MNs doesn’t drive toxicity. Sphinganine drove toxicity regardless of free delivery or delivery by liposome. Interestingly, the liposomal delivery did introduce more lipid label into neuronal neurites independent of APOE status. When sphinganine liposomes were delivered bound to APOE, neurites did appear to degenerate more, despite no changes in measured toxicity. This indicates that the method of lipid delivery matters, but it will be important to understand how this alters physiology. It’s probable that other aspects of neuronal function (e.g., ER stress response) are dependent on the format of lipid exposure. Additionally, more chronic treatments might reveal differences in toxicity, whereas these could have been missed with these acute toxic doses.

### Interpretation of bulk CNS lipidomics

Our findings have implications for interpreting bulk lipidomics of tissue in animal models and patients. Specially, we found that cortical neurons, MNs, astrocytes and microglia feature unique cytoplasmic and nuclear ceramide composition. Thus, disease reported changes in ceramide species may suggest the loss or enrichment of a particular cell type. For example, the accumulation of C24 and C24:1 ceramide isoforms in the spinal cords of ALS patients (Dodge et al., 2015) may be indicative of increased glial activation.

Additionally, our RNA-seq data indicate that changes in lipid signatures can alter cytokines and chemokines. CCL2 and similar factors can alter immune cell infiltration, possibly altering the cellular milieu through increased CNS immune cells. Our data show that microglia differ significantly from other CNS cell-types in ceramide signature. It is probable that sufficient immune cell infiltration could alter lipidomic signatures with consequences for interpreting tissue lipidomic signatures in stroke, multiple sclerosis, and other neurodegenerative diseases.

Based on our *in vitro* labelling data that revealed acute lipid changes separate from iPSC differentiation effects, it makes sense for more studies to explore *in vivo* labeling approaches. It is likely that lipids generated prior to disease onset or through alternative synthesis pathways often create noise when attempting to determine disease and therapeutic effects in pre-clinical models. Labeling would allow both isolation of defined timepoints and pathways of lipid synthesis.

### Conclusions

Together our data reinforce the notion that glia are the lipid handling cells of the brain, with astrocytes having particularly large ceramide synthesis capacity. Microglia did have significant synthesis capacity as well and uniquely synthesized longer chain ceramides. Astrocytes were differentially resilient to ceramide/sphinganine related toxicity, which might be associated with their function in synthesizing lipids and transferring them to other cell-types in the brain. Future studies are necessary to extend these findings *in vivo*. Additionally, understanding selective lipotoxicity vulnerable cell-types and cell-type specific lipid synthesis, could contribute significantly to our understanding of neurodegenerative diseases with lipid dysregulation. It may be important to measure both free and apolipoprotein associated lipids to better understand contexts where lipids become toxic in disease.

## Supporting information

Venn_List_Astrocyte FB1 GCSi changes

Venn_List Microglia genes FB1-down GCSi-up

MN astrocyte microglia common GCSi gene changes

Pathway analysis MNs astrocytes microglia FB1 and GCSi

Document1_Venn_List Astrocyte FB1-up GCSi-down

Document2_Venn_List Astrocyte FB1-down GCSi-up

Venn_List_FB1 Astrocyte Microglia common gene changes

Microglia FB1 and GCSi gene changes

## Acknowledgements

We thank Pablo Sardi and Aurora Pujol for their input on the manuscript, and BrainXell for technical support.

## Notes

### Competing Interest Statement

All authors were employees of Sanofi at the time of the work

