## Supplemental Figures for "Unravelling neuronal and glial differences in ceramide composition, synthesis, and sensitivity to toxicity"

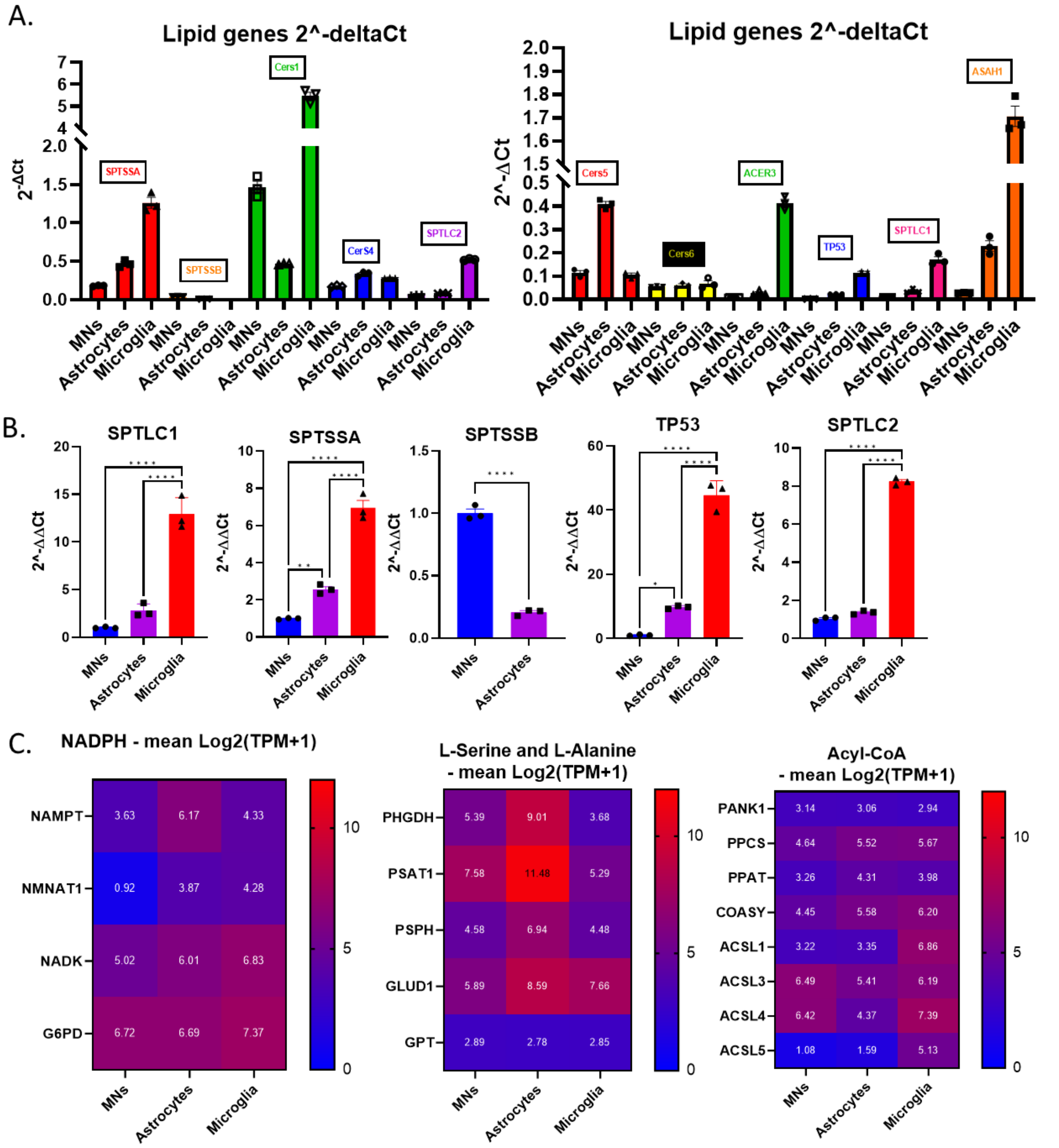


**Supplemental Figure 1. qPCR -2$^{-\Delta Ct}$ and -2$^{-\Delta\Delta Ct}$ characterization of ceramide synthesis pathway in iPSC-derived motor neurons (MNs), astrocytes, and microglia. A and B)** iPSC-derived cells were lysed and analyzed by taqman qPCR for the expression of genes involved in ceramide synthesis and ceramidases. **A)** Expression was analyzed by -2$^{-\Delta Ct}$ for a rough comparison of expression across different genes. **B)** Additional -2$^{-\Delta\Delta Ct}$ analysis comparing gene expression across cell types, SPTSSB was undetectable in microglia (extension of Figure 2). All gene expression was normalized to GAPDH controls. (One-way Anova, Tukey's multiple comparisons test, $^{*}p$<0.05, $^{**}p$<0.01, $^{****}p$<0.0001; $n$ = 3 replicate wells, 3 qPCR reactions per well). **C)** Heat maps of the ceramide synthesis anteome, an extension of Figure 2 A-F.

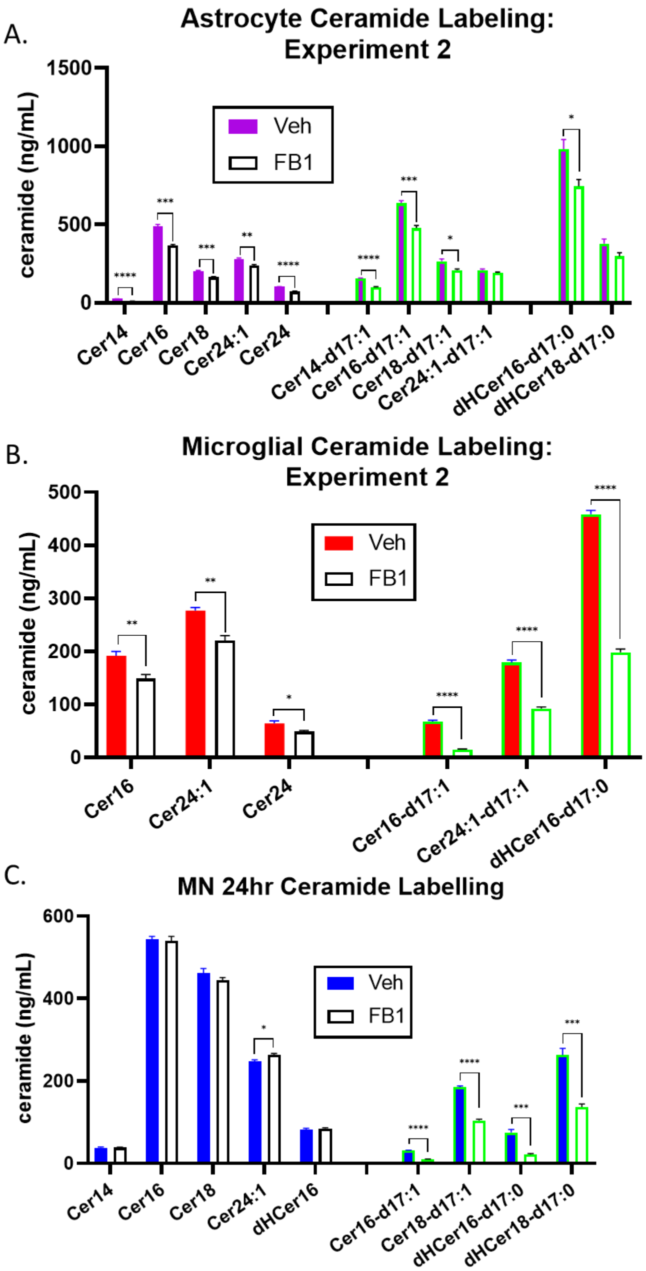


**Supplemental Figure 2. Ceramide Profiles in iPSC-derived glia and motor neurons (MNs) labeled with d17:0-sphinganine. A-C)** By introducing a traceable sphinganine (d17:0) precursor, in the presence or absence of 3 μM FB1, we were able to track the active CerS-dependent ceramide synthesis in each cell type over a 2-hour period. **A and B)** Second experimental repeats of 2-hour labelling in astrocytes and microglia. **C)** MNs had minimal labelling after exposure to the d17:0-sphinganine for 2 hours, so we performed labeling for 24-hours to determine which ceramides would be produced over a longer time-period (Comparisons of each ceramide species by Student's T-test, *****P*<0.0001, ****P*<0.001, ***P*<0.01 **P*<0.05; *n* = 4 wells).

| Percent reduction by FB1 treatment | | | | | |
|---|---|---|---|---|---|
| lipid | Microglia 2hr | Astrocytes 2hr | MN 2hr | | MN 24hr |
| Cer14 | - | - | 0.75 | | -1.32 |
| Cer16 | 27.73 | *9.53 | -2.96 | | 0.75 |
| dHCer16 | - | - | -3.75 | | -4.62 |
| dHCer16-d17:1 | *69.91 | *45.48 | 11.40 | | *71.79 |
| Cer16-d17:1 | *100.00 | *26.22 | - | | *68.32 |
| Cer18 | - | *58.05 | -4.19 | | 3.59 |
| Cer18-d17:1 | - | - | - | | *43.52 |
| dHCer18-d17:1 | - | *100.00 | *47.58 | | *47.96 |
| Cer24:1 | 22.59 | 9.95 | -5.90 | | *-6.53 |
| Cer24:1-d17:1 | *46.53 | -9.19 | - | | - |
| Cer24 | 22.33 | - | - | | - |

| Percent labelled | | | | | |
|---|---|---|---|---|---|
| lipid | Microglia 2hr | Astrocytes 2hr | MNs 2hr | | MNs 24hr |
| Cer14 | - | - | 0.00 | | 0.00 |
| Cer16 | 15.35 | 41.81 | 0.00 | | 5.50 |
| dHCer16 | 100.00 | 100.00 | 54.72 | | 47.99 |
| Cer18 | - | 0.00 | 0.00 | | 28.62 |
| dHCer18 | - | 100.00 | 100.00 | | 100.00 |
| Cer24 | 0.00 | - | - | | - |
| Cer24:1 | 26.54 | 13.59 | 0.00 | | 0.00 |

**Supplemental Figure 3. Calculations of the percent of d17:0-sphinganine labeling and the percentage of labeling inhibited by FB1 treatment.** By introducing a traceable sphinganine (d17:0) precursor, in the presence or absence of 3 μM FB1, we were able to track the active CerS-dependent ceramide synthesis in each cell type over 2-hours or 24-hours. We chart both the reduction in lipids (labeled and unlabeled) in response to FB1 and the percent of each lipid that was labeled in cells not treated with FB1.

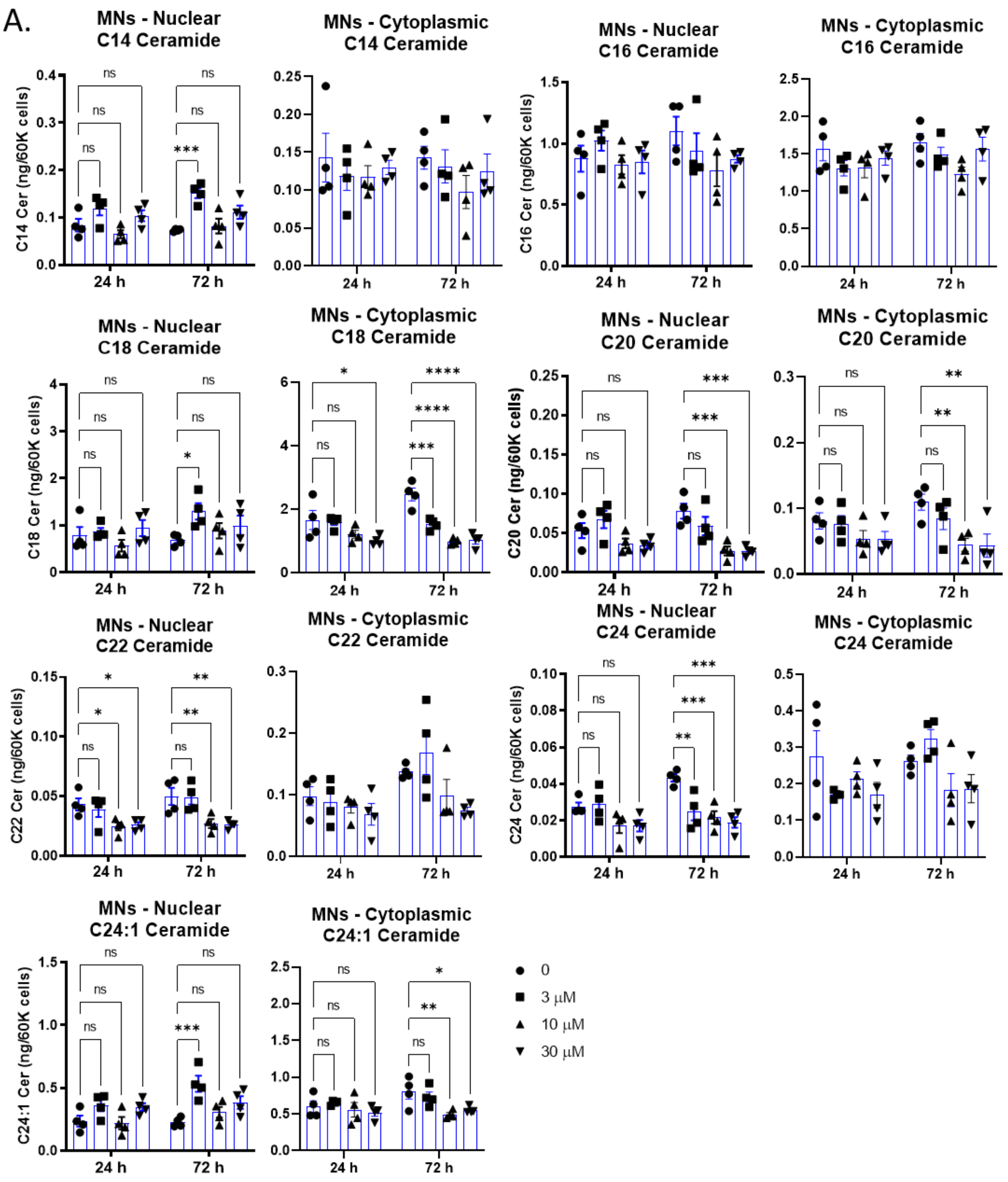

A.
MNs - Nuclear C14 Ceramide
MNs - Cytoplasmic C14 Ceramide
MNs - Nuclear C16 Ceramide
MNs - Cytoplasmic C16 Ceramide
MNs - Nuclear C18 Ceramide
MNs - Cytoplasmic C18 Ceramide
MNs - Nuclear C20 Ceramide
MNs - Cytoplasmic C20 Ceramide
MNs - Nuclear C22 Ceramide
MNs - Cytoplasmic C22 Ceramide
MNs - Nuclear C24 Ceramide
MNs - Cytoplasmic C24 Ceramide
MNs - Nuclear C24:1 Ceramide
MNs - Cytoplasmic C24:1 Ceramide
C14 Cer (ng/60K cells)
C16 Cer (ng/60K cells)
C18 Cer (ng/60K cells)
C20 Cer (ng/60K cells)
C22 Cer (ng/60K cells)
C24 Cer (ng/60K cells)
C24:1 Cer (ng/60K cells)
24 h
72 h
0
3 µM
10 µM
30 µM

B.

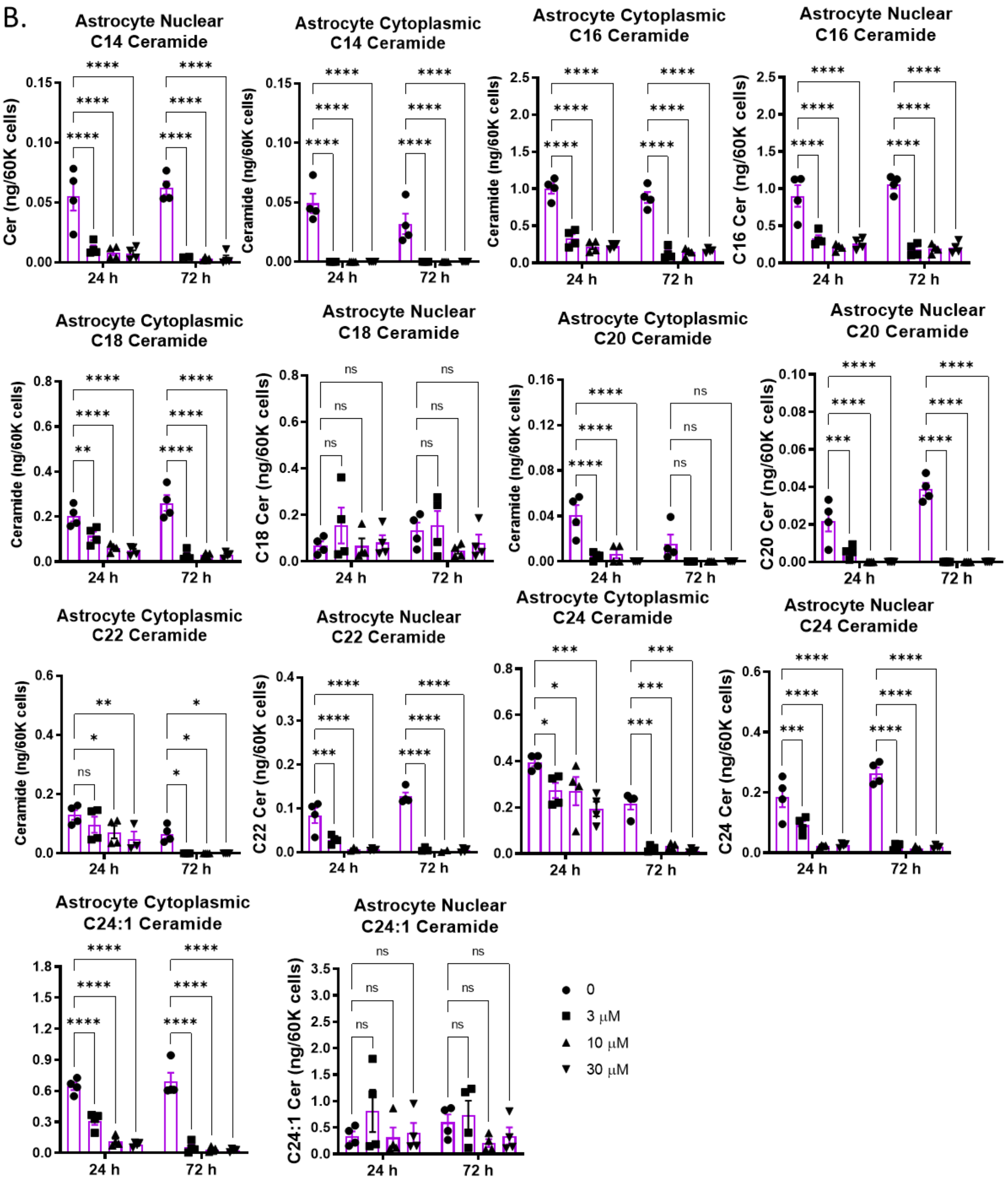
Astrocyte Nuclear C14 Ceramide
Cer (ng/60K cells)
Astrocyte Cytoplasmic C14 Ceramide
Ceramide (ng/60K cells)
Astrocyte Cytoplasmic C16 Ceramide
Ceramide (ng/60K cells)
Astrocyte Nuclear C16 Ceramide
C16 Cer (ng/60K cells)
Astrocyte Cytoplasmic C18 Ceramide
Ceramide (ng/60K cells)
Astrocyte Nuclear C18 Ceramide
C18 Cer (ng/60K cells)
Astrocyte Cytoplasmic C20 Ceramide
Ceramide (ng/60K cells)
Astrocyte Nuclear C20 Ceramide
C20 Cer (ng/60K cells)
Astrocyte Cytoplasmic C22 Ceramide
Ceramide (ng/60K cells)
Astrocyte Nuclear C22 Ceramide
C22 Cer (ng/60K cells)
Astrocyte Cytoplasmic C24 Ceramide
Astrocyte Nuclear C24 Ceramide
C24 Cer (ng/60K cells)
Astrocyte Cytoplasmic C24:1 Ceramide
Astrocyte Nuclear C24:1 Ceramide
C24:1 Cer (ng/60K cells)
24 h
72 h
0
3 µM
10 µM
30 µM

C.

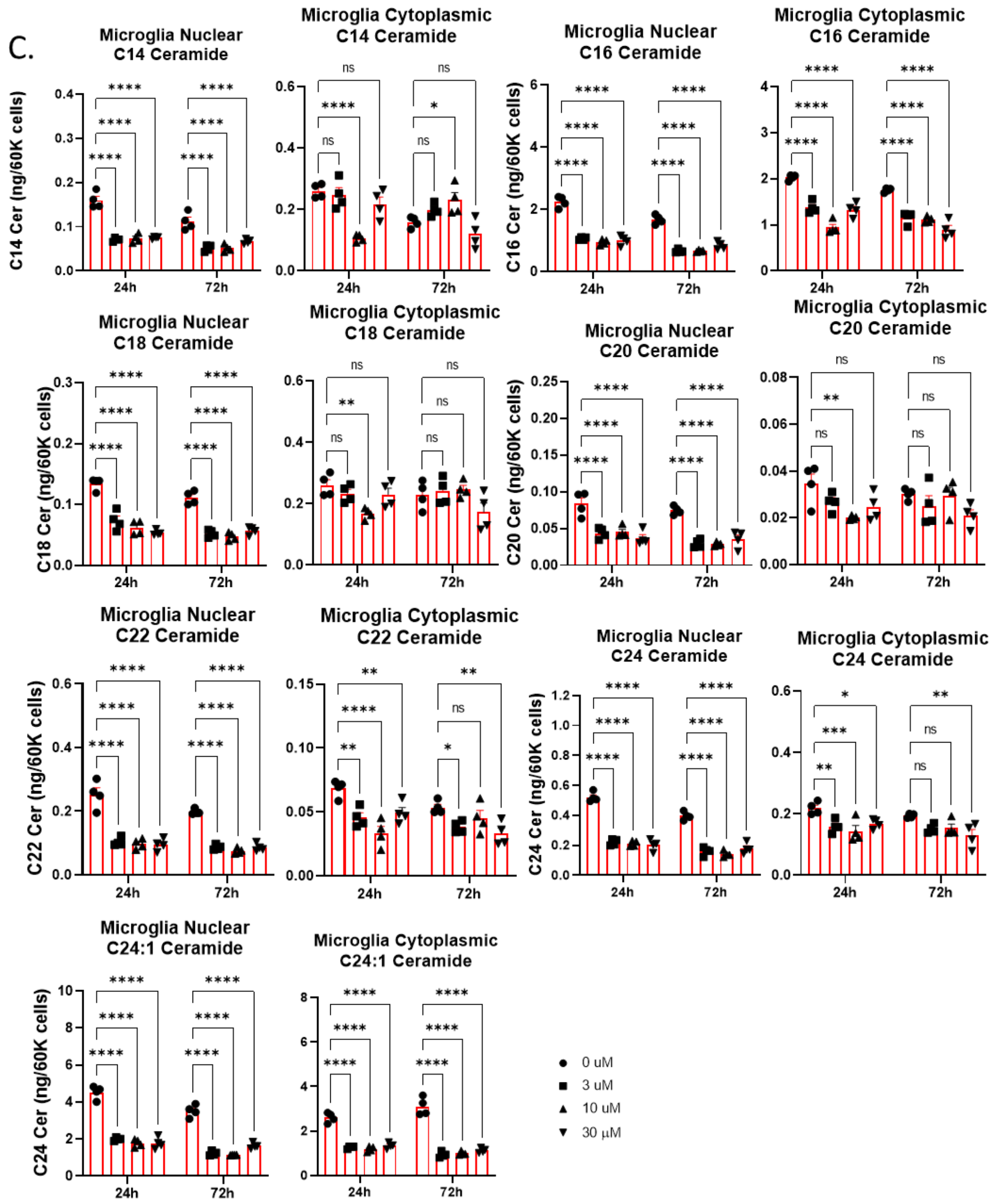


**Supplemental Figure 4. Ceramide Profiles of iPSC derived neural cell types treated with FB1.** Astroglial and microglial ceramide profiles were more significantly impacted by CerS inhibition than motor neurons. **A)** Lipidomics from cytoplasmic and nuclear fractions of motor neurons (MNs) treated with 0-30 µM FB1. **B)** Lipidomics from cytoplasmic and nuclear fractions of astrocytes treated with 0-30 µM FB1. **C)** Lipidomics form cytoplasmic and nuclear fractions of microglia treated with 0-30 µM FB1. (*****P*<0.0001, ****P*<0.001, ***P*<0.01 **P*<0.05; 2-way ANOVA; *n* = 4 replicate wells).

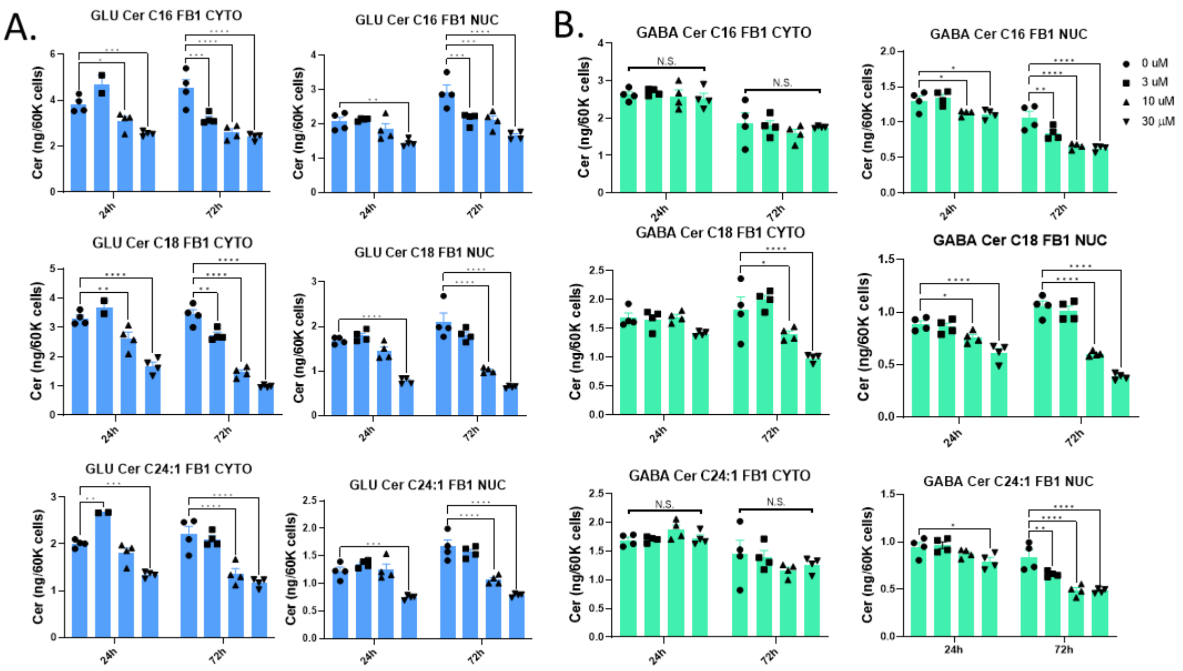


**Supplemental Figure 5. Ceramide Profiles of iPSC-derived cortical neuronal cell-types treated with FB1.** Glutamatergic **(A)** and GABAergic **(B)** cortical neuronal cytoplasmic and nuclear ceramide profiles after treatment with 0-30 µM FB1. (*****P*<0.0001, ****P*<0.001, ***P*<0.01 **P*<0.05; 2-way ANOVA; *n* = 4 technical replicates).

A.

| IPA pathway changes in astrocytes after FB1 treatment | | | |
|---|---|---|---|
| | Ingenuity Canonical Pathways | -log(p-value) | Ratio |
| 1 | ABRA Signaling Pathway | 9.21E+00 | 2.50E-01 |
| 2 | Hepatic Fibrosis / Hepatic Stellate Cell Activation | 8.76E+00 | 1.75E-01 |
| 3 | Agranulocyte Adhesion and Diapedesis | 6.30E+00 | 1.48E-01 |
| 4 | Pulmonary Fibrosis Idiopathic Signaling Pathway | 6.18E+00 | 1.26E-01 |
| 5 | Neuroinflammation Signaling Pathway | 6.08E+00 | 1.26E-01 |
| 6 | Calcium Signaling | 5.40E+00 | 1.36E-01 |
| 7 | Osteoarthritis Pathway | 5.21E+00 | 1.31E-01 |
| 8 | Ferroptosis Signaling Pathway | 4.98E+00 | 1.59E-01 |
| 9 | Axonal Guidance Signaling | 4.87E+00 | 1.02E-01 |
| 10 | Superpathway of Cholesterol Biosynthesis | 4.79E+00 | 3.10E-01 |

B.

| IPA pathway changes in astrocytes after GCSi treatment | | | |
|---|---|---|---|
| | Ingenuity Canonical Pathways | -log(p-value) | Ratio |
| 1 | LXR/RXR Activation | 8.88E+00 | 1.14E-01 |
| 2 | Acute Phase Response Signaling | 5.06E+00 | 6.49E-02 |
| 3 | HMGB1 Signaling | 4.74E+00 | 6.59E-02 |
| 4 | TNFR2 Signaling | 4.18E+00 | 1.56E-01 |
| 5 | Atherosclerosis Signaling | 4.07E+00 | 6.77E-02 |
| 6 | Agranulocyte Adhesion and Diapedesis | 3.84E+00 | 5.24E-02 |
| 7 | Airway Pathology in Chronic Obstructive Pulmonary Disease | 3.69E+00 | 6.78E-02 |
| 8 | ABRA Signaling Pathway | 3.60E+00 | 7.61E-02 |
| 9 | NOD1/2 Signaling Pathway | 3.58E+00 | 5.29E-02 |
| 10 | Hepatic Cholestasis | 3.54E+00 | 5.24E-02 |

C.

| IPA pathway changes in microglia after FB1 treatment | | | |
|---|---|---|---|
| | Ingenuity Canonical Pathways | -log(p-value) | Ratio |
| 1 | CREB Signaling in Neurons | 8.29E+00 | 7.74E-02 |
| 2 | Granulocyte Adhesion and Diapedesis | 7.11E+00 | 1.16E-01 |
| 3 | Agranulocyte Adhesion and Diapedesis | 6.31E+00 | 1.05E-01 |
| 4 | Breast Cancer Regulation by Stathmin1 | 5.98E+00 | 6.90E-02 |
| 5 | S100 Family Signaling Pathway | 5.97E+00 | 6.35E-02 |
| 6 | Hepatic Fibrosis / Hepatic Stellate Cell Activation | 5.11E+00 | 9.79E-02 |
| 7 | Role of IL-17A in Psoriasis | 4.39E+00 | 3.57E-01 |
| 8 | G-Protein Coupled Receptor Signaling | 4.24E+00 | 5.83E-02 |
| 9 | Airway Pathology in Chronic Obstructive Pulmonary Disease | 4.20E+00 | 1.10E-01 |
| 10 | Phagosome Formation | 3.42E+00 | 5.47E-02 |

D.

| IPA pathway changes in microglia after GCSi treatment | | | |
|---|---|---|---|
| | Ingenuity Canonical Pathways | -log(p-value) | Ratio |
| 1 | S100 Family Signaling Pathway | 1.11E+01 | 6.48E-02 |
| 2 | Hepatic Fibrosis / Hepatic Stellate Cell Activation | 9.52E+00 | 1.13E-01 |
| 3 | CREB Signaling in Neurons | 8.69E+00 | 6.43E-02 |
| 4 | Phagosome Formation | 8.52E+00 | 6.04E-02 |
| 5 | Granulocyte Adhesion and Diapedesis | 6.73E+00 | 9.52E-02 |
| 6 | Agranulocyte Adhesion and Diapedesis | 6.72E+00 | 9.05E-02 |
| 7 | Pulmonary Fibrosis Idiopathic Signaling Pathway | 6.62E+00 | 7.36E-02 |
| 8 | Breast Cancer Regulation by Stathmin1 | 5.99E+00 | 5.56E-02 |
| 9 | G-Protein Coupled Receptor Signaling | 5.66E+00 | 5.12E-02 |
| 10 | Gustation Pathway | 4.98E+00 | 7.88E-02 |

E.

| IPA pathway changes in both astrocytes and microglia after FB1 treatment | | | |
|---|---|---|---|
| | Ingenuity Canonical Pathways | -log(p-value) | Ratio |
| 1 | Hepatic Fibrosis / Hepatic Stellate Cell Activation | 2.74E+00 | 2.58E-02 |
| 2 | Superpathway of Cholesterol Biosynthesis | 2.13E+00 | 6.90E-02 |
| 3 | ABRA Signaling Pathway | 2.09E+00 | 3.26E-02 |
| 4 | Granulocyte Adhesion and Diapedesis | 1.99E+00 | 2.12E-02 |
| 5 | Melatonin Degradation II | 1.75E+00 | 2.50E-01 |
| 6 | Zymosterol Biosynthesis | 1.58E+00 | 1.67E-01 |
| 7 | NAD Biosynthesis III | 1.58E+00 | 1.67E-01 |
| 8 | LPS/IL-1 Mediated Inhibition of RXR Function | 1.56E+00 | 1.57E-02 |
| 9 | NAD Biosynthesis from 2-amino-3-carboxymuconate Semialdehyde | 1.51E+00 | 1.43E-01 |
| 10 | NAD Salvage Pathway III | 1.51E+00 | 1.43E-01 |

F.

| IPA pathway changes in motor neurons after GCSi treatment | | | |
|---|---|---|---|
| | Ingenuity Canonical Pathways | -log(p-value) | Ratio |
| 1 | Glutamate Degradation II | 1.84E+00 | 3.33E-01 |
| 2 | Aspartate Biosynthesis | 1.84E+00 | 3.33E-01 |
| 3 | L-cysteine Degradation I | 1.72E+00 | 2.50E-01 |
| 4 | Neuroprotective Role of THOP1 in Alzheimer's Disease | 1.68E+00 | 2.48E-02 |
| 5 | LXR/RXR Activation | 1.66E+00 | 2.44E-02 |
| 6 | FXR/RXR Activation | 1.63E+00 | 2.38E-02 |
| 7 | Osteoarthritis Pathway | 1.55E+00 | 1.69E-02 |
| 8 | Role of Cytokines in Mediating Communication between Immune Cells | 1.55E+00 | 3.70E-02 |
| 9 | Serotonin and Melatonin Biosynthesis | 1.54E+00 | 1.67E-01 |
| 10 | Aspartate Degradation II | 1.48E+00 | 1.43E-01 |

G.

| IPA pathway changes in astrocytes after both FB1 and GCSi treatment | | | |
|---|---|---|---|
| | Ingenuity Canonical Pathways | -log(p-value) | Ratio |
| 1 | Agranulocyte Adhesion and Diapedesis | 3.01E+00 | 2.86E-02 |
| 2 | Inhibition of Matrix Metalloproteases | 2.92E+00 | 7.69E-02 |
| 3 | ABRA Signaling Pathway | 2.81E+00 | 4.35E-02 |
| 4 | Pulmonary Fibrosis Idiopathic Signaling Pathway | 2.70E+00 | 2.15E-02 |
| 5 | Apelin Cardiac Fibroblast Signaling Pathway | 2.17E+00 | 8.70E-02 |
| 6 | Role Of Chondrocytes In Rheumatoid Arthritis Signaling Pathway | 2.14E+00 | 2.84E-02 |
| 7 | Granulocyte Adhesion and Diapedesis | 1.71E+00 | 2.12E-02 |
| 8 | WNK Renal Signaling Pathway | 1.70E+00 | 2.83E-02 |
| 9 | Leukocyte Extravasation Signaling | 1.68E+00 | 2.07E-02 |
| 10 | ID1 Signaling Pathway | 1.63E+00 | 1.99E-02 |

H.

| IPA pathway changes in microglia after both FB1 and GCSi treatment | | | |
|---|---|---|---|
| | Ingenuity Canonical Pathways | -log(p-value) | Ratio |
| 1 | Granulocyte Adhesion and Diapedesis | 6.78E+00 | 5.29E-02 |
| 2 | Agranulocyte Adhesion and Diapedesis | 6.36E+00 | 4.76E-02 |
| 3 | Tumor Microenvironment Pathway | 4.99E+00 | 4.47E-02 |
| 4 | Hepatic Fibrosis / Hepatic Stellate Cell Activation | 4.73E+00 | 4.12E-02 |
| 5 | S100 Family Signaling Pathway | 4.35E+00 | 1.94E-02 |
| 6 | CREB Signaling in Neurons | 4.25E+00 | 2.14E-02 |
| 7 | Axonal Guidance Signaling | 3.09E+00 | 1.96E-02 |
| 8 | Role Of Chondrocytes In Rheumatoid Arthritis Signaling Pathway | 2.85E+00 | 3.55E-02 |
| 9 | Inhibition of Matrix Metalloproteases | 2.82E+00 | 7.69E-02 |
| 10 | Heparan Sulfate Biosynthesis | 2.78E+00 | 4.60E-02 |

**Figure 6. IPA pathway analysis of bulk RNAseq data from astrocytes, microglia, and motor neurons treated with FB1 and GCSi.** List of top 10 pathway changes as sorted by the -log(p-value).

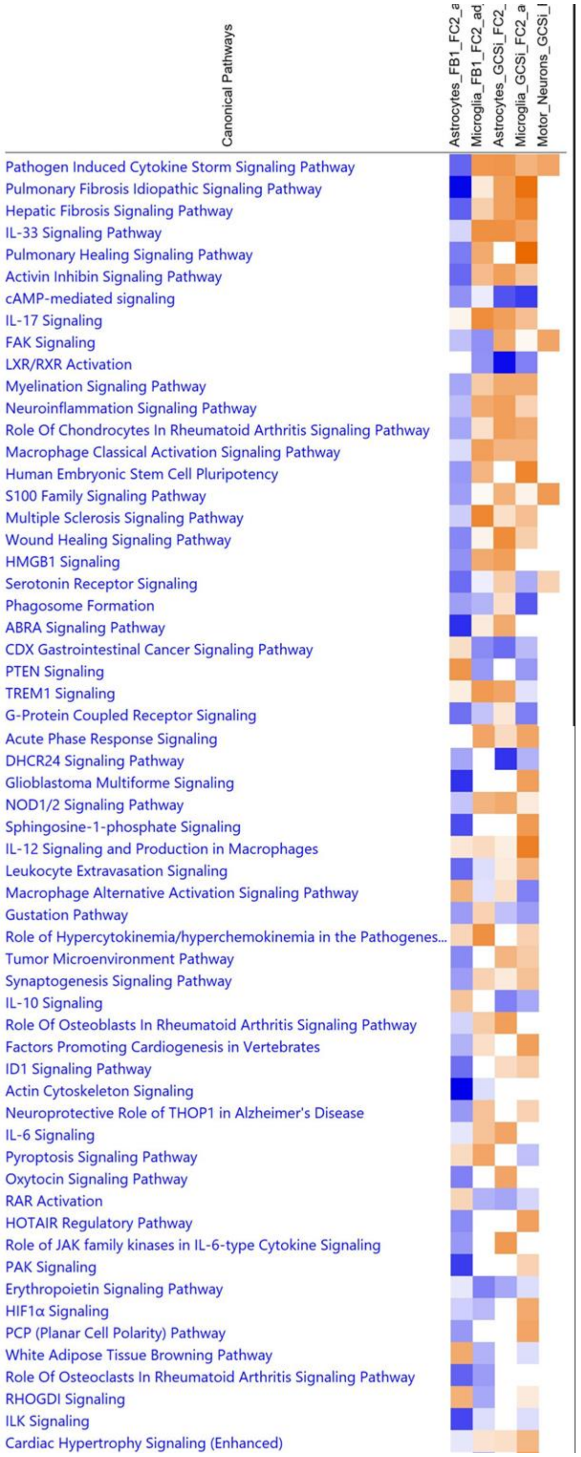


**Figure 7. Common pathway changes across treatments and cell-types.** List of top pathway changes as sorted by absolute value of effect sizes across conditions.

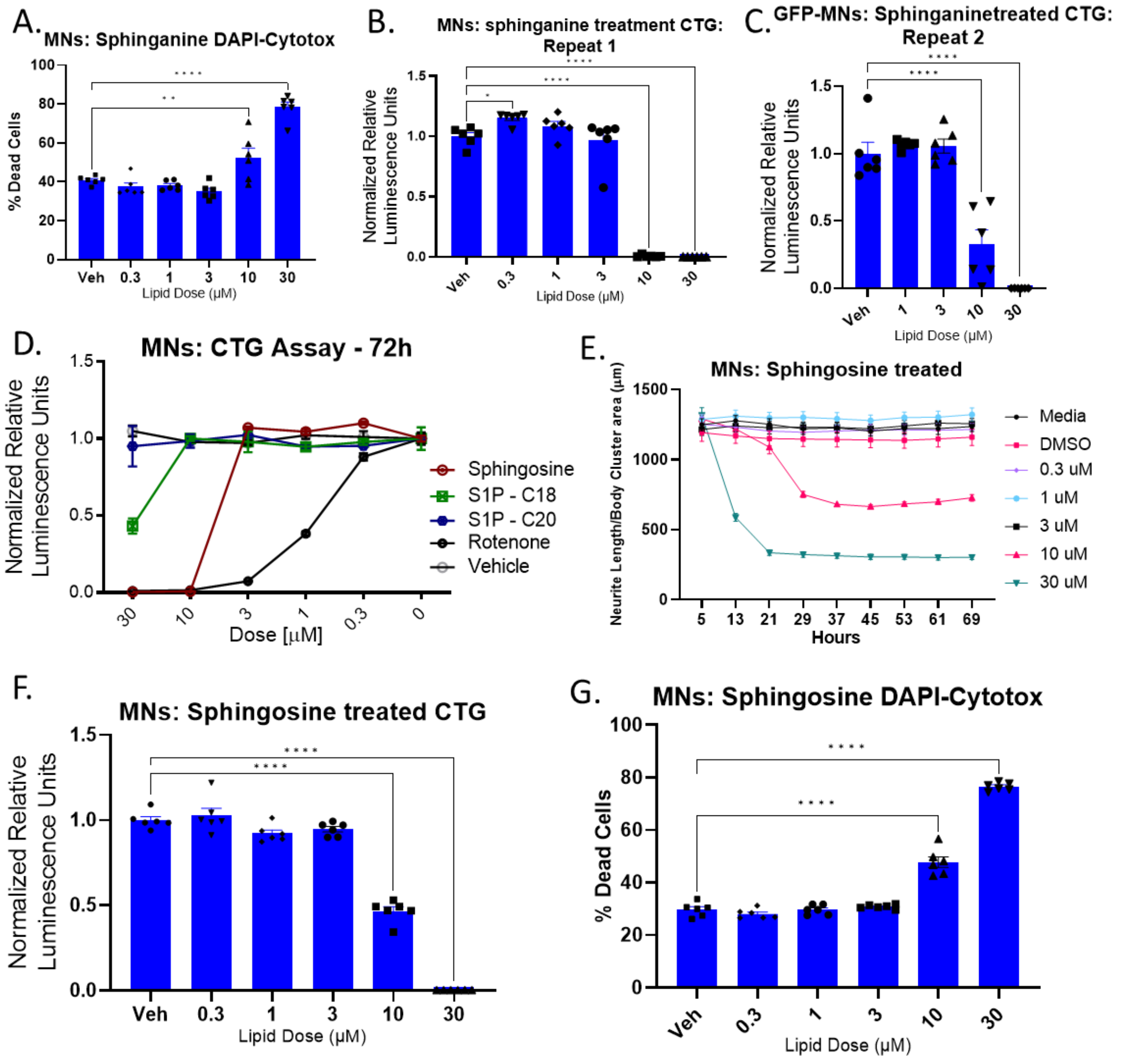


**Supplemental Figure 8. Toxicity of multiple sphingolipids to motor neurons (MNs). A)** Confirmation of CTG and neurite analysis (Figures 7G and J) toxicity measures after sphinganine treatment by live nuclear counts and cytotox green dead cell counts. Graphed as percentage of total nuclei that were cytotox$^{+}$. **B and C)** Repeats of MN sphinganine toxicity experiment CTG readouts of cell viability. One repeat was performed on GFP$^{+}$ MNs and was consistent with 10 µM being the first toxic dose. **D)** Increasing doses of Rotenone, C20 sphingosine-1-phosphate (S1P – C20), C18 sphingosine-1-phosphate (S1P – C18), and sphingosine are toxic to motor neurons as determined by cell titer glo (CTG) measures (performed at BrainXell). **E and F)** Sphingosine was toxic at 10 µM doses as shown by neurite analysis (**E**) and CTG (**F**). **G)** Cytotox and total nuclear staining confirms findings of sphingosine toxicity. Data analyzed by one-way ANOVA with Bonferroni's multiple comparisons test, ****$p<0.0001$, ***$p<0.001$, **$p<0.01$, *$P<0.05$.

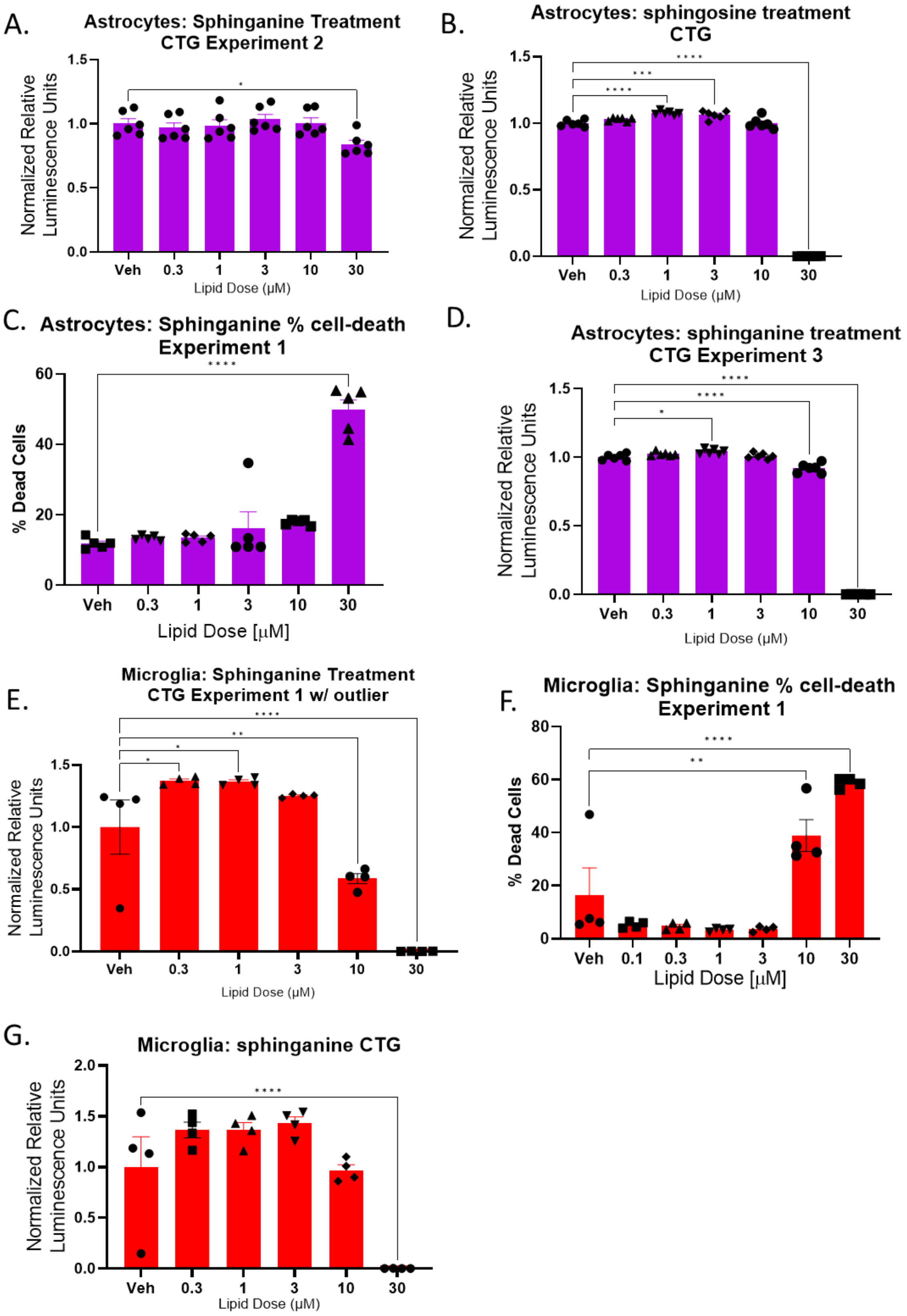
A.
Astrocytes: Sphinganine Treatment
CTG Experiment 2
Normalized Relative Luminescence Units
Lipid Dose (μM)
B.
Astrocytes: sphingosine treatment
CTG
Normalized Relative Luminescence Units
Lipid Dose (μM)
C.
Astrocytes: Sphinganine % cell-death
Experiment 1
% Dead Cells
Lipid Dose [μM]
D.
Astrocytes: sphinganine treatment
CTG Experiment 3
Normalized Relative Luminescence Units
Lipid Dose (μM)
E.
Microglia: Sphinganine Treatment
CTG Experiment 1 w/ outlier
Normalized Relative Luminescence Units
Lipid Dose (μM)
F.
Microglia: Sphinganine % cell-death
Experiment 1
% Dead Cells
Lipid Dose [μM]
G.
Microglia: sphinganine CTG
Normalized Relative Luminescence Units
Lipid Dose (μM)
Veh
0.1
0.3
1
3
10
30

**Supplemental Figure 9. Multiple sphingolipids are toxic to glia. A)** Second repeat of sphinganine toxicity to astrocytes (first repeat Figure 7E). **B)** Astrocytes treated with sphingosine were similarly resistant to 10 μM dose as measured by CTG. **C)** Confirmation of sphinganine toxicity in astrocytes by cytotox$^{+}$ counts as percentage of total nuclear counts. **D)** Third repeat of sphinganine toxicity to astrocytes measured by CTG. **E)** Repeat of sphinganine toxicity to microglia measured by CTG. **F)** Confirmation of sphinganine toxicity (Figure 7I) in microglia by cytotox$^{+}$ counts as percentage of total nuclear counts. Data analyze by one-way ANOVA with Bonferroni's multiple comparisons test, ****$p$<0.0001, ***$p$<0.001, **$p$<0.01, *$P$<0.05.

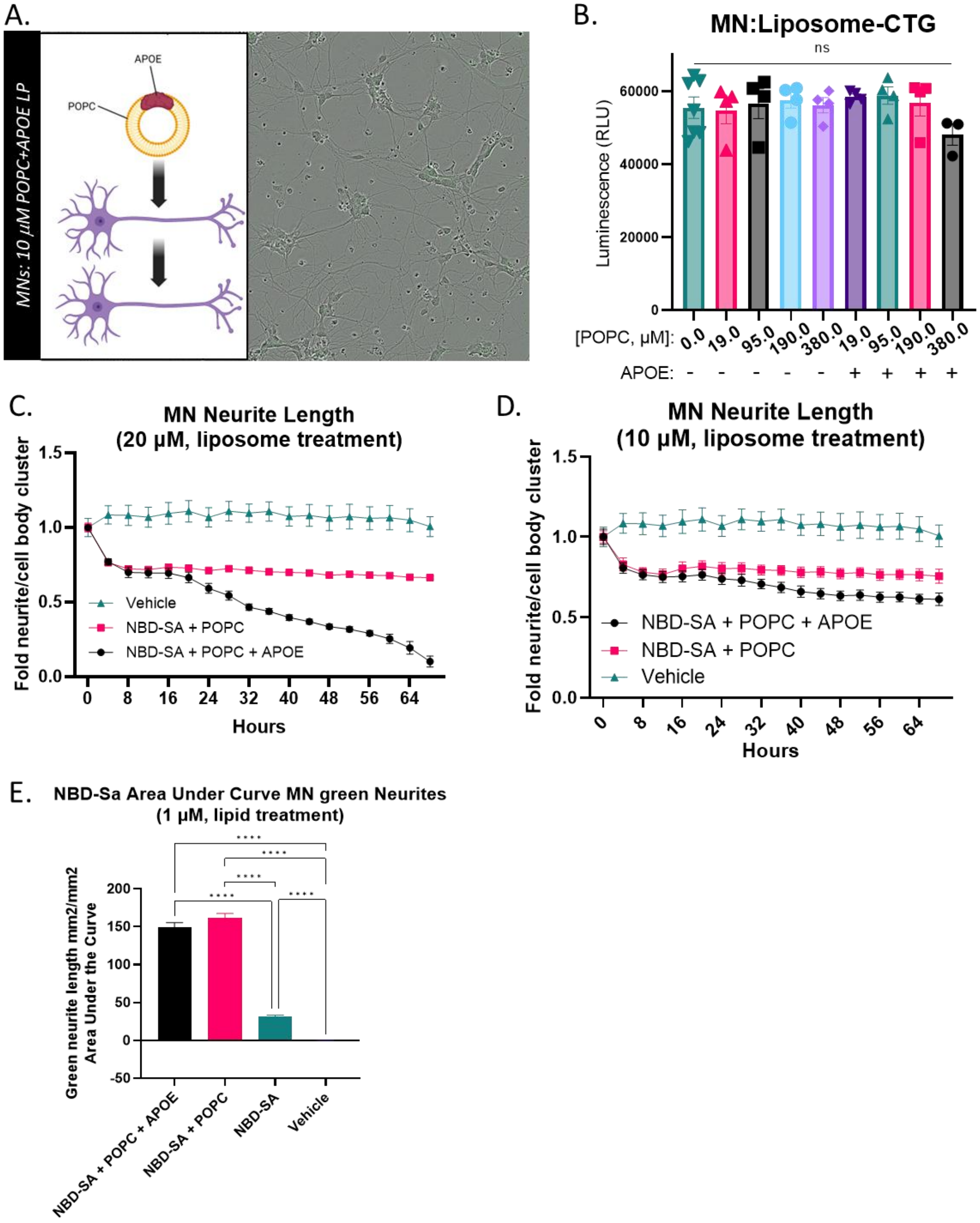
A.
MNs: 10 μM POPC+APOE LP
APOE
POPC
B.
MN:Liposome-CTG
ns
Luminescence (RLU)
60000
40000
20000
0
[POPC, μM]: 0.0 19.0 95.0 190.0 380.0 19.0 95.0 190.0 380.0
APOE: - - - - - + + + +
C.
MN Neurite Length
(20 μM, liposome treatment)
Fold neurite/cell body cluster
1.5
1.0
0.5
0.0
Vehicle
NBD-SA + POPC
NBD-SA + POPC + APOE
0 8 16 24 32 40 48 56 64
Hours
D.
MN Neurite Length
(10 μM, liposome treatment)
Fold neurite/cell body cluster
1.5
1.0
0.5
0.0
NBD-SA + POPC + APOE
NBD-SA + POPC
Vehicle
0 8 16 24 32 40 48 56 64
Hours
E.
NBD-Sa Area Under Curve MN green Neurites
(1 μM, lipid treatment)
Green neurite length mm2/mm2
Area Under the Curve
200
150
100
50
0
-50
****
NBD-SA + POPC + APOE
NBD-SA + POPC
NBD-SA
Vehicle

**Supplemental Figure 10. Liposomal delivery of sphinganine to motor neurons (Figure 8 continued). A-E)** To determine if a more physiological preparation of sphinganine would alter delivery, we made liposomes with NBD-labelled (green) sphinganine and liposomes lacking sphinganine. **A)** Images of motor neuron (MN) cultures treated with liposomes made up of POPC and APOE (no NBD-sphinganine), showing no visible green or toxicity. **B)** CTG analysis of MN viability shows no significant impact of liposomes without NBD-sphinganine. **C and D)** Phase neurite analysis of MNs treated with 20 μM and 10 μM NBD-sphinganine in different liposomal formats. **E)** Area under the curve analysis of Figure 8B, showing significant increases in NBD signal when NBD-sphinganine is delivered in a liposomal versus free format but no significant change resulting from $APOE^{+}$ liposomes (****$P$<0.0001, One-way ANOVA, Tukey's multiple comparisons test).
