## Supplementary material for "Unravelling neuronal and glial differences in ceramide composition, synthesis, and sensitivity to toxicity": Venn_List_Astrocyte FB1 GCSi changes

| GeneID | name |
|---|---|
| ENSG0000 | PDK4 |
| ENSG0000 | DNAH9 |
| ENSG0000 | DLEC1 |
| ENSG0000 | BIRC3 |
| ENSG0000 | EFCAB1 |
| ENSG0000 | LMO3 |
| ENSG0000 | PTPRN |
| ENSG0000 | SREBF1 |
| ENSG0000 | MPP4 |
| ENSG0000 | SLCO1A2 |
| ENSG0000 | NKAIN4 |
| ENSG0000 | ODAD1 |
| ENSG0000 | TFPI2 |
| ENSG0000 | ACTA2 |
| ENSG0000 | DKK1 |
| ENSG0000 | CCL2 |
| ENSG0000 | RSPH4A |
| ENSG0000 | SERPINI2 |
| ENSG0000 | RGS4 |
| ENSG0000 | IRAG2 |
| ENSG0000 | ADGB |
| ENSG0000 | SGK1 |
| ENSG0000 | CCN2 |
| ENSG0000 | CCDC170 |
| ENSG0000 | EGR1 |
| ENSG0000 | DUSP4 |
| ENSG0000 | DNAI1 |
| ENSG0000 | MATN4 |
| ENSG0000 | WNK4 |
| ENSG0000 | MASP1 |
| ENSG0000 | CPA4 |
| ENSG0000 | TNNI3 |
| ENSG0000 | NALF2 |
| ENSG0000 | KLHDC7B |
| ENSG0000 | STOML3 |
| ENSG0000 | DYDC2 |
| ENSG0000 | SYT6 |
| ENSG0000 | EGR4 |
| ENSG0000 | GPNMB |
| ENSG0000 | WDR38 |
| ENSG0000 | TTC29 |
| ENSG0000 | THBS1 |
| ENSG0000 | ITGB7 |
| ENSG0000 | AK7 |
| ENSG0000 | MYLK3 |
| ENSG0000 | LRRC46 |

ENSG0000 SLC14A1
ENSG0000 PMAIP1
ENSG0000 CFAP74
ENSG0000 CCN1
ENSG0000 TAGLN
ENSG0000 KIAA1755
ENSG0000 CAPSL
ENSG0000 DYNLT5
ENSG0000 NRSN1
ENSG0000 CHST9
ENSG0000 DNAAF1
ENSG0000 USP43
ENSG0000 PTPRN2
ENSG0000 CFAP161
ENSG0000 FBXO32
ENSG0000 VWA5B1
ENSG0000 DRC7
ENSG0000 TPPP3
ENSG0000 GAB3
ENSG0000 ICOSLG
ENSG0000 CFAP157
ENSG0000 LRRC71
ENSG0000 AQP5
ENSG0000 UBXN10
ENSG0000 ATF3
ENSG0000 KCNF1
ENSG0000 GABRG1
ENSG0000 EFHB
ENSG0000 CFAP100
ENSG0000 ZNF474
ENSG0000 PI16
ENSG0000 SYTL3
ENSG0000 PDZRN4
ENSG0000 CFAP52
ENSG0000 MMP10
ENSG0000 DNAAF3
ENSG0000 ANGPTL4
ENSG0000 TEKT1
ENSG0000 STXBP6
ENSG0000 RSPH10B2
ENSG0000 FRMPD2
ENSG0000 IL16
ENSG0000 EFCAB12
ENSG0000 HPSE2
ENSG0000 CCDC13-AS1
ENSG0000 CHRNA9
ENSG0000 VWA3A

| | |
|---|---|
| ENSG0000 | PLEKHD1 |
| ENSG0000 | SPHK1 |
| ENSG0000 | MAP3K19 |
| ENSG0000 | GCNT4 |
| ENSG0000 | FIBIN |
| ENSG0000 | ODF3B |
| ENSG0000 | ERICH3 |
| ENSG0000 | EGR3 |
| ENSG0000 | C3orf80 |
| ENSG0000 | NME9 |
| ENSG0000 | CFAP65 |
| ENSG0000 | KIAA2012 |
| ENSG0000 | HOATZ |
| ENSG0000 | DNAH2 |
| ENSG0000 | SDR42E2 |
| ENSG0000 | CLDN5 |
| ENSG0000 | MORN5 |
| ENSG0000 | MYBL1 |
| ENSG0000 | KLHL32 |
| ENSG0000 | CFAP73 |
| ENSG0000 | DNAJB13 |
| ENSG0000 | LRRC74B |
| ENSG0000 | PRELP |
| ENSG0000 | KIF19 |
| ENSG0000 | CFAP299 |
| ENSG0000 | PLN |
| ENSG0000 | MMP17 |
| ENSG0000 | Y_RNA |
| ENSG0000 | IQANK1 |
| ENSG0000 | ERICH2 |
| ENSG0000 | BTBD17 |
| ENSG0000 | GMNC |
| ENSG0000 | CFAP99 |
| ENSG0000 | CFAP45 |
| ENSG0000 | C10orf105 |
| ENSG0000 | SIAH3 |
| ENSG0000 | WARS2-IT1 |
| ENSG0000 | ZNF812P |
| ENSG0000 | None |
| ENSG0000 | None |
| ENSG0000 | MYOSLID |
| ENSG0000 | TGFB2-AS1 |
| ENSG0000 | None |
| ENSG0000 | LINC00707 |
| ENSG0000 | VSIG8 |
| ENSG0000 | GSTA1 |
| ENSG0000 | None |

| | |
|---|---|
| ENSG0000 | None |
| ENSG0000 | C8orf34-AS1 |
| ENSG0000 | ZNF474-AS1 |
| ENSG0000 | STK19B |
| ENSG0000 | None |
| ENSG0000 | None |
| ENSG0000 | None |
| ENSG0000 | AGAP12P |
| ENSG0000 | CCDC177 |
| ENSG0000 | GAS2L2 |
| ENSG0000 | FLJ16779 |
| ENSG0000 | None |
| ENSG0000 | None |
