## Supplementary material for "Unravelling neuronal and glial differences in ceramide composition, synthesis, and sensitivity to toxicity": Venn_List Microglia genes FB1-down GCSi-up

| geneid | name |
|---|---|
| ENSG0000 | MYH16 |
| ENSG0000 | NTSR1 |
| ENSG0000 | F13A1 |
| ENSG0000 | SCN7A |
| ENSG0000 | ABCC12 |
| ENSG0000 | HMGA2 |
| ENSG0000 | MMP21 |
| ENSG0000 | STEAP1 |
| ENSG0000 | HTRA3 |
| ENSG0000 | SULT1B1 |
| ENSG0000 | RFLNB |
| ENSG0000 | ARSI |
| ENSG0000 | CLDN5 |
| ENSG0000 | NOTUM |
| ENSG0000 | C5orf52 |
| ENSG0000 | SP6 |
| ENSG0000 | None |
| ENSG0000 | RPSAP52 |
| ENSG0000 | LINC02454 |
| ENSG0000 | None |
| ENSG0000 | H4C1 |

| GeneID | Treatment | Treatment | Treatment | Treatment | name |
|---|---|---|---|---|---|
| ENSG0000 | -1.94531 | -3.8512 | 0.001088 | 0.026086 | MYH16 |
| ENSG0000 | -1.59435 | -3.01958 | 0.093712 | 0.473866 | HSPB6 |
| ENSG0000 | -1.34423 | -2.53896 | 0.02106 | 0.202859 | ITGA2B |
| ENSG0000 | -1.06376 | -2.09037 | 1.01E-11 | 2.45E-09 | ETV1 |
| ENSG0000 | -1.13021 | -2.18891 | 0.005242 | 0.080785 | SEMA3G |
| ENSG0000 | -2.40964 | -5.31344 | 0.062692 | 0.383859 | ISL1 |
| ENSG0000 | -1.32254 | -2.50106 | 0.040539 | 0.303305 | HGF |
| ENSG0000 | -2.92359 | -7.58732 | 0.027885 | 0.241401 | SLC7A9 |
| ENSG0000 | -1.634 | -3.10372 | 0.007428 | 0.10248 | SOX30 |
| ENSG0000 | -1.06906 | -2.09807 | 0.003985 | 0.066367 | ADAMTS6 |
| ENSG0000 | -1.27645 | -2.42242 | 0.00637 | 0.091955 | NEXMIF |
| ENSG0000 | -1.01434 | -2.01998 | 6.55E-37 | 2.30E-33 | MSMO1 |
| ENSG0000 | -5.71474 | -52.518 | 0.001649 | 0.035714 | POU1F1 |
| ENSG0000 | -2.95086 | -7.73209 | 0.024875 | 0.225802 | SNAP91 |
| ENSG0000 | -1.50458 | -2.83742 | 0.00332 | 0.057869 | SYT1 |
| ENSG0000 | -2.85121 | -7.21608 | 0.076689 | 0.425382 | STON1-GTF2A1L |
| ENSG0000 | -4.67841 | -25.606 | 0.035015 | 0.278621 | MAOB |
| ENSG0000 | -3.59777 | -12.107 | 0.014935 | 0.160991 | MYO3B |
| ENSG0000 | -5.04665 | -33.0515 | 0.010506 | 0.127955 | WSCD2 |
| ENSG0000 | -1.68876 | -3.22379 | 4.20E-09 | 6.21E-07 | NMRK2 |
| ENSG0000 | -1.82026 | -3.53144 | 0.053936 | 0.355307 | ADCY2 |
| ENSG0000 | -3.36217 | -10.2829 | 0.000289 | 0.009292 | CDH17 |
| ENSG0000 | -1.76022 | -3.38749 | 0.016615 | 0.172604 | COBLL1 |
| ENSG0000 | -2.387 | -5.23069 | 0.069482 | 0.405762 | APOB |
| ENSG0000 | -5.69643 | -51.8556 | 7.20E-05 | 0.003145 | HSD17B2 |
| ENSG0000 | -3.61277 | -12.2336 | 0.0002 | 0.006945 | REM1 |
| ENSG0000 | -1.33385 | -2.52075 | 0.071022 | 0.409896 | FER1L4 |
| ENSG0000 | -1.52548 | -2.87883 | 0.076292 | 0.424503 | TMEM40 |
| ENSG0000 | -4.70732 | -26.1243 | 0.005161 | 0.079808 | SIRPG |
| ENSG0000 | -4.09202 | -17.0538 | 0.002569 | 0.048526 | CHGB |
| ENSG0000 | -1.40786 | -2.65343 | 0.041001 | 0.305617 | HEPH |
| ENSG0000 | -3.31931 | -9.98188 | 0.014184 | 0.155285 | THPO |
| ENSG0000 | -1.04927 | -2.06948 | 5.64E-10 | 9.89E-08 | TGFB2 |
| ENSG0000 | -1.08732 | -2.12479 | 8.69E-22 | 7.17E-19 | SORBS1 |
| ENSG0000 | -1.25125 | -2.38047 | 4.75E-13 | 1.46E-10 | DSP |
| ENSG0000 | -2.2638 | -4.80255 | 0.012402 | 0.141875 | SLC7A4 |
| ENSG0000 | -5.02392 | -32.535 | 0.011163 | 0.132786 | PNPLA5 |
| ENSG0000 | -3.42518 | -10.7419 | 0.009188 | 0.118295 | GRAP2 |
| ENSG0000 | -1.05834 | -2.08253 | 8.48E-07 | 7.36E-05 | TRIM9 |
| ENSG0000 | -1.92774 | -3.80457 | 1.35E-27 | 2.60E-24 | NTSR1 |
| ENSG0000 | -2.76569 | -6.80075 | 0.046091 | 0.326547 | BMX |
| ENSG0000 | -1.70447 | -3.25909 | 0.039085 | 0.29686 | ASB9 |
| ENSG0000 | -3.61942 | -12.2901 | 0.064352 | 0.390104 | RAB40AL |
| ENSG0000 | -1.50325 | -2.8348 | 0.054734 | 0.357945 | CRYM |
| ENSG0000 | -1.41293 | -2.66277 | 0.075336 | 0.421944 | KLC3 |
| ENSG0000 | -1.00816 | -2.01135 | 0.009695 | 0.122115 | EPHX3 |

| | | | | | |
|---|---|---|---|---|---|
| ENSG0000 | -1.10719 | -2.15426 | 4.50E-08 | 5.20E-06 | CD79A |
| ENSG0000 | -1.59932 | -3.03 | 0.003598 | 0.061374 | ATP1A3 |
| ENSG0000 | -1.7463 | -3.35496 | 0.000509 | 0.014601 | SIGLEC6 |
| ENSG0000 | -5.46994 | -44.3216 | 0.000148 | 0.005586 | MAG |
| ENSG0000 | -1.48143 | -2.79225 | 2.62E-12 | 7.06E-10 | CRHR2 |
| ENSG0000 | -2.40338 | -5.2904 | 4.14E-16 | 1.94E-13 | KCNA1 |
| ENSG0000 | -1.16779 | -2.24667 | 4.31E-09 | 6.34E-07 | ST8SIA1 |
| ENSG0000 | -1.06881 | -2.09771 | 0.010397 | 0.127127 | MAK |
| ENSG0000 | -1.54739 | -2.92288 | 0.095958 | 0.479543 | IMPG1 |
| ENSG0000 | -1.10693 | -2.15386 | 5.41E-64 | 5.06E-60 | HMGCS1 |
| ENSG0000 | -1.91739 | -3.77738 | 0.087454 | 0.456032 | FGF1 |
| ENSG0000 | -3.88841 | -14.8091 | 0.002615 | 0.048905 | CDX1 |
| ENSG0000 | -1.14715 | -2.21475 | 2.67E-11 | 5.90E-09 | HYAL1 |
| ENSG0000 | -4.29949 | -19.6914 | 0.013339 | 0.148909 | PLSCR4 |
| ENSG0000 | -1.59931 | -3.02998 | 7.95E-07 | 6.94E-05 | VIPR1 |
| ENSG0000 | -1.0209 | -2.02918 | 0.013315 | 0.148849 | SLC4A3 |
| ENSG0000 | -1.2932 | -2.45071 | 0.044264 | 0.319658 | DNAH6 |
| ENSG0000 | -1.98412 | -3.95622 | 5.70E-15 | 2.35E-12 | IL1R2 |
| ENSG0000 | -5.04552 | -33.0257 | 0.001993 | 0.040982 | RGS4 |
| ENSG0000 | -1.82844 | -3.55153 | 7.45E-14 | 2.61E-11 | SLC16A7 |
| ENSG0000 | -2.43064 | -5.39131 | 0.018041 | 0.182817 | TRIM67 |
| ENSG0000 | -1.20719 | -2.30887 | 0.027139 | 0.237737 | SLC46A2 |
| ENSG0000 | -1.44828 | -2.72883 | 4.93E-08 | 5.66E-06 | EPCAM |
| ENSG0000 | -2.76762 | -6.80982 | 0.096513 | 0.480642 | HOXB5 |
| ENSG0000 | -2.13236 | -4.38434 | 0.059554 | 0.37387 | MYOT |
| ENSG0000 | -1.38483 | -2.61141 | 0.082747 | 0.443526 | ZSCAN18 |
| ENSG0000 | -2.97558 | -7.86571 | 0.005118 | 0.079358 | PDZRN3 |
| ENSG0000 | -1.94929 | -3.86183 | 0.007939 | 0.107063 | CSMD2 |
| ENSG0000 | -3.45421 | -10.9602 | 0.012885 | 0.145491 | AKR1D1 |
| ENSG0000 | -1.89226 | -3.71216 | 0.079519 | 0.433177 | RAB9B |
| ENSG0000 | -1.27305 | -2.41672 | 0.067346 | 0.39937 | HIF3A |
| ENSG0000 | -1.39058 | -2.62183 | 0.00599 | 0.088193 | F13A1 |
| ENSG0000 | -2.55841 | -5.89059 | 0.089178 | 0.460401 | GRM4 |
| ENSG0000 | -1.25892 | -2.39316 | 0.002787 | 0.051102 | KIF25 |
| ENSG0000 | -3.68737 | -12.8828 | 0.076788 | 0.425589 | MYH2 |
| ENSG0000 | -1.88799 | -3.7012 | 1.25E-05 | 0.000729 | KIRREL2 |
| ENSG0000 | -2.17557 | -4.51766 | 3.67E-06 | 0.000253 | ACKR4 |
| ENSG0000 | -1.32928 | -2.51277 | 2.60E-08 | 3.21E-06 | RNASE1 |
| ENSG0000 | -5.50381 | -45.3747 | 0.002319 | 0.045333 | PRRG3 |
| ENSG0000 | -3.14355 | -8.83697 | 0.028383 | 0.244079 | SULT4A1 |
| ENSG0000 | -2.59846 | -6.0564 | 0.016305 | 0.171098 | CRB3 |
| ENSG0000 | -5.47003 | -44.3244 | 0.004181 | 0.068618 | TEX101 |
| ENSG0000 | -5.02567 | -32.5746 | 0.006851 | 0.0969 | None |
| ENSG0000 | -2.62278 | -6.15935 | 0.039498 | 0.298618 | RAB25 |
| ENSG0000 | -4.96729 | -31.2827 | 0.000942 | 0.023394 | SEC14L4 |
| ENSG0000 | -2.86739 | -7.29745 | 1.91E-19 | 1.37E-16 | LYVE1 |
| ENSG0000 | -3.75016 | -13.4558 | 0.035165 | 0.279177 | TEX15 |

| | | | | | |
|---|---|---|---|---|---|
| ENSG0000 | -1.60259 | -3.03688 | 0.004364 | 0.070841 | HRH4 |
| ENSG0000 | -2.54515 | -5.83667 | 0.101519 | 0.493758 | RERG |
| ENSG0000 | -2.73268 | -6.64691 | 0.006214 | 0.090545 | TCN1 |
| ENSG0000 | -1.60062 | -3.03273 | 3.08E-29 | 8.63E-26 | ADAM19 |
| ENSG0000 | -1.23186 | -2.3487 | 0.055573 | 0.3612 | GALNT5 |
| ENSG0000 | -1.00158 | -2.00219 | 0.001132 | 0.026978 | SCN7A |
| ENSG0000 | -1.67442 | -3.19191 | 3.26E-14 | 1.19E-11 | HCRTR2 |
| ENSG0000 | -1.47555 | -2.78089 | 0.076232 | 0.424334 | TUBB2B |
| ENSG0000 | -2.04902 | -4.13826 | 0.070842 | 0.40959 | LHCGR |
| ENSG0000 | -5.05414 | -33.2237 | 0.000513 | 0.014681 | SLC3A1 |
| ENSG0000 | -1.28682 | -2.4399 | 0.017264 | 0.177553 | ZNF365 |
| ENSG0000 | -1.02806 | -2.03927 | 1.91E-06 | 0.000149 | AK7 |
| ENSG0000 | -4.76645 | -27.2173 | 0.036047 | 0.282659 | FAM181A |
| ENSG0000 | -4.33217 | -20.1426 | 0.020967 | 0.202432 | RLBP1 |
| ENSG0000 | -2.84488 | -7.18448 | 3.54E-09 | 5.28E-07 | ABCC12 |
| ENSG0000 | -4.27509 | -19.3612 | 0.01477 | 0.159644 | SLC25A52 |
| ENSG0000 | -1.26755 | -2.40753 | 4.50E-66 | 1.26E-61 | TNFRSF11A |
| ENSG0000 | -2.75412 | -6.74641 | 0.102361 | 0.49639 | CBLN2 |
| ENSG0000 | -5.56807 | -47.4412 | 0.000637 | 0.01724 | CIB3 |
| ENSG0000 | -1.02967 | -2.04155 | 0.076569 | 0.424948 | HUNK |
| ENSG0000 | -1.32429 | -2.50409 | 6.78E-09 | 9.42E-07 | KCNA10 |
| ENSG0000 | -2.82599 | -7.09099 | 0.09788 | 0.484462 | S100A7 |
| ENSG0000 | -3.15393 | -8.90077 | 0.027548 | 0.24012 | ACTA1 |
| ENSG0000 | -2.43732 | -5.41636 | 0.001622 | 0.03536 | NYAP2 |
| ENSG0000 | -4.73408 | -26.6133 | 0.007974 | 0.10738 | GADL1 |
| ENSG0000 | -1.48685 | -2.80275 | 0.008603 | 0.112707 | CAND2 |
| ENSG0000 | -5.02618 | -32.5859 | 0.002402 | 0.046494 | TMEM108 |
| ENSG0000 | -1.618 | -3.0695 | 7.17E-08 | 7.95E-06 | CXCL14 |
| ENSG0000 | -5.89066 | -59.3288 | 0.000148 | 0.00558 | TAAR1 |
| ENSG0000 | -1.67371 | -3.19034 | 0.037333 | 0.289041 | ZAN |
| ENSG0000 | -1.07644 | -2.10883 | 8.71E-08 | 9.45E-06 | GPC3 |
| ENSG0000 | -2.55437 | -5.87412 | 0.094365 | 0.47552 | SNTG1 |
| ENSG0000 | -1.50632 | -2.84085 | 0.049215 | 0.338506 | LCN9 |
| ENSG0000 | -4.7526 | -26.9573 | 0.005781 | 0.086358 | ZP1 |
| ENSG0000 | -1.096 | -2.13762 | 3.60E-24 | 3.61E-21 | HMGA2 |
| ENSG0000 | -5.90666 | -59.9905 | 0.001803 | 0.038207 | FREM2 |
| ENSG0000 | -1.19433 | -2.28839 | 0.020973 | 0.202432 | CACNA1C |
| ENSG0000 | -1.26492 | -2.40313 | 0.096084 | 0.479543 | BTBD11 |
| ENSG0000 | -1.00683 | -2.00948 | 2.58E-05 | 0.001336 | ADAMTS12 |
| ENSG0000 | -1.69904 | -3.24684 | 1.73E-05 | 0.000955 | SPOCK1 |
| ENSG0000 | -2.29075 | -4.8931 | 0.085466 | 0.450949 | DMP1 |
| ENSG0000 | -3.68713 | -12.8806 | 0.079026 | 0.431664 | SLC16A12 |
| ENSG0000 | -2.06033 | -4.17083 | 0.008795 | 0.114427 | CNTNAP4 |
| ENSG0000 | -2.55013 | -5.85685 | 0.076465 | 0.424948 | FRMD1 |
| ENSG0000 | -1.80215 | -3.48739 | 0.002303 | 0.045117 | MMP21 |
| ENSG0000 | -1.08099 | -2.11549 | 0.068914 | 0.40388 | GRIP1 |
| ENSG0000 | -2.53678 | -5.80292 | 0.087411 | 0.456032 | LHFPL4 |

| | | | | | |
|---|---|---|---|---|---|
| ENSG0000 | -5.06587 | -33.4949 | 0.00443 | 0.071539 | KLHL40 |
| ENSG0000 | -2.58126 | -5.98463 | 0.008045 | 0.107759 | STEAP2 |
| ENSG0000 | -1.3761 | -2.59565 | 1.06E-05 | 0.000631 | GALNT14 |
| ENSG0000 | -1.23808 | -2.35884 | 7.00E-05 | 0.003077 | ESYT3 |
| ENSG0000 | -1.32606 | -2.50718 | 1.13E-13 | 3.76E-11 | CD1A |
| ENSG0000 | -1.79965 | -3.48137 | 0.084298 | 0.447815 | TGM7 |
| ENSG0000 | -4.72407 | -26.4293 | 0.001799 | 0.038172 | FCN2 |
| ENSG0000 | -3.11682 | -8.67474 | 0.026107 | 0.232313 | PTGER1 |
| ENSG0000 | -1.26416 | -2.40188 | 0.024865 | 0.225802 | NPHS1 |
| ENSG0000 | -1.49164 | -2.81209 | 0.007108 | 0.099291 | PLXDC1 |
| ENSG0000 | -4.9435 | -30.7711 | 0.005579 | 0.084038 | BICDL2 |
| ENSG0000 | -3.06441 | -8.36525 | 0.03233 | 0.265181 | C1QTNF7 |
| ENSG0000 | -1.72301 | -3.30124 | 0.02063 | 0.200573 | DAPL1 |
| ENSG0000 | -1.27031 | -2.41213 | 0.019962 | 0.196118 | TAFA4 |
| ENSG0000 | -1.09544 | -2.13678 | 0.002673 | 0.049663 | SPTA1 |
| ENSG0000 | -2.0628 | -4.17796 | 0.044201 | 0.319545 | C3orf49 |
| ENSG0000 | -4.64015 | -24.9358 | 0.00846 | 0.111401 | ERICH6 |
| ENSG0000 | -3.2409 | -9.45381 | 0.015765 | 0.167376 | DNAJC5G |
| ENSG0000 | -4.98357 | -31.6377 | 0.005948 | 0.087791 | EMCN |
| ENSG0000 | -1.27652 | -2.42255 | 1.95E-28 | 4.57E-25 | SLC9B2 |
| ENSG0000 | -1.70235 | -3.25431 | 0.022886 | 0.214424 | ASB5 |
| ENSG0000 | -2.94386 | -7.69466 | 0.075082 | 0.421702 | HTR4 |
| ENSG0000 | -1.90951 | -3.75681 | 1.31E-07 | 1.37E-05 | IL22RA2 |
| ENSG0000 | -1.11067 | -2.15946 | 5.06E-09 | 7.28E-07 | STEAP1 |
| ENSG0000 | -2.48814 | -5.61053 | 0.094786 | 0.476823 | INTS4P1 |
| ENSG0000 | -1.66894 | -3.17981 | 1.39E-27 | 2.60E-24 | COL1A2 |
| ENSG0000 | -1.49032 | -2.80952 | 0.06126 | 0.379145 | TMEM184A |
| ENSG0000 | -1.15835 | -2.23202 | 0.021142 | 0.203298 | TMC1 |
| ENSG0000 | -2.37069 | -5.17189 | 0.067274 | 0.399189 | CFAP47 |
| ENSG0000 | -2.36638 | -5.15644 | 1.89E-24 | 2.04E-21 | PKNOX2 |
| ENSG0000 | -1.54654 | -2.92116 | 0.000311 | 0.00983 | SLC39A2 |
| ENSG0000 | -1.39145 | -2.62342 | 0.081713 | 0.44018 | ERICH6B |
| ENSG0000 | -1.5392 | -2.90632 | 0.096053 | 0.479543 | PKD1L2 |
| ENSG0000 | -1.16567 | -2.24338 | 0.042748 | 0.312783 | FAM86GP |
| ENSG0000 | -2.7603 | -6.77536 | 0.055473 | 0.3612 | LDHC |
| ENSG0000 | -1.83077 | -3.55728 | 0.102358 | 0.49639 | NAALADL1 |
| ENSG0000 | -5.53466 | -46.3553 | 0.000316 | 0.009944 | OR1F1 |
| ENSG0000 | -1.54538 | -2.91881 | 2.57E-05 | 0.001333 | DEGS2 |
| ENSG0000 | -2.00951 | -4.02645 | 0.005422 | 0.082428 | KCNG4 |
| ENSG0000 | -1.6257 | -3.08591 | 0.002623 | 0.049022 | PYDC1 |
| ENSG0000 | -1.31005 | -2.4795 | 1.42E-24 | 1.59E-21 | HTRA3 |
| ENSG0000 | -1.24988 | -2.37821 | 0.052722 | 0.351758 | KRT15 |
| ENSG0000 | -5.00757 | -32.1683 | 0.002908 | 0.052795 | COL6A5 |
| ENSG0000 | -1.04221 | -2.05938 | 0.000182 | 0.006544 | MRGPRF |
| ENSG0000 | -3.5631 | -11.8195 | 0.032754 | 0.267145 | SLC22A13 |
| ENSG0000 | -3.5522 | -11.7305 | 0.036859 | 0.286384 | XKR3 |
| ENSG0000 | -2.72911 | -6.63046 | 0.047195 | 0.331201 | CCR9 |

| | | | | | |
|---|---|---|---|---|---|
| ENSG0000 | -2.56235 | -5.90671 | 6.31E-07 | 5.69E-05 | SULT1B1 |
| ENSG0000 | -1.31369 | -2.48576 | 0.000154 | 0.005771 | WFIKKN2 |
| ENSG0000 | -1.0179 | -2.02497 | 4.10E-10 | 7.33E-08 | SLCO4C1 |
| ENSG0000 | -1.56465 | -2.95805 | 4.60E-05 | 0.002187 | TNK1 |
| ENSG0000 | -4.29957 | -19.6924 | 0.022188 | 0.21004 | ODAPH |
| ENSG0000 | -2.54254 | -5.82615 | 0.088192 | 0.458008 | OR10P1 |
| ENSG0000 | -5.02402 | -32.5372 | 0.009416 | 0.119884 | DMBT1L1 |
| ENSG0000 | -5.26832 | -38.5408 | 0.001221 | 0.028554 | TCIM |
| ENSG0000 | -5.08254 | -33.8841 | 0.002893 | 0.052595 | LINC01561 |
| ENSG0000 | -4.73534 | -26.6367 | 0.010019 | 0.12473 | GJD4 |
| ENSG0000 | -1.28021 | -2.42874 | 0.025233 | 0.227798 | CD19 |
| ENSG0000 | -1.88271 | -3.68766 | 0.066136 | 0.394946 | ERICH5 |
| ENSG0000 | -3.1938 | -9.15016 | 0.07014 | 0.407741 | NHLH2 |
| ENSG0000 | -1.20166 | -2.30004 | 2.79E-25 | 3.40E-22 | CD163L1 |
| ENSG0000 | -3.12621 | -8.73136 | 0.026837 | 0.236122 | VN1R1 |
| ENSG0000 | -1.14014 | -2.20402 | 2.12E-07 | 2.08E-05 | TMEM151B |
| ENSG0000 | -1.21632 | -2.32353 | 0.000596 | 0.016344 | DNAJC22 |
| ENSG0000 | -1.37169 | -2.58773 | 0.021946 | 0.208456 | ARMC10P1 |
| ENSG0000 | -3.69355 | -12.9381 | 0.040559 | 0.30337 | GP5 |
| ENSG0000 | -2.75007 | -6.72751 | 0.05551 | 0.3612 | C12orf42 |
| ENSG0000 | -6.01259 | -64.5611 | 0.002111 | 0.042658 | LINC00311 |
| ENSG0000 | -2.59753 | -6.05249 | 1.11E-13 | 3.75E-11 | PCARE |
| ENSG0000 | -1.3429 | -2.53661 | 0.039829 | 0.300459 | HTR1D |
| ENSG0000 | -3.63252 | -12.4022 | 0.008128 | 0.108248 | RNF151 |
| ENSG0000 | -1.0641 | -2.09087 | 0.003118 | 0.055568 | FCER1A |
| ENSG0000 | -1.34987 | -2.54889 | 4.46E-07 | 4.11E-05 | SSC5D |
| ENSG0000 | -2.18498 | -4.54722 | 0.029019 | 0.247931 | None |
| ENSG0000 | -5.28311 | -38.9381 | 0.004538 | 0.072669 | ADGRD2 |
| ENSG0000 | -1.76172 | -3.39103 | 0.02787 | 0.241344 | MAP3K15 |
| ENSG0000 | -1.85215 | -3.61037 | 3.66E-28 | 7.89E-25 | F2R |
| ENSG0000 | -5.00816 | -32.1815 | 0.001567 | 0.034543 | CFAP65 |
| ENSG0000 | -5.71006 | -52.348 | 0.001598 | 0.035011 | ODF3L2 |
| ENSG0000 | -3.71771 | -13.1565 | 0.031684 | 0.262001 | None |
| ENSG0000 | -2.68817 | -6.44493 | 0.062639 | 0.383785 | OR7E115P |
| ENSG0000 | -1.01148 | -2.01597 | 1.86E-05 | 0.001011 | SPNS2 |
| ENSG0000 | -1.21376 | -2.31941 | 0.056683 | 0.364344 | RPSAP19 |
| ENSG0000 | -1.22166 | -2.33214 | 1.34E-05 | 0.000768 | EPHA10 |
| ENSG0000 | -5.69832 | -51.9237 | 6.13E-05 | 0.002769 | SH2D7 |
| ENSG0000 | -1.02878 | -2.0403 | 4.70E-26 | 6.59E-23 | RFLNB |
| ENSG0000 | -2.27289 | -4.8329 | 0.059519 | 0.37387 | ACP7 |
| ENSG0000 | -1.16146 | -2.23684 | 0.074275 | 0.418932 | SLC35F3 |
| ENSG0000 | -1.89189 | -3.7112 | 0.009229 | 0.118441 | EMILIN3 |
| ENSG0000 | -1.2899 | -2.44512 | 0.007597 | 0.104041 | ARSI |
| ENSG0000 | -1.41726 | -2.67078 | 0.001449 | 0.032647 | CLDN5 |
| ENSG0000 | -4.33043 | -20.1182 | 0.026879 | 0.236122 | CNTN2 |
| ENSG0000 | -3.39382 | -10.511 | 0.010594 | 0.128632 | DGAT2L6 |
| ENSG0000 | -1.03814 | -2.05358 | 1.92E-20 | 1.49E-17 | TACSTD2 |

| | | | | | |
|---|---|---|---|---|---|
| ENSG0000 | -5.31391 | -39.7783 | 0.000288 | 0.009277 | KRT76 |
| ENSG0000 | -1.44115 | -2.71538 | 2.79E-14 | 1.04E-11 | NOTUM |
| ENSG0000 | -1.93394 | -3.82098 | 3.27E-10 | 6.03E-08 | PRKG1 |
| ENSG0000 | -1.25378 | -2.38465 | 0.001258 | 0.029193 | SYN3 |
| ENSG0000 | -2.20718 | -4.61771 | 0.000121 | 0.004764 | KCNQ5 |
| ENSG0000 | -3.75555 | -13.5062 | 0.00417 | 0.068521 | CCIN |
| ENSG0000 | -3.7664 | -13.6081 | 0.069495 | 0.405762 | NAP1L3 |
| ENSG0000 | -1.27808 | -2.42517 | 0.000393 | 0.011804 | None |
| ENSG0000 | -4.94671 | -30.8395 | 0.005023 | 0.078103 | LHFPL3 |
| ENSG0000 | -5.5189 | -45.8517 | 0.001005 | 0.024551 | DPPA3 |
| ENSG0000 | -1.79065 | -3.45971 | 0.094246 | 0.475366 | C5orf52 |
| ENSG0000 | -1.63328 | -3.10217 | 0.001088 | 0.026086 | KRT18P59 |
| ENSG0000 | -1.35936 | -2.56572 | 0.101174 | 0.493021 | KBTBD12 |
| ENSG0000 | -1.00987 | -2.01373 | 0.005547 | 0.083679 | DNAJB13 |
| ENSG0000 | -1.20137 | -2.29957 | 0.066052 | 0.394778 | HS6ST1P1 |
| ENSG0000 | -2.04334 | -4.12199 | 4.45E-26 | 6.56E-23 | RUFY4 |
| ENSG0000 | -3.53089 | -11.5586 | 0.020878 | 0.201793 | PPP3R2 |
| ENSG0000 | -6.42655 | -86.0168 | 2.38E-05 | 0.00125 | INSC |
| ENSG0000 | -2.74993 | -6.72686 | 0.074238 | 0.418808 | DUSP21 |
| ENSG0000 | -1.73512 | -3.32908 | 1.08E-18 | 6.57E-16 | SP6 |
| ENSG0000 | -4.37135 | -20.697 | 0.035237 | 0.27959 | NKAPL |
| ENSG0000 | -3.36496 | -10.3027 | 0.009793 | 0.122962 | FAM180A |
| ENSG0000 | -2.85773 | -7.24874 | 0.00778 | 0.105622 | THEM5 |
| ENSG0000 | -5.27141 | -38.6236 | 0.002938 | 0.053202 | SPTSSB |
| ENSG0000 | -1.98467 | -3.95771 | 3.90E-27 | 6.83E-24 | COL27A1 |
| ENSG0000 | -2.66828 | -6.35673 | 0.064802 | 0.39156 | None |
| ENSG0000 | -4.32713 | -20.0723 | 0.019678 | 0.194239 | HCAR1 |
| ENSG0000 | -1.05891 | -2.08336 | 1.35E-06 | 0.00011 | TPM2 |
| ENSG0000 | -4.68259 | -25.6803 | 0.015329 | 0.164229 | Y_RNA |
| ENSG0000 | -3.87508 | -14.6729 | 0.00406 | 0.067309 | Y_RNA |
| ENSG0000 | -3.75947 | -13.543 | 0.046352 | 0.327901 | RNU6-824P |
| ENSG0000 | -2.56286 | -5.90879 | 0.065877 | 0.3942 | Y_RNA |
| ENSG0000 | -4.36322 | -20.5806 | 0.029474 | 0.250144 | Y_RNA |
| ENSG0000 | -4.68566 | -25.735 | 0.007074 | 0.098889 | RNU1-91P |
| ENSG0000 | -3.22233 | -9.3329 | 0.008372 | 0.110775 | IGBP1-AS1 |
| ENSG0000 | -4.03481 | -16.3907 | 0.000274 | 0.008959 | NPY4R |
| ENSG0000 | -2.94509 | -7.70122 | 0.05356 | 0.354183 | LINC02872 |
| ENSG0000 | -2.72414 | -6.60765 | 0.06826 | 0.401728 | ZSCAN5C |
| ENSG0000 | -3.68881 | -12.8956 | 0.060448 | 0.377006 | PSORS1C1 |
| ENSG0000 | -1.87075 | -3.65723 | 0.032778 | 0.267264 | None |
| ENSG0000 | -1.4087 | -2.65497 | 0.084701 | 0.449024 | GRXCR2 |
| ENSG0000 | -5.04907 | -33.1071 | 0.006856 | 0.096916 | LINC01121 |
| ENSG0000 | -5.30162 | -39.441 | 0.000452 | 0.013392 | SLCO6A1 |
| ENSG0000 | -6.03817 | -65.7161 | 1.54E-05 | 0.000864 | SMIM45 |
| ENSG0000 | -2.52192 | -5.74348 | 0.077142 | 0.426609 | RNASE13 |
| ENSG0000 | -2.82315 | -7.07705 | 0.089871 | 0.462447 | SNORA22B |
| ENSG0000 | -3.24147 | -9.4576 | 0.037027 | 0.287534 | SNORA69 |

| | | | | | |
|---|---|---|---|---|---|
| ENSG0000 | -5.1202 | -34.7804 | 0.008912 | 0.11536 | None |
| ENSG0000 | -5.08356 | -33.9082 | 0.002746 | 0.050576 | Y_RNA |
| ENSG0000 | -4.2988 | -19.6819 | 0.011435 | 0.135044 | None |
| ENSG0000 | -4.68598 | -25.7406 | 0.006052 | 0.088945 | MIR199A1 |
| ENSG0000 | -5.00732 | -32.1627 | 0.002149 | 0.043129 | SNORD72 |
| ENSG0000 | -4.30406 | -19.7538 | 0.038391 | 0.293495 | None |
| ENSG0000 | -1.69904 | -3.24685 | 0.041489 | 0.307425 | LPAL2 |
| ENSG0000 | -2.14066 | -4.40965 | 0.078106 | 0.42918 | None |
| ENSG0000 | -1.59219 | -3.01507 | 0.041953 | 0.309264 | PDCL3P5 |
| ENSG0000 | -3.58538 | -12.0035 | 0.014955 | 0.161028 | SCYL2P1 |
| ENSG0000 | -1.93426 | -3.82183 | 0.08328 | 0.444766 | NUDCP1 |
| ENSG0000 | -1.00845 | -2.01175 | 0.01784 | 0.181508 | ARHGEF35 |
| ENSG0000 | -1.62192 | -3.07785 | 0.070678 | 0.409452 | None |
| ENSG0000 | -4.96306 | -31.1911 | 0.005779 | 0.086358 | RANP6 |
| ENSG0000 | -1.08622 | -2.12317 | 0.036625 | 0.285123 | TTLL13P |
| ENSG0000 | -2.26645 | -4.81137 | 0.04842 | 0.335541 | RPL11P3 |
| ENSG0000 | -1.32769 | -2.51001 | 0.007202 | 0.100302 | RPS16P2 |
| ENSG0000 | -1.14329 | -2.20885 | 0.005822 | 0.086637 | RPL21P75 |
| ENSG0000 | -4.32032 | -19.9777 | 0.0084 | 0.111091 | RPL3P6 |
| ENSG0000 | -3.22055 | -9.32142 | 0.035798 | 0.281812 | KCTD9P4 |
| ENSG0000 | -2.82289 | -7.07577 | 0.08816 | 0.457928 | ZNF705EP |
| ENSG0000 | -2.14281 | -4.41622 | 0.054379 | 0.356624 | GTF2IRD2P1 |
| ENSG0000 | -1.23338 | -2.35117 | 0.032082 | 0.264219 | LRRC37A11P |
| ENSG0000 | -5.05454 | -33.2329 | 0.000539 | 0.01522 | HMGN2P15 |
| ENSG0000 | -1.74835 | -3.35975 | 0.088131 | 0.457928 | None |
| ENSG0000 | -5.45689 | -43.9225 | 0.000177 | 0.006396 | EEF1A1P29 |
| ENSG0000 | -5.07208 | -33.6393 | 0.00073 | 0.019215 | GPX1P2 |
| ENSG0000 | -5.47943 | -44.6142 | 0.000457 | 0.013487 | ZNF663P |
| ENSG0000 | -2.7697 | -6.81965 | 0.06192 | 0.381277 | ANKRD20A11P |
| ENSG0000 | -4.73487 | -26.6279 | 0.008541 | 0.112203 | RPL22P12 |
| ENSG0000 | -5.08401 | -33.9188 | 0.003476 | 0.059815 | None |
| ENSG0000 | -4.56014 | -23.5905 | 0.047956 | 0.333531 | RPS3AP2 |
| ENSG0000 | -5.47919 | -44.6068 | 0.000697 | 0.018643 | CATSPERZ |
| ENSG0000 | -1.32245 | -2.5009 | 0.046459 | 0.328202 | None |
| ENSG0000 | -1.12169 | -2.17601 | 0.002855 | 0.052069 | ZSCAN12P1 |
| ENSG0000 | -2.45889 | -5.49792 | 0.066795 | 0.397273 | HNRNPA1P12 |
| ENSG0000 | -5.28502 | -38.9896 | 0.002543 | 0.048123 | None |
| ENSG0000 | -6.01651 | -64.7366 | 1.43E-05 | 0.000811 | IGBP1-AS2 |
| ENSG0000 | -2.05817 | -4.16457 | 0.014741 | 0.15952 | SNORA11F |
| ENSG0000 | -5.38933 | -41.913 | 0.010784 | 0.130093 | MIR548L |
| ENSG0000 | -1.51807 | -2.86407 | 0.062088 | 0.381826 | RNU6ATAC18P |
| ENSG0000 | -5.27088 | -38.6094 | 0.000529 | 0.014974 | MIR1303 |
| ENSG0000 | -5.64143 | -49.916 | 0.011745 | 0.137097 | RNU6ATAC10P |
| ENSG0000 | -1.37299 | -2.59006 | 0.001222 | 0.028554 | HMSD |
| ENSG0000 | -2.97107 | -7.84118 | 0.099015 | 0.488096 | RNU4-86P |
| ENSG0000 | -4.30117 | -19.7142 | 0.045979 | 0.32584 | RN7SKP151 |
| ENSG0000 | -3.72032 | -13.1804 | 0.039718 | 0.299801 | RNU2-39P |

| | | | | | |
|---|---|---|---|---|---|
| ENSG0000 | -6.05135 | -66.319 | 0.000669 | 0.017926 | RNU2-70P |
| ENSG0000 | -4.89573 | -29.7687 | 0.04583 | 0.325192 | RNU2-40P |
| ENSG0000 | -5.06651 | -33.5097 | 0.003156 | 0.055955 | MEG9 |
| ENSG0000 | -4.31815 | -19.9477 | 0.008033 | 0.107759 | RBM7P1 |
| ENSG0000 | -1.71874 | -3.2915 | 0.093465 | 0.473126 | CKMT1A |
| ENSG0000 | -6.07858 | -67.5827 | 6.95E-06 | 0.000442 | None |
| ENSG0000 | -1.14675 | -2.21414 | 0.04923 | 0.338525 | LINC02593 |
| ENSG0000 | -6.15571 | -71.2941 | 1.33E-06 | 0.000109 | LINP1 |
| ENSG0000 | -3.69056 | -12.9113 | 0.046753 | 0.329572 | LINC01870 |
| ENSG0000 | -4.7729 | -27.3391 | 0.004052 | 0.067259 | NPM1P19 |
| ENSG0000 | -3.098 | -8.56231 | 0.019681 | 0.194239 | LINC01449 |
| ENSG0000 | -1.03492 | -2.049 | 0.045372 | 0.323855 | RABGEF1P3 |
| ENSG0000 | -1.00449 | -2.00624 | 0.078978 | 0.431653 | PRR29 |
| ENSG0000 | -1.97137 | -3.92139 | 0.079129 | 0.431979 | None |
| ENSG0000 | -2.27451 | -4.83832 | 0.085778 | 0.452 | None |
| ENSG0000 | -3.69331 | -12.9359 | 0.042351 | 0.310887 | OR4F2P |
| ENSG0000 | -1.44388 | -2.72051 | 0.006749 | 0.095843 | RPS4XP16 |
| ENSG0000 | -1.47492 | -2.77967 | 0.035776 | 0.2818 | LINC01770 |
| ENSG0000 | -3.77437 | -13.6835 | 0.092456 | 0.46995 | TNPO1P1 |
| ENSG0000 | -1.69768 | -3.24379 | 0.075327 | 0.421944 | None |
| ENSG0000 | -1.50886 | -2.84585 | 0.063184 | 0.385606 | C2CD4D |
| ENSG0000 | -3.61563 | -12.2578 | 0.070118 | 0.407698 | EIF2S2P5 |
| ENSG0000 | -1.62388 | -3.08202 | 0.00413 | 0.068178 | HNRNPA1P35 |
| ENSG0000 | -5.02565 | -32.574 | 0.006354 | 0.091817 | ADGRF5-AS1 |
| ENSG0000 | -2.28094 | -4.85994 | 0.094447 | 0.475722 | LINC02816 |
| ENSG0000 | -4.35907 | -20.5216 | 0.02067 | 0.200768 | None |
| ENSG0000 | -3.6196 | -12.2916 | 0.064101 | 0.389002 | SLC25A39P1 |
| ENSG0000 | -3.31624 | -9.96062 | 0.059091 | 0.372122 | LINC01733 |
| ENSG0000 | -1.19077 | -2.28275 | 0.092037 | 0.468775 | None |
| ENSG0000 | -1.55245 | -2.93315 | 0.028876 | 0.246932 | ZKSCAN8P1 |
| ENSG0000 | -3.61056 | -12.2148 | 0.078639 | 0.43081 | RPL15P14 |
| ENSG0000 | -1.33685 | -2.526 | 0.037703 | 0.29045 | None |
| ENSG0000 | -2.80793 | -7.00279 | 0.069858 | 0.406746 | FUNDC2P4 |
| ENSG0000 | -1.67376 | -3.19045 | 7.32E-05 | 0.003164 | LINC00511 |
| ENSG0000 | -1.16783 | -2.24674 | 0.021375 | 0.204227 | GTF2IP7 |
| ENSG0000 | -2.27645 | -4.84485 | 1.35E-07 | 1.40E-05 | None |
| ENSG0000 | -5.28381 | -38.9569 | 0.013124 | 0.147365 | None |
| ENSG0000 | -4.70827 | -26.1415 | 0.01328 | 0.148519 | None |
| ENSG0000 | -3.37585 | -10.3808 | 0.030467 | 0.2555 | WDR35-DT |
| ENSG0000 | -2.74553 | -6.70638 | 0.027615 | 0.240403 | LINC01537 |
| ENSG0000 | -5.32038 | -39.957 | 0.001228 | 0.028629 | HSD11B1-AS1 |
| ENSG0000 | -4.30463 | -19.7617 | 0.025703 | 0.230266 | None |
| ENSG0000 | -1.45334 | -2.73842 | 0.008088 | 0.108039 | DNMBP-AS1 |
| ENSG0000 | -3.72497 | -13.2229 | 0.060562 | 0.377151 | None |
| ENSG0000 | -4.73615 | -26.6516 | 0.028174 | 0.242929 | None |
| ENSG0000 | -4.30323 | -19.7425 | 0.037846 | 0.291063 | TEX46 |
| ENSG0000 | -2.05846 | -4.16542 | 0.03162 | 0.261789 | PSPC1P1 |

| | | | | | |
|---|---|---|---|---|---|
| ENSG0000 | -4.20916 | -18.4962 | 0.032247 | 0.265016 | LINC02344 |
| ENSG0000 | -4.96735 | -31.2839 | 0.000884 | 0.022356 | None |
| ENSG0000 | -1.82439 | -3.54158 | 0.057695 | 0.367494 | None |
| ENSG0000 | -1.69692 | -3.24209 | 0.066137 | 0.394946 | None |
| ENSG0000 | -3.0191 | -8.10661 | 0.054138 | 0.355874 | None |
| ENSG0000 | -2.54976 | -5.85536 | 0.065865 | 0.3942 | None |
| ENSG0000 | -1.31304 | -2.48465 | 0.089816 | 0.462246 | ITPRIP-AS1 |
| ENSG0000 | -1.08894 | -2.12717 | 0.088762 | 0.459368 | None |
| ENSG0000 | -4.99329 | -31.8515 | 0.001841 | 0.038748 | None |
| ENSG0000 | -1.62096 | -3.0758 | 0.046149 | 0.326848 | RPL13AP19 |
| ENSG0000 | -5.48979 | -44.9358 | 0.002458 | 0.047189 | None |
| ENSG0000 | -3.75561 | -13.5067 | 0.041284 | 0.30667 | DUTP1 |
| ENSG0000 | -1.31868 | -2.49438 | 0.081369 | 0.439415 | PPP1R12BP1 |
| ENSG0000 | -3.6598 | -12.6389 | 0.050245 | 0.342628 | RLIMP1 |
| ENSG0000 | -3.1191 | -8.68848 | 0.013461 | 0.149709 | LINC01204 |
| ENSG0000 | -5.53335 | -46.3132 | 0.000356 | 0.010964 | None |
| ENSG0000 | -5.23165 | -37.5738 | 0.002604 | 0.048831 | RPS21P1 |
| ENSG0000 | -2.91336 | -7.53372 | 0.082784 | 0.443556 | KHSRPP1 |
| ENSG0000 | -4.76431 | -27.1769 | 0.002441 | 0.047056 | LINC01435 |
| ENSG0000 | -4.6363 | -24.8694 | 0.011346 | 0.134294 | SNAP47-AS1 |
| ENSG0000 | -3.24256 | -9.4647 | 0.040852 | 0.304828 | HSPD1P6 |
| ENSG0000 | -6.10523 | -68.8427 | 4.52E-06 | 0.000303 | TOMM20P4 |
| ENSG0000 | -2.37312 | -5.1806 | 0.081615 | 0.440149 | HCG24 |
| ENSG0000 | -1.38298 | -2.60807 | 0.056698 | 0.364344 | None |
| ENSG0000 | -1.83388 | -3.56495 | 0.063577 | 0.387157 | |
| ENSG0000 | -2.7505 | -6.7295 | 0.070994 | 0.409896 | YY1P1 |
| ENSG0000 | -1.12147 | -2.17569 | 0.089288 | 0.460713 | None |
| ENSG0000 | -2.56928 | -5.93512 | 0.093512 | 0.473279 | SRGAP2-AS1 |
| ENSG0000 | -1.28486 | -2.43658 | 0.012452 | 0.142328 | LINC01614 |
| ENSG0000 | -4.27582 | -19.3709 | 0.012469 | 0.142347 | PPIHP1 |
| ENSG0000 | -1.495 | -2.81864 | 2.00E-06 | 0.000155 | DPP4-DT |
| ENSG0000 | -1.57414 | -2.97759 | 0.073643 | 0.417044 | RPL7P32 |
| ENSG0000 | -5.02443 | -32.5465 | 0.004691 | 0.07442 | NEK2-DT |
| ENSG0000 | -2.49315 | -5.63006 | 0.059556 | 0.37387 | NPM1P9 |
| ENSG0000 | -5.2337 | -37.627 | 0.00034 | 0.010579 | SPATA20P1 |
| ENSG0000 | -4.66059 | -25.2917 | 0.013448 | 0.149626 | FARP1-AS1 |
| ENSG0000 | -2.47792 | -5.57093 | 0.095012 | 0.477426 | IGKV1OR2-108 |
| ENSG0000 | -1.99473 | -3.9854 | 3.45E-08 | 4.16E-06 | TIMM8AP1 |
| ENSG0000 | -3.08768 | -8.50128 | 0.060653 | 0.37761 | RNASEH1P2 |
| ENSG0000 | -5.01037 | -32.2308 | 0.001857 | 0.039018 | RPS6KA2-AS1 |
| ENSG0000 | -3.68686 | -12.8782 | 0.08154 | 0.439912 | None |
| ENSG0000 | -5.64283 | -49.9646 | 0.007625 | 0.104225 | None |
| ENSG0000 | -3.32494 | -10.0209 | 0.022171 | 0.209951 | None |
| ENSG0000 | -3.12883 | -8.74726 | 0.00923 | 0.118441 | FAAHP1 |
| ENSG0000 | -2.96935 | -7.83181 | 0.052081 | 0.349565 | RPLP1P13 |
| ENSG0000 | -1.4524 | -2.73663 | 0.00037 | 0.01124 | TMEM114 |
| ENSG0000 | -5.30659 | -39.577 | 0.016356 | 0.171453 | None |

| | | | | | |
|---|---|---|---|---|---|
| ENSG0000 | -4.27258 | -19.3275 | 0.016994 | 0.17544 | GXYLT1P3 |
| ENSG0000 | -1.05864 | -2.08296 | 0.04296 | 0.313861 | TGFB2-AS1 |
| ENSG0000 | -1.68675 | -3.2193 | 2.97E-06 | 0.000213 | |
| ENSG0000 | -6.30711 | -79.1823 | 6.61E-07 | 5.89E-05 | CSPG4BP |
| ENSG0000 | -3.74947 | -13.4494 | 0.039307 | 0.297703 | None |
| ENSG0000 | -2.76248 | -6.78562 | 0.038351 | 0.293281 | KIZ-AS1 |
| ENSG0000 | -3.50147 | -11.3252 | 0.008772 | 0.114231 | None |
| ENSG0000 | -3.62146 | -12.3074 | 0.061452 | 0.379832 | None |
| ENSG0000 | -1.39433 | -2.62867 | 0.002646 | 0.049343 | None |
| ENSG0000 | -2.40765 | -5.30608 | 0.011146 | 0.132747 | RPL23AP93 |
| ENSG0000 | -3.43594 | -10.8224 | 0.011482 | 0.135487 | None |
| ENSG0000 | -5.25975 | -38.3127 | 0.022899 | 0.214443 | None |
| ENSG0000 | -3.48827 | -11.2221 | 0.020091 | 0.197108 | MROH3P |
| ENSG0000 | -3.78059 | -13.7426 | 0.002006 | 0.041184 | PHC2-AS1 |
| ENSG0000 | -3.41423 | -10.6607 | 0.018118 | 0.183197 | TLR8-AS1 |
| ENSG0000 | -2.49294 | -5.62923 | 0.087772 | 0.45661 | None |
| ENSG0000 | -3.72654 | -13.2373 | 0.070229 | 0.408005 | None |
| ENSG0000 | -2.97935 | -7.88629 | 0.098105 | 0.485317 | CCNQP1 |
| ENSG0000 | -4.70924 | -26.1591 | 0.0085 | 0.111825 | RPSAP13 |
| ENSG0000 | -5.12029 | -34.7825 | 0.02026 | 0.198278 | BTF3P6 |
| ENSG0000 | -2.95256 | -7.74123 | 0.026797 | 0.235994 | TPT1P1 |
| ENSG0000 | -2.17114 | -4.50378 | 0.060754 | 0.377827 | None |
| ENSG0000 | -1.86236 | -3.63602 | 0.052298 | 0.350512 | PDE4DIPP7 |
| ENSG0000 | -1.02245 | -2.03136 | 0.083915 | 0.446748 | ST13P18 |
| ENSG0000 | -4.78284 | -27.5282 | 0.010135 | 0.125519 | None |
| ENSG0000 | -2.57644 | -5.96467 | 0.088741 | 0.459368 | None |
| ENSG0000 | -2.89902 | -7.45921 | 0.047498 | 0.331589 | LINC02765 |
| ENSG0000 | -4.20587 | -18.4541 | 0.037686 | 0.290428 | PPP1R2P1 |
| ENSG0000 | -4.29806 | -19.6719 | 0.013751 | 0.151969 | NDUFB1P2 |
| ENSG0000 | -1.25803 | -2.3917 | 0.081263 | 0.439181 | C2CD4D-AS1 |
| ENSG0000 | -1.37876 | -2.60045 | 1.64E-06 | 0.000132 | LINC01135 |
| ENSG0000 | -5.01128 | -32.2511 | 0.000901 | 0.022633 | LINC02623 |
| ENSG0000 | -2.47745 | -5.56911 | 0.019838 | 0.195307 | SHISA8 |
| ENSG0000 | -4.68033 | -25.6401 | 0.02206 | 0.209249 | None |
| ENSG0000 | -1.07012 | -2.09961 | 0.01185 | 0.137978 | ID2-AS1 |
| ENSG0000 | -1.43579 | -2.70531 | 0.096067 | 0.479543 | None |
| ENSG0000 | -5.77516 | -54.7642 | 0.000284 | 0.009174 | None |
| ENSG0000 | -2.95185 | -7.7374 | 0.040743 | 0.304339 | MRPL53P1 |
| ENSG0000 | -5.67111 | -50.9536 | 0.000261 | 0.008669 | EPN2-AS1 |
| ENSG0000 | -5.04321 | -32.9729 | 0.002135 | 0.042986 | None |
| ENSG0000 | -1.38443 | -2.61069 | 7.89E-08 | 8.64E-06 | None |
| ENSG0000 | -1.38365 | -2.60928 | 6.42E-06 | 0.000412 | LINC01814 |
| ENSG0000 | -1.60774 | -3.04773 | 0.091366 | 0.466767 | LINC02889 |
| ENSG0000 | -2.53895 | -5.81165 | 0.097753 | 0.484095 | None |
| ENSG0000 | -3.49228 | -11.2533 | 0.011315 | 0.134064 | None |
| ENSG0000 | -1.66883 | -3.17956 | 0.099627 | 0.489137 | EEF1E1P1 |
| ENSG0000 | -1.58876 | -3.0079 | 0.034302 | 0.27521 | None |

| | | | | | |
|---|---|---|---|---|---|
| ENSG0000 | -5.81166 | -56.1673 | 7.02E-05 | 0.003083 | RRAS2P1 |
| ENSG0000 | -3.38134 | -10.4204 | 0.035976 | 0.282501 | SLC35G4 |
| ENSG0000 | -2.74896 | -6.72231 | 0.031097 | 0.258363 | None |
| ENSG0000 | -1.64062 | -3.11801 | 3.21E-08 | 3.90E-06 | RPL21P133 |
| ENSG0000 | -5.53501 | -46.3664 | 0.000996 | 0.024403 | KLF2P4 |
| ENSG0000 | -1.15583 | -2.22813 | 0.037335 | 0.289041 | LINC01844 |
| ENSG0000 | -1.50459 | -2.83745 | 0.001033 | 0.025097 | None |
| ENSG0000 | -1.47353 | -2.777 | 0.089004 | 0.460012 | LINC01554 |
| ENSG0000 | -2.71813 | -6.58021 | 0.064895 | 0.391713 | CLCA4-AS1 |
| ENSG0000 | -3.2607 | -9.58451 | 0.081121 | 0.438874 | LINC01141 |
| ENSG0000 | -5.6788 | -51.2259 | 3.46E-05 | 0.001713 | CARD11-AS1 |
| ENSG0000 | -5.69486 | -51.7992 | 0.001618 | 0.035285 | RAI1-AS1 |
| ENSG0000 | -3.68964 | -12.9031 | 0.053423 | 0.353809 | RPSAP11 |
| ENSG0000 | -1.74434 | -3.35041 | 0.07653 | 0.424948 | LINC01275 |
| ENSG0000 | -5.26501 | -38.4525 | 0.008001 | 0.10754 | None |
| ENSG0000 | -2.90599 | -7.49532 | 0.022745 | 0.213588 | IFNWP2 |
| ENSG0000 | -2.75189 | -6.73599 | 0.086071 | 0.453116 | EEF1A1P30 |
| ENSG0000 | -5.35102 | -40.8149 | 0.012821 | 0.144955 | EEF1A1P31 |
| ENSG0000 | -2.49797 | -5.64888 | 0.044023 | 0.318739 | DDX39BP2 |
| ENSG0000 | -2.06392 | -4.18121 | 0.048044 | 0.333808 | ABCA17P |
| ENSG0000 | -1.01804 | -2.02517 | 0.050293 | 0.34281 | None |
| ENSG0000 | -6.1705 | -72.0285 | 0.000184 | 0.006594 | UBE2V2P3 |
| ENSG0000 | -1.13488 | -2.19601 | 0.078478 | 0.430287 | None |
| ENSG0000 | -1.33362 | -2.52035 | 0.000486 | 0.014148 | None |
| ENSG0000 | -5.32373 | -40.0499 | 0.006337 | 0.09167 | SNORD121A |
| ENSG0000 | -2.00373 | -4.01035 | 0.097299 | 0.483207 | RPL21P10 |
| ENSG0000 | -2.49737 | -5.64656 | 0.082492 | 0.442666 | PCDHA13 |
| ENSG0000 | -1.41012 | -2.65759 | 0.058857 | 0.37168 | RN7SL535P |
| ENSG0000 | -4.27605 | -19.374 | 0.011828 | 0.137773 | RPL21P39 |
| ENSG0000 | -5.5215 | -45.9342 | 0.00307 | 0.054912 | LINC02917 |
| ENSG0000 | -5.46901 | -44.293 | 0.002939 | 0.053202 | None |
| ENSG0000 | -2.75328 | -6.74246 | 0.075199 | 0.421937 | RN7SL351P |
| ENSG0000 | -3.71844 | -13.1632 | 0.033754 | 0.273001 | RN7SL263P |
| ENSG0000 | -3.96426 | -15.6085 | 0.001104 | 0.026421 | None |
| ENSG0000 | -3.75951 | -13.5433 | 0.054812 | 0.35837 | RPL12P37 |
| ENSG0000 | -3.48275 | -11.1793 | 0.013848 | 0.152866 | RN7SL559P |
| ENSG0000 | -1.42153 | -2.67869 | 0.041065 | 0.306012 | RN7SL521P |
| ENSG0000 | -3.68962 | -12.9029 | 0.058752 | 0.371494 | RN7SL798P |
| ENSG0000 | -2.86633 | -7.29209 | 0.025332 | 0.228247 | None |
| ENSG0000 | -5.10027 | -34.3031 | 0.002142 | 0.04309 | None |
| ENSG0000 | -3.61082 | -12.217 | 0.07818 | 0.429379 | LINC00635 |
| ENSG0000 | -2.73601 | -6.66227 | 0.063174 | 0.385606 | INMT |
| ENSG0000 | -2.49313 | -5.62997 | 0.060339 | 0.376814 | RPSAP52 |
| ENSG0000 | -3.66118 | -12.651 | 0.047647 | 0.332282 | None |
| ENSG0000 | -4.80153 | -27.8871 | 0.027438 | 0.239832 | DBIL5P2 |
| ENSG0000 | -2.16687 | -4.49048 | 0.051578 | 0.347406 | RPL21P11 |
| ENSG0000 | -5.48767 | -44.8698 | 0.002536 | 0.048066 | CCDC137P |

| | | | | | |
|---|---|---|---|---|---|
| ENSG0000 | -5.56955 | -47.4899 | 0.000944 | 0.023421 | RPL12P21 |
| ENSG0000 | -4.75663 | -27.0327 | 0.010617 | 0.128688 | RPL12P7 |
| ENSG0000 | -1.05112 | -2.07214 | 0.009398 | 0.119705 | None |
| ENSG0000 | -1.17662 | -2.26046 | 0.089312 | 0.46075 | MCCC1-AS1 |
| ENSG0000 | -5.73662 | -53.3206 | 0.000235 | 0.007941 | LINC02086 |
| ENSG0000 | -1.31756 | -2.49245 | 0.027115 | 0.237598 | None |
| ENSG0000 | -4.95347 | -30.9843 | 0.018238 | 0.183945 | CYP2U1-AS1 |
| ENSG0000 | -1.86588 | -3.64491 | 0.046891 | 0.330258 | None |
| ENSG0000 | -5.07177 | -33.6321 | 0.014097 | 0.15458 | DIAPH1-AS1 |
| ENSG0000 | -5.7205 | -52.7282 | 7.22E-05 | 0.003147 | RGMB-AS1 |
| ENSG0000 | -2.50169 | -5.66349 | 0.019789 | 0.195035 | FAM174A-DT |
| ENSG0000 | -5.33515 | -40.3682 | 0.000885 | 0.022356 | None |
| ENSG0000 | -1.63645 | -3.109 | 0.046162 | 0.326848 | LINC02014 |
| ENSG0000 | -5.24922 | -38.034 | 0.003358 | 0.058326 | ARL2BPP6 |
| ENSG0000 | -4.88609 | -29.5706 | 3.00E-05 | 0.001523 | None |
| ENSG0000 | -3.1339 | -8.77806 | 0.044729 | 0.321836 | LINC02485 |
| ENSG0000 | -5.88849 | -59.2395 | 0.003145 | 0.055826 | LINC02117 |
| ENSG0000 | -1.68771 | -3.22146 | 0.040406 | 0.303034 | None |
| ENSG0000 | -2.95521 | -7.75542 | 0.078482 | 0.430287 | OXCT1-AS1 |
| ENSG0000 | -4.709 | -26.1547 | 0.005585 | 0.084086 | C4orf54 |
| ENSG0000 | -2.9482 | -7.71784 | 0.021275 | 0.203663 | None |
| ENSG0000 | -1.79992 | -3.48201 | 0.063653 | 0.387289 | None |
| ENSG0000 | -2.17799 | -4.52523 | 0.063629 | 0.38723 | None |
| ENSG0000 | -1.74226 | -3.34559 | 0.082102 | 0.441502 | None |
| ENSG0000 | -5.67832 | -51.2088 | 0.010566 | 0.128407 | ENPP7P1 |
| ENSG0000 | -5.52261 | -45.9697 | 0.000496 | 0.014306 | None |
| ENSG0000 | -3.62346 | -12.3246 | 0.010562 | 0.128407 | CASC11 |
| ENSG0000 | -4.30241 | -19.7313 | 0.028854 | 0.246864 | LINC01179 |
| ENSG0000 | -2.95395 | -7.74868 | 0.043408 | 0.316001 | LINC01258 |
| ENSG0000 | -5.28388 | -38.9588 | 0.004974 | 0.077638 | None |
| ENSG0000 | -2.12157 | -4.35168 | 0.010313 | 0.126488 | PKD2L2-DT |
| ENSG0000 | -4.68062 | -25.6453 | 0.020583 | 0.200393 | None |
| ENSG0000 | -1.43291 | -2.6999 | 0.067042 | 0.398283 | None |
| ENSG0000 | -2.98048 | -7.89246 | 0.051621 | 0.347553 | None |
| ENSG0000 | -2.87162 | -7.31887 | 0.07409 | 0.418226 | None |
| ENSG0000 | -2.73745 | -6.66892 | 0.021858 | 0.207757 | None |
| ENSG0000 | -4.68613 | -25.7433 | 0.005964 | 0.087947 | PRODH2 |
| ENSG0000 | -1.939 | -3.8344 | 0.010041 | 0.124843 | None |
| ENSG0000 | -1.1522 | -2.22253 | 0.100561 | 0.491655 | None |
| ENSG0000 | -1.53982 | -2.90757 | 0.086097 | 0.453167 | H3P14 |
| ENSG0000 | -2.52314 | -5.7483 | 0.087001 | 0.455033 | LINC01337 |
| ENSG0000 | -4.74234 | -26.7662 | 0.021157 | 0.203301 | NIPAL4-DT |
| ENSG0000 | -1.01724 | -2.02405 | 0.035545 | 0.280697 | None |
| ENSG0000 | -3.68814 | -12.8897 | 0.066843 | 0.397474 | HSPA8P19 |
| ENSG0000 | -4.27777 | -19.3971 | 0.008065 | 0.10782 | SNORA70 |
| ENSG0000 | -3.07874 | -8.44878 | 0.029059 | 0.248116 | RNU4ATAC18P |
| ENSG0000 | -5.00986 | -32.2195 | 0.001387 | 0.031599 | SCARNA5 |

| | | | | | |
|---|---|---|---|---|---|
| ENSG0000 | -1.35157 | -2.55189 | 0.079912 | 0.434556 | RNU6-415P |
| ENSG0000 | -5.74352 | -53.5761 | 0.005199 | 0.080304 | RNU6-703P |
| ENSG0000 | -1.30385 | -2.46887 | 0.020248 | 0.198278 | Y_RNA |
| ENSG0000 | -4.24832 | -19.0051 | 0.015978 | 0.16893 | RNU6-731P |
| ENSG0000 | -2.98056 | -7.89291 | 0.042132 | 0.30979 | RNU6-1238P |
| ENSG0000 | -5.5237 | -46.0043 | 1.58E-05 | 0.000883 | None |
| ENSG0000 | -3.68003 | -12.8174 | 0.022989 | 0.214653 | None |
| ENSG0000 | -2.25928 | -4.78752 | 0.100379 | 0.491431 | LINC01605 |
| ENSG0000 | -5.34712 | -40.7045 | 0.005337 | 0.081625 | None |
| ENSG0000 | -4.3014 | -19.7175 | 0.02055 | 0.200208 | None |
| ENSG0000 | -4.73705 | -26.6682 | 0.03556 | 0.280728 | None |
| ENSG0000 | -1.5829 | -2.99572 | 0.016711 | 0.173287 | LINC01932 |
| ENSG0000 | -1.61269 | -3.05821 | 0.022447 | 0.211564 | None |
| ENSG0000 | -1.8028 | -3.48896 | 0.053103 | 0.352688 | None |
| ENSG0000 | -4.94022 | -30.7012 | 0.007408 | 0.102351 | None |
| ENSG0000 | -3.61595 | -12.2606 | 0.069593 | 0.405995 | None |
| ENSG0000 | -1.3588 | -2.56471 | 0.096146 | 0.479685 | LINC02365 |
| ENSG0000 | -4.30389 | -19.7515 | 0.047346 | 0.33135 | None |
| ENSG0000 | -1.46322 | -2.75723 | 0.028317 | 0.24371 | None |
| ENSG0000 | -1.37522 | -2.59408 | 0.091919 | 0.468345 | LINC02547 |
| ENSG0000 | -1.55988 | -2.94829 | 0.026693 | 0.235347 | None |
| ENSG0000 | -1.25952 | -2.39417 | 0.055046 | 0.359232 | None |
| ENSG0000 | -2.29573 | -4.91002 | 0.10107 | 0.492753 | FAR1-IT1 |
| ENSG0000 | -1.67985 | -3.20395 | 0.050405 | 0.343479 | None |
| ENSG0000 | -3.26724 | -9.62803 | 0.03097 | 0.257997 | None |
| ENSG0000 | -1.03802 | -2.0534 | 0.01005 | 0.124846 | None |
| ENSG0000 | -1.51714 | -2.86222 | 0.101753 | 0.494637 | NOX5 |
| ENSG0000 | -2.11353 | -4.32748 | 0.097657 | 0.484088 | |
| ENSG0000 | -5.7077 | -52.2622 | 0.003056 | 0.054774 | None |
| ENSG0000 | -2.04701 | -4.13249 | 0.099207 | 0.488247 | None |
| ENSG0000 | -2.46958 | -5.53883 | 0.089406 | 0.461066 | None |
| ENSG0000 | -5.8994 | -59.6892 | 5.19E-05 | 0.002427 | None |
| ENSG0000 | -5.48493 | -44.7846 | 8.99E-05 | 0.003752 | None |
| ENSG0000 | -1.00176 | -2.00244 | 0.015721 | 0.167152 | LINC02454 |
| ENSG0000 | -5.28509 | -38.9916 | 0.001123 | 0.026818 | None |
| ENSG0000 | -3.7504 | -13.4581 | 0.035415 | 0.280214 | ARPC3P4 |
| ENSG0000 | -4.7337 | -26.6063 | 0.013348 | 0.148909 | IQSEC3P1 |
| ENSG0000 | -2.92867 | -7.61408 | 0.036579 | 0.28485 | None |
| ENSG0000 | -5.01208 | -32.269 | 0.000661 | 0.017731 | None |
| ENSG0000 | -4.25097 | -19.0401 | 0.013208 | 0.14795 | GPR142 |
| ENSG0000 | -5.4705 | -44.3387 | 0.003036 | 0.054514 | None |
| ENSG0000 | -2.48905 | -5.6141 | 0.075664 | 0.422599 | None |
| ENSG0000 | -1.47123 | -2.77258 | 0.035636 | 0.281086 | None |
| ENSG0000 | -6.01794 | -64.8009 | 6.47E-06 | 0.000414 | OR7E47P |
| ENSG0000 | -3.41767 | -10.6862 | 0.043211 | 0.315054 | None |
| ENSG0000 | -6.46287 | -88.2099 | 4.86E-05 | 0.002297 | None |
| ENSG0000 | -4.70767 | -26.1306 | 0.019493 | 0.193079 | None |

| | | | | | |
|---|---|---|---|---|---|
| ENSG0000 | -2.46575 | -5.52415 | 0.102379 | 0.49639 | R3HDM2-DT |
| ENSG0000 | -3.75958 | -13.544 | 0.054942 | 0.359057 | None |
| ENSG0000 | -1.63854 | -3.1135 | 0.083853 | 0.446748 | None |
| ENSG0000 | -5.59181 | -48.2284 | 0.001367 | 0.03123 | None |
| ENSG0000 | -2.19073 | -4.56538 | 0.006887 | 0.097263 | None |
| ENSG0000 | -4.68745 | -25.767 | 0.006439 | 0.092715 | None |
| ENSG0000 | -3.74888 | -13.4439 | 0.033885 | 0.273596 | None |
| ENSG0000 | -1.22048 | -2.33024 | 0.03577 | 0.2818 | None |
| ENSG0000 | -1.87553 | -3.66935 | 0.090875 | 0.464966 | CHMP4BP1 |
| ENSG0000 | -2.23911 | -4.72107 | 0.101937 | 0.495276 | LINC02207 |
| ENSG0000 | -5.28341 | -38.9463 | 0.003285 | 0.057453 | |
| ENSG0000 | -1.9284 | -3.80633 | 0.044713 | 0.321836 | None |
| ENSG0000 | -5.46161 | -44.0664 | 0.000554 | 0.015497 | LINC01579 |
| ENSG0000 | -5.58366 | -47.9567 | 0.000992 | 0.024353 | PSMB3P1 |
| ENSG0000 | -3.20347 | -9.21171 | 0.041124 | 0.306287 | ACTN1-DT |
| ENSG0000 | -2.76096 | -6.77849 | 0.050669 | 0.344179 | COILP1 |
| ENSG0000 | -2.62071 | -6.15053 | 0.046997 | 0.330296 | DDX18P1 |
| ENSG0000 | -1.80049 | -3.48338 | 0.00926 | 0.11854 | None |
| ENSG0000 | -3.69141 | -12.9189 | 0.060237 | 0.376514 | None |
| ENSG0000 | -4.93391 | -30.5671 | 0.013071 | 0.146945 | None |
| ENSG0000 | -2.48401 | -5.5945 | 0.073848 | 0.417555 | None |
| ENSG0000 | -3.94579 | -15.41 | 0.001618 | 0.035285 | LINC02895 |
| ENSG0000 | -1.49985 | -2.82814 | 0.057142 | 0.365845 | CTXND1 |
| ENSG0000 | -5.42789 | -43.0485 | 0.001176 | 0.027738 | UBE2Q2L |
| ENSG0000 | -1.19316 | -2.28654 | 0.087091 | 0.455099 | None |
| ENSG0000 | -6.4703 | -88.6656 | 4.04E-05 | 0.001952 | TYRO3P |
| ENSG0000 | -2.6541 | -6.29455 | 0.103319 | 0.499006 | PRELID1P4 |
| ENSG0000 | -3.69044 | -12.9102 | 0.047582 | 0.331999 | |
| ENSG0000 | -2.53392 | -5.79144 | 0.102582 | 0.496606 | None |
| ENSG0000 | -4.61638 | -24.5285 | 0.009525 | 0.12083 | None |
| ENSG0000 | -1.52216 | -2.87221 | 0.084733 | 0.449024 | ANP32BP1 |
| ENSG0000 | -1.55718 | -2.94278 | 0.097701 | 0.484088 | None |
| ENSG0000 | -1.18308 | -2.27061 | 0.090692 | 0.464489 | None |
| ENSG0000 | -4.26598 | -19.2392 | 0.055015 | 0.359134 | None |
| ENSG0000 | -4.74042 | -26.7306 | 0.02757 | 0.24023 | None |
| ENSG0000 | -1.85505 | -3.61764 | 0.062582 | 0.383515 | None |
| ENSG0000 | -5.30185 | -39.4472 | 0.003104 | 0.055347 | None |
| ENSG0000 | -3.02897 | -8.1623 | 0.053433 | 0.353809 | None |
| ENSG0000 | -2.80766 | -7.0015 | 0.096731 | 0.481153 | None |
| ENSG0000 | -1.40668 | -2.65127 | 0.09649 | 0.480632 | LINC02166 |
| ENSG0000 | -2.7302 | -6.63546 | 0.083005 | 0.444059 | ITFG1-AS1 |
| ENSG0000 | -4.98474 | -31.6633 | 0.010731 | 0.129737 | LINC01544 |
| ENSG0000 | -4.7238 | -26.4243 | 0.001641 | 0.035562 | None |
| ENSG0000 | -5.00644 | -32.1431 | 0.00294 | 0.053202 | None |
| ENSG0000 | -3.11023 | -8.6352 | 0.036034 | 0.282659 | None |
| ENSG0000 | -4.92877 | -30.4584 | 0.029755 | 0.251692 | None |
| ENSG0000 | -1.16619 | -2.24418 | 0.093646 | 0.473648 | GOLGA6FP |

| | | | | | |
|---|---|---|---|---|---|
| ENSG0000 | -1.45197 | -2.73581 | 0.009468 | 0.120272 | None |
| ENSG0000 | -4.31869 | -19.9551 | 0.008361 | 0.110726 | None |
| ENSG0000 | -1.61711 | -3.0676 | 0.032685 | 0.266787 | None |
| ENSG0000 | -2.71415 | -6.56209 | 0.081682 | 0.440172 | None |
| ENSG0000 | -3.65679 | -12.6126 | 0.056379 | 0.363201 | None |
| ENSG0000 | -2.93265 | -7.63513 | 0.052346 | 0.350532 | None |
| ENSG0000 | -4.29857 | -19.6788 | 0.016346 | 0.171453 | LINC02544 |
| ENSG0000 | -2.90379 | -7.48389 | 0.031842 | 0.263077 | None |
| ENSG0000 | -2.19222 | -4.57009 | 0.009879 | 0.123597 | EFCAB6-DT |
| ENSG0000 | -1.6212 | -3.0763 | 0.053578 | 0.354183 | None |
| ENSG0000 | -2.99946 | -7.99701 | 0.049465 | 0.339558 | None |
| ENSG0000 | -1.25649 | -2.38914 | 0.075546 | 0.422449 | POLG-DT |
| ENSG0000 | -3.72022 | -13.1795 | 0.039387 | 0.297942 | CCNYL7 |
| ENSG0000 | -2.78086 | -6.87263 | 0.073033 | 0.415203 | |
| ENSG0000 | -1.08476 | -2.12103 | 0.103571 | 0.499118 | None |
| ENSG0000 | -3.76138 | -13.5609 | 0.053311 | 0.35328 | HPR |
| ENSG0000 | -5.79895 | -55.6746 | 0.000453 | 0.013392 | GOSR2-DT |
| ENSG0000 | -1.0258 | -2.03609 | 0.027722 | 0.240823 | None |
| ENSG0000 | -5.26847 | -38.5449 | 0.004748 | 0.075104 | None |
| ENSG0000 | -1.80475 | -3.49369 | 0.022032 | 0.209127 | LINC02861 |
| ENSG0000 | -4.19332 | -18.2943 | 0.02078 | 0.201052 | None |
| ENSG0000 | -3.11231 | -8.64765 | 0.025889 | 0.231261 | MCUR1P1 |
| ENSG0000 | -4.63602 | -24.8645 | 0.011587 | 0.135924 | None |
| ENSG0000 | -4.97808 | -31.5175 | 0.013725 | 0.151816 | PCDHGA4 |
| ENSG0000 | -1.65993 | -3.16001 | 0.095561 | 0.478897 | MTCO1P40 |
| ENSG0000 | -5.30395 | -39.5046 | 0.000698 | 0.018654 | RPL23AP86 |
| ENSG0000 | -1.87247 | -3.66159 | 0.070836 | 0.40959 | KARS1P3 |
| ENSG0000 | -5.36222 | -41.133 | 0.002297 | 0.045026 | None |
| ENSG0000 | -1.04663 | -2.0657 | 0.024338 | 0.222596 | None |
| ENSG0000 | -4.92901 | -30.4636 | 0.023481 | 0.217509 | RN7SL155P |
| ENSG0000 | -4.68524 | -25.7276 | 0.007507 | 0.103114 | MIR3155A |
| ENSG0000 | -4.63157 | -24.7881 | 0.020642 | 0.200618 | None |
| ENSG0000 | -4.39279 | -21.0069 | 0.000394 | 0.011804 | DLGAP1-AS3 |
| ENSG0000 | -4.79536 | -27.7681 | 0.01675 | 0.173526 | RN7SL336P |
| ENSG0000 | -3.76048 | -13.5524 | 0.051801 | 0.348264 | None |
| ENSG0000 | -2.24882 | -4.75295 | 0.084599 | 0.448824 | ANXA8L1 |
| ENSG0000 | -1.49948 | -2.82742 | 0.008573 | 0.112421 | None |
| ENSG0000 | -4.36578 | -20.6172 | 0.026055 | 0.232078 | MIR548AC |
| ENSG0000 | -1.19351 | -2.28709 | 0.089032 | 0.460047 | MIR4653 |
| ENSG0000 | -6.07017 | -67.1897 | 7.22E-05 | 0.003147 | MIR3183 |
| ENSG0000 | -2.27998 | -4.85671 | 0.000633 | 0.017166 | NPY4R2 |
| ENSG0000 | -3.72669 | -13.2387 | 0.055713 | 0.361611 | MIR5094 |
| ENSG0000 | -1.57003 | -2.9691 | 0.011745 | 0.137097 | GJA5 |
| ENSG0000 | -1.24293 | -2.36678 | 0.00035 | 0.010842 | None |
| ENSG0000 | -3.27081 | -9.65186 | 0.008441 | 0.111336 | ARGFXP2 |
| ENSG0000 | -4.24589 | -18.9732 | 0.019062 | 0.190142 | None |
| ENSG0000 | -2.50553 | -5.67857 | 0.067707 | 0.400406 | None |

| | | | | | |
|---|---|---|---|---|---|
| ENSG0000 | -4.73037 | -26.5451 | 0.007699 | 0.104857 | None |
| ENSG0000 | -1.26504 | -2.40333 | 0.00666 | 0.094732 | SNHG25 |
| ENSG0000 | -1.77154 | -3.41419 | 0.03912 | 0.296962 | MIR3677 |
| ENSG0000 | -5.0057 | -32.1267 | 0.00552 | 0.083425 | None |
| ENSG0000 | -5.51806 | -45.8248 | 8.84E-05 | 0.0037 | None |
| ENSG0000 | -3.75746 | -13.5241 | 0.043641 | 0.317106 | None |
| ENSG0000 | -3.75243 | -13.4771 | 0.043291 | 0.315477 | None |
| ENSG0000 | -1.10756 | -2.1548 | 0.00017 | 0.00619 | PTGES3L |
| ENSG0000 | -4.36626 | -20.6241 | 0.030497 | 0.2555 | None |
| ENSG0000 | -4.72982 | -26.5349 | 0.006571 | 0.093943 | RNF157-AS1 |
| ENSG0000 | -1.03197 | -2.04482 | 0.020375 | 0.199158 | |
| ENSG0000 | -1.10049 | -2.14427 | 0.091499 | 0.466965 | LINC01775 |
| ENSG0000 | -2.76025 | -6.77511 | 0.051195 | 0.345828 | WHSC1L2P |
| ENSG0000 | -4.34707 | -20.3515 | 0.010437 | 0.127398 | None |
| ENSG0000 | -1.06576 | -2.09327 | 0.01697 | 0.175252 | KCNJ2-AS1 |
| ENSG0000 | -3.10936 | -8.63 | 0.070656 | 0.409452 | LINC02073 |
| ENSG0000 | -5.05267 | -33.19 | 0.033861 | 0.273596 | None |
| ENSG0000 | -4.77029 | -27.2899 | 0.025343 | 0.228273 | LINC01864 |
| ENSG0000 | -2.77681 | -6.85334 | 0.029313 | 0.249301 | None |
| ENSG0000 | -6.68943 | -103.209 | 1.06E-06 | 8.88E-05 | None |
| ENSG0000 | -2.39132 | -5.24637 | 0.010138 | 0.125519 | None |
| ENSG0000 | -1.28069 | -2.42955 | 0.038666 | 0.294876 | None |
| ENSG0000 | -4.65882 | -25.2607 | 0.012438 | 0.142228 | SNX6P1 |
| ENSG0000 | -3.68706 | -12.88 | 0.078684 | 0.430812 | None |
| ENSG0000 | -5.25316 | -38.1381 | 0.001359 | 0.03109 | BNIP3P27 |
| ENSG0000 | -5.64339 | -49.9839 | 0.000489 | 0.014181 | None |
| ENSG0000 | -1.25755 | -2.3909 | 0.031088 | 0.258363 | None |
| ENSG0000 | -3.53606 | -11.6 | 0.034574 | 0.276303 | HSFX2 |
| ENSG0000 | -4.70881 | -26.1512 | 0.004933 | 0.077218 | None |
| ENSG0000 | -2.42603 | -5.37414 | 0.089452 | 0.461136 | ESPNP |
| ENSG0000 | -5.00488 | -32.1084 | 0.004328 | 0.070536 | None |
| ENSG0000 | -4.26845 | -19.2722 | 0.035797 | 0.281812 | SLC6A21P |
| ENSG0000 | -2.52522 | -5.75661 | 0.070287 | 0.408142 | None |
| ENSG0000 | -5.90696 | -60.003 | 1.78E-05 | 0.000976 | None |
| ENSG0000 | -3.14934 | -8.87248 | 0.021038 | 0.202847 | None |
| ENSG0000 | -4.73146 | -26.5651 | 0.008813 | 0.114599 | None |
| ENSG0000 | -1.31585 | -2.48949 | 0.066419 | 0.396045 | None |
| ENSG0000 | -1.51314 | -2.8543 | 0.09229 | 0.469554 | |
| ENSG0000 | -4.16222 | -17.9042 | 0.059733 | 0.374653 | None |
| ENSG0000 | -1.09514 | -2.13634 | 0.027344 | 0.239303 | None |
| ENSG0000 | -1.49424 | -2.81716 | 0.019858 | 0.195437 | |
| ENSG0000 | -4.33011 | -20.1137 | 0.026037 | 0.232078 | None |
| ENSG0000 | -3.64926 | -12.5469 | 0.075315 | 0.421944 | None |
| ENSG0000 | -3.6286 | -12.3685 | 0.025548 | 0.229665 | None |
| ENSG0000 | -5.57351 | -47.6205 | 0.005322 | 0.081526 | None |
| ENSG0000 | -1.20043 | -2.29808 | 0.013971 | 0.1538 | None |
| ENSG0000 | -4.65879 | -25.2602 | 0.017241 | 0.17747 | None |

| | | | | | |
|---|---|---|---|---|---|
| ENSG0000 | -4.67749 | -25.5897 | 0.043921 | 0.318458 | MAGOH3P |
| ENSG0000 | -4.33835 | -20.229 | 0.035814 | 0.281862 | None |
| ENSG0000 | -1.50253 | -2.83339 | 0.039111 | 0.296962 | BBIP1P1 |
| ENSG0000 | -4.67773 | -25.5939 | 0.041575 | 0.307573 | None |
| ENSG0000 | -2.97064 | -7.83886 | 0.062676 | 0.383859 | None |
| ENSG0000 | -5.0723 | -33.6445 | 0.00451 | 0.072364 | SYNPO2L-AS1 |
| ENSG0000 | -1.96552 | -3.90553 | 0.103404 | 0.499006 | None |
| ENSG0000 | -1.16432 | -2.24127 | 0.058671 | 0.371248 | None |
| ENSG0000 | -5.25072 | -38.0735 | 0.002354 | 0.045793 | |
| ENSG0000 | -2.95257 | -7.74127 | 0.015444 | 0.165122 | ANKH-DT |
| ENSG0000 | -5.24518 | -37.9278 | 0.005151 | 0.079691 | None |
| ENSG0000 | -1.9199 | -3.78398 | 0.082011 | 0.441294 | None |
| ENSG0000 | -1.14542 | -2.21211 | 0.091812 | 0.468026 | IER3-AS1 |
| ENSG0000 | -1.37772 | -2.59857 | 0.053959 | 0.355307 | None |
| ENSG0000 | -3.14622 | -8.85335 | 0.057273 | 0.366436 | None |
| ENSG0000 | -5.36353 | -41.1703 | 0.005687 | 0.085395 | None |
| ENSG0000 | -5.03103 | -32.6956 | 0.010151 | 0.125519 | None |
| ENSG0000 | -2.18576 | -4.54967 | 0.072471 | 0.414086 | RNU6-88P |
| ENSG0000 | -5.06981 | -33.5865 | 0.003098 | 0.055285 | None |
| ENSG0000 | -2.85802 | -7.25021 | 0.023275 | 0.216315 | None |
| ENSG0000 | -1.54196 | -2.91191 | 0.008725 | 0.113776 | None |
| ENSG0000 | -2.1909 | -4.56591 | 0.072634 | 0.414361 | CNNM3-DT |
| ENSG0000 | -1.51821 | -2.86436 | 0.054884 | 0.358761 | |
| ENSG0000 | -1.20735 | -2.30913 | 0.002188 | 0.043654 | LINC02725 |
| ENSG0000 | -4.26864 | -19.2748 | 0.034563 | 0.276303 | None |
| ENSG0000 | -3.62629 | -12.3487 | 0.055086 | 0.359329 | None |
| ENSG0000 | -1.74776 | -3.35837 | 0.022633 | 0.212849 | None |
| ENSG0000 | -3.07536 | -8.42897 | 0.067079 | 0.398283 | None |
| ENSG0000 | -3.53147 | -11.5632 | 0.037751 | 0.290716 | None |
| ENSG0000 | -5.77445 | -54.7373 | 0.008283 | 0.109896 | SPDYE10 |
| ENSG0000 | -1.02467 | -2.0345 | 0.056186 | 0.363029 | None |
| ENSG0000 | -2.83504 | -7.13562 | 0.076556 | 0.424948 | None |
| ENSG0000 | -2.91798 | -7.55785 | 0.077191 | 0.426614 | GXYLT1P4 |
| ENSG0000 | -1.33799 | -2.52799 | 0.099138 | 0.488247 | None |
| ENSG0000 | -1.185 | -2.27364 | 0.008647 | 0.11299 | None |
| ENSG0000 | -2.76243 | -6.7854 | 0.057993 | 0.368519 | None |
| ENSG0000 | -3.72475 | -13.2209 | 0.047678 | 0.332422 | |
| ENSG0000 | -2.25975 | -4.78908 | 0.097801 | 0.484239 | None |
| ENSG0000 | -3.0438 | -8.24658 | 0.044829 | 0.322163 | HNRNPCL2 |
| ENSG0000 | -5.50354 | -45.366 | 0.001674 | 0.036107 | OR7G15P |
| ENSG0000 | -2.87143 | -7.31791 | 0.081326 | 0.439266 | None |
| ENSG0000 | -4.26625 | -19.2428 | 0.052475 | 0.350911 | None |
| ENSG0000 | -4.78387 | -27.5478 | 0.008326 | 0.110365 | None |
| ENSG0000 | -5.06786 | -33.5412 | 0.00753 | 0.103278 | None |
| ENSG0000 | -2.13661 | -4.39726 | 0.013726 | 0.151816 | None |
| ENSG0000 | -5.04656 | -33.0496 | 0.002519 | 0.047837 | None |
| ENSG0000 | -2.57638 | -5.96441 | 0.056986 | 0.365179 | |

| | | | | | |
|---|---|---|---|---|---|
| ENSG0000 | -2.40583 | -5.29941 | 0.06207 | 0.381799 | HYDIN2 |
| ENSG0000 | -3.04701 | -8.26497 | 0.036385 | 0.284161 | H4C4 |
| ENSG0000 | -1.92908 | -3.80813 | 0.057166 | 0.365853 | None |
| ENSG0000 | -2.06101 | -4.17279 | 0.077195 | 0.426614 | None |
| ENSG0000 | -3.72111 | -13.1876 | 0.042608 | 0.312204 | None |
| ENSG0000 | -4.69268 | -25.8605 | 0.001582 | 0.034771 | None |
| ENSG0000 | -4.68485 | -25.7205 | 0.018086 | 0.183197 | None |
| ENSG0000 | -2.2881 | -4.88414 | 0.095169 | 0.4777 | None |
| ENSG0000 | -1.68035 | -3.20505 | 0.008892 | 0.115257 | None |
| ENSG0000 | -3.66637 | -12.6966 | 0.039026 | 0.296593 | None |
| ENSG0000 | -4.73504 | -26.6311 | 0.001271 | 0.029329 | None |
| ENSG0000 | -2.75864 | -6.76757 | 0.048214 | 0.334738 | None |
| ENSG0000 | -3.62718 | -12.3564 | 0.053954 | 0.355307 | H4C1 |
| ENSG0000 | -4.36575 | -20.6168 | 0.034153 | 0.274418 | |
| ENSG0000 | -1.60819 | -3.04869 | 0.027747 | 0.24088 | H2AC17 |
| ENSG0000 | -1.64914 | -3.13647 | 0.057772 | 0.367699 | None |
| ENSG0000 | -4.68497 | -25.7227 | 0.008045 | 0.107759 | None |
| ENSG0000 | -1.99703 | -3.99178 | 0.086901 | 0.454835 | None |
| ENSG0000 | -1.21152 | -2.31582 | 0.079741 | 0.433964 | None |
| ENSG0000 | -2.54136 | -5.82139 | 0.065346 | 0.392891 | |
| ENSG0000 | -5.24366 | -37.8877 | 0.000472 | 0.013809 | None |
| ENSG0000 | -1.43835 | -2.71011 | 0.073351 | 0.416145 | None |
| ENSG0000 | -3.02553 | -8.14281 | 0.045726 | 0.324785 | None |
| ENSG0000 | -5.35186 | -40.8386 | 0.015725 | 0.167152 | None |
| ENSG0000 | -1.51774 | -2.86343 | 0.02743 | 0.239832 | None |
| ENSG0000 | -3.2286 | -9.37355 | 0.050591 | 0.344007 | None |
| ENSG0000 | -3.68857 | -12.8935 | 0.066598 | 0.396777 | None |
| ENSG0000 | -5.28844 | -39.0821 | 0.000343 | 0.010647 | None |
| ENSG0000 | -5.34267 | -40.5792 | 0.00609 | 0.089249 | None |
| ENSG0000 | -2.94982 | -7.72654 | 0.031174 | 0.2587 | None |
| ENSG0000 | -4.75924 | -27.0815 | 0.007858 | 0.106273 | None |
| ENSG0000 | -4.30376 | -19.7497 | 0.045576 | 0.324253 | None |
| ENSG0000 | -5.32201 | -40.0022 | 0.002201 | 0.043801 | None |
| ENSG0000 | -5.48322 | -44.7316 | 0.007062 | 0.098845 | None |
| ENSG0000 | -4.26609 | -19.2407 | 0.053949 | 0.355307 | None |
| ENSG0000 | -5.68115 | -51.3094 | 0.001734 | 0.037163 | None |
| ENSG0000 | -2.68569 | -6.43387 | 0.047348 | 0.33135 | None |
| ENSG0000 | -5.02657 | -32.5947 | 0.001959 | 0.040492 | |
| ENSG0000 | -4.87909 | -29.4274 | 0.025134 | 0.227125 | None |
| ENSG0000 | -3.23404 | -9.40897 | 0.029966 | 0.252637 | ERC2-IT1 |
| ENSG0000 | -4.31946 | -19.9658 | 0.010113 | 0.125459 | METTL14-DT |
| ENSG0000 | -1.60596 | -3.04399 | 0.034412 | 0.27586 | None |
| ENSG0000 | -3.10077 | -8.57876 | 0.049989 | 0.341759 | None |
| ENSG0000 | -3.75152 | -13.4685 | 0.001514 | 0.033761 | None |
| ENSG0000 | -5.23641 | -37.698 | 0.001527 | 0.033893 | U4 |
| ENSG0000 | -5.07655 | -33.7438 | 0.000883 | 0.022356 | None |
| ENSG0000 | -5.72146 | -52.7633 | 0.000462 | 0.01358 | LINC01902 |

| | | | | | |
|---|---|---|---|---|---|
| ENSG0000 | -5.77925 | -54.9195 | 0.000283 | 0.009169 | None |
| ENSG0000 | -2.24406 | -4.73727 | 0.086648 | 0.454425 | Y_RNA |

| GeneID | Treatment | Treatment | Treatment | Treatment | name |
|---|---|---|---|---|---|
| ENSG0000 | 1.125644 | 2.181989 | 0.004058 | 0.034537 | MYH16 |
| ENSG0000 | 1.13678 | 2.198896 | 1.48E-14 | 1.95E-12 | TMEM132A |
| ENSG0000 | 2.489453 | 5.61565 | 3.11E-11 | 2.45E-09 | CACNA1G |
| ENSG0000 | 1.636226 | 3.108516 | 6.11E-10 | 3.88E-08 | SYN1 |
| ENSG0000 | 1.747656 | 3.358125 | 1.53E-05 | 0.000349 | SLC6A13 |
| ENSG0000 | 1.014215 | 2.019803 | 7.04E-17 | 1.23E-14 | SLC38A5 |
| ENSG0000 | 1.387331 | 2.615943 | 4.55E-09 | 2.46E-07 | ATP1A2 |
| ENSG0000 | 1.117641 | 2.169919 | 7.82E-21 | 2.22E-18 | EHD2 |
| ENSG0000 | 6.281693 | 77.7997 | 6.72E-05 | 0.001251 | INSRR |
| ENSG0000 | 1.245699 | 2.371334 | 3.41E-23 | 1.20E-20 | ANK1 |
| ENSG0000 | 1.661367 | 3.16316 | 6.29E-16 | 1.00E-13 | TNC |
| ENSG0000 | 4.542028 | 23.29628 | 0.003146 | 0.028565 | DKK3 |
| ENSG0000 | 1.327379 | 2.509464 | 5.26E-32 | 4.70E-29 | PTPRN |
| ENSG0000 | 1.86293 | 3.637458 | 0.005903 | 0.046009 | KCNH2 |
| ENSG0000 | 3.759623 | 13.54439 | 0.004108 | 0.034855 | NGFR |
| ENSG0000 | 1.228531 | 2.343283 | 1.22E-07 | 4.79E-06 | HIPK2 |
| ENSG0000 | 1.14839 | 2.216663 | 2.79E-05 | 0.000582 | SLC9A3R2 |
| ENSG0000 | 1.607046 | 3.046274 | 0.005359 | 0.042853 | GLP2R |
| ENSG0000 | 1.372382 | 2.588976 | 1.08E-11 | 9.25E-10 | EML1 |
| ENSG0000 | 1.146443 | 2.213674 | 8.66E-11 | 6.27E-09 | NAV3 |
| ENSG0000 | 6.480759 | 89.31057 | 0.000245 | 0.003663 | SCT |
| ENSG0000 | 5.250826 | 38.07643 | 0.001198 | 0.013441 | FNDC8 |
| ENSG0000 | 1.558149 | 2.944757 | 4.94E-17 | 8.78E-15 | FRY |
| ENSG0000 | 1.231685 | 2.348411 | 1.66E-18 | 3.46E-16 | NOTCH3 |
| ENSG0000 | 1.121298 | 2.175426 | 0.005243 | 0.042127 | SNCB |
| ENSG0000 | 1.459735 | 2.750578 | 7.15E-31 | 6.02E-28 | FOSL2 |
| ENSG0000 | 2.127986 | 4.37107 | 0.000959 | 0.011174 | FAP |
| ENSG0000 | 1.000805 | 2.001117 | 3.84E-23 | 1.34E-20 | SYNJ2 |
| ENSG0000 | 1.24554 | 2.371073 | 6.80E-21 | 1.94E-18 | COL5A3 |
| ENSG0000 | 2.240386 | 4.725233 | 0.000356 | 0.004969 | IGSF9B |
| ENSG0000 | 1.277448 | 2.424098 | 8.92E-05 | 0.00159 | CXCL2 |
| ENSG0000 | 1.683623 | 3.212336 | 0.005492 | 0.043711 | MECOM |
| ENSG0000 | 1.001676 | 2.002325 | 2.17E-13 | 2.42E-11 | TFAP2C |
| ENSG0000 | 1.175082 | 2.258058 | 1.10E-25 | 5.00E-23 | P3H2 |
| ENSG0000 | 1.093827 | 2.134395 | 9.58E-14 | 1.12E-11 | PITPNM2 |
| ENSG0000 | 4.704794 | 26.07859 | 7.14E-05 | 0.001317 | IL11 |
| ENSG0000 | 1.239511 | 2.361186 | 0.000489 | 0.006455 | SERPIND1 |
| ENSG0000 | 1.156799 | 2.229622 | 3.40E-09 | 1.88E-07 | CARD10 |
| ENSG0000 | 1.316834 | 2.491188 | 5.49E-13 | 5.78E-11 | CPNE6 |
| ENSG0000 | 1.108588 | 2.156345 | 0.002303 | 0.022363 | SLA2 |
| ENSG0000 | 1.492486 | 2.813734 | 1.82E-21 | 5.42E-19 | NTSR1 |
| ENSG0000 | 1.095749 | 2.13724 | 2.14E-05 | 0.000462 | EEF1A2 |
| ENSG0000 | 1.758083 | 3.382483 | 0.004851 | 0.039825 | WFDC2 |
| ENSG0000 | 5.581848 | 47.89648 | 0.002168 | 0.021315 | RS1 |
| ENSG0000 | 1.139764 | 2.20345 | 7.15E-08 | 2.96E-06 | GABRE |
| ENSG0000 | 1.088629 | 2.126718 | 7.48E-10 | 4.66E-08 | SRPX2 |

| | | | | | |
|---|---|---|---|---|---|
| ENSG0000 | 1.116063 | 2.167546 | 2.37E-05 | 0.000503 | FOXF1 |
| ENSG0000 | 3.196012 | 9.164221 | 2.63E-14 | 3.35E-12 | CORO2B |
| ENSG0000 | 2.198019 | 4.588488 | 2.49E-08 | 1.15E-06 | CCN4 |
| ENSG0000 | 3.844497 | 14.36511 | 0.005536 | 0.043997 | CGB3 |
| ENSG0000 | 2.368917 | 5.165532 | 5.92E-10 | 3.77E-08 | TFPI2 |
| ENSG0000 | 1.007348 | 2.010212 | 0.00511 | 0.041392 | CAV2 |
| ENSG0000 | 1.524544 | 2.876957 | 5.21E-25 | 2.16E-22 | CAV1 |
| ENSG0000 | 1.054567 | 2.077095 | 9.41E-11 | 6.75E-09 | MET |
| ENSG0000 | 1.033596 | 2.04712 | 3.94E-22 | 1.25E-19 | STX1A |
| ENSG0000 | 1.58888 | 3.008157 | 0.000814 | 0.0098 | CHCHD2 |
| ENSG0000 | 1.039853 | 2.056018 | 2.92E-06 | 8.28E-05 | CCL24 |
| ENSG0000 | 1.088601 | 2.126678 | 2.79E-35 | 3.47E-32 | SERPINE1 |
| ENSG0000 | 1.657169 | 3.153969 | 3.91E-05 | 0.000778 | AEBP1 |
| ENSG0000 | 2.354822 | 5.115311 | 0.003946 | 0.033816 | KCNT1 |
| ENSG0000 | 1.603478 | 3.03875 | 2.47E-13 | 2.73E-11 | GLIS3 |
| ENSG0000 | 1.584881 | 2.99983 | 2.97E-08 | 1.35E-06 | WNT3 |
| ENSG0000 | 1.74474 | 3.351345 | 3.66E-25 | 1.54E-22 | CCL7 |
| ENSG0000 | 1.473915 | 2.777746 | 0.000166 | 0.002667 | EFNB3 |
| ENSG0000 | 2.819125 | 7.057342 | 1.43E-18 | 3.00E-16 | AREG |
| ENSG0000 | 6.269491 | 77.14446 | 0.000425 | 0.005761 | NKX3-2 |
| ENSG0000 | 2.087466 | 4.250009 | 0.006273 | 0.048129 | CRYAB |
| ENSG0000 | 5.589178 | 48.14045 | 0.000194 | 0.003011 | B3GAT1 |
| ENSG0000 | 1.704585 | 3.259351 | 2.62E-18 | 5.40E-16 | SLC1A2 |
| ENSG0000 | 1.578221 | 2.986015 | 2.82E-08 | 1.29E-06 | WNT5B |
| ENSG0000 | 1.250135 | 2.378636 | 2.82E-12 | 2.65E-10 | PDGFRB |
| ENSG0000 | 1.248063 | 2.375224 | 0.003709 | 0.032323 | AMOTL2 |
| ENSG0000 | 1.421708 | 2.679026 | 1.81E-13 | 2.05E-11 | COL7A1 |
| ENSG0000 | 1.50219 | 2.832724 | 0.002726 | 0.025542 | C3orf52 |
| ENSG0000 | 2.106715 | 4.307095 | 0.002697 | 0.025331 | PRRX1 |
| ENSG0000 | 1.318333 | 2.493778 | 0.003249 | 0.029309 | ADGRL2 |
| ENSG0000 | 1.581375 | 2.99255 | 7.65E-11 | 5.63E-09 | MMP8 |
| ENSG0000 | 3.613395 | 12.23884 | 0.00031 | 0.004428 | GRIA2 |
| ENSG0000 | 2.4295 | 5.387067 | 0.004273 | 0.035994 | ADGRB2 |
| ENSG0000 | 1.365105 | 2.57595 | 0.003312 | 0.029737 | TMEM54 |
| ENSG0000 | 1.168753 | 2.248174 | 4.56E-16 | 7.42E-14 | CXCR4 |
| ENSG0000 | 1.408428 | 2.654477 | 1.49E-27 | 8.71E-25 | INHBA |
| ENSG0000 | 3.109016 | 8.627939 | 4.71E-18 | 9.41E-16 | TWIST1 |
| ENSG0000 | 1.877795 | 3.67513 | 3.04E-11 | 2.41E-09 | CHST3 |
| ENSG0000 | 1.655258 | 3.149795 | 0.00015 | 0.002449 | PLP1 |
| ENSG0000 | 5.205392 | 36.89598 | 0.001061 | 0.012134 | PI3 |
| ENSG0000 | 1.947082 | 3.855939 | 0.003114 | 0.028293 | SLPI |
| ENSG0000 | 1.043443 | 2.061141 | 1.03E-24 | 4.20E-22 | PMEPA1 |
| ENSG0000 | 1.347552 | 2.544799 | 0.001193 | 0.013393 | F13A1 |
| ENSG0000 | 2.782249 | 6.879238 | 4.92E-07 | 1.71E-05 | EREG |
| ENSG0000 | 1.617583 | 3.068606 | 2.30E-24 | 8.89E-22 | MYRF |
| ENSG0000 | 1.507955 | 2.844065 | 1.93E-23 | 6.99E-21 | WNT1 |
| ENSG0000 | 1.050189 | 2.070801 | 7.26E-10 | 4.54E-08 | EFNB2 |

| | | | | | |
|---|---|---|---|---|---|
| ENSG0000 | 1.322592 | 2.501151 | 6.42E-06 | 0.000164 | SOX9 |
| ENSG0000 | 1.52143 | 2.870755 | 8.30E-30 | 6.24E-27 | HS3ST3B1 |
| ENSG0000 | 1.179816 | 2.265479 | 3.44E-40 | 6.56E-37 | IL1B |
| ENSG0000 | 1.864003 | 3.640163 | 9.08E-18 | 1.78E-15 | FOSB |
| ENSG0000 | 6.008752 | 64.38945 | 0.000486 | 0.006426 | FOXA2 |
| ENSG0000 | 1.059959 | 2.084873 | 8.39E-06 | 0.000208 | CCR7 |
| ENSG0000 | 1.544303 | 2.91663 | 0.001409 | 0.01523 | FLRT1 |
| ENSG0000 | 1.106831 | 2.15372 | 1.90E-15 | 2.84E-13 | A4GALT |
| ENSG0000 | 1.056132 | 2.079349 | 7.52E-14 | 8.96E-12 | PODXL |
| ENSG0000 | 1.177866 | 2.262419 | 1.87E-06 | 5.56E-05 | FLNC |
| ENSG0000 | 2.936523 | 7.65564 | 1.66E-05 | 0.000373 | ISLR |
| ENSG0000 | 1.024868 | 2.034773 | 7.86E-08 | 3.22E-06 | COL5A1 |
| ENSG0000 | 1.317255 | 2.491915 | 1.35E-05 | 0.000312 | LAMA5 |
| ENSG0000 | 1.524974 | 2.877815 | 0.000245 | 0.003661 | ULBP3 |
| ENSG0000 | 2.399698 | 5.276926 | 1.44E-54 | 5.14E-51 | GFPT2 |
| ENSG0000 | 1.916908 | 3.77613 | 7.15E-33 | 6.82E-30 | NT5E |
| ENSG0000 | 5.70708 | 52.2399 | 0.000581 | 0.00743 | RNASE2CP |
| ENSG0000 | 1.521085 | 2.870067 | 5.53E-09 | 2.93E-07 | SCN7A |
| ENSG0000 | 5.455836 | 43.89047 | 0.001315 | 0.014421 | SLC28A2 |
| ENSG0000 | 1.477191 | 2.784062 | 6.02E-30 | 4.65E-27 | STRA6 |
| ENSG0000 | 1.527016 | 2.881891 | 0.002883 | 0.026591 | ARHGAP29 |
| ENSG0000 | 1.263313 | 2.400464 | 0.001317 | 0.014436 | FGF2 |
| ENSG0000 | 1.141294 | 2.205787 | 2.46E-52 | 7.04E-49 | SLC39A8 |
| ENSG0000 | 1.24043 | 2.36269 | 4.45E-07 | 1.56E-05 | RBP5 |
| ENSG0000 | 1.285296 | 2.43732 | 7.63E-15 | 1.03E-12 | JDP2 |
| ENSG0000 | 1.635283 | 3.106486 | 3.06E-10 | 2.02E-08 | ABCC12 |
| ENSG0000 | 3.583141 | 11.98486 | 0.005571 | 0.044231 | GATA6 |
| ENSG0000 | 1.843962 | 3.589945 | 0.003641 | 0.031885 | MYOM3 |
| ENSG0000 | 1.261414 | 2.397306 | 6.76E-10 | 4.26E-08 | HSPG2 |
| ENSG0000 | 2.431942 | 5.396192 | 1.13E-09 | 6.81E-08 | TINAGL1 |
| ENSG0000 | 1.706584 | 3.263871 | 4.24E-10 | 2.75E-08 | KCNN3 |
| ENSG0000 | 3.875645 | 14.67863 | 0.000111 | 0.001909 | ACKR3 |
| ENSG0000 | 1.805599 | 3.495742 | 2.69E-13 | 2.95E-11 | TAGLN3 |
| ENSG0000 | 1.753052 | 3.370709 | 4.82E-10 | 3.11E-08 | UCN2 |
| ENSG0000 | 1.52775 | 2.883358 | 7.89E-139 | 2.26E-134 | RNF145 |
| ENSG0000 | 1.283066 | 2.433556 | 0.000434 | 0.005863 | IGFBP3 |
| ENSG0000 | 1.342662 | 2.536188 | 2.26E-06 | 6.56E-05 | SLC16A2 |
| ENSG0000 | 1.321983 | 2.500096 | 0.000701 | 0.008634 | NFIB |
| ENSG0000 | 1.177011 | 2.261079 | 4.55E-05 | 0.000885 | CRB2 |
| ENSG0000 | 5.465674 | 44.19078 | 0.001434 | 0.01546 | FAM13C |
| ENSG0000 | 2.184229 | 4.54484 | 0.0006 | 0.00762 | ANKRD1 |
| ENSG0000 | 1.29736 | 2.457787 | 0.003769 | 0.03276 | C20orf144 |
| ENSG0000 | 1.613594 | 3.060133 | 1.40E-55 | 5.74E-52 | HMGA2 |
| ENSG0000 | 3.193403 | 9.147664 | 5.41E-05 | 0.001028 | MMP3 |
| ENSG0000 | 1.509967 | 2.848036 | 0.002991 | 0.027424 | ADGRL3 |
| ENSG0000 | 1.102682 | 2.147535 | 6.41E-11 | 4.75E-09 | SLC7A11 |
| ENSG0000 | 3.863708 | 14.55767 | 0.003062 | 0.027915 | ENKUR |

| | | | | | |
|---|---|---|---|---|---|
| ENSG0000 | 1.460317 | 2.751689 | 2.14E-06 | 6.28E-05 | PTPN14 |
| ENSG0000 | 1.52136 | 2.870616 | 3.81E-09 | 2.09E-07 | BMP6 |
| ENSG0000 | 1.148759 | 2.217231 | 1.14E-53 | 3.61E-50 | LPCAT1 |
| ENSG0000 | 1.501818 | 2.831994 | 1.13E-05 | 0.000266 | PTPRD |
| ENSG0000 | 1.461505 | 2.753955 | 0.001767 | 0.018134 | MMP21 |
| ENSG0000 | 1.01485 | 2.020693 | 0.000959 | 0.011169 | JAM2 |
| ENSG0000 | 1.245577 | 2.371134 | 3.43E-12 | 3.18E-10 | PPM1J |
| ENSG0000 | 1.05033 | 2.071003 | 1.76E-06 | 5.30E-05 | PXYLP1 |
| ENSG0000 | 5.749027 | 53.78109 | 0.000107 | 0.00185 | KCNS2 |
| ENSG0000 | 1.922562 | 3.790957 | 0.000212 | 0.003249 | NMNAT2 |
| ENSG0000 | 2.060675 | 4.171814 | 1.64E-12 | 1.59E-10 | KCNJ15 |
| ENSG0000 | 2.288162 | 4.884335 | 0.001076 | 0.012285 | SLC34A2 |
| ENSG0000 | 1.34759 | 2.544866 | 0.00569 | 0.044823 | CLSTN2 |
| ENSG0000 | 1.014224 | 2.019816 | 0.000164 | 0.002635 | SHROOM4 |
| ENSG0000 | 2.883612 | 7.379957 | 0.001572 | 0.016563 | NRG2 |
| ENSG0000 | 1.174623 | 2.257338 | 1.21E-15 | 1.87E-13 | B4GALT5 |
| ENSG0000 | 1.324266 | 2.504054 | 1.78E-06 | 5.35E-05 | CACHD1 |
| ENSG0000 | 1.4813 | 2.792002 | 7.42E-05 | 0.001364 | LAD1 |
| ENSG0000 | 5.029812 | 32.66812 | 9.25E-11 | 6.64E-09 | STC1 |
| ENSG0000 | 1.125528 | 2.181814 | 0.002231 | 0.021822 | FTCD |
| ENSG0000 | 1.112875 | 2.162762 | 5.51E-37 | 7.50E-34 | TAL1 |
| ENSG0000 | 1.150983 | 2.220651 | 2.75E-06 | 7.84E-05 | PDPN |
| ENSG0000 | 1.43416 | 2.702248 | 6.32E-10 | 3.99E-08 | RFTN2 |
| ENSG0000 | 5.83671 | 57.15111 | 4.33E-05 | 0.000848 | LRRTM1 |
| ENSG0000 | 1.217155 | 2.324878 | 0.001104 | 0.012575 | IGFBP7 |
| ENSG0000 | 3.954718 | 15.50561 | 0.004956 | 0.040411 | EOMES |
| ENSG0000 | 2.615725 | 6.129312 | 0.003772 | 0.032776 | ADAMTS9 |
| ENSG0000 | 2.309575 | 4.95737 | 5.61E-31 | 4.86E-28 | CXCL3 |
| ENSG0000 | 3.158236 | 8.927375 | 4.63E-88 | 6.62E-84 | CXCL5 |
| ENSG0000 | 1.220625 | 2.330477 | 5.67E-07 | 1.93E-05 | PLXNB1 |
| ENSG0000 | 1.418225 | 2.672565 | 8.04E-11 | 5.87E-09 | ITGA2 |
| ENSG0000 | 1.716795 | 3.287054 | 0.00023 | 0.003486 | EDIL3 |
| ENSG0000 | 5.174062 | 36.10337 | 0.003031 | 0.027677 | DACT2 |
| ENSG0000 | 1.422571 | 2.680628 | 7.82E-14 | 9.29E-12 | IL31RA |
| ENSG0000 | 5.80671 | 55.97499 | 0.000859 | 0.01025 | SMAD5-AS1 |
| ENSG0000 | 1.524429 | 2.876728 | 1.82E-20 | 4.81E-18 | STEAP1 |
| ENSG0000 | 1.470889 | 2.771927 | 8.10E-13 | 8.25E-11 | AQP3 |
| ENSG0000 | 1.839184 | 3.578077 | 1.24E-17 | 2.37E-15 | DACT1 |
| ENSG0000 | 1.145847 | 2.21276 | 3.78E-06 | 0.000103 | PACSIN3 |
| ENSG0000 | 1.204505 | 2.304581 | 3.81E-27 | 2.06E-24 | HTRA1 |
| ENSG0000 | 1.452223 | 2.736294 | 2.59E-09 | 1.46E-07 | RRAD |
| ENSG0000 | 1.155753 | 2.228006 | 3.91E-37 | 5.89E-34 | ANPEP |
| ENSG0000 | 1.219964 | 2.329409 | 1.77E-08 | 8.56E-07 | NAV2 |
| ENSG0000 | 1.283784 | 2.434767 | 0.001829 | 0.018636 | IGF2 |
| ENSG0000 | 1.604048 | 3.039951 | 2.91E-05 | 0.000604 | GNG8 |
| ENSG0000 | 1.837639 | 3.574246 | 0.001889 | 0.019116 | AQP2 |
| ENSG0000 | 1.379518 | 2.601814 | 0.001537 | 0.016257 | CDC42EP5 |

| | | | | | |
|---|---|---|---|---|---|
| ENSG0000 | 1.345721 | 2.541571 | 0.00151 | 0.016039 | TMEM88 |
| ENSG0000 | 1.066926 | 2.094965 | 0.000238 | 0.003572 | PNOC |
| ENSG0000 | 1.167882 | 2.246816 | 0.004512 | 0.037595 | AXIN2 |
| ENSG0000 | 1.032939 | 2.046189 | 0.00012 | 0.002043 | ATOH8 |
| ENSG0000 | 1.080218 | 2.114355 | 0.002195 | 0.021527 | IRS1 |
| ENSG0000 | 1.41428 | 2.665267 | 0.00191 | 0.019288 | RAB3B |
| ENSG0000 | 1.399699 | 2.638465 | 3.41E-14 | 4.26E-12 | NPR1 |
| ENSG0000 | 1.063687 | 2.090266 | 0.00165 | 0.017251 | CHRNA5 |
| ENSG0000 | 1.012826 | 2.01786 | 3.27E-18 | 6.69E-16 | WNT10B |
| ENSG0000 | 2.257783 | 4.782559 | 2.19E-33 | 2.32E-30 | TM4SF1 |
| ENSG0000 | 1.382868 | 2.607862 | 1.57E-15 | 2.37E-13 | NLGN2 |
| ENSG0000 | 4.043552 | 16.49037 | 0.001633 | 0.017093 | HSD17B13 |
| ENSG0000 | 5.488134 | 44.88414 | 0.000942 | 0.011021 | SYT9 |
| ENSG0000 | 1.580206 | 2.990126 | 2.96E-07 | 1.08E-05 | GPR37 |
| ENSG0000 | 1.140745 | 2.204948 | 1.28E-20 | 3.52E-18 | HTRA3 |
| ENSG0000 | 1.120079 | 2.173589 | 1.72E-19 | 4.11E-17 | MTSS1 |
| ENSG0000 | 1.571816 | 2.972786 | 0.000735 | 0.008977 | LRRC8E |
| ENSG0000 | 1.090731 | 2.129819 | 2.21E-09 | 1.26E-07 | HOPX |
| ENSG0000 | 1.208683 | 2.311265 | 3.53E-06 | 9.68E-05 | LPAR3 |
| ENSG0000 | 1.488726 | 2.80641 | 3.24E-05 | 0.000662 | SYNPO |
| ENSG0000 | 2.421513 | 5.357325 | 2.24E-22 | 7.44E-20 | ID4 |
| ENSG0000 | 4.66488 | 25.36699 | 0.003501 | 0.030975 | PURG |
| ENSG0000 | 1.347267 | 2.544297 | 1.22E-05 | 0.000286 | GXYLT2 |
| ENSG0000 | 4.882926 | 29.50579 | 0.0046 | 0.03822 | ADAMTS20 |
| ENSG0000 | 1.350227 | 2.549523 | 0.001859 | 0.018898 | ZNF483 |
| ENSG0000 | 1.846974 | 3.597448 | 5.24E-09 | 2.79E-07 | SULT1B1 |
| ENSG0000 | 4.321093 | 19.98842 | 0.00064 | 0.008055 | C9orf131 |
| ENSG0000 | 1.080244 | 2.114394 | 4.48E-13 | 4.75E-11 | MARCKSL1 |
| ENSG0000 | 2.160455 | 4.470558 | 2.04E-06 | 6.01E-05 | CHST1 |
| ENSG0000 | 1.351917 | 2.552511 | 7.97E-29 | 5.43E-26 | CLCF1 |
| ENSG0000 | 4.836458 | 28.57057 | 0.004783 | 0.039402 | None |
| ENSG0000 | 1.71785 | 3.289459 | 0.003624 | 0.031768 | CRYBG2 |
| ENSG0000 | 1.590474 | 3.011483 | 0.004444 | 0.037157 | FOXG1 |
| ENSG0000 | 5.796255 | 55.57078 | 2.45E-05 | 0.000519 | IRX5 |
| ENSG0000 | 5.665458 | 50.75429 | 0.000306 | 0.004385 | GPR4 |
| ENSG0000 | 4.258857 | 19.14449 | 0.002762 | 0.025791 | None |
| ENSG0000 | 1.122957 | 2.177929 | 8.55E-05 | 0.001531 | CTXN1 |
| ENSG0000 | 1.453256 | 2.738254 | 4.21E-60 | 2.41E-56 | THBD |
| ENSG0000 | 1.01165 | 2.016216 | 3.32E-06 | 9.22E-05 | VWA1 |
| ENSG0000 | 1.062511 | 2.088564 | 8.24E-08 | 3.37E-06 | ZFPM1 |
| ENSG0000 | 1.6297 | 3.094487 | 0.00012 | 0.002039 | CITED4 |
| ENSG0000 | 1.532707 | 2.893283 | 0.000241 | 0.003602 | MAPK15 |
| ENSG0000 | 2.330088 | 5.028359 | 1.96E-08 | 9.36E-07 | RGMA |
| ENSG0000 | 1.227849 | 2.342176 | 0.005198 | 0.041885 | CRIP2 |
| ENSG0000 | 1.08267 | 2.117953 | 4.45E-06 | 0.000119 | GJC1 |
| ENSG0000 | 5.735797 | 53.29015 | 0.000103 | 0.001803 | NKX2-5 |
| ENSG0000 | 5.452161 | 43.77883 | 0.001757 | 0.018063 | CABCOCO1 |

| | | | | | |
|---|---|---|---|---|---|
| ENSG0000 | 1.672133 | 3.186854 | 0.000222 | 0.003381 | GRIN2A |
| ENSG0000 | 3.941493 | 15.36412 | 0.004878 | 0.039932 | COL18A1-AS1 |
| ENSG0000 | 1.000162 | 2.000225 | 3.33E-27 | 1.87E-24 | RFLNB |
| ENSG0000 | 1.304934 | 2.470724 | 0.002147 | 0.021178 | KIRREL1 |
| ENSG0000 | 1.857403 | 3.623547 | 1.26E-06 | 3.95E-05 | ARSI |
| ENSG0000 | 1.190951 | 2.283031 | 0.005719 | 0.044939 | NPW |
| ENSG0000 | 1.555217 | 2.938779 | 1.54E-05 | 0.00035 | CLDN5 |
| ENSG0000 | 1.245332 | 2.37073 | 0.000233 | 0.003506 | PKP3 |
| ENSG0000 | 1.339811 | 2.531182 | 1.54E-25 | 6.88E-23 | TAFA3 |
| ENSG0000 | 5.392756 | 42.01276 | 0.002468 | 0.023558 | LINC00313 |
| ENSG0000 | 1.482882 | 2.795065 | 6.02E-22 | 1.85E-19 | NOTUM |
| ENSG0000 | 5.574209 | 47.64354 | 7.59E-05 | 0.001389 | NTF3 |
| ENSG0000 | 1.514577 | 2.85715 | 0.001533 | 0.016218 | MATN1-AS1 |
| ENSG0000 | 5.821666 | 56.55827 | 0.000348 | 0.004877 | |
| ENSG0000 | 1.05946 | 2.084151 | 5.54E-08 | 2.36E-06 | HPDL |
| ENSG0000 | 1.139496 | 2.203041 | 6.21E-06 | 0.00016 | TEAD1 |
| ENSG0000 | 1.253552 | 2.384277 | 9.01E-06 | 0.000219 | C11orf96 |
| ENSG0000 | 1.026929 | 2.037682 | 4.56E-09 | 2.46E-07 | PLEKHN1 |
| ENSG0000 | 1.391191 | 2.622952 | 4.07E-05 | 0.000806 | PERM1 |
| ENSG0000 | 2.618428 | 6.140805 | 0.000184 | 0.002879 | C5orf52 |
| ENSG0000 | 5.284827 | 38.98444 | 0.000592 | 0.007541 | C9orf153 |
| ENSG0000 | 1.560949 | 2.950479 | 3.85E-05 | 0.000768 | NCCRP1 |
| ENSG0000 | 1.876109 | 3.670836 | 0.002833 | 0.026277 | HMX2 |
| ENSG0000 | 1.199231 | 2.296172 | 5.09E-06 | 0.000134 | RPSAP47 |
| ENSG0000 | 1.070858 | 2.100683 | 1.76E-09 | 1.03E-07 | SP6 |
| ENSG0000 | 1.326816 | 2.508484 | 0.003687 | 0.032178 | CLDN4 |
| ENSG0000 | 5.197515 | 36.69509 | 0.000488 | 0.006438 | SH2D5 |
| ENSG0000 | 1.052255 | 2.073768 | 0.000159 | 0.002565 | NCR1 |
| ENSG0000 | 5.712593 | 52.43992 | 0.001782 | 0.018241 | MYT1 |
| ENSG0000 | 1.070336 | 2.099922 | 7.30E-12 | 6.44E-10 | RFX8 |
| ENSG0000 | 2.701946 | 6.506792 | 0.000475 | 0.006304 | H4C3 |
| ENSG0000 | 1.286398 | 2.439183 | 0.003688 | 0.032178 | HMGA2-AS1 |
| ENSG0000 | 4.759746 | 27.09108 | 0.002729 | 0.025542 | FAM177B |
| ENSG0000 | 5.876463 | 58.74781 | 0.000174 | 0.002765 | ZNF560 |
| ENSG0000 | 3.600853 | 12.13291 | 0.005523 | 0.043933 | MYL4 |
| ENSG0000 | 1.768278 | 3.406471 | 0.005976 | 0.0464 | ABCA4 |
| ENSG0000 | 1.647107 | 3.132049 | 3.65E-19 | 8.21E-17 | SMOC1 |
| ENSG0000 | 1.717358 | 3.288338 | 4.47E-12 | 4.04E-10 | F5 |
| ENSG0000 | 1.610807 | 3.054227 | 1.09E-28 | 7.25E-26 | APCDD1L |
| ENSG0000 | 1.673236 | 3.189292 | 2.23E-05 | 0.00048 | HSD3BP5 |
| ENSG0000 | 4.939517 | 30.68618 | 0.003304 | 0.029682 | CSAG1 |
| ENSG0000 | 1.351722 | 2.552166 | 3.33E-05 | 0.000678 | SNORD104 |
| ENSG0000 | 6.897254 | 119.2012 | 0.000846 | 0.010134 | Y_RNA |
| ENSG0000 | 4.985233 | 31.67413 | 0.003289 | 0.029598 | SNORA63D |
| ENSG0000 | 4.965753 | 31.24932 | 0.005038 | 0.040925 | Y_RNA |
| ENSG0000 | 6.590537 | 96.37166 | 5.28E-05 | 0.00101 | RNU6-1082P |
| ENSG0000 | 2.846198 | 7.19103 | 0.000798 | 0.009642 | Y_RNA |

| | | | | | |
|---|---|---|---|---|---|
| ENSG0000 | 6.197169 | 73.37258 | 0.004219 | 0.035624 | Y_RNA |
| ENSG0000 | 4.349434 | 20.38498 | 0.00594 | 0.046203 | RNY1P13 |
| ENSG0000 | 1.315774 | 2.489358 | 3.56E-11 | 2.77E-09 | PLPP4 |
| ENSG0000 | 1.368745 | 2.582458 | 0.000231 | 0.003489 | CLLU1-AS1 |
| ENSG0000 | 1.780179 | 3.434689 | 0.00015 | 0.002449 | LINC01602 |
| ENSG0000 | 1.062919 | 2.089154 | 1.01E-11 | 8.74E-10 | ADGRG1 |
| ENSG0000 | 4.987401 | 31.72177 | 0.004024 | 0.034349 | Y_RNA |
| ENSG0000 | 5.645028 | 50.04061 | 0.006397 | 0.048922 | U3 |
| ENSG0000 | 5.079921 | 33.82273 | 0.002046 | 0.020396 | Y_RNA |
| ENSG0000 | 5.488574 | 44.89783 | 0.000903 | 0.010654 | MIR645 |
| ENSG0000 | 5.884706 | 59.08442 | 0.00084 | 0.010075 | SNORD105 |
| ENSG0000 | 1.471625 | 2.773341 | 1.49E-09 | 8.79E-08 | MT-TY |
| ENSG0000 | 3.826246 | 14.18453 | 0.002783 | 0.025928 | MT-TS1 |
| ENSG0000 | 1.703561 | 3.257038 | 0.000395 | 0.005413 | MT-TG |
| ENSG0000 | 1.263541 | 2.400843 | 0.006052 | 0.046864 | MT-TH |
| ENSG0000 | 1.311427 | 2.481869 | 1.11E-05 | 0.000263 | MT-TL2 |
| ENSG0000 | 1.280722 | 2.429605 | 0.000287 | 0.004181 | MT-TE |
| ENSG0000 | 1.524898 | 2.877663 | 0.000583 | 0.007452 | MT-TT |
| ENSG0000 | 2.194921 | 4.578645 | 0.001136 | 0.012891 | TRGV1 |
| ENSG0000 | 2.482744 | 5.589596 | 0.005163 | 0.041659 | TRAJ23 |
| ENSG0000 | 2.15055 | 4.439969 | 0.005095 | 0.041305 | SNORA3B |
| ENSG0000 | 1.575057 | 2.979473 | 0.002822 | 0.026194 | None |
| ENSG0000 | 3.902301 | 14.95235 | 0.002732 | 0.025549 | MAGEA12 |
| ENSG0000 | 3.290057 | 9.78151 | 0.001633 | 0.017093 | None |
| ENSG0000 | 1.171595 | 2.252606 | 0.000178 | 0.002808 | S1PR3 |
| ENSG0000 | 5.361716 | 41.11852 | 0.001192 | 0.013388 | RAB5CP1 |
| ENSG0000 | 1.774175 | 3.420424 | 0.000932 | 0.010926 | MPRIPP1 |
| ENSG0000 | 6.164505 | 71.73003 | 0.000297 | 0.004281 | MARCKSL1P2 |
| ENSG0000 | 4.140608 | 17.63791 | 0.00142 | 0.015328 | RPL31P52 |
| ENSG0000 | 4.544767 | 23.34055 | 0.003638 | 0.031869 | TAFA5 |
| ENSG0000 | 6.137542 | 70.40186 | 0.00303 | 0.027677 | None |
| ENSG0000 | 4.881397 | 29.47453 | 0.001169 | 0.01319 | None |
| ENSG0000 | 5.505543 | 45.42903 | 0.000801 | 0.009656 | MIR548I2 |
| ENSG0000 | 6.245088 | 75.85058 | 6.53E-05 | 0.001219 | LINC02860 |
| ENSG0000 | 1.069023 | 2.098012 | 2.53E-06 | 7.27E-05 | None |
| ENSG0000 | 4.916128 | 30.19271 | 0.000648 | 0.00813 | RN7SKP150 |
| ENSG0000 | 2.44316 | 5.438317 | 0.003564 | 0.03133 | None |
| ENSG0000 | 1.111536 | 2.160756 | 0.001619 | 0.016981 | None |
| ENSG0000 | 5.80696 | 55.98469 | 0.001729 | 0.017871 | ARHGEF2-AS1 |
| ENSG0000 | 5.128817 | 34.98869 | 0.00385 | 0.033227 | OLFM5P |
| ENSG0000 | 1.527167 | 2.882194 | 3.11E-06 | 8.72E-05 | None |
| ENSG0000 | 5.61173 | 48.89889 | 0.002416 | 0.023218 | ARL14EPP1 |
| ENSG0000 | 1.247583 | 2.374433 | 1.12E-07 | 4.45E-06 | MTND1P23 |
| ENSG0000 | 5.233587 | 37.62415 | 0.004406 | 0.036894 | None |
| ENSG0000 | 1.437402 | 2.708327 | 0.003692 | 0.032197 | CFLAR-AS1 |
| ENSG0000 | 6.930951 | 122.0181 | 1.04E-05 | 0.000247 | RPL22P3 |
| ENSG0000 | 5.505586 | 45.43041 | 0.000555 | 0.007164 | USF1P1 |

| | | | | | |
|---|---|---|---|---|---|
| ENSG0000 | 1.289762 | 2.444878 | 0.004492 | 0.037499 | None |
| ENSG0000 | 5.910793 | 60.16251 | 0.0013 | 0.014275 | SMCR5 |
| ENSG0000 | 5.249633 | 38.04495 | 0.000771 | 0.00935 | MTND5P1 |
| ENSG0000 | 5.424356 | 42.94314 | 0.000902 | 0.01065 | None |
| ENSG0000 | 6.911562 | 120.3892 | 0.000446 | 0.005995 | MTND1P11 |
| ENSG0000 | 3.050972 | 8.287702 | 0.000196 | 0.003037 | LYPLAL1-AS1 |
| ENSG0000 | 6.180999 | 72.55481 | 0.003899 | 0.033522 | MTND1P9 |
| ENSG0000 | 5.537089 | 46.43332 | 0.000287 | 0.004178 | None |
| ENSG0000 | 4.967648 | 31.2904 | 0.005529 | 0.043967 | PSMA6P2 |
| ENSG0000 | 1.299493 | 2.461424 | 0.002316 | 0.022426 | HLA-DRB6 |
| ENSG0000 | 4.855364 | 28.94744 | 0.001959 | 0.019719 | None |
| ENSG0000 | 1.002454 | 2.003406 | 0.00102 | 0.011751 | ATXN1-AS1 |
| ENSG0000 | 5.211172 | 37.04412 | 0.002183 | 0.021446 | None |
| ENSG0000 | 1.088088 | 2.125921 | 0.000225 | 0.003416 | None |
| ENSG0000 | 2.619189 | 6.144045 | 7.81E-06 | 0.000196 | |
| ENSG0000 | 5.100689 | 34.31314 | 0.005646 | 0.044628 | STK24P1 |
| ENSG0000 | 5.723446 | 52.83587 | 0.001226 | 0.013671 | TRBV5-4 |
| ENSG0000 | 1.339824 | 2.531204 | 0.000212 | 0.003246 | None |
| ENSG0000 | 5.118392 | 34.73678 | 0.00602 | 0.046643 | None |
| ENSG0000 | 1.547174 | 2.922442 | 4.27E-37 | 6.11E-34 | APCDD1L-DT |
| ENSG0000 | 1.041113 | 2.057815 | 4.60E-05 | 0.000893 | EMSLR |
| ENSG0000 | 5.06526 | 33.48076 | 0.006082 | 0.046997 | None |
| ENSG0000 | 1.106352 | 2.153006 | 0.001901 | 0.01921 | None |
| ENSG0000 | 1.249524 | 2.37763 | 0.000543 | 0.007035 | SLCO4A1-AS1 |
| ENSG0000 | 2.753128 | 6.741773 | 0.003283 | 0.029577 | CRYZP1 |
| ENSG0000 | 5.830112 | 56.89036 | 0.000158 | 0.002561 | None |
| ENSG0000 | 5.979118 | 63.08033 | 0.000292 | 0.004221 | CSNK1G2P1 |
| ENSG0000 | 1.373845 | 2.591604 | 0.002723 | 0.025517 | None |
| ENSG0000 | 5.220943 | 37.29584 | 0.000211 | 0.003244 | None |
| ENSG0000 | 5.562217 | 47.24918 | 0.00074 | 0.009026 | NSRP1P1 |
| ENSG0000 | 5.995277 | 63.79084 | 0.000509 | 0.006679 | SNRPFP4 |
| ENSG0000 | 5.491038 | 44.97459 | 0.000226 | 0.00343 | None |
| ENSG0000 | 6.18976 | 72.99672 | 7.42E-06 | 0.000187 | COX7CP1 |
| ENSG0000 | 5.75267 | 53.91706 | 0.00591 | 0.046035 | None |
| ENSG0000 | 5.613535 | 48.96012 | 3.65E-05 | 0.000734 | EHMT2-AS1 |
| ENSG0000 | 6.388331 | 83.76824 | 0.000983 | 0.011414 | VDAC1P11 |
| ENSG0000 | 1.37068 | 2.585925 | 0.000734 | 0.008973 | DGUOK-AS1 |
| ENSG0000 | 6.324914 | 80.16576 | 7.02E-05 | 0.0013 | None |
| ENSG0000 | 5.590653 | 48.18971 | 0.001427 | 0.015392 | RNU7-38P |
| ENSG0000 | 2.151822 | 4.443888 | 6.23E-08 | 2.61E-06 | PCDHGC5 |
| ENSG0000 | 5.207066 | 36.93883 | 0.004412 | 0.036909 | None |
| ENSG0000 | 1.006494 | 2.009023 | 0.002503 | 0.023815 | EGFL8 |
| ENSG0000 | 5.473621 | 44.43489 | 0.000449 | 0.006028 | None |
| ENSG0000 | 1.66084 | 3.162006 | 0.000836 | 0.010034 | None |
| ENSG0000 | 5.396332 | 42.11705 | 0.002324 | 0.02248 | None |
| ENSG0000 | 2.533977 | 5.791661 | 0.004411 | 0.036909 | RPSAP52 |
| ENSG0000 | 6.557533 | 94.192 | 0.000258 | 0.00382 | None |

| | | | | | |
|---|---|---|---|---|---|
| ENSG0000 | 5.450484 | 43.72795 | 0.000525 | 0.00686 | PDE6B-AS1 |
| ENSG0000 | 3.556971 | 11.76942 | 0.005342 | 0.04278 | GATA2-AS1 |
| ENSG0000 | 5.411339 | 42.55743 | 0.000316 | 0.004498 | DUXAP10 |
| ENSG0000 | 5.206482 | 36.92387 | 0.000986 | 0.011423 | None |
| ENSG0000 | 4.867735 | 29.19673 | 0.002823 | 0.026194 | RPL12P33 |
| ENSG0000 | 1.059216 | 2.083798 | 0.000247 | 0.003681 | WNT5A-AS1 |
| ENSG0000 | 5.583056 | 47.9366 | 0.006434 | 0.049172 | None |
| ENSG0000 | 3.829579 | 14.21733 | 0.005193 | 0.041856 | None |
| ENSG0000 | 1.888634 | 3.702845 | 4.65E-12 | 4.16E-10 | TMEM158 |
| ENSG0000 | 1.225335 | 2.338097 | 5.77E-13 | 6.02E-11 | SHANK3 |
| ENSG0000 | 6.892875 | 118.8398 | 1.88E-05 | 0.000414 | None |
| ENSG0000 | 5.465254 | 44.17793 | 0.000106 | 0.001846 | None |
| ENSG0000 | 5.388492 | 41.88878 | 0.001622 | 0.017005 | RNU7-18P |
| ENSG0000 | 3.091463 | 8.523599 | 0.005844 | 0.045629 | SNORA74D |
| ENSG0000 | 5.35272 | 40.86292 | 0.003739 | 0.032537 | RNU4ATAC12P |
| ENSG0000 | 4.951649 | 30.94532 | 0.001147 | 0.012984 | USP12P1 |
| ENSG0000 | 1.11031 | 2.158921 | 3.74E-06 | 0.000102 | TRNP1 |
| ENSG0000 | 4.768715 | 27.26003 | 0.002602 | 0.024564 | None |
| ENSG0000 | 1.38528 | 2.612226 | 0.001358 | 0.014777 | ALG1L10P |
| ENSG0000 | 5.164692 | 35.86966 | 0.000593 | 0.007552 | None |
| ENSG0000 | 4.936937 | 30.63135 | 0.002093 | 0.020765 | LINC02749 |
| ENSG0000 | 6.163642 | 71.68713 | 0.000186 | 0.002911 | None |
| ENSG0000 | 1.188198 | 2.27868 | 1.45E-09 | 8.58E-08 | MEX3A |
| ENSG0000 | 1.391456 | 2.623433 | 0.002101 | 0.020793 | None |
| ENSG0000 | 4.556452 | 23.53037 | 0.006109 | 0.047134 | LINC02551 |
| ENSG0000 | 1.249335 | 2.377318 | 7.51E-05 | 0.001376 | SNHG9 |
| ENSG0000 | 5.379484 | 41.62803 | 0.005973 | 0.046399 | DDX18P5 |
| ENSG0000 | 1.033379 | 2.046812 | 8.95E-09 | 4.55E-07 | None |
| ENSG0000 | 5.240135 | 37.79529 | 0.005045 | 0.040975 | ELOCP31 |
| ENSG0000 | 1.149669 | 2.21863 | 0.000395 | 0.005413 | SBNO1-AS1 |
| ENSG0000 | 5.522391 | 45.96268 | 0.004643 | 0.038462 | RPL7AP3 |
| ENSG0000 | 1.074707 | 2.106295 | 0.001999 | 0.020018 | LINC02454 |
| ENSG0000 | 5.10159 | 34.33456 | 0.003655 | 0.031981 | None |
| ENSG0000 | 1.681904 | 3.208511 | 0.00143 | 0.015413 | SLC6A12-AS1 |
| ENSG0000 | 2.877061 | 7.346521 | 0.003556 | 0.031281 | None |
| ENSG0000 | 1.085197 | 2.121665 | 2.44E-08 | 1.14E-06 | MGAM |
| ENSG0000 | 5.271099 | 38.61524 | 0.001355 | 0.014755 | None |
| ENSG0000 | 6.558182 | 94.23441 | 0.000356 | 0.004963 | None |
| ENSG0000 | 6.915729 | 120.7374 | 0.000277 | 0.004054 | None |
| ENSG0000 | 5.925807 | 60.79189 | 0.004347 | 0.036466 | None |
| ENSG0000 | 1.1329 | 2.192991 | 3.81E-66 | 2.72E-62 | CLEC5A |
| ENSG0000 | 1.004739 | 2.00658 | 0.000211 | 0.003244 | LINC00641 |
| ENSG0000 | 5.774581 | 54.74219 | 0.000156 | 0.002534 | None |
| ENSG0000 | 6.356009 | 81.91232 | 1.14E-05 | 0.000268 | None |
| ENSG0000 | 1.113881 | 2.164271 | 4.61E-06 | 0.000123 | None |
| ENSG0000 | 1.808665 | 3.503179 | 0.000715 | 0.008781 | KIF23-AS1 |
| ENSG0000 | 5.180854 | 36.27376 | 0.001738 | 0.017937 | GCSHP2 |

| | | | | | |
|---|---|---|---|---|---|
| ENSG0000 | 2.165713 | 4.486881 | 0.004086 | 0.034734 | None |
| ENSG0000 | 1.307397 | 2.474946 | 8.77E-11 | 6.34E-09 | SNHG19 |
| ENSG0000 | 5.769161 | 54.53693 | 0.000437 | 0.005895 | TPRKBP2 |
| ENSG0000 | 1.064748 | 2.091805 | 0.006539 | 0.049788 | None |
| ENSG0000 | 6.017604 | 64.78573 | 0.002271 | 0.022123 | ATP5MFP6 |
| ENSG0000 | 5.60034 | 48.51437 | 0.001795 | 0.018353 | TNRC6B-DT |
| ENSG0000 | 4.829396 | 28.43107 | 0.002713 | 0.025431 | None |
| ENSG0000 | 4.661017 | 25.29915 | 0.003356 | 0.030002 | GOLGA8T |
| ENSG0000 | 4.859643 | 29.03342 | 0.001568 | 0.016527 | None |
| ENSG0000 | 1.551787 | 2.931801 | 2.29E-20 | 6.01E-18 | TPBGL |
| ENSG0000 | 1.341464 | 2.534084 | 0.000255 | 0.003786 | None |
| ENSG0000 | 6.445857 | 87.17585 | 6.99E-05 | 0.001296 | None |
| ENSG0000 | 1.012909 | 2.017976 | 0.005916 | 0.046069 | SPON1 |
| ENSG0000 | 5.686343 | 51.49437 | 0.001161 | 0.013121 | MRPS21P9 |
| ENSG0000 | 6.704561 | 104.2975 | 6.04E-05 | 0.001136 | RYKP1 |
| ENSG0000 | 4.864224 | 29.12576 | 0.003512 | 0.031052 | ABHD17AP5 |
| ENSG0000 | 1.521374 | 2.870644 | 0.00579 | 0.045314 | None |
| ENSG0000 | 2.040965 | 4.115208 | 0.001229 | 0.013683 | None |
| ENSG0000 | 2.895102 | 7.438964 | 0.005249 | 0.042142 | MIR4648 |
| ENSG0000 | 6.655988 | 100.8444 | 0.002505 | 0.023831 | None |
| ENSG0000 | 5.047494 | 33.07098 | 0.003307 | 0.029703 | MIR5188 |
| ENSG0000 | 1.643795 | 3.124867 | 0.003205 | 0.028973 | None |
| ENSG0000 | 6.30176 | 78.88943 | 2.69E-06 | 7.69E-05 | None |
| ENSG0000 | 1.246738 | 2.373043 | 3.98E-17 | 7.12E-15 | None |
| ENSG0000 | 5.870919 | 58.52248 | 0.000397 | 0.005437 | None |
| ENSG0000 | 6.395593 | 84.19091 | 0.000295 | 0.004256 | None |
| ENSG0000 | 5.196712 | 36.67467 | 0.002206 | 0.021604 | None |
| ENSG0000 | 2.895864 | 7.442894 | 0.000987 | 0.011437 | None |
| ENSG0000 | 1.392863 | 2.625994 | 0.00349 | 0.030906 | None |
| ENSG0000 | 6.253083 | 76.2721 | 1.85E-05 | 0.00041 | GABRQ |
| ENSG0000 | 5.460453 | 44.03117 | 0.000681 | 0.00845 | AIRN |
| ENSG0000 | 1.156128 | 2.228585 | 2.98E-05 | 0.000616 | LINC01711 |
| ENSG0000 | 6.822839 | 113.2086 | 0.000273 | 0.004009 | None |
| ENSG0000 | 6.258014 | 76.53322 | 0.002776 | 0.025893 | RMRP |
| ENSG0000 | 3.370095 | 10.3395 | 0.006532 | 0.049755 | HEATR9 |
| ENSG0000 | 5.66073 | 50.58823 | 0.000575 | 0.007379 | None |
| ENSG0000 | 7.397592 | 168.6153 | 3.70E-08 | 1.65E-06 | None |
| ENSG0000 | 4.427018 | 21.51122 | 0.002973 | 0.027304 | None |
| ENSG0000 | 6.513971 | 91.39044 | 6.82E-05 | 0.001268 | None |
| ENSG0000 | 5.391485 | 41.97578 | 0.006184 | 0.047602 | None |
| ENSG0000 | 1.248273 | 2.375569 | 0.003365 | 0.030036 | None |
| ENSG0000 | 1.374055 | 2.591981 | 0.000603 | 0.007658 | MIR4787 |
| ENSG0000 | 5.147026 | 35.43311 | 0.001456 | 0.015625 | None |
| ENSG0000 | 1.074729 | 2.106327 | 0.001279 | 0.014092 | None |
| ENSG0000 | 3.775847 | 13.69756 | 0.006368 | 0.048741 | None |
| ENSG0000 | 6.090623 | 68.14912 | 0.001223 | 0.013665 | H4C6 |
| ENSG0000 | 1.597378 | 3.025929 | 6.53E-05 | 0.001219 | None |

| | | | | | |
|---|---|---|---|---|---|
| ENSG0000 | 5.614329 | 48.98708 | 0.001291 | 0.01421 | MIR7152 |
| ENSG0000 | 4.479613 | 22.30991 | 0.00023 | 0.003486 | H2AC12 |
| ENSG0000 | 7.134152 | 140.4733 | 7.52E-05 | 0.001377 | H4C13 |
| ENSG0000 | 1.360821 | 2.568313 | 1.07E-07 | 4.28E-06 | LINC02340 |
| ENSG0000 | 5.143264 | 35.34083 | 0.006245 | 0.04798 | None |
| ENSG0000 | 4.774028 | 27.3606 | 0.001539 | 0.016269 | H3C11 |
| ENSG0000 | 1.491944 | 2.812678 | 0.000744 | 0.009068 | H2BC9 |
| ENSG0000 | 5.6646 | 50.72411 | 0.000419 | 0.005689 | None |
| ENSG0000 | 2.162599 | 4.477206 | 0.004788 | 0.039427 | DACH1 |
| ENSG0000 | 5.923058 | 60.67616 | 0.002014 | 0.020133 | Metazoa_SRP |
| ENSG0000 | 6.276534 | 77.522 | 0.003983 | 0.034095 | None |
| ENSG0000 | 5.721444 | 52.76262 | 0.000303 | 0.004346 | None |
| ENSG0000 | 1.25038 | 2.37904 | 0.005024 | 0.040834 | H4C5 |
| ENSG0000 | 5.860127 | 58.08636 | 0.000154 | 0.002506 | None |
| ENSG0000 | 7.930294 | 243.925 | 3.94E-07 | 1.39E-05 | H3C7 |
| ENSG0000 | 5.836062 | 57.12548 | 0.006068 | 0.046938 | None |
| ENSG0000 | 5.852852 | 57.79418 | 6.76E-05 | 0.001258 | RN7SL113P |
| ENSG0000 | 4.398193 | 21.08571 | 0.006008 | 0.046598 | H4C1 |
| ENSG0000 | 2.778074 | 6.859361 | 0.003164 | 0.02865 | None |
| ENSG0000 | 7.313641 | 159.0836 | 8.57E-06 | 0.000211 | None |
| ENSG0000 | 1.209858 | 2.313149 | 0.003671 | 0.032095 | None |
| ENSG0000 | 1.252143 | 2.381949 | 0.001112 | 0.012634 | None |
| ENSG0000 | 4.298897 | 19.68326 | 0.003006 | 0.027507 | None |
| ENSG0000 | 1.039303 | 2.055234 | 0.005589 | 0.044333 | None |
| ENSG0000 | 2.920459 | 7.570868 | 0.000767 | 0.009306 | None |
| ENSG0000 | 5.183597 | 36.34279 | 0.001493 | 0.015901 | None |
| ENSG0000 | 2.145912 | 4.425721 | 0.000968 | 0.011259 | None |
| ENSG0000 | 6.512853 | 91.3196 | 0.000805 | 0.009696 | None |
| ENSG0000 | 1.058225 | 2.082368 | 0.002782 | 0.025928 | None |
| ENSG0000 | 2.319955 | 4.993167 | 0.00069 | 0.00854 | |
| ENSG0000 | 5.846817 | 57.55293 | 0.004216 | 0.035612 | None |
| ENSG0000 | 1.285961 | 2.438444 | 0.000131 | 0.002203 | None |
| ENSG0000 | 4.608047 | 24.3871 | 0.003029 | 0.027677 | None |
| ENSG0000 | 1.288101 | 2.442063 | 0.002834 | 0.026277 | SLFNL1-AS1 |
| ENSG0000 | 4.961141 | 31.14958 | 0.002952 | 0.027146 | DPRXP3 |
| ENSG0000 | 5.675433 | 51.10642 | 0.001771 | 0.018152 | None |
| ENSG0000 | 5.059128 | 33.33876 | 0.002244 | 0.021913 | LINC02452 |
