## Supplementary material for "Unravelling neuronal and glial differences in ceramide composition, synthesis, and sensitivity to toxicity": MN astrocyte microglia common GCSi gene changes

| GeneID | name | Common GCSi changes in Astrocytes and Microglia |
|---|---|---|
| ENSG0000 | PTPRN | |
| ENSG0000 | IL11 | |
| ENSG0000 | TFPI2 | |
| ENSG0000 | SERPINE1 | |
| ENSG0000 | ADGB | |
| ENSG0000 | CCDC170 | |
| ENSG0000 | SLPI | |
| ENSG0000 | GFPT2 | |
| ENSG0000 | AK7 | |
| ENSG0000 | FBXO15 | |
| ENSG0000 | BANK1 | |
| ENSG0000 | PDZRN4 | |
| ENSG0000 | DCDC1 | |
| ENSG0000 | ZNF483 | |
| ENSG0000 | CLCF1 | |
| ENSG0000 | PCED1B | |
| ENSG0000 | CABCOCO1 | |
| ENSG0000 | CLDN5 | |
| ENSG0000 | SH2D5 | |
| ENSG0000 | DTHD1 | |
| ENSG0000 | SMOC1 | |
| ENSG0000 | None | |
| ENSG0000 | None | |
| ENSG0000 | SPON1 | |

| GeneID | name | Common GCSi gene changes in Microglia and MNs |
| --- | --- | --- |
| ENSG0000 | FAP | |
| ENSG0000 | IL18RAP | |
| ENSG0000 | GIMAP4 | |
| ENSG0000 | ASB15 | |
| ENSG0000 | CD1B | |
| ENSG0000 | IGF2 | |
| ENSG0000 | S100Z | |
| ENSG0000 | None | |
| ENSG0000 | TAFA3 | |
| ENSG0000 | ACTBP11 | |
| ENSG0000 | MYL4 | |
| ENSG0000 | None | |
| ENSG0000 | HLA-DRB6 | |
| ENSG0000 | None | |
| ENSG0000 | None | |
| ENSG0000 | None | |

| GeneID | name | Common GCSi gene changes in Astrocytes and MNs |
|---|---|---|
| ENSG0000 | PRRG3 | |
| ENSG0000 | None | |

| GeneID | Treatment | Treatment | Treatment | Treatment | name |
|---|---|---|---|---|---|
| ENSG0000 | -1.16653 | -2.24471 | 3.21E-05 | 0.000478 | PDK4 |
| ENSG0000 | 1.100447 | 2.144212 | 3.33E-15 | 7.66E-13 | TNFRSF12A |
| ENSG0000 | -1.24455 | -2.36944 | 0.002302 | 0.015921 | DNAH9 |
| ENSG0000 | -1.16074 | -2.23571 | 1.78E-10 | 1.33E-08 | DLEC1 |
| ENSG0000 | -1.2906 | -2.4463 | 0.007281 | 0.038655 | PLEKHG6 |
| ENSG0000 | -1.07957 | -2.1134 | 5.09E-10 | 3.45E-08 | SERPINB1 |
| ENSG0000 | 1.21005 | 2.313457 | 7.58E-05 | 0.000986 | BIRC3 |
| ENSG0000 | -1.50181 | -2.83199 | 1.07E-09 | 6.59E-08 | TYMP |
| ENSG0000 | -1.11024 | -2.15881 | 3.73E-12 | 3.98E-10 | EFCAB1 |
| ENSG0000 | -1.134 | -2.19466 | 5.18E-06 | 0.000102 | LMO3 |
| ENSG0000 | -1.54397 | -2.91596 | 3.18E-10 | 2.23E-08 | ENTPD2 |
| ENSG0000 | 1.090425 | 2.129368 | 1.54E-07 | 4.95E-06 | PTPRN |
| ENSG0000 | -1.03027 | -2.0424 | 1.24E-09 | 7.40E-08 | TRAF1 |
| ENSG0000 | -1.12434 | -2.18002 | 0.00955 | 0.047697 | SPAG4 |
| ENSG0000 | -1.7656 | -3.40017 | 4.18E-07 | 1.17E-05 | ZMYND12 |
| ENSG0000 | 1.506003 | 2.840221 | 4.20E-21 | 3.45E-18 | SREBF1 |
| ENSG0000 | -1.30103 | -2.46405 | 2.54E-05 | 0.000394 | ADCYAP1R1 |
| ENSG0000 | 1.054077 | 2.076389 | 0.005059 | 0.029193 | ATP8B1 |
| ENSG0000 | 1.054422 | 2.076886 | 9.68E-06 | 0.000175 | MPP4 |
| ENSG0000 | -1.29272 | -2.4499 | 0.002701 | 0.018019 | KCNK2 |
| ENSG0000 | -1.2822 | -2.43209 | 0.000677 | 0.00602 | SLCO1A2 |
| ENSG0000 | 1.513387 | 2.854795 | 1.78E-10 | 1.33E-08 | ICAM1 |
| ENSG0000 | -1.36201 | -2.57043 | 7.39E-06 | 0.000139 | ABCC6 |
| ENSG0000 | -1.69234 | -3.2318 | 1.44E-20 | 9.59E-18 | ADA2 |
| ENSG0000 | 2.469383 | 5.538069 | 1.12E-07 | 3.74E-06 | IL11 |
| ENSG0000 | 1.577564 | 2.984654 | 2.87E-29 | 1.64E-25 | SCD |
| ENSG0000 | -1.62608 | -3.08672 | 5.17E-07 | 1.41E-05 | RASL10A |
| ENSG0000 | -1.22701 | -2.34081 | 1.91E-08 | 8.20E-07 | ISM2 |
| ENSG0000 | -1.58221 | -2.99429 | 4.55E-27 | 1.31E-23 | EFS |
| ENSG0000 | -1.24137 | -2.36422 | 3.87E-09 | 1.99E-07 | NKAIN4 |
| ENSG0000 | -1.26012 | -2.39515 | 0.000974 | 0.00812 | RGCC |
| ENSG0000 | 1.168175 | 2.247272 | 6.10E-10 | 4.04E-08 | RELB |
| ENSG0000 | -1.06235 | -2.08834 | 9.19E-07 | 2.33E-05 | PEX11G |
| ENSG0000 | 1.317504 | 2.492345 | 0.006039 | 0.033622 | ICAM5 |
| ENSG0000 | -1.7932 | -3.46582 | 8.78E-10 | 5.52E-08 | ODAD1 |
| ENSG0000 | 1.244811 | 2.369875 | 1.15E-17 | 4.31E-15 | TFPI2 |
| ENSG0000 | -1.01469 | -2.02047 | 4.21E-05 | 0.0006 | DNAH11 |
| ENSG0000 | 1.788596 | 3.454786 | 5.72E-18 | 2.41E-15 | SERPINE1 |
| ENSG0000 | -1.08063 | -2.11496 | 0.002279 | 0.015796 | VSIR |
| ENSG0000 | 1.150505 | 2.219916 | 1.64E-06 | 3.79E-05 | ACTA2 |
| ENSG0000 | 1.281685 | 2.431228 | 2.87E-06 | 6.17E-05 | DKK1 |
| ENSG0000 | 1.37118 | 2.586821 | 6.70E-14 | 1.07E-11 | CCL2 |
| ENSG0000 | 1.379297 | 2.601416 | 0.001467 | 0.011244 | VWF |
| ENSG0000 | -1.03753 | -2.05272 | 3.35E-23 | 5.27E-20 | RSPH4A |
| ENSG0000 | -1.24636 | -2.37242 | 1.21E-12 | 1.39E-10 | MDFI |
| ENSG0000 | -1.02953 | -2.04135 | 0.002622 | 0.017588 | SERPINI2 |

| | | | | | |
|---|---|---|---|---|---|
| ENSG0000 | -1.34229 | -2.53554 | 0.001101 | 0.008975 | CFAP92 |
| ENSG0000 | 1.178138 | 2.262845 | 2.80E-09 | 1.51E-07 | IL1R1 |
| ENSG0000 | 1.175926 | 2.259379 | 7.21E-22 | 7.33E-19 | GADD45A |
| ENSG0000 | 2.174913 | 4.515585 | 1.47E-18 | 6.85E-16 | RGS4 |
| ENSG0000 | -1.31129 | -2.48163 | 0.000106 | 0.001296 | IRAG2 |
| ENSG0000 | -1.58704 | -3.00432 | 0.00506 | 0.029193 | ADGB |
| ENSG0000 | 1.311658 | 2.482267 | 1.24E-20 | 8.60E-18 | TNFAIP3 |
| ENSG0000 | 1.363292 | 2.572716 | 8.93E-21 | 6.88E-18 | SGK1 |
| ENSG0000 | 1.105419 | 2.151613 | 1.58E-05 | 0.000264 | CCN2 |
| ENSG0000 | -1.07571 | -2.10776 | 8.03E-08 | 2.81E-06 | ECRG4 |
| ENSG0000 | 1.18038 | 2.266365 | 1.96E-08 | 8.34E-07 | DUSP1 |
| ENSG0000 | -1.13843 | -2.20142 | 1.46E-19 | 7.84E-17 | CCDC170 |
| ENSG0000 | 1.604952 | 3.041856 | 2.57E-16 | 7.29E-14 | EGR1 |
| ENSG0000 | 1.592829 | 3.016404 | 3.79E-29 | 1.64E-25 | DUSP4 |
| ENSG0000 | -1.27364 | -2.41771 | 1.04E-06 | 2.59E-05 | EPHX2 |
| ENSG0000 | 1.222835 | 2.334049 | 2.03E-06 | 4.55E-05 | SV2C |
| ENSG0000 | -1.64824 | -3.13452 | 9.81E-21 | 7.06E-18 | DNAI1 |
| ENSG0000 | 1.904321 | 3.743327 | 7.81E-22 | 7.50E-19 | EGR2 |
| ENSG0000 | -1.13631 | -2.19818 | 0.00124 | 0.009825 | NFE2 |
| ENSG0000 | -1.21806 | -2.32634 | 0.00484 | 0.028169 | SLPI |
| ENSG0000 | -1.40341 | -2.64526 | 1.02E-05 | 0.000183 | KCNS1 |
| ENSG0000 | -1.01215 | -2.01691 | 1.85E-05 | 0.000301 | MATN4 |
| ENSG0000 | -1.07972 | -2.11362 | 2.64E-06 | 5.74E-05 | TOX2 |
| ENSG0000 | 1.027337 | 2.038259 | 9.83E-05 | 0.001219 | CHST8 |
| ENSG0000 | 1.05892 | 2.083371 | 0.000127 | 0.001505 | GRPR |
| ENSG0000 | 1.258038 | 2.391703 | 8.05E-11 | 6.63E-09 | WNK4 |
| ENSG0000 | -1.07993 | -2.11394 | 6.17E-15 | 1.38E-12 | MASP1 |
| ENSG0000 | 1.214656 | 2.320854 | 2.44E-14 | 4.54E-12 | LIF |
| ENSG0000 | 1.791326 | 3.461329 | 1.05E-13 | 1.63E-11 | CPA4 |
| ENSG0000 | 1.556878 | 2.942165 | 6.80E-10 | 4.47E-08 | VGF |
| ENSG0000 | -2.00088 | -4.00243 | 0.000668 | 0.005968 | TNNI3 |
| ENSG0000 | -1.02068 | -2.02888 | 0.001378 | 0.010674 | PRRG3 |
| ENSG0000 | 1.098386 | 2.14115 | 0.002031 | 0.014429 | NALF2 |
| ENSG0000 | -4.17499 | -18.0633 | 0.008149 | 0.042113 | KLHDC7B |
| ENSG0000 | -1.36299 | -2.57218 | 0.000163 | 0.001859 | CALY |
| ENSG0000 | -1.15179 | -2.2219 | 1.31E-08 | 5.87E-07 | THEMIS2 |
| ENSG0000 | 1.019768 | 2.027593 | 5.49E-14 | 8.95E-12 | GFPT2 |
| ENSG0000 | -1.30444 | -2.46987 | 9.22E-13 | 1.09E-10 | PPP1R1B |
| ENSG0000 | -1.31136 | -2.48175 | 9.64E-11 | 7.86E-09 | RGS22 |
| ENSG0000 | -1.09261 | -2.13259 | 2.36E-05 | 0.000369 | ANKEF1 |
| ENSG0000 | 1.492986 | 2.81471 | 4.54E-23 | 6.54E-20 | CHI3L1 |
| ENSG0000 | -1.22291 | -2.33418 | 0.003241 | 0.020676 | STOML3 |
| ENSG0000 | -1.14713 | -2.21473 | 0.000435 | 0.004221 | SFTPD |
| ENSG0000 | -2.05513 | -4.15581 | 1.69E-10 | 1.28E-08 | DYDC2 |
| ENSG0000 | -1.43658 | -2.70678 | 1.16E-07 | 3.84E-06 | SYT6 |
| ENSG0000 | -1.12561 | -2.18193 | 3.71E-09 | 1.92E-07 | MYCN |
| ENSG0000 | -10.2102 | -1184.65 | 0.000251 | 0.002686 | SAA2 |

| | | | | | |
|---|---|---|---|---|---|
| ENSG0000 | 2.061046 | 4.172888 | 3.71E-18 | 1.60E-15 | SPOCD1 |
| ENSG0000 | 1.002531 | 2.003512 | 1.44E-11 | 1.38E-09 | ETS1 |
| ENSG0000 | 2.087594 | 4.250386 | 4.31E-06 | 8.69E-05 | EGR4 |
| ENSG0000 | -1.06407 | -2.09082 | 5.25E-08 | 1.93E-06 | AGT |
| ENSG0000 | -1.03566 | -2.05005 | 0.002044 | 0.014497 | PCDH8 |
| ENSG0000 | -1.6916 | -3.23014 | 7.02E-09 | 3.33E-07 | CNMD |
| ENSG0000 | 2.089025 | 4.254603 | 1.11E-06 | 2.73E-05 | GPNMB |
| ENSG0000 | -1.77908 | -3.43208 | 5.29E-07 | 1.44E-05 | MYCBPAP |
| ENSG0000 | -1.12309 | -2.17813 | 0.0037 | 0.022912 | CHAD |
| ENSG0000 | -1.03796 | -2.05332 | 2.96E-10 | 2.08E-08 | WDR38 |
| ENSG0000 | 1.135619 | 2.197128 | 1.33E-15 | 3.27E-13 | IER3 |
| ENSG0000 | -1.45602 | -2.7435 | 1.09E-09 | 6.64E-08 | TTC29 |
| ENSG0000 | -1.41206 | -2.66116 | 3.36E-07 | 9.62E-06 | SULF1 |
| ENSG0000 | -1.01524 | -2.02124 | 8.68E-10 | 5.50E-08 | CFAP300 |
| ENSG0000 | 1.160816 | 2.235839 | 1.58E-10 | 1.21E-08 | THBS1 |
| ENSG0000 | 1.001483 | 2.002057 | 4.16E-08 | 1.59E-06 | DUSP6 |
| ENSG0000 | -1.1312 | -2.19041 | 6.69E-08 | 2.41E-06 | ITGB7 |
| ENSG0000 | -1.23533 | -2.35435 | 1.05E-11 | 1.03E-09 | MORN3 |
| ENSG0000 | -1.15309 | -2.2239 | 8.83E-20 | 5.09E-17 | AK7 |
| ENSG0000 | -1.33268 | -2.5187 | 1.04E-05 | 0.000185 | WDR93 |
| ENSG0000 | -1.23948 | -2.36113 | 1.46E-05 | 0.000248 | MYLK3 |
| ENSG0000 | -1.36978 | -2.58432 | 0.001952 | 0.014004 | IRF8 |
| ENSG0000 | -1.02859 | -2.04002 | 6.51E-11 | 5.51E-09 | LRRC46 |
| ENSG0000 | -1.38914 | -2.61922 | 0.003238 | 0.020668 | SLC14A1 |
| ENSG0000 | -1.66081 | -3.16195 | 0.007223 | 0.03842 | SLC13A5 |
| ENSG0000 | -1.14093 | -2.20524 | 0.000215 | 0.002348 | FBXO15 |
| ENSG0000 | 1.581609 | 2.993036 | 0.007304 | 0.038723 | PMAIP1 |
| ENSG0000 | -1.63483 | -3.1055 | 1.90E-13 | 2.76E-11 | CFAP74 |
| ENSG0000 | 1.041503 | 2.058371 | 5.87E-09 | 2.82E-07 | CCN1 |
| ENSG0000 | -1.06958 | -2.09882 | 1.96E-07 | 6.16E-06 | RXRG |
| ENSG0000 | 1.034074 | 2.047799 | 0.001483 | 0.011334 | KCNH1 |
| ENSG0000 | -1.74754 | -3.35787 | 1.31E-13 | 1.99E-11 | ANKRD53 |
| ENSG0000 | -1.68307 | -3.2111 | 6.64E-17 | 1.98E-14 | SCN1A |
| ENSG0000 | -1.78334 | -3.44223 | 3.97E-07 | 1.11E-05 | DDIT4L |
| ENSG0000 | -1.01374 | -2.01913 | 0.001659 | 0.012343 | RNF175 |
| ENSG0000 | -1.31572 | -2.48926 | 5.07E-08 | 1.88E-06 | SLC25A48 |
| ENSG0000 | -1.34752 | -2.54474 | 0.000206 | 0.002267 | SLC22A3 |
| ENSG0000 | 1.028834 | 2.040374 | 1.69E-21 | 1.54E-18 | CDKN2B |
| ENSG0000 | -2.11475 | -4.33115 | 0.001231 | 0.00978 | NKX6-2 |
| ENSG0000 | 1.059155 | 2.08371 | 2.19E-13 | 3.15E-11 | ADAM12 |
| ENSG0000 | 1.235243 | 2.35421 | 0.000102 | 0.001254 | P4HA3 |
| ENSG0000 | -1.00083 | -2.00115 | 4.77E-08 | 1.78E-06 | ADAM33 |
| ENSG0000 | 1.113192 | 2.163237 | 2.77E-05 | 0.000424 | TAGLN |
| ENSG0000 | -1.31465 | -2.48742 | 8.30E-05 | 0.001061 | KIAA1755 |
| ENSG0000 | -1.42926 | -2.69308 | 0.009867 | 0.048844 | DOC2A |
| ENSG0000 | 1.202326 | 2.301103 | 5.71E-07 | 1.54E-05 | IL18 |
| ENSG0000 | -1.32624 | -2.50749 | 0.000101 | 0.001248 | CAPSL |

| | | | | | |
|---|---|---|---|---|---|
| ENSG0000 | -1.0317 | -2.04443 | 9.21E-17 | 2.70E-14 | DYNLT5 |
| ENSG0000 | -1.36236 | -2.57105 | 0.002345 | 0.016144 | NRSN1 |
| ENSG0000 | -1.34479 | -2.53993 | 0.005164 | 0.029667 | BANK1 |
| ENSG0000 | -1.20208 | -2.30072 | 0.004405 | 0.026216 | FAM81B |
| ENSG0000 | -1.46933 | -2.76893 | 3.50E-21 | 3.02E-18 | CHST9 |
| ENSG0000 | -1.26667 | -2.40605 | 8.04E-16 | 2.11E-13 | DNAAF1 |
| ENSG0000 | -1.04572 | -2.0644 | 2.34E-11 | 2.14E-09 | PITPNC1 |
| ENSG0000 | -1.21489 | -2.32123 | 2.23E-13 | 3.19E-11 | USP43 |
| ENSG0000 | -1.01438 | -2.02004 | 0.00401 | 0.024367 | RSPH10B |
| ENSG0000 | -1.09768 | -2.1401 | 1.50E-11 | 1.44E-09 | PTPRN2 |
| ENSG0000 | -1.54832 | -2.92476 | 5.55E-10 | 3.75E-08 | SPAG17 |
| ENSG0000 | -1.24819 | -2.37543 | 9.05E-10 | 5.63E-08 | CFAP70 |
| ENSG0000 | -1.78441 | -3.44477 | 5.02E-07 | 1.37E-05 | CFAP161 |
| ENSG0000 | -1.04869 | -2.06864 | 2.58E-18 | 1.15E-15 | FBXO32 |
| ENSG0000 | 1.116232 | 2.167801 | 4.59E-13 | 5.83E-11 | NRG1 |
| ENSG0000 | -1.45772 | -2.74674 | 7.51E-10 | 4.82E-08 | GRHL3 |
| ENSG0000 | -1.74143 | -3.34365 | 2.00E-14 | 3.83E-12 | LRRC43 |
| ENSG0000 | -1.03643 | -2.05115 | 0.007455 | 0.039315 | XDH |
| ENSG0000 | -1.12402 | -2.17954 | 0.003534 | 0.022085 | H2BC5 |
| ENSG0000 | -1.04234 | -2.05956 | 0.000113 | 0.001369 | CATIP |
| ENSG0000 | -1.39576 | -2.63127 | 3.78E-05 | 0.000549 | VWA5B1 |
| ENSG0000 | -1.00875 | -2.01217 | 1.02E-11 | 1.01E-09 | None |
| ENSG0000 | -1.42215 | -2.67984 | 8.54E-23 | 1.13E-19 | DRC7 |
| ENSG0000 | -1.37294 | -2.58998 | 1.66E-15 | 4.04E-13 | TPPP3 |
| ENSG0000 | 1.002789 | 2.003871 | 1.76E-09 | 1.01E-07 | ABCG1 |
| ENSG0000 | 1.058909 | 2.083355 | 0.001467 | 0.011246 | GAB3 |
| ENSG0000 | 1.432644 | 2.699409 | 0.000307 | 0.003174 | ICOSLG |
| ENSG0000 | -1.18021 | -2.2661 | 6.70E-22 | 7.24E-19 | CFAP157 |
| ENSG0000 | -1.80332 | -3.49023 | 2.29E-07 | 7.05E-06 | LRRC71 |
| ENSG0000 | -1.04614 | -2.065 | 5.41E-05 | 0.00074 | LRRC56 |
| ENSG0000 | -1.72294 | -3.30108 | 0.000407 | 0.00399 | AQP5 |
| ENSG0000 | -1.7264 | -3.30902 | 1.90E-08 | 8.19E-07 | UBXN10 |
| ENSG0000 | 1.031216 | 2.043746 | 8.93E-12 | 8.98E-10 | ATF3 |
| ENSG0000 | 1.024646 | 2.03446 | 9.53E-06 | 0.000173 | KCNF1 |
| ENSG0000 | -3.4916 | -11.248 | 3.92E-05 | 0.000566 | GABRG1 |
| ENSG0000 | -1.01689 | -2.02356 | 0.003536 | 0.022091 | DCST2 |
| ENSG0000 | -1.20839 | -2.3108 | 1.00E-05 | 0.000181 | EFHB |
| ENSG0000 | 1.133951 | 2.19459 | 8.00E-05 | 0.00103 | CDCP1 |
| ENSG0000 | -1.66098 | -3.16231 | 1.15E-19 | 6.42E-17 | CFAP100 |
| ENSG0000 | -1.09918 | -2.14233 | 1.71E-09 | 9.89E-08 | ZNF474 |
| ENSG0000 | -1.15771 | -2.23104 | 3.03E-05 | 0.000457 | PI16 |
| ENSG0000 | -1.10818 | -2.15573 | 4.86E-09 | 2.41E-07 | SYTL3 |
| ENSG0000 | 1.274639 | 2.419383 | 1.42E-12 | 1.62E-10 | DLC1 |
| ENSG0000 | -1.05822 | -2.08236 | 1.34E-08 | 5.95E-07 | HEPACAM |
| ENSG0000 | -1.03603 | -2.05058 | 0.003318 | 0.021062 | PDZRN4 |
| ENSG0000 | -1.09665 | -2.13858 | 0.003513 | 0.02199 | TMEM130 |
| ENSG0000 | -1.61527 | -3.06369 | 6.03E-07 | 1.61E-05 | A2ML1 |

| | | | | | |
|---|---|---|---|---|---|
| ENSG0000 | -1.19516 | -2.2897 | 1.02E-15 | 2.59E-13 | CFAP52 |
| ENSG0000 | 1.200072 | 2.297511 | 0.002238 | 0.015574 | MMP10 |
| ENSG0000 | -1.66419 | -3.16936 | 4.07E-07 | 1.14E-05 | SGSM1 |
| ENSG0000 | -1.29842 | -2.4596 | 2.88E-19 | 1.46E-16 | DNAAF3 |
| ENSG0000 | -2.17155 | -4.50506 | 3.97E-08 | 1.52E-06 | ANGPTL4 |
| ENSG0000 | -1.65311 | -3.14511 | 6.51E-18 | 2.62E-15 | TEKT1 |
| ENSG0000 | -1.51172 | -2.8515 | 0.004623 | 0.027212 | SCNN1B |
| ENSG0000 | -1.04928 | -2.0695 | 2.05E-05 | 0.000328 | LGI3 |
| ENSG0000 | -1.05931 | -2.08394 | 1.16E-05 | 0.000205 | VWA3B |
| ENSG0000 | 1.322673 | 2.501292 | 1.10E-12 | 1.28E-10 | STXBP6 |
| ENSG0000 | 1.180722 | 2.266902 | 5.57E-07 | 1.51E-05 | VXN |
| ENSG0000 | 1.082118 | 2.117142 | 8.33E-12 | 8.42E-10 | EFNA1 |
| ENSG0000 | -1.40689 | -2.65165 | 0.007599 | 0.039902 | RSPH10B2 |
| ENSG0000 | 2.035768 | 4.10041 | 0.002274 | 0.015767 | CXCL8 |
| ENSG0000 | -1.49365 | -2.816 | 1.51E-06 | 3.54E-05 | FRMPD2 |
| ENSG0000 | 1.141278 | 2.205763 | 5.84E-13 | 7.21E-11 | FOS |
| ENSG0000 | 1.128634 | 2.186516 | 1.38E-12 | 1.57E-10 | METTL7B |
| ENSG0000 | -1.06611 | -2.09378 | 1.19E-07 | 3.92E-06 | DCDC1 |
| ENSG0000 | -1.07725 | -2.11001 | 5.40E-11 | 4.64E-09 | GRIK1 |
| ENSG0000 | 1.260301 | 2.395457 | 5.18E-17 | 1.63E-14 | JUNB |
| ENSG0000 | -1.70337 | -3.25661 | 0.001722 | 0.012704 | DNAI2 |
| ENSG0000 | -1.05944 | -2.08412 | 3.80E-09 | 1.95E-07 | LRRC34 |
| ENSG0000 | -1.79259 | -3.46437 | 0.000599 | 0.005454 | IL16 |
| ENSG0000 | -1.14342 | -2.20905 | 4.04E-24 | 6.98E-21 | EFCAB12 |
| ENSG0000 | -1.41945 | -2.67484 | 0.000458 | 0.004403 | HPSE2 |
| ENSG0000 | -1.18981 | -2.28123 | 7.72E-05 | 0.001 | ZNF483 |
| ENSG0000 | -6.12302 | -69.6968 | 0.00758 | 0.039834 | SAA1 |
| ENSG0000 | -1.44718 | -2.72674 | 4.86E-06 | 9.64E-05 | FAM166C |
| ENSG0000 | -1.32756 | -2.50977 | 1.09E-05 | 0.000193 | CCDC13-AS1 |
| ENSG0000 | -1.77446 | -3.4211 | 1.45E-06 | 3.44E-05 | SNX31 |
| ENSG0000 | 1.396293 | 2.632244 | 0.000335 | 0.003389 | CHRNA9 |
| ENSG0000 | 1.622283 | 3.078618 | 2.35E-06 | 5.19E-05 | CMKLR1 |
| ENSG0000 | -1.55086 | -2.92992 | 0.002149 | 0.015096 | DNAH12 |
| ENSG0000 | 1.129631 | 2.188027 | 4.63E-05 | 0.00065 | ASPHD1 |
| ENSG0000 | -1.21151 | -2.3158 | 2.12E-09 | 1.18E-07 | FAM182B |
| ENSG0000 | -1.43846 | -2.71032 | 2.88E-20 | 1.78E-17 | VWA3A |
| ENSG0000 | -1.00497 | -2.00691 | 0.00285 | 0.018767 | GRAMD2A |
| ENSG0000 | 1.277117 | 2.423542 | 1.03E-28 | 3.55E-25 | LPL |
| ENSG0000 | 1.457078 | 2.745517 | 5.52E-17 | 1.70E-14 | CLCF1 |
| ENSG0000 | 1.893023 | 3.714125 | 8.69E-18 | 3.34E-15 | FOSL1 |
| ENSG0000 | -1.20487 | -2.30517 | 0.000567 | 0.005215 | TEX26 |
| ENSG0000 | -1.39188 | -2.62421 | 0.001537 | 0.01162 | LINC01106 |
| ENSG0000 | -1.06058 | -2.08576 | 0.005449 | 0.030996 | PLEKHD1 |
| ENSG0000 | 1.159949 | 2.234495 | 0.005915 | 0.033064 | SPHK1 |
| ENSG0000 | -1.37656 | -2.59649 | 1.77E-09 | 1.02E-07 | PRR18 |
| ENSG0000 | -1.43991 | -2.71303 | 4.56E-25 | 8.75E-22 | MAP3K19 |
| ENSG0000 | 1.04107 | 2.057753 | 2.42E-14 | 4.54E-12 | MYO1D |

| | | | | | |
|---|---|---|---|---|---|
| ENSG0000 | 1.380439 | 2.603476 | 1.54E-11 | 1.46E-09 | BDNF |
| ENSG0000 | 1.039736 | 2.055851 | 0.000131 | 0.001559 | GCNT4 |
| ENSG0000 | -1.28973 | -2.44482 | 8.13E-20 | 4.85E-17 | FIBIN |
| ENSG0000 | -1.81585 | -3.52068 | 1.87E-14 | 3.63E-12 | ODF3B |
| ENSG0000 | -1.38962 | -2.6201 | 1.75E-13 | 2.59E-11 | FAR2P2 |
| ENSG0000 | -1.08119 | -2.11578 | 8.34E-08 | 2.91E-06 | ERICH3 |
| ENSG0000 | -1.34755 | -2.54479 | 0.00726 | 0.038554 | DAND5 |
| ENSG0000 | 1.594231 | 3.019334 | 2.64E-25 | 5.69E-22 | EGR3 |
| ENSG0000 | -1.00589 | -2.00818 | 2.88E-14 | 5.24E-12 | SLC17A8 |
| ENSG0000 | -1.03523 | -2.04944 | 0.008579 | 0.043848 | PCED1B |
| ENSG0000 | 1.873076 | 3.663128 | 0.003472 | 0.021793 | C3orf80 |
| ENSG0000 | -1.12615 | -2.18276 | 3.87E-06 | 7.97E-05 | KCNE1 |
| ENSG0000 | -1.13966 | -2.2033 | 1.06E-05 | 0.000189 | TSPYL5 |
| ENSG0000 | 1.052581 | 2.074238 | 6.02E-13 | 7.38E-11 | ARSJ |
| ENSG0000 | -1.56301 | -2.95469 | 1.41E-14 | 2.87E-12 | FAM83H |
| ENSG0000 | -1.37639 | -2.59618 | 2.44E-14 | 4.54E-12 | NME9 |
| ENSG0000 | -1.35314 | -2.55467 | 1.54E-10 | 1.19E-08 | CFAP65 |
| ENSG0000 | -2.88541 | -7.38917 | 3.43E-06 | 7.18E-05 | ATG9B |
| ENSG0000 | 1.10302 | 2.148038 | 1.33E-08 | 5.95E-07 | GPR3 |
| ENSG0000 | -1.02129 | -2.02973 | 2.26E-07 | 6.98E-06 | HHIPL1 |
| ENSG0000 | -1.19963 | -2.2968 | 2.34E-05 | 0.000367 | KIAA2012 |
| ENSG0000 | -1.86039 | -3.63105 | 0.006969 | 0.037447 | NXPH3 |
| ENSG0000 | -1.23112 | -2.34749 | 0.002023 | 0.014394 | CSMD1 |
| ENSG0000 | -1.07058 | -2.10028 | 4.25E-08 | 1.61E-06 | CABCOCO1 |
| ENSG0000 | 1.12602 | 2.182559 | 0.000161 | 0.001838 | TNFAIP8L3 |
| ENSG0000 | -1.78144 | -3.4377 | 2.00E-05 | 0.000321 | HOATZ |
| ENSG0000 | -1.36576 | -2.57712 | 5.66E-10 | 3.79E-08 | LRRC55 |
| ENSG0000 | -1.28976 | -2.44487 | 0.000222 | 0.002413 | DNAH2 |
| ENSG0000 | -1.29066 | -2.4464 | 8.71E-06 | 0.000161 | SDR42E2 |
| ENSG0000 | -2.19749 | -4.58681 | 1.19E-26 | 2.93E-23 | CLDN5 |
| ENSG0000 | -1.04959 | -2.06994 | 3.70E-09 | 1.92E-07 | KCNQ3 |
| ENSG0000 | 1.282488 | 2.432581 | 4.35E-12 | 4.61E-10 | MAFF |
| ENSG0000 | -1.15118 | -2.22096 | 3.84E-07 | 1.08E-05 | EFCAB10 |
| ENSG0000 | -1.11779 | -2.17014 | 8.44E-08 | 2.93E-06 | MORN5 |
| ENSG0000 | 1.198987 | 2.295784 | 9.65E-07 | 2.43E-05 | MYBL1 |
| ENSG0000 | -1.82219 | -3.53619 | 3.39E-14 | 5.98E-12 | KLHL32 |
| ENSG0000 | -1.75555 | -3.37655 | 1.39E-06 | 3.30E-05 | LINC00643 |
| ENSG0000 | -1.1092 | -2.15727 | 9.30E-06 | 0.00017 | PDE2A |
| ENSG0000 | -1.49119 | -2.81121 | 5.97E-19 | 2.95E-16 | CFAP73 |
| ENSG0000 | 1.341858 | 2.534776 | 2.94E-14 | 5.29E-12 | SPRY4 |
| ENSG0000 | -1.45454 | -2.7407 | 3.49E-05 | 0.000512 | DNAJB13 |
| ENSG0000 | -1.49817 | -2.82485 | 1.76E-15 | 4.22E-13 | DIPK1C |
| ENSG0000 | -1.31057 | -2.4804 | 0.000564 | 0.005191 | LRRC74B |
| ENSG0000 | -1.4027 | -2.64396 | 0.000141 | 0.001647 | SNHG28 |
| ENSG0000 | 1.272116 | 2.415156 | 4.05E-16 | 1.09E-13 | SPRED3 |
| ENSG0000 | -1.32826 | -2.51099 | 0.000965 | 0.008068 | PRELP |
| ENSG0000 | -1.41755 | -2.67131 | 1.01E-06 | 2.53E-05 | HACD4 |

| | | | | | |
|---|---|---|---|---|---|
| ENSG0000 | 1.666634 | 3.174731 | 7.44E-05 | 0.000971 | SH2D5 |
| ENSG0000 | -2.26042 | -4.79132 | 1.11E-08 | 5.05E-07 | KIF19 |
| ENSG0000 | -1.05398 | -2.07625 | 0.008564 | 0.043787 | SEMA4A |
| ENSG0000 | -1.2884 | -2.44258 | 1.56E-06 | 3.64E-05 | ARL9 |
| ENSG0000 | 1.024183 | 2.033807 | 4.42E-09 | 2.21E-07 | HRH1 |
| ENSG0000 | -1.02625 | -2.03672 | 0.000565 | 0.005201 | DTHD1 |
| ENSG0000 | -1.06219 | -2.0881 | 2.62E-13 | 3.65E-11 | CFAP43 |
| ENSG0000 | -1.51379 | -2.85559 | 0.000876 | 0.007477 | CFAP299 |
| ENSG0000 | -1.90617 | -3.74813 | 0.004889 | 0.028408 | HES5 |
| ENSG0000 | -1.3361 | -2.52467 | 0.002559 | 0.017233 | LEKR1 |
| ENSG0000 | 2.377378 | 5.195917 | 0.000228 | 0.002472 | PLN |
| ENSG0000 | 1.9691 | 3.915238 | 7.46E-30 | 6.45E-26 | ARC |
| ENSG0000 | -1.23841 | -2.35938 | 2.82E-06 | 6.07E-05 | MMP17 |
| ENSG0000 | -1.09401 | -2.13466 | 2.18E-11 | 2.03E-09 | DLL1 |
| ENSG0000 | -1.39496 | -2.62981 | 1.75E-05 | 0.000287 | SMOC1 |
| ENSG0000 | -1.80789 | -3.50131 | 0.000567 | 0.00521 | Y_RNA |
| ENSG0000 | -1.12416 | -2.17974 | 0.007054 | 0.037765 | TDRKH-AS1 |
| ENSG0000 | -1.40555 | -2.64919 | 1.87E-07 | 5.92E-06 | IQANK1 |
| ENSG0000 | -1.11584 | -2.16721 | 3.19E-07 | 9.19E-06 | PRRT1 |
| ENSG0000 | -1.62404 | -3.08237 | 0.003372 | 0.021338 | ERICH2 |
| ENSG0000 | -1.24038 | -2.36261 | 7.03E-12 | 7.19E-10 | CYP21A1P |
| ENSG0000 | -1.28951 | -2.44445 | 2.91E-06 | 6.24E-05 | BTBD17 |
| ENSG0000 | -1.531 | -2.88987 | 8.62E-08 | 2.98E-06 | LRRC10B |
| ENSG0000 | -1.60031 | -3.03208 | 0.000417 | 0.004069 | MT1M |
| ENSG0000 | -1.27981 | -2.42806 | 0.000376 | 0.003735 | GMNC |
| ENSG0000 | -1.01351 | -2.01882 | 3.89E-09 | 1.99E-07 | NYNRIN |
| ENSG0000 | -1.37763 | -2.59841 | 1.42E-17 | 5.14E-15 | CFAP99 |
| ENSG0000 | -1.03489 | -2.04896 | 0.00515 | 0.029646 | MIR149 |
| ENSG0000 | -1.17872 | -2.26376 | 0.008354 | 0.04299 | None |
| ENSG0000 | -1.33208 | -2.51765 | 1.45E-30 | 2.50E-26 | CFAP45 |
| ENSG0000 | -1.55801 | -2.94448 | 7.17E-08 | 2.56E-06 | C10orf105 |
| ENSG0000 | -1.05165 | -2.0729 | 0.000186 | 0.002085 | COL28A1 |
| ENSG0000 | -1.0904 | -2.12933 | 4.71E-13 | 5.94E-11 | C5orf49 |
| ENSG0000 | -1.42829 | -2.69128 | 8.64E-06 | 0.00016 | SIAH3 |
| ENSG0000 | -1.046 | -2.06479 | 0.000196 | 0.002174 | None |
| ENSG0000 | -1.78061 | -3.43571 | 0.002356 | 0.016194 | LINC01422 |
| ENSG0000 | -1.16991 | -2.24998 | 0.00168 | 0.012456 | WARS2-IT1 |
| ENSG0000 | 1.586721 | 3.003658 | 1.65E-06 | 3.79E-05 | ZNF812P |
| ENSG0000 | -2.13348 | -4.38774 | 0.000722 | 0.006344 | LINC00092 |
| ENSG0000 | 1.125619 | 2.181951 | 0.008118 | 0.042 | None |
| ENSG0000 | 1.022212 | 2.031031 | 0.004251 | 0.025504 | None |
| ENSG0000 | -3.38223 | -10.4268 | 0.006056 | 0.033672 | ORM2 |
| ENSG0000 | -1.28695 | -2.44012 | 1.09E-09 | 6.64E-08 | None |
| ENSG0000 | -1.48816 | -2.8053 | 1.54E-20 | 9.89E-18 | PROB1 |
| ENSG0000 | -1.26414 | -2.40184 | 0.000377 | 0.003744 | FRMD3-AS1 |
| ENSG0000 | -1.26141 | -2.39729 | 0.004763 | 0.027871 | None |
| ENSG0000 | -4.48299 | -22.3621 | 0.003018 | 0.019614 | ORM1 |

| | | | | | |
|---|---|---|---|---|---|
| ENSG0000 | 1.721197 | 3.297098 | 0.000144 | 0.001674 | MYOSLID |
| ENSG0000 | -1.51193 | -2.85191 | 0.005751 | 0.032381 | None |
| ENSG0000 | -1.19783 | -2.29395 | 3.46E-08 | 1.36E-06 | CYP21A2 |
| ENSG0000 | -1.01647 | -2.02296 | 9.03E-07 | 2.29E-05 | TGFB2-AS1 |
| ENSG0000 | -1.72416 | -3.30387 | 0.006179 | 0.034156 | TMEM229A |
| ENSG0000 | -1.77025 | -3.41113 | 0.001731 | 0.012759 | OBI1-AS1 |
| ENSG0000 | -1.21601 | -2.32303 | 3.56E-05 | 0.00052 | DAAM2-AS1 |
| ENSG0000 | -1.3219 | -2.49995 | 0.007251 | 0.03852 | DNM1P51 |
| ENSG0000 | -1.19805 | -2.2943 | 0.001529 | 0.011587 | DPY19L2P4 |
| ENSG0000 | -1.18143 | -2.26801 | 2.89E-07 | 8.46E-06 | DOCK7-DT |
| ENSG0000 | -1.14728 | -2.21497 | 0.00684 | 0.036917 | None |
| ENSG0000 | -2.13596 | -4.3953 | 3.05E-05 | 0.000458 | None |
| ENSG0000 | -1.70522 | -3.26078 | 0.009402 | 0.04712 | None |
| ENSG0000 | -1.30922 | -2.47807 | 0.000245 | 0.002634 | CT75 |
| ENSG0000 | -1.66714 | -3.17584 | 0.000121 | 0.001442 | LRIG2-DT |
| ENSG0000 | 1.488611 | 2.806186 | 2.41E-05 | 0.000377 | LINC00707 |
| ENSG0000 | -1.40947 | -2.6564 | 0.000929 | 0.007815 | RPS27P25 |
| ENSG0000 | -1.51882 | -2.86557 | 0.000112 | 0.001356 | None |
| ENSG0000 | -1.099 | -2.14205 | 0.000522 | 0.004881 | SMKR1 |
| ENSG0000 | -1.68006 | -3.20441 | 0.000465 | 0.004463 | VSIG8 |
| ENSG0000 | 1.391119 | 2.62282 | 2.10E-07 | 6.53E-06 | |
| ENSG0000 | 1.323907 | 2.503432 | 1.61E-06 | 3.72E-05 | GSTA1 |
| ENSG0000 | -1.05751 | -2.08134 | 1.07E-14 | 2.21E-12 | CCDC13 |
| ENSG0000 | -1.00065 | -2.0009 | 1.65E-05 | 0.000273 | USP2-AS1 |
| ENSG0000 | -1.08687 | -2.12412 | 0.002295 | 0.015887 | None |
| ENSG0000 | -1.25072 | -2.3796 | 1.86E-07 | 5.92E-06 | TNXA |
| ENSG0000 | -1.32723 | -2.5092 | 4.17E-06 | 8.46E-05 | None |
| ENSG0000 | -1.04824 | -2.06801 | 0.004166 | 0.02506 | ALG1L9P |
| ENSG0000 | -1.05083 | -2.07172 | 3.45E-05 | 0.000505 | C8orf34-AS1 |
| ENSG0000 | -1.01494 | -2.02082 | 0.00242 | 0.016508 | TICAM2-AS1 |
| ENSG0000 | -1.3026 | -2.46673 | 5.78E-05 | 0.000784 | None |
| ENSG0000 | -1.22253 | -2.33355 | 0.004037 | 0.024488 | ZNF474-AS1 |
| ENSG0000 | -1.06969 | -2.09899 | 1.51E-05 | 0.000254 | None |
| ENSG0000 | -1.18782 | -2.27808 | 0.004252 | 0.025504 | None |
| ENSG0000 | -1.20792 | -2.31005 | 0.000455 | 0.004381 | STK19B |
| ENSG0000 | -1.25861 | -2.39264 | 0.003101 | 0.020034 | KBTBD11-AS1 |
| ENSG0000 | 1.091105 | 2.130371 | 0.001016 | 0.008396 | LINC02732 |
| ENSG0000 | -1.92966 | -3.80966 | 0.002475 | 0.016799 | RASSF10-DT |
| ENSG0000 | 1.080583 | 2.11489 | 1.22E-06 | 2.96E-05 | None |
| ENSG0000 | -1.39745 | -2.63435 | 0.002541 | 0.017139 | B3GAT1-DT |
| ENSG0000 | -1.00758 | -2.01054 | 4.56E-08 | 1.71E-06 | TRIL |
| ENSG0000 | -1.7832 | -3.44188 | 3.23E-05 | 0.00048 | None |
| ENSG0000 | 1.733677 | 3.325743 | 0.003541 | 0.022109 | PPP1R14B-AS1 |
| ENSG0000 | -2.92247 | -7.58144 | 0.007748 | 0.040531 | HP |
| ENSG0000 | 1.833678 | 3.564447 | 0.001321 | 0.010354 | None |
| ENSG0000 | -1.95477 | -3.87653 | 0.000117 | 0.001405 | None |
| ENSG0000 | -1.01303 | -2.01815 | 8.61E-07 | 2.21E-05 | LINC01579 |

| | | | | | |
|---|---|---|---|---|---|
| ENSG0000 | -3.11403 | -8.65798 | 2.14E-06 | 4.77E-05 | None |
| ENSG0000 | -1.36575 | -2.57711 | 0.000627 | 0.005662 | LINC02352 |
| ENSG0000 | -1.17452 | -2.25717 | 0.00154 | 0.011631 | None |
| ENSG0000 | -1.08213 | -2.11716 | 4.34E-06 | 8.74E-05 | None |
| ENSG0000 | -1.148 | -2.21606 | 1.72E-05 | 0.000284 | None |
| ENSG0000 | -1.16661 | -2.24484 | 0.000538 | 0.004999 | None |
| ENSG0000 | -1.00328 | -2.00455 | 8.64E-05 | 0.0011 | TSPAN5-DT |
| ENSG0000 | -1.82339 | -3.53912 | 0.000317 | 0.003252 | |
| ENSG0000 | -1.5124 | -2.85284 | 0.001573 | 0.011847 | LINC02175 |
| ENSG0000 | -1.4398 | -2.71283 | 4.32E-07 | 1.20E-05 | SPON1 |
| ENSG0000 | -1.31678 | -2.4911 | 0.005861 | 0.032853 | None |
| ENSG0000 | -1.10966 | -2.15795 | 0.002266 | 0.015729 | TMEM220-AS1 |
| ENSG0000 | -1.1106 | -2.15935 | 0.008058 | 0.041727 | AGAP12P |
| ENSG0000 | 1.678717 | 3.201432 | 3.12E-06 | 6.60E-05 | GJA5 |
| ENSG0000 | -1.21162 | -2.31597 | 9.15E-21 | 6.88E-18 | MYO15B |
| ENSG0000 | -1.4933 | -2.81533 | 1.91E-10 | 1.41E-08 | CCDC177 |
| ENSG0000 | -1.53145 | -2.89076 | 0.00025 | 0.002678 | None |
| ENSG0000 | -1.81028 | -3.5071 | 0.001445 | 0.011098 | MEI4 |
| ENSG0000 | 1.039005 | 2.05481 | 8.40E-06 | 0.000156 | None |
| ENSG0000 | -1.69882 | -3.24636 | 0.000211 | 0.00231 | GAS2L2 |
| ENSG0000 | -1.32705 | -2.50889 | 0.002212 | 0.015428 | None |
| ENSG0000 | -1.50578 | -2.83978 | 0.002123 | 0.014957 | LINC01607 |
| ENSG0000 | -1.4838 | -2.79685 | 0.000316 | 0.003249 | None |
| ENSG0000 | -1.05698 | -2.08057 | 9.61E-07 | 2.43E-05 | MESTIT1 |
| ENSG0000 | -1.26018 | -2.39526 | 0.009844 | 0.048756 | None |
| ENSG0000 | -1.26575 | -2.40452 | 6.70E-07 | 1.77E-05 | ADIRF-AS1 |
| ENSG0000 | -1.1859 | -2.27506 | 0.002359 | 0.016202 | None |
| ENSG0000 | -1.48001 | -2.78951 | 0.003119 | 0.020112 | LENG9 |
| ENSG0000 | -1.16426 | -2.24118 | 1.10E-06 | 2.73E-05 | FLJ16779 |
| ENSG0000 | -1.4376 | -2.7087 | 3.05E-14 | 5.44E-12 | None |
| ENSG0000 | 2.270464 | 4.824784 | 0.000171 | 0.001936 | None |
| ENSG0000 | -1.99948 | -3.99856 | 1.19E-07 | 3.92E-06 | None |
| ENSG0000 | -1.17813 | -2.26283 | 0.007941 | 0.041261 | None |
| ENSG0000 | 1.158074 | 2.231594 | 1.36E-09 | 8.06E-08 | None |

| GeneID | Treatment | Treatment | Treatment | Treatment | name |
|---|---|---|---|---|---|
| ENSG0000 | 1.125644 | 2.181989 | 0.004058 | 0.034537 | MYH16 |
| ENSG0000 | -1.38298 | -2.60806 | 3.90E-20 | 9.86E-18 | CD38 |
| ENSG0000 | 1.13678 | 2.198896 | 1.48E-14 | 1.95E-12 | TMEM132A |
| ENSG0000 | 2.489453 | 5.61565 | 3.11E-11 | 2.45E-09 | CACNA1G |
| ENSG0000 | -1.12324 | -2.17836 | 1.73E-38 | 2.91E-35 | SCIN |
| ENSG0000 | 1.636226 | 3.108516 | 6.11E-10 | 3.88E-08 | SYN1 |
| ENSG0000 | 1.747656 | 3.358125 | 1.53E-05 | 0.000349 | SLC6A13 |
| ENSG0000 | -1.26716 | -2.40688 | 1.23E-06 | 3.85E-05 | GABRA3 |
| ENSG0000 | -1.91755 | -3.7778 | 3.98E-08 | 1.77E-06 | IGF1 |
| ENSG0000 | 1.014215 | 2.019803 | 7.04E-17 | 1.23E-14 | SLC38A5 |
| ENSG0000 | 1.387331 | 2.615943 | 4.55E-09 | 2.46E-07 | ATP1A2 |
| ENSG0000 | -2.11832 | -4.34187 | 0.00552 | 0.043924 | HGF |
| ENSG0000 | 1.117641 | 2.169919 | 7.82E-21 | 2.22E-18 | EHD2 |
| ENSG0000 | 6.281693 | 77.7997 | 6.72E-05 | 0.001251 | INSRR |
| ENSG0000 | 1.245699 | 2.371334 | 3.41E-23 | 1.20E-20 | ANK1 |
| ENSG0000 | 1.661367 | 3.16316 | 6.29E-16 | 1.00E-13 | TNC |
| ENSG0000 | 4.542028 | 23.29628 | 0.003146 | 0.028565 | DKK3 |
| ENSG0000 | 1.327379 | 2.509464 | 5.26E-32 | 4.70E-29 | PTPRN |
| ENSG0000 | 1.86293 | 3.637458 | 0.005903 | 0.046009 | KCNH2 |
| ENSG0000 | 3.759623 | 13.54439 | 0.004108 | 0.034855 | NGFR |
| ENSG0000 | 1.228531 | 2.343283 | 1.22E-07 | 4.79E-06 | HIPK2 |
| ENSG0000 | 1.14839 | 2.216663 | 2.79E-05 | 0.000582 | SLC9A3R2 |
| ENSG0000 | 1.607046 | 3.046274 | 0.005359 | 0.042853 | GLP2R |
| ENSG0000 | 1.372382 | 2.588976 | 1.08E-11 | 9.25E-10 | EML1 |
| ENSG0000 | 1.146443 | 2.213674 | 8.66E-11 | 6.27E-09 | NAV3 |
| ENSG0000 | 6.480759 | 89.31057 | 0.000245 | 0.003663 | SCT |
| ENSG0000 | -1.09877 | -2.14172 | 3.27E-11 | 2.57E-09 | FCGR2B |
| ENSG0000 | -2.03906 | -4.10977 | 5.94E-15 | 8.17E-13 | IRAG1 |
| ENSG0000 | 5.250826 | 38.07643 | 0.001198 | 0.013441 | FNDC8 |
| ENSG0000 | 1.558149 | 2.944757 | 4.94E-17 | 8.78E-15 | FRY |
| ENSG0000 | 1.231685 | 2.348411 | 1.66E-18 | 3.46E-16 | NOTCH3 |
| ENSG0000 | 1.121298 | 2.175426 | 0.005243 | 0.042127 | SNCB |
| ENSG0000 | 1.459735 | 2.750578 | 7.15E-31 | 6.02E-28 | FOSL2 |
| ENSG0000 | 2.127986 | 4.37107 | 0.000959 | 0.011174 | FAP |
| ENSG0000 | 1.000805 | 2.001117 | 3.84E-23 | 1.34E-20 | SYNJ2 |
| ENSG0000 | 1.24554 | 2.371073 | 6.80E-21 | 1.94E-18 | COL5A3 |
| ENSG0000 | 2.240386 | 4.725233 | 0.000356 | 0.004969 | IGSF9B |
| ENSG0000 | 1.277448 | 2.424098 | 8.92E-05 | 0.00159 | CXCL2 |
| ENSG0000 | 1.683623 | 3.212336 | 0.005492 | 0.043711 | MECOM |
| ENSG0000 | -1.09185 | -2.13147 | 1.10E-06 | 3.48E-05 | HSD17B14 |
| ENSG0000 | -6.10172 | -68.6755 | 3.43E-06 | 9.48E-05 | CETP |
| ENSG0000 | 1.001676 | 2.002325 | 2.17E-13 | 2.42E-11 | TFAP2C |
| ENSG0000 | -4.02163 | -16.2416 | 0.003045 | 0.02777 | CHGB |
| ENSG0000 | 1.175082 | 2.258058 | 1.10E-25 | 5.00E-23 | P3H2 |
| ENSG0000 | 1.093827 | 2.134395 | 9.58E-14 | 1.12E-11 | PITPNM2 |
| ENSG0000 | -1.17347 | -2.25554 | 9.84E-12 | 8.53E-10 | RPGRIP1 |

| | | | | | |
|---|---|---|---|---|---|
| ENSG0000 | 4.704794 | 26.07859 | 7.14E-05 | 0.001317 | IL11 |
| ENSG0000 | 1.239511 | 2.361186 | 0.000489 | 0.006455 | SERPIND1 |
| ENSG0000 | 1.156799 | 2.229622 | 3.40E-09 | 1.88E-07 | CARD10 |
| ENSG0000 | 1.316834 | 2.491188 | 5.49E-13 | 5.78E-11 | CPNE6 |
| ENSG0000 | 1.108588 | 2.156345 | 0.002303 | 0.022363 | SLA2 |
| ENSG0000 | 1.492486 | 2.813734 | 1.82E-21 | 5.42E-19 | NTSR1 |
| ENSG0000 | 1.095749 | 2.13724 | 2.14E-05 | 0.000462 | EEF1A2 |
| ENSG0000 | -1.10328 | -2.14842 | 2.30E-15 | 3.37E-13 | ISM1 |
| ENSG0000 | -1.06772 | -2.09611 | 1.18E-24 | 4.75E-22 | MYL9 |
| ENSG0000 | 1.758083 | 3.382483 | 0.004851 | 0.039825 | WFDC2 |
| ENSG0000 | 5.581848 | 47.89648 | 0.002168 | 0.021315 | RS1 |
| ENSG0000 | 1.139764 | 2.20345 | 7.15E-08 | 2.96E-06 | GABRE |
| ENSG0000 | 1.088629 | 2.126718 | 7.48E-10 | 4.66E-08 | SRPX2 |
| ENSG0000 | 1.116063 | 2.167546 | 2.37E-05 | 0.000503 | FOXF1 |
| ENSG0000 | 3.196012 | 9.164221 | 2.63E-14 | 3.35E-12 | CORO2B |
| ENSG0000 | 2.198019 | 4.588488 | 2.49E-08 | 1.15E-06 | CCN4 |
| ENSG0000 | 3.844497 | 14.36511 | 0.005536 | 0.043997 | CGB3 |
| ENSG0000 | -1.26604 | -2.405 | 4.22E-05 | 0.000831 | RETN |
| ENSG0000 | -5.38592 | -41.8143 | 0.001105 | 0.012578 | SLC1A6 |
| ENSG0000 | -3.00438 | -8.02432 | 1.20E-07 | 4.74E-06 | SIGLEC8 |
| ENSG0000 | -1.00496 | -2.00688 | 2.38E-08 | 1.11E-06 | LILRB5 |
| ENSG0000 | -1.14687 | -2.21433 | 0.005228 | 0.042056 | PDE4C |
| ENSG0000 | 2.368917 | 5.165532 | 5.92E-10 | 3.77E-08 | TFPI2 |
| ENSG0000 | 1.007348 | 2.010212 | 0.00511 | 0.041392 | CAV2 |
| ENSG0000 | 1.524544 | 2.876957 | 5.21E-25 | 2.16E-22 | CAV1 |
| ENSG0000 | 1.054567 | 2.077095 | 9.41E-11 | 6.75E-09 | MET |
| ENSG0000 | 1.033596 | 2.04712 | 3.94E-22 | 1.25E-19 | STX1A |
| ENSG0000 | 1.58888 | 3.008157 | 0.000814 | 0.0098 | CHCHD2 |
| ENSG0000 | 1.039853 | 2.056018 | 2.92E-06 | 8.28E-05 | CCL24 |
| ENSG0000 | 1.088601 | 2.126678 | 2.79E-35 | 3.47E-32 | SERPINE1 |
| ENSG0000 | 1.657169 | 3.153969 | 3.91E-05 | 0.000778 | AEBP1 |
| ENSG0000 | -1.1685 | -2.24778 | 0.006434 | 0.049172 | TNFSF8 |
| ENSG0000 | 2.354822 | 5.115311 | 0.003946 | 0.033816 | KCNT1 |
| ENSG0000 | 1.603478 | 3.03875 | 2.47E-13 | 2.73E-11 | GLIS3 |
| ENSG0000 | 1.584881 | 2.99983 | 2.97E-08 | 1.35E-06 | WNT3 |
| ENSG0000 | 1.74474 | 3.351345 | 3.66E-25 | 1.54E-22 | CCL7 |
| ENSG0000 | 1.473915 | 2.777746 | 0.000166 | 0.002667 | EFNB3 |
| ENSG0000 | 2.819125 | 7.057342 | 1.43E-18 | 3.00E-16 | AREG |
| ENSG0000 | -1.23076 | -2.34691 | 1.71E-06 | 5.18E-05 | TBC1D19 |
| ENSG0000 | -3.94714 | -15.4244 | 0.002668 | 0.02507 | CLNK |
| ENSG0000 | 6.269491 | 77.14446 | 0.000425 | 0.005761 | NKX3-2 |
| ENSG0000 | 2.087466 | 4.250009 | 0.006273 | 0.048129 | CRYAB |
| ENSG0000 | 5.589178 | 48.14045 | 0.000194 | 0.003011 | B3GAT1 |
| ENSG0000 | -1.60272 | -3.03715 | 7.62E-40 | 1.36E-36 | MS4A6A |
| ENSG0000 | -1.33655 | -2.52547 | 3.40E-18 | 6.89E-16 | MS4A4A |
| ENSG0000 | 1.704585 | 3.259351 | 2.62E-18 | 5.40E-16 | SLC1A2 |
| ENSG0000 | 1.578221 | 2.986015 | 2.82E-08 | 1.29E-06 | WNT5B |

| | | | | | |
|---|---|---|---|---|---|
| ENSG0000 | -1.59501 | -3.02097 | 0.0012 | 0.013459 | TREML2 |
| ENSG0000 | -4.88605 | -29.5698 | 0.004404 | 0.036889 | NR2E1 |
| ENSG0000 | -1.10127 | -2.14544 | 1.15E-06 | 3.63E-05 | ADGRG6 |
| ENSG0000 | -1.01722 | -2.02401 | 8.17E-13 | 8.29E-11 | LY86 |
| ENSG0000 | 1.250135 | 2.378636 | 2.82E-12 | 2.65E-10 | PDGFRB |
| ENSG0000 | 1.248063 | 2.375224 | 0.003709 | 0.032323 | AMOTL2 |
| ENSG0000 | 1.421708 | 2.679026 | 1.81E-13 | 2.05E-11 | COL7A1 |
| ENSG0000 | 1.50219 | 2.832724 | 0.002726 | 0.025542 | C3orf52 |
| ENSG0000 | -5.71818 | -52.6434 | 2.95E-05 | 0.000611 | IL18RAP |
| ENSG0000 | 2.106715 | 4.307095 | 0.002697 | 0.025331 | PRRX1 |
| ENSG0000 | -1.54804 | -2.92419 | 2.99E-13 | 3.25E-11 | OLFML3 |
| ENSG0000 | -1.1792 | -2.26451 | 1.67E-05 | 0.000375 | CD2 |
| ENSG0000 | 1.318333 | 2.493778 | 0.003249 | 0.029309 | ADGRL2 |
| ENSG0000 | -1.25028 | -2.37888 | 5.05E-14 | 6.14E-12 | GBP1 |
| ENSG0000 | 1.581375 | 2.99255 | 7.65E-11 | 5.63E-09 | MMP8 |
| ENSG0000 | -5.39274 | -42.0123 | 0.000235 | 0.003545 | ADGB |
| ENSG0000 | 3.613395 | 12.23884 | 0.00031 | 0.004428 | GRIA2 |
| ENSG0000 | -1.0056 | -2.00778 | 1.57E-16 | 2.64E-14 | CCDC170 |
| ENSG0000 | -1.0053 | -2.00736 | 1.11E-10 | 7.87E-09 | ACAT2 |
| ENSG0000 | -1.29517 | -2.45406 | 3.09E-07 | 1.12E-05 | KCNJ5 |
| ENSG0000 | -1.77473 | -3.42173 | 9.68E-08 | 3.90E-06 | GJB6 |
| ENSG0000 | 2.4295 | 5.387067 | 0.004273 | 0.035994 | ADGRB2 |
| ENSG0000 | 1.365105 | 2.57595 | 0.003312 | 0.029737 | TMEM54 |
| ENSG0000 | 1.168753 | 2.248174 | 4.56E-16 | 7.42E-14 | CXCR4 |
| ENSG0000 | -1.08457 | -2.12074 | 2.61E-17 | 4.87E-15 | XPNPEP2 |
| ENSG0000 | -1.64045 | -3.11763 | 3.87E-17 | 6.97E-15 | LRRC39 |
| ENSG0000 | 1.408428 | 2.654477 | 1.49E-27 | 8.71E-25 | INHBA |
| ENSG0000 | 3.109016 | 8.627939 | 4.71E-18 | 9.41E-16 | TWIST1 |
| ENSG0000 | 1.877795 | 3.67513 | 3.04E-11 | 2.41E-09 | CHST3 |
| ENSG0000 | 1.655258 | 3.149795 | 0.00015 | 0.002449 | PLP1 |
| ENSG0000 | 5.205392 | 36.89598 | 0.001061 | 0.012134 | PI3 |
| ENSG0000 | 1.947082 | 3.855939 | 0.003114 | 0.028293 | SLPI |
| ENSG0000 | 1.043443 | 2.061141 | 1.03E-24 | 4.20E-22 | PMEPA1 |
| ENSG0000 | -1.15012 | -2.21932 | 0.002282 | 0.02221 | BCAS4 |
| ENSG0000 | 1.347552 | 2.544799 | 0.001193 | 0.013393 | F13A1 |
| ENSG0000 | 2.782249 | 6.879238 | 4.92E-07 | 1.71E-05 | EREG |
| ENSG0000 | 1.617583 | 3.068606 | 2.30E-24 | 8.89E-22 | MYRF |
| ENSG0000 | 1.507955 | 2.844065 | 1.93E-23 | 6.99E-21 | WNT1 |
| ENSG0000 | -1.6803 | -3.20495 | 3.93E-06 | 0.000107 | MT1G |
| ENSG0000 | 1.050189 | 2.070801 | 7.26E-10 | 4.54E-08 | EFNB2 |
| ENSG0000 | 1.322592 | 2.501151 | 6.42E-06 | 0.000164 | SOX9 |
| ENSG0000 | 1.52143 | 2.870755 | 8.30E-30 | 6.24E-27 | HS3ST3B1 |
| ENSG0000 | 1.179816 | 2.265479 | 3.44E-40 | 6.56E-37 | IL1B |
| ENSG0000 | 1.864003 | 3.640163 | 9.08E-18 | 1.78E-15 | FOSB |
| ENSG0000 | 6.008752 | 64.38945 | 0.000486 | 0.006426 | FOXA2 |
| ENSG0000 | -1.13083 | -2.18984 | 4.13E-11 | 3.17E-09 | LAMP5 |
| ENSG0000 | -1.07454 | -2.10605 | 0.000308 | 0.004404 | MCF2L |

| | | | | | |
|---|---|---|---|---|---|
| ENSG0000 | 1.059959 | 2.084873 | 8.39E-06 | 0.000208 | CCR7 |
| ENSG0000 | 1.544303 | 2.91663 | 0.001409 | 0.01523 | FLRT1 |
| ENSG0000 | -2.44216 | -5.43456 | 0.000882 | 0.010461 | RGS13 |
| ENSG0000 | -2.78547 | -6.89461 | 0.000908 | 0.010697 | TRPV5 |
| ENSG0000 | -2.83129 | -7.11712 | 1.04E-07 | 4.16E-06 | F2RL3 |
| ENSG0000 | 1.106831 | 2.15372 | 1.90E-15 | 2.84E-13 | A4GALT |
| ENSG0000 | 1.056132 | 2.079349 | 7.52E-14 | 8.96E-12 | PODXL |
| ENSG0000 | 1.177866 | 2.262419 | 1.87E-06 | 5.56E-05 | FLNC |
| ENSG0000 | 2.936523 | 7.65564 | 1.66E-05 | 0.000373 | ISLR |
| ENSG0000 | 1.024868 | 2.034773 | 7.86E-08 | 3.22E-06 | COL5A1 |
| ENSG0000 | 1.317255 | 2.491915 | 1.35E-05 | 0.000312 | LAMA5 |
| ENSG0000 | 1.524974 | 2.877815 | 0.000245 | 0.003661 | ULBP3 |
| ENSG0000 | -5.33147 | -40.2655 | 0.005244 | 0.042127 | TEX101 |
| ENSG0000 | 2.399698 | 5.276926 | 1.44E-54 | 5.14E-51 | GFPT2 |
| ENSG0000 | -2.79443 | -6.93758 | 0.003938 | 0.033764 | ACY3 |
| ENSG0000 | -1.08274 | -2.11806 | 3.11E-40 | 6.35E-37 | CHIT1 |
| ENSG0000 | -1.01674 | -2.02335 | 2.26E-12 | 2.13E-10 | PDE6B |
| ENSG0000 | -1.58534 | -3.00078 | 3.69E-21 | 1.08E-18 | LGALS12 |
| ENSG0000 | -1.14381 | -2.20963 | 0.000138 | 0.002295 | PLAAT4 |
| ENSG0000 | -1.13141 | -2.19072 | 2.47E-25 | 1.06E-22 | GIMAP4 |
| ENSG0000 | -4.85162 | -28.8723 | 0.002633 | 0.024799 | SPINK5 |
| ENSG0000 | -1.03255 | -2.04564 | 3.46E-05 | 0.000701 | LYVE1 |
| ENSG0000 | -1.12746 | -2.18474 | 5.35E-06 | 0.00014 | RAB33A |
| ENSG0000 | -1.0164 | -2.02286 | 5.54E-13 | 5.81E-11 | SDS |
| ENSG0000 | 1.916908 | 3.77613 | 7.15E-33 | 6.82E-30 | NT5E |
| ENSG0000 | -1.07375 | -2.10489 | 2.37E-34 | 2.83E-31 | PRR5L |
| ENSG0000 | -1.0282 | -2.03947 | 2.23E-06 | 6.49E-05 | ELF5 |
| ENSG0000 | -5.16404 | -35.8535 | 0.002397 | 0.023052 | EDAR |
| ENSG0000 | -1.14184 | -2.20662 | 0.001246 | 0.013815 | EDNRB |
| ENSG0000 | -1.39722 | -2.63393 | 0.000375 | 0.005189 | CIDEB |
| ENSG0000 | 5.70708 | 52.2399 | 0.000581 | 0.00743 | RNASE2CP |
| ENSG0000 | 1.521085 | 2.870067 | 5.53E-09 | 2.93E-07 | SCN7A |
| ENSG0000 | -1.10638 | -2.15304 | 3.68E-12 | 3.39E-10 | IL10 |
| ENSG0000 | -5.17984 | -36.2483 | 0.001521 | 0.016121 | ALDOB |
| ENSG0000 | -6.04087 | -65.8391 | 2.12E-05 | 0.000459 | POU2F3 |
| ENSG0000 | -1.38828 | -2.61767 | 3.18E-06 | 8.89E-05 | CASP5 |
| ENSG0000 | 5.455836 | 43.89047 | 0.001315 | 0.014421 | SLC28A2 |
| ENSG0000 | 1.477191 | 2.784062 | 6.02E-30 | 4.65E-27 | STRA6 |
| ENSG0000 | -4.55335 | -23.4799 | 0.002053 | 0.020446 | BCL2L10 |
| ENSG0000 | -1.11719 | -2.16924 | 5.13E-05 | 0.000984 | IFI44L |
| ENSG0000 | 1.527016 | 2.881891 | 0.002883 | 0.026591 | ARHGAP29 |
| ENSG0000 | -1.57406 | -2.97742 | 7.45E-06 | 0.000188 | DNASE2B |
| ENSG0000 | -3.9875 | -15.862 | 0.003039 | 0.027739 | LHCGR |
| ENSG0000 | -4.84983 | -28.8366 | 0.003453 | 0.030663 | BTBD16 |
| ENSG0000 | -1.92085 | -3.78647 | 0.002525 | 0.023983 | MYPN |
| ENSG0000 | -1.43501 | -2.70384 | 1.01E-21 | 3.08E-19 | SLC40A1 |
| ENSG0000 | 1.263313 | 2.400464 | 0.001317 | 0.014436 | FGF2 |

| | | | | | |
|---|---|---|---|---|---|
| ENSG0000 | 1.141294 | 2.205787 | 2.46E-52 | 7.04E-49 | SLC39A8 |
| ENSG0000 | 1.24043 | 2.36269 | 4.45E-07 | 1.56E-05 | RBP5 |
| ENSG0000 | -1.36732 | -2.57992 | 1.63E-32 | 1.50E-29 | AMDHD1 |
| ENSG0000 | -5.7378 | -53.3642 | 0.000863 | 0.010303 | ESR2 |
| ENSG0000 | 1.285296 | 2.43732 | 7.63E-15 | 1.03E-12 | JDP2 |
| ENSG0000 | -1.04165 | -2.05858 | 2.15E-06 | 6.30E-05 | AK7 |
| ENSG0000 | 1.635283 | 3.106486 | 3.06E-10 | 2.02E-08 | ABCC12 |
| ENSG0000 | -1.17589 | -2.25932 | 0.000408 | 0.005564 | CDH13 |
| ENSG0000 | 3.583141 | 11.98486 | 0.005571 | 0.044231 | GATA6 |
| ENSG0000 | -1.22504 | -2.33762 | 1.55E-05 | 0.000352 | FBXO15 |
| ENSG0000 | -5.3839 | -41.7558 | 0.001407 | 0.015224 | CBLN2 |
| ENSG0000 | 1.843962 | 3.589945 | 0.003641 | 0.031885 | MYOM3 |
| ENSG0000 | 1.261414 | 2.397306 | 6.76E-10 | 4.26E-08 | HSPG2 |
| ENSG0000 | 2.431942 | 5.396192 | 1.13E-09 | 6.81E-08 | TINAGL1 |
| ENSG0000 | -1.59039 | -3.01132 | 1.28E-13 | 1.47E-11 | ADCY10 |
| ENSG0000 | 1.706584 | 3.263871 | 4.24E-10 | 2.75E-08 | KCNN3 |
| ENSG0000 | -5.87036 | -58.4999 | 0.006345 | 0.048612 | PKLR |
| ENSG0000 | 3.875645 | 14.67863 | 0.000111 | 0.001909 | ACKR3 |
| ENSG0000 | -2.10242 | -4.29428 | 0.000166 | 0.002658 | TRPM8 |
| ENSG0000 | -1.21958 | -2.3288 | 2.27E-08 | 1.07E-06 | ITGA9 |
| ENSG0000 | -4.90218 | -29.9023 | 0.000985 | 0.011421 | SLC22A14 |
| ENSG0000 | 1.805599 | 3.495742 | 2.69E-13 | 2.95E-11 | TAGLN3 |
| ENSG0000 | 1.753052 | 3.370709 | 4.82E-10 | 3.11E-08 | UCN2 |
| ENSG0000 | -1.05205 | -2.07347 | 0.00013 | 0.00219 | PLAC8 |
| ENSG0000 | -1.77209 | -3.41549 | 9.38E-05 | 0.001664 | UGT3A1 |
| ENSG0000 | -1.40299 | -2.64448 | 0.00355 | 0.031238 | CRHBP |
| ENSG0000 | 1.52775 | 2.883358 | 7.89E-139 | 2.26E-134 | RNF145 |
| ENSG0000 | -1.32057 | -2.49766 | 4.50E-08 | 1.95E-06 | KCNMB1 |
| ENSG0000 | 1.283066 | 2.433556 | 0.000434 | 0.005863 | IGFBP3 |
| ENSG0000 | -4.51529 | -22.8685 | 0.00447 | 0.03734 | ASB15 |
| ENSG0000 | 1.342662 | 2.536188 | 2.26E-06 | 6.56E-05 | SLC16A2 |
| ENSG0000 | 1.321983 | 2.500096 | 0.000701 | 0.008634 | NFIB |
| ENSG0000 | 1.177011 | 2.261079 | 4.55E-05 | 0.000885 | CRB2 |
| ENSG0000 | 5.465674 | 44.19078 | 0.001434 | 0.01546 | FAM13C |
| ENSG0000 | -5.16535 | -35.886 | 0.00378 | 0.032824 | RGR |
| ENSG0000 | 2.184229 | 4.54484 | 0.0006 | 0.00762 | ANKRD1 |
| ENSG0000 | -1.90952 | -3.75683 | 0.000119 | 0.002029 | FEZ1 |
| ENSG0000 | -1.01422 | -2.01981 | 0.000896 | 0.010585 | MPZL2 |
| ENSG0000 | 1.29736 | 2.457787 | 0.003769 | 0.03276 | C20orf144 |
| ENSG0000 | 1.613594 | 3.060133 | 1.40E-55 | 5.74E-52 | HMGA2 |
| ENSG0000 | 3.193403 | 9.147664 | 5.41E-05 | 0.001028 | MMP3 |
| ENSG0000 | -1.26733 | -2.40715 | 5.69E-14 | 6.87E-12 | FCGR1A |
| ENSG0000 | 1.509967 | 2.848036 | 0.002991 | 0.027424 | ADGRL3 |
| ENSG0000 | -6.0325 | -65.4583 | 4.33E-05 | 0.000849 | SPATA4 |
| ENSG0000 | -5.76811 | -54.4971 | 0.002307 | 0.022375 | FREM2 |
| ENSG0000 | 1.102682 | 2.147535 | 6.41E-11 | 4.75E-09 | SLC7A11 |
| ENSG0000 | 3.863708 | 14.55767 | 0.003062 | 0.027915 | ENKUR |

| | | | | | |
|---|---|---|---|---|---|
| ENSG0000 | -1.22492 | -2.33742 | 7.98E-09 | 4.08E-07 | TDO2 |
| ENSG0000 | 1.460317 | 2.751689 | 2.14E-06 | 6.28E-05 | PTPN14 |
| ENSG0000 | -1.22563 | -2.33858 | 0.000244 | 0.003652 | ANKRD22 |
| ENSG0000 | -1.1146 | -2.16534 | 0.002212 | 0.021652 | BANK1 |
| ENSG0000 | 1.52136 | 2.870616 | 3.81E-09 | 2.09E-07 | BMP6 |
| ENSG0000 | -5.13143 | -35.052 | 0.003843 | 0.033213 | FEZF2 |
| ENSG0000 | 1.148759 | 2.217231 | 1.14E-53 | 3.61E-50 | LPCAT1 |
| ENSG0000 | 1.501818 | 2.831994 | 1.13E-05 | 0.000266 | PTPRD |
| ENSG0000 | -5.38295 | -41.7281 | 0.003939 | 0.033768 | SPHKAP |
| ENSG0000 | -1.35994 | -2.56675 | 4.47E-09 | 2.42E-07 | ANKRD29 |
| ENSG0000 | -5.3828 | -41.7239 | 0.001494 | 0.015901 | CERS3 |
| ENSG0000 | 1.461505 | 2.753955 | 0.001767 | 0.018134 | MMP21 |
| ENSG0000 | 1.01485 | 2.020693 | 0.000959 | 0.011169 | JAM2 |
| ENSG0000 | 1.245577 | 2.371134 | 3.43E-12 | 3.18E-10 | PPM1J |
| ENSG0000 | 1.05033 | 2.071003 | 1.76E-06 | 5.30E-05 | PXYLP1 |
| ENSG0000 | 5.749027 | 53.78109 | 0.000107 | 0.00185 | KCNS2 |
| ENSG0000 | 1.922562 | 3.790957 | 0.000212 | 0.003249 | NMNAT2 |
| ENSG0000 | -1.04599 | -2.06478 | 0.000114 | 0.001956 | CACNA2D3 |
| ENSG0000 | 2.060675 | 4.171814 | 1.64E-12 | 1.59E-10 | KCNJ15 |
| ENSG0000 | 2.288162 | 4.884335 | 0.001076 | 0.012285 | SLC34A2 |
| ENSG0000 | 1.34759 | 2.544866 | 0.00569 | 0.044823 | CLSTN2 |
| ENSG0000 | 1.014224 | 2.019816 | 0.000164 | 0.002635 | SHROOM4 |
| ENSG0000 | 2.883612 | 7.379957 | 0.001572 | 0.016563 | NRG2 |
| ENSG0000 | 1.174623 | 2.257338 | 1.21E-15 | 1.87E-13 | B4GALT5 |
| ENSG0000 | -1.04611 | -2.06496 | 5.67E-09 | 2.99E-07 | CD1B |
| ENSG0000 | 1.324266 | 2.504054 | 1.78E-06 | 5.35E-05 | CACHD1 |
| ENSG0000 | 1.4813 | 2.792002 | 7.42E-05 | 0.001364 | LAD1 |
| ENSG0000 | 5.029812 | 32.66812 | 9.25E-11 | 6.64E-09 | STC1 |
| ENSG0000 | -5.09881 | -34.2684 | 0.001768 | 0.018137 | PGLYRP3 |
| ENSG0000 | 1.125528 | 2.181814 | 0.002231 | 0.021822 | FTCD |
| ENSG0000 | -3.72502 | -13.2234 | 0.004028 | 0.034349 | SHANK2 |
| ENSG0000 | 1.112875 | 2.162762 | 5.51E-37 | 7.50E-34 | TAL1 |
| ENSG0000 | 1.150983 | 2.220651 | 2.75E-06 | 7.84E-05 | PDPN |
| ENSG0000 | 1.43416 | 2.702248 | 6.32E-10 | 3.99E-08 | RFTN2 |
| ENSG0000 | 5.83671 | 57.15111 | 4.33E-05 | 0.000848 | LRRTM1 |
| ENSG0000 | -4.95036 | -30.9176 | 0.003931 | 0.033728 | ALMS1P1 |
| ENSG0000 | -1.72279 | -3.30075 | 0.00121 | 0.013546 | VSNL1 |
| ENSG0000 | -4.8865 | -29.5789 | 0.002985 | 0.027375 | SLC16A14 |
| ENSG0000 | 1.217155 | 2.324878 | 0.001104 | 0.012575 | IGFBP7 |
| ENSG0000 | 3.954718 | 15.50561 | 0.004956 | 0.040411 | EOMES |
| ENSG0000 | -2.00767 | -4.02131 | 3.36E-26 | 1.63E-23 | MNDA |
| ENSG0000 | -5.59933 | -48.4804 | 0.002049 | 0.020418 | ALB |
| ENSG0000 | 2.615725 | 6.129312 | 0.003772 | 0.032776 | ADAMTS9 |
| ENSG0000 | 2.309575 | 4.95737 | 5.61E-31 | 4.86E-28 | CXCL3 |
| ENSG0000 | 3.158236 | 8.927375 | 4.63E-88 | 6.62E-84 | CXCL5 |
| ENSG0000 | 1.220625 | 2.330477 | 5.67E-07 | 1.93E-05 | PLXNB1 |
| ENSG0000 | -1.80392 | -3.49168 | 0.002006 | 0.020072 | NPY1R |

| | | | | | |
|---|---|---|---|---|---|
| ENSG0000 | 1.418225 | 2.672565 | 8.04E-11 | 5.87E-09 | ITGA2 |
| ENSG0000 | 1.716795 | 3.287054 | 0.00023 | 0.003486 | EDIL3 |
| ENSG0000 | -5.55945 | -47.1587 | 0.000775 | 0.009394 | HTR4 |
| ENSG0000 | 5.174062 | 36.10337 | 0.003031 | 0.027677 | DACT2 |
| ENSG0000 | 1.422571 | 2.680628 | 7.82E-14 | 9.29E-12 | IL31RA |
| ENSG0000 | 5.80671 | 55.97499 | 0.000859 | 0.01025 | SMAD5-AS1 |
| ENSG0000 | 1.524429 | 2.876728 | 1.82E-20 | 4.81E-18 | STEAP1 |
| ENSG0000 | -1.15211 | -2.22239 | 1.05E-10 | 7.45E-09 | TMEM71 |
| ENSG0000 | -5.14851 | -35.4696 | 0.002811 | 0.026118 | ASB11 |
| ENSG0000 | 1.470889 | 2.771927 | 8.10E-13 | 8.25E-11 | AQP3 |
| ENSG0000 | -2.04392 | -4.12364 | 1.98E-25 | 8.71E-23 | FOLR2 |
| ENSG0000 | -1.01797 | -2.02507 | 1.86E-09 | 1.08E-07 | TMEM63C |
| ENSG0000 | -1.42874 | -2.69211 | 0.001585 | 0.016682 | AKR1E2 |
| ENSG0000 | 1.839184 | 3.578077 | 1.24E-17 | 2.37E-15 | DACT1 |
| ENSG0000 | -1.64152 | -3.11995 | 0.000997 | 0.011539 | PPP1R36 |
| ENSG0000 | 1.145847 | 2.21276 | 3.78E-06 | 0.000103 | PACSIN3 |
| ENSG0000 | -1.07404 | -2.10532 | 0.000541 | 0.007016 | PDZRN4 |
| ENSG0000 | 1.204505 | 2.304581 | 3.81E-27 | 2.06E-24 | HTRA1 |
| ENSG0000 | -5.10904 | -34.5124 | 0.006295 | 0.048272 | None |
| ENSG0000 | -1.03136 | -2.04394 | 1.67E-12 | 1.61E-10 | PLD4 |
| ENSG0000 | 1.452223 | 2.736294 | 2.59E-09 | 1.46E-07 | RRAD |
| ENSG0000 | 1.155753 | 2.228006 | 3.91E-37 | 5.89E-34 | ANPEP |
| ENSG0000 | 1.219964 | 2.329409 | 1.77E-08 | 8.56E-07 | NAV2 |
| ENSG0000 | 1.283784 | 2.434767 | 0.001829 | 0.018636 | IGF2 |
| ENSG0000 | 1.604048 | 3.039951 | 2.91E-05 | 0.000604 | GNG8 |
| ENSG0000 | -1.12157 | -2.17584 | 2.29E-26 | 1.15E-23 | TUBA1A |
| ENSG0000 | 1.837639 | 3.574246 | 0.001889 | 0.019116 | AQP2 |
| ENSG0000 | 1.379518 | 2.601814 | 0.001537 | 0.016257 | CDC42EP5 |
| ENSG0000 | 1.345721 | 2.541571 | 0.00151 | 0.016039 | TMEM88 |
| ENSG0000 | -1.55954 | -2.9476 | 3.14E-05 | 0.000644 | GSDMA |
| ENSG0000 | -1.22151 | -2.33191 | 1.77E-28 | 1.15E-25 | RAB3IL1 |
| ENSG0000 | 1.066926 | 2.094965 | 0.000238 | 0.003572 | PNOC |
| ENSG0000 | -1.20066 | -2.29845 | 0.00085 | 0.010172 | DEGS2 |
| ENSG0000 | 1.167882 | 2.246816 | 0.004512 | 0.037595 | AXIN2 |
| ENSG0000 | 1.032939 | 2.046189 | 0.00012 | 0.002043 | ATOH8 |
| ENSG0000 | -3.53524 | -11.5935 | 0.004872 | 0.039917 | CTRB2 |
| ENSG0000 | 1.080218 | 2.114355 | 0.002195 | 0.021527 | IRS1 |
| ENSG0000 | 1.41428 | 2.665267 | 0.00191 | 0.019288 | RAB3B |
| ENSG0000 | -2.52679 | -5.76288 | 2.77E-19 | 6.39E-17 | P2RY12 |
| ENSG0000 | -1.06535 | -2.09268 | 1.55E-05 | 0.000351 | RNASE2 |
| ENSG0000 | 1.399699 | 2.638465 | 3.41E-14 | 4.26E-12 | NPR1 |
| ENSG0000 | 1.063687 | 2.090266 | 0.00165 | 0.017251 | CHRNA5 |
| ENSG0000 | -1.11078 | -2.15962 | 1.33E-06 | 4.17E-05 | MT1E |
| ENSG0000 | 1.012826 | 2.01786 | 3.27E-18 | 6.69E-16 | WNT10B |
| ENSG0000 | 2.257783 | 4.782559 | 2.19E-33 | 2.32E-30 | TM4SF1 |
| ENSG0000 | 1.382868 | 2.607862 | 1.57E-15 | 2.37E-13 | NLGN2 |
| ENSG0000 | 4.043552 | 16.49037 | 0.001633 | 0.017093 | HSD17B13 |

| | | | | | |
|---|---|---|---|---|---|
| ENSG0000 | 5.488134 | 44.88414 | 0.000942 | 0.011021 | SYT9 |
| ENSG0000 | 1.580206 | 2.990126 | 2.96E-07 | 1.08E-05 | GPR37 |
| ENSG0000 | 1.140745 | 2.204948 | 1.28E-20 | 3.52E-18 | HTRA3 |
| ENSG0000 | 1.120079 | 2.173589 | 1.72E-19 | 4.11E-17 | MTSS1 |
| ENSG0000 | -4.54865 | -23.4035 | 0.005899 | 0.045986 | PLA2G1B |
| ENSG0000 | -1.20115 | -2.29923 | 0.006197 | 0.047647 | DCDC1 |
| ENSG0000 | 1.571816 | 2.972786 | 0.000735 | 0.008977 | LRRC8E |
| ENSG0000 | -1.38737 | -2.61601 | 1.19E-09 | 7.13E-08 | TMEM37 |
| ENSG0000 | 1.090731 | 2.129819 | 2.21E-09 | 1.26E-07 | HOPX |
| ENSG0000 | 1.208683 | 2.311265 | 3.53E-06 | 9.68E-05 | LPAR3 |
| ENSG0000 | -1.0601 | -2.08508 | 4.00E-13 | 4.31E-11 | PTCRA |
| ENSG0000 | -1.10793 | -2.15536 | 3.74E-07 | 1.33E-05 | S100Z |
| ENSG0000 | 1.488726 | 2.80641 | 3.24E-05 | 0.000662 | SYNPO |
| ENSG0000 | 2.421513 | 5.357325 | 2.24E-22 | 7.44E-20 | ID4 |
| ENSG0000 | 4.66488 | 25.36699 | 0.003501 | 0.030975 | PURG |
| ENSG0000 | -4.86901 | -29.2226 | 0.003793 | 0.032904 | COL6A5 |
| ENSG0000 | -1.41892 | -2.67386 | 1.20E-12 | 1.19E-10 | RAB37 |
| ENSG0000 | 1.347267 | 2.544297 | 1.22E-05 | 0.000286 | GXYLT2 |
| ENSG0000 | -1.19165 | -2.28414 | 1.05E-06 | 3.33E-05 | HPSE |
| ENSG0000 | 4.882926 | 29.50579 | 0.0046 | 0.03822 | ADAMTS20 |
| ENSG0000 | 1.350227 | 2.549523 | 0.001859 | 0.018898 | ZNF483 |
| ENSG0000 | 1.846974 | 3.597448 | 5.24E-09 | 2.79E-07 | SULT1B1 |
| ENSG0000 | -1.18918 | -2.28023 | 7.98E-18 | 1.57E-15 | GPR160 |
| ENSG0000 | -1.12569 | -2.18206 | 9.46E-12 | 8.25E-10 | SLCO4C1 |
| ENSG0000 | 4.321093 | 19.98842 | 0.00064 | 0.008055 | C9orf131 |
| ENSG0000 | -1.213 | -2.31819 | 7.49E-24 | 2.82E-21 | TLR10 |
| ENSG0000 | -4.58471 | -23.9959 | 0.002437 | 0.023354 | IGDCC3 |
| ENSG0000 | -1.32808 | -2.51069 | 4.20E-08 | 1.84E-06 | ADGRE1 |
| ENSG0000 | 1.080244 | 2.114394 | 4.48E-13 | 4.75E-11 | MARCKSL1 |
| ENSG0000 | -1.35639 | -2.56044 | 2.14E-07 | 8.03E-06 | C1orf127 |
| ENSG0000 | 2.160455 | 4.470558 | 2.04E-06 | 6.01E-05 | CHST1 |
| ENSG0000 | -1.70561 | -3.26167 | 2.46E-05 | 0.00052 | SCUBE2 |
| ENSG0000 | 1.351917 | 2.552511 | 7.97E-29 | 5.43E-26 | CLCF1 |
| ENSG0000 | -6.18122 | -72.5659 | 8.06E-06 | 0.000201 | GPR152 |
| ENSG0000 | -5.73825 | -53.3808 | 6.30E-05 | 0.001182 | CCDC197 |
| ENSG0000 | -1.54418 | -2.91637 | 1.08E-29 | 7.93E-27 | GAPT |
| ENSG0000 | 4.836458 | 28.57057 | 0.004783 | 0.039402 | None |
| ENSG0000 | 1.71785 | 3.289459 | 0.003624 | 0.031768 | CRYBG2 |
| ENSG0000 | 1.590474 | 3.011483 | 0.004444 | 0.037157 | FOXG1 |
| ENSG0000 | -5.91929 | -60.5181 | 3.64E-05 | 0.000733 | HORMAD2 |
| ENSG0000 | 5.796255 | 55.57078 | 2.45E-05 | 0.000519 | IRX5 |
| ENSG0000 | 5.665458 | 50.75429 | 0.000306 | 0.004385 | GPR4 |
| ENSG0000 | 4.258857 | 19.14449 | 0.002762 | 0.025791 | None |
| ENSG0000 | -5.87887 | -58.8459 | 4.61E-05 | 0.000893 | SPATA31C2 |
| ENSG0000 | 1.122957 | 2.177929 | 8.55E-05 | 0.001531 | CTXN1 |
| ENSG0000 | -2.46404 | -5.51761 | 0.001888 | 0.019105 | CD28 |
| ENSG0000 | -1.00508 | -2.00705 | 8.39E-26 | 3.87E-23 | MAF |

| | | | | | |
|---|---|---|---|---|---|
| ENSG0000 | 1.453256 | 2.738254 | 4.21E-60 | 2.41E-56 | THBD |
| ENSG0000 | -1.04909 | -2.06922 | 4.16E-38 | 6.62E-35 | FUCA1 |
| ENSG0000 | 1.01165 | 2.016216 | 3.32E-06 | 9.22E-05 | VWA1 |
| ENSG0000 | 1.062511 | 2.088564 | 8.24E-08 | 3.37E-06 | ZFPM1 |
| ENSG0000 | -2.0068 | -4.01891 | 0.003846 | 0.033216 | ALOX15B |
| ENSG0000 | -1.41178 | -2.66065 | 0.000171 | 0.002724 | FCER1A |
| ENSG0000 | -1.50615 | -2.8405 | 1.36E-06 | 4.23E-05 | PCED1B |
| ENSG0000 | 1.6297 | 3.094487 | 0.00012 | 0.002039 | CITED4 |
| ENSG0000 | -1.53198 | -2.89183 | 5.16E-07 | 1.78E-05 | NAIPP2 |
| ENSG0000 | -5.14456 | -35.3725 | 0.005715 | 0.044924 | ADGRD2 |
| ENSG0000 | -1.03487 | -2.04893 | 2.16E-06 | 6.31E-05 | SSTR2 |
| ENSG0000 | 1.532707 | 2.893283 | 0.000241 | 0.003602 | MAPK15 |
| ENSG0000 | -1.09588 | -2.13743 | 1.01E-05 | 0.000242 | C6orf223 |
| ENSG0000 | -1.19486 | -2.28923 | 4.08E-12 | 3.73E-10 | P2RY13 |
| ENSG0000 | -1.37132 | -2.58707 | 0.00029 | 0.004213 | LINC02724 |
| ENSG0000 | 2.330088 | 5.028359 | 1.96E-08 | 9.36E-07 | RGMA |
| ENSG0000 | -5.09054 | -34.0725 | 0.005637 | 0.044603 | PDSS1P1 |
| ENSG0000 | -1.26977 | -2.41123 | 0.003153 | 0.02858 | DNM1P46 |
| ENSG0000 | -1.12221 | -2.1768 | 0.001702 | 0.017657 | TSPAN10 |
| ENSG0000 | 1.227849 | 2.342176 | 0.005198 | 0.041885 | CRIP2 |
| ENSG0000 | 1.08267 | 2.117953 | 4.45E-06 | 0.000119 | GJC1 |
| ENSG0000 | 5.735797 | 53.29015 | 0.000103 | 0.001803 | NKX2-5 |
| ENSG0000 | -1.04685 | -2.06602 | 0.000179 | 0.002821 | EPHA10 |
| ENSG0000 | 5.452161 | 43.77883 | 0.001757 | 0.018063 | CABCOCO1 |
| ENSG0000 | 1.672133 | 3.186854 | 0.000222 | 0.003381 | GRIN2A |
| ENSG0000 | 3.941493 | 15.36412 | 0.004878 | 0.039932 | COL18A1-AS1 |
| ENSG0000 | 1.000162 | 2.000225 | 3.33E-27 | 1.87E-24 | RFLNB |
| ENSG0000 | 1.304934 | 2.470724 | 0.002147 | 0.021178 | KIRREL1 |
| ENSG0000 | 1.857403 | 3.623547 | 1.26E-06 | 3.95E-05 | ARSI |
| ENSG0000 | -4.90699 | -30.0021 | 0.00602 | 0.046643 | SRARP |
| ENSG0000 | 1.190951 | 2.283031 | 0.005719 | 0.044939 | NPW |
| ENSG0000 | 1.555217 | 2.938779 | 1.54E-05 | 0.00035 | CLDN5 |
| ENSG0000 | 1.245332 | 2.37073 | 0.000233 | 0.003506 | PKP3 |
| ENSG0000 | 1.339811 | 2.531182 | 1.54E-25 | 6.88E-23 | TAFA3 |
| ENSG0000 | 5.392756 | 42.01276 | 0.002468 | 0.023558 | LINC00313 |
| ENSG0000 | 1.482882 | 2.795065 | 6.02E-22 | 1.85E-19 | NOTUM |
| ENSG0000 | -1.07973 | -2.11364 | 7.87E-06 | 0.000197 | INKA1 |
| ENSG0000 | 5.574209 | 47.64354 | 7.59E-05 | 0.001389 | NTF3 |
| ENSG0000 | -1.04437 | -2.06246 | 0.004673 | 0.038643 | PMEL |
| ENSG0000 | -1.32001 | -2.49667 | 0.000963 | 0.011208 | SYN3 |
| ENSG0000 | -1.09617 | -2.13787 | 1.75E-06 | 5.27E-05 | CALHM1 |
| ENSG0000 | 1.514577 | 2.85715 | 0.001533 | 0.016218 | MATN1-AS1 |
| ENSG0000 | -1.06357 | -2.0901 | 1.11E-22 | 3.82E-20 | CD300LF |
| ENSG0000 | -1.11844 | -2.17113 | 1.37E-09 | 8.15E-08 | FFAR4 |
| ENSG0000 | 5.821666 | 56.55827 | 0.000348 | 0.004877 | |
| ENSG0000 | -1.07854 | -2.1119 | 5.74E-05 | 0.001087 | CYP4X1 |
| ENSG0000 | 1.05946 | 2.084151 | 5.54E-08 | 2.36E-06 | HPDL |

| | | | | | |
|---|---|---|---|---|---|
| ENSG0000 | 1.139496 | 2.203041 | 6.21E-06 | 0.00016 | TEAD1 |
| ENSG0000 | -1.48805 | -2.8051 | 7.34E-05 | 0.00135 | None |
| ENSG0000 | 1.253552 | 2.384277 | 9.01E-06 | 0.000219 | C11orf96 |
| ENSG0000 | -5.38035 | -41.653 | 0.001343 | 0.014664 | DPPA3 |
| ENSG0000 | 1.026929 | 2.037682 | 4.56E-09 | 2.46E-07 | PLEKHN1 |
| ENSG0000 | 1.391191 | 2.622952 | 4.07E-05 | 0.000806 | PERM1 |
| ENSG0000 | 2.618428 | 6.140805 | 0.000184 | 0.002879 | C5orf52 |
| ENSG0000 | 5.284827 | 38.98444 | 0.000592 | 0.007541 | C9orf153 |
| ENSG0000 | -1.07445 | -2.10592 | 0.002985 | 0.027375 | SELL |
| ENSG0000 | -4.92319 | -30.3408 | 0.001942 | 0.019584 | ACTBP11 |
| ENSG0000 | 1.560949 | 2.950479 | 3.85E-05 | 0.000768 | NCCRP1 |
| ENSG0000 | -1.77287 | -3.41733 | 0.006552 | 0.049859 | RHCE |
| ENSG0000 | 1.876109 | 3.670836 | 0.002833 | 0.026277 | HMX2 |
| ENSG0000 | 1.199231 | 2.296172 | 5.09E-06 | 0.000134 | RPSAP47 |
| ENSG0000 | 1.070858 | 2.100683 | 1.76E-09 | 1.03E-07 | SP6 |
| ENSG0000 | 1.326816 | 2.508484 | 0.003687 | 0.032178 | CLDN4 |
| ENSG0000 | 5.197515 | 36.69509 | 0.000488 | 0.006438 | SH2D5 |
| ENSG0000 | 1.052255 | 2.073768 | 0.000159 | 0.002565 | NCR1 |
| ENSG0000 | 5.712593 | 52.43992 | 0.001782 | 0.018241 | MYT1 |
| ENSG0000 | -1.02943 | -2.04122 | 1.53E-06 | 4.71E-05 | GREB1 |
| ENSG0000 | 1.070336 | 2.099922 | 7.30E-12 | 6.44E-10 | RFX8 |
| ENSG0000 | -1.09823 | -2.14092 | 9.71E-06 | 0.000234 | CRACDL |
| ENSG0000 | -5.76049 | -54.21 | 7.49E-05 | 0.001374 | DTHD1 |
| ENSG0000 | 2.701946 | 6.506792 | 0.000475 | 0.006304 | H4C3 |
| ENSG0000 | -5.14802 | -35.4576 | 0.000355 | 0.004961 | ZSCAN5B |
| ENSG0000 | 1.286398 | 2.439183 | 0.003688 | 0.032178 | HMGA2-AS1 |
| ENSG0000 | -1.08619 | -2.12312 | 1.04E-05 | 0.000247 | COL13A1 |
| ENSG0000 | 4.759746 | 27.09108 | 0.002729 | 0.025542 | FAM177B |
| ENSG0000 | -1.29906 | -2.46069 | 4.08E-09 | 2.23E-07 | FCGR1B |
| ENSG0000 | 5.876463 | 58.74781 | 0.000174 | 0.002765 | ZNF560 |
| ENSG0000 | 3.600853 | 12.13291 | 0.005523 | 0.043933 | MYL4 |
| ENSG0000 | 1.768278 | 3.406471 | 0.005976 | 0.0464 | ABCA4 |
| ENSG0000 | 1.647107 | 3.132049 | 3.65E-19 | 8.21E-17 | SMOC1 |
| ENSG0000 | 1.717358 | 3.288338 | 4.47E-12 | 4.04E-10 | F5 |
| ENSG0000 | 1.610807 | 3.054227 | 1.09E-28 | 7.25E-26 | APCDD1L |
| ENSG0000 | 1.673236 | 3.189292 | 2.23E-05 | 0.00048 | HSD3BP5 |
| ENSG0000 | 4.939517 | 30.68618 | 0.003304 | 0.029682 | CSAG1 |
| ENSG0000 | -5.4974 | -45.1734 | 0.00321 | 0.028999 | Y_RNA |
| ENSG0000 | 1.351722 | 2.552166 | 3.33E-05 | 0.000678 | SNORD104 |
| ENSG0000 | 6.897254 | 119.2012 | 0.000846 | 0.010134 | Y_RNA |
| ENSG0000 | 4.985233 | 31.67413 | 0.003289 | 0.029598 | SNORA63D |
| ENSG0000 | 4.965753 | 31.24932 | 0.005038 | 0.040925 | Y_RNA |
| ENSG0000 | 6.590537 | 96.37166 | 5.28E-05 | 0.00101 | RNU6-1082P |
| ENSG0000 | 2.846198 | 7.19103 | 0.000798 | 0.009642 | Y_RNA |
| ENSG0000 | 6.197169 | 73.37258 | 0.004219 | 0.035624 | Y_RNA |
| ENSG0000 | -4.59077 | -24.0968 | 0.006013 | 0.046608 | RNU4-28P |
| ENSG0000 | -4.91031 | -30.0713 | 0.004137 | 0.035059 | RN7SKP239 |

| | | | | | |
|---|---|---|---|---|---|
| ENSG0000 | 4.349434 | 20.38498 | 0.00594 | 0.046203 | RNY1P13 |
| ENSG0000 | -5.20347 | -36.8469 | 0.004316 | 0.036248 | RNU6-61P |
| ENSG0000 | -4.92438 | -30.3659 | 0.002905 | 0.026761 | LINC00501 |
| ENSG0000 | -1.04572 | -2.0644 | 1.26E-07 | 4.91E-06 | CR1 |
| ENSG0000 | 1.315774 | 2.489358 | 3.56E-11 | 2.77E-09 | PLPP4 |
| ENSG0000 | -4.52917 | -23.0896 | 0.005846 | 0.045629 | |
| ENSG0000 | -1.46405 | -2.75882 | 1.83E-06 | 5.47E-05 | GGTA1 |
| ENSG0000 | -5.10925 | -34.5175 | 0.005047 | 0.040975 | MAMDC2-AS1 |
| ENSG0000 | 1.368745 | 2.582458 | 0.000231 | 0.003489 | CLLU1-AS1 |
| ENSG0000 | -4.87372 | -29.3182 | 0.001226 | 0.013671 | HSP90AA4P |
| ENSG0000 | 1.780179 | 3.434689 | 0.00015 | 0.002449 | LINC01602 |
| ENSG0000 | -4.91218 | -30.1102 | 0.000546 | 0.007072 | LINC01460 |
| ENSG0000 | 1.062919 | 2.089154 | 1.01E-11 | 8.74E-10 | ADGRG1 |
| ENSG0000 | -2.00776 | -4.02158 | 0.0004 | 0.005469 | MT1H |
| ENSG0000 | -1.11701 | -2.16896 | 2.46E-07 | 9.11E-06 | COLQ |
| ENSG0000 | 4.987401 | 31.72177 | 0.004024 | 0.034349 | Y_RNA |
| ENSG0000 | 5.645028 | 50.04061 | 0.006397 | 0.048922 | U3 |
| ENSG0000 | -5.11458 | -34.6451 | 0.001983 | 0.019899 | RNU6-30P |
| ENSG0000 | 5.079921 | 33.82273 | 0.002046 | 0.020396 | Y_RNA |
| ENSG0000 | 5.488574 | 44.89783 | 0.000903 | 0.010654 | MIR645 |
| ENSG0000 | 5.884706 | 59.08442 | 0.00084 | 0.010075 | SNORD105 |
| ENSG0000 | 1.471625 | 2.773341 | 1.49E-09 | 8.79E-08 | MT-TY |
| ENSG0000 | 3.826246 | 14.18453 | 0.002783 | 0.025928 | MT-TS1 |
| ENSG0000 | 1.703561 | 3.257038 | 0.000395 | 0.005413 | MT-TG |
| ENSG0000 | 1.263541 | 2.400843 | 0.006052 | 0.046864 | MT-TH |
| ENSG0000 | 1.311427 | 2.481869 | 1.11E-05 | 0.000263 | MT-TL2 |
| ENSG0000 | 1.280722 | 2.429605 | 0.000287 | 0.004181 | MT-TE |
| ENSG0000 | 1.524898 | 2.877663 | 0.000583 | 0.007452 | MT-TT |
| ENSG0000 | 2.194921 | 4.578645 | 0.001136 | 0.012891 | TRGV1 |
| ENSG0000 | 2.482744 | 5.589596 | 0.005163 | 0.041659 | TRAJ23 |
| ENSG0000 | 2.15055 | 4.439969 | 0.005095 | 0.041305 | SNORA3B |
| ENSG0000 | 1.575057 | 2.979473 | 0.002822 | 0.026194 | None |
| ENSG0000 | -3.08632 | -8.49325 | 0.003004 | 0.027497 | PDCL3P5 |
| ENSG0000 | -5.98982 | -63.5499 | 1.27E-05 | 0.000294 | FABP5P10 |
| ENSG0000 | -1.29932 | -2.46113 | 0.003819 | 0.033102 | ARHGEF35 |
| ENSG0000 | 3.902301 | 14.95235 | 0.002732 | 0.025549 | MAGEA12 |
| ENSG0000 | -5.32311 | -40.0328 | 0.000854 | 0.010205 | None |
| ENSG0000 | 3.290057 | 9.78151 | 0.001633 | 0.017093 | None |
| ENSG0000 | 1.171595 | 2.252606 | 0.000178 | 0.002808 | S1PR3 |
| ENSG0000 | 5.361716 | 41.11852 | 0.001192 | 0.013388 | RAB5CP1 |
| ENSG0000 | -1.05484 | -2.07749 | 0.006128 | 0.047243 | RPL7AP64 |
| ENSG0000 | -1.72814 | -3.31301 | 0.00145 | 0.01558 | None |
| ENSG0000 | 1.774175 | 3.420424 | 0.000932 | 0.010926 | MPRIPP1 |
| ENSG0000 | -6.07163 | -67.2576 | 0.0007 | 0.008634 | FAM66B |
| ENSG0000 | -5.14538 | -35.3927 | 0.002421 | 0.023223 | None |
| ENSG0000 | 6.164505 | 71.73003 | 0.000297 | 0.004281 | MARCKSL1P2 |
| ENSG0000 | -5.52675 | -46.1016 | 0.002021 | 0.020181 | DDX18P3 |

| | | | | | |
|---|---|---|---|---|---|
| ENSG0000 | -1.15783 | -2.23121 | 0.006177 | 0.047562 | None |
| ENSG0000 | 4.140608 | 17.63791 | 0.00142 | 0.015328 | RPL31P52 |
| ENSG0000 | -5.34064 | -40.5221 | 0.00095 | 0.011079 | CATSPERZ |
| ENSG0000 | 4.544767 | 23.34055 | 0.003638 | 0.031869 | TAFA5 |
| ENSG0000 | 6.137542 | 70.40186 | 0.00303 | 0.027677 | None |
| ENSG0000 | 4.881397 | 29.47453 | 0.001169 | 0.01319 | None |
| ENSG0000 | 5.505543 | 45.42903 | 0.000801 | 0.009656 | MIR548I2 |
| ENSG0000 | -6.25436 | -76.3397 | 7.45E-06 | 0.000188 | EBF2 |
| ENSG0000 | 6.245088 | 75.85058 | 6.53E-05 | 0.001219 | LINC02860 |
| ENSG0000 | 1.069023 | 2.098012 | 2.53E-06 | 7.27E-05 | None |
| ENSG0000 | -5.6076 | -48.7591 | 0.001849 | 0.018803 | RNU4-86P |
| ENSG0000 | 4.916128 | 30.19271 | 0.000648 | 0.00813 | RN7SKP150 |
| ENSG0000 | -5.14693 | -35.4307 | 0.001803 | 0.018421 | RNU4-78P |
| ENSG0000 | -5.22213 | -37.3265 | 0.003525 | 0.031118 | None |
| ENSG0000 | 2.44316 | 5.438317 | 0.003564 | 0.03133 | None |
| ENSG0000 | -4.58744 | -24.0412 | 0.003112 | 0.028293 | EEF1A1P1 |
| ENSG0000 | 1.111536 | 2.160756 | 0.001619 | 0.016981 | None |
| ENSG0000 | -1.13179 | -2.1913 | 1.99E-07 | 7.51E-06 | LINC01857 |
| ENSG0000 | 5.80696 | 55.98469 | 0.001729 | 0.017871 | ARHGEF2-AS1 |
| ENSG0000 | 5.128817 | 34.98869 | 0.00385 | 0.033227 | OLFM5P |
| ENSG0000 | -1.01844 | -2.02572 | 1.41E-07 | 5.47E-06 | None |
| ENSG0000 | -5.08574 | -33.9594 | 0.001158 | 0.013091 | MCCD1P2 |
| ENSG0000 | 1.527167 | 2.882194 | 3.11E-06 | 8.72E-05 | None |
| ENSG0000 | -5.61182 | -48.9019 | 0.000441 | 0.005933 | SLC31A1P1 |
| ENSG0000 | -1.01885 | -2.02631 | 0.000617 | 0.007799 | EML4-AS1 |
| ENSG0000 | 5.61173 | 48.89889 | 0.002416 | 0.023218 | ARL14EPP1 |
| ENSG0000 | -1.27511 | -2.42017 | 0.000809 | 0.009738 | LINC01645 |
| ENSG0000 | -1.41258 | -2.66213 | 0.000165 | 0.002656 | None |
| ENSG0000 | -1.2892 | -2.44392 | 4.82E-14 | 5.89E-12 | HSPA7 |
| ENSG0000 | -4.90232 | -29.9051 | 0.00102 | 0.011752 | None |
| ENSG0000 | -5.41117 | -42.5523 | 0.00052 | 0.006806 | None |
| ENSG0000 | -2.70912 | -6.53921 | 0.000262 | 0.003861 | DBH-AS1 |
| ENSG0000 | -1.17785 | -2.26239 | 0.000176 | 0.002793 | None |
| ENSG0000 | 1.247583 | 2.374433 | 1.12E-07 | 4.45E-06 | MTND1P23 |
| ENSG0000 | -4.83415 | -28.5249 | 0.002736 | 0.025571 | LAMP5-AS1 |
| ENSG0000 | -5.1447 | -35.3761 | 0.004912 | 0.040141 | None |
| ENSG0000 | -5.94663 | -61.6758 | 0.000713 | 0.008752 | LINC01733 |
| ENSG0000 | 5.233587 | 37.62415 | 0.004406 | 0.036894 | None |
| ENSG0000 | 1.437402 | 2.708327 | 0.003692 | 0.032197 | CFLAR-AS1 |
| ENSG0000 | -1.08669 | -2.12387 | 0.002147 | 0.021178 | LINC02542 |
| ENSG0000 | 6.930951 | 122.0181 | 1.04E-05 | 0.000247 | RPL22P3 |
| ENSG0000 | 5.505586 | 45.43041 | 0.000555 | 0.007164 | USF1P1 |
| ENSG0000 | -5.56915 | -47.4769 | 0.003012 | 0.027549 | PSMD10P2 |
| ENSG0000 | 1.289762 | 2.444878 | 0.004492 | 0.037499 | None |
| ENSG0000 | 5.910793 | 60.16251 | 0.0013 | 0.014275 | SMCR5 |
| ENSG0000 | -1.71288 | -3.27816 | 2.14E-14 | 2.74E-12 | |
| ENSG0000 | -1.43098 | -2.6963 | 2.26E-05 | 0.000485 | LINC02611 |

| | | | | | |
|---|---|---|---|---|---|
| ENSG0000 | -1.16388 | -2.24059 | 0.003993 | 0.034134 | RHEBP2 |
| ENSG0000 | -4.80942 | -28.0401 | 0.005746 | 0.045091 | None |
| ENSG0000 | -4.82879 | -28.4192 | 0.001228 | 0.013676 | None |
| ENSG0000 | 5.249633 | 38.04495 | 0.000771 | 0.00935 | MTND5P1 |
| ENSG0000 | 5.424356 | 42.94314 | 0.000902 | 0.01065 | None |
| ENSG0000 | -3.8783 | -14.7056 | 0.001421 | 0.015333 | None |
| ENSG0000 | -2.3558 | -5.11878 | 0.002435 | 0.02334 | None |
| ENSG0000 | 6.911562 | 120.3892 | 0.000446 | 0.005995 | MTND1P11 |
| ENSG0000 | 3.050972 | 8.287702 | 0.000196 | 0.003037 | LYPLAL1-AS1 |
| ENSG0000 | -5.09182 | -34.1028 | 0.004101 | 0.034828 | None |
| ENSG0000 | -4.55156 | -23.4507 | 0.003808 | 0.033031 | None |
| ENSG0000 | -4.90624 | -29.9865 | 0.002076 | 0.020637 | THRB-AS1 |
| ENSG0000 | 6.180999 | 72.55481 | 0.003899 | 0.033522 | MTND1P9 |
| ENSG0000 | 5.537089 | 46.43332 | 0.000287 | 0.004178 | None |
| ENSG0000 | 4.967648 | 31.2904 | 0.005529 | 0.043967 | PSMA6P2 |
| ENSG0000 | -1.87274 | -3.66228 | 0.001777 | 0.01821 | CCDC26 |
| ENSG0000 | -1.87936 | -3.67912 | 9.07E-13 | 9.14E-11 | PGA4 |
| ENSG0000 | 1.299493 | 2.461424 | 0.002316 | 0.022426 | HLA-DRB6 |
| ENSG0000 | -5.3948 | -42.0723 | 0.000499 | 0.006565 | None |
| ENSG0000 | -1.25631 | -2.38884 | 0.00066 | 0.008256 | MBNL1-AS1 |
| ENSG0000 | -1.71569 | -3.28453 | 4.73E-05 | 0.000915 | LINC01150 |
| ENSG0000 | 4.855364 | 28.94744 | 0.001959 | 0.019719 | None |
| ENSG0000 | 1.002454 | 2.003406 | 0.00102 | 0.011751 | ATXN1-AS1 |
| ENSG0000 | 5.211172 | 37.04412 | 0.002183 | 0.021446 | None |
| ENSG0000 | 1.088088 | 2.125921 | 0.000225 | 0.003416 | None |
| ENSG0000 | 2.619189 | 6.144045 | 7.81E-06 | 0.000196 | |
| ENSG0000 | 5.100689 | 34.31314 | 0.005646 | 0.044628 | STK24P1 |
| ENSG0000 | 5.723446 | 52.83587 | 0.001226 | 0.013671 | TRBV5-4 |
| ENSG0000 | -4.83512 | -28.5441 | 0.002493 | 0.023753 | TPRG1-AS2 |
| ENSG0000 | -5.31636 | -39.8458 | 0.000109 | 0.001878 | LINC01381 |
| ENSG0000 | -3.73271 | -13.294 | 0.002443 | 0.023399 | None |
| ENSG0000 | 1.339824 | 2.531204 | 0.000212 | 0.003246 | None |
| ENSG0000 | 5.118392 | 34.73678 | 0.00602 | 0.046643 | None |
| ENSG0000 | -4.90386 | -29.9371 | 0.002325 | 0.022483 | None |
| ENSG0000 | -5.19381 | -36.6009 | 0.000698 | 0.008616 | SRGAP2-AS1 |
| ENSG0000 | 1.547174 | 2.922442 | 4.27E-37 | 6.11E-34 | APCDD1L-DT |
| ENSG0000 | -4.88611 | -29.5709 | 0.004104 | 0.03483 | SEPTIN14P21 |
| ENSG0000 | -1.09189 | -2.13153 | 7.96E-05 | 0.001442 | LINC00278 |
| ENSG0000 | -1.00087 | -2.00121 | 0.002043 | 0.020378 | EMBP1 |
| ENSG0000 | -4.85625 | -28.9653 | 0.001359 | 0.014784 | RPS6KA2-IT1 |
| ENSG0000 | -5.36557 | -41.2285 | 0.002702 | 0.025352 | VN1R54P |
| ENSG0000 | -4.91028 | -30.0705 | 0.004096 | 0.034795 | FTLP8 |
| ENSG0000 | -1.81715 | -3.52383 | 3.71E-05 | 0.000745 | TMEM114 |
| ENSG0000 | -1.1912 | -2.28342 | 0.004679 | 0.038679 | None |
| ENSG0000 | -4.8678 | -29.198 | 0.003571 | 0.031369 | |
| ENSG0000 | 1.041113 | 2.057815 | 4.60E-05 | 0.000893 | EMSLR |
| ENSG0000 | -1.25224 | -2.38212 | 4.90E-06 | 0.00013 | LINC02642 |

| | | | | | |
|---|---|---|---|---|---|
| ENSG0000 | 5.06526 | 33.48076 | 0.006082 | 0.046997 | None |
| ENSG0000 | -1.14528 | -2.21188 | 5.14E-05 | 0.000984 | None |
| ENSG0000 | 1.106352 | 2.153006 | 0.001901 | 0.01921 | None |
| ENSG0000 | 1.249524 | 2.37763 | 0.000543 | 0.007035 | SLCO4A1-AS1 |
| ENSG0000 | -4.92997 | -30.4837 | 0.005742 | 0.045087 | GSTO3P |
| ENSG0000 | -1.03004 | -2.04208 | 3.32E-07 | 1.20E-05 | LYRM9 |
| ENSG0000 | -5.40089 | -42.2504 | 0.003931 | 0.033728 | NUCKS1P1 |
| ENSG0000 | 2.753128 | 6.741773 | 0.003283 | 0.029577 | CRYZP1 |
| ENSG0000 | -2.2755 | -4.84167 | 0.005726 | 0.044973 | None |
| ENSG0000 | -1.38958 | -2.62002 | 0.003325 | 0.029814 | ZFY-AS1 |
| ENSG0000 | -3.53211 | -11.5683 | 0.004062 | 0.034553 | LNCARSR |
| ENSG0000 | -6.12074 | -69.5867 | 4.52E-05 | 0.000881 | MROH3P |
| ENSG0000 | -3.58485 | -11.999 | 0.005193 | 0.041856 | FAM238A |
| ENSG0000 | -5.54287 | -46.6199 | 2.59E-05 | 0.000546 | None |
| ENSG0000 | -5.60299 | -48.6036 | 0.001866 | 0.01895 | CCNQP1 |
| ENSG0000 | -4.86561 | -29.1537 | 0.005891 | 0.045949 | None |
| ENSG0000 | -1.20706 | -2.30867 | 0.001242 | 0.013777 | TPRG1-AS1 |
| ENSG0000 | -5.60716 | -48.7442 | 0.000168 | 0.002689 | LINC01191 |
| ENSG0000 | 5.830112 | 56.89036 | 0.000158 | 0.002561 | None |
| ENSG0000 | 5.979118 | 63.08033 | 0.000292 | 0.004221 | CSNK1G2P1 |
| ENSG0000 | -1.65634 | -3.15216 | 0.003528 | 0.03113 | H2BP2 |
| ENSG0000 | 1.373845 | 2.591604 | 0.002723 | 0.025517 | None |
| ENSG0000 | -5.47495 | -44.4759 | 0.004754 | 0.039185 | None |
| ENSG0000 | -5.52137 | -45.9303 | 0.005827 | 0.045525 | None |
| ENSG0000 | 5.220943 | 37.29584 | 0.000211 | 0.003244 | None |
| ENSG0000 | 5.562217 | 47.24918 | 0.00074 | 0.009026 | NSRP1P1 |
| ENSG0000 | 5.995277 | 63.79084 | 0.000509 | 0.006679 | SNRPFP4 |
| ENSG0000 | -1.0085 | -2.01182 | 0.00373 | 0.032491 | LINC00484 |
| ENSG0000 | 5.491038 | 44.97459 | 0.000226 | 0.00343 | None |
| ENSG0000 | 6.18976 | 72.99672 | 7.42E-06 | 0.000187 | COX7CP1 |
| ENSG0000 | -1.24051 | -2.36282 | 7.98E-09 | 4.08E-07 | None |
| ENSG0000 | 5.75267 | 53.91706 | 0.00591 | 0.046035 | None |
| ENSG0000 | -4.56909 | -23.7374 | 0.003066 | 0.027941 | None |
| ENSG0000 | -5.41274 | -42.5989 | 0.000748 | 0.009104 | LINC02087 |
| ENSG0000 | 5.613535 | 48.96012 | 3.65E-05 | 0.000734 | EHMT2-AS1 |
| ENSG0000 | -5.40064 | -42.2429 | 0.00352 | 0.031087 | NR2F1-AS1 |
| ENSG0000 | -1.26514 | -2.40351 | 0.003686 | 0.032178 | DCLRE1CP1 |
| ENSG0000 | -4.91936 | -30.2603 | 0.001181 | 0.013295 | PKN2-AS1 |
| ENSG0000 | 6.388331 | 83.76824 | 0.000983 | 0.011414 | VDAC1P11 |
| ENSG0000 | -5.5123 | -45.6424 | 0.00166 | 0.01732 | ZNF101P1 |
| ENSG0000 | 1.37068 | 2.585925 | 0.000734 | 0.008973 | DGUOK-AS1 |
| ENSG0000 | 6.324914 | 80.16576 | 7.02E-05 | 0.0013 | None |
| ENSG0000 | 5.590653 | 48.18971 | 0.001427 | 0.015392 | RNU7-38P |
| ENSG0000 | -1.09216 | -2.13193 | 0.002622 | 0.02471 | MTATP8P1 |
| ENSG0000 | 2.151822 | 4.443888 | 6.23E-08 | 2.61E-06 | PCDHGC5 |
| ENSG0000 | -6.01416 | -64.6312 | 0.000359 | 0.005002 | None |
| ENSG0000 | -2.10498 | -4.30191 | 2.92E-07 | 1.07E-05 | TDGF1 |

| | | | | | |
|---|---|---|---|---|---|
| ENSG0000 | -4.96171 | -31.1619 | 0.002823 | 0.026194 | None |
| ENSG0000 | 5.207066 | 36.93883 | 0.004412 | 0.036909 | None |
| ENSG0000 | 1.006494 | 2.009023 | 0.002503 | 0.023815 | EGFL8 |
| ENSG0000 | 5.473621 | 44.43489 | 0.000449 | 0.006028 | None |
| ENSG0000 | -5.84388 | -57.436 | 0.005223 | 0.042039 | None |
| ENSG0000 | 1.66084 | 3.162006 | 0.000836 | 0.010034 | None |
| ENSG0000 | 5.396332 | 42.11705 | 0.002324 | 0.02248 | None |
| ENSG0000 | 2.533977 | 5.791661 | 0.004411 | 0.036909 | RPSAP52 |
| ENSG0000 | -5.58446 | -47.9834 | 0.002512 | 0.023878 | None |
| ENSG0000 | -4.85684 | -28.9771 | 0.001722 | 0.017823 | DENND6A-DT |
| ENSG0000 | 6.557533 | 94.192 | 0.000258 | 0.00382 | None |
| ENSG0000 | -5.77329 | -54.6932 | 7.24E-05 | 0.001334 | RPL12P32 |
| ENSG0000 | -5.43099 | -43.1412 | 0.001263 | 0.013961 | RPL12P21 |
| ENSG0000 | 5.450484 | 43.72795 | 0.000525 | 0.00686 | PDE6B-AS1 |
| ENSG0000 | -1.24488 | -2.37 | 0.002983 | 0.027375 | None |
| ENSG0000 | -5.59389 | -48.2978 | 9.65E-05 | 0.001702 | H3P13 |
| ENSG0000 | 3.556971 | 11.76942 | 0.005342 | 0.04278 | GATA2-AS1 |
| ENSG0000 | 5.411339 | 42.55743 | 0.000316 | 0.004498 | DUXAP10 |
| ENSG0000 | -5.16216 | -35.8067 | 0.001546 | 0.016315 | RPS2P45 |
| ENSG0000 | 5.206482 | 36.92387 | 0.000986 | 0.011423 | None |
| ENSG0000 | 4.867735 | 29.19673 | 0.002823 | 0.026194 | RPL12P33 |
| ENSG0000 | 1.059216 | 2.083798 | 0.000247 | 0.003681 | WNT5A-AS1 |
| ENSG0000 | -1.17236 | -2.2538 | 0.005597 | 0.044375 | None |
| ENSG0000 | -1.45336 | -2.73845 | 7.44E-10 | 4.65E-08 | FCGR2C |
| ENSG0000 | -5.5704 | -47.518 | 0.000142 | 0.002347 | LINC02273 |
| ENSG0000 | -1.08964 | -2.1282 | 1.15E-17 | 2.22E-15 | LINC00900 |
| ENSG0000 | -1.11905 | -2.17204 | 6.01E-14 | 7.19E-12 | USP51 |
| ENSG0000 | -1.48279 | -2.79488 | 4.08E-13 | 4.37E-11 | PCED1B-AS1 |
| ENSG0000 | -1.66342 | -3.16767 | 0.001384 | 0.015028 | NAIPP3 |
| ENSG0000 | 5.583056 | 47.9366 | 0.006434 | 0.049172 | None |
| ENSG0000 | -2.46045 | -5.50388 | 1.48E-06 | 4.54E-05 | None |
| ENSG0000 | -5.18151 | -36.2902 | 0.002096 | 0.020779 | None |
| ENSG0000 | -4.8522 | -28.884 | 0.00242 | 0.023223 | LINC02071 |
| ENSG0000 | -1.36515 | -2.57603 | 8.41E-11 | 6.10E-09 | NAIP |
| ENSG0000 | -5.35406 | -40.9009 | 0.000727 | 0.008908 | LINC01470 |
| ENSG0000 | 3.829579 | 14.21733 | 0.005193 | 0.041856 | None |
| ENSG0000 | 1.888634 | 3.702845 | 4.65E-12 | 4.16E-10 | TMEM158 |
| ENSG0000 | -1.21891 | -2.32771 | 2.45E-06 | 7.07E-05 | ARPIN-AP3S2 |
| ENSG0000 | -5.71043 | -52.3612 | 2.31E-05 | 0.000493 | None |
| ENSG0000 | -1.20027 | -2.29782 | 0.000701 | 0.008638 | NAIPP1 |
| ENSG0000 | -1.63455 | -3.10491 | 2.22E-56 | 1.06E-52 | SELENOP |
| ENSG0000 | 1.225335 | 2.338097 | 5.77E-13 | 6.02E-11 | SHANK3 |
| ENSG0000 | 6.892875 | 118.8398 | 1.88E-05 | 0.000414 | None |
| ENSG0000 | 5.465254 | 44.17793 | 0.000106 | 0.001846 | None |
| ENSG0000 | -5.33438 | -40.3468 | 0.00571 | 0.044909 | RNU6-377P |
| ENSG0000 | 5.388492 | 41.88878 | 0.001622 | 0.017005 | RNU7-18P |
| ENSG0000 | 3.091463 | 8.523599 | 0.005844 | 0.045629 | SNORA74D |

| | | | | | |
|---|---|---|---|---|---|
| ENSG0000 | 5.35272 | 40.86292 | 0.003739 | 0.032537 | RNU4ATAC12P |
| ENSG0000 | -5.60496 | -48.67 | 0.006391 | 0.048892 | RNU6-703P |
| ENSG0000 | -1.71832 | -3.29053 | 8.27E-20 | 1.99E-17 | None |
| ENSG0000 | -4.83477 | -28.5372 | 0.002963 | 0.02722 | LINC00967 |
| ENSG0000 | -5.87732 | -58.7827 | 0.000254 | 0.003775 | None |
| ENSG0000 | -6.3666 | -82.5161 | 2.02E-06 | 5.97E-05 | None |
| ENSG0000 | -4.93325 | -30.5531 | 0.00535 | 0.042818 | None |
| ENSG0000 | 4.951649 | 30.94532 | 0.001147 | 0.012984 | USP12P1 |
| ENSG0000 | -1.03776 | -2.05304 | 0.00425 | 0.035836 | None |
| ENSG0000 | -6.23841 | -75.5004 | 0.000258 | 0.003821 | None |
| ENSG0000 | 1.11031 | 2.158921 | 3.74E-06 | 0.000102 | TRNP1 |
| ENSG0000 | -5.35319 | -40.8762 | 0.00043 | 0.005813 | None |
| ENSG0000 | 4.768715 | 27.26003 | 0.002602 | 0.024564 | None |
| ENSG0000 | -1.77877 | -3.43132 | 0.001518 | 0.0161 | None |
| ENSG0000 | -4.58303 | -23.9678 | 0.002261 | 0.022061 | None |
| ENSG0000 | -5.29203 | -39.1797 | 0.001309 | 0.014367 | None |
| ENSG0000 | -4.03801 | -16.4271 | 0.000949 | 0.011076 | None |
| ENSG0000 | 1.38528 | 2.612226 | 0.001358 | 0.014777 | ALG1L10P |
| ENSG0000 | 5.164692 | 35.86966 | 0.000593 | 0.007552 | None |
| ENSG0000 | -3.5456 | -11.6771 | 0.004114 | 0.034901 | None |
| ENSG0000 | 4.936937 | 30.63135 | 0.002093 | 0.020765 | LINC02749 |
| ENSG0000 | -5.07796 | -33.7767 | 0.002552 | 0.024173 | None |
| ENSG0000 | 6.163642 | 71.68713 | 0.000186 | 0.002911 | None |
| ENSG0000 | 1.188198 | 2.27868 | 1.45E-09 | 8.58E-08 | MEX3A |
| ENSG0000 | 1.391456 | 2.623433 | 0.002101 | 0.020793 | None |
| ENSG0000 | 4.556452 | 23.53037 | 0.006109 | 0.047134 | LINC02551 |
| ENSG0000 | -6.27594 | -77.4902 | 2.96E-06 | 8.37E-05 | LINC02705 |
| ENSG0000 | -5.31234 | -39.7352 | 0.000516 | 0.006755 | None |
| ENSG0000 | 1.249335 | 2.377318 | 7.51E-05 | 0.001376 | SNHG9 |
| ENSG0000 | 5.379484 | 41.62803 | 0.005973 | 0.046399 | DDX18P5 |
| ENSG0000 | -4.95935 | -31.111 | 0.001899 | 0.019195 | OR5BA1P |
| ENSG0000 | -1.32194 | -2.50002 | 5.77E-09 | 3.04E-07 | None |
| ENSG0000 | -5.56914 | -47.4765 | 0.003851 | 0.033227 | None |
| ENSG0000 | 1.033379 | 2.046812 | 8.95E-09 | 4.55E-07 | None |
| ENSG0000 | 5.240135 | 37.79529 | 0.005045 | 0.040975 | ELOCP31 |
| ENSG0000 | 1.149669 | 2.21863 | 0.000395 | 0.005413 | SBNO1-AS1 |
| ENSG0000 | 5.522391 | 45.96268 | 0.004643 | 0.038462 | RPL7AP3 |
| ENSG0000 | 1.074707 | 2.106295 | 0.001999 | 0.020018 | LINC02454 |
| ENSG0000 | 5.10159 | 34.33456 | 0.003655 | 0.031981 | None |
| ENSG0000 | 1.681904 | 3.208511 | 0.00143 | 0.015413 | SLC6A12-AS1 |
| ENSG0000 | 2.877061 | 7.346521 | 0.003556 | 0.031281 | None |
| ENSG0000 | -4.54548 | -23.3521 | 0.001252 | 0.013867 | None |
| ENSG0000 | -1.26315 | -2.40019 | 0.004921 | 0.040193 | NHLRC4 |
| ENSG0000 | -5.51969 | -45.8766 | 0.000395 | 0.005413 | |
| ENSG0000 | -5.1796 | -36.2421 | 0.002035 | 0.020312 | LINC02388 |
| ENSG0000 | 1.085197 | 2.121665 | 2.44E-08 | 1.14E-06 | MGAM |
| ENSG0000 | -2.7249 | -6.61113 | 0.001827 | 0.018626 | None |

| | | | | | |
|---|---|---|---|---|---|
| ENSG0000 | 5.271099 | 38.61524 | 0.001355 | 0.014755 | None |
| ENSG0000 | -4.61116 | -24.4398 | 0.005672 | 0.044765 | None |
| ENSG0000 | -5.32961 | -40.2136 | 0.006081 | 0.046997 | None |
| ENSG0000 | -5.6142 | -48.9826 | 0.00095 | 0.011079 | RPL36A-HNRNPH2 |
| ENSG0000 | 6.558182 | 94.23441 | 0.000356 | 0.004963 | None |
| ENSG0000 | 6.915729 | 120.7374 | 0.000277 | 0.004054 | None |
| ENSG0000 | -1.88072 | -3.68259 | 4.06E-08 | 1.79E-06 | OTOAP1 |
| ENSG0000 | -5.2006 | -36.7737 | 0.002265 | 0.022076 | LINC02404 |
| ENSG0000 | -6.03227 | -65.4478 | 0.000359 | 0.005004 | None |
| ENSG0000 | 5.925807 | 60.79189 | 0.004347 | 0.036466 | None |
| ENSG0000 | 1.1329 | 2.192991 | 3.81E-66 | 2.72E-62 | CLEC5A |
| ENSG0000 | -6.51401 | -91.3931 | 1.81E-07 | 6.88E-06 | None |
| ENSG0000 | 1.004739 | 2.00658 | 0.000211 | 0.003244 | LINC00641 |
| ENSG0000 | -3.55602 | -11.7617 | 0.005218 | 0.042019 | LINC01500 |
| ENSG0000 | -1.39941 | -2.63795 | 6.33E-09 | 3.28E-07 | LINC01629 |
| ENSG0000 | 5.774581 | 54.74219 | 0.000156 | 0.002534 | None |
| ENSG0000 | 6.356009 | 81.91232 | 1.14E-05 | 0.000268 | None |
| ENSG0000 | -2.91698 | -7.55266 | 0.001544 | 0.016306 | None |
| ENSG0000 | -1.2786 | -2.42603 | 0.002577 | 0.024377 | FPGT-TNNI3K |
| ENSG0000 | -1.22084 | -2.33082 | 0.003021 | 0.027627 | LINC00639 |
| ENSG0000 | -2.47965 | -5.57762 | 0.000275 | 0.004029 | None |
| ENSG0000 | 1.113881 | 2.164271 | 4.61E-06 | 0.000123 | None |
| ENSG0000 | -5.96512 | -62.4712 | 3.98E-05 | 0.00079 | DCAF13P3 |
| ENSG0000 | 1.808665 | 3.503179 | 0.000715 | 0.008781 | KIF23-AS1 |
| ENSG0000 | 5.180854 | 36.27376 | 0.001738 | 0.017937 | GCSHP2 |
| ENSG0000 | -4.87029 | -29.2484 | 0.002667 | 0.02507 | None |
| ENSG0000 | 2.165713 | 4.486881 | 0.004086 | 0.034734 | None |
| ENSG0000 | -4.54954 | -23.4179 | 0.004789 | 0.039427 | None |
| ENSG0000 | -5.51478 | -45.7208 | 0.000705 | 0.008676 | None |
| ENSG0000 | -5.33618 | -40.397 | 0.001298 | 0.01427 | None |
| ENSG0000 | -4.52963 | -23.0969 | 0.005639 | 0.044603 | MRPS21P7 |
| ENSG0000 | -5.57073 | -47.5289 | 0.000111 | 0.001909 | None |
| ENSG0000 | -5.44251 | -43.4868 | 0.001284 | 0.01414 | None |
| ENSG0000 | 1.307397 | 2.474946 | 8.77E-11 | 6.34E-09 | SNHG19 |
| ENSG0000 | 5.769161 | 54.53693 | 0.000437 | 0.005895 | TPRKBP2 |
| ENSG0000 | 1.064748 | 2.091805 | 0.006539 | 0.049788 | None |
| ENSG0000 | -5.33856 | -40.4639 | 0.000616 | 0.007797 | None |
| ENSG0000 | 6.017604 | 64.78573 | 0.002271 | 0.022123 | ATP5MFP6 |
| ENSG0000 | -5.33953 | -40.4909 | 0.000843 | 0.01011 | None |
| ENSG0000 | 5.60034 | 48.51437 | 0.001795 | 0.018353 | TNRC6B-DT |
| ENSG0000 | 4.829396 | 28.43107 | 0.002713 | 0.025431 | None |
| ENSG0000 | 4.661017 | 25.29915 | 0.003356 | 0.030002 | GOLGA8T |
| ENSG0000 | 4.859643 | 29.03342 | 0.001568 | 0.016527 | None |
| ENSG0000 | 1.551787 | 2.931801 | 2.29E-20 | 6.01E-18 | TPBGL |
| ENSG0000 | -5.78931 | -55.3039 | 0.00038 | 0.005242 | None |
| ENSG0000 | -5.12991 | -35.0153 | 0.005971 | 0.046397 | None |
| ENSG0000 | 1.341464 | 2.534084 | 0.000255 | 0.003786 | None |

| | | | | | |
|---|---|---|---|---|---|
| ENSG0000 | 6.445857 | 87.17585 | 6.99E-05 | 0.001296 | None |
| ENSG0000 | 1.012909 | 2.017976 | 0.005916 | 0.046069 | SPON1 |
| ENSG0000 | 5.686343 | 51.49437 | 0.001161 | 0.013121 | MRPS21P9 |
| ENSG0000 | -1.16465 | -2.24179 | 0.000717 | 0.008798 | MYZAP |
| ENSG0000 | 6.704561 | 104.2975 | 6.04E-05 | 0.001136 | RYKP1 |
| ENSG0000 | 4.864224 | 29.12576 | 0.003512 | 0.031052 | ABHD17AP5 |
| ENSG0000 | -4.15386 | -17.8007 | 0.004822 | 0.039654 | MAPK8IP1P2 |
| ENSG0000 | 1.521374 | 2.870644 | 0.00579 | 0.045314 | None |
| ENSG0000 | -5.81134 | -56.1551 | 0.000948 | 0.01107 | LINC02864 |
| ENSG0000 | 2.040965 | 4.115208 | 0.001229 | 0.013683 | None |
| ENSG0000 | -4.58983 | -24.0812 | 0.005217 | 0.042019 | None |
| ENSG0000 | -5.70169 | -52.045 | 1.99E-05 | 0.000435 | MIR3189 |
| ENSG0000 | -5.34237 | -40.5708 | 0.000419 | 0.005689 | None |
| ENSG0000 | 2.895102 | 7.438964 | 0.005249 | 0.042142 | MIR4648 |
| ENSG0000 | -4.88761 | -29.6017 | 0.001033 | 0.011885 | None |
| ENSG0000 | 6.655988 | 100.8444 | 0.002505 | 0.023831 | None |
| ENSG0000 | -4.90816 | -30.0264 | 0.003791 | 0.032899 | RN7SL45P |
| ENSG0000 | 5.047494 | 33.07098 | 0.003307 | 0.029703 | MIR5188 |
| ENSG0000 | 1.643795 | 3.124867 | 0.003205 | 0.028973 | None |
| ENSG0000 | -1.55757 | -2.94357 | 2.11E-09 | 1.21E-07 | FCGR1CP |
| ENSG0000 | 6.30176 | 78.88943 | 2.69E-06 | 7.69E-05 | None |
| ENSG0000 | -4.88585 | -29.5657 | 0.006255 | 0.048021 | None |
| ENSG0000 | 1.246738 | 2.373043 | 3.98E-17 | 7.12E-15 | None |
| ENSG0000 | 5.870919 | 58.52248 | 0.000397 | 0.005437 | None |
| ENSG0000 | 6.395593 | 84.19091 | 0.000295 | 0.004256 | None |
| ENSG0000 | -2.00607 | -4.01688 | 1.72E-09 | 1.01E-07 | PTGES3L |
| ENSG0000 | -6.21725 | -74.401 | 7.91E-06 | 0.000198 | None |
| ENSG0000 | -1.55005 | -2.92826 | 0.001639 | 0.017144 | KCNJ2-AS1 |
| ENSG0000 | 5.196712 | 36.67467 | 0.002206 | 0.021604 | None |
| ENSG0000 | -5.72369 | -52.8446 | 0.000876 | 0.010417 | LINC02073 |
| ENSG0000 | 2.895864 | 7.442894 | 0.000987 | 0.011437 | None |
| ENSG0000 | -4.51459 | -22.8574 | 0.0043 | 0.036159 | None |
| ENSG0000 | -3.64699 | -12.5272 | 0.006571 | 0.049958 | MIR4527HG |
| ENSG0000 | 1.392863 | 2.625994 | 0.00349 | 0.030906 | None |
| ENSG0000 | -5.53952 | -46.5118 | 0.006254 | 0.048021 | LINC01533 |
| ENSG0000 | 6.253083 | 76.2721 | 1.85E-05 | 0.00041 | GABRQ |
| ENSG0000 | -4.95088 | -30.9289 | 0.001145 | 0.012971 | None |
| ENSG0000 | -5.35726 | -40.9917 | 0.000107 | 0.00185 | None |
| ENSG0000 | -4.55014 | -23.4276 | 0.005783 | 0.045274 | EEF1A1P7 |
| ENSG0000 | 5.460453 | 44.03117 | 0.000681 | 0.00845 | AIRN |
| ENSG0000 | -5.8471 | -57.5644 | 4.10E-05 | 0.00081 | None |
| ENSG0000 | -5.50484 | -45.4068 | 0.000671 | 0.008356 | None |
| ENSG0000 | -5.69896 | -51.9465 | 0.000352 | 0.004929 | None |
| ENSG0000 | 1.156128 | 2.228585 | 2.98E-05 | 0.000616 | LINC01711 |
| ENSG0000 | 6.822839 | 113.2086 | 0.000273 | 0.004009 | None |
| ENSG0000 | 6.258014 | 76.53322 | 0.002776 | 0.025893 | RMRP |
| ENSG0000 | -2.80007 | -6.96475 | 0.001704 | 0.017662 | None |

| | | | | | |
|---|---|---|---|---|---|
| ENSG0000 | -2.50275 | -5.66763 | 1.14E-11 | 9.63E-10 | LINC01480 |
| ENSG0000 | 3.370095 | 10.3395 | 0.006532 | 0.049755 | HEATR9 |
| ENSG0000 | 5.66073 | 50.58823 | 0.000575 | 0.007379 | None |
| ENSG0000 | 7.397592 | 168.6153 | 3.70E-08 | 1.65E-06 | None |
| ENSG0000 | 4.427018 | 21.51122 | 0.002973 | 0.027304 | None |
| ENSG0000 | 6.513971 | 91.39044 | 6.82E-05 | 0.001268 | None |
| ENSG0000 | 5.391485 | 41.97578 | 0.006184 | 0.047602 | None |
| ENSG0000 | 1.248273 | 2.375569 | 0.003365 | 0.030036 | None |
| ENSG0000 | 1.374055 | 2.591981 | 0.000603 | 0.007658 | MIR4787 |
| ENSG0000 | -5.17318 | -36.0814 | 0.000272 | 0.003995 | None |
| ENSG0000 | 5.147026 | 35.43311 | 0.001456 | 0.015625 | None |
| ENSG0000 | -1.16012 | -2.23476 | 0.00493 | 0.040229 | C2orf15 |
| ENSG0000 | 1.074729 | 2.106327 | 0.001279 | 0.014092 | None |
| ENSG0000 | -3.8603 | -14.5233 | 0.002462 | 0.02352 | None |
| ENSG0000 | -5.89847 | -59.6507 | 2.61E-05 | 0.000547 | OR6L2P |
| ENSG0000 | -5.57034 | -47.5161 | 0.003403 | 0.030293 | SFTPD-AS1 |
| ENSG0000 | -3.66302 | -12.6672 | 0.001279 | 0.014092 | None |
| ENSG0000 | 3.775847 | 13.69756 | 0.006368 | 0.048741 | None |
| ENSG0000 | -5.38248 | -41.7147 | 0.000263 | 0.003881 | MIR7843 |
| ENSG0000 | 6.090623 | 68.14912 | 0.001223 | 0.013665 | H4C6 |
| ENSG0000 | 1.597378 | 3.025929 | 6.53E-05 | 0.001219 | None |
| ENSG0000 | -5.51824 | -45.8308 | 0.000461 | 0.006156 | CCL23 |
| ENSG0000 | 5.614329 | 48.98708 | 0.001291 | 0.01421 | MIR7152 |
| ENSG0000 | 4.479613 | 22.30991 | 0.00023 | 0.003486 | H2AC12 |
| ENSG0000 | 7.134152 | 140.4733 | 7.52E-05 | 0.001377 | H4C13 |
| ENSG0000 | 1.360821 | 2.568313 | 1.07E-07 | 4.28E-06 | LINC02340 |
| ENSG0000 | 5.143264 | 35.34083 | 0.006245 | 0.04798 | None |
| ENSG0000 | 4.774028 | 27.3606 | 0.001539 | 0.016269 | H3C11 |
| ENSG0000 | -1.08195 | -2.11689 | 4.21E-14 | 5.19E-12 | FCGBP |
| ENSG0000 | -4.90286 | -29.9163 | 0.001215 | 0.013587 | None |
| ENSG0000 | 1.491944 | 2.812678 | 0.000744 | 0.009068 | H2BC9 |
| ENSG0000 | -5.36499 | -41.2118 | 0.002187 | 0.021475 | OR7G15P |
| ENSG0000 | -5.43346 | -43.215 | 0.003069 | 0.027956 | None |
| ENSG0000 | 5.6646 | 50.72411 | 0.000419 | 0.005689 | None |
| ENSG0000 | 2.162599 | 4.477206 | 0.004788 | 0.039427 | DACH1 |
| ENSG0000 | 5.923058 | 60.67616 | 0.002014 | 0.020133 | Metazoa_SRP |
| ENSG0000 | 6.276534 | 77.522 | 0.003983 | 0.034095 | None |
| ENSG0000 | 5.721444 | 52.76262 | 0.000303 | 0.004346 | None |
| ENSG0000 | 1.25038 | 2.37904 | 0.005024 | 0.040834 | H4C5 |
| ENSG0000 | 5.860127 | 58.08636 | 0.000154 | 0.002506 | None |
| ENSG0000 | -4.60737 | -24.3757 | 0.004091 | 0.034765 | None |
| ENSG0000 | -4.88902 | -29.6307 | 0.001882 | 0.019075 | None |
| ENSG0000 | 7.930294 | 243.925 | 3.94E-07 | 1.39E-05 | H3C7 |
| ENSG0000 | 5.836062 | 57.12548 | 0.006068 | 0.046938 | None |
| ENSG0000 | 5.852852 | 57.79418 | 6.76E-05 | 0.001258 | RN7SL113P |
| ENSG0000 | -5.37148 | -41.3976 | 0.003873 | 0.033365 | None |
| ENSG0000 | 4.398193 | 21.08571 | 0.006008 | 0.046598 | H4C1 |

| | | | | | |
|---|---|---|---|---|---|
| ENSG0000 | 2.778074 | 6.859361 | 0.003164 | 0.02865 | None |
| ENSG0000 | 7.313641 | 159.0836 | 8.57E-06 | 0.000211 | None |
| ENSG0000 | 1.209858 | 2.313149 | 0.003671 | 0.032095 | None |
| ENSG0000 | 1.252143 | 2.381949 | 0.001112 | 0.012634 | None |
| ENSG0000 | 4.298897 | 19.68326 | 0.003006 | 0.027507 | None |
| ENSG0000 | 1.039303 | 2.055234 | 0.005589 | 0.044333 | None |
| ENSG0000 | 2.920459 | 7.570868 | 0.000767 | 0.009306 | None |
| ENSG0000 | -5.60448 | -48.6538 | 0.004495 | 0.037504 | None |
| ENSG0000 | -5.85264 | -57.7856 | 0.000394 | 0.005413 | None |
| ENSG0000 | 5.183597 | 36.34279 | 0.001493 | 0.015901 | None |
| ENSG0000 | 2.145912 | 4.425721 | 0.000968 | 0.011259 | None |
| ENSG0000 | 6.512853 | 91.3196 | 0.000805 | 0.009696 | None |
| ENSG0000 | -5.4482 | -43.6589 | 0.00219 | 0.021497 | None |
| ENSG0000 | 1.058225 | 2.082368 | 0.002782 | 0.025928 | None |
| ENSG0000 | -5.60609 | -48.7081 | 0.000112 | 0.001922 | None |
| ENSG0000 | 2.319955 | 4.993167 | 0.00069 | 0.00854 | |
| ENSG0000 | 5.846817 | 57.55293 | 0.004216 | 0.035612 | None |
| ENSG0000 | -5.7668 | -54.4477 | 0.00101 | 0.011661 | None |
| ENSG0000 | 1.285961 | 2.438444 | 0.000131 | 0.002203 | None |
| ENSG0000 | -1.79856 | -3.47872 | 5.32E-07 | 1.83E-05 | LINC01374 |
| ENSG0000 | -5.36627 | -41.2484 | 0.002847 | 0.026347 | None |
| ENSG0000 | 4.608047 | 24.3871 | 0.003029 | 0.027677 | None |
| ENSG0000 | 1.288101 | 2.442063 | 0.002834 | 0.026277 | SLFNL1-AS1 |
| ENSG0000 | 4.961141 | 31.14958 | 0.002952 | 0.027146 | DPRXP3 |
| ENSG0000 | -5.20349 | -36.8474 | 0.004337 | 0.036387 | None |
| ENSG0000 | -1.59157 | -3.01377 | 1.12E-70 | 1.07E-66 | ADORA3 |
| ENSG0000 | 5.675433 | 51.10642 | 0.001771 | 0.018152 | None |
| ENSG0000 | 5.059128 | 33.33876 | 0.002244 | 0.021913 | LINC02452 |

| GeneID | Treatment | Treatment | Treatment | Treatment | name |
|---|---|---|---|---|---|
| ENSG0000 | 1.237761 | 2.358323 | 5.22E-06 | 4.98E-05 | HOXA11 |
| ENSG0000 | 1.250518 | 2.379268 | 0.000197 | 0.001092 | CEACAM7 |
| ENSG0000 | 1.318781 | 2.494553 | 8.84E-06 | 7.79E-05 | ZPBP |
| ENSG0000 | -1.1475 | -2.21529 | 5.00E-10 | 2.40E-08 | UTS2 |
| ENSG0000 | 1.248558 | 2.376038 | 0.000188 | 0.001046 | SPO11 |
| ENSG0000 | -1.00863 | -2.012 | 4.26E-26 | 1.09E-22 | NTN1 |
| ENSG0000 | 1.378477 | 2.599937 | 3.06E-05 | 0.000224 | FAP |
| ENSG0000 | -1.04649 | -2.0655 | 3.26E-21 | 3.27E-18 | SLC1A3 |
| ENSG0000 | 1.324511 | 2.504479 | 0.002026 | 0.007958 | AFP |
| ENSG0000 | 1.107467 | 2.154671 | 0.004182 | 0.014651 | F11 |
| ENSG0000 | -1.0087 | -2.0121 | 1.08E-26 | 3.84E-23 | CCDC80 |
| ENSG0000 | 1.287116 | 2.440398 | 9.21E-06 | 8.05E-05 | GGTLC2 |
| ENSG0000 | -1.11646 | -2.16815 | 1.53E-44 | 4.91E-40 | SPARC |
| ENSG0000 | 1.00447 | 2.006207 | 0.011889 | 0.034904 | IL18RAP |
| ENSG0000 | 1.100873 | 2.144844 | 0.005582 | 0.018595 | CASQ2 |
| ENSG0000 | 1.014722 | 2.020514 | 0.010713 | 0.031991 | LYZL1 |
| ENSG0000 | 1.039689 | 2.055784 | 0.000443 | 0.00218 | FMOD |
| ENSG0000 | 1.049388 | 2.069652 | 0.011477 | 0.0339 | LAX1 |
| ENSG0000 | 1.386833 | 2.615041 | 0.0094 | 0.028706 | NME2P1 |
| ENSG0000 | 1.414728 | 2.666096 | 7.89E-05 | 0.000501 | FAM124B |
| ENSG0000 | 1.317002 | 2.491478 | 0.01105 | 0.032854 | MAGEA10 |
| ENSG0000 | 1.007036 | 2.009778 | 0.000718 | 0.003289 | XG |
| ENSG0000 | 1.390671 | 2.622006 | 0.002383 | 0.009105 | BANF2 |
| ENSG0000 | 1.309177 | 2.478002 | 0.003468 | 0.012527 | RNASE1 |
| ENSG0000 | -1.0359 | -2.0504 | 2.74E-20 | 2.21E-17 | PRRG3 |
| ENSG0000 | -1.3833 | -2.60864 | 2.28E-11 | 1.92E-09 | H19 |
| ENSG0000 | 1.043789 | 2.061635 | 0.001303 | 0.005466 | G6PC1 |
| ENSG0000 | 1.160604 | 2.23551 | 1.66E-05 | 0.000133 | KRT34 |
| ENSG0000 | 1.304325 | 2.469682 | 0.000307 | 0.001595 | USP29 |
| ENSG0000 | -1.01572 | -2.02191 | 7.85E-22 | 9.02E-19 | VSTM2L |
| ENSG0000 | 1.074411 | 2.105862 | 0.000468 | 0.002288 | RNF128 |
| ENSG0000 | 1.092865 | 2.132971 | 0.009879 | 0.029902 | GIMAP4 |
| ENSG0000 | 1.007661 | 2.010649 | 0.003026 | 0.011159 | CD180 |
| ENSG0000 | 1.08057 | 2.114871 | 2.45E-05 | 0.000186 | IL36RN |
| ENSG0000 | 1.214316 | 2.320307 | 9.93E-08 | 1.81E-06 | TPH2 |
| ENSG0000 | -1.2587 | -2.3928 | 0.004919 | 0.016722 | CTRL |
| ENSG0000 | 2.174031 | 4.512826 | 0.000398 | 0.001993 | S100A8 |
| ENSG0000 | 1.019576 | 2.027323 | 0.002772 | 0.010369 | ASB15 |
| ENSG0000 | 1.20675 | 2.308171 | 0.000189 | 0.00105 | RAB19 |
| ENSG0000 | -1.17638 | -2.26009 | 9.65E-11 | 6.40E-09 | GPC3 |
| ENSG0000 | 1.604866 | 3.041675 | 0.001781 | 0.007138 | ODF1 |
| ENSG0000 | 1.356038 | 2.559812 | 0.005482 | 0.018319 | LYZL4 |
| ENSG0000 | 1.139216 | 2.202613 | 0.004399 | 0.015255 | CD1B |
| ENSG0000 | 1.309443 | 2.478459 | 0.013676 | 0.039198 | PRAC1 |
| ENSG0000 | 1.108837 | 2.156717 | 0.000397 | 0.001988 | TFF1 |
| ENSG0000 | 1.017138 | 2.0239 | 0.000373 | 0.001887 | LY6K |

| | | | | | |
|---|---|---|---|---|---|
| ENSG0000 | 1.077984 | 2.111083 | 0.0016 | 0.006518 | S100A11 |
| ENSG0000 | 1.079293 | 2.113 | 0.006324 | 0.020635 | CTLA4 |
| ENSG0000 | 1.107669 | 2.154972 | 3.94E-05 | 0.000277 | FBXO40 |
| ENSG0000 | 1.014077 | 2.019611 | 2.82E-05 | 0.000209 | FBP1 |
| ENSG0000 | 1.119048 | 2.172036 | 0.017496 | 0.04797 | MAGEC3 |
| ENSG0000 | 1.236127 | 2.355653 | 0.013191 | 0.038061 | CCDC103 |
| ENSG0000 | 1.066866 | 2.094877 | 0.01145 | 0.033837 | PRR15L |
| ENSG0000 | -1.20679 | -2.30824 | 1.20E-37 | 1.93E-33 | IGF2 |
| ENSG0000 | 1.414288 | 2.665281 | 0.003457 | 0.01249 | PPDPFL |
| ENSG0000 | 1.017142 | 2.023905 | 0.016822 | 0.046447 | GOT1L1 |
| ENSG0000 | 1.13702 | 2.199262 | 0.0019 | 0.007546 | ELSPBP1 |
| ENSG0000 | 2.043646 | 4.122861 | 0.000149 | 0.000864 | S100Z |
| ENSG0000 | 1.005721 | 2.007947 | 0.008092 | 0.02533 | TEX37 |
| ENSG0000 | 1.111302 | 2.160406 | 1.48E-05 | 0.000121 | KRT2 |
| ENSG0000 | 1.124896 | 2.180858 | 6.61E-05 | 0.000431 | C11orf86 |
| ENSG0000 | 1.067365 | 2.095602 | 6.24E-05 | 0.000409 | SPATA3 |
| ENSG0000 | 1.040535 | 2.056991 | 0.011838 | 0.034785 | None |
| ENSG0000 | 1.337815 | 2.527682 | 0.013581 | 0.038977 | RPL35P9 |
| ENSG0000 | 1.193181 | 2.286564 | 0.003364 | 0.01222 | TEX36 |
| ENSG0000 | 1.009511 | 2.013228 | 0.004984 | 0.016912 | ANKS4B |
| ENSG0000 | 1.030391 | 2.042578 | 0.004194 | 0.014684 | OR10P1 |
| ENSG0000 | 1.038912 | 2.054677 | 0.000535 | 0.002562 | GOLGA8DP |
| ENSG0000 | 1.029225 | 2.040928 | 0.016311 | 0.045303 | None |
| ENSG0000 | 1.015587 | 2.021726 | 0.00232 | 0.008904 | OR10AC1 |
| ENSG0000 | 1.197505 | 2.293426 | 0.002549 | 0.009641 | KRT18P28 |
| ENSG0000 | -1.05879 | -2.08318 | 0.001256 | 0.005304 | ZFP42 |
| ENSG0000 | 1.075613 | 2.107618 | 0.000857 | 0.003824 | C12orf40 |
| ENSG0000 | 1.36852 | 2.582056 | 0.004521 | 0.015599 | OR51A9P |
| ENSG0000 | 1.036697 | 2.051526 | 0.000716 | 0.003281 | ANKRD30B |
| ENSG0000 | 1.432639 | 2.699401 | 0.013569 | 0.038959 | ADIPOQ |
| ENSG0000 | 1.029511 | 2.041333 | 0.001886 | 0.007501 | IFNL1 |
| ENSG0000 | 1.912995 | 3.7659 | 0.001505 | 0.006187 | OR2V2 |
| ENSG0000 | 1.095162 | 2.136371 | 0.001533 | 0.006281 | SPDYE4 |
| ENSG0000 | 1.224873 | 2.337348 | 0.00543 | 0.018171 | PSG9 |
| ENSG0000 | 1.586284 | 3.00275 | 2.37E-05 | 0.000181 | S100A7A |
| ENSG0000 | 1.113782 | 2.164122 | 5.64E-07 | 7.66E-06 | STING1 |
| ENSG0000 | 1.022273 | 2.031116 | 1.67E-06 | 1.90E-05 | TAFA3 |
| ENSG0000 | 1.397822 | 2.635035 | 0.006353 | 0.020715 | LCE3A |
| ENSG0000 | 1.227461 | 2.341546 | 0.013609 | 0.039041 | CCIN |
| ENSG0000 | 1.020552 | 2.028695 | 0.000655 | 0.003039 | SLC36A2 |
| ENSG0000 | 1.249927 | 2.378293 | 5.13E-05 | 0.000346 | POTEG |
| ENSG0000 | 1.685019 | 3.215447 | 0.00152 | 0.00624 | C2orf78 |
| ENSG0000 | 1.070308 | 2.099881 | 0.001397 | 0.005802 | ACTBP11 |
| ENSG0000 | 1.150283 | 2.219574 | 0.001011 | 0.004399 | NKAPL |
| ENSG0000 | 1.010154 | 2.014126 | 0.000613 | 0.002871 | BECN2 |
| ENSG0000 | 1.203461 | 2.302914 | 0.017239 | 0.047402 | PATE2 |
| ENSG0000 | 1.155534 | 2.227667 | 1.82E-05 | 0.000144 | HCAR1 |

| | | | | | |
|---|---|---|---|---|---|
| ENSG0000 | 1.15905 | 2.233104 | 0.010081 | 0.030407 | LINC00477 |
| ENSG0000 | 1.075059 | 2.106808 | 0.001575 | 0.006429 | MYL4 |
| ENSG0000 | -1.07307 | -2.10391 | 0.012764 | 0.037019 | MIR324 |
| ENSG0000 | 1.241533 | 2.364496 | 0.006121 | 0.020079 | RNY3P12 |
| ENSG0000 | 1.029286 | 2.041014 | 0.010438 | 0.031283 | None |
| ENSG0000 | 1.290523 | 2.446168 | 0.009111 | 0.02796 | HSD3B2 |
| ENSG0000 | 1.331384 | 2.51644 | 0.005663 | 0.018818 | None |
| ENSG0000 | 1.114137 | 2.164654 | 0.002687 | 0.010099 | PRAMEF20 |
| ENSG0000 | 1.699561 | 3.24802 | 0.00235 | 0.008999 | PRAMEF17 |
| ENSG0000 | 1.413015 | 2.66293 | 0.002197 | 0.008514 | PRAMEF9 |
| ENSG0000 | 1.087444 | 2.124973 | 0.001943 | 0.007687 | SPATA31A1 |
| ENSG0000 | -1.06034 | -2.08543 | 7.48E-08 | 1.43E-06 | CFI |
| ENSG0000 | 1.241775 | 2.364893 | 0.002537 | 0.009602 | KRTAP12-3 |
| ENSG0000 | 1.144234 | 2.210287 | 0.001256 | 0.005305 | IGLL3P |
| ENSG0000 | 1.418581 | 2.673225 | 0.000504 | 0.00243 | MIR183 |
| ENSG0000 | 1.128305 | 2.186018 | 0.01099 | 0.032716 | IGLV1-44 |
| ENSG0000 | 1.455551 | 2.742614 | 0.001417 | 0.005877 | TRBV4-1 |
| ENSG0000 | -1.06895 | -2.09791 | 9.99E-05 | 0.000613 | SNORD17 |
| ENSG0000 | 1.470739 | 2.771638 | 0.009001 | 0.027692 | HMGN2P8 |
| ENSG0000 | 1.133723 | 2.194242 | 0.006915 | 0.022236 | OR5S1P |
| ENSG0000 | 1.044521 | 2.062681 | 0.000672 | 0.003106 | None |
| ENSG0000 | 1.510438 | 2.848966 | 0.0055 | 0.01837 | STRADBP1 |
| ENSG0000 | 1.093171 | 2.133425 | 0.014492 | 0.041092 | None |
| ENSG0000 | 1.551184 | 2.930575 | 0.001483 | 0.006109 | RPSAP46 |
| ENSG0000 | 1.245572 | 2.371126 | 0.000891 | 0.003953 | RPS2P17 |
| ENSG0000 | 1.801388 | 3.485554 | 0.004155 | 0.014569 | None |
| ENSG0000 | 1.227408 | 2.34146 | 0.013288 | 0.03828 | PPIAP68 |
| ENSG0000 | 1.481635 | 2.79265 | 1.63E-06 | 1.86E-05 | KRT8P37 |
| ENSG0000 | 1.057941 | 2.081958 | 0.016525 | 0.045759 | NHP2P2 |
| ENSG0000 | 1.327907 | 2.510381 | 0.014168 | 0.040369 | None |
| ENSG0000 | 1.075627 | 2.107638 | 0.01038 | 0.031138 | DNAJB1P1 |
| ENSG0000 | 1.031701 | 2.044434 | 0.009074 | 0.027869 | ZNF99 |
| ENSG0000 | 1.327645 | 2.509926 | 0.014341 | 0.040746 | KRT18P38 |
| ENSG0000 | 1.106419 | 2.153106 | 0.014852 | 0.041922 | TPT1P5 |
| ENSG0000 | 1.301998 | 2.465702 | 0.002452 | 0.00933 | None |
| ENSG0000 | 1.045053 | 2.063442 | 0.015572 | 0.043552 | CTAGE16P |
| ENSG0000 | 1.155203 | 2.227157 | 0.001575 | 0.006429 | None |
| ENSG0000 | 1.149704 | 2.218683 | 1.38E-05 | 0.000114 | None |
| ENSG0000 | 1.115558 | 2.166787 | 0.003912 | 0.013849 | None |
| ENSG0000 | 1.207318 | 2.30908 | 0.007611 | 0.024103 | C1QBPP1 |
| ENSG0000 | 1.302698 | 2.466897 | 0.014965 | 0.042187 | FAM90A24P |
| ENSG0000 | 1.205289 | 2.305835 | 0.014557 | 0.041242 | LL22NC01-81G9.3 |
| ENSG0000 | 1.069152 | 2.098199 | 0.0033 | 0.012026 | None |
| ENSG0000 | 1.066571 | 2.094449 | 0.003424 | 0.012387 | FAM230B |
| ENSG0000 | 1.663289 | 3.167379 | 0.00029 | 0.001517 | FRG1JP |
| ENSG0000 | 1.450266 | 2.732585 | 0.015035 | 0.042334 | KRT18P3 |
| ENSG0000 | 1.416736 | 2.669808 | 0.007561 | 0.023961 | NBPF7P |

| | | | | | |
|---|---|---|---|---|---|
| ENSG0000 | 1.102387 | 2.147096 | 0.000623 | 0.002912 | |
| ENSG0000 | 1.022789 | 2.031843 | 0.008274 | 0.025821 | FAM8A6P |
| ENSG0000 | 1.314581 | 2.487302 | 0.015079 | 0.042438 | CNN3P1 |
| ENSG0000 | 1.050117 | 2.070698 | 0.007084 | 0.022686 | None |
| ENSG0000 | 1.049271 | 2.069484 | 0.016504 | 0.045716 | RPS20P2 |
| ENSG0000 | -1.22097 | -2.33104 | 0.006926 | 0.022267 | MIR1285-1 |
| ENSG0000 | 1.326183 | 2.507383 | 0.000928 | 0.004085 | DPP3P2 |
| ENSG0000 | 1.145214 | 2.21179 | 0.00458 | 0.015757 | None |
| ENSG0000 | 1.923217 | 3.792679 | 0.000805 | 0.003623 | None |
| ENSG0000 | 1.03542 | 2.049711 | 2.44E-07 | 3.84E-06 | MEG9 |
| ENSG0000 | 1.56401 | 2.956745 | 0.001449 | 0.005992 | None |
| ENSG0000 | 1.462095 | 2.755082 | 0.005444 | 0.018206 | NDUFB4P8 |
| ENSG0000 | 1.021652 | 2.030243 | 0.000189 | 0.001054 | None |
| ENSG0000 | 1.165464 | 2.243054 | 0.008777 | 0.027126 | LINC00691 |
| ENSG0000 | 1.0331 | 2.046417 | 0.001086 | 0.004672 | UBTFL8 |
| ENSG0000 | 1.703195 | 3.256214 | 2.35E-08 | 5.57E-07 | LINC01280 |
| ENSG0000 | 1.567064 | 2.963012 | 8.71E-05 | 0.000545 | None |
| ENSG0000 | 1.11549 | 2.166686 | 0.017288 | 0.047515 | None |
| ENSG0000 | 1.327304 | 2.509333 | 0.002284 | 0.008783 | CCND3P1 |
| ENSG0000 | 1.376699 | 2.596736 | 0.010675 | 0.031909 | RPL36P16 |
| ENSG0000 | 1.528393 | 2.884644 | 0.001065 | 0.004594 | MEP1AP4 |
| ENSG0000 | 1.066423 | 2.094234 | 0.0009 | 0.003988 | TUBB8P6 |
| ENSG0000 | 1.719437 | 3.293079 | 0.00843 | 0.026202 | LINC01627 |
| ENSG0000 | 1.374496 | 2.592773 | 0.001753 | 0.007042 | None |
| ENSG0000 | 1.131643 | 2.191081 | 0.003032 | 0.011179 | None |
| ENSG0000 | 1.326697 | 2.508277 | 0.012379 | 0.036098 | TUBB4BP2 |
| ENSG0000 | 1.378265 | 2.599556 | 0.001939 | 0.007671 | None |
| ENSG0000 | 1.071379 | 2.101441 | 0.002086 | 0.008153 | RPL3P7 |
| ENSG0000 | 1.12306 | 2.178084 | 0.00023 | 0.001247 | None |
| ENSG0000 | 1.277615 | 2.424378 | 0.002273 | 0.00875 | ASS1P11 |
| ENSG0000 | 1.420964 | 2.677644 | 0.00763 | 0.024154 | HBAP1 |
| ENSG0000 | 1.026676 | 2.037324 | 0.013432 | 0.038619 | None |
| ENSG0000 | 1.342875 | 2.536562 | 0.001424 | 0.005899 | LINC01504 |
| ENSG0000 | 1.483099 | 2.795487 | 0.00219 | 0.008493 | RPL35P5 |
| ENSG0000 | 1.301939 | 2.465601 | 0.006797 | 0.021919 | LINC01729 |
| ENSG0000 | 1.312788 | 2.484212 | 0.000256 | 0.001362 | None |
| ENSG0000 | 1.235388 | 2.354447 | 0.001558 | 0.006366 | PSG8-AS1 |
| ENSG0000 | 1.530701 | 2.889262 | 0.002011 | 0.007906 | None |
| ENSG0000 | 1.476547 | 2.782818 | 0.000253 | 0.001351 | LINC02794 |
| ENSG0000 | 1.147093 | 2.214672 | 0.002864 | 0.010663 | None |
| ENSG0000 | 1.317245 | 2.491898 | 0.0145 | 0.041111 | LINC02848 |
| ENSG0000 | 1.032328 | 2.045322 | 0.00871 | 0.026962 | OAZ1P1 |
| ENSG0000 | 1.125773 | 2.182184 | 0.006961 | 0.022369 | ZNF877P |
| ENSG0000 | 1.18942 | 2.28061 | 0.000631 | 0.002944 | None |
| ENSG0000 | -1.05575 | -2.0788 | 0.01464 | 0.041425 | None |
| ENSG0000 | 1.106025 | 2.152517 | 0.003532 | 0.012717 | None |
| ENSG0000 | 1.014367 | 2.020017 | 0.004851 | 0.016537 | YAE1-DT |

| | | | | | |
|---|---|---|---|---|---|
| ENSG0000 | 1.504117 | 2.83651 | 0.000544 | 0.002597 | CNN2P1 |
| ENSG0000 | 1.389349 | 2.619605 | 0.006321 | 0.020631 | None |
| ENSG0000 | 1.242023 | 2.365299 | 0.009072 | 0.027864 | RAB28P3 |
| ENSG0000 | 1.03228 | 2.045253 | 0.002123 | 0.008273 | HPN-AS1 |
| ENSG0000 | 1.381929 | 2.606166 | 0.010364 | 0.031105 | None |
| ENSG0000 | 1.078163 | 2.111346 | 0.000336 | 0.001722 | None |
| ENSG0000 | 1.309441 | 2.478455 | 0.01517 | 0.042639 | LINC02088 |
| ENSG0000 | 1.309191 | 2.478025 | 0.010501 | 0.031447 | None |
| ENSG0000 | 1.160938 | 2.236028 | 0.002397 | 0.009152 | None |
| ENSG0000 | 1.269889 | 2.41143 | 0.007165 | 0.022913 | POLR2MP1 |
| ENSG0000 | 1.174745 | 2.25753 | 0.007979 | 0.025047 | None |
| ENSG0000 | 1.094788 | 2.135817 | 0.005606 | 0.018658 | LINC01696 |
| ENSG0000 | 1.237777 | 2.358349 | 0.008073 | 0.025289 | None |
| ENSG0000 | 1.424163 | 2.683587 | 0.018103 | 0.049235 | None |
| ENSG0000 | 1.124719 | 2.180591 | 0.012935 | 0.03746 | TBX18-AS1 |
| ENSG0000 | 1.07598 | 2.108154 | 0.016053 | 0.044707 | PRELID1P6 |
| ENSG0000 | 1.44946 | 2.731058 | 0.000732 | 0.003344 | None |
| ENSG0000 | 1.46444 | 2.759564 | 0.002238 | 0.008643 | ITPKB-IT1 |
| ENSG0000 | 1.13683 | 2.198973 | 0.009841 | 0.02981 | CSTP1 |
| ENSG0000 | 1.167966 | 2.246947 | 0.002198 | 0.008515 | None |
| ENSG0000 | 1.462917 | 2.756652 | 0.003679 | 0.013176 | None |
| ENSG0000 | 1.807345 | 3.499976 | 0.002421 | 0.009231 | None |
| ENSG0000 | 1.313443 | 2.48534 | 0.002638 | 0.009936 | None |
| ENSG0000 | 1.479183 | 2.787908 | 0.000363 | 0.001843 | None |
| ENSG0000 | 1.257408 | 2.390658 | 0.005574 | 0.018572 | ST6GALNAC2P1 |
| ENSG0000 | 1.160202 | 2.234887 | 0.017272 | 0.047484 | LINC02607 |
| ENSG0000 | 1.173141 | 2.255021 | 0.007531 | 0.023879 | YY1P2 |
| ENSG0000 | 2.128778 | 4.373469 | 0.00125 | 0.00528 | RPS7P4 |
| ENSG0000 | 1.444038 | 2.720813 | 8.77E-05 | 0.000549 | POM121L8P |
| ENSG0000 | 1.052544 | 2.074184 | 0.004297 | 0.014966 | PES1P2 |
| ENSG0000 | 1.225315 | 2.338065 | 0.008341 | 0.025983 | LINC02829 |
| ENSG0000 | 1.54504 | 2.918121 | 0.007543 | 0.023908 | None |
| ENSG0000 | 1.202264 | 2.301005 | 0.011989 | 0.035137 | HLA-DRB6 |
| ENSG0000 | 1.267969 | 2.408223 | 0.002838 | 0.010578 | MIR5689HG |
| ENSG0000 | 1.459561 | 2.750247 | 5.32E-05 | 0.000357 | LINC02766 |
| ENSG0000 | 1.104889 | 2.150823 | 0.010544 | 0.031558 | PHKG1P4 |
| ENSG0000 | 1.491323 | 2.811466 | 0.005153 | 0.017382 | LINC01523 |
| ENSG0000 | 1.036907 | 2.051824 | 0.012617 | 0.036656 | None |
| ENSG0000 | 1.272982 | 2.416606 | 0.00055 | 0.00262 | MTND5P2 |
| ENSG0000 | 1.00705 | 2.009797 | 0.004302 | 0.01498 | KRT8P21 |
| ENSG0000 | 1.14728 | 2.214959 | 0.013987 | 0.039953 | ZNF492 |
| ENSG0000 | 1.140139 | 2.204023 | 0.012839 | 0.03721 | SLC25A38P1 |
| ENSG0000 | 1.566472 | 2.961795 | 0.00548 | 0.018316 | KRT18P26 |
| ENSG0000 | 1.266215 | 2.405297 | 0.015169 | 0.042639 | IMMTP1 |
| ENSG0000 | 1.461318 | 2.753598 | 0.006428 | 0.020913 | LINC02865 |
| ENSG0000 | 1.315954 | 2.48967 | 0.002196 | 0.00851 | TEX53 |
| ENSG0000 | 1.057293 | 2.081022 | 0.014084 | 0.04017 | DLEC1P1 |

| | | | | | |
|---|---|---|---|---|---|
| ENSG0000 | 1.123655 | 2.178984 | 0.00842 | 0.026176 | None |
| ENSG0000 | 1.119621 | 2.172899 | 0.000403 | 0.002013 | CLUHP5 |
| ENSG0000 | 1.210209 | 2.313711 | 0.000737 | 0.003362 | LINC01756 |
| ENSG0000 | 1.148705 | 2.217148 | 2.90E-05 | 0.000214 | NOS2P3 |
| ENSG0000 | 1.074561 | 2.106081 | 0.008353 | 0.02601 | LIMD1-AS1 |
| ENSG0000 | 1.217608 | 2.325608 | 0.008998 | 0.02769 | RGPD4-AS1 |
| ENSG0000 | 1.230806 | 2.34698 | 0.00104 | 0.004507 | None |
| ENSG0000 | 1.554561 | 2.937443 | 0.011593 | 0.034181 | None |
| ENSG0000 | 1.672195 | 3.186991 | 0.0091 | 0.02793 | None |
| ENSG0000 | 1.669472 | 3.180981 | 0.000186 | 0.001037 | None |
| ENSG0000 | 1.093257 | 2.133551 | 0.003985 | 0.014056 | None |
| ENSG0000 | 1.006871 | 2.009549 | 0.015174 | 0.042646 | None |
| ENSG0000 | 1.144719 | 2.211031 | 0.003498 | 0.012605 | TRMT1P1 |
| ENSG0000 | 1.088903 | 2.127123 | 0.006408 | 0.02086 | None |
| ENSG0000 | 1.226119 | 2.339369 | 4.21E-05 | 0.000292 | KRT8P10 |
| ENSG0000 | 1.48996 | 2.808811 | 0.001772 | 0.007105 | LINC00396 |
| ENSG0000 | 1.43103 | 2.696392 | 0.004428 | 0.015331 | GOT2P1 |
| ENSG0000 | 1.114538 | 2.165256 | 0.011057 | 0.03287 | PRKD3-DT |
| ENSG0000 | 1.118376 | 2.171024 | 0.005725 | 0.018987 | LARGE-IT1 |
| ENSG0000 | 1.007205 | 2.010013 | 0.002848 | 0.010608 | None |
| ENSG0000 | 1.139276 | 2.202704 | 0.003418 | 0.012375 | CNN2P10 |
| ENSG0000 | 1.183452 | 2.271196 | 0.008355 | 0.026013 | None |
| ENSG0000 | 1.183848 | 2.27182 | 0.010689 | 0.031937 | None |
| ENSG0000 | 1.177776 | 2.262278 | 0.002102 | 0.008201 | None |
| ENSG0000 | 1.158171 | 2.231743 | 0.008082 | 0.025309 | RSU1P2 |
| ENSG0000 | 1.235427 | 2.35451 | 0.010258 | 0.030829 | PRCPP1 |
| ENSG0000 | 1.155031 | 2.226892 | 0.012476 | 0.036334 | LINC01646 |
| ENSG0000 | 1.094956 | 2.136066 | 0.000268 | 0.00142 | LINC01656 |
| ENSG0000 | 1.329699 | 2.513502 | 0.012311 | 0.035935 | None |
| ENSG0000 | 1.086092 | 2.122982 | 0.00557 | 0.018563 | None |
| ENSG0000 | 1.046049 | 2.064867 | 0.002925 | 0.010846 | PHGR1 |
| ENSG0000 | 1.127823 | 2.185288 | 9.85E-05 | 0.000606 | LINC01271 |
| ENSG0000 | 1.153769 | 2.224944 | 0.01526 | 0.042847 | UBE2E2-DT |
| ENSG0000 | 1.163437 | 2.239904 | 0.003624 | 0.013002 | None |
| ENSG0000 | 1.166013 | 2.243907 | 0.015877 | 0.044296 | PTPRT-DT |
| ENSG0000 | 1.026574 | 2.03718 | 0.002338 | 0.008963 | ANAPC1P1 |
| ENSG0000 | 1.01963 | 2.0274 | 0.001228 | 0.005197 | LINC01865 |
| ENSG0000 | 1.155436 | 2.227517 | 0.001829 | 0.007301 | LINC02579 |
| ENSG0000 | 1.437445 | 2.708408 | 0.002444 | 0.009306 | None |
| ENSG0000 | 1.177816 | 2.26234 | 0.008383 | 0.026082 | None |
| ENSG0000 | 1.235762 | 2.355056 | 0.01782 | 0.048672 | PPP1R2P2 |
| ENSG0000 | 1.367755 | 2.580687 | 0.015023 | 0.042306 | None |
| ENSG0000 | 1.091809 | 2.131412 | 0.013536 | 0.038885 | LINC02765 |
| ENSG0000 | 1.174943 | 2.257839 | 0.015013 | 0.04229 | BIN2P1 |
| ENSG0000 | 1.319014 | 2.494955 | 0.015369 | 0.043085 | RAD1P1 |
| ENSG0000 | 1.251139 | 2.380293 | 0.01603 | 0.044653 | None |
| ENSG0000 | 1.00886 | 2.01232 | 0.000385 | 0.00194 | GULOP |

| | | | | | |
|---|---|---|---|---|---|
| ENSG0000 | 1.292377 | 2.449313 | 0.015108 | 0.042505 | LINC00676 |
| ENSG0000 | 1.065508 | 2.092906 | 0.001371 | 0.005709 | None |
| ENSG0000 | 1.224298 | 2.336418 | 0.018072 | 0.04917 | None |
| ENSG0000 | 1.223424 | 2.335002 | 0.008083 | 0.025309 | IFNA12P |
| ENSG0000 | 1.027049 | 2.037852 | 0.012888 | 0.03733 | CHCHD2P8 |
| ENSG0000 | 1.023185 | 2.032401 | 0.013942 | 0.039841 | LINC01055 |
| ENSG0000 | 1.500052 | 2.828529 | 1.62E-05 | 0.00013 | ZSWIM5P1 |
| ENSG0000 | 1.135038 | 2.196244 | 0.005855 | 0.019333 | PHACTR2-AS1 |
| ENSG0000 | 1.112126 | 2.16164 | 0.010838 | 0.032329 | KIF3AP1 |
| ENSG0000 | 1.194476 | 2.288617 | 0.003399 | 0.012321 | C10orf71-AS1 |
| ENSG0000 | 1.769967 | 3.410462 | 0.00027 | 0.001428 | ANKRD54P1 |
| ENSG0000 | 1.18828 | 2.278809 | 0.000263 | 0.001398 | KLF2P4 |
| ENSG0000 | 1.044532 | 2.062697 | 0.001097 | 0.004711 | EN2-DT |
| ENSG0000 | 1.462252 | 2.75538 | 0.000523 | 0.002511 | LINC02810 |
| ENSG0000 | 1.182286 | 2.269361 | 0.017947 | 0.048914 | LINC01010 |
| ENSG0000 | 1.528558 | 2.884974 | 0.002522 | 0.009558 | None |
| ENSG0000 | 1.555653 | 2.939667 | 0.007891 | 0.024815 | LINC01563 |
| ENSG0000 | 1.001646 | 2.002283 | 0.000412 | 0.002054 | DDX11L5 |
| ENSG0000 | 1.007313 | 2.010164 | 0.007801 | 0.02457 | PHKG1P3 |
| ENSG0000 | 1.133664 | 2.194153 | 4.15E-06 | 4.11E-05 | LINC01141 |
| ENSG0000 | 1.41606 | 2.668557 | 0.002185 | 0.008479 | None |
| ENSG0000 | 1.228322 | 2.342944 | 0.011828 | 0.03477 | GYG1P2 |
| ENSG0000 | 1.249638 | 2.377817 | 0.002288 | 0.008794 | PRKAR1B-AS1 |
| ENSG0000 | 1.230063 | 2.345773 | 0.003884 | 0.013766 | CARD11-AS1 |
| ENSG0000 | 1.069473 | 2.098666 | 0.000709 | 0.003256 | GPAT2P1 |
| ENSG0000 | 1.58991 | 3.010305 | 0.003771 | 0.013434 | None |
| ENSG0000 | 1.165958 | 2.243822 | 0.015278 | 0.042887 | None |
| ENSG0000 | 1.104301 | 2.149947 | 0.00289 | 0.010741 | None |
| ENSG0000 | -1.27677 | -2.42296 | 0.004971 | 0.01687 | RPL28P2 |
| ENSG0000 | 1.038381 | 2.053921 | 0.004039 | 0.014219 | SPATA31D2P |
| ENSG0000 | 1.682099 | 3.208945 | 2.81E-05 | 0.000209 | None |
| ENSG0000 | 1.45255 | 2.736913 | 0.007465 | 0.023713 | AKR1B1P4 |
| ENSG0000 | 1.471938 | 2.773943 | 0.007892 | 0.024815 | None |
| ENSG0000 | 1.220316 | 2.329978 | 0.001422 | 0.005896 | None |
| ENSG0000 | 1.720552 | 3.295626 | 0.001237 | 0.005233 | CRIP1P2 |
| ENSG0000 | 1.113164 | 2.163196 | 0.000927 | 0.004084 | None |
| ENSG0000 | 1.261453 | 2.397371 | 0.003764 | 0.013418 | IMP3P2 |
| ENSG0000 | 1.986755 | 3.963446 | 0.001297 | 0.005445 | None |
| ENSG0000 | 1.403379 | 2.645204 | 0.010393 | 0.031166 | PSMD12P1 |
| ENSG0000 | 1.16548 | 2.243078 | 0.011952 | 0.035051 | RPS19P3 |
| ENSG0000 | 1.947857 | 3.85801 | 0.001855 | 0.007388 | None |
| ENSG0000 | 1.186869 | 2.276582 | 0.017457 | 0.047876 | None |
| ENSG0000 | 1.1128 | 2.162649 | 0.014698 | 0.041566 | KRTAP4-7 |
| ENSG0000 | 1.141725 | 2.206446 | 0.009535 | 0.029049 | |
| ENSG0000 | 1.152976 | 2.223721 | 0.01044 | 0.031286 | WDR82P2 |
| ENSG0000 | 1.222027 | 2.332743 | 0.012153 | 0.035528 | LASTR |
| ENSG0000 | 1.520982 | 2.869864 | 0.00077 | 0.003487 | RPL13AP23 |

| | | | | | |
|---|---|---|---|---|---|
| ENSG0000 | 1.299085 | 2.460727 | 0.008407 | 0.026142 | RN7SL364P |
| ENSG0000 | 1.006289 | 2.008738 | 0.016633 | 0.046016 | WBP1LP1 |
| ENSG0000 | 1.132675 | 2.19265 | 0.011635 | 0.034293 | ITGB5-AS1 |
| ENSG0000 | 1.070787 | 2.100579 | 0.008514 | 0.02645 | None |
| ENSG0000 | 1.32448 | 2.504427 | 0.001098 | 0.004711 | MTHFD2P1 |
| ENSG0000 | 1.360824 | 2.568319 | 0.01551 | 0.043419 | None |
| ENSG0000 | 1.181849 | 2.268673 | 0.003207 | 0.011738 | WDR45P1 |
| ENSG0000 | 1.061317 | 2.086836 | 0.000576 | 0.002724 | None |
| ENSG0000 | 1.704056 | 3.258156 | 1.34E-06 | 1.58E-05 | LINC02753 |
| ENSG0000 | 1.358112 | 2.563495 | 0.008661 | 0.026839 | None |
| ENSG0000 | 1.164469 | 2.241508 | 0.000592 | 0.002788 | None |
| ENSG0000 | 1.166026 | 2.243928 | 1.93E-05 | 0.000151 | LINC01411 |
| ENSG0000 | 1.347867 | 2.545355 | 0.001163 | 0.004953 | AK4P2 |
| ENSG0000 | 1.910415 | 3.759173 | 0.00059 | 0.002779 | OR52V1P |
| ENSG0000 | 1.398913 | 2.637028 | 0.006519 | 0.02116 | KRT19P3 |
| ENSG0000 | 1.309373 | 2.478339 | 0.007661 | 0.024221 | HSPE1P10 |
| ENSG0000 | 1.050898 | 2.071819 | 0.003341 | 0.012154 | ITGA2-AS1 |
| ENSG0000 | 1.103232 | 2.148355 | 0.013865 | 0.039654 | LINC00536 |
| ENSG0000 | 1.90756 | 3.751741 | 0.00081 | 0.003641 | None |
| ENSG0000 | 1.183063 | 2.270583 | 0.016447 | 0.045593 | CRYZP2 |
| ENSG0000 | 1.530503 | 2.888865 | 0.000877 | 0.0039 | ZBED1P1 |
| ENSG0000 | 1.754865 | 3.374948 | 0.002841 | 0.010587 | CRLF3P2 |
| ENSG0000 | 1.695545 | 3.238992 | 0.000813 | 0.003654 | None |
| ENSG0000 | 1.158336 | 2.231999 | 0.003307 | 0.012044 | LINC02125 |
| ENSG0000 | 1.342346 | 2.535632 | 0.006897 | 0.02219 | KNOP1P5 |
| ENSG0000 | 1.046953 | 2.066162 | 0.002802 | 0.010457 | None |
| ENSG0000 | 1.122884 | 2.177819 | 0.002407 | 0.009183 | None |
| ENSG0000 | 1.223162 | 2.334578 | 0.001091 | 0.004687 | None |
| ENSG0000 | 1.164537 | 2.241613 | 5.32E-05 | 0.000357 | None |
| ENSG0000 | 1.461742 | 2.754408 | 8.54E-05 | 0.000536 | LINC01843 |
| ENSG0000 | 1.307698 | 2.475463 | 0.013676 | 0.039198 | MTCYBP35 |
| ENSG0000 | 1.321585 | 2.499405 | 0.001328 | 0.005557 | None |
| ENSG0000 | 1.060153 | 2.085152 | 7.48E-06 | 6.77E-05 | WWC2-AS2 |
| ENSG0000 | 1.195181 | 2.289735 | 0.003016 | 0.011127 | None |
| ENSG0000 | 1.197256 | 2.293031 | 0.001802 | 0.00721 | LINC02230 |
| ENSG0000 | 1.018348 | 2.025598 | 0.001016 | 0.004417 | C5orf64-AS1 |
| ENSG0000 | 1.440339 | 2.713847 | 0.002542 | 0.009619 | None |
| ENSG0000 | 1.158428 | 2.232141 | 0.01625 | 0.045152 | TRBV6-7 |
| ENSG0000 | 1.12497 | 2.18097 | 0.011354 | 0.0336 | LINC01845 |
| ENSG0000 | 1.159507 | 2.233811 | 0.006531 | 0.021194 | AFG3L2P1 |
| ENSG0000 | 1.26405 | 2.40169 | 0.01518 | 0.042657 | LINC02844 |
| ENSG0000 | 2.165021 | 4.48473 | 0.000593 | 0.00279 | None |
| ENSG0000 | 1.434511 | 2.702906 | 0.006775 | 0.021865 | MARK2P11 |
| ENSG0000 | 1.554004 | 2.93631 | 0.000533 | 0.002552 | IGLVV-66 |
| ENSG0000 | 1.019751 | 2.027569 | 0.009957 | 0.030094 | LINC02055 |
| ENSG0000 | 1.391578 | 2.623655 | 0.000811 | 0.003647 | None |
| ENSG0000 | 1.268712 | 2.409463 | 0.008351 | 0.026005 | None |

| | | | | | |
|---|---|---|---|---|---|
| ENSG0000 | 1.28569 | 2.437987 | 0.009722 | 0.029531 | |
| ENSG0000 | 1.07884 | 2.112337 | 1.35E-05 | 0.000111 | |
| ENSG0000 | 1.380792 | 2.604113 | 0.008011 | 0.02513 | None |
| ENSG0000 | 1.71661 | 3.286632 | 0.000561 | 0.002661 | None |
| ENSG0000 | 1.235436 | 2.354524 | 0.00052 | 0.002499 | LINC02752 |
| ENSG0000 | 1.055517 | 2.078463 | 0.008843 | 0.027308 | LINC02584 |
| ENSG0000 | 1.174391 | 2.256976 | 0.006037 | 0.01985 | None |
| ENSG0000 | -1.21473 | -2.32097 | 0.00055 | 0.002619 | ATP5PBP5 |
| ENSG0000 | 1.352298 | 2.553185 | 0.003451 | 0.012474 | None |
| ENSG0000 | 1.584948 | 2.99997 | 0.003484 | 0.012571 | None |
| ENSG0000 | 1.294482 | 2.45289 | 0.008786 | 0.027152 | None |
| ENSG0000 | 1.393036 | 2.626308 | 0.000715 | 0.003278 | LINC02690 |
| ENSG0000 | 1.853126 | 3.612823 | 4.73E-06 | 4.59E-05 | None |
| ENSG0000 | 1.055743 | 2.078789 | 0.013573 | 0.038965 | LY6G6E |
| ENSG0000 | 1.527093 | 2.882046 | 0.000649 | 0.003018 | None |
| ENSG0000 | 1.306196 | 2.472886 | 0.007313 | 0.023306 | None |
| ENSG0000 | 1.027294 | 2.038198 | 0.000119 | 0.000709 | GAPDH-DT |
| ENSG0000 | 1.080066 | 2.114132 | 0.014278 | 0.040608 | None |
| ENSG0000 | 1.18726 | 2.277198 | 0.008315 | 0.025925 | None |
| ENSG0000 | 1.021142 | 2.029525 | 0.01484 | 0.041894 | None |
| ENSG0000 | 1.30378 | 2.468749 | 0.009177 | 0.028132 | None |
| ENSG0000 | 1.027109 | 2.037936 | 0.009724 | 0.029535 | None |
| ENSG0000 | 1.416429 | 2.66924 | 0.014779 | 0.041751 | LINC01154 |
| ENSG0000 | 1.191269 | 2.283535 | 0.000971 | 0.004253 | None |
| ENSG0000 | 1.272876 | 2.416428 | 0.000228 | 0.001235 | KRT73-AS1 |
| ENSG0000 | 2.320579 | 4.995325 | 9.51E-06 | 8.29E-05 | None |
| ENSG0000 | 1.103289 | 2.148439 | 0.004649 | 0.015948 | KRT8P19 |
| ENSG0000 | 1.075846 | 2.107957 | 0.012208 | 0.035675 | None |
| ENSG0000 | 1.014671 | 2.020442 | 0.00177 | 0.007098 | AQP5-AS1 |
| ENSG0000 | 1.099057 | 2.142147 | 0.00487 | 0.016592 | LINC02356 |
| ENSG0000 | 1.543421 | 2.914849 | 0.004897 | 0.016661 | None |
| ENSG0000 | -1.13674 | -2.19883 | 0.014419 | 0.040929 | None |
| ENSG0000 | 1.442556 | 2.71802 | 0.006008 | 0.019763 | LINC02457 |
| ENSG0000 | 1.259841 | 2.394693 | 0.007706 | 0.024333 | KRT128P |
| ENSG0000 | 1.04559 | 2.06421 | 0.008751 | 0.027053 | None |
| ENSG0000 | 1.42481 | 2.684792 | 0.001026 | 0.004454 | CLUHP8 |
| ENSG0000 | 1.588749 | 3.007883 | 0.007022 | 0.022529 | None |
| ENSG0000 | 1.319343 | 2.495525 | 0.014106 | 0.040218 | None |
| ENSG0000 | 1.07096 | 2.100831 | 0.012663 | 0.036777 | BLZF2P |
| ENSG0000 | 1.092477 | 2.132398 | 0.004917 | 0.016718 | C20orf141 |
| ENSG0000 | 1.277908 | 2.424872 | 0.004184 | 0.014653 | None |
| ENSG0000 | 1.100131 | 2.143742 | 0.00214 | 0.008328 | KRT8P2 |
| ENSG0000 | 1.308198 | 2.47632 | 0.01508 | 0.042438 | None |
| ENSG0000 | 1.190431 | 2.282208 | 0.014426 | 0.040945 | COX5AP2 |
| ENSG0000 | 1.237437 | 2.357794 | 0.000242 | 0.001301 | None |
| ENSG0000 | 1.031728 | 2.044471 | 0.001984 | 0.007818 | LINC02285 |
| ENSG0000 | 1.343385 | 2.53746 | 0.001494 | 0.006149 | LINC01595 |

| | | | | | |
|---|---|---|---|---|---|
| ENSG0000 | 1.267887 | 2.408086 | 0.00447 | 0.015453 | LIPC-AS1 |
| ENSG0000 | -1.02903 | -2.04065 | 0.016892 | 0.0466 | None |
| ENSG0000 | 1.92199 | 3.789453 | 0.00043 | 0.002128 | GEMIN8P1 |
| ENSG0000 | 1.204688 | 2.304874 | 0.00115 | 0.00491 | KRT8P24 |
| ENSG0000 | 1.36499 | 2.575745 | 0.012116 | 0.035439 | None |
| ENSG0000 | 1.057359 | 2.081119 | 0.001275 | 0.005371 | |
| ENSG0000 | 1.137468 | 2.199945 | 0.009835 | 0.029799 | None |
| ENSG0000 | 1.237334 | 2.357624 | 0.000158 | 0.000907 | None |
| ENSG0000 | 1.413455 | 2.663744 | 0.016056 | 0.04471 | None |
| ENSG0000 | 1.048547 | 2.068445 | 0.000107 | 0.000651 | GOLGA2P11 |
| ENSG0000 | 1.127515 | 2.184821 | 0.000352 | 0.001797 | CA5AP1 |
| ENSG0000 | 1.096204 | 2.137914 | 0.009627 | 0.029291 | None |
| ENSG0000 | 1.160182 | 2.234857 | 0.000697 | 0.003206 | None |
| ENSG0000 | 1.085843 | 2.122615 | 0.000123 | 0.00073 | None |
| ENSG0000 | 1.300865 | 2.463766 | 5.60E-05 | 0.000373 | None |
| ENSG0000 | 1.268468 | 2.409057 | 0.002258 | 0.008702 | GEMIN8P2 |
| ENSG0000 | 1.429449 | 2.693439 | 0.008284 | 0.025843 | None |
| ENSG0000 | 1.171542 | 2.252523 | 0.000965 | 0.00423 | TLE7 |
| ENSG0000 | 1.205405 | 2.30602 | 0.004634 | 0.015909 | None |
| ENSG0000 | 1.683707 | 3.212523 | 0.000282 | 0.001481 | None |
| ENSG0000 | 1.668714 | 3.17931 | 0.017977 | 0.048958 | None |
| ENSG0000 | 1.149507 | 2.218381 | 5.09E-05 | 0.000344 | LINC01960 |
| ENSG0000 | 1.012709 | 2.017696 | 0.000177 | 0.000994 | None |
| ENSG0000 | 1.439406 | 2.712092 | 0.002832 | 0.01056 | None |
| ENSG0000 | 1.112186 | 2.16173 | 0.000144 | 0.000836 | None |
| ENSG0000 | 1.270548 | 2.412532 | 0.014806 | 0.04182 | None |
| ENSG0000 | 1.528966 | 2.885789 | 6.14E-05 | 0.000404 | CORO1A-AS1 |
| ENSG0000 | 1.148263 | 2.216469 | 0.006934 | 0.022287 | LINC00558 |
| ENSG0000 | 1.080159 | 2.11427 | 0.008892 | 0.027422 | SLC25A1P4 |
| ENSG0000 | 1.058555 | 2.082845 | 0.000591 | 0.002782 | None |
| ENSG0000 | 1.571417 | 2.971964 | 2.38E-05 | 0.000181 | LINC01996 |
| ENSG0000 | 1.702557 | 3.254773 | 0.0002 | 0.001107 | None |
| ENSG0000 | 1.025986 | 2.036351 | 0.000773 | 0.003501 | MAPK8IP1P1 |
| ENSG0000 | 1.332732 | 2.518792 | 0.00057 | 0.002699 | None |
| ENSG0000 | 1.125983 | 2.182502 | 4.51E-06 | 4.42E-05 | None |
| ENSG0000 | 1.685375 | 3.216239 | 0.000765 | 0.003469 | None |
| ENSG0000 | 1.127841 | 2.185315 | 0.009319 | 0.028496 | UBBP4 |
| ENSG0000 | 1.001466 | 2.002033 | 0.000792 | 0.003573 | None |
| ENSG0000 | 1.165568 | 2.243215 | 0.013367 | 0.038462 | MIR4456 |
| ENSG0000 | 1.042165 | 2.059316 | 3.89E-07 | 5.63E-06 | None |
| ENSG0000 | 1.152597 | 2.223137 | 0.013638 | 0.039108 | None |
| ENSG0000 | 1.463798 | 2.758335 | 0.000888 | 0.00394 | KRT18P8 |
| ENSG0000 | 1.069098 | 2.098121 | 0.006649 | 0.021505 | MIR4710 |
| ENSG0000 | 1.431504 | 2.697278 | 0.008053 | 0.025245 | None |
| ENSG0000 | 1.139371 | 2.202849 | 0.016963 | 0.04675 | |
| ENSG0000 | 1.393756 | 2.627618 | 0.000574 | 0.002711 | None |
| ENSG0000 | 1.305857 | 2.472306 | 0.002972 | 0.010992 | ARL2BPP1 |

| | | | | | |
|---|---|---|---|---|---|
| ENSG0000 | 1.772924 | 3.417459 | 0.003867 | 0.013722 | SLC9A3R1-AS1 |
| ENSG0000 | 1.359163 | 2.565363 | 0.001082 | 0.00466 | GRAMD4P7 |
| ENSG0000 | 1.088002 | 2.125794 | 0.001807 | 0.00723 | None |
| ENSG0000 | 1.131679 | 2.191136 | 0.017349 | 0.04766 | None |
| ENSG0000 | -1.2312 | -2.34762 | 0.010993 | 0.03272 | None |
| ENSG0000 | 1.446788 | 2.726005 | 0.008807 | 0.027214 | LINC01443 |
| ENSG0000 | 1.196705 | 2.292156 | 0.006063 | 0.019928 | IGBP1P2 |
| ENSG0000 | 1.608122 | 3.048547 | 0.00175 | 0.007033 | AKR1B1P7 |
| ENSG0000 | 1.294745 | 2.453336 | 8.28E-05 | 0.000522 | LINC01841 |
| ENSG0000 | 1.59665 | 3.024401 | 0.008726 | 0.026996 | FTLP5 |
| ENSG0000 | 1.206615 | 2.307954 | 6.71E-05 | 0.000436 | SCAT1 |
| ENSG0000 | 1.15062 | 2.220092 | 0.000926 | 0.00408 | None |
| ENSG0000 | 1.48457 | 2.798338 | 0.01369 | 0.039224 | None |
| ENSG0000 | 1.309933 | 2.4793 | 0.006082 | 0.019982 | SMCO4P1 |
| ENSG0000 | 1.173695 | 2.255887 | 0.002946 | 0.010916 | None |
| ENSG0000 | 1.092375 | 2.132247 | 0.001087 | 0.004677 | FAM215A |
| ENSG0000 | 1.006032 | 2.00838 | 0.014713 | 0.041601 | NFE2L3P1 |
| ENSG0000 | 1.000351 | 2.000486 | 0.010206 | 0.030712 | None |
| ENSG0000 | 1.250272 | 2.378862 | 0.003399 | 0.012321 | None |
| ENSG0000 | 1.154121 | 2.225487 | 2.19E-06 | 2.39E-05 | None |
| ENSG0000 | 1.51617 | 2.860308 | 0.003198 | 0.011713 | |
| ENSG0000 | 1.032776 | 2.045958 | 0.014952 | 0.042155 | None |
| ENSG0000 | 1.621089 | 3.076071 | 0.005098 | 0.017216 | None |
| ENSG0000 | 1.062207 | 2.088123 | 0.000787 | 0.003556 | ZSCAN5DP |
| ENSG0000 | 1.314625 | 2.487377 | 0.000397 | 0.001987 | LINC01785 |
| ENSG0000 | 1.434392 | 2.702682 | 0.011344 | 0.033572 | |
| ENSG0000 | 3.419482 | 10.69958 | 3.65E-05 | 0.000259 | None |
| ENSG0000 | 1.377356 | 2.597918 | 0.018095 | 0.049225 | None |
| ENSG0000 | 1.322859 | 2.501613 | 0.005766 | 0.019096 | None |
| ENSG0000 | 1.050986 | 2.071946 | 0.011312 | 0.033495 | SIGLEC27P |
| ENSG0000 | 1.32087 | 2.498168 | 0.004446 | 0.015386 | TP53TG3HP |
| ENSG0000 | 1.708044 | 3.267176 | 0.000343 | 0.001754 | LINC00221 |
| ENSG0000 | 1.506054 | 2.840322 | 0.001629 | 0.006622 | None |
| ENSG0000 | 1.720172 | 3.294757 | 0.001569 | 0.006405 | IGLCOR22-2 |
| ENSG0000 | 1.308235 | 2.476383 | 0.0167 | 0.04617 | IGLVIVOR22-1 |
| ENSG0000 | 1.05144 | 2.072598 | 0.003899 | 0.013812 | KRT18P9 |
| ENSG0000 | 1.516612 | 2.861184 | 0.005431 | 0.018172 | None |
| ENSG0000 | 1.199248 | 2.2962 | 0.00152 | 0.00624 | None |
| ENSG0000 | 1.232296 | 2.349406 | 0.010145 | 0.030579 | None |
| ENSG0000 | 1.271271 | 2.413741 | 0.017397 | 0.04775 | None |
| ENSG0000 | 1.35398 | 2.556163 | 4.04E-05 | 0.000282 | None |
| ENSG0000 | 1.43806 | 2.709562 | 2.45E-06 | 2.63E-05 | None |
| ENSG0000 | 1.007443 | 2.010345 | 0.000564 | 0.002674 | None |
| ENSG0000 | 1.223739 | 2.335513 | 0.004509 | 0.015566 | None |
| ENSG0000 | 1.005472 | 2.0076 | 0.004053 | 0.014257 | None |
| ENSG0000 | 1.014439 | 2.020117 | 0.005123 | 0.017295 | None |
| ENSG0000 | 1.273257 | 2.417067 | 0.007767 | 0.024493 | OR7M1P |

| | | | | | |
|---|---|---|---|---|---|
| ENSG0000 | 1.324573 | 2.504588 | 0.00047 | 0.002296 | MICE |
| ENSG0000 | 1.726136 | 3.308405 | 1.02E-05 | 8.76E-05 | None |
| ENSG0000 | 1.339588 | 2.530791 | 0.000124 | 0.000736 | None |
| ENSG0000 | 1.330137 | 2.514265 | 0.015444 | 0.043266 | None |
| ENSG0000 | 1.462833 | 2.75649 | 0.001006 | 0.004382 | GRAMD4P4 |
| ENSG0000 | 1.222812 | 2.334013 | 0.003169 | 0.011627 | None |
| ENSG0000 | 1.352159 | 2.55294 | 0.001785 | 0.007151 | TBC1D3I |
| ENSG0000 | 1.526608 | 2.881076 | 0.000238 | 0.001283 | None |
| ENSG0000 | 1.420997 | 2.677704 | 0.015048 | 0.042363 | MIR6753 |
| ENSG0000 | 1.163214 | 2.239557 | 0.004036 | 0.014213 | |
| ENSG0000 | 2.406997 | 5.303693 | 0.000543 | 0.002594 | MIR6809 |
| ENSG0000 | 1.272266 | 2.415406 | 0.000626 | 0.002921 | MT1IP |
| ENSG0000 | 1.29592 | 2.455335 | 0.004028 | 0.014187 | CCL15 |
| ENSG0000 | 1.298828 | 2.460289 | 0.018114 | 0.049245 | HTR1DP1 |
| ENSG0000 | 1.078775 | 2.112242 | 0.001765 | 0.007082 | None |
| ENSG0000 | 1.416752 | 2.669838 | 0.001521 | 0.006244 | None |
| ENSG0000 | 1.22406 | 2.336032 | 0.007698 | 0.024314 | None |
| ENSG0000 | 1.024664 | 2.034485 | 0.001667 | 0.006755 | None |
| ENSG0000 | 1.459667 | 2.750448 | 0.000284 | 0.001488 | LINC02227 |
| ENSG0000 | 1.020716 | 2.028926 | 0.004728 | 0.016176 | None |
| ENSG0000 | 1.754856 | 3.374925 | 0.000422 | 0.002096 | None |
| ENSG0000 | 1.005606 | 2.007787 | 0.000131 | 0.000771 | GOLGA8IP |
| ENSG0000 | 1.027011 | 2.037798 | 0.004744 | 0.016222 | None |
| ENSG0000 | 1.520601 | 2.869106 | 0.003183 | 0.011668 | Metazoa_SRP |
| ENSG0000 | 1.396801 | 2.633171 | 0.011845 | 0.034798 | FAM30B |
| ENSG0000 | 1.193351 | 2.286832 | 0.001214 | 0.005144 | None |
| ENSG0000 | 1.005352 | 2.007434 | 0.00628 | 0.020527 | None |
| ENSG0000 | -1.1771 | -2.26122 | 0.014028 | 0.040048 | None |
| ENSG0000 | -1.12446 | -2.1802 | 0.000469 | 0.002292 | None |
| ENSG0000 | 1.558279 | 2.945024 | 0.000525 | 0.002519 | None |
| ENSG0000 | 1.149348 | 2.218137 | 0.003151 | 0.011565 | None |
| ENSG0000 | 1.536754 | 2.901409 | 0.001604 | 0.006531 | None |
| ENSG0000 | 1.753246 | 3.371163 | 3.43E-05 | 0.000247 | None |
| ENSG0000 | 1.291544 | 2.447899 | 0.005116 | 0.017278 | None |
| ENSG0000 | 1.285874 | 2.438297 | 0.001367 | 0.005694 | None |
| ENSG0000 | 1.598891 | 3.029104 | 0.001451 | 0.005998 | None |
| ENSG0000 | -1.14303 | -2.20845 | 0.005913 | 0.019494 | None |
| ENSG0000 | 1.805914 | 3.496506 | 0.002015 | 0.007921 | None |
| ENSG0000 | 1.267419 | 2.407305 | 0.000732 | 0.003343 | None |
| ENSG0000 | 1.221517 | 2.331918 | 0.0146 | 0.041329 | LINC02033 |
| ENSG0000 | 1.386766 | 2.614919 | 0.000535 | 0.002562 | None |
| ENSG0000 | 1.287147 | 2.440449 | 0.001011 | 0.0044 | |
| ENSG0000 | 1.72885 | 3.314635 | 0.00019 | 0.001056 | None |
| ENSG0000 | 1.043137 | 2.060703 | 0.001326 | 0.005549 | None |
| ENSG0000 | 1.146938 | 2.214434 | 0.017756 | 0.048543 | None |
| ENSG0000 | 1.286822 | 2.4399 | 0.009523 | 0.02902 | PCAT5 |
| ENSG0000 | 1.01225 | 2.017055 | 0.008932 | 0.027525 | None |

| | | | | | |
|---|---|---|---|---|---|
| ENSG0000 | 1.103612 | 2.14892 | 0.009699 | 0.029471 | LINC01395 |
| ENSG0000 | 1.012512 | 2.017421 | 0.001458 | 0.006021 | TRG-AS1 |
| ENSG0000 | 1.166529 | 2.244709 | 0.004514 | 0.01558 | LSINCT5 |
| ENSG0000 | 1.907416 | 3.751365 | 2.77E-05 | 0.000206 | |
| ENSG0000 | 1.076081 | 2.108301 | 2.00E-05 | 0.000156 | LINC01230 |
| ENSG0000 | 1.148731 | 2.217187 | 0.001725 | 0.006945 | None |
| ENSG0000 | 1.089664 | 2.128245 | 0.003482 | 0.012565 | None |
| ENSG0000 | 1.382262 | 2.606768 | 0.007428 | 0.023613 | None |
| ENSG0000 | 1.136347 | 2.198237 | 0.009399 | 0.028705 | AGGF1P10 |
| ENSG0000 | 1.054714 | 2.077306 | 0.006734 | 0.021749 | MPP7-DT |
| ENSG0000 | 1.391885 | 2.624214 | 0.003797 | 0.013506 | |
| ENSG0000 | 1.62417 | 3.082647 | 0.003666 | 0.013132 | None |
| ENSG0000 | 1.778068 | 3.429665 | 2.91E-05 | 0.000214 | None |
