## Supplementary material for "Unravelling neuronal and glial differences in ceramide composition, synthesis, and sensitivity to toxicity": Pathway analysis MNs astrocytes microglia FB1 and GCSi

IPA Astrocytes FB1 FC2 adj0.05 1820

| | Ingenuity Canonical Pathways | -log(p-value) | Ratio | z-score | Molecules |
|---|---|---|---|---|---|
| 1 | ABRA Signaling Pathway | 9.21E+00 | 2.50E-01 | -2.711 | ACTA2,ACTB,ACTC1,ACTG2,CALD1,CCN2,EGR1,FOSB,GATA4,IGF1,MYH11,MYH9,MYL1,MYL9,PPARGC1A,PPP1R14A,SMTN,SRF,TAGLN,TPM1,TPM2,TPM4,VEGFA |
| 2 | Hepatic Fibrosis / Hepatic Stellate Cell Activation | 8.76E+00 | 1.75E-01 | NaN | A2M,ACTA2,CCL2,CCN2,COL12A1,COL21A1,COL26A1,COL5A1,COL5A3,COL6A6,COL9A1,EDN1,IGF1,IGF2,IGFBP3,IL1A,MYH10,MYH11,MYH2,MYH7,MYH9,MYL1,MYL6,MYL9,MYO1A,NGFR,PDGFB,PDGFD,PDGFRA,PDGFRB,TGFB3,TGFBR2,TLR4,VEGFA |
| 3 | Agranulocyte Adhesion and Diapedesis | 6.30E+00 | 1.48E-01 | NaN | ACKR3,ACTA2,ACTB,ACTC1,ACTG2,CCL2,CLDN19,CLDN5,CXCL10,CXCL12,CXCL14,IL1A,IL33,ITGB7,MMP10,MMP15,MMP17,MMP19,MMP24,MYH10,MYH11,MYH2,MYH7,MYH9,MYL1,MYL6,MYL9,MYO1A,PODXL,SELP,SELPLG |
| 4 | Pulmonary Fibrosis Idiopathic Signaling Pathway | 6.18E+00 | 1.26E-01 | -3.244 | ACTA2,ACTB,ACTC1,ACTG2,AREG,BBC3,CCN2,COL12A1,COL21A1,COL26A1,COL5A1,COL5A3,COL6A6,COL9A1,CXCL12,EDN1,EGR1,EIF4EBP1,EIF4EBP3,FGF18,FZD10,MMP10,MMP15,MMP17,MMP19,MMP24,PDGFB,PDGFD,PDGFRA,PDGFRB,PIK3CG,PIK3R3,PMAIP1,SRF,TGFB3,TGFBR2,THBS1,TP53,WNT1,WNT2B,WNT7B |
| 5 | Neuroinflammation Signaling Pathway | 6.08E+00 | 1.26E-01 | -0.87 | ATF4,BACE2,BIRC3,CASP1,CCL2,CD86,CNTF,CREB3L1,CXCL10,CXCL12,GABRA2,GABRA3,GABRA4,GABRB2,GABRG1,GABRQ,HLA-DMB,HLA-DQA1,HLA-DQB1,HMOX1,KCNJ3,KCNJ6,MAPK15,MAPK4,NCF2,NFATC2,P2RX7,PIK3CG,PIK3R3,PLA2G3,PLA2G4A,PLA2G4C,PLCG2,S100B,TGFB3,TGFBR2,TICAM2,TLR4,TLR5,WNT1 |
| 6 | Calcium Signaling | 5.40E+00 | 1.36E-01 | -0.333 | ACTA2,ACTC1,AKAP5,ATF4,CACNG4,CHRNA3,CHRNA9,CREB3L1,MYH10,MYH11,MYH2,MYH7,MYH9,MYL1,MYL6,MYL9,MYO1A,NFATC2,RCAN1,SLC8A2,SLC8A3,TNNC1,TNNI1,TNNI3,TNNT1,TNNT3,TPM1,TPM2,TPM4,TRPC4 |
| 7 | Osteoarthritis Pathway | 5.21E+00 | 1.31E-01 | 0 | ACAN,ADAMTS4,ALPL,ATF4,BMP2,CASP1,CASP10,CASP4,CASP5,CEBPB,CREB3L1,DDIT4,DDR2,DKK1,FGF18,FZD10,GREM1,ITGA9,ITGB3,ITGB7,MATN3,MMP10,P2RX7,PPARG,PPARGC1A,RUNX2,S1PR3,SPHK1,TGFBR2,TLR4,VEGFA |
| 8 | Ferroptosis Signaling Pathway | 4.98E+00 | 1.59E-01 | 0.655 | ANGPTL4,ATF4,CARS1,CBS/LOC102724560,CDKN1A,CHAC1,CTH,FTH1,FTL,H2BC26,HMGCR,HMOX1,LINC00472,SLC1A5,SLC38A1,SLC3A2,SLC7A11,SQSTM1,STEAP3,TP53,TXNRD1 |
| 9 | Axonal Guidance Signaling | 4.87E+00 | 1.02E-01 | NaN | ADAM19,ADAMTS12,ADAMTS16,ADAMTS2,ADAMTS4,ADAMTS8,ADAMTS9,BMP2,BMP4,CXCL12,EFNA5,EPHA1,EPHA4,EPHA6,EPHA8,EPHB6,FZD10,IGF1,ITGA9,ITGB3,ITGB7,L1CAM,MMP10,MMP15,MMP17,MMP24,MYL1,MYL6,MYL9,NFATC2,NGFR,NTN4,NTRK3,PAK3,PAPPA,PDGFB,PDGFD,PIK3CG,PIK3R3,PLCE1,PLCG2,SEMA3F,SHANK2,TUBB3,TUBB4B,TUBB6,UNC5A,UNC5B,VEGFA,WNT1,WNT2B,WNT7B |
| 10 | Superpathway of Cholesterol Biosynthesis | 4.79E+00 | 3.10E-01 | -3 | ACAT2,DHCR24,DHCR7,HMGCR,HMGCS1,IDI1,MSMO1,MVD,MVK |
| | Dilated Cardiomyopathy Signaling Pathway | 4.62E+00 | 1.47E-01 | 2.828 | ACTA2,ACTB,ACTC1,ACTG2,CACNG4,CNN1,MYH10,MYH11,MYH2,MYH7,MYH9,MYL1,MYL6,MYL9,MYO1A,PLN,TNNC1,TNNI1,TNNI3,TNNT1,TNNT3,TPM1 |
| | Myelination Signaling Pathway | 4.59E+00 | 1.13E-01 | -1.151 | ARHGAP6,ARHGAP8/PRR5-ARHGAP8,ARHGAP9,ATF4,BMP2,BMP4,CREB3L1,EIF4EBP1,FZD10,GPR17,HMGCR,IDI1,IGF1,IRS1,ITGB3,ITGB7,LAMB1,LMNB1,LYN,mir-23,MTHFD2,NFATC2,NGFR,OLIG2,PDGFB,PDGFD,PDGFRA,PDGFRB,PIK3CG,PIK3R3,PLCG2,PLP1,SLC16A1,SREBF1,WNT1,WNT2B,WNT7B |
| | DNA damage-induced 14-3-3σ Signaling | 4.53E+00 | 2.39E-01 | 2.714 | CASP1,CASP10,CASP4,CASP5,CCNB1,CCNB2,CCNE2,CDK1,CDKN1A,E2F1,TP53 |
| | tRNA Charging | 4.45E+00 | 2.56E-01 | 3.162 | CARS1,EPRS1,GARS1,IARS1,MARS1,NARS1,SARS1,TARS1,WARS1,YARS1 |
| | Phagosome Formation | 4.44E+00 | 9.21E-02 | -1.25 | ACKR3,ADGRA2,ADGRL3,ADRA1A,ADRB2,APLNR,CD36,CMKLR2,CNR1,COLEC12,F2RL2,FZD10,GAB2,GALR1,GPR137C,GPR157,GPR160,GPR17,GPR183,GPR37L1,GPRC5A,GRM2,HMOX1,HRH4,HTR7,ITGA9,ITGB3,ITGB7,LYN,MAPK15,MAPK4,MTNR1B,MYH10,MYH11,MYH2,MYH7,MYH9,MYL1,MYL6,MYL9,MYLK3,MYO1A,NPY1R,P2RY12,PAK3,PIK3CG,PIK3R3,PLA2G3,PLA2G4A,PLA2G4C,PLCG2,PLD5,PLD6,PNPLA3,PTGER4,RAPGEF4,S1PR1,S1PR3,SCARA3,SPHK1,TACR1,TLR4,TLR5,VAV1 |
| | Mevalonate Pathway I | 4.26E+00 | 4.29E-01 | -2.449 | ACAT2,HMGCR,HMGCS1,IDI1,MVD,MVK |
| | Role Of Osteoblasts In Rheumatoid Arthritis Signaling | 4.11E+00 | 1.19E-01 | -0.557 | ALPL,BMP2,BMP4,CNTF,CTSC,CTSV,CXCL12,DKK1,DKK2,FZD10,IGF1,IL17B,IL1A,IL33,MMP10,MMP15,MMP17,MMP19,MMP24,PIK3CG,PIK3R3,RUNX2,SFRP4,TGFB3,TNFSF9,VEGFA,WNT1,WNT2B,WNT7B |
| | Actin Cytoskeleton Signaling | 4.11E+00 | 1.19E-01 | -3.3 | ACTA2,ACTB,ACTC1,ACTG2,DIAPH3,FGD3,FGF13,FGF18,FGF7,FLNA,ITGA9,ITGB3,ITGB7,MYH10,MYH11,MYH2,MYH7,MYH9,MYL1,MYL6,MYL9,MYLK3,MYO1A,PAK3,PDGFB,PDGFD,PIK3CG,PIK3R3,VAV1 |
| | ILK Signaling | 3.94E+00 | 1.24E-01 | -2.4 | ACTA2,ACTB,ACTC1,ACTG2,ATF4,BMP2,CREB3L1,FLNA,IRS1,ITGB3,ITGB7,LIMS2,MYH10,MYH11,MYH2,MYH7,MYH9,MYL1,MYL6,MYL9,MYO1A,PIK3CG,PIK3R3,TGFB1I1,VEGFA |
| | Leukocyte Extravasation Signaling | 3.81E+00 | 1.24E-01 | -1.964 | ACTA2,ACTB,ACTC1,ACTG2,ARHGAP6,ARHGAP8/PRR5-ARHGAP8,ARHGAP9,CLDN19,CLDN5,CXCL12,MMP10,MMP15,MMP17,MMP19,MMP24,MYL6,NCF2,PIK3CG,PIK3R3,PLCG2,RAPGEF4,SELPLG,THY1,VAV1 |
| | Role Of Chondrocytes In Rheumatoid Arthritis Signalir | 3.58E+00 | 1.35E-01 | -1.147 | ADAMTS4,CASP1,CCL2,CEBPA,CEBPB,CEBPG,CXCL12,EGR1,IL1A,IL33,mir-23,MMP10,MMP15,MMP17,MMP19,MMP24,NGFR,RUNX2,VEGFA |
| | Superpathway of Geranylgeranyldiphosphate Biosynt | 3.55E+00 | 3.33E-01 | -2.449 | ACAT2,HMGCR,HMGCS1,IDI1,MVD,MVK |
| | Tight Junction Signaling | 3.47E+00 | 1.23E-01 | NaN | ACTA2,ACTB,ACTC1,ACTG2,ARHGEF2,CEBPA,CLDN19,CLDN5,F2RL2,MYH10,MYH11,MYH2,MYH7,MYH9,MYL1,MYL6,MYL9,MYO1A,NGFR,TGFB3,TGFBR2,TJP2 |
| | RHOA Signaling | 3.35E+00 | 1.37E-01 | -2.673 | ACTA2,ACTB,ACTC1,ACTG2,ANLN,ARHGAP6,ARHGAP8/PRR5-ARHGAP8,ARHGAP9,CIT,EPHA1,IGF1,MYL1,MYL6,MYL9,MYLK3,PLEKHG5,SEMA3F |
| | S100 Family Signaling Pathway | 3.17E+00 | 8.29E-02 | -1.25 | ACKR3,ADGRA2,ADGRL3,ADRA1A,ADRB2,AHNAK,APLNR,AREG,ATF4,CACNG4,CD36,CDKN1A,CMKLR2,CNR1,CREB3L1,F2RL2,FZD10,GALR1,GPR137C,GPR157,GPR160,GPR17,GPR183,GPR37L1,GPRC5A,GRM2,HBEGF,HRH4,HTR7,MAPK15,MAPK4,MCAM,MMP10,MMP15,MMP17,MMP19,MMP24,MTNR1B,MYH9,NCF2,NPY1R,P2RY12,PIK3CG,PIK3R3,PLA2G3,PLA2G4A,PLA2G4C,PLCE1,PLCG2,PTGER4,S100B,S1PR1,S1PR3,TACR1,TGFB3,TLR4,TP53,TPM1,TPM2,TPM4,VEGFA,WNT1,WNT2B,WNT7B |
| | Pathogen Induced Cytokine Storm Signaling Pathway | 3.16E+00 | 9.70E-02 | -2 | ADRA1A,CASP1,CCL2,CNTF,COL12A1,COL21A1,COL26A1,COL5A1,COL5A3,COL6A6,COL9A1,CXCL10,CXCL12,CXCL14,FGB,FTH1,FTL,HLA-DMB,HLA-DQA1,HLA-DQB1,HMGCR,IL17B,IL1A,IL33,MAPK15,MAPK4,NGFR,P2RX7,STX11,STXBP2,TGFB3,TLR4,TLR5,TNFRSF10B,TNFSF9,VEGFA |
| | ID1 Signaling Pathway | 3.15E+00 | 1.14E-01 | -1.877 | ATF3,BMP2,BMP4,CCN2,CDKN1A,CDKN2A,CHRNA3,CHRNA9,E2F1,EGR1,IGF2,KIF11,LYN,MCAM,NGFR,PIK3CG,PIK3R3,PTGER4,RNF125,TGFB3,TGFBR2,TP53,VEGFA |
| | Macrophage Alternative Activation Signaling Pathway | 3.14E+00 | 1.16E-01 | 1.706 | ATF4,CD36,CEBPB,CREB3L1,EIF4EBP1,HLA-DMB,HLA-DQA1,HLA-DQB1,IL1A,IL33,IRS1,KDM6B,KLF4,PIK3CG,PIK3R3,PPARG,PPARGC1A,SREBF1,TGFB3,THBS1,TLR4,TSC22D3 |
| | Tumor Microenvironment Pathway | 3.07E+00 | 1.17E-01 | -1.528 | CCL2,CXCL12,FGF13,FGF18,FGF7,IGF1,IGF2,ITGB3,MMP10,MMP15,MMP17,MMP19,MMP24,PDGFB,PDGFD,PIK3CG,PIK3R3,SLC16A1,TGFB3,TSLP,VEGFA |
| | Ephrin A Signaling | 3.04E+00 | 1.91E-01 | NaN | EFNA5,EPHA1,EPHA4,EPHA6,EPHA8,NGFR,PIK3CG,PIK3R3,VAV1 |
| | p53 Signaling | 3.04E+00 | 1.43E-01 | 2.309 | BBC3,CCNG1,CDKN1A,CDKN2A,E2F1,GADD45B,PIK3CG,PIK3R3,PMAIP1,SERPINE2,THBS1,TNFRSF10A,TNFRSF10B,TP53 |
| | HIF1α Signaling | 2.95E+00 | 1.11E-01 | -0.626 | ADM,CDKN1A,EDN1,EIF4EBP1,HK2,HMOX1,HSPA2,HSPA9,IGF1,IGF2,MKNK2,MMP10,MMP15,MMP17,MMP19,MMP24,NCF2,PDGFB,PIK3CG,PIK3R3,PLCG2,TP53,VEGFA |
| | Hepatic Fibrosis Signaling Pathway | 2.95E+00 | 9.22E-02 | -2.058 | ACTA2,ATF4,CACNG4,CCL2,CCN2,CEBPB,CNR1,COL5A3,CREB3L1,EDN1,FTH1,FTL,FZD10,IL1A,IL33,ITGA9,ITGB3,ITGB7,MYL1,MYL6,MYL9,MYLK3,NCF2,NGFR,PDGFB,PDGFD,PDGFRA,PDGFRB,PIK3CG,PIK3R3,PLCG2,PPARG,TGFB3,TGFBR2,TLR4,VEGFA,WNT1,WNT2B,WNT7B |
| | Glioblastoma Multiforme Signaling | 2.94E+00 | 1.17E-01 | -2.668 | CCND3,CCNE2,CDKN1A,CDKN2A,E2F1,FZD10,IGF1,IGF2,PDGFB,PDGFD,PDGFRA,PDGFRB,PIK3CG,PIK3R3,PLCE1,PLCG2,TP53,WNT1,WNT2B,WNT7B |
| | STAT3 Pathway | 2.93E+00 | 1.26E-01 | -1.069 | CDKN1A,GHR,IFNLR1,IGF1,IL17RE,IL1A,IL22RA1,IL27RA,IL2RB,IL31RA,NGFR,NTRK3,PDGFB,PDGFRA,PDGFRB,TGFBR2,VEGFA |
| | Endocannabinoid Cancer Inhibition Pathway | 2.93E+00 | 1.22E-01 | 2.183 | ATF3,ATF4,CASP1,CASP10,CASP4,CASP5,CCND3,CCNE2,CDKN1A,CNR1,CREB3L1,DDIT3,NUPR1,PIK3CG,PIK3R3,TRIB3,TWIST1,VEGFA |
| | Unfolded protein response | 2.91E+00 | 1.44E-01 | 2.309 | ATF4,CEBPA,CEBPB,CEBPG,DDIT3,DNAJB13,ERN1,HSPA2,HSPA9,PPARG,PPP1R15A,SREBF1,XBP1 |
| | Pulmonary Healing Signaling Pathway | 2.85E+00 | 1.11E-01 | -1.706 | ACKR3,BMP4,CCNB1,CXCL12,FGF7,FOXM1,FZD10,HBEGF,LYN,MMP10,MMP15,MMP17,MMP19,MMP24,NGFR,TGFB3,THBS1,TLR4,VEGFA,WNT1,WNT2B,WNT7B |
| | GADD45 Signaling | 2.83E+00 | 1.67E-01 | 1.897 | CCNB1,CCND3,CCNE2,CDK1,CDKN1A,GADD45B,mir-130,TGFB3,TGFBR2,TP53 |
| | Dopamine Degradation | 2.83E+00 | 2.19E-01 | 1.134 | ALDH2,ALDH3A1,ALDH4A1,MAOB,SMOX,SULT1A1,SULT1C2 |
| | Superpathway of Serine and Glycine Biosynthesis I | 2.82E+00 | 6.00E-01 | NaN | PHGDH,PSAT1,SHMT2 |
| | Wound Healing Signaling Pathway | 2.81E+00 | 1.03E-01 | -1.569 | ACTA2,CALCA,CEBPB,CNTF,COL12A1,COL21A1,COL26A1,COL5A1,COL5A3,COL6A6,COL9A1,FGF7,HBEGF,IL17B,IL1A,IL33,LAMB1,MMP10,NGFR,PDGFB,PDGFD,TAC1,TGFB3,TGFBR2,TNFSF9,VEGFA |
| | Growth Hormone Signaling | 2.79E+00 | 1.55E-01 | -1 | A2M,CEBPA,GHR,IGF1,IGF2,IGFBP3,IRS1,PIK3CG,PIK3R3,PLCG2,SRF |
| | Pyroptosis Signaling Pathway | 2.78E+00 | 1.40E-01 | 0.832 | CASP1,CASP4,CASP5,GAS5,GBP1,GBP2,GBP5,IL1A,NGFR,P2RX7,PTGER4,TLR4,TLR5 |
| | Sphingosine-1-phosphate Signaling | 2.62E+00 | 1.25E-01 | -2.324 | CASP1,CASP10,CASP4,CASP5,PDGFB,PDGFD,PDGFRA,PDGFRB,PIK3CG,PIK3R3,PLCE1,PLCG2,S1PR1,S1PR3,SPHK1 |
| | Apelin Cardiomyocyte Signaling Pathway | 2.53E+00 | 1.31E-01 | -1.387 | APLNR,MAPK15,MAPK4,MYL1,MYL6,MYL9,PIK3CG,PIK3R3,PLCE1,PLCG2,PLN,SLC8A2,SLC8A3 |
| | Cysteine Biosynthesis/Homocysteine Degradation | 2.52E+00 | 1.00E+00 | NaN | CBS/LOC102724560,CTH |
| | Th2 Pathway | 2.46E+00 | 1.17E-01 | 1.5 | CD86,HLA-DMB,HLA-DQA1,HLA-DQB1,ICOSLG/LOC102723996,IKZF1,IL2RB,IL33,JAG2,NFATC2,PIK3CG,PIK3R3,S1PR1,TGFBR2,TSLP,VAV1 |
| | Glioma Signaling | 2.45E+00 | 1.20E-01 | -1.667 | CCND3,CDKN1A,CDKN2A,CDKN2C,E2F1,IGF1,IGF2,PDGFB,PDGFD,PDGFRA,PDGFRB,PIK3CG,PIK3R3,PLCG2,TP53 |
| | Cellular Effects of Sildenafil (Viagra) | 2.44E+00 | 1.13E-01 | NaN | ACTA2,ACTB,ACTC1,ACTG2,CACNG4,MYH10,MYH11,MYH2,MYH7,MYH9,MYL1,MYL6,MYL9,MYO1A,PLCE1,PLCG2,SLC4A5 |
| | Iron homeostasis signaling pathway | 2.42E+00 | 1.16E-01 | NaN | ALAS2,ATP6V0D2,BMP2,BMP4,CP,ERFE,FTH1,FTL,GDF15,HMOX1,HSPA9,PDGFB,PDGFRA,PDGFRB,STEAP3,TFR2 |
| | Cardiac Hypertrophy Signaling (Enhanced) | 2.41E+00 | 8.30E-02 | -0.324 | ADRA1A,ADRB2,CACNG4,CNTF,DIAPH3,EDN1,EIF4EBP1,FGF13,FGF18,FGF7,FZD10,GATA4,GHR,IFNLR1,IGF1,IL17B,IL17RE,IL1A,IL22RA1,IL27RA,IL2RB,IL31RA,IL33,ITGA9,ITGB3,ITGB7,MAP3K15,MKNK2,NFATC2,NGFR,NKX2-5,PIK3CG,PIK3R3,PLCE1,PLCG2,PLD6,PLN,RCAN1,SRF,TGFB3,TGFBR2,TNFSF9,WNT1,WNT2B,WNT7B |
| | Serotonin Receptor Signaling | 2.40E+00 | 8.53E-02 | -1.897 | ADRA1A,ADRB2,ATF4,BST1,CACNG4,CCNE2,CREB3L1,EDN1,EIF4EBP1,FGB,HMOX1,HTR7,IL33,ITGB3,KCNA5,KCNJ3,KCNJ6,LYN,MAOB,MYLK3,PAK3,PIK3CG,PIK3R3,PLA2G3,PLA2G4A,PLA2G4C,PLCE1,PLCG2,PLD5,PLD6,PLN,PNPLA3,PPARGC1A,SMOX,SQSTM1,SRF,TGFB3,TNNI1,TNNI3,TRPC4 |
| | MSP-RON Signaling Pathway | 2.39E+00 | 1.55E-01 | NaN | ACTA2,ACTB,ACTC1,ACTG2,CCL2,KLK5,PIK3CG,PIK3R3,TLR4 |
| | GP6 Signaling Pathway | 2.38E+00 | 1.18E-01 | -1.604 | COL12A1,COL21A1,COL26A1,COL5A1,COL5A3,COL6A6,COL9A1,FGB,ITGB3,LAMB1,LYN,PIK3CG,PIK3R3,PLCG2,VAV1 |
| | Senescence Pathway | 2.37E+00 | 9.36E-02 | 0.2 | ASXL3,ATF3,CACNG4,CAPN6,CCNB1,CCNB2,CCND3,CCNE2,CDK1,CDKN1A,CDKN2A,CEBPB,DHCR24,E2F1,EIF4EBP1,GADD45B,GATA4,IL1A,MAPK15,MAPK4,NFATC2,PDK4,PIK3CG,PIK3R3,SQSTM1,TGFB3,TGFBR2,TP53 |
| | Factors Promoting Cardiogenesis in Vertebrates | 2.35E+00 | 1.11E-01 | -1 | ATF4,BMP2,BMP4,CREB3L1,DKK1,FZD10,GATA4,MYH7,NKX2-5,PLCE1,PLCG2,TDGF1,TGFB3,TGFBR2,WNT1,WNT2B,WNT7B |
| | Bladder Cancer Signaling | 2.34E+00 | 1.21E-01 | NaN | CDKN1A,CDKN2A,E2F1,FGF13,FGF18,FGF7,MMP10,MMP15,MMP17,MMP19,MMP24,THBS1,TP53,VEGFA |
| | Inhibition of Matrix Metalloproteases | 2.32E+00 | 1.79E-01 | 1.134 | A2M,MMP10,MMP15,MMP17,MMP19,MMP24,TFPI2 |
| | Trehalose Degradation II (Trehalase) | 2.31E+00 | 4.29E-01 | NaN | HK2,HKDC1,TREH |
| | Role of Macrophages, Fibroblasts and Endothelial Cel | 2.28E+00 | 9.04E-02 | NaN | ADAMTS4,ATF4,CCL2,CEBPA,CEBPB,CEBPG,CREB3L1,CXCL12,DKK1,DKK2,FZD10,IL16,IL1A,IL32,IL33,NFATC2,NGFR,PDGFB,PDGFD,PIK3CG,PIK3R3,PLCE1,PLCG2,SFRP4,TLR4,TLR5,VEGFA,WNT1,WNT2B,WNT7B |
| | Fcγ Receptor-mediated Phagocytosis in Macrophages | 2.28E+00 | 1.28E-01 | 0.302 | ACTA2,ACTB,ACTC1,ACTG2,GAB2,HMOX1,LYN,PIK3CG,PIK3R3,PLD5,PLD6,VAV1 |
| | Cell Cycle: G2/M DNA Damage Checkpoint Regulation | 2.26E+00 | 1.60E-01 | 2.121 | CCNB1,CCNB2,CDK1,CDKN1A,CDKN2A,PKMYT1,TOP2A,TP53 |
| | Phenylalanine Degradation IV (Mammalian, via Side C | 2.24E+00 | 2.86E-01 | 1 | ALDH2,GOT1,MAOB,SMOX |
| | Putrescine Degradation III | 2.23E+00 | 2.27E-01 | 0.447 | ALDH2,ALDH3A1,ALDH4A1,MAOB,SMOX |
| | p38 MAPK Signaling | 2.21E+00 | 1.17E-01 | 0 | ATF4,CREB3L1,DDIT3,IL1A,IL33,MKNK2,PLA2G3,PLA2G4A,PLA2G4C,SRF,TAB2,TGFB3,TGFBR2,TP53 |
| | Th1 and Th2 Activation Pathway | 2.18E+00 | 1.05E-01 | NaN | CD86,HLA-DMB,HLA-DQA1,HLA-DQB1,ICOSLG/LOC102723996,IKZF1,IL27RA,IL2RB,IL33,JAG2,NFATC2,NFIL3,PIK3CG,PIK3R3,S1PR1,TGFBR2,TSLP,VAV1 |
| | Aryl Hydrocarbon Receptor Signaling | 2.18E+00 | 1.07E-01 | -0.333 | ALDH1L2,ALDH2,ALDH3A1,ALDH4A1,CCND3,CCNE2,CDKN1A,CDKN2A,CYP1A1,DCT,E2F1,GSTA1,GSTT2/GSTT2B,IL1A,NCOA7,TGFB3,TP53 |
| | Cyclins and Cell Cycle Regulation | 2.18E+00 | 1.29E-01 | -2.111 | CCNB1,CCNB2,CCND3,CCNE2,CDK1,CDKN1A,CDKN2A,CDKN2C,E2F1,TGFB3,TP53 |
| | Role of Hypercytokinemia/hyperchemokinemia in the | 2.14E+00 | 1.28E-01 | 0.905 | AREG,CASP1,CCL2,CXCL10,IFNLR1,IL1A,IL33,ISG20,OAS3,S1PR1,TLR4 |
| | Role of Osteoblasts, Osteoclasts and Chondrocytes in | 2.13E+00 | 9.65E-02 | NaN | ADAMTS4,ALPL,BIRC3,BMP2,BMP4,DKK1,DKK2,FZD10,IGF1,IL1A,IL33,ITGB3,NFATC2,NGFR,PIK3CG,PIK3R3,RUNX2,SFRP4,TAB2,WNT1,WNT2B,WNT7B |
| | Human Embryonic Stem Cell Pluripotency | 2.13E+00 | 9.95E-02 | -1.342 | BMP2,BMP4,DUSP9,FZD10,KLF4,NTRK3,PDGFB,PDGFD,PDGFRA,PDGFRB,PIK3CG,PIK3R3,SPHK1,TBX3,TDGF1,TGFB3,TGFBR2,WNT1,WNT2B,WNT7B |
| | Caveolar-mediated Endocytosis Signaling | 2.11E+00 | 1.33E-01 | NaN | ACTA2,ACTB,ACTC1,ACTG2,CAVIN1,CD55,FLNA,ITGA9,ITGB3,ITGB7 |
| | PDGF Signaling | 2.10E+00 | 1.26E-01 | 0 | CSNK2A3,INPP5B,PDGFB,PDGFD,PDGFRA,PDGFRB,PIK3CG,PIK3R3,PLCG2,SPHK1,SRF |
| | Granulocyte Adhesion and Diapedesis | 2.09E+00 | 1.01E-01 | NaN | ACKR3,CCL2,CLDN19,CLDN5,CXCL10,CXCL12,CXCL14,HRH4,IL1A,IL33,MMP10,MMP15,MMP17,MMP19,MMP24,NGFR,SELP,SELPLG,THY1 |
| | Activin Inhibin Signaling Pathway | 2.07E+00 | 9.68E-02 | -1.964 | CCN2,CDKN1A,CEBPA,CEBPB,COL5A3,FBXO32,FOSB,FOSL2,FSTL3,IL1A,IL33,MAPK15,MAPK4,NGFR,PIK3CG,PIK3R3,PPARG,TDGF1,TGFB3,TLR4,TLR5 |
| | Serine Biosynthesis | 2.06E+00 | 6.67E-01 | NaN | PHGDH,PSAT1 |
| | Transcriptional Regulatory Network in Embryonic Ster | 2.05E+00 | 1.04E-01 | -1.213 | BMP4,FZD10,GATA4,IGF1,IGF2,KLF4,L1CAM,LHX5,PIK3CG,PIK3R3,TBX3,TGFB3,TGFBR2,TP53,WNT1,WNT2B,WNT7B |
| | NAD Signaling Pathway | 2.05E+00 | 1.06E-01 | 0.5 | BST1,CEBPB,ERN1,H1-0,H2BC26,IGF1,NMNAT2,PDGFB,PIK3CG,PIK3R3,PPARGC1A,SLC7A5,SREBF1,TP53,TRPM2,VEGFA |
| | Pancreatic Adenocarcinoma Signaling | 2.03E+00 | 1.11E-01 | 1.667 | CCNE2,CDKN1A,CDKN2A,E2F1,HBEGF,HMOX1,PIK3CG,PIK3R3,PLD5,PLD6,TGFB3,TGFBR2,TP53,VEGFA |
| | RHOGDI Signaling | 2.01E+00 | 9.55E-02 | 1.732 | ACTA2,ACTB,ACTC1,ACTG2,ARHGAP6,ARHGAP8/PRR5-ARHGAP8,ARHGAP9,ARHGEF2,ITGA9,ITGB3,ITGB7,MYH10,MYH11,MYH2,MYH7,MYH9,MYL1,MYL6,MYL9,MYO1A,PAK3 |
| | Folate Transformations I | 1.97E+00 | 3.33E-01 | NaN | MTHFD1L,MTHFD2,SHMT2 |
| | Multiple Sclerosis Signaling Pathway | 1.96E+00 | 9.46E-02 | -0.655 | CAPN6,CASP1,CNTF,GAS5,HLA-DMB,HLA-DQA1,HLA-DQB1,IL17B,IL1A,IL33,MASP1,NFATC2,P2RX7,PLP1,PPARG,SLC8A2,SLC8A3,TGFB3,TLR4,TLR5,TNFSF9 |
| | PAK Signaling | 1.92E+00 | 1.11E-01 | -2.53 | ITGA9,ITGB3,ITGB7,MYL1,MYL6,MYL9,PAK3,PDGFB,PDGFD,PDGFRA,PDGFRB,PIK3CG,PIK3R3 |
| | Oxytocin Signaling Pathway | 1.90E+00 | 8.87E-02 | -1.633 | ATF4,CACNG4,CD36,CREB3L1,CTH,KCNA5,KCNT2,MAPK15,MAPK4,MYH10,MYH11,MYH2,MYH7,MYH9,MYL1,MYL6,MYL9,MYO1A,PIK3CG,PIK3R3,PLA2G3,PLA2G4A,PLA2G4C,PPARG,PPP1R14A |
| | Airway Pathology in Chronic Obstructive Pulmonary D | 1.89E+00 | 1.10E-01 | NaN | CCL2,CNTF,FGF13,FGF18,FGF7,IL17B,IL1A,IL33,LCN9,PTGDS,TGFB3,TNFSF9,TSLP |
| | Phospholipases | 1.89E+00 | 1.30E-01 | 0.707 | HMOX1,PLA2G3,PLA2G4A,PLA2G4C,PLCE1,PLCG2,PLD5,PLD6,PNPLA3 |
| | Cardiomyocyte Differentiation via BMP Receptors | 1.84E+00 | 1.85E-01 | NaN | BMP2,BMP4,GATA4,MYH7,NKX2-5 |
| | Tryptophan Degradation X (Mammalian, via Tryptami | 1.84E+00 | 1.85E-01 | 0.447 | ALDH2,ALDH3A1,ALDH4A1,MAOB,SMOX |
| | Apelin Liver Signaling Pathway | 1.84E+00 | 1.85E-01 | -2.236 | APLNR,COL5A3,EDN1,IRS1,PDGFRB |
| | Atherosclerosis Signaling | 1.84E+00 | 1.05E-01 | NaN | CCL2,CD36,COL5A3,CXCL12,IL1A,IL33,PDGFB,PDGFD,PLA2G3,PLA2G4A,PLA2G4C,PNPLA3,SELP,SELPLG |
| | ERK/MAPK Signaling | 1.82E+00 | 9.30E-02 | 0.258 | ATF4,CREB3L1,DUSP4,DUSP9,EIF4EBP1,ITGA9,ITGB3,ITGB7,MKNK2,PAK3,PIK3CG,PIK3R3,PLA2G3,PLA2G4A,PLA2G4C,PLCG2,PPARG,PPP1R14A,RAPGEF4,SRF |
| | Macrophage Classical Activation Signaling Pathway | 1.79E+00 | 9.52E-02 | -0.471 | CD86,CNTF,CXCL10,GBP2,HLA-DMB,HLA-DQA1,HLA-DQB1,IL17B,IL1A,IL33,mir-130,mir-27,MT-CO1,NCF2,PPARG,TGFB3,TLR4,TNFSF9 |
| | Heme Degradation | 1.78E+00 | 5.00E-01 | NaN | BLVRB,HMOX1 |
| | Melatonin Degradation II | 1.78E+00 | 5.00E-01 | NaN | MAOB,SMOX |
| | Arginine Degradation I (Arginase Pathway) | 1.78E+00 | 5.00E-01 | NaN | ALDH4A1,ARG2 |
| | Neurovascular Coupling Signaling Pathway | 1.77E+00 | 9.05E-02 | -0.218 | CACNG4,CTH,ENTPD3,GABRA2,GABRA3,GABRA4,GABRB2,GABRG1,GABRQ,KCNJ3,KCNJ4,KCNJ6,KCNMA1,KCNMB1,PLA2G3,PLA2G4A,PLA2G4C,PNPLA3,PTGER4,SLC16A1,SLCO2A1 |
| | Role of PKR in Interferon Induction and Antiviral Resp | 1.76E+00 | 1.03E-01 | -0.302 | ATF3,CASP1,CASP5,COLEC12,E2F1,HSPA2,HSPA9,PDGFB,PDGFD,PDGFRB,SCARA3,TAB2,TLR4,TP53 |
| | HOTAIR Regulatory Pathway | 1.76E+00 | 9.82E-02 | -1.5 | CDKN1A,FOXM1,mir-130,MMP10,MMP15,MMP17,MMP19,MMP24,PIK3CG,PIK3R3,SRF,TLR4,TWIST1,WNT1,WNT2B,WNT7B |
| | Breast Cancer Regulation by Stathmin1 | 1.74E+00 | 7.58E-02 | -0.457 | ACKR3,ADGRA2,ADGRL3,ADRA1A,ADRB2,APLNR,ARHGEF2,ATF4,CCND3,CDK1,CMKLR2,CNR1,CREB3L1,E2F1,F2RL2,FOXM1,FZD10,GALR1,GPR137C,GPR157,GPR160,GPR17,GPR183,GPR37L1,GPRC5A,GRM2,HRH4,HTR7,IGF1,MTNR1B,NPY1R,P2RY12,PDGFB,PIK3CG,PIK3R3,PPP1R14A,PTGER4,S1PR1,S1PR3,TACR1,TP53,TUBB3,TUBB4B,TUBB6,VEGFA |
| | VEGF Signaling | 1.70E+00 | 1.11E-01 | -0.333 | ACTA2,ACTB,ACTC1,ACTG2,EIF1,EIF2S2,PIK3CG,PIK3R3,PLCG2,SH2D2A,VEGFA |
| | Endothelin-1 Signaling | 1.68E+00 | 9.28E-02 | 0.728 | CASP1,CASP10,CASP4,CASP5,EDN1,HMOX1,MAPK15,MAPK4,PIK3CG,PIK3R3,PLA2G3,PLA2G4A,PLA2G4C,PLCE1,PLCG2,PLD5,PLD6,PNPLA3 |
| | Inflammasome pathway | 1.67E+00 | 2.00E-01 | 1 | CASP1,CASP5,P2RX7,TLR4 |
| | IL-10 Signaling | 1.65E+00 | 9.74E-02 | 1.291 | ARG2,ATF4,BLVRB,BMP2,CD86,CDKN1A,CREB3L1,DDIT4,HLA-DMB,HLA-DQA1,HLA-DQB1,HMOX1,IL1A,IL33,TLR4 |
| | Role of p14/p19ARF in Tumor Suppression | 1.65E+00 | 1.67E-01 | 1.342 | CDKN2A,E2F1,PIK3CG,PIK3R3,TP53 |
| | Mechanisms of Viral Exit from Host Cells | 1.63E+00 | 1.46E-01 | NaN | ACTA2,ACTB,ACTC1,ACTG2,CHMP4C,LMNB1 |
| | Ribonucleotide Reductase Signaling Pathway | 1.60E+00 | 9.41E-02 | -1 | ACTB,ATF4,CDK1,CDKN1A,CDKN2A,CREB3L1,E2F1,EIF4EBP1,FOXM1,PIK3CG,PIK3R3,RRM2,THBS1,TP53,VASH2,VEGFA |
| | Endoplasmic Reticulum Stress Pathway | 1.59E+00 | 1.90E-01 | 2 | ATF4,DDIT3,ERN1,XBP1 |
| | Tetrahydrofolate Salvage from 5,10-methenyltetrahy | 1.57E+00 | 4.00E-01 | NaN | MTHFD1L,MTHFD2 |
| | Folate Polyglutamylation | 1.57E+00 | 4.00E-01 | NaN | MTHFD1L,SHMT2 |
| | PI3K/AKT Signaling | 1.57E+00 | 9.00E-02 | 0.447 | CDKN1A,EIF4EBP1,GAB2,GDF15,GHR,IFNLR1,IL17RE,IL22RA1,IL27RA,IL2RB,IL31RA,INPP5B,ITGA9,ITGB3,ITGB7,PIK3CG,PIK3R3,TP53 |
| | Virus Entry via Endocytic Pathways | 1.54E+00 | 1.02E-01 | NaN | ACTA2,ACTB,ACTC1,ACTG2,CD55,FLNA,FOLR1,ITGB3,ITGB7,PIK3CG,PIK3R3,PLCG2 |
| | Autophagy | 1.53E+00 | 8.80E-02 | 0.943 | ATF4,CREB3L1,DDIT3,E2F1,ERN1,IGF1,IRS1,NGFR,PDGFB,PIK3CG,PIK3R3,SLC1A5,SLC3A2,SLC7A5,SQSTM1,TLR4,TP53,VEGFA,WIPI1 |
| | Cholesterol Biosynthesis I | 1.51E+00 | 2.31E-01 | NaN | DHCR24,DHCR7,MSMO1 |
| | γ-glutamyl Cycle | 1.51E+00 | 2.31E-01 | NaN | CHAC1,GGT5,OPLAH |
| | Cholesterol Biosynthesis II (via 24,25-dihydrolanoster | 1.51E+00 | 2.31E-01 | NaN | DHCR24,DHCR7,MSMO1 |
| | Cholesterol Biosynthesis III (via Desmosterol) | 1.51E+00 | 2.31E-01 | NaN | DHCR24,DHCR7,MSMO1 |
| | Remodeling of Epithelial Adherens Junctions | 1.49E+00 | 1.18E-01 | NaN | ACTA2,ACTB,ACTC1,ACTG2,TUBB3,TUBB4B,TUBB6,ZYX |
| | Cell Cycle: G1/S Checkpoint Regulation | 1.49E+00 | 1.18E-01 | 1.89 | CCND3,CCNE2,CDKN1A,CDKN2A,CDKN2C,E2F1,TGFB3,TP53 |
| | Role Of Osteoclasts In Rheumatoid Arthritis Signaling | 1.46E+00 | 8.09E-02 | -2.041 | ADAM19,ATF4,BIRC3,COL12A1,COL21A1,COL26A1,COL5A1,COL5A3,COL6A6,COL9A1,CREB3L1,ITGB3,MMP10,MMP15,MMP17,MMP19,MMP24,NGFR,PIK3CG,PIK3R3,PLCG2,SFRP4,SRF,TAB2,TLR4 |
| | Apelin Cardiac Fibroblast Signaling Pathway | 1.46E+00 | 1.74E-01 | 0 | APLNR,CCN2,SPHK1,TGFB3 |
| | cAMP-mediated signaling | 1.44E+00 | 8.47E-02 | -1.414 | ADRB2,AKAP5,APLNR,ATF4,CNGA3,CNR1,CREB3L1,DUSP4,DUSP9,GPR17,GRM2,HTR7,NPY1R,P2RY12,PLD6,PTGER4,RAPGEF4,RGS4,S1PR1,S1PR3 |
| | SNARE Signaling Pathway | 1.44E+00 | 9.56E-02 | -2.496 | MYH10,MYH11,MYH2,MYH7,MYH9,MYL1,MYL6,MYL9,MYO1A,STXBP2,STXBP6,SYT12,SYT6 |
| | Inhibition of Angiogenesis by TSP1 | 1.44E+00 | 1.47E-01 | -0.447 | CD36,TGFBR2,THBS1,TP53,VEGFA |
| | G-Protein Coupled Receptor Signaling | 1.43E+00 | 7.11E-02 | -1.857 | ACKR3,ADGRA2,ADGRL3,ADRA1A,ADRB2,APLNR,ATF4,CMKLR2,CNGA3,CNR1,CREB3L1,DUSP4,DUSP9,F2RL2,FZD10,GALR1,GPR137C,GPR157,GPR160,GPR17,GPR183,GPR37L1,GPRC5A,GRM2,HRH4,HTR7,MAP3K15,MEGF6,MTNR1B,MYL1,MYL6,MYL9,MYLK3,NFATC2,NPY1R,P2RY12,PAK3,PIK3CG,PIK3R3,PLCE1,PLCG2,PLD6,PTGER4,RAPGEF4,RGS16,RGS4,S1PR1,S1PR3,SRF,TACR1 |
| | CREB Signaling in Neurons | 1.43E+00 | 7.25E-02 | -1.372 | ACKR3,ADGRA2,ADGRL3,ADRA1A,ADRB2,APLNR,ATF4,CACNG4,CMKLR2,CNR1,CREB3L1,F2RL2,FZD10,GALR1,GHR,GPR137C,GPR157,GPR160,GPR17,GPR183,GPR37L1,GPRC5A,GRM2,HRH4,HTR7,IGF1,MTNR1B,NGFR,NPY1R,NTRK3,P2RY12,PDGFB,PDGFRA,PDGFRB,PIK3CG,PIK3R3,PLCE1,PLCG2,PTGER4,S1PR1,S1PR3,TACR1,TGFBR2,VEGFA |
| | Clathrin-mediated Endocytosis Signaling | 1.42E+00 | 8.65E-02 | NaN | ACTA2,ACTB,ACTC1,ACTG2,CSNK2A3,FGF13,FGF18,FGF7,IGF1,ITGB3,ITGB7,LDLR,PDGFB,PDGFD,PIK3CG,PIK3R3,SH3KBP1,VEGFA |
| | PTEN Signaling | 1.42E+00 | 9.27E-02 | 2.333 | CDKN1A,CSNK2A3,GHR,INPP5B,ITGA9,ITGB3,ITGB7,NGFR,NTRK3,PDGFRA,PDGFRB,PIK3CG,PIK3R3,TGFBR2 |
| | Arginine Degradation VI (Arginase 2 Pathway) | 1.41E+00 | 3.33E-01 | NaN | ARG2,PYCR1 |
| | Selenocysteine Biosynthesis II (Archaea and Eukaryot | 1.41E+00 | 3.33E-01 | NaN | SARS1,SEPHS2 |
| | Kinetochore Metaphase Signaling Pathway | 1.38E+00 | 9.91E-02 | -2.121 | BUB1B,CCNB1,CDK1,CENPA,CENPK,CENPO,KNL1,MXD3,PPP1R14A,REC8,SPC24 |
| | LPS/IL-1 Mediated Inhibition of RXR Function | 1.38E+00 | 8.24E-02 | 0 | ALDH1L2,ALDH2,ALDH3A1,ALDH4A1,CPT1A,FMO2,GSTA1,GSTT2/GSTT2B,HMGCS1,HS3ST5,IL1A,IL33,MAOB,NGFR,PPARGC1A,SLCO1A2,SMOX,SREBF1,SULT1A1,SULT1C2,TLR4 |
| | Tryptophan Degradation III (Eukaryotic) | 1.34E+00 | 1.60E-01 | NaN | ACAT2,HACD1,KMO,TBXAS1 |
| | Choline Biosynthesis III | 1.34E+00 | 2.00E-01 | NaN | HMOX1,PLD5,PLD6 |
| | Germ Cell-Sertoli Cell Junction Signaling | 1.32E+00 | 8.82E-02 | NaN | A2M,ACTA2,ACTB,ACTC1,ACTG2,MAP3K15,PAK3,PIK3CG,PIK3R3,TGFB3,TGFBR2,TUBB3,TUBB4B,TUBB6,ZYX |

| | | | | |
|---|---|---|---|---|
| Necroptosis Signaling Pathway | 1.32E+00 | 8.97E-02 | 1.069 | BIRC3,CAPN6,CASP1,CASP10,NGFR,PLA2G3,PLA2G4A,PLA2G4C,TAB2,TIMM44,TLR4,TNFRSF10A,TNFRSF10B,TP53 |
| Noradrenaline and Adrenaline Degradation | 1.30E+00 | 1.35E-01 | 0.447 | ALDH2,ALDH3A1,ALDH4A1,MAOB,SMOX |
| Regulation of Actin-based Motility by Rho | 1.29E+00 | 9.57E-02 | -2.828 | ACTA2,ACTB,ACTC1,ACTG2,ITGA9,ITGB3,ITGB7,MYL1,MYL6,MYL9,PAK3 |
| Estrogen-mediated S-phase Entry | 1.29E+00 | 1.54E-01 | -2 | CCNE2,CDK1,CDKN1A,E2F1 |
| Role of JAK2 in Hormone-like Cytokine Signaling | 1.28E+00 | 1.13E-01 | -1.89 | BIRC3,CCND3,GHR,IGF1,IRS1,POMC,VEGFA |
| CDX Gastrointestinal Cancer Signaling Pathway | 1.27E+00 | 8.42E-02 | 0.728 | BLM,BMP2,BMP4,CNTF,FZD10,HBEGF,IL17B,IL1A,IL33,NCF2,PIK3CG,PIK3R3,TGFB3,TNFSF9,WNT1,WNT2B,WNT7B |
| Ephrin Receptor Signaling | 1.27E+00 | 8.42E-02 | 0 | ATF4,CREB3L1,CXCL12,EFNA5,EPHA1,EPHA4,EPHA6,EPHA8,EPHB6,ITGA9,ITGB3,ITGB7,PAK3,PDGFB,PDGFD,PIK3CG,VEGFA |
| Melanoma Signaling | 1.26E+00 | 1.20E-01 | -1.342 | CDKN1A,CDKN2A,E2F1,PIK3CG,PIK3R3,TP53 |
| MYC Mediated Apoptosis Signaling | 1.26E+00 | 1.20E-01 | 0.816 | ADRB2,BBC3,CDKN2A,NGFR,PMAIP1,TP53 |
| Thio-molybdenum Cofactor Biosynthesis | 1.26E+00 | 1.00E+00 | NaN | MOCOS |
| L-cysteine Degradation II | 1.26E+00 | 1.00E+00 | NaN | CTH |
| Asparagine Biosynthesis I | 1.26E+00 | 1.00E+00 | NaN | ASNS |
| Immunogenic Cell Death Signaling Pathway | 1.23E+00 | 1.00E-01 | 0.333 | ATF4,CASP1,CXCL10,DDIT3,HSPA2,HSPA9,NGFR,P2RX7,TLR4 |
| IGF-1 Signaling | 1.20E+00 | 9.52E-02 | -0.816 | CCN1,CCN2,CSNK2A3,GRB10,IGF1,IGFBP3,IRS1,PIK3CG,PIK3R3,SRF |
| NRF2-mediated Oxidative Stress Response | 1.20E+00 | 8.02E-02 | 2.828 | ACTA2,ACTB,ACTC1,ACTG2,ATF4,CYP1A1,DNAJB13,ENC1,FTH1,FTL,GSTA1,GSTT2/GSTT2B,HERPUD1,HMOX1,MAFG,PIK3CG,PIK3R3,SQSTM1,TXNRD1 |
| Histidine Degradation III | 1.17E+00 | 2.50E-01 | NaN | MTHFD1L,MTHFD2 |
| Paxillin Signaling | 1.16E+00 | 9.35E-02 | -1.89 | ACTA2,ACTB,ACTC1,ACTG2,ITGA9,ITGB3,ITGB7,PAK3,PIK3CG,PIK3R3 |
| Semaphorin Neuronal Repulsive Signaling Pathway | 1.16E+00 | 8.67E-02 | -2.309 | ACAN,ITGA9,ITGB3,ITGB7,MYL1,MYL6,MYL9,PAK3,PIK3CG,PIK3R3,PLCG2,SEMA3F,TP53 |
| Regulation of the Epithelial-Mesenchymal Transition I | 1.15E+00 | 8.21E-02 | NaN | EGR1,FGF13,FGF18,FGF7,FZD10,JAG2,PDGFD,PDGFRB,PIK3CG,PIK3R3,TGFB3,TGFBR2,TWIST1,WNT1,WNT2B,WNT7B |
| Th1 Pathway | 1.14E+00 | 9.02E-02 | 0.905 | CD86,HLA-DMB,HLA-DQA1,HLA-DQB1,ICOSLG/LOC102723996,IL27RA,NFATC2,NFIL3,PIK3CG,PIK3R3,VAV1 |
| Glutaryl-CoA Degradation | 1.14E+00 | 1.67E-01 | NaN | ACAT2,HACD1,TBXAS1 |
| DHCR24 Signaling Pathway | 1.13E+00 | 8.76E-02 | -1.155 | DHCR24,DHCR7,HMGCR,HMOX1,IGF1,PDGFB,PDGFD,PIK3CG,PIK3R3,POMC,SREBF1,TP53 |
| IL-15 Production | 1.12E+00 | 8.94E-02 | -0.905 | DDR2,EPHA1,EPHA4,EPHA6,EPHA8,KIT,LYN,NTRK3,PDGFRA,PDGFRB,TEK |
| Integrin Signaling | 1.12E+00 | 8.02E-02 | -1.807 | ACTA2,ACTB,ACTC1,ACTG2,CAPN6,ITGA9,ITGB3,ITGB7,MYL9,MYLK3,PAK3,PDGFB,PIK3CG,PIK3R3,PLCG2,TSPAN2,ZYX |
| Death Receptor Signaling | 1.09E+00 | 9.38E-02 | -0.333 | ACTA2,ACTB,ACTC1,ACTG2,BIRC3,CASP10,TNFRSF10A,TNFRSF10B,TNFRSF25 |
| Salvage Pathways of Pyrimidine Deoxyribonucleotides | 1.08E+00 | 2.22E-01 | NaN | APOBEC3B,UPP1 |
| IL-33 Signaling Pathway | 1.06E+00 | 8.06E-02 | -0.535 | AREG,ATF4,CASP1,CASP10,CASP4,CASP5,CCL2,CREB3L1,H2BC26,IL1A,IL33,KIT,PIK3CG,PIK3R3,PLCG2 |
| Role of Pattern Recognition Receptors in Recognition | 1.06E+00 | 8.33E-02 | 0.816 | CASP1,CNTF,IL17B,IL1A,IL33,OAS3,PIK3CG,PIK3R3,PLCG2,TGFB3,TLR4,TLR5,TNFSF9 |
| eNOS Signaling | 1.06E+00 | 8.33E-02 | 0.707 | AQP1,AQP3,AQP5,CHRNA3,CHRNA9,CNGA3,HSPA2,HSPA9,PIK3CG,PIK3R3,PLCG2,SLC7A1,VEGFA |
| Eicosanoid Signaling | 1.05E+00 | 1.00E-01 | 0 | PLA2G3,PLA2G4A,PLA2G4C,PNPLA3,PTGDS,PTGER4,TBXAS1 |
| TR/RXR Activation | 1.05E+00 | 9.52E-02 | NaN | LDLR,NRGN,PIK3CG,PIK3R3,PPARGC1A,SREBF1,SYT12,TBL1Y |
| Hereditary Breast Cancer Signaling | 1.04E+00 | 8.45E-02 | NaN | ACTB,BARD1,BLM,CCNB1,CCND3,CDK1,CDKN1A,E2F1,GADD45B,PIK3CG,PIK3R3,TP53 |
| IL-17 Signaling | 1.04E+00 | 8.02E-02 | 0.258 | CCL2,CEBPB,CNTF,IL17B,IL17RE,IL1A,IL33,mir-23,PIK3CG,PIK3R3,RGS16,TAB2,TGFB3,TNFSF9,VEGFA |
| MIF Regulation of Innate Immunity | 1.04E+00 | 1.14E-01 | 0.447 | PLA2G3,PLA2G4A,PLA2G4C,TLR4,TP53 |
| Androgen Biosynthesis | 1.03E+00 | 1.50E-01 | NaN | GSTA1,HSD17B14,HSD3B7 |
| Antioxidant Action of Vitamin C | 1.02E+00 | 8.77E-02 | -0.707 | HMOX1,PLA2G3,PLA2G4A,PLA2G4C,PLCE1,PLCG2,PLD5,PLD6,PNPLA3,TXNRD1 |
| NOD1/2 Signaling Pathway | 1.01E+00 | 7.94E-02 | -0.775 | BIRC3,CASP1,CCL2,CNTF,ERN1,HSPA2,HSPA9,IL17B,IL1A,IL33,TAB2,TGFB3,TLR4,TLR5,TNFSF9 |
| ATM Signaling | 1.01E+00 | 9.00E-02 | 1.633 | ATF4,BLM,CCNB1,CCNB2,CDK1,CDKN1A,CREB3L1,GADD45B,TP53 |
| Basal Cell Carcinoma Signaling | 1.00E+00 | 9.72E-02 | -1 | BMP2,BMP4,FZD10,TP53,WNT1,WNT2B,WNT7B |
| Serotonin Degradation | 1.00E+00 | 9.72E-02 | 1.134 | ALDH2,ALDH3A1,ALDH4A1,MAOB,SMOX,SULT1A1,SULT1C2 |
| Synaptogenesis Signaling Pathway | 9.98E-01 | 7.30E-02 | -1.279 | ATF4,CHN1,CREB3L1,EFNA5,EIF4EBP1,EIF4EBP3,EPHA1,EPHA4,EPHA6,EPHA8,EPHB6,GRM2,LYN,NLGN4X,NRXN1,PIK3CG,PIK3R3,PLCG2,STXBP2,STXBP6,SYT12,SYT6,THBS1 |
| Embryonic Stem Cell Differentiation into Cardiac Linea | 9.95E-01 | 2.00E-01 | NaN | GATA4,NKX2-5 |
| Prostanoid Biosynthesis | 9.95E-01 | 2.00E-01 | NaN | PTGDS,TBXAS1 |
| Sertoli Cell-Sertoli Cell Junction Signaling | 9.89E-01 | 7.77E-02 | NaN | A2M,ACTA2,ACTB,ACTC1,ACTG2,CLDN19,CLDN5,ITGA9,ITGB3,ITGB7,MAP3K15,TGFB3,TJP2,TUBB3,TUBB4B,TUBB6 |
| Xenobiotic Metabolism AHR Signaling Pathway | 9.81E-01 | 9.20E-02 | 0.707 | ALDH1L2,ALDH2,ALDH3A1,ALDH4A1,CYP1A1,GSTA1,GSTT2/GSTT2B,IL1A |
| IL-13 Signaling Pathway | 9.80E-01 | 8.62E-02 | -1.265 | CD36,FOSL2,HBEGF,IL33,LYN,PIK3CG,PIK3R3,TGFB3,TP53,TSLP |
| Airway Inflammation in Asthma | 9.80E-01 | 1.21E-01 | NaN | CCL2,IL33,TGFB3,TSLP |
| Histamine Degradation | 9.78E-01 | 1.43E-01 | NaN | ALDH2,ALDH3A1,ALDH4A1 |
| Fatty Acid α-oxidation | 9.78E-01 | 1.43E-01 | NaN | ALDH2,ALDH3A1,ALDH4A1 |
| Role of Tissue Factor in Cancer | 9.75E-01 | 7.73E-02 | -2 | CCN1,CCN2,EGR1,FGB,HBEGF,ITGB3,LYN,MAPK15,MAPK4,NGFR,PIK3CG,PIK3R3,TGFB3,TGFBR2,TP53,VEGFA |
| Apelin Pancreas Signaling Pathway | 9.74E-01 | 1.09E-01 | -1.342 | APLNR,DDIT3,ERN1,PIK3CG,PIK3R3 |
| 4-hydroxyproline Degradation I | 9.72E-01 | 5.00E-01 | NaN | ALDH4A1 |
| Glycine Biosynthesis I | 9.72E-01 | 5.00E-01 | NaN | SHMT2 |
| Regulation Of The Epithelial Mesenchymal Transition | 9.69E-01 | 7.81E-02 | -1.732 | EGR1,FGF13,FGF18,FGF7,NGFR,PDGFB,PDGFD,PDGFRA,PDGFRB,PIK3CG,PIK3R3,TGFB3,TGFBR2,TNFSF9,TWIST1 |
| Neutrophil Extracellular Trap Signaling Pathway | 9.60E-01 | 7.02E-02 | 2.646 | ARG2,CASP1,CASP10,CASP4,CASP5,COL12A1,COL21A1,COL26A1,COL5A1,COL5A3,COL6A6,COL9A1,LYN,NCF2,NDUFA4L2,PIK3CG,PIK3R3,PLA2G3,PLA2G4A,PLA2G4C,PLCE1,PLCG2,PNPLA3,SELP,SELPLG,TIMM44,TLR4,VSTM1 |
| Role of IL-17F in Allergic Inflammatory Airway Disease | 9.44E-01 | 1.06E-01 | NaN | ATF4,CCL2,CREB3L1,CXCL10,IGF1 |
| Molecular Mechanisms of Cancer | 9.40E-01 | 6.89E-02 | NaN | ARHGEF2,BBC3,BIRC3,BMP2,BMP4,CASP10,CCND3,CCNE2,CDK1,CDKN1A,CDKN2A,CDKN2C,E2F1,FZD10,GAB2,IRS1,ITGA9,ITGB3,ITGB7,PAK3,PIK3CG,PIK3R3,PMAIP1,RASGRF2,TAB2,TGFB3,TGFBR2,TP53,WNT1,WNT2B,WNT7B |
| Glucocorticoid Receptor Signaling | 9.38E-01 | 6.70E-02 | NaN | A2M,ACTB,ADRB2,ARG2,CCL2,CDKN1A,CEBPA,GHR,HLA-DMB,HLA-DQA1,HLA-DQB1,HSPA2,HSPA9,IFNLR1,IGF1,IL17RE,IL1A,IL22RA1,IL27RA,IL2RB,IL31RA,KRT77,KRT80,NDUFA4L2,NFATC2,PCK2,PDK4,PIK3CG,PIK3R3,PLA2G3,PLA2G4A,PLA2G4C,POMC,PPARG,SGK1,TAF4B,TGFB3,TGFBR2,TSC22D3 |
| MSP-RON Signaling In Macrophages Pathway | 9.26E-01 | 8.40E-02 | -1 | ATF4,CREB3L1,GAB2,HLA-DMB,HLA-DQA1,HLA-DQB1,KLK5,PIK3CG,PIK3R3,TLR4 |
| Ketogenesis | 9.24E-01 | 1.82E-01 | NaN | ACAT2,HMGCS1 |
| Glycine Betaine Degradation | 9.24E-01 | 1.82E-01 | NaN | DMGDH,SHMT2 |
| EIF2 Signaling | 9.18E-01 | 7.49E-02 | -1.265 | ACTA2,ACTB,ACTC1,ACTG2,ATF3,ATF4,ATF5,DDIT3,EIF1,EIF2S2,PIK3CG,PIK3R3,PPP1R15A,SREBF1,TRIB3,VEGFA,WARS1 |
| Regulation of Cellular Mechanics by Calpain Protease | 9.18E-01 | 8.89E-02 | NaN | CAPN6,CCND3,CCNE2,CDK1,CNGA3,ITGA9,ITGB3,ITGB7 |
| PFKFB4 Signaling Pathway | 9.15E-01 | 1.04E-01 | 0 | ATF4,CREB3L1,HK2,TGFB3,TP53 |
| GDNF Family Ligand-Receptor Interactions | 9.10E-01 | 9.21E-02 | 0 | DOK5,DOK7,IRS1,PDLIM7,PIK3CG,PIK3R3,PLCG2 |
| Macropinocytosis Signaling | 9.10E-01 | 9.21E-02 | NaN | ITGB3,ITGB7,PDGFB,PDGFD,PIK3CG,PIK3R3,PLCG2 |
| Crosstalk between Dendritic Cells and Natural Killer Ce | 8.98E-01 | 8.79E-02 | NaN | ACTA2,ACTB,ACTC1,ACTG2,CD86,FSCN2,IL2RB,TLR4 |
| Apelin Adipocyte Signaling Pathway | 8.98E-01 | 8.79E-02 | -1.414 | APLNR,GPX3,GSTA1,GSTT2/GSTT2B,MAPK15,MAPK4,NCF2,PPARGC1A |
| TREM1 Signaling | 8.88E-01 | 9.09E-02 | 0.378 | CASP1,CASP5,CCL2,CD86,PLCG2,TLR4,TLR5 |
| Chronic Myeloid Leukemia Signaling | 8.80E-01 | 7.19E-02 | -2.236 | CCND3,CDKN1A,CDKN2A,CEBPA,E2F1,FOXM1,FZD10,GAB2,KLF4,NFATC2,PIK3CG,PIK3R3,PLCE1,PLCG2,TGFB3,TGFBR2,TP53,WNT1,WNT2B,WNT7B |
| MIF-mediated Glucocorticoid Regulation | 8.76E-01 | 1.11E-01 | 1 | PLA2G3,PLA2G4A,PLA2G4C,TLR4 |
| PPAR Signaling | 8.75E-01 | 8.41E-02 | 2.333 | IL1A,IL33,NGFR,PDGFB,PDGFD,PDGFRA,PDGFRB,PPARG,PPARGC1A |
| PD-1, PD-L1 cancer immunotherapy pathway | 8.75E-01 | 8.41E-02 | -1.89 | CSNK2A3,HLA-DMB,HLA-DQA1,HLA-DQB1,IL2RB,NGFR,PIK3CG,PIK3R3,TGFB3 |
| VDR/RXR Activation | 8.67E-01 | 8.97E-02 | 0 | CDKN1A,CEBPA,CEBPB,CXCL10,IGFBP3,KLF4,RUNX2 |
| Neurotrophin/TRK Signaling | 8.67E-01 | 8.97E-02 | 0 | ATF4,CREB3L1,NGFR,NTRK3,PIK3CG,PIK3R3,SORCS1 |
| FAT10 Cancer Signaling Pathway | 8.60E-01 | 1.00E-01 | -1.342 | ACKR3,NGFR,TGFB3,TGFBR2,TP53 |
| GDP-glucose Biosynthesis | 8.60E-01 | 1.67E-01 | NaN | HK2,HKDC1 |
| Mineralocorticoid Biosynthesis | 8.60E-01 | 1.67E-01 | NaN | GSTA1,HSD3B7 |
| LXR/RXR Activation | 8.58E-01 | 8.13E-02 | 0 | ARG2,CCL2,CD36,HMGCR,IL1A,IL33,LDLR,NGFR,SREBF1,TLR4 |
| Protein Kinase A Signaling | 8.52E-01 | 6.81E-02 | 0 | AKAP5,ATF4,CNGA3,CREB3L1,DUSP4,DUSP5,DUSP9,FLNA,H1-0,MYH10,MYH2,MYL1,MYL6,MYL9,MYLK3,NFATC2,NGFR,PHKG1,PLCE1,PLCG2,PLD6,PLN,PPP1R14A,PTPRN,TGFB3,TGFBR2,TNNI1,TNNI3 |
| RAR Activation | 8.47E-01 | 6.78E-02 | 0.928 | ACTB,ATF4,CDKN1A,CNGA3,CNTF,CREB3L1,CSNK2A3,CYP26A1,DHRS3,DKK1,DKK2,GATA4,IGFBP3,IL17B,IL1A,IL33,KIT,LAMB1,PIK3CG,PIK3R3,PLD6,PPARG,PPARGC1A,PRDM16,RPE65,TGFB3,TGFBR2,TNFSF9,VEGFA |
| Role of MAPK Signaling in Inhibiting the Pathogenesis | 8.47E-01 | 8.86E-02 | -0.378 | CCL2,CXCL10,PLA2G3,PLA2G4A,PLA2G4C,PNPLA3,TLR4 |
| Role of NANOG in Mammalian Embryonic Stem Cell P | 8.42E-01 | 8.06E-02 | NaN | BMP2,BMP4,FZD10,GATA4,PIK3CG,PIK3R3,TP53,WNT1,WNT2B,WNT7B |
| Role of BRCA1 in DNA Damage Response | 8.26E-01 | 8.75E-02 | 0.447 | ACTB,BARD1,BLM,BRIP1,CDKN1A,E2F1,TP53 |
| Gustation Pathway | 8.26E-01 | 7.39E-02 | -1.291 | CACNG4,CD36,GABRA2,GABRA3,GABRA4,GABRB2,GABRG1,GABRQ,P2RX3,P2RX7,P2RY12,SCN1B,SCN4A,SCN7A,SLC5A1 |
| WNT/Ca+ pathway | 8.15E-01 | 9.09E-02 | -0.447 | ATF4,CREB3L1,FZD10,NFATC2,PLCE1,PLCG2 |
| Ethanol Degradation IV | 8.09E-01 | 1.20E-01 | NaN | ALDH2,ALDH3A1,ALDH4A1 |
| Proline Degradation | 8.08E-01 | 3.33E-01 | NaN | ALDH4A1 |
| L-cysteine Degradation III | 8.08E-01 | 3.33E-01 | NaN | GOT1 |
| Biotin-carboxyl Carrier Protein Assembly | 8.08E-01 | 3.33E-01 | NaN | HLCS |
| Glutamate Degradation II | 8.08E-01 | 3.33E-01 | NaN | GOT1 |
| Aspartate Biosynthesis | 8.08E-01 | 3.33E-01 | NaN | GOT1 |
| CLEAR Signaling Pathway | 8.07E-01 | 7.02E-02 | 2.982 | ATF4,ATP6V0D2,CREB3L1,DDIT4,GHR,IGF1,MAPK15,MAPK4,NGFR,NTRK3,PDGFB,PDGFRA,PDGFRB,PPARGC1A,SESN2,TGFBR2,TLR4,TLR5,TP53,VEGFA |
| Glucose and Glucose-1-phosphate Degradation | 8.03E-01 | 1.54E-01 | NaN | HK2,HKDC1 |
| NAD biosynthesis II (from tryptophan) | 8.03E-01 | 1.54E-01 | NaN | KMO,NMNAT2 |
| Glucocorticoid Biosynthesis | 8.03E-01 | 1.54E-01 | NaN | GSTA1,HSD3B7 |
| UDP-N-acetyl-D-galactosamine Biosynthesis II | 8.03E-01 | 1.54E-01 | NaN | HK2,HKDC1 |
| Mitotic Roles of Polo-Like Kinase | 7.93E-01 | 8.96E-02 | -2.236 | CCNB1,CCNB2,CDK1,KIF11,KIF23,PKMYT1 |
| Melanocyte Development and Pigmentation Signaling | 7.70E-01 | 8.16E-02 | 0 | ATF4,CREB3L1,DCT,KIT,PIK3CG,PIK3R3,PLCG2,POMC |
| IL-6 Signaling | 7.66E-01 | 7.75E-02 | -0.333 | A2M,CEBPB,CSNK2A3,IL1A,IL33,NGFR,PIK3CG,PIK3R3,SRF,VEGFA |
| Sperm Motility | 7.62E-01 | 7.00E-02 | 0 | CNGA3,DDR2,EPHA1,EPHA4,EPHA6,EPHA8,KIT,LYN,NTRK3,PDGFRA,PDGFRB,PLA2G3,PLA2G4A,PLA2G4C,PLCE1,PLCG2,PNPLA3,TEK |
| Agrin Interactions at Neuromuscular Junction | 7.52E-01 | 8.70E-02 | -2.236 | ACTA2,ACTB,ACTC1,ACTG2,LAMB1,PAK3 |
| VEGF Family Ligand-Receptor Interactions | 7.50E-01 | 8.33E-02 | 1.134 | PIK3CG,PIK3R3,PLA2G3,PLA2G4A,PLA2G4C,PLCG2,VEGFA |
| AMPK Signaling | 7.49E-01 | 7.02E-02 | -0.302 | ACTB,ADRA1A,ADRB2,AK7,ATF4,CDKN1A,CHRNA3,CHRNA9,CPT1A,CREB3L1,EIF4EBP1,HMGCR,IRS1,PCK2,PIK3CG,PIK3R3,PPARGC1A |
| CDK5 Signaling | 7.43E-01 | 7.83E-02 | 0.333 | CABLES1,CDK5R1,EGR1,FOSB,LAMB1,MAPK15,MAPK4,NGFR,PPP1R14A |
| Xenobiotic Metabolism Signaling | 7.39E-01 | 6.85E-02 | NaN | ALDH1L2,ALDH2,ALDH3A1,ALDH4A1,CYP1A1,FMO2,FTL,GSTA1,GSTT2/GSTT2B,HMOX1,HS3ST5,IL1A,MAOB,MAP3K15,PIK3CG,PIK3R3,PPARGC1A,SMOX,SULT1A1,SULT1C2 |
| Estrogen Receptor Signaling | 7.32E-01 | 6.60E-02 | -1.633 | ARG2,ATF4,CACNG4,CDKN1A,CREB3L1,EIF4EBP1,FBXO32,IGF1,IGF2,MMP10,MMP15,MMP17,MMP19,MMP24,MYL1,MYL6,MYL9,NDUFA4L2,PIK3CG,PIK3R3,PLCE1,PLCG2,PPARGC1A,RUNX2,TBL1Y,TP53,VEGFA |
| Amyotrophic Lateral Sclerosis Signaling | 7.28E-01 | 7.76E-02 | 1.414 | BIRC3,CACNG4,CAPN6,CASP1,IGF1,PIK3CG,PIK3R3,TP53,VEGFA |
| Cardiac Hypertrophy Signaling | 7.22E-01 | 6.90E-02 | -2 | ADRA1A,ADRB2,CACNG4,GATA4,IGF1,IRS1,MAP3K15,MYL1,MYL6,MYL9,NKX2-5,PIK3CG,PIK3R3,PLCE1,PLCG2,SRF,TGFB3,TGFBR2 |
| D-myo-inositol-5-phosphate Metabolism | 7.21E-01 | 7.14E-02 | -0.302 | ALPL,CA3,DUSP4,DUSP5,DUSP9,EYA4,INPP5B,MTMR8,PLCE1,PLCG2,PPFIA2,PPP1R14A,PSPH,PTPRN |
| Cell Cycle Control of Chromosomal Replication | 7.16E-01 | 8.93E-02 | -1.342 | CDC6,CDK1,MCM4,RPA4,TOP2A |
| FGF Signaling | 7.15E-01 | 8.14E-02 | 0.816 | ATF4,CREB3L1,FGF13,FGF18,FGF7,PIK3CG,PIK3R3 |
| Neuregulin Signaling | 7.14E-01 | 7.69E-02 | 0 | AREG,CDK5R1,ERRFI1,HBEGF,ITGA9,ITGB3,ITGB7,PIK3R3,PLCG2 |
| Glutathione Redox Reactions I | 7.06E-01 | 1.07E-01 | NaN | GPX3,GSTA1,GSTT2/GSTT2B |
| CDP-diacylglycerol Biosynthesis I | 7.06E-01 | 1.07E-01 | NaN | CDS1,GPAT3,PNPLA3 |
| Fc Epsilon RI Signaling | 6.99E-01 | 7.63E-02 | 1 | INPP5B,LYN,PIK3CG,PIK3R3,PLA2G3,PLA2G4A,PLA2G4C,PLCG2,VAV1 |
| Gap Junction Signaling | 6.99E-01 | 7.07E-02 | NaN | ACTA2,ACTB,ACTC1,ACTG2,CTH,GJB2,PIK3CG,PIK3R3,PLCE1,PLCG2,TJP2,TUBB3,TUBB4B,TUBB6 |
| Spermine and Spermidine Degradation I | 6.94E-01 | 2.50E-01 | NaN | SMOX |
| Phenylethylamine Degradation I | 6.94E-01 | 2.50E-01 | NaN | ALDH2 |
| Eumelanin Biosynthesis | 6.94E-01 | 2.50E-01 | NaN | DCT |
| Proline Biosynthesis I | 6.94E-01 | 2.50E-01 | NaN | PYCR1 |
| L-cysteine Degradation I | 6.94E-01 | 2.50E-01 | NaN | GOT1 |
| Oxytocin In Brain Signaling Pathway | 6.88E-01 | 7.04E-02 | -1.387 | ATF4,CACNG4,CASP1,CASP5,CREB3L1,DDIT3,KCNJ13,PIK3CG,PIK3R3,PLA2G3,PLA2G4A,PLA2G4C,PNPLA3,TLR4 |
| Cancer Drug Resistance By Drug Efflux | 6.74E-01 | 8.62E-02 | -0.447 | mir-130,mir-27,PIK3CG,PIK3R3,TP53 |
| Superpathway of Inositol Phosphate Compounds | 6.65E-01 | 6.84E-02 | -0.277 | ALPL,CA3,DUSP4,DUSP5,DUSP9,EYA4,INPP5B,MTMR8,PIK3CG,PIK3R3,PLCE1,PLCG2,PPFIA2,PPP1R14A,PSPH,PTPRN |
| Polyamine Regulation in Colon Cancer | 6.54E-01 | 8.47E-02 | -1 | CSNK2A3,PPARG,SLC3A2,SMOX,TP53 |
| Phosphatidylglycerol Biosynthesis II (Non-plastidic) | 6.46E-01 | 1.00E-01 | NaN | CDS1,GPAT3,PNPLA3 |
| GPCR-Mediated Integration of Enteroendocrine Signa | 6.40E-01 | 8.00E-02 | 0 | ADRB2,CALCA,GALR1,PLCE1,PLCG2,SST |
| PCP (Planar Cell Polarity) Pathway | 6.34E-01 | 8.33E-02 | -1.342 | FZD10,RSPO3,WNT1,WNT2B,WNT7B |
| Adipogenesis pathway | 6.32E-01 | 7.19E-02 | 0.632 | BMP2,BMP4,CEBPA,CEBPB,DDIT3,FZD10,PPARG,SREBF1,TP53,XBP1 |
| Colorectal Cancer Metastasis Signaling | 6.30E-01 | 6.64E-02 | -2 | FZD10,MMP10,MMP15,MMP17,MMP19,MMP24,PIK3CG,PIK3R3,PTGER4,TGFB3,TGFBR2,TLR4,TLR5,TP53,VEGFA,WNT1,WNT2B,WNT7B |
| Aldosterone Signaling in Epithelial Cells | 6.27E-01 | 6.98E-02 | -0.447 | CRYAB,DNAJB13,HSPA12A,HSPA2,HSPA9,KCNMA1,KCNMB1,PIK3CG,PIK3R3,PLCE1,PLCG2,SGK1 |
| Hypoxia Signaling in the Cardiovascular System | 6.23E-01 | 7.89E-02 | NaN | ATF4,CREB3L1,EDN1,TP53,UBE2C,VEGFA |
| Isoleucine Degradation I | 6.22E-01 | 1.18E-01 | NaN | ACAT2,BCAT1 |
| Sonic Hedgehog Signaling | 6.19E-01 | 9.68E-02 | NaN | CCNB1,CCNB2,CDK1 |
| 3-phosphoinositide Biosynthesis | 6.16E-01 | 6.80E-02 | -0.302 | ALPL,CA3,DUSP4,DUSP5,DUSP9,EYA4,INPP5B,MTMR8,PIK3CG,PIK3R3,PPFIA2,PPP1R14A,PSPH,PTPRN |
| Proline Biosynthesis II (from Arginine) | 6.09E-01 | 2.00E-01 | NaN | PYCR1 |
| Tetrapyrrole Biosynthesis II | 6.09E-01 | 2.00E-01 | NaN | ALAS2 |
| Trans, trans-farnesyl Diphosphate Biosynthesis | 6.09E-01 | 2.00E-01 | NaN | IDI1 |

| | | | | |
|---|---|---|---|---|
| Lactose Degradation III | 6.09E-01 | 2.00E-01 | NaN | GLB1L3 |
| dTMP De Novo Biosynthesis | 6.09E-01 | 2.00E-01 | NaN | SHMT2 |
| Ovarian Cancer Signaling | 6.00E-01 | 6.96E-02 | -0.447 | CDKN2A,E2F1,EDN1,FZD10,PIK3CG,PIK3R3,TP53,VEGFA,WNT1,WNT2B,WNT7B |
| Toll-like Receptor Signaling | 5.91E-01 | 7.69E-02 | -0.447 | IL1A,IL33,TAB2,TICAM2,TLR4,TLR5 |
| Xenobiotic Metabolism PXR Signaling Pathway | 5.81E-01 | 6.74E-02 | 0.277 | ALDH1L2,ALDH2,ALDH3A1,ALDH4A1,GSTA1,GSTT2/GSTT2B,HS3ST5,MAOB,PPARGC1A,PPP1R14A,SMOX,SULT1A1,SULT1C2 |
| Thyroid Cancer Signaling | 5.75E-01 | 7.59E-02 | -0.816 | IGF1,IRS1,NTRK3,PIK3CG,PIK3R3,TP53 |
| Prolactin Signaling | 5.75E-01 | 7.37E-02 | 0 | CEBPB,IRS1,KCNMA1,KCNMB1,PIK3CG,PIK3R3,PLCG2 |
| TGF-β Signaling | 5.61E-01 | 7.29E-02 | -0.447 | BMP2,BMP4,NKX2-5,RUNX2,TGFB3,TGFBR2,TGIF1 |
| Opioid Signaling Pathway | 5.56E-01 | 6.43E-02 | -0.535 | ATF4,CACNG4,CREB3L1,EGR4,FOSB,KCNJ3,KCNJ6,LYN,MAPK15,MAPK4,PIK3CG,POMC,RGS16,RGS4,RGS5,RGS8,SCN7A,SRF |
| Salvage Pathways of Pyrimidine Ribonucleotides | 5.48E-01 | 7.22E-02 | -1.89 | AK7,APOBEC3B,CDK1,MAK,PAK3,SGK1,UPP1 |
| D-myo-inositol (1,4,5,6)-Tetrakisphosphate Biosynthe | 5.45E-01 | 6.67E-02 | -0.333 | ALPL,CA3,DUSP4,DUSP5,DUSP9,EYA4,INPP5B,MTMR8,PPFIA2,PPP1R14A,PSPH,PTPRN |
| D-myo-inositol (3,4,5,6)-tetrakisphosphate Biosynthe | 5.45E-01 | 6.67E-02 | -0.333 | ALPL,CA3,DUSP4,DUSP5,DUSP9,EYA4,INPP5B,MTMR8,PPFIA2,PPP1R14A,PSPH,PTPRN |
| Urea Cycle | 5.42E-01 | 1.67E-01 | NaN | ARG2 |
| Tryptophan Degradation to 2-amino-3-carboxymucor | 5.42E-01 | 1.67E-01 | NaN | KMO |
| Zymosterol Biosynthesis | 5.42E-01 | 1.67E-01 | NaN | MSMO1 |
| NAD Biosynthesis III | 5.42E-01 | 1.67E-01 | NaN | NMNAT2 |
| Prostate Cancer Signaling | 5.40E-01 | 7.02E-02 | NaN | ATF4,CCNE2,CDKN1A,CREB3L1,E2F1,PIK3CG,PIK3R3,TP53 |
| DNA Methylation and Transcriptional Repression Sign | 5.35E-01 | 7.14E-02 | -1.342 | CDK1,CEBPB,E2F1,GADD45B,TCF19,TP53,UHRF1 |
| Natural Killer Cell Signaling | 5.34E-01 | 6.57E-02 | 1.155 | COL5A3,HSPA2,HSPA9,IL2RB,KLRB1,MAP3K15,NFATC2,PAK3,PIK3CG,PIK3R3,PLCG2,ULBP1,VAV1 |
| FLT3 Signaling in Hematopoietic Progenitor Cells | 5.31E-01 | 7.32E-02 | 0.447 | ATF4,CREB3L1,EIF4EBP1,GAB2,PIK3CG,PIK3R3 |
| BEX2 Signaling Pathway | 5.31E-01 | 7.32E-02 | -0.816 | BBC3,BEX2,CDKN1A,LGALS1,NGFR,VEGFA |
| Signaling by Rho Family GTPases | 5.25E-01 | 6.37E-02 | -2.138 | ACTA2,ACTB,ACTC1,ACTG2,ARHGEF2,CIT,DIAPH3,ITGA9,ITGB3,ITGB7,MYL1,MYL6,MYL9,NCF2,PAK3,PIK3CG,PIK3R3 |
| Adrenomedullin signaling pathway | 5.25E-01 | 6.53E-02 | 0.302 | ADM,CEBPB,CTH,IL1A,IL33,MAPK15,MAPK4,MYLK3,PIK3CG,PIK3R3,PLCE1,PLCG2,PPARG |
| HGF Signaling | 5.23E-01 | 6.82E-02 | 0 | CDKN1A,CDKN2A,ITGA9,ITGB3,ITGB7,MAP3K15,PIK3CG,PIK3R3,PLCG2 |
| IL-9 Signaling | 5.21E-01 | 8.57E-02 | NaN | IRS1,PIK3CG,PIK3R3 |
| Ethanol Degradation II | 5.21E-01 | 8.57E-02 | NaN | ALDH2,ALDH3A1,ALDH4A1 |
| Estrogen-Dependent Breast Cancer Signaling | 5.17E-01 | 7.23E-02 | 0 | ATF4,CREB3L1,HSD17B14,IGF1,PIK3CG,PIK3R3 |
| Chondroitin Sulfate Biosynthesis (Late Stages) | 5.14E-01 | 7.84E-02 | 0 | CHST9,HS3ST5,SULT1A1,SULT1C2 |
| Amyloid Processing | 5.14E-01 | 7.84E-02 | NaN | BACE2,CAPN6,CDK5R1,MARK1 |
| HMGB1 Signaling | 5.08E-01 | 6.59E-02 | -1.414 | CCL2,CNTF,IL17B,IL1A,IL33,NGFR,PIK3CG,PIK3R3,TGFB3,TLR4,TNFSF9 |
| Role of MAPK Signaling in the Pathogenesis of Influen | 5.03E-01 | 7.14E-02 | NaN | CCL2,CXCL10,PLA2G3,PLA2G4A,PLA2G4C,PNPLA3 |
| PEDF Signaling | 5.03E-01 | 7.14E-02 | 0.816 | ARHGAP22,PIK3CG,PIK3R3,PPARG,SRF,TP53 |
| Superpathway of Melatonin Degradation | 4.99E-01 | 7.35E-02 | 1.342 | CYP1A1,MAOB,SMOX,SULT1A1,SULT1C2 |
| Neuropathic Pain Signaling In Dorsal Horn Neurons | 4.97E-01 | 6.93E-02 | -0.378 | GRM2,PIK3CG,PIK3R3,PLCE1,PLCG2,TAC1,TACR1 |
| UVB-Induced MAPK Signaling | 4.96E-01 | 7.69E-02 | 1 | EIF4EBP1,PIK3CG,PIK3R3,TP53 |
| Corticotropin Releasing Hormone Signaling | 4.90E-01 | 6.58E-02 | 0.378 | ATF4,CACNG4,CNR1,CREB3L1,CTH,NR4A1,PLCG2,POMC,UCN,VEGFA |
| Insulin Secretion Signaling Pathway | 4.88E-01 | 6.25E-02 | 0.775 | ATF4,CACNG4,CREB3L1,EIF1,EIF2S2,EIF4EBP1,EIF4EBP3,GHR,KLF11,LYN,MAFA,PCSK1,PIK3CG,PIK3R3,PLCE1,PLCG2,RAPGEF4 |
| NAD Biosynthesis from 2-amino-3-carboxymuconate | 4.86E-01 | 1.43E-01 | NaN | NMNAT2 |
| Ceramide Biosynthesis | 4.86E-01 | 1.43E-01 | NaN | SPTLC3 |
| Thioredoxin Pathway | 4.86E-01 | 1.43E-01 | NaN | TXNRD1 |
| NAD Salvage Pathway III | 4.86E-01 | 1.43E-01 | NaN | NMNAT2 |
| Adenine and Adenosine Salvage III | 4.86E-01 | 1.43E-01 | NaN | ADARB2 |
| Aspartate Degradation II | 4.86E-01 | 1.43E-01 | NaN | GOT1 |
| Role of JAK1 and JAK3 in γc Cytokine Signaling | 4.84E-01 | 7.25E-02 | NaN | IL2RB,IRS1,PIK3CG,PIK3R3,TSLP |
| NGF Signaling | 4.71E-01 | 6.67E-02 | 1.134 | ATF4,CREB3L1,MAP3K15,NGFR,PIK3CG,PIK3R3,PLCG2,TP53 |
| Regulation Of The Epithelial Mesenchymal Transition | 4.64E-01 | 6.90E-02 | 0 | FZD10,JAG2,TWIST1,WNT1,WNT2B,WNT7B |
| Reelin Signaling in Neurons | 4.61E-01 | 6.52E-02 | 0.333 | ARHGEF2,CDK5R1,CNR1,DCX,LYN,MAPK8IP2,PDK4,PIK3CG,PIK3R3 |
| White Adipose Tissue Browning Pathway | 4.61E-01 | 6.52E-02 | 1.89 | ATF4,CACNG4,CEBPB,CREB3L1,PPARG,PPARGC1A,PRDM16,SLC16A1,VEGFA |
| Role of NFAT in Cardiac Hypertrophy | 4.60E-01 | 6.25E-02 | -0.832 | AKAP5,CACNG4,GATA4,IGF1,NKX2-5,PIK3CG,PIK3R3,PLCE1,PLCG2,RCAN1,SLC8A2,SLC8A3,TGFB3,TGFBR2 |
| Hepatic Cholestasis | 4.47E-01 | 6.28E-02 | NaN | CNTF,HSD3B7,IL17B,IL1A,IL33,NGFR,SLCO1A2,SREBF1,TGFB3,TJP2,TLR4,TNFSF9 |
| 3-phosphoinositide Degradation | 4.47E-01 | 6.28E-02 | -0.333 | ALPL,CA3,DUSP4,DUSP5,DUSP9,EYA4,INPP5B,MTMR8,PPFIA2,PPP1R14A,PSPH,PTPRN |
| WNT/β-catenin Signaling | 4.45E-01 | 6.32E-02 | 0.632 | CDKN2A,DKK1,DKK2,FZD10,SFRP4,TGFB3,TGFBR2,TP53,WNT1,WNT2B,WNT7B |
| Gαi Signaling | 4.41E-01 | 6.43E-02 | -1.667 | APLNR,CNR1,GPR17,GRM2,NPY1R,P2RY12,RGS4,S1PR1,S1PR3 |
| Antigen Presentation Pathway | 4.41E-01 | 7.69E-02 | NaN | HLA-DMB,HLA-DQA1,HLA-DQB1 |
| Differential Regulation of Cytokine Production in Inte: | 4.41E-01 | 8.70E-02 | NaN | CCL2,IL1A |
| The Visual Cycle | 4.41E-01 | 8.70E-02 | NaN | DHRS3,RPE65 |
| Pyrimidine Deoxyribonucleotides De Novo Biosynthes | 4.41E-01 | 8.70E-02 | NaN | AK7,RRM2 |
| Vitamin-C Transport | 4.41E-01 | 8.70E-02 | NaN | GJB2,TXNRD1 |
| WNK Renal Signaling Pathway | 4.40E-01 | 6.60E-02 | -1.633 | BIRC3,IGF1,IRS1,PIK3CG,PIK3R3,SGK1,WNK4 |
| Xenobiotic Metabolism CAR Signaling Pathway | 4.39E-01 | 6.25E-02 | 0.577 | ALDH1L2,ALDH2,ALDH3A1,ALDH4A1,CYP1A1,FMO2,GSTA1,GSTT2/GSTT2B,HS3ST5,PPARGC1A,SULT1A1,SULT1C2 |
| HER-2 Signaling in Breast Cancer | 4.37E-01 | 6.17E-02 | 1.069 | AREG,CCNE2,CDKN1A,HBEGF,IL1A,ITGB3,ITGB7,LYN,MT-CO1,MT-CO3,PIK3CG,PIK3R3,TP53,TP53RK |
| IL-8 Signaling | 4.34E-01 | 6.19E-02 | 0.577 | CCND3,EIF4EBP1,HBEGF,HMOX1,ITGB3,MYL9,NCF2,PIK3CG,PIK3R3,PLD5,PLD6,TEK,VEGFA |
| EGF Signaling | 4.32E-01 | 7.14E-02 | NaN | CSNK2A3,PIK3CG,PIK3R3,SRF |
| Erythropoietin Signaling Pathway | 4.20E-01 | 6.21E-02 | -0.302 | BIRC3,CNTF,IL17B,IL1A,IL33,PIK3CG,PIK3R3,TGFB3,TNFSF9,TP53,WNT1 |
| Triacylglycerol Degradation | 4.18E-01 | 7.02E-02 | NaN | ALDH2,CEL,PNPLA3,RPE65 |
| Ceramide Signaling | 4.17E-01 | 6.59E-02 | -0.816 | NGFR,PIK3CG,PIK3R3,S1PR1,S1PR3,SPHK1 |
| RANK Signaling in Osteoclasts | 4.17E-01 | 6.59E-02 | -0.447 | BIRC3,MAP3K15,NFATC2,PIK3CG,PIK3R3,TAB2 |
| Huntington's Disease Signaling | 4.13E-01 | 6.01E-02 | -1.732 | ATF4,CAPN6,CASP1,CASP10,CASP4,CASP5,CDK5R1,CREB3L1,HAP1,HSPA2,HSPA9,IGF1,PIK3CG,PIK3R3,RPH3A,SGK1,TP53 |
| Gαs Signaling | 4.11E-01 | 6.35E-02 | 0 | ADRB2,ATF4,CNGA3,CNR1,CREB3L1,HTR7,PTGER4,RAPGEF4 |
| Role of CHK Proteins in Cell Cycle Checkpoint Control | 4.04E-01 | 6.90E-02 | 2 | CDK1,CDKN1A,E2F1,TP53 |
| Endocannabinoid Developing Neuron Pathway | 4.02E-01 | 6.30E-02 | -0.378 | ATF4,CNR1,CREB3L1,MAPK15,MAPK4,PIK3CG,PIK3R3,STMN2 |
| Pathogenesis of Multiple Sclerosis | 4.00E-01 | 1.11E-01 | NaN | CXCL10 |
| Sucrose Degradation V (Mammalian) | 4.00E-01 | 1.11E-01 | NaN | ALDOC |
| Citrulline Biosynthesis | 4.00E-01 | 1.11E-01 | NaN | ARG2 |
| Leucine Degradation I | 4.00E-01 | 1.11E-01 | NaN | BCAT1 |
| Heme Biosynthesis II | 4.00E-01 | 1.11E-01 | NaN | ALAS2 |
| Purine Ribonucleosides Degradation to Ribose-1-phos | 4.00E-01 | 1.11E-01 | NaN | ADARB2 |
| Glycolysis I | 3.96E-01 | 8.00E-02 | NaN | ALDOC,ENO4 |
| Gluconeogenesis I | 3.96E-01 | 8.00E-02 | NaN | ALDOC,ENO4 |
| ERBB Signaling | 3.95E-01 | 6.45E-02 | 0 | AREG,HBEGF,PAK3,PIK3CG,PIK3R3,PLCG2 |
| Leptin Signaling in Obesity | 3.92E-01 | 6.58E-02 | NaN | PIK3CG,PIK3R3,PLCE1,PLCG2,POMC |
| Chondroitin Sulfate Biosynthesis | 3.90E-01 | 6.78E-02 | 0 | CHST9,HS3ST5,SULT1A1,SULT1C2 |
| Intrinsic Prothrombin Activation Pathway | 3.90E-01 | 7.14E-02 | NaN | COL5A3,FGB,KLK5 |
| Superpathway of Methionine Degradation | 3.90E-01 | 7.14E-02 | NaN | CBS/LOC102724560,CTH,GOT1 |
| Non-Small Cell Lung Cancer Signaling | 3.84E-01 | 6.38E-02 | -1 | CCND3,CDKN2A,E2F1,PIK3CG,PIK3R3,TP53 |
| Antiproliferative Role of Somatostatin Receptor 2 | 3.80E-01 | 6.49E-02 | -1 | CDKN1A,CTH,PIK3CG,PIK3R3,SST |
| Retinoic acid Mediated Apoptosis Signaling | 3.77E-01 | 6.67E-02 | 2 | TNFRSF10A,TNFRSF10B,TNFRSF10C,TNFRSF10D |
| SPINK1 Pancreatic Cancer Pathway | 3.77E-01 | 6.67E-02 | 2 | CPA4,KLK5,TGFBR2,TMPRSS15 |
| IL-12 Signaling and Production in Macrophages | 3.75E-01 | 5.93E-02 | 0.535 | CEBPA,CEBPB,COL5A3,DDR2,NFATC2,NFIL3,PIK3CG,PIK3R3,PPARG,SELP,TGFB3,THBS1,TICAM2,TLR4 |
| D-myo-inositol (1,4,5)-Trisphosphate Biosynthesis | 3.75E-01 | 7.69E-02 | NaN | PLCE1,PLCG2 |
| Mitochondrial Dysfunction | 3.69E-01 | 5.81E-02 | -0.447 | ARG2,ATF4,BACE2,BBC3,CACNG4,CAPN6,CREB3L1,GPX3,GSTA1,GSTT2/GSTT2B,HAP1,MAOB,MT-CO1,MT-CO3,NDUFA4L2,PIK3CG,PIK3R3,PPARG,PPARGC1A,TP53 |
| Ketolysis | 3.65E-01 | 1.00E-01 | NaN | ACAT2 |
| Small Cell Lung Cancer Signaling | 3.64E-01 | 6.25E-02 | NaN | CCND3,CCNE2,E2F1,PIK3CG,PIK3R3,TP53 |
| Acute Phase Response Signaling | 3.60E-01 | 5.95E-02 | 0 | A2M,CEBPB,CP,FGB,FTL,HMOX1,IL1A,IL33,NGFR,PIK3CG,PIK3R3 |
| BER (Base Excision Repair) Pathway | 3.59E-01 | 6.82E-02 | NaN | CDKN2A,E2F1,TP53 |
| Renal Cell Carcinoma Signaling | 3.58E-01 | 6.33E-02 | NaN | PAK3,PDGFB,PIK3CG,PIK3R3,VEGFA |
| Role of JAK family kinases in IL-6-type Cytokine Signali | 3.58E-01 | 6.33E-02 | -1.342 | CNTF,IL27RA,POMC,TGFB3,VEGFA |
| IL-2 Signaling | 3.52E-01 | 6.45E-02 | NaN | CSNK2A3,IL2RB,PIK3CG,PIK3R3 |
| P2Y Purigenic Receptor Signaling Pathway | 3.50E-01 | 6.02E-02 | 0 | ATF4,CREB3L1,ITGB3,P2RY12,PIK3CG,PIK3R3,PLCE1,PLCG2 |
| Heparan Sulfate Biosynthesis (Late Stages) | 3.48E-01 | 6.25E-02 | 0 | ALDH2,HS3ST5,RPE65,SULT1A1,SULT1C2 |
| MicroRNA Biogenesis Signaling Pathway | 3.46E-01 | 5.88E-02 | -0.302 | BMP4,EXOSC5,EXOSC6,HMOX1,IGF1,PDGFB,PIK3CG,PIK3R3,TGFB3,TP53,VEGFA |
| Coronavirus Replication Pathway | 3.45E-01 | 6.67E-02 | NaN | TUBB3,TUBB4B,TUBB6 |
| Thrombopoietin Signaling | 3.40E-01 | 6.35E-02 | 1 | GAB2,PIK3CG,PIK3R3,PLCG2 |
| Gαq Signaling | 3.40E-01 | 5.88E-02 | 0.333 | ADRA1A,HMOX1,NFATC2,PIK3CG,PIK3R3,PLCG2,PLD5,PLD6,RGS16,RGS4 |
| Type II Diabetes Mellitus Signaling | 3.33E-01 | 5.88E-02 | -1.134 | CACNG4,CD36,CEBPB,IRS1,MAFA,NGFR,PIK3CG,PIK3R3,PPARG |
| Cysteine Biosynthesis III (mammalia) | 3.21E-01 | 6.90E-02 | NaN | CBS/LOC102724560,CTH |
| RAC Signaling | 3.18E-01 | 5.84E-02 | 0.447 | CDK5R1,ITGA9,ITGB3,ITGB7,NCF2,PAK3,PIK3CG,PIK3R3 |
| Retinol Biosynthesis | 3.18E-01 | 6.38E-02 | NaN | CEL,DHRS3,PNPLA3 |
| PXR/RXR Activation | 3.18E-01 | 6.15E-02 | -2 | CPT1A,GSTA1,PCK2,PPARGC1A |
| Induction of Apoptosis by HIV1 | 3.18E-01 | 6.15E-02 | 1 | BBC3,BIRC3,NGFR,TP53 |
| Pyridoxal 5'-phosphate Salvage Pathway | 3.18E-01 | 6.15E-02 | -2 | CDK1,MAK,PAK3,SGK1 |
| Nitric Oxide Signaling in the Cardiovascular System | 3.10E-01 | 5.83E-02 | 0 | ARG2,CACNG4,PIK3CG,PIK3R3,PLN,SLC7A1,VEGFA |
| Dermatan Sulfate Biosynthesis (Late Stages) | 3.06E-01 | 6.25E-02 | NaN | HS3ST5,SULT1A1,SULT1C2 |
| Neuroprotective Role of THOP1 in Alzheimer's Disease | 3.02E-01 | 5.79E-02 | -1.342 | KLK5,LONP1,MASP1,SST,TAC1,TMPRSS15,TMPRSS7 |
| Insulin Receptor Signaling | 2.97E-01 | 5.71E-02 | -1.134 | EIF4EBP1,GRB10,INPP5B,IRS1,PIK3CG,PIK3R3,PPP1R14A,SGK1 |
| MSP-RON Signaling In Cancer Cells Pathway | 2.97E-01 | 5.71E-02 | -0.378 | ACTA2,ATF4,CREB3L1,FLNA,KLK5,PIK3CG,PIK3R3,VEGFA |
| Circadian Rhythm Signaling | 2.95E-01 | 5.60E-02 | NaN | ATF4,CACNG4,CREB3L1,CTH,EIF4EBP1,EIF4EBP3,FBXL21P,LYN,MKNK2,MTNR1B,NFIL3,NGFR,PLCE1,PLCG2,PTGDS |
| Apoptosis Signaling | 2.93E-01 | 5.77E-02 | 0 | BIRC3,CAPN6,CASP10,CDK1,PLCG2,TP53 |
| Oleate Biosynthesis II (Animals) | 2.84E-01 | 7.69E-02 | NaN | FADS2 |
| Heparan Sulfate Biosynthesis | 2.81E-01 | 5.75E-02 | 0 | ALDH2,HS3ST5,RPE65,SULT1A1,SULT1C2 |
| Synaptic Long Term Depression | 2.78E-01 | 5.56E-02 | -0.632 | CACNG4,CTH,GRM2,IGF1,LYN,PLA2G3,PLA2G4A,PLA2G4C,PLCE1,PLCG2,PNPLA3 |
| Regulation of eIF4 and p70S6K Signaling | 2.70E-01 | 5.52E-02 | NaN | EIF1,EIF2S2,EIF4EBP1,EIF4EBP3,IRS1,ITGA9,ITGB3,ITGB7,PIK3CG,PIK3R3 |
| Phagosome Maturation | 2.67E-01 | 5.52E-02 | NaN | ATP6V0D2,CTSC,CTSV,DYNC1I1,NCF2,TUBB3,TUBB4B,TUBB6,TUBE1 |
| Telomerase Signaling | 2.62E-01 | 5.56E-02 | 0 | CDKN1A,E2F1,IL2RB,PIK3CG,PIK3R3,TP53 |
| 14-3-3-mediated Signaling | 2.59E-01 | 5.51E-02 | 0 | PIK3CG,PIK3R3,PLCE1,PLCG2,TUBB3,TUBB4B,TUBB6 |
| Netrin Signaling | 2.50E-01 | 5.56E-02 | -1 | CACNG4,NFATC2,UNC5A,UNC5B |
| 4-1BB Signaling in T Lymphocytes | 2.49E-01 | 5.88E-02 | NaN | TNFRSF9,TNFSF9 |
| Acute Myeloid Leukemia Signaling | 2.49E-01 | 5.49E-02 | -1 | CEBPA,EIF4EBP1,KIT,PIK3CG,PIK3R3 |
| BMP signaling pathway | 2.49E-01 | 5.49E-02 | 0 | BMP2,BMP4,GREM1,NKX2-5,RUNX2 |
| Superpathway of Citrulline Metabolism | 2.43E-01 | 6.67E-02 | NaN | ARG2 |
| Leukotriene Biosynthesis | 2.43E-01 | 6.67E-02 | NaN | GGT5 |
| Role of Cytokines in Mediating Communication betwe | 2.40E-01 | 5.56E-02 | NaN | IL1A,IL32,IL33 |
| Coagulation System | 2.37E-01 | 5.71E-02 | NaN | A2M,FGB |
| Granzyme B Signaling | 2.26E-01 | 6.25E-02 | NaN | LMNB1 |
| Extrinsic Prothrombin Activation Pathway | 2.26E-01 | 6.25E-02 | NaN | FGB |
| Bile Acid Biosynthesis, Neutral Pathway | 2.26E-01 | 6.25E-02 | NaN | HSD3B7 |
| D-myo-inositol (1,4,5)-trisphosphate Degradation | 2.26E-01 | 6.25E-02 | NaN | INPP5B |
| Adenosine Nucleotides Degradation II | 2.26E-01 | 6.25E-02 | NaN | ADARB2 |
| Parkinson's Signaling | 2.26E-01 | 6.25E-02 | NaN | UCHL1 |

| | | | | |
|---|---|---|---|---|
| 1D-myo-inositol Hexakisphosphate Biosynthesis II (M | 2.10E-01 | 5.88E-02 | NaN | INPP5B |
| D-myo-inositol (1,3,4)-trisphosphate Biosynthesis | 2.10E-01 | 5.88E-02 | NaN | INPP5B |
| Differential Regulation of Cytokine Production in Mac | 1.95E-01 | 5.56E-02 | NaN | CCL2 |
| CSDE1 Signaling Pathway | 0.00E+00 | 3.57E-02 | NaN | CCL2,CCND3 |
| Oxytocin In Spinal Neurons Signaling Pathway | 0.00E+00 | 2.86E-02 | NaN | CTH |
| Synaptic Long Term Potentiation | 0.00E+00 | 4.55E-02 | 0.447 | ATF4,CREB3L1,GRM2,PLCE1,PLCG2,PPP1R14A |
| Chaperone Mediated Autophagy Signaling Pathway | 0.00E+00 | 2.95E-02 | -0.229 | ARHGAP8/PRR5-ARHGAP8,ATP6V0D2,BBC3,CASP1,EEF1A2,HK2,HSPA2,HSPA9,MMP10,MMP15,MMP17,MMP19,MMP24,NFATC2,PIK3CG,PIK3R3,RCAN1,TP53,UCHL1 |
| PPARα/RXRα Activation | 0.00E+00 | 4.10E-02 | 1.633 | CD36,GHR,IRS1,PLCE1,PLCG2,PPARGC1A,TGFB3,TGFBR2 |
| FXR/RXR Activation | 0.00E+00 | 4.76E-02 | NaN | IL1A,IL33,PCK2,PPARG,PPARGC1A,SREBF1 |
| α-Adrenergic Signaling | 0.00E+00 | 4.59E-02 | -0.447 | ADRA1A,PHKG1,PLCG2,SLC8A2,SLC8A3 |
| Role of RIG1-like Receptors in Antiviral Innate Immuni | 0.00E+00 | 2.17E-02 | NaN | CASP10 |
| Role of NFAT in Regulation of the Immune Response | 0.00E+00 | 1.25E-02 | 0.905 | AKAP5,CD86,GATA4,HLA-DMB,HLA-DQA1,HLA-DQB1,IGLV1-50,LYN,NFATC2,PIK3CG,PIK3R3,PLCG2,RCAN1 |
| FcγRIIB Signaling in B Lymphocytes | 0.00E+00 | 1.13E-02 | 1 | CACNG4,IGLV1-50,LYN,PIK3CG,PIK3R3,PLCG2 |
| LPS-stimulated MAPK Signaling | 0.00E+00 | 4.71E-02 | 0 | PIK3CG,PIK3R3,SRF,TLR4 |
| NF-κB Activation by Viruses | 0.00E+00 | 3.85E-02 | NaN | ITGB3,PIK3CG,PIK3R3 |
| CCR5 Signaling in Macrophages | 0.00E+00 | 4.01E-03 | NaN | CACNG4,PLCG2 |
| IL-22 Signaling | 0.00E+00 | 4.17E-02 | NaN | IL22RA1 |
| CD40 Signaling | 0.00E+00 | 2.99E-02 | NaN | PIK3CG,PIK3R3 |
| Calcium-induced T Lymphocyte Apoptosis | 0.00E+00 | 1.09E-02 | 0.447 | HLA-DMB,HLA-DQA1,HLA-DQB1,NFATC2,NR4A1 |
| CD27 Signaling in Lymphocytes | 0.00E+00 | 1.75E-02 | NaN | MAP3K15 |
| IL-3 Signaling | 0.00E+00 | 3.80E-02 | NaN | GAB2,PIK3CG,PIK3R3 |
| Lymphotoxin β Receptor Signaling | 0.00E+00 | 3.64E-02 | NaN | PIK3CG,PIK3R3 |
| fMLP Signaling in Neutrophils | 0.00E+00 | 3.05E-02 | 0 | NCF2,NFATC2,PIK3CG,PIK3R3 |
| CXCR4 Signaling | 0.00E+00 | 5.36E-02 | -1.667 | CXCL12,EGR1,LYN,MYL1,MYL6,MYL9,PAK3,PIK3CG,PIK3R3 |
| CTLA4 Signaling in Cytotoxic T Lymphocytes | 0.00E+00 | 2.30E-02 | -0.832 | CD86,HLA-DMB,HLA-DQA1,HLA-DQB1,HMOX1,LYN,NFATC2,PIK3CG,PIK3R3,PLCG2,PLD5,PLD6,TGFB3,VAV1 |
| B Cell Activating Factor Signaling | 0.00E+00 | 2.33E-02 | NaN | NFATC2 |
| T Helper Cell Differentiation | 0.00E+00 | 1.49E-02 | NaN | CD86,HLA-DMB,HLA-DQA1,HLA-DQB1,ICOSLG/LOC102723996,NGFR,TGFBR2 |
| CCR3 Signaling in Eosinophils | 0.00E+00 | 4.48E-02 | 0 | PAK3,PIK3CG,PIK3R3,PLA2G3,PLA2G4A,PLA2G4C |
| CD28 Signaling in T Helper Cells | 0.00E+00 | 1.55E-02 | -0.447 | CD86,HLA-DMB,HLA-DQA1,HLA-DQB1,NFATC2,PIK3CG,PIK3R3,VAV1 |
| IL-15 Signaling | 0.00E+00 | 7.58E-03 | NaN | IGLV1-50,IL2RB,PIK3CG,PIK3R3 |
| Dendritic Cell Maturation | 0.00E+00 | 3.02E-02 | 0.243 | ATF4,CD86,COL5A3,CREB3L1,DDR2,FSCN2,HLA-DMB,HLA-DQA1,HLA-DQB1,IL1A,IL32,IL33,NGFR,PIK3CG,PIK3R3,PLCE1,PLCG2,TLR4 |
| Melatonin Signaling | 0.00E+00 | 4.17E-02 | NaN | MTNR1B,PLCE1,PLCG2 |
| Angiopoietin Signaling | 0.00E+00 | 5.26E-02 | NaN | PAK3,PIK3CG,PIK3R3,TEK |
| Relaxin Signaling | 0.00E+00 | 3.23E-02 | NaN | CTH,PIK3CG,PIK3R3,PLD6,VEGFA |
| CNTF Signaling | 0.00E+00 | 5.26E-02 | NaN | CNTF,PIK3CG,PIK3R3 |
| Renin-Angiotensin Signaling | 0.00E+00 | 4.13E-02 | -0.447 | CCL2,PAK3,PIK3CG,PIK3R3,PLCG2 |
| Docosahexaenoic Acid (DHA) Signaling | 0.00E+00 | 5.26E-02 | NaN | PIK3CG,PIK3R3 |
| Semaphorin Signaling in Neurons | 0.00E+00 | 1.64E-02 | NaN | PAK3 |
| Thrombin Signaling | 0.00E+00 | 4.44E-02 | 1.134 | ARHGEF2,F2RL2,GATA4,MYL1,MYL6,MYL9,PIK3CG,PIK3R3,PLCE1,PLCG2 |
| ICOS-ICOSL Signaling in T Helper Cells | 0.00E+00 | 2.17E-02 | 1.897 | CD86,GAB2,HLA-DMB,HLA-DQA1,HLA-DQB1,ICOSLG/LOC102723996,IL2RB,NFATC2,PIK3CG,PIK3R3,VAV1 |
| Cell Cycle Regulation by BTG Family Proteins | 0.00E+00 | 5.26E-02 | NaN | CCNE2,E2F1 |
| Maturity Onset Diabetes of Young (MODY) Signaling | 0.00E+00 | 5.06E-02 | NaN | APOL6,CACNG4,CEL,KLF11 |
| GNRH Signaling | 0.00E+00 | 3.66E-02 | -0.447 | ATF4,CACNG4,CREB3L1,EGR1,HBEGF,MAP3K15,PAK3 |
| Cholecystokinin/Gastrin-mediated Signaling | 0.00E+00 | 4.20E-02 | -1.342 | EPHA4,IL1A,IL33,SRF,SST |
| Androgen Signaling | 0.00E+00 | 1.78E-02 | NaN | CACNG4,TAF4B,TGFB1I1 |
| Role of OCT4 in Mammalian Embryonic Stem Cell Plur | 0.00E+00 | 2.17E-02 | NaN | TP53 |
| Type I Diabetes Mellitus Signaling | 0.00E+00 | 1.18E-02 | NaN | CD86,HLA-DMB,HLA-DQA1,HLA-DQB1,NGFR,PTPRN |
| Endometrial Cancer Signaling | 0.00E+00 | 5.00E-02 | NaN | PIK3CG,PIK3R3,TP53 |
| Allograft Rejection Signaling | 0.00E+00 | 8.21E-03 | NaN | CD86,HLA-DMB,HLA-DQA1,HLA-DQB1 |
| Autoimmune Thyroid Disease Signaling | 0.00E+00 | 8.73E-03 | NaN | CD86,HLA-DMB,HLA-DQA1,HLA-DQB1 |
| Graft-versus-Host Disease Signaling | 0.00E+00 | 1.35E-02 | NaN | CD86,HLA-DMB,HLA-DQA1,HLA-DQB1,IL1A,IL33 |
| Production of Nitric Oxide and Reactive Oxygen Speci | 0.00E+00 | 4.71E-02 | 1 | ARG2,MAP3K15,NCF2,NGFR,PIK3CG,PIK3R3,PLCG2,PPP1R14A,TLR4 |
| Gα12/13 Signaling | 0.00E+00 | 5.26E-02 | -0.378 | F2RL2,MYL1,MYL6,MYL9,PIK3CG,PIK3R3,VAV1 |
| p70S6K Signaling | 0.00E+00 | 1.38E-02 | 0.378 | F2RL2,IGLV1-50,IRS1,LYN,PIK3CG,PIK3R3,PLCE1,PLCG2 |
| ERK5 Signaling | 0.00E+00 | 5.41E-02 | NaN | ATF4,CREB3L1,SGK1,SH2D2A |
| mTOR Signaling | 0.00E+00 | 5.14E-02 | 0.816 | ARHGAP8/PRR5-ARHGAP8,DDIT4,EIF4EBP1,EIF4EBP3,HMOX1,IRS1,PIK3CG,PIK3R3,PLD5,PLD6,VEGFA |
| G Beta Gamma Signaling | 0.00E+00 | 4.65E-02 | 0 | CACNG4,HBEGF,KCNJ3,KCNJ6,PIK3CG,PLCG2 |
| G Protein Signaling Mediated by Tubby | 0.00E+00 | 4.27E-03 | NaN | PLCG2,VAV1 |
| Communication between Innate and Adaptive Immun | 0.00E+00 | 7.50E-03 | NaN | CD86,CXCL10,IGLV1-50,IL1A,IL33,TLR4,TLR5 |
| Systemic Lupus Erythematosus Signaling | 0.00E+00 | 8.43E-03 | NaN | CD86,IGLV1-50,IL1A,IL33,LYN,NFATC2,PIK3CG,PIK3R3,PLCG2 |
| CDC42 Signaling | 0.00E+00 | 2.26E-02 | -1.89 | DIAPH3,FGD3,HLA-DMB,HLA-DQA1,HLA-DQB1,ITGA9,ITGB3,ITGB7,MYL1,MYL6,MYL9,PAK3,VAV1 |
| FAK Signaling | 0.00E+00 | 5.19E-02 | -0.816 | ACKR3,ADGRA2,ADGRL3,ADRA1A,ADRB2,APLNR,CAPN6,CMKLR2,CNR1,COL5A3,EFNA5,F2RL2,FZD10,GALR1,GHR,GPR137C,GPR157,GPR160,GPR17,GPR183,GPR37L1,GPRC5A,GRM2,HRH4,HTR7,IFNLR1,IL17RE,IL22RA1,IL27RA,IL2RB,IL31RA,IL33,ITGA9,ITGB3,ITGB7,KIT,MAPK4,MTNR1B,NPY1R,P2RY12,PAK3,PDGFRA,PDGFRB,PIK3CG,PIK3R3,PLCG2,PTGER4,S1PR1,S1PR3,SH2D2A,TACR1,TGFB3,TGFBR2,TP53 |
| April Mediated Signaling | 0.00E+00 | 2.38E-02 | NaN | NFATC2 |
| Phospholipase C Signaling | 0.00E+00 | 1.96E-02 | -0.535 | AHNAK,ARHGEF2,ATF4,CREB3L1,HMOX1,IGLV1-50,ITGA9,ITGB3,ITGB7,LYN,MYL1,MYL6,MYL9,NFATC2,PLA2G3,PLA2G4A,PLA2G4C,PLCE1,PLCG2,PLD5,PLD6,PPP1R14A |
| Altered T Cell and B Cell Signaling in Rheumatoid Arth | 0.00E+00 | 9.57E-03 | NaN | CD86,HLA-DMB,HLA-DQA1,HLA-DQB1,IGLV1-50,IL1A,IL33,TLR4,TLR5 |
| Glioma Invasiveness Signaling | 0.00E+00 | 4.11E-02 | NaN | ITGB3,PIK3CG,PIK3R3 |
| B Cell Development | 0.00E+00 | 1.02E-02 | NaN | CD86,HLA-DMB,HLA-DQA1,HLA-DQB1,IGLV1-50 |
| IL-1 Signaling | 0.00E+00 | 2.08E-02 | NaN | IL1A,TAB2 |
| Regulation of IL-2 Expression in Activated and Anergic | 0.00E+00 | 1.08E-02 | 1.342 | NFATC2,PLCG2,TGFB3,TGFBR2,VAV1 |
| Granzyme A Signaling | 0.00E+00 | 5.33E-02 | 0 | CASP1,H1-0,LMNB1,NDUFA4L2 |
| Role of WNT/GSK-3β Signaling in the Pathogenesis of | 0.00E+00 | 5.13E-02 | -1 | FZD10,WNT1,WNT2B,WNT7B |
| TWEAK Signaling | 0.00E+00 | 5.41E-02 | NaN | BIRC3,TNFRSF25 |
| NUR77 Signaling in T Lymphocytes | 0.00E+00 | 9.75E-03 | NaN | CD86,HLA-DMB,HLA-DQA1,HLA-DQB1,NR4A1 |
| PKCθ Signaling in T Lymphocytes | 0.00E+00 | 1.97E-02 | 1.265 | CACNG4,CD86,HLA-DMB,HLA-DQA1,HLA-DQB1,MAP3K15,NFATC2,PIK3CG,PIK3R3,PLCG2,VAV1 |
| TNFR1 Signaling | 0.00E+00 | 3.92E-02 | NaN | BIRC3,PAK3 |
| TNFR2 Signaling | 0.00E+00 | 3.12E-02 | NaN | BIRC3 |
| Role of PI3K/AKT Signaling in the Pathogenesis of Infl | 0.00E+00 | 3.03E-02 | NaN | PIK3CG,PIK3R3 |
| Antiproliferative Role of TOB in T Cell Signaling | 0.00E+00 | 7.04E-03 | NaN | CCNE2,TGFB3,TGFBR2 |
| OX40 Signaling Pathway | 0.00E+00 | 6.28E-03 | NaN | HLA-DMB,HLA-DQA1,HLA-DQB1 |
| PI3K Signaling in B Lymphocytes | 0.00E+00 | 2.03E-02 | 1.897 | ATF3,ATF4,ATF5,IGLV1-50,IRS1,LYN,NFATC2,PIK3CG,PLCE1,PLCG2,TLR4,VAV1 |
| Assembly of RNA Polymerase II Complex | 0.00E+00 | 2.00E-02 | NaN | TAF4B |
| IL-17A Signaling in Airway Cells | 0.00E+00 | 2.99E-02 | NaN | PIK3CG,PIK3R3 |
| Role of IL-17A in Arthritis | 0.00E+00 | 5.26E-02 | NaN | CCL2,PIK3CG,PIK3R3 |
| IL-17A Signaling in Fibroblasts | 0.00E+00 | 5.26E-02 | NaN | CCL2,CEBPB |
| IL-17A Signaling in Gastric Cells | 0.00E+00 | 3.85E-02 | NaN | CXCL10 |
| Actin Nucleation by ARP-WASP Complex | 0.00E+00 | 3.23E-02 | NaN | ITGA9,ITGB3,ITGB7 |
| Dopamine-DARPP32 Feedback in cAMP Signaling | 0.00E+00 | 4.84E-02 | 1 | ATF4,CACNG4,CREB3L1,KCNJ3,KCNJ4,KCNJ6,PLCE1,PLCG2,PPP1R14A |
| Mouse Embryonic Stem Cell Pluripotency | 0.00E+00 | 4.81E-02 | -1.342 | BMP4,FZD10,PIK3CG,PIK3R3,TP53 |
| Hematopoiesis from Pluripotent Stem Cells | 0.00E+00 | 2.25E-03 | NaN | IL1A |
| iNOS Signaling | 0.00E+00 | 4.26E-02 | NaN | HMGA1,TLR4 |
| nNOS Signaling in Neurons | 0.00E+00 | 2.13E-02 | NaN | CAPN6 |
| nNOS Signaling in Skeletal Muscle Cells | 0.00E+00 | 2.08E-02 | NaN | CACNG4 |
| Ephrin B Signaling | 0.00E+00 | 4.17E-02 | NaN | CXCL12,EPHB6,VAV1 |
| ERB2-ERBB3 Signaling | 0.00E+00 | 3.08E-02 | NaN | PIK3CG,PIK3R3 |
| ERBB4 Signaling | 0.00E+00 | 4.41E-02 | NaN | PIK3CG,PIK3R3,PLCG2 |
| Bupropion Degradation | 0.00E+00 | 4.00E-02 | NaN | CYP1A1 |
| Retinoate Biosynthesis I | 0.00E+00 | 2.38E-02 | NaN | DHRS3 |
| Thyroid Hormone Metabolism II (via Conjugation and | 0.00E+00 | 4.88E-02 | NaN | SULT1A1,SULT1C2 |
| Nicotine Degradation III | 0.00E+00 | 1.69E-02 | NaN | CYP1A1 |
| Pyrimidine Ribonucleotides De Novo Biosynthesis | 0.00E+00 | 4.76E-02 | NaN | AK7,ENTPD3 |
| Ubiquinol-10 Biosynthesis (Eukaryotic) | 0.00E+00 | 3.23E-02 | NaN | CYP26A1 |
| tRNA Splicing | 0.00E+00 | 2.17E-02 | NaN | PLD6 |
| Melatonin Degradation I | 0.00E+00 | 4.76E-02 | NaN | CYP1A1,SULT1A1,SULT1C2 |
| γ-linolenate Biosynthesis II (Animals) | 0.00E+00 | 5.26E-02 | NaN | FADS2 |
| NAD Salvage Pathway II | 0.00E+00 | 3.70E-02 | NaN | NMNAT2 |
| Estrogen Biosynthesis | 0.00E+00 | 4.26E-02 | NaN | CYP1A1,HSD17B14 |
| Dermatan Sulfate Biosynthesis | 0.00E+00 | 4.92E-02 | NaN | HS3ST5,SULT1A1,SULT1C2 |
| Glutathione-mediated Detoxification | 0.00E+00 | 5.41E-02 | NaN | GSTA1,GSTT2/GSTT2B |
| Nicotine Degradation II | 0.00E+00 | 4.48E-02 | NaN | CYP1A1,FMO2,INMT |
| Valine Degradation I | 0.00E+00 | 5.00E-02 | NaN | BCAT1 |
| Superpathway of D-myo-inositol (1,4,5)-trisphosphat | 0.00E+00 | 4.55E-02 | NaN | INPP5B |
| Purine Nucleotides Degradation II (Aerobic) | 0.00E+00 | 5.26E-02 | NaN | ADARB2 |
| Mitochondrial L-carnitine Shuttle Pathway | 0.00E+00 | 5.26E-02 | NaN | CPT1A |
| Triacylglycerol Biosynthesis | 0.00E+00 | 3.64E-02 | NaN | GPAT3,PNPLA3 |
| Pyrimidine Ribonucleotides Interconversion | 0.00E+00 | 5.13E-02 | NaN | AK7,ENTPD3 |
| Histidine Degradation VI | 0.00E+00 | 3.57E-02 | NaN | CYP26A1 |
| Stearate Biosynthesis I (Animals) | 0.00E+00 | 2.90E-02 | NaN | DHCR24,TBXAS1 |
| Pregnenolone Biosynthesis | 0.00E+00 | 3.70E-02 | NaN | CYP26A1 |
| Acetone Degradation I (to Methylglyoxal) | 0.00E+00 | 2.38E-02 | NaN | CYP1A1 |
| Oxidative Ethanol Degradation III | 0.00E+00 | 5.26E-02 | NaN | ALDH2,ALDH3A1,ALDH4A1 |
| Epithelial Adherens Junction Signaling | 0.00E+00 | 5.10E-02 | -2.121 | KIF23,MYH10,MYH2,NANOS1,PAK3,RACGAP1,TGFBR2,TNS1 |
| TEC Kinase Signaling | 0.00E+00 | 2.94E-02 | 0 | ACTA2,ACTB,ACTC1,ACTG2,ITGA9,ITGB3,ITGB7,LYN,PAK3,PIK3CG,PIK3R3,PLCG2,TLR4,TNFRSF10A,TNFRSF10B,TNFRSF25,VAV1 |
| UVA-Induced MAPK Signaling | 0.00E+00 | 5.10E-02 | 0 | PIK3CG,PIK3R3,PLCE1,PLCG2,TP53 |
| UVC-Induced MAPK Signaling | 0.00E+00 | 1.96E-02 | NaN | TP53 |
| Oxidative Phosphorylation | 0.00E+00 | 1.80E-02 | NaN | MT-CO1,MT-CO3 |
| HIPPO signaling | 0.00E+00 | 2.33E-02 | NaN | PPP1R14A,TJP2 |
| Nucleotide Excision Repair Pathway | 0.00E+00 | 2.86E-02 | NaN | RPA4 |
| SAPK/JNK Signaling | 0.00E+00 | 1.39E-02 | -0.816 | DUSP4,IRS1,MAPK8IP2,PIK3CG,PIK3R3,SH2D2A,TP53 |
| Cardiac β-adrenergic Signaling | 0.00E+00 | 3.89E-02 | NaN | AKAP5,CACNG4,PLD6,PLN,PPP1R14A,SLC8A2,SLC8A3 |
| Protein Ubiquitination Pathway | 0.00E+00 | 3.30E-02 | NaN | BIRC3,CRYAB,DNAJB13,HSPA12A,HSPA2,HSPA9,UBE2C,UCHL1,USP43 |
| JAK/STAT Signaling | 0.00E+00 | 4.88E-02 | 0 | CDKN1A,CEBPB,PIK3CG,PIK3R3 |
| GABA Receptor Signaling | 0.00E+00 | 5.30E-02 | NaN | CACNG4,GABRA2,GABRA3,GABRA4,GABRB2,GABRG1,GABRQ |
| IL-4 Signaling | 0.00E+00 | 3.47E-02 | -0.894 | ARHGAP8/PRR5-ARHGAP8,ATF4,COL12A1,COL21A1,COL26A1,COL5A1,COL5A3,COL6A6,COL9A1,CREB3L1,EIF4EBP1,HLA-DMB,HLA-DQA1,HLA-DQB1,IRS1,NFATC2,NFIL3,PIK3CG,PIK3R3,TGFB3 |
| B Cell Receptor Signaling | 0.00E+00 | 2.05E-02 | 0.905 | ATF4,CREB3L1,EGR1,GAB2,IGLV1-50,INPP5B,LYN,MAP3K15,NFATC2,PIK3CG,PIK3R3,PLCG2,VAV1 |
| Phototransduction Pathway | 0.00E+00 | 1.85E-02 | NaN | CNGA3 |
| Chemokine Signaling | 0.00E+00 | 4.94E-02 | 0 | CCL2,CXCL12,PIK3CG,PLCG2 |
| Dopamine Receptor Signaling | 0.00E+00 | 3.75E-02 | NaN | MAOB,PPP1R14A,SMOX |
| Glutamate Receptor Signaling | 0.00E+00 | 3.03E-02 | NaN | GRM2,SLC38A1 |
| Notch Signaling | 0.00E+00 | 5.26E-02 | NaN | HEY2,JAG2 |

| | | | | |
|---|---|---|---|---|
| NF-κB Signaling | 0.00E+00 | 2.98E-02 | -1.5 | BMP2,BMP4,CSNK2A3,GHR,IL1A,IL33,NGFR,NTRK3,PDGFRA,PDGFRB,PIK3CG,PIK3R3,PLCG2,TAB2,TGFBR2,TLR4,TLR5 |
| T Cell Receptor Signaling | 0.00E+00 | 1.79E-02 | 1.265 | CD86,DUSP5,HLA-DMB,HLA-DQA1,HLA-DQB1,ICOSLG/LOC102723996,NFATC2,PIK3CG,PIK3R3,PLCG2,VAV1 |
| GPCR-Mediated Nutrient Sensing in Enteroendocrine | 0.00E+00 | 3.39E-02 | 0 | CACNG4,PLCE1,PLCG2,RAPGEF4 |
| Sumoylation Pathway | 0.00E+00 | 2.91E-02 | NaN | CEBPA,ISG20,TP53 |
| IL-7 Signaling Pathway | 0.00E+00 | 5.13E-02 | 0 | CCND3,LYN,PIK3CG,PIK3R3 |
| Sirtuin Signaling Pathway | 0.00E+00 | 4.78E-02 | 1.897 | ARG2,CPT1A,E2F1,GADD45B,H1-0,MAPK15,MAPK4,NDUFA4L2,PCK2,PPARG,PPARGC1A,SREBF1,TIMM44,TP53 |
| Th17 Activation Pathway | 0.00E+00 | 4.13E-03 | NaN | NFATC2,PTGER4 |
| NER (Nucleotide Excision Repair, Enhanced Pathway) | 0.00E+00 | 3.33E-02 | NaN | RPA4,TCEA1,TOP2A |
| SPINK1 General Cancer Pathway | 0.00E+00 | 2.90E-02 | NaN | PIK3CG,PIK3R3 |
| Endocannabinoid Neuronal Synapse Pathway | 0.00E+00 | 5.37E-02 | -1.414 | CACNG4,CNR1,KCNJ3,KCNJ6,MAPK15,MAPK4,PLCE1,PLCG2 |
| Apelin Endothelial Signaling Pathway | 0.00E+00 | 3.55E-02 | -1.342 | APLNR,CCL2,PIK3CG,PIK3R3,TEK |
| Apelin Muscle Signaling Pathway | 0.00E+00 | 4.17E-02 | NaN | APLNR,PPARGC1A |
| BAG2 Signaling Pathway | 0.00E+00 | 4.76E-02 | 1 | CDKN1A,HSPA2,HSPA9,TP53 |
| FAT10 Signaling Pathway | 0.00E+00 | 1.79E-02 | NaN | SQSTM1 |
| T Cell Exhaustion Signaling Pathway | 0.00E+00 | 1.76E-02 | -1.134 | CD86,HLA-DMB,HLA-DQA1,HLA-DQB1,NFATC2,PIK3CG,PIK3R3,PLCG2,TGFBR2,VEGFA |
| IL-23 Signaling Pathway | 0.00E+00 | 4.35E-02 | NaN | PIK3CG,PIK3R3 |
| Systemic Lupus Erythematosus In T Cell Signaling Path | 0.00E+00 | 2.64E-02 | -1 | ATF4,CASP1,CASP10,CASP4,CASP5,CD86,CREB3L1,HLA-DMB,HLA-DQA1,HLA-DQB1,ICOSLG/LOC102723996,NFATC2,PIK3CG,PIK3R3,S1PR3,SELPLG,VAV1 |
| Systemic Lupus Erythematosus In B Cell Signaling Path | 0.00E+00 | 2.21E-02 | -0.775 | CCND3,CNTF,IGLV1-50,IL17B,IL1A,IL33,INPP5B,ISG20,LYN,NFATC2,PIK3CG,PIK3R3,PLCG2,TGFB3,TNFSF9,VAV1 |
| Inhibition of ARE-Mediated mRNA Degradation Pathw | 0.00E+00 | 4.32E-02 | -1.89 | EXOSC5,EXOSC6,MAPK15,MAPK4,NGFR,POMC,TNFSF9 |
| Xenobiotic Metabolism General Signaling Pathway | 0.00E+00 | 4.90E-02 | 0.378 | FTL,GSTA1,GSTT2/GSTT2B,HMOX1,MAP3K15,PIK3CG,PIK3R3 |
| Coronavirus Pathogenesis Pathway | 0.00E+00 | 4.90E-02 | 1.265 | ATF4,CASP1,CCL2,CCNE2,DDIT3,E2F1,EEF1A2,OAS3,TGFBR2,TP53 |
| GM-CSF Signaling | 0.00E+00 | 4.29E-02 | NaN | LYN,PIK3CG,PIK3R3 |
| Complement System | 0.00E+00 | 5.41E-02 | NaN | CD55,MASP1 |
| Role of MAPK Signaling in Promoting the Pathogenesi | 0.00E+00 | 4.39E-02 | 0.447 | ATP6V0D2,PLA2G3,PLA2G4A,PLA2G4C,PNPLA3 |

IPA_Astrocytes_GCSi_FC2_adj0.05_409genes

| | Ingenuity Canonical Pathways | -log(p-value) | Ratio | z-score | Molecules |
|---|---|---|---|---|---|
| 1 | LXR/RXR Activation | 8.88E+00 | 1.14E-01 | -3.162 | ABCG1,AGT,CCL2,IL18,IL1R1,LPL,ORM1,ORM2,RELB,RXRG,SAA1,SAA2,SCD,SREBF1 |
| 2 | Acute Phase Response Signaling | 5.06E+00 | 6.49E-02 | 0.816 | AGT,FOS,HP,IL18,IL1R1,ORM1,ORM2,RELB,SAA1,SAA2,SERPINE1,VWF |
| 3 | HMGB1 Signaling | 4.74E+00 | 6.59E-02 | 2.121 | CCL2,CLCF1,CXCL8,FOS,ICAM1,IL11,IL18,IL1R1,LIF,RELB,SERPINE1 |
| 4 | TNFR2 Signaling | 4.18E+00 | 1.56E-01 | NaN | BIRC3,FOS,RELB,TNFAIP3,TRAF1 |
| 5 | Atherosclerosis Signaling | 4.07E+00 | 6.77E-02 | NaN | CCL2,CXCL8,ICAM1,IL18,LPL,ORM1,ORM2,RELB,TNFRSF12A |
| 6 | Agranulocyte Adhesion and Diapedesis | 3.84E+00 | 5.24E-02 | NaN | ACTA2,CCL2,CLDN5,CXCL8,HRH1,ICAM1,IL18,IL1R1,ITGB7,MMP10,MMP17 |
| 7 | Airway Pathology in Chronic Obstructive Pulmonary Diseas | 3.69E+00 | 6.78E-02 | NaN | CCL2,CLCF1,CXCL8,IL11,IL18,LIF,ORM1,ORM2 |
| 8 | ABRA Signaling Pathway | 3.60E+00 | 7.61E-02 | 1.89 | ACTA2,CCN2,EGR1,FOS,JUNB,LIF,TAGLN |
| 9 | NOD1/2 Signaling Pathway | 3.58E+00 | 5.29E-02 | 1.897 | BIRC3,CCL2,CLCF1,CXCL8,FOS,IL11,IL18,LIF,TAB2,TNFAIP3 |
| 10 | Hepatic Cholestasis | 3.54E+00 | 5.24E-02 | NaN | ATP8B1,CLCF1,CXCL8,IL11,IL18,IL1R1,LIF,RELB,SLCO1A2,SREBF1 |
| | FXR/RXR Activation | 3.49E+00 | 6.35E-02 | NaN | AGT,IL18,LPL,ORM1,ORM2,SAA1,SAA2,SREBF1 |
| | Hepatic Fibrosis / Hepatic Stellate Cell Activation | 3.49E+00 | 5.15E-02 | NaN | ACTA2,AGT,CCL2,CCN2,COL28A1,CXCL8,ICAM1,IL1R1,RELB,SERPINE1 |
| | Toll-like Receptor Signaling | 3.18E+00 | 7.69E-02 | NaN | FOS,IL18,RELB,TAB2,TNFAIP3,TRAF1 |
| | Role Of Chondrocytes In Rheumatoid Arthritis Signaling Pat | 3.16E+00 | 5.67E-02 | 2.121 | CCL2,CXCL8,EGR1,FOS,IL18,IL1R1,MMP10,MMP17 |
| | Granulocyte Adhesion and Diapedesis | 2.95E+00 | 4.76E-02 | NaN | CCL2,CLDN5,CXCL8,HRH1,ICAM1,IL18,IL1R1,MMP10,MMP17 |
| | TWEAK Signaling | 2.82E+00 | 1.08E-01 | NaN | BIRC3,RELB,TNFRSF12A,TRAF1 |
| | Pulmonary Fibrosis Idiopathic Signaling Pathway | 2.76E+00 | 3.68E-02 | 2.111 | ACTA2,CCN2,COL28A1,EGR1,FOS,IL11,MMP10,MMP17,PMAIP1,RELB,SERPINE1,THBS1 |
| | Inhibition of Matrix Metalloproteases | 2.73E+00 | 1.03E-01 | 0 | ADAM12,MMP10,MMP17,TFPI2 |
| | CD40 Signaling | 2.67E+00 | 7.46E-02 | 0 | FOS,ICAM1,RELB,TNFAIP3,TRAF1 |
| | Wound Healing Signaling Pathway | 2.62E+00 | 3.97E-02 | 2.53 | ACTA2,CLCF1,COL28A1,CXCL8,FOS,IL11,IL18,IL1R1,LIF,MMP10 |
| | DHCR24 Signaling Pathway | 2.57E+00 | 5.11E-02 | -2.646 | AGT,ORM1,ORM2,RXRG,SAA1,SAA2,SREBF1 |
| | Erythropoietin Signaling Pathway | 2.54E+00 | 4.52E-02 | -1.134 | BIRC3,CLCF1,CXCL8,FOS,IL11,IL18,LIF,RELB |
| | Apelin Cardiac Fibroblast Signaling Pathway | 2.45E+00 | 1.30E-01 | NaN | CCN2,SERPINE1,SPHK1 |
| | Role of IL-17F in Allergic Inflammatory Airway Diseases | 2.43E+00 | 8.51E-02 | NaN | CCL2,CXCL8,IL11,RELB |
| | TREM1 Signaling | 2.41E+00 | 6.49E-02 | 2 | CCL2,CXCL8,ICAM1,IL18,RELB |
| | IL-17 Signaling | 2.39E+00 | 4.28E-02 | 2.121 | CCL2,CLCF1,CXCL8,FOS,IL11,IL18,LIF,TAB2 |
| | Macrophage Alternative Activation Signaling Pathway | 2.36E+00 | 4.23E-02 | 0.707 | DUSP1,FOS,IL18,LPL,RELB,RXRG,SREBF1,THBS1 |
| | Role of JAK family kinases in IL-6-type Cytokine Signaling | 2.36E+00 | 6.33E-02 | 2.236 | CLCF1,FOS,IL11,JUNB,LIF |
| | TNFR1 Signaling | 2.30E+00 | 7.84E-02 | NaN | BIRC3,FOS,RELB,TNFAIP3 |
| | Hepatic Fibrosis Signaling Pathway | 2.28E+00 | 3.07E-02 | 2.111 | ACTA2,AGT,CCL2,CCN2,CXCL8,FOS,ICAM1,IL18,IL1R1,ITGB7,MYLK3,RELB,SERPINE1 |
| | CDX Gastrointestinal Cancer Signaling Pathway | 2.19E+00 | 3.96E-02 | -1.89 | CLCF1,CXCL8,DLL1,FOS,IL11,IL18,LIF,RELB |
| | Activin Inhibin Signaling Pathway | 2.02E+00 | 3.69E-02 | 2.121 | CCN2,FBXO32,FOS,FOSL1,IL11,IL18,IL1R1,SERPINE1 |
| | Role Of Osteoclasts In Rheumatoid Arthritis Signaling Pathv | 2.01E+00 | 3.24E-02 | 0 | ADAM12,ADAM33,BIRC3,COL28A1,FOS,IL18,IL1R1,MMP10,MMP17,TAB2 |
| | Multiple Sclerosis Signaling Pathway | 1.96E+00 | 3.60E-02 | 0.707 | CLCF1,CXCL8,DUSP1,DUSP6,IL11,IL18,LIF,MASP1 |
| | Mineralocorticoid Biosynthesis | 1.96E+00 | 1.67E-01 | NaN | CYP21A2,GSTA1 |
| | Tumor Microenvironment Pathway | 1.95E+00 | 3.91E-02 | 1.633 | CCL2,CXCL8,FOS,ICAM1,MMP10,MMP17,RELB |
| | Neuroinflammation Signaling Pathway | 1.94E+00 | 3.15E-02 | 2.121 | BDNF,BIRC3,CCL2,CXCL8,FOS,GABRG1,ICAM1,IL18,IL1R1,RELB |
| | Glucocorticoid Biosynthesis | 1.89E+00 | 1.54E-01 | NaN | CYP21A2,GSTA1 |
| | IL-33 Signaling Pathway | 1.87E+00 | 3.76E-02 | 2.449 | CCL2,CXCL8,FOS,H2BC5,ICAM1,ICAM5,IL18 |
| | Myelination Signaling Pathway | 1.85E+00 | 3.06E-02 | 1.897 | BDNF,DLC1,EGR2,FOS,HES5,ITGB7,NRG1,RXRG,SCD,SREBF1 |
| | RAR Activation | 1.85E+00 | 2.80E-02 | -1.155 | CDKN2B,CLCF1,CXCL8,DKK1,DUSP1,FOS,IL11,IL18,LIF,PDE2A,RELB,RXRG |
| | IL-17A Signaling in Fibroblasts | 1.84E+00 | 7.89E-02 | NaN | CCL2,FOS,RELB |
| | WNK Renal Signaling Pathway | 1.83E+00 | 4.72E-02 | 0 | AGT,BIRC3,SCNN1B,SGK1,WNK4 |
| | Osteoarthritis Pathway | 1.81E+00 | 3.39E-02 | 1.633 | CNMD,CXCL8,DKK1,IL1R1,ITGB7,MMP10,RELB,SPHK1 |
| | ERK5 Signaling | 1.74E+00 | 5.41E-02 | 2 | FOS,FOSL1,LIF,SGK1 |
| | Role Of Osteoblasts In Rheumatoid Arthritis Signaling Pathv | 1.74E+00 | 3.28E-02 | 2.121 | CLCF1,CXCL8,DKK1,IL11,IL18,LIF,MMP10,MMP17 |
| | IL-10 Signaling | 1.73E+00 | 3.90E-02 | -1.633 | DUSP1,FOS,ICAM1,IL18,IL1R1,RELB |
| | April Mediated Signaling | 1.72E+00 | 7.14E-02 | NaN | FOS,RELB,TRAF1 |
| | Adenosine Nucleotides Degradation II | 1.71E+00 | 1.25E-01 | NaN | ADA2,XDH |
| | Role of Pattern Recognition Receptors in Recognition of Ba | 1.71E+00 | 3.85E-02 | NaN | CLCF1,CXCL8,IL11,IL18,LIF,RELB |
| | B Cell Activating Factor Signaling | 1.69E+00 | 6.98E-02 | NaN | FOS,RELB,TRAF1 |
| | Senescence Pathway | 1.68E+00 | 3.01E-02 | 0.333 | ATF3,CDKN2B,CXCL8,ETS1,GADD45A,PDK4,SAA1,SAA2,SERPINE1 |
| | Purine Nucleotides Degradation II (Aerobic) | 1.57E+00 | 1.05E-01 | NaN | ADA2,XDH |
| | ERK/MAPK Signaling | 1.56E+00 | 3.26E-02 | -0.816 | DUSP1,DUSP4,DUSP6,ETS1,FOS,ITGB7,MYCN |
| | IL-6 Signaling | 1.51E+00 | 3.88E-02 | 2 | CXCL8,FOS,IL18,IL1R1,RELB |
| | RANK Signaling in Osteoclasts | 1.46E+00 | 4.40E-02 | NaN | BIRC3,FOS,RELB,TAB2 |
| | Role of Osteoblasts, Osteoclasts and Chondrocytes in Rheu | 1.44E+00 | 3.07E-02 | NaN | BIRC3,DKK1,FOS,IL11,IL18,IL1R1,TAB2 |
| | D-myo-inositol (1,4,5,6)-Tetrakisphosphate Biosynthesis | 1.44E+00 | 3.33E-02 | 0 | DUSP1,DUSP4,DUSP6,EPHX2,PPP1R1B,PTPRN |
| | D-myo-inositol (3,4,5,6)-tetrakisphosphate Biosynthesis | 1.44E+00 | 3.33E-02 | 0 | DUSP1,DUSP4,DUSP6,EPHX2,PPP1R1B,PTPRN |
| | Role of Macrophages, Fibroblasts and Endothelial Cells in R | 1.43E+00 | 2.71E-02 | NaN | CCL2,CXCL8,DKK1,FOS,ICAM1,IL16,IL18,IL1R1,TRAF1 |
| | Role of PKR in Interferon Induction and Antiviral Response | 1.42E+00 | 3.68E-02 | 1 | ATF3,FOS,IL18,RELB,TAB2 |
| | Small Cell Lung Cancer Signaling | 1.38E+00 | 4.17E-02 | NaN | CDKN2B,RELB,RXRG,TRAF1 |
| | IL-1 Signaling | 1.38E+00 | 4.17E-02 | NaN | FOS,IL1R1,RELB,TAB2 |
| | Role of IL-17A in Arthritis | 1.38E+00 | 5.26E-02 | NaN | CCL2,CXCL8,RELB |
| | IL-12 Signaling and Production in Macrophages | 1.38E+00 | 2.97E-02 | 0.378 | FOS,IL18,IRF8,ORM1,ORM2,RELB,THBS1 |
| | NRF2-mediated Oxidative Stress Response | 1.37E+00 | 2.95E-02 | 0 | ACTA2,DNAJB13,FOS,FOSL1,GSTA1,JUNB,MAFF |
| | Macrophage Classical Activation Signaling Pathway | 1.36E+00 | 3.17E-02 | 1.633 | CLCF1,CXCL8,IL11,IL18,IRF8,LIF |
| | 3-phosphoinositide Degradation | 1.34E+00 | 3.14E-02 | 0 | DUSP1,DUSP4,DUSP6,EPHX2,PPP1R1B,PTPRN |

| | | | | |
|---|---|---|---|---|
| Leukocyte Extravasation Signaling | 1.32E+00 | 3.11E-02 | 0.447 | ACTA2,CLDN5,DLC1,ICAM1,MMP10,MMP17 |
| IL-17A Signaling in Gastric Cells | 1.32E+00 | 7.69E-02 | NaN | CXCL8,FOS |
| D-myo-inositol-5-phosphate Metabolism | 1.29E+00 | 3.06E-02 | 0 | DUSP1,DUSP4,DUSP6,EPHX2,PPP1R1B,PTPRN |
| Protein Kinase A Signaling | 1.27E+00 | 2.43E-02 | -2.646 | DUSP1,DUSP4,DUSP6,MYLK3,PDE2A,PLN,PPP1R1B,PTPRN,RELB,TNNI3 |
| Adrenomedullin signaling pathway | 1.27E+00 | 3.02E-02 | 0 | FOS,IL18,KCNQ3,MYLK3,RELB,RXRG |
| Glucocorticoid Receptor Signaling | 1.27E+00 | 2.23E-02 | NaN | AGT,CCL2,CXCL8,DUSP1,FOS,HP,ICAM1,IL1R1,PDK4,RXRG,SERPINE1,SGK1,SLPI |
| ID1 Signaling Pathway | 1.25E+00 | 2.99E-02 | 0.816 | ATF3,CCN2,CHRNA9,EGR1,ETS1,LIF |
| PPAR Signaling | 1.24E+00 | 3.74E-02 | NaN | FOS,IL18,IL1R1,RELB |
| Induction of Apoptosis by HIV1 | 1.24E+00 | 4.62E-02 | NaN | BIRC3,RELB,TRAF1 |
| Gustation Pathway | 1.23E+00 | 2.96E-02 | -0.816 | ENTPD2,GABRG1,KCNQ3,LPL,SCN1A,SCNN1B |
| Coronavirus Pathogenesis Pathway | 1.23E+00 | 2.94E-02 | 1.633 | AGT,CCL2,CXCL8,FOS,RELB,SERPINE1 |
| 3-phosphoinositide Biosynthesis | 1.21E+00 | 2.91E-02 | 0 | DUSP1,DUSP4,DUSP6,EPHX2,PPP1R1B,PTPRN |
| Pathogen Induced Cytokine Storm Signaling Pathway | 1.18E+00 | 2.43E-02 | 2.333 | CCL2,CLCF1,COL28A1,CXCL8,FOS,IL11,IL18,IL1R1,LIF |
| Bladder Cancer Signaling | 1.14E+00 | 3.45E-02 | NaN | CXCL8,MMP10,MMP17,THBS1 |
| Airway Inflammation in Asthma | 1.14E+00 | 6.06E-02 | NaN | CCL2,CXCL8 |
| 4-1BB Signaling in T Lymphocytes | 1.11E+00 | 5.88E-02 | NaN | RELB,TRAF1 |
| UDP-N-acetyl-D-glucosamine Biosynthesis II | 1.11E+00 | 1.67E-01 | NaN | GFPT2 |
| p38 MAPK Signaling | 1.10E+00 | 3.33E-02 | 0 | DUSP1,IL18,IL1R1,TAB2 |
| Coagulation System | 1.09E+00 | 5.71E-02 | NaN | SERPINE1,VWF |
| Renin-Angiotensin Signaling | 1.09E+00 | 3.31E-02 | NaN | AGT,CCL2,FOS,RELB |
| Th1 Pathway | 1.08E+00 | 3.28E-02 | 1 | DLL1,ICAM1,ICOSLG/LOC102723996,IL18 |
| Aldosterone Signaling in Epithelial Cells | 1.07E+00 | 2.91E-02 | NaN | DNAJB13,DUSP1,LRRC55,SCNN1B,SGK1 |
| VDR/RXR Activation | 1.05E+00 | 3.85E-02 | NaN | GADD45A,RXRG,SERPINB1 |
| Adenine and Adenosine Salvage III | 1.04E+00 | 1.43E-01 | NaN | ADA2 |
| Thyroid Cancer Signaling | 1.04E+00 | 3.80E-02 | NaN | BDNF,CXCL8,FOS |
| Notch Signaling | 1.03E+00 | 5.26E-02 | NaN | DLL1,HES5 |
| G-Protein Coupled Receptor Signaling | 1.01E+00 | 1.99E-02 | 0.535 | ADCYAP1R1,CMKLR1,DUSP1,DUSP4,DUSP6,FOS,GPR3,GRPR,HRH1,KCNQ3,MYLK3,PDE2A,RELB,RGS4 |
| Pyrimidine Ribonucleotides Interconversion | 1.01E+00 | 5.13E-02 | NaN | AK7,ENTPD2 |
| Superpathway of Inositol Phosphate Compounds | 1.00E+00 | 2.56E-02 | 0 | DUSP1,DUSP4,DUSP6,EPHX2,PPP1R1B,PTPRN |
| Sirtuin Signaling Pathway | 9.83E-01 | 2.39E-02 | -1.342 | ATG9B,CXCL8,DUSP6,GADD45A,MYCN,RELB,SREBF1 |
| TR/RXR Activation | 9.77E-01 | 3.57E-02 | NaN | HP,RXRG,SREBF1 |
| Pyrimidine Ribonucleotides De Novo Biosynthesis | 9.57E-01 | 4.76E-02 | NaN | AK7,ENTPD2 |
| Role of Hypercytokinemia/hyperchemokinemia in the Path | 9.55E-01 | 3.49E-02 | NaN | CCL2,CXCL8,IL18 |
| Purine Ribonucleosides Degradation to Ribose-1-phosphate | 9.38E-01 | 1.11E-01 | NaN | ADA2 |
| Salvage Pathways of Pyrimidine Deoxyribonucleotides | 9.38E-01 | 1.11E-01 | NaN | TYMP |
| Production of Nitric Oxide and Reactive Oxygen Species in | 9.28E-01 | 2.62E-02 | -1 | FOS,IRF8,ORM1,ORM2,RELB |
| MIF Regulation of Innate Immunity | 9.24E-01 | 4.55E-02 | NaN | FOS,RELB |
| Cardiac Hypertrophy Signaling (Enhanced) | 9.23E-01 | 2.03E-02 | 0.707 | AGT,CLCF1,CXCL8,IL11,IL18,IL1R1,ITGB7,LIF,PDE2A,PLN,RELB |
| MSP-RON Signaling In Cancer Cells Pathway | 9.13E-01 | 2.86E-02 | NaN | ACTA2,ETS1,FOS,RELB |
| Apelin Endothelial Signaling Pathway | 9.05E-01 | 2.84E-02 | NaN | CCL2,FOS,ICAM1,RELB |
| Ceramide Signaling | 9.01E-01 | 3.30E-02 | NaN | FOS,RELB,SPHK1 |
| Crosstalk between Dendritic Cells and Natural Killer Cells | 9.01E-01 | 3.30E-02 | NaN | ACTA2,IL18,RELB |
| iNOS Signaling | 8.77E-01 | 4.26E-02 | NaN | FOS,RELB |
| LPS/IL-1 Mediated Inhibition of RXR Function | 8.76E-01 | 2.35E-02 | 0 | ABCG1,GSTA1,IL18,IL1R1,SLCO1A2,SREBF1 |
| Death Receptor Signaling | 8.51E-01 | 3.12E-02 | NaN | ACTA2,BIRC3,RELB |
| Dilated Cardiomyopathy Signaling Pathway | 8.35E-01 | 2.67E-02 | NaN | ACTA2,PDE2A,PLN,TNNI3 |
| p53 Signaling | 8.32E-01 | 3.06E-02 | NaN | GADD45A,PMAIP1,THBS1 |
| Role of Tissue Factor in Cancer | 8.24E-01 | 2.42E-02 | 2.236 | CCN1,CCN2,CXCL8,EGR1,FOS |
| Glycogen Degradation II | 8.22E-01 | 8.33E-02 | NaN | TYMP |
| Neuropathic Pain Signaling In Dorsal Horn Neurons | 8.04E-01 | 2.97E-02 | NaN | BDNF,FOS,KCNQ3 |
| Oleate Biosynthesis II (Animals) | 7.90E-01 | 7.69E-02 | NaN | SCD |
| Guanosine Nucleotides Degradation III | 7.90E-01 | 7.69E-02 | NaN | XDH |
| Role of Cytokines in Mediating Communication between In | 7.81E-01 | 3.70E-02 | NaN | CXCL8,IL18 |
| Aryl Hydrocarbon Receptor Signaling | 7.72E-01 | 2.52E-02 | NaN | FOS,GSTA1,RELB,RXRG |
| IGF-1 Signaling | 7.70E-01 | 2.86E-02 | NaN | CCN1,CCN2,FOS |
| Lymphotoxin β Receptor Signaling | 7.69E-01 | 3.64E-02 | NaN | RELB,TRAF1 |
| Role of IL-17A in Psoriasis | 7.61E-01 | 7.14E-02 | NaN | CXCL8 |
| Glycogen Degradation III | 7.61E-01 | 7.14E-02 | NaN | TYMP |
| Urate Biosynthesis/Inosine 5'-phosphate Degradation | 7.61E-01 | 7.14E-02 | NaN | XDH |
| CSDE1 Signaling Pathway | 7.57E-01 | 3.57E-02 | NaN | CCL2,FOS |
| HOTAIR Regulatory Pathway | 7.45E-01 | 2.45E-02 | NaN | ICAM1,MMP10,MMP17,RELB |
| CD27 Signaling in Lymphocytes | 7.45E-01 | 3.51E-02 | NaN | FOS,RELB |
| MSP-RON Signaling Pathway | 7.33E-01 | 3.45E-02 | NaN | ACTA2,CCL2 |
| EIF2 Signaling | 7.11E-01 | 2.20E-02 | -2 | ACTA2,ATF3,MYCN,NKX6-2,SREBF1 |
| HER-2 Signaling in Breast Cancer | 7.11E-01 | 2.20E-02 | 1.342 | ETS1,FOS,ITGB7,NRG1,RELB |
| GADD45 Signaling | 7.11E-01 | 3.33E-02 | NaN | GADD45A,RELB |
| CDK5 Signaling | 6.90E-01 | 2.61E-02 | NaN | BDNF,EGR1,PPP1R1B |
| Th1 and Th2 Activation Pathway | 6.90E-01 | 2.33E-02 | NaN | DLL1,ICAM1,ICOSLG/LOC102723996,IL18 |
| Role of JAK2 in Hormone-like Cytokine Signaling | 6.89E-01 | 3.23E-02 | NaN | BIRC3,FOS |
| cAMP-mediated signaling | 6.66E-01 | 2.12E-02 | -2.236 | DUSP1,DUSP4,DUSP6,PDE2A,RGS4 |
| Differential Regulation of Cytokine Production in Macropha | 6.63E-01 | 5.56E-02 | NaN | CCL2 |
| PXR/RXR Activation | 6.59E-01 | 3.08E-02 | NaN | GSTA1,SCD |

| | | | | |
|---|---|---|---|---|
| Tight Junction Signaling | 6.49E-01 | 2.23E-02 | NaN | ACTA2,CLDN5,FOS,RELB |
| Glutamate Receptor Signaling | 6.49E-01 | 3.03E-02 | NaN | GRIK1,SLC17A8 |
| Cell Cycle: G1/S Checkpoint Regulation | 6.30E-01 | 2.94E-02 | NaN | CDKN2B,NRG1 |
| RHOA Signaling | 6.28E-01 | 2.42E-02 | NaN | ACTA2,DLC1,MYLK3 |
| Androgen Biosynthesis | 6.23E-01 | 5.00E-02 | NaN | GSTA1 |
| Inflammasome pathway | 6.23E-01 | 5.00E-02 | NaN | IL18 |
| Agrin Interactions at Neuromuscular Junction | 6.21E-01 | 2.90E-02 | NaN | ACTA2,NRG1 |
| Regulation Of The Epithelial Mesenchymal Transition By Gr | 5.82E-01 | 2.08E-02 | NaN | EGR1,ETS1,FOS,RELB |
| HGF Signaling | 5.77E-01 | 2.27E-02 | NaN | ETS1,FOS,ITGB7 |
| Differential Regulation of Cytokine Production in Intestinal | 5.70E-01 | 4.35E-02 | NaN | CCL2 |
| Pyrimidine Deoxyribonucleotides De Novo Biosynthesis I | 5.70E-01 | 4.35E-02 | NaN | AK7 |
| Caveolar-mediated Endocytosis Signaling | 5.70E-01 | 2.67E-02 | NaN | ACTA2,ITGB7 |
| Tumoricidal Function of Hepatic Natural Killer Cells | 5.55E-01 | 4.17E-02 | NaN | ICAM1 |
| Th2 Pathway | 5.48E-01 | 2.19E-02 | NaN | DLL1,ICAM1,ICOSLG/LOC102723996 |
| Neurotrophin/TRK Signaling | 5.46E-01 | 2.56E-02 | NaN | BDNF,FOS |
| White Adipose Tissue Browning Pathway | 5.43E-01 | 2.17E-02 | NaN | BDNF,RXRG,VGF |
| ILK Signaling | 5.39E-01 | 1.99E-02 | NaN | ACTA2,FOS,ITGB7,RELB |
| Renal Cell Carcinoma Signaling | 5.38E-01 | 2.53E-02 | NaN | ETS1,FOS |
| Role of MAPK Signaling in Inhibiting the Pathogenesis of Inf | 5.38E-01 | 2.53E-02 | NaN | CCL2,CXCL8 |
| Adipogenesis pathway | 5.37E-01 | 2.16E-02 | NaN | EGR2,LPL,SREBF1 |
| Dopamine Receptor Signaling | 5.31E-01 | 2.50E-02 | NaN | CALY,PPP1R1B |
| Role of JAK1, JAK2 and TYK2 in Interferon Signaling | 5.25E-01 | 3.85E-02 | NaN | RELB |
| Chemokine Signaling | 5.24E-01 | 2.47E-02 | NaN | CCL2,FOS |
| JAK/STAT Signaling | 5.16E-01 | 2.44E-02 | NaN | FOS,RELB |
| Apelin Liver Signaling Pathway | 5.12E-01 | 3.70E-02 | NaN | AGT |
| Estrogen-Dependent Breast Cancer Signaling | 5.09E-01 | 2.41E-02 | NaN | FOS,RELB |
| Clathrin-mediated Endocytosis Signaling | 5.08E-01 | 1.92E-02 | NaN | ACTA2,ITGB7,ORM1,ORM2 |
| PEDF Signaling | 5.02E-01 | 2.38E-02 | NaN | BDNF,RELB |
| Glutathione Redox Reactions I | 4.99E-01 | 3.57E-02 | NaN | GSTA1 |
| LPS-stimulated MAPK Signaling | 4.96E-01 | 2.35E-02 | NaN | FOS,RELB |
| PDGF Signaling | 4.82E-01 | 2.30E-02 | NaN | FOS,SPHK1 |
| Xenobiotic Metabolism AHR Signaling Pathway | 4.82E-01 | 2.30E-02 | NaN | GSTA1,RELB |
| Cellular Effects of Sildenafil (Viagra) | 4.80E-01 | 2.00E-02 | NaN | ACTA2,KCNQ3,PDE2A |
| Unfolded protein response | 4.63E-01 | 2.22E-02 | NaN | DNAJB13,SREBF1 |
| Calcium Signaling | 4.60E-01 | 1.82E-02 | NaN | ACTA2,CHRNA9,GRIK1,TNNI3 |
| Relaxin Signaling | 4.57E-01 | 1.94E-02 | NaN | FOS,PDE2A,RELB |
| S100 Family Signaling Pathway | 4.53E-01 | 1.55E-02 | 1.732 | ADCYAP1R1,BDNF,CMKLR1,CXCL8,DLC1,FOS,GPR3,GRPR,HRH1,IL18,MMP10,MMP17 |
| Pyroptosis Signaling Pathway | 4.45E-01 | 2.15E-02 | NaN | IL18,IL1R1 |
| ERBB Signaling | 4.45E-01 | 2.15E-02 | NaN | FOS,NRG1 |
| Prolactin Signaling | 4.34E-01 | 2.11E-02 | NaN | FOS,LRRC55 |
| Inhibition of Angiogenesis by TSP1 | 4.31E-01 | 2.94E-02 | NaN | THBS1 |
| TGF-β Signaling | 4.28E-01 | 2.08E-02 | NaN | FOS,SERPINE1 |
| Salvage Pathways of Pyrimidine Ribonucleotides | 4.22E-01 | 2.06E-02 | NaN | AK7,SGK1 |
| IL-9 Signaling | 4.21E-01 | 2.86E-02 | NaN | RELB |
| Axonal Guidance Signaling | 4.14E-01 | 1.57E-02 | NaN | ADAM12,ADAM33,BDNF,EFNA1,ITGB7,MMP10,MMP17,SEMA4A |
| MIF-mediated Glucocorticoid Regulation | 4.12E-01 | 2.78E-02 | NaN | RELB |
| Glutathione-mediated Detoxification | 4.02E-01 | 2.70E-02 | NaN | GSTA1 |
| Complement System | 4.02E-01 | 2.70E-02 | NaN | MASP1 |
| Gαq Signaling | 3.93E-01 | 1.76E-02 | NaN | HRH1,RELB,RGS4 |
| Sumoylation Pathway | 3.91E-01 | 1.94E-02 | NaN | ETS1,FOS |
| Molecular Mechanisms of Cancer | 3.90E-01 | 1.56E-02 | NaN | BIRC3,CDKN2B,FOS,ITGB7,PMAIP1,RELB,TAB2 |
| Apoptosis Signaling | 3.86E-01 | 1.92E-02 | NaN | BIRC3,RELB |
| Synaptogenesis Signaling Pathway | 3.74E-01 | 1.59E-02 | 0.447 | BDNF,EFNA1,STXBP6,SYT6,THBS1 |
| Paxillin Signaling | 3.71E-01 | 1.87E-02 | NaN | ACTA2,ITGB7 |
| Mechanisms of Viral Exit from Host Cells | 3.69E-01 | 2.44E-02 | NaN | ACTA2 |
| Oncostatin M Signaling | 3.53E-01 | 2.33E-02 | NaN | CHI3L1 |
| Serotonin Receptor Signaling | 3.48E-01 | 1.49E-02 | 1.134 | BDNF,CXCL8,KCNQ3,MYLK3,PLN,TNNI3,VWF |
| Regulation of Actin-based Motility by Rho | 3.35E-01 | 1.74E-02 | NaN | ACTA2,ITGB7 |
| Role of RIG1-like Receptors in Antiviral Innate Immunity | 3.32E-01 | 2.17E-02 | NaN | RELB |
| Role of OCT4 in Mammalian Embryonic Stem Cell Pluripote | 3.32E-01 | 2.17E-02 | NaN | FBXO15 |
| tRNA Splicing | 3.32E-01 | 2.17E-02 | NaN | PDE2A |
| Apelin Pancreas Signaling Pathway | 3.32E-01 | 2.17E-02 | NaN | RELB |
| IL-23 Signaling Pathway | 3.32E-01 | 2.17E-02 | NaN | RELB |
| IL-13 Signaling Pathway | 3.31E-01 | 1.72E-02 | NaN | CHI3L1,DUSP1 |
| Amyotrophic Lateral Sclerosis Signaling | 3.31E-01 | 1.72E-02 | NaN | BIRC3,GRIK1 |
| Neuregulin Signaling | 3.27E-01 | 1.71E-02 | NaN | ITGB7,NRG1 |
| Ephrin A Signaling | 3.25E-01 | 2.13E-02 | NaN | EFNA1 |
| Retinol Biosynthesis | 3.25E-01 | 2.13E-02 | NaN | LPL |
| Virus Entry via Endocytic Pathways | 3.23E-01 | 1.69E-02 | NaN | ACTA2,ITGB7 |
| Estrogen Receptor Signaling | 3.21E-01 | 1.47E-02 | 0.447 | AGT,FBXO32,FOS,MMP10,MMP17,RELB |
| GNRH Signaling | 3.20E-01 | 1.57E-02 | NaN | EGR1,FOS,RELB |
| Cholecystokinin/Gastrin-mediated Signaling | 3.19E-01 | 1.68E-02 | NaN | FOS,IL18 |

| | | | | | |
|---|---|---|---|---|---|
| MSP-RON Signaling In Macrophages Pathway | 3.19E-01 | 1.68E-02 | NaN | | FOS,RELB |
| PFKFB4 Signaling Pathway | 3.19E-01 | 2.08E-02 | NaN | | XDH |
| Nitric Oxide Signaling in the Cardiovascular System | 3.15E-01 | 1.67E-02 | NaN | | PDE2A,PLN |
| Signaling by Rho Family GTPases | 3.11E-01 | 1.50E-02 | NaN | | ACTA2,FOS,ITGB7,RELB |
| Neuroprotective Role of THOP1 in Alzheimer's Disease | 3.11E-01 | 1.65E-02 | NaN | | AGT,MASP1 |
| PPARα/RXRα Activation | 3.08E-01 | 1.54E-02 | NaN | | IL1R1,LPL,RELB |
| Regulation of the Epithelial-Mesenchymal Transition Pathw | 3.08E-01 | 1.54E-02 | NaN | | EGR1,ETS1,RELB |
| MYC Mediated Apoptosis Signaling | 3.06E-01 | 2.00E-02 | NaN | | PMAIP1 |
| Cell Cycle: G2/M DNA Damage Checkpoint Regulation | 3.06E-01 | 2.00E-02 | NaN | | GADD45A |
| FAT10 Cancer Signaling Pathway | 3.06E-01 | 2.00E-02 | NaN | | RELB |
| Colorectal Cancer Metastasis Signaling | 3.01E-01 | 1.48E-02 | NaN | | FOS,MMP10,MMP17,RELB |
| Chondroitin Sulfate Biosynthesis (Late Stages) | 3.00E-01 | 1.96E-02 | NaN | | CHST9 |
| UVC-Induced MAPK Signaling | 3.00E-01 | 1.96E-02 | NaN | | FOS |
| Gap Junction Signaling | 2.99E-01 | 1.52E-02 | NaN | | ACTA2,GJA5,GRIK1 |
| Pulmonary Healing Signaling Pathway | 2.96E-01 | 1.51E-02 | NaN | | MMP10,MMP17,THBS1 |
| UVB-Induced MAPK Signaling | 2.94E-01 | 1.92E-02 | NaN | | FOS |
| PI3K/AKT Signaling | 2.93E-01 | 1.50E-02 | NaN | | IL1R1,ITGB7,RELB |
| Pancreatic Adenocarcinoma Signaling | 2.93E-01 | 1.59E-02 | NaN | | CDKN2B,RELB |
| Human Embryonic Stem Cell Pluripotency | 2.90E-01 | 1.49E-02 | NaN | | BDNF,LIF,SPHK1 |
| Sertoli Cell-Sertoli Cell Junction Signaling | 2.76E-01 | 1.46E-02 | NaN | | ACTA2,CLDN5,ITGB7 |
| Oxytocin Signaling Pathway | 2.75E-01 | 1.42E-02 | | 2 | CXCL8,FOS,LPL,RELB |
| EGF Signaling | 2.73E-01 | 1.79E-02 | NaN | | FOS |
| GABA Receptor Signaling | 2.72E-01 | 1.52E-02 | NaN | | GABRG1,KCNQ3 |
| Ferroptosis Signaling Pathway | 2.72E-01 | 1.52E-02 | NaN | | ANGPTL4,H2BC5 |
| HIF1α Signaling | 2.71E-01 | 1.44E-02 | NaN | | MMP10,MMP17,SERPINE1 |
| P2Y Purigenic Receptor Signaling Pathway | 2.69E-01 | 1.50E-02 | NaN | | FOS,RELB |
| Triacylglycerol Degradation | 2.68E-01 | 1.75E-02 | NaN | | LPL |
| IL-8 Signaling | 2.66E-01 | 1.43E-02 | NaN | | CXCL8,FOS,ICAM1 |
| Integrin Signaling | 2.61E-01 | 1.42E-02 | NaN | | ACTA2,ITGB7,MYLK3 |
| SNARE Signaling Pathway | 2.59E-01 | 1.47E-02 | NaN | | STXBP6,SYT6 |
| Polyamine Regulation in Colon Cancer | 2.58E-01 | 1.69E-02 | NaN | | FOS |
| Chondroitin Sulfate Biosynthesis | 2.58E-01 | 1.69E-02 | NaN | | CHST9 |
| RAC Signaling | 2.56E-01 | 1.46E-02 | NaN | | ITGB7,RELB |
| Retinoic acid Mediated Apoptosis Signaling | 2.53E-01 | 1.67E-02 | NaN | | RXRG |
| PCP (Planar Cell Polarity) Pathway | 2.53E-01 | 1.67E-02 | NaN | | JUNB |
| SPINK1 Pancreatic Cancer Pathway | 2.53E-01 | 1.67E-02 | NaN | | CPA4 |
| Insulin Receptor Signaling | 2.47E-01 | 1.43E-02 | NaN | | SCNN1B,SGK1 |
| IL-2 Signaling | 2.44E-01 | 1.61E-02 | NaN | | FOS |
| RHOGDI Signaling | 2.41E-01 | 1.36E-02 | NaN | | ACTA2,DLC1,ITGB7 |
| Thrombopoietin Signaling | 2.39E-01 | 1.59E-02 | NaN | | FOS |
| Activation of IRF by Cytosolic Pattern Recognition Receptor | 2.31E-01 | 1.54E-02 | NaN | | RELB |
| ERB2-ERBB3 Signaling | 2.31E-01 | 1.54E-02 | NaN | | NRG1 |
| Pyridoxal 5'-phosphate Salvage Pathway | 2.31E-01 | 1.54E-02 | NaN | | SGK1 |
| Role of PI3K/AKT Signaling in the Pathogenesis of Influenza | 2.27E-01 | 1.52E-02 | NaN | | RELB |
| WNT/Ca+ pathway | 2.27E-01 | 1.52E-02 | NaN | | RELB |
| IL-17A Signaling in Airway Cells | 2.23E-01 | 1.49E-02 | NaN | | RELB |
| ERBB4 Signaling | 2.19E-01 | 1.47E-02 | NaN | | NRG1 |
| Remodeling of Epithelial Adherens Junctions | 2.19E-01 | 1.47E-02 | NaN | | ACTA2 |
| Phospholipases | 2.15E-01 | 1.45E-02 | NaN | | LPL |
| SPINK1 General Cancer Pathway | 2.15E-01 | 1.45E-02 | NaN | | MT1M |
| GM-CSF Signaling | 2.11E-01 | 1.43E-02 | NaN | | ETS1 |
| Growth Hormone Signaling | 2.08E-01 | 1.41E-02 | NaN | | FOS |
| NAD Signaling Pathway | 0.00E+00 | 1.32E-02 | NaN | | H2BC5,SREBF1 |
| Neurovascular Coupling Signaling Pathway | 0.00E+00 | 1.29E-02 | NaN | | ENTPD2,GABRG1,LRRC55 |
| Ribonucleotide Reductase Signaling Pathway | 0.00E+00 | 1.18E-02 | NaN | | FOS,THBS1 |
| Natural Killer Cell Signaling | 0.00E+00 | 1.01E-02 | NaN | | IL18,RELB |
| Neutrophil Extracellular Trap Signaling Pathway | 0.00E+00 | 7.52E-03 | NaN | | COL28A1,CXCL8,SERPINB1 |
| Circadian Rhythm Signaling | 0.00E+00 | 1.12E-02 | NaN | | ADCYAP1R1,BDNF,GRPR |
| Actin Cytoskeleton Signaling | 0.00E+00 | 1.23E-02 | NaN | | ACTA2,ITGB7,MYLK3 |
| Huntington's Disease Signaling | 0.00E+00 | 7.07E-03 | NaN | | BDNF,SGK1 |
| Chaperone Mediated Autophagy Signaling Pathway | 0.00E+00 | 4.67E-03 | NaN | | IL18,MMP10,MMP17 |
| Mitochondrial Dysfunction | 0.00E+00 | 2.91E-03 | NaN | | GSTA1 |
| Role of BRCA1 in DNA Damage Response | 0.00E+00 | 1.25E-02 | NaN | | GADD45A |
| 14-3-3-mediated Signaling | 0.00E+00 | 7.87E-03 | NaN | | FOS |
| Fcγ Receptor-mediated Phagocytosis in Macrophages and I | 0.00E+00 | 1.06E-02 | NaN | | ACTA2 |
| Role of NFAT in Regulation of the Immune Response | 0.00E+00 | 1.93E-03 | NaN | | FOS,RELB |
| NF-κB Activation by Viruses | 0.00E+00 | 1.28E-02 | NaN | | RELB |
| CCR5 Signaling in Macrophages | 0.00E+00 | 2.00E-03 | NaN | | FOS |
| IL-3 Signaling | 0.00E+00 | 1.27E-02 | NaN | | FOS |
| fMLP Signaling in Neutrophils | 0.00E+00 | 7.63E-03 | NaN | | RELB |
| CXCR4 Signaling | 0.00E+00 | 1.19E-02 | NaN | | EGR1,FOS |
| CTLA4 Signaling in Cytotoxic T Lymphocytes | 0.00E+00 | 1.64E-03 | NaN | | FOS |

| | | | | |
|---|---|---|---|---|
| IL-15 Production | 0.00E+00 | 8.13E-03 | NaN | RELB |
| T Helper Cell Differentiation | 0.00E+00 | 4.25E-03 | NaN | ICOSLG/LOC102723996,IL18 |
| CD28 Signaling in T Helper Cells | 0.00E+00 | 3.87E-03 | NaN | FOS,RELB |
| IL-15 Signaling | 0.00E+00 | 3.79E-03 | NaN | CXCL8,RELB |
| Dendritic Cell Maturation | 0.00E+00 | 6.71E-03 | 1 | ICAM1,IL18,IRF8,RELB |
| Reelin Signaling in Neurons | 0.00E+00 | 7.25E-03 | NaN | PDK4 |
| Angiopoietin Signaling | 0.00E+00 | 1.32E-02 | NaN | RELB |
| Endothelin-1 Signaling | 0.00E+00 | 5.15E-03 | NaN | FOS |
| Factors Promoting Cardiogenesis in Vertebrates | 0.00E+00 | 1.31E-02 | NaN | DKK1,GJA5 |
| Thrombin Signaling | 0.00E+00 | 4.44E-03 | NaN | RELB |
| ICOS-ICOSL Signaling in T Helper Cells | 0.00E+00 | 3.94E-03 | NaN | ICOSLG/LOC102723996,RELB |
| Corticotropin Releasing Hormone Signaling | 0.00E+00 | 1.32E-02 | NaN | BDNF,FOS |
| DNA Methylation and Transcriptional Repression Signaling | 0.00E+00 | 1.02E-02 | NaN | GADD45A |
| ATM Signaling | 0.00E+00 | 1.00E-02 | NaN | GADD45A |
| Androgen Signaling | 0.00E+00 | 5.92E-03 | NaN | RELB |
| Germ Cell-Sertoli Cell Junction Signaling | 0.00E+00 | 5.88E-03 | NaN | ACTA2 |
| Role of NANOG in Mammalian Embryonic Stem Cell Pluripc | 0.00E+00 | 8.06E-03 | NaN | LIF |
| CREB Signaling in Neurons | 0.00E+00 | 9.88E-03 | 0.816 | ADCYAP1R1,CMKLR1,GPR3,GRIK1,GRPR,HRH1 |
| Prostate Cancer Signaling | 0.00E+00 | 8.77E-03 | NaN | RELB |
| Type I Diabetes Mellitus Signaling | 0.00E+00 | 5.89E-03 | NaN | IL1R1,PTPRN,RELB |
| Glioma Signaling | 0.00E+00 | 8.00E-03 | NaN | CDKN2B |
| Acute Myeloid Leukemia Signaling | 0.00E+00 | 1.10E-02 | NaN | RELB |
| Graft-versus-Host Disease Signaling | 0.00E+00 | 2.24E-03 | NaN | IL18 |
| Type II Diabetes Mellitus Signaling | 0.00E+00 | 6.54E-03 | NaN | RELB |
| Chronic Myeloid Leukemia Signaling | 0.00E+00 | 1.08E-02 | NaN | FOS,IRF8,RELB |
| Non-Small Cell Lung Cancer Signaling | 0.00E+00 | 1.06E-02 | NaN | RXRG |
| Gα12/13 Signaling | 0.00E+00 | 7.52E-03 | NaN | RELB |
| p70S6K Signaling | 0.00E+00 | 1.72E-03 | NaN | AGT |
| Communication between Innate and Adaptive Immune Cel | 0.00E+00 | 2.14E-03 | NaN | CXCL8,IL18 |
| Sphingosine-1-phosphate Signaling | 0.00E+00 | 8.33E-03 | NaN | SPHK1 |
| Systemic Lupus Erythematosus Signaling | 0.00E+00 | 1.87E-03 | NaN | FOS,IL18 |
| CDC42 Signaling | 0.00E+00 | 3.47E-03 | NaN | FOS,ITGB7 |
| FAK Signaling | 0.00E+00 | 9.61E-03 | 1.897 | ADCYAP1R1,CMKLR1,EFNA1,ETS1,FOS,GPR3,GRPR,HRH1,IL1R1,ITGB7 |
| AMPK Signaling | 0.00E+00 | 8.26E-03 | NaN | AK7,CHRNA9 |
| PAK Signaling | 0.00E+00 | 8.55E-03 | NaN | ITGB7 |
| Hereditary Breast Cancer Signaling | 0.00E+00 | 7.04E-03 | NaN | GADD45A |
| Phospholipase C Signaling | 0.00E+00 | 1.78E-03 | NaN | ITGB7,RELB |
| Altered T Cell and B Cell Signaling in Rheumatoid Arthritis | 0.00E+00 | 2.13E-03 | NaN | IL18,RELB |
| Regulation of eIF4 and p70S6K Signaling | 0.00E+00 | 5.52E-03 | NaN | ITGB7 |
| Role of NFAT in Cardiac Hypertrophy | 0.00E+00 | 8.93E-03 | NaN | IL11,LIF |
| Breast Cancer Regulation by Stathmin1 | 0.00E+00 | 8.42E-03 | 1.342 | ADCYAP1R1,CMKLR1,GPR3,GRPR,HRH1 |
| Regulation of IL-2 Expression in Activated and Anergic T Lyr | 0.00E+00 | 4.33E-03 | NaN | FOS,RELB |
| PKCθ Signaling in T Lymphocytes | 0.00E+00 | 3.58E-03 | NaN | FOS,RELB |
| Role of MAPK Signaling in the Pathogenesis of Influenza | 0.00E+00 | 1.19E-02 | NaN | CCL2 |
| OX40 Signaling Pathway | 0.00E+00 | 2.09E-03 | NaN | RELB |
| PI3K Signaling in B Lymphocytes | 0.00E+00 | 5.08E-03 | NaN | ATF3,FOS,RELB |
| Cyclins and Cell Cycle Regulation | 0.00E+00 | 1.18E-02 | NaN | CDKN2B |
| Actin Nucleation by ARP-WASP Complex | 0.00E+00 | 1.08E-02 | NaN | ITGB7 |
| Dopamine-DARPP32 Feedback in cAMP Signaling | 0.00E+00 | 1.08E-02 | NaN | CALY,PPP1R1B |
| NGF Signaling | 0.00E+00 | 8.33E-03 | NaN | RELB |
| Telomerase Signaling | 0.00E+00 | 9.26E-03 | NaN | ETS1 |
| Mouse Embryonic Stem Cell Pluripotency | 0.00E+00 | 9.62E-03 | NaN | LIF |
| Hematopoiesis from Pluripotent Stem Cells | 0.00E+00 | 6.74E-03 | NaN | CXCL8,IL11,LIF |
| Transcriptional Regulatory Network in Embryonic Stem Cell | 0.00E+00 | 6.10E-03 | NaN | LIF |
| eNOS Signaling | 0.00E+00 | 1.28E-02 | NaN | AQP5,CHRNA9 |
| VEGF Family Ligand-Receptor Interactions | 0.00E+00 | 1.19E-02 | NaN | FOS |
| GDNF Family Ligand-Receptor Interactions | 0.00E+00 | 1.32E-02 | NaN | FOS |
| Antioxidant Action of Vitamin C | 0.00E+00 | 8.77E-03 | NaN | RELB |
| Epithelial Adherens Junction Signaling | 0.00E+00 | 6.37E-03 | NaN | DLL1 |
| Gαi Signaling | 0.00E+00 | 7.14E-03 | NaN | RGS4 |
| Regulation of Cellular Mechanics by Calpain Protease | 0.00E+00 | 1.11E-02 | NaN | ITGB7 |
| Sperm Motility | 0.00E+00 | 3.89E-03 | NaN | PDE2A |
| TEC Kinase Signaling | 0.00E+00 | 6.92E-03 | NaN | ACTA2,FOS,ITGB7,RELB |
| UVA-Induced MAPK Signaling | 0.00E+00 | 1.02E-02 | NaN | FOS |
| STAT3 Pathway | 0.00E+00 | 7.41E-03 | NaN | IL1R1 |
| SAPK/JNK Signaling | 0.00E+00 | 3.98E-03 | NaN | DUSP4,GADD45A |
| PTEN Signaling | 0.00E+00 | 1.32E-02 | NaN | ITGB7,RELB |
| Cardiac β-adrenergic Signaling | 0.00E+00 | 1.11E-02 | NaN | PDE2A,PLN |
| Protein Ubiquitination Pathway | 0.00E+00 | 1.10E-02 | NaN | BIRC3,DNAJB13,USP43 |
| Xenobiotic Metabolism Signaling | 0.00E+00 | 6.85E-03 | NaN | GSTA1,RELB |
| IL-4 Signaling | 0.00E+00 | 1.73E-03 | NaN | COL28A1 |
| B Cell Receptor Signaling | 0.00E+00 | 4.72E-03 | NaN | EGR1,ETS1,RELB |

| | | | | |
|---|---|---|---|---|
| WNT/β-catenin Signaling | 0.00E+00 | 5.75E-03 | NaN | DKK1 |
| NF-κB Signaling | 0.00E+00 | 8.76E-03 | 0.447 | IL18,IL1R1,RELB,TAB2,TNFAIP3 |
| VEGF Signaling | 0.00E+00 | 1.01E-02 | NaN | ACTA2 |
| T Cell Receptor Signaling | 0.00E+00 | 8.12E-03 | 1 | DUSP6,FOS,ICAM1,ICOSLG/LOC102723996,RELB |
| BMP signaling pathway | 0.00E+00 | 1.10E-02 | NaN | RELB |
| GPCR-Mediated Integration of Enteroendocrine Signaling E | 0.00E+00 | 1.33E-02 | NaN | GRPR |
| Phagosome Formation | 0.00E+00 | 1.15E-02 | 0.707 | ADCYAP1R1,CMKLR1,GPR3,GRPR,HRH1,ITGB7,MYLK3,SPHK1 |
| Autophagy | 0.00E+00 | 9.26E-03 | NaN | ATG9B,FOS |
| Macropinocytosis Signaling | 0.00E+00 | 1.32E-02 | NaN | ITGB7 |
| GP6 Signaling Pathway | 0.00E+00 | 7.87E-03 | NaN | COL28A1 |
| Opioid Signaling Pathway | 0.00E+00 | 1.07E-02 | NaN | EGR4,FOS,RGS4 |
| Iron homeostasis signaling pathway | 0.00E+00 | 7.25E-03 | NaN | HP |
| Th17 Activation Pathway | 0.00E+00 | 4.13E-03 | NaN | IL1R1,RELB |
| Endocannabinoid Cancer Inhibition Pathway | 0.00E+00 | 6.80E-03 | NaN | ATF3 |
| Apelin Adipocyte Signaling Pathway | 0.00E+00 | 1.10E-02 | NaN | GSTA1 |
| Apelin Cardiomyocyte Signaling Pathway | 0.00E+00 | 1.01E-02 | NaN | PLN |
| BAG2 Signaling Pathway | 0.00E+00 | 1.19E-02 | NaN | RELB |
| T Cell Exhaustion Signaling Pathway | 0.00E+00 | 1.76E-03 | NaN | FOS |
| Systemic Lupus Erythematosus In T Cell Signaling Pathway | 0.00E+00 | 4.67E-03 | NaN | FOS,GADD45A,ICOSLG/LOC102723996 |
| Systemic Lupus Erythematosus In B Cell Signaling Pathway | 0.00E+00 | 1.10E-02 | 1.89 | CLCF1,CXCL8,FOS,IL11,IL18,LIF,RELB,TRAF1 |
| BEX2 Signaling Pathway | 0.00E+00 | 1.22E-02 | NaN | RELB |
| Necroptosis Signaling Pathway | 0.00E+00 | 1.28E-02 | NaN | BIRC3,TAB2 |
| Xenobiotic Metabolism General Signaling Pathway | 0.00E+00 | 6.99E-03 | NaN | GSTA1 |
| Xenobiotic Metabolism CAR Signaling Pathway | 0.00E+00 | 5.21E-03 | NaN | GSTA1 |
| Xenobiotic Metabolism PXR Signaling Pathway | 0.00E+00 | 5.18E-03 | NaN | GSTA1 |
| Insulin Secretion Signaling Pathway | 0.00E+00 | 3.68E-03 | NaN | SCNN1B |
| Semaphorin Neuronal Repulsive Signaling Pathway | 0.00E+00 | 6.67E-03 | NaN | ITGB7 |
| Regulation Of The Epithelial Mesenchymal Transition In De | 0.00E+00 | 1.15E-02 | NaN | RELB |
| Ephrin Receptor Signaling | 0.00E+00 | 9.90E-03 | NaN | EFNA1,ITGB7 |

IPA Microglia FB1 FC2 adj0.05 1734

| | Ingenuity Canonical Pathways | -log(p-value) | Ratio | z-score | Molecules |
|---|---|---|---|---|---|
| 1 | CREB Signaling in Neurons | 8.29E+00 | 7.74E-02 | -0.146 | ACKR4,ADCY2,ADCY5,ADGRB2,ADGRD2,ADGRL3,ADRA2A,BDKRB2,BMP6,BMPR1B,CACNA1B,CACNA1C,CCR7,CCR9,CREB5,CRHR2,DRD4,F2R,FZD6,GLP2R,GPR142,GRIA2,GRIK3,GRM1,GRM4,HCAR1,HCAR2,HCAR3,HCRTR2,HGF,HRH4,HTR1D,HTR4,LGR5,LHCGR,NGFR,NOTUM,NPY4R/NPY4R2,NTRK1,NTSR1,PTGER1,TAAR1,TGFB2,TNFRSF11A,TSHR,VIPR1,VN1R1 |
| 2 | Granulocyte Adhesion and Diapedesis | 7.11E+00 | 1.16E-01 | NaN | ACKR4,CCL19,CCL20,CCL26,CCL4,CCL8,CCR7,CCR9,CLDN16,CLDN4,CLDN5,CXCL14,CXCL3,CXCL5,CXCL8,CXCR4,HRH4,IL1B,IL1R2,MMP21,MMP3,NGFR |
| 3 | Agranulocyte Adhesion and Diapedesis | 6.31E+00 | 1.05E-01 | NaN | ACKR4,ACTA1,CCL19,CCL20,CCL26,CCL4,CCL8,CCR7,CCR9,CLDN16,CLDN4,CLDN5,CXCL14,CXCL3,CXCL5,CXCL8,CXCR4,IL1B,MMP21,MMP3,MYH2,MYL4 |
| 4 | Breast Cancer Regulation by Stathmin1 | 5.98E+00 | 6.90E-02 | -0.469 | ACKR4,ADGRB2,ADGRD2,ADGRL3,ADRA2A,ARHGEF10,BDKRB2,BMP6,CCR7,CCR9,CREB5,CRHR2,DRD4,F2R,FZD6,GLP2R,GPR142,GRM1,GRM4,HCAR1,HCAR2,HCAR3,HCRTR2,HGF,HRH4,HTR1D,HTR4,LGR5,LHCGR,NPY4R/NPY4R2,NTSR1,PPP1R14D,PPP2R2C,PTGER1,TAAR1,TGFB2,TSHR,TUBB2B,TUBB4A,VIPR1,VN1R1 |
| 5 | S100 Family Signaling Pathway | 5.97E+00 | 6.35E-02 | 0.143 | ACKR4,ADGRB2,ADGRD2,ADGRL3,ADRA2A,AREG,BDKRB2,CACNA1B,CACNA1C,CCL20,CCL4,CCR7,CCR9,CREB5,CRHR2,CXCL8,DRD4,EREG,F2R,FZD6,GLP2R,GPR142,GRM1,GRM4,HCAR1,HCAR2,HCAR3,HCRTR2,HRH4,HTR1D,HTR4,IL1B,IL23A,LGR5,LHCGR,MMP21,MMP3,NOTUM,NPY4R/NPY4R2,NTF3,NTSR1,PTGER1,S100A7,TAAR1,TGFB2,TPM2,TSHR,VIPR1,VN1R1 |
| 6 | Hepatic Fibrosis / Hepatic Stellate Cell Activation | 5.11E+00 | 9.79E-02 | NaN | AGT,CCN2,CCR7,COL1A2,COL27A1,COL6A5,COL8A1,CXCL3,CXCL8,EDN1,FGF1,HGF,IGF2,IL1B,IL1R2,MYH2,MYL4,NGFR,TGFB2 |
| 7 | Role of IL-17A in Psoriasis | 4.39E+00 | 3.57E-01 | 1.342 | CCL20,CXCL3,CXCL5,CXCL8,S100A7 |
| 8 | G-Protein Coupled Receptor Signaling | 4.24E+00 | 5.83E-02 | -0.781 | ACKR4,ADCY2,ADCY5,ADGRB2,ADGRD2,ADGRL3,ADRA2A,BDKRB2,CCR7,CCR9,CREB5,CRHR2,DRD4,F2R,FOXG1,FZD6,GLP2R,GPR142,GRM1,GRM4,HCAR1,HCAR2,HCAR3,HCRTR2,HRH4,HTR1D,HTR4,LGR5,LHCGR,MAP3K15,MYL11,MYL4,NPY4R/NPY4R2,NTSR1,PPP3R2,PTGER1,RGS4,TAAR1,TSHR,VIPR1,VN1R1 |
| 9 | Airway Pathology in Chronic Obstructive Pulmonary Disease | 4.20E+00 | 1.10E-01 | NaN | CCL20,CXCL3,CXCL8,FGF1,FGF13,IL11,IL17C,IL17D,IL1B,IL23A,LCN9,PAEP,TGFB2 |
| 10 | Phagosome Formation | 3.42E+00 | 5.47E-02 | -0.973 | ACKR4,ADGRB2,ADGRD2,ADGRL3,ADRA2A,BDKRB2,CCR7,CCR9,CRHR2,DRD4,F2R,FCER1A,FCER2,FZD6,GLP2R,GPR142,GRM1,GRM4,HCAR1,HCAR2,HCAR3,HCRTR2,HRH4,HTR1D,HTR4,ITGA2B,LGR5,LHCGR,MYH2,MYL4,NPY4R/NPY4R2,NTSR1,PTGER1,SCARA3,TAAR1,TSHR,VIPR1,VN1R1 |
| | Pathogen Induced Cytokine Storm Signaling Pathway | 2.98E+00 | 6.20E-02 | 2.294 | CCL19,CCL20,CCL26,CCL4,CCL8,COL1A2,COL27A1,COL6A5,COL8A1,CXCL14,CXCL3,CXCL5,CXCL8,CXCR4,IL11,IL17C,IL17D,IL1B,IL1R2,IL23A,NGFR,NLRP12,TGFB2 |
| | Tumor Microenvironment Pathway | 2.93E+00 | 7.82E-02 | 0 | COL1A2,CXCL8,CXCR4,FGF1,FGF13,FOXG1,HGF,HLA-G,IGF2,IL1B,MMP21,MMP3,TGFB2,TNC |
| | Embryonic Stem Cell Differentiation into Cardiac Lineages | 2.54E+00 | 3.00E-01 | NaN | HOXB5,ISL1,NKX2-5 |
| | Role of IL-17F in Allergic Inflammatory Airway Diseases | 2.54E+00 | 1.28E-01 | 2.236 | CCL4,CREB5,CXCL5,CXCL8,IL11,IL1B |
| | Dopamine-DARPP32 Feedback in cAMP Signaling | 2.34E+00 | 6.99E-02 | -1 | ADCY2,ADCY5,CACNA1B,CACNA1C,CALY,CREB5,DRD4,KCNJ4,NOTUM,PPP1R14D,PPP2R2C,PPP3R2,PRKG1 |
| | cAMP-mediated signaling | 2.23E+00 | 6.36E-02 | -0.258 | ADCY2,ADCY5,ADRA2A,CREB5,DRD4,GRM4,HCAR2,HCAR3,HTR1D,HTR4,LHCGR,PPP3R2,RGS4,TAAR1,VIPR1 |
| | Cellular Effects of Sildenafil (Viagra) | 2.21E+00 | 7.33E-02 | NaN | ACTA1,ADCY2,ADCY5,CACNA1B,CACNA1C,MYH2,MYL11,MYL4,NOTUM,PRKG1,SLC4A10 |
| | Multiple Sclerosis Signaling Pathway | 2.09E+00 | 6.31E-02 | 2.673 | CXCL8,HLA-G,IL11,IL17C,IL17D,IL1B,IL23A,KCNG4,MOG,NLRP1,NLRP12,PLP1,SLC4A1,TGFB2 |
| | STAT3 Pathway | 2.08E+00 | 7.41E-02 | 0.707 | BMP6,BMPR1B,HGF,IL1B,IL1R2,IL22RA2,NGFR,NTRK1,TGFB2,TNFRSF11A |
| | SNARE Signaling Pathway | 2.06E+00 | 7.35E-02 | 0 | ADCY2,ADCY5,CACNA1B,MYH2,MYL4,RAB9B,SNAP91,SYN3,SYT1,SYT13 |
| | Dopamine Receptor Signaling | 1.97E+00 | 8.75E-02 | NaN | ADCY2,ADCY5,CALY,DRD4,MAOB,PPP1R14D,PPP2R2C |
| | Hepatic Cholestasis | 1.85E+00 | 6.28E-02 | NaN | ADCY2,ADCY5,CXCL8,FABP6,IL11,IL17C,IL17D,IL1B,IL1R2,INS,NGFR,TGFB2 |
| | Gαs Signaling | 1.83E+00 | 7.14E-02 | -0.333 | ADCY2,ADCY5,CREB5,HCAR2,HCAR3,HTR4,LHCGR,TAAR1,VIPR1 |
| | Synaptogenesis Signaling Pathway | 1.78E+00 | 5.40E-02 | 1 | ADCY2,ADCY5,CACNA1B,CDH17,CDH2,CDH6,CREB5,EPHA10,GRIA2,GRM1,GRM4,MAP1B,SYN3,SYT1,SYT13,THBS4,VLDLR |
| | Airway Inflammation in Asthma | 1.77E+00 | 1.21E-01 | NaN | CCL20,CXCL8,IL23A,TGFB2 |
| | Ceramide Biosynthesis | 1.76E+00 | 2.86E-01 | NaN | DEGS2,SPTSSB |
| | NAD Salvage Pathway III | 1.76E+00 | 2.86E-01 | NaN | NMNAT2,NMRK2 |
| | Coagulation System | 1.68E+00 | 1.14E-01 | -2 | BDKRB2,F13A1,F2R,SERPINA5 |
| | Pulmonary Fibrosis Idiopathic Signaling Pathway | 1.65E+00 | 5.21E-02 | 0.5 | ACTA1,AREG,CCN2,CDH2,COL1A2,COL27A1,COL6A5,COL8A1,EDN1,F2R,FOXG1,FZD6,IL11,IL1B,MMP21,MMP3,TGFB2 |
| | Role of Osteoblasts, Osteoclasts and Chondrocytes in Rheumato | 1.65E+00 | 5.70E-02 | NaN | BMP6,BMPR1B,DKK3,FZD6,IL11,IL1B,IL1R2,MMP3,NGFR,PPP3R2,SFRP1,SFRP2,TNFRSF11A |
| | White Adipose Tissue Browning Pathway | 1.60E+00 | 6.52E-02 | -1 | ADCY2,ADCY5,CACNA1B,CACNA1C,CREB5,FCER1A,LDHC,PRKG1,RUNX1T1 |
| | Cardiac Hypertrophy Signaling (Enhanced) | 1.58E+00 | 4.61E-02 | 0.626 | ADCY2,ADCY5,ADRA2A,AGT,CACNA1B,CACNA1C,CXCL8,EDN1,FGF1,FGF13,FZD6,IL11,IL17C,IL17D,IL1B,IL1R2,IL22RA2,ITGA2B,MAP3K15,NGFR,NKX2-5,NOTUM,PPP3R2,PRKG1,TGFB2 |
| | Role Of Osteoclasts In Rheumatoid Arthritis Signaling Pathway | 1.56E+00 | 5.18E-02 | -1.291 | ADAM19,COL1A2,COL27A1,COL6A5,COL8A1,CREB5,FOXG1,IL1B,IL1R2,MMP21,MMP3,NGFR,PPP3R2,SFRP1,SFRP2,TNFRSF11A |
| | Citrulline Biosynthesis | 1.54E+00 | 2.22E-01 | NaN | OTC,PRODH2 |
| | Citrulline Degradation | 1.52E+00 | 1.00E+00 | NaN | OTC |
| | LXR/RXR Activation | 1.47E+00 | 6.50E-02 | -1.414 | ABCA1,ABCG1,AGT,APOB,HPR,IL1B,IL1R2,NGFR |
| | Aldosterone Signaling in Epithelial Cells | 1.42E+00 | 5.81E-02 | NaN | CRYAB,DNAJB13,DNAJC22,DNAJC5G,DNAJC6,HSPA1A/HSPA1B,HSPA6,HSPB6,NOTUM,SLC12A1 |
| | Endocannabinoid Neuronal Synapse Pathway | 1.42E+00 | 6.04E-02 | 0 | ADCY2,ADCY5,CACNA1B,CACNA1C,DNAH3,GRIA2,GRM1,NOTUM,PPP3R2 |
| | FXR/RXR Activation | 1.41E+00 | 6.35E-02 | NaN | AGT,APOB,FABP6,FOXA2,HPR,IL1B,INS,VLDLR |
| | BEX2 Signaling Pathway | 1.41E+00 | 7.32E-02 | -0.816 | BEX2,CDH2,MAP2,NGFR,NHLH2,PPP2R2C |
| | Axonal Guidance Signaling | 1.40E+00 | 4.51E-02 | NaN | ADAM19,ADAMTS12,ADAMTS6,ADAMTS9,BMP6,CXCR4,EPHA10,FZD6,ITGA2B,MAG,MMP21,MMP3,MYL11,MYL4,NGFR,NOTUM,NTF3,NTRK1,PAPPA,PPP3R2,SEMA3G,TUBB2B,TUBB4A |
| | Gap Junction Signaling | 1.40E+00 | 5.56E-02 | NaN | ACTA1,ADCY2,ADCY5,GJA5,GRIA2,GRIK3,NOTUM,PPP3R2,PRKG1,TUBB2B,TUBB4A |
| | NAD Signaling Pathway | 1.38E+00 | 5.96E-02 | -0.333 | BMP6,HGF,INS,LDHC,NMNAT2,NMRK2,NT5E,SLC22A13,TGFB2 |
| | Wound Healing Signaling Pathway | 1.35E+00 | 5.16E-02 | 0.277 | COL1A2,COL27A1,COL6A5,COL8A1,CXCL8,IL11,IL17C,IL17D,IL1B,IL1R2,LAMA5,NGFR,TGFB2 |
| | CDX Gastrointestinal Cancer Signaling Pathway | 1.35E+00 | 5.45E-02 | -1.508 | BMP6,CDH17,CDX1,CXCL8,FZD6,GATA6,IL11,IL17C,IL17D,IL1B,TGFB2 |
| | Role of Hypercytokinemia/hyperchemokinemia in the Pathogene | 1.33E+00 | 6.98E-02 | 2.449 | AREG,CCL4,CXCL3,CXCL8,IL1B,ISG20 |
| | Tight Junction Signaling | 1.33E+00 | 5.59E-02 | NaN | ACTA1,CLDN16,CLDN4,CLDN5,CRB3,MYH2,MYL4,NGFR,PPP2R2C,TGFB2 |
| | NAD Salvage Pathway II | 1.32E+00 | 1.11E-01 | NaN | NMNAT2,NMRK2,NT5E |
| | Apelin Liver Signaling Pathway | 1.32E+00 | 1.11E-01 | NaN | AGT,COL1A2,EDN1 |
| | Synaptic Long Term Potentiation | 1.32E+00 | 6.06E-02 | -0.378 | CACNA1C,CREB5,GRIA2,GRM1,GRM4,NOTUM,PPP1R14D,PPP3R2 |
| | LPS/IL-1 Mediated Inhibition of RXR Function | 1.31E+00 | 5.10E-02 | -0.447 | ABCA1,ABCG1,ALDH1L2,CHST1,FABP6,FMO1,HMGCS1,IL1B,IL1R2,MAOB,NGFR,SULT1B1,SULT4A1 |
| | Heparan Sulfate Biosynthesis | 1.31E+00 | 6.90E-02 | -0.447 | B3GAT1,CHST1,NOTUM,SULT1B1,SULT4A1,TNFAIP6 |
| | Glutamate Receptor Signaling | 1.30E+00 | 7.58E-02 | NaN | GRIA2,GRIK3,GRIP1,GRM1,GRM4 |
| | Sertoli Cell-Sertoli Cell Junction Signaling | 1.30E+00 | 5.34E-02 | NaN | ACTA1,CLDN16,CLDN4,CLDN5,ITGA2B,MAP3K15,PRKG1,SORBS1,SPTA1,TUBB2B,TUBB4A |
| | HIF1α Signaling | 1.27E+00 | 5.29E-02 | -0.905 | BMP6,EDN1,HGF,HSPA1A/HSPA1B,HSPA6,IGF2,LDHC,MMP21,MMP3,PPP3R2,TGFB2 |
| | IL-17 Signaling | 1.22E+00 | 5.35E-02 | 2.53 | CCL20,CXCL3,CXCL5,CXCL8,IL11,IL17C,IL17D,IL1B,MMP3,TGFB2 |
| | CDK5 Signaling | 1.21E+00 | 6.09E-02 | 1.134 | ADCY2,ADCY5,FOSB,LAMA5,NGFR,PPP1R14D,PPP2R2C |
| | Gαi Signaling | 1.19E+00 | 5.71E-02 | 0.707 | ADCY2,ADCY5,ADRA2A,DRD4,GRM4,HCAR2,HTR1D,RGS4 |
| | Role Of Chondrocytes In Rheumatoid Arthritis Signaling Pathway | 1.18E+00 | 5.67E-02 | 0.707 | CXCL8,CXCR4,IL1B,IL1R2,MMP21,MMP3,NGFR,PPP3R2 |
| | Activin Inhibin Signaling Pathway | 1.17E+00 | 5.07E-02 | 1.508 | CCN2,COL1A2,CXCR4,FOSB,IL11,IL1B,IL1R2,NGFR,POU1F1,SOX17,TGFB2 |
| | Role Of Osteoblasts In Rheumatoid Arthritis Signaling Pathway | 1.16E+00 | 4.92E-02 | 1.155 | BMPR1B,CXCL8,FZD6,IL11,IL17C,IL17D,IL1B,MMP21,MMP3,SFRP1,SFRP2,TGFB2 |
| | Superpathway of Citrulline Metabolism | 1.13E+00 | 1.33E-01 | NaN | OTC,PRODH2 |
| | Neuroprotective Role of THOP1 in Alzheimer's Disease | 1.12E+00 | 5.79E-02 | 1.342 | AGT,HLA-G,HTRA3,PRSS16,PRSS21,PRSS23,PRSS33 |
| | GPCR-Mediated Integration of Enteroendocrine Signaling Exemp | 1.11E+00 | 6.67E-02 | 0.447 | ADCY2,ADCY5,GLP2R,NOTUM,VIPR1 |
| | Role of Cytokines in Mediating Communication between Immun | 1.09E+00 | 7.41E-02 | NaN | CXCL8,IL1B,IL23A,IL32 |
| | Bile Acid Biosynthesis, Neutral Pathway | 1.08E+00 | 1.25E-01 | NaN | AKR1C1/AKR1C2,AKR1D1 |
| | WNT/β-catenin Signaling | 1.07E+00 | 5.17E-02 | -1.414 | CDH2,DKK3,FZD6,PPP2R2C,SFRP1,SFRP2,SOX17,SOX5,TGFB2 |
| | TREM1 Signaling | 1.07E+00 | 6.49E-02 | 2.236 | CXCL3,CXCL8,IL1B,NLRP12,TREM1 |
| | Proline Degradation | 1.05E+00 | 3.33E-01 | NaN | PRODH2 |
| | Gustation Pathway | 1.04E+00 | 4.93E-02 | 1 | ADCY2,ADCY5,CACNA1B,CACNA1C,GABRA2,GABRG3,GABRQ,NOTUM,SCN3A,SCN7A |
| | Maturity Onset Diabetes of Young (MODY) Signaling | 1.03E+00 | 6.33E-02 | NaN | APOB,CACNA1B,CACNA1C,FOXA2,INS |
| | Thyroid Cancer Signaling | 1.03E+00 | 6.33E-02 | 2.236 | CXCL8,CXCR4,INS,NTF3,NTRK1 |
| | GP6 Signaling Pathway | 1.03E+00 | 5.51E-02 | -0.816 | COL1A2,COL27A1,COL6A5,COL8A1,GRAP2,ITGA2B,LAMA5 |
| | Role of IL-17A in Arthritis | 1.02E+00 | 7.02E-02 | NaN | CCL20,CXCL3,CXCL5,CXCL8 |
| | Triacylglycerol Degradation | 1.02E+00 | 7.02E-02 | NaN | CES1,NOTUM,PNPLA5,TNFAIP6 |
| | Factors Promoting Cardiogenesis in Vertebrates | 1.02E+00 | 5.23E-02 | 0.707 | BMP6,BMPR1B,CREB5,FZD6,GJA5,NKX2-5,NOTUM,TGFB2 |
| | Heparan Sulfate Biosynthesis (Late Stages) | 1.02E+00 | 6.25E-02 | -1 | CHST1,NOTUM,SULT1B1,SULT4A1,TNFAIP6 |
| | Neurovascular Coupling Signaling Pathway | 1.01E+00 | 4.74E-02 | 1.508 | BDKRB2,CACNA1B,CACNA1C,CYP4Z1,GABRA2,GABRG3,GABRQ,GRIA2,GRM1,KCNJ4,PRKG1 |
| | Complement System | 9.94E-01 | 8.11E-02 | NaN | C1S,CFH,CFI |
| | Cardiac Hypertrophy Signaling | 9.91E-01 | 4.60E-02 | -0.632 | ADCY2,ADCY5,ADRA2A,CACNA1B,CACNA1C,MAP3K15,MYL11,MYL4,NKX2-5,NOTUM,PPP3R2,TGFB2 |
| | Differential Regulation of Cytokine Production in Macrophages a | 9.90E-01 | 1.11E-01 | NaN | CCL4,IL1B |
| | FAK Signaling | 9.83E-01 | 3.75E-02 | -1.441 | ACKR4,ADGRB2,ADGRD2,ADGRL3,ADRA2A,BDKRB2,CCR7,CCR9,CD3G,COL1A2,CRHR2,DRD4,F2R,FZD6,GLP2R,GPR142,GRM1,GRM4,HCAR1,HCAR2,HCAR3,HCRTR2,HRH4,HTR1D,HTR4,IL1R2,IL22RA2,ITGA2B,LGR5,LHCGR,NPY4R/NPY4R2,NTSR1,PTGER1,TAAR1,TGFB2,TRBV5-4,TSHR,VIPR1,VN1R1 |
| | Chondroitin Sulfate Biosynthesis | 9.82E-01 | 6.78E-02 | 0 | B3GAT1,CHST1,SULT1B1,SULT4A1 |
| | GABA Receptor Signaling | 9.68E-01 | 5.30E-02 | NaN | ADCY2,ADCY5,CACNA1B,CACNA1C,GABRA2,GABRG3,GABRQ |
| | Dermatan Sulfate Biosynthesis | 9.43E-01 | 6.56E-02 | 0 | B3GAT1,CHST1,SULT1B1,SULT4A1 |
| | Inhibition of Matrix Metalloproteases | 9.42E-01 | 7.69E-02 | NaN | MMP21,MMP3,TFPI2 |
| | Taurine Biosynthesis | 9.35E-01 | 2.50E-01 | NaN | FMO1 |
| | Melatonin Degradation II | 9.35E-01 | 2.50E-01 | NaN | MAOB |
| | Inflammasome pathway | 9.13E-01 | 1.00E-01 | NaN | IL1B,NLRP1 |
| | NOD1/2 Signaling Pathway | 9.10E-01 | 4.76E-02 | 1.667 | CXCL8,HSPA1A/HSPA1B,HSPA6,IL11,IL17C,IL17D,IL1B,NLRP12,TGFB2 |
| | Serotonin Receptor Signaling | 9.01E-01 | 4.05E-02 | -0.229 | ADCY2,ADCY5,ADRA2A,CACNA1B,CACNA1C,CCR7,CREB5,CXCL8,EDN1,F13A1,GAP43,HTR1D,HTR4,IL1B,ITGA2B,MAOB,NOTUM,TGFB2,TGM7 |
| | RHOGDI Signaling | 8.72E-01 | 4.55E-02 | -1.134 | ACTA1,ARHGEF10,CDH17,CDH2,CDH6,GRIP1,ITGA2B,MYH2,MYL11,MYL4 |
| | Calcium Signaling | 8.72E-01 | 4.55E-02 | 0.816 | ACTA1,CACNA1B,CACNA1C,CHRNB2,CREB5,GRIA2,MYH2,MYL4,PPP3R2,TPM2 |
| | Unfolded protein response | 8.58E-01 | 5.56E-02 | NaN | DNAJB13,DNAJC5G,DNAJC6,HSPA1A/HSPA1B,HSPA6 |
| | Proline Biosynthesis II (from Arginine) | 8.45E-01 | 2.00E-01 | NaN | OTC |
| | Creatine-phosphate Biosynthesis | 8.45E-01 | 2.00E-01 | NaN | CKMT1A/CKMT1B |
| | Apelin Adipocyte Signaling Pathway | 8.44E-01 | 5.49E-02 | 1 | ADCY2,ADCY5,GPX8,INS,PTGES |
| | ABRA Signaling Pathway | 8.30E-01 | 5.43E-02 | 0.447 | ACTA1,CCN2,FOSB,PPP3R2,TPM2 |
| | Hepatic Fibrosis Signaling Pathway | 8.29E-01 | 4.02E-02 | 1.069 | AGT,CACNA1B,CACNA1C,CCN2,COL1A2,CREB5,CXCL8,EDN1,FZD6,IL1B,IL1R2,INS,ITGA2B,MYL11,MYL4,NGFR,TGFB2 |
| | Synaptic Long Term Depression | 8.24E-01 | 4.55E-02 | -0.333 | CACNA1B,CACNA1C,GRIA2,GRM1,GRM4,NOTUM,PPP1R17,PPP2R2C,PRKG1 |
| | Remodeling of Epithelial Adherens Junctions | 8.19E-01 | 5.88E-02 | NaN | ACTA1,HGF,TUBB2B,TUBB4A |
| | Differential Regulation of Cytokine Production in Intestinal Epith | 8.14E-01 | 8.70E-02 | NaN | CCL4,IL1B |
| | Pyrimidine Deoxyribonucleotides De Novo Biosynthesis I | 8.14E-01 | 8.70E-02 | NaN | AK7,AK8 |
| | Apelin Cardiac Fibroblast Signaling Pathway | 8.14E-01 | 8.70E-02 | NaN | CCN2,TGFB2 |
| | Arginine Biosynthesis IV | 7.72E-01 | 1.67E-01 | NaN | OTC |
| | Pyruvate Fermentation to Lactate | 7.72E-01 | 1.67E-01 | NaN | LDHC |
| | Urea Cycle | 7.72E-01 | 1.67E-01 | NaN | OTC |
| | Glycine Cleavage Complex | 7.72E-01 | 1.67E-01 | NaN | GLDC |
| | UDP-N-acetyl-D-glucosamine Biosynthesis II | 7.72E-01 | 1.67E-01 | NaN | GFPT2 |
| | Zymosterol Biosynthesis | 7.72E-01 | 1.67E-01 | NaN | MSMO1 |
| | NAD Biosynthesis III | 7.72E-01 | 1.67E-01 | NaN | NMNAT2 |
| | Estrogen Biosynthesis | 7.68E-01 | 6.38E-02 | NaN | AKR1C1/AKR1C2,HSD17B13,HSD17B2 |
| | Dilated Cardiomyopathy Signaling Pathway | 7.63E-01 | 4.67E-02 | 0 | ACTA1,ADCY2,ADCY5,CACNA1B,CACNA1C,MYH2,MYL4 |
| | Netrin Signaling | 7.56E-01 | 5.56E-02 | 0 | CACNA1B,CACNA1C,PPP3R2,PRKG1 |
| | Dermatan Sulfate Biosynthesis (Late Stages) | 7.49E-01 | 6.25E-02 | NaN | CHST1,SULT1B1,SULT4A1 |
| | Clathrin-mediated Endocytosis Signaling | 7.37E-01 | 4.33E-02 | NaN | ACTA1,APOB,F2R,FGF1,FGF13,INS,PPP3R2,SH3BP4,SNAP91 |
| | IL-17A Signaling in Gastric Cells | 7.30E-01 | 7.69E-02 | NaN | CCL20,CXCL8 |
| | Caveolar-mediated Endocytosis Signaling | 7.13E-01 | 5.33E-02 | NaN | ACTA1,HLA-G,INS,ITGA2B |
| | FAT10 Cancer Signaling Pathway | 7.13E-01 | 6.00E-02 | NaN | CXCR4,NGFR,TGFB2 |
| | NAD Biosynthesis from 2-amino-3-carboxymuconate Semialdehy | 7.12E-01 | 1.43E-01 | NaN | NMNAT2 |
| | Glycoaminoglycan-protein Linkage Region Biosynthesis | 7.12E-01 | 1.43E-01 | NaN | B3GAT1 |
| | eNOS Signaling | 7.05E-01 | 4.49E-02 | -0.378 | ADCY2,ADCY5,BDKRB2,CHRNB2,HSPA1A/HSPA1B,HSPA6,PRKG1 |
| | Cardiomyocyte Differentiation via BMP Receptors | 7.05E-01 | 7.41E-02 | NaN | BMPR1B,NKX2-5 |

| | | | | |
|---|---|---|---|---|
| Leptin Signaling in Obesity | 6.99E-01 | 5.26E-02 | NaN | ADCY2,ADCY5,INS,NOTUM |
| Chondroitin Sulfate Biosynthesis (Late Stages) | 6.96E-01 | 5.88E-02 | NaN | CHST1,SULT1B1,SULT4A1 |
| Glutathione Redox Reactions I | 6.81E-01 | 7.14E-02 | NaN | GPX8,PTGES |
| Role of Macrophages, Fibroblasts and Endothelial Cells in Rheum | 6.74E-01 | 3.92E-02 | NaN | CREB5,CXCL8,DKK3,FZD6,IL1B,IL1R2,IL32,MMP3,NGFR,NOTUM,PPP3R2,SFRP1,SFRP2 |
| Neurotrophin/TRK Signaling | 6.73E-01 | 5.13E-02 | NaN | CREB5,NGFR,NTF3,NTRK1 |
| Macrophage Classical Activation Signaling Pathway | 6.62E-01 | 4.23E-02 | 2.121 | CCL20,CXCL8,IL11,IL17C,IL17D,IL1B,IL23A,TGFB2 |
| Gα12/13 Signaling | 6.61E-01 | 4.51E-02 | 0 | CDH17,CDH2,CDH6,F2R,MYL11,MYL4 |
| Atherosclerosis Signaling | 6.61E-01 | 4.51E-02 | NaN | APOB,COL1A2,CXCL8,CXCR4,IL1B,MMP3 |
| Superpathway of Cholesterol Biosynthesis | 6.58E-01 | 6.90E-02 | NaN | HMGCS1,MSMO1 |
| Regulation Of The Epithelial Mesenchymal Transition By Growth | 6.38E-01 | 4.17E-02 | 0 | CDH2,FGF1,FGF13,HGF,HMGA2,NGFR,TGFB2,TWIST1 |
| RAR Activation | 6.35E-01 | 3.74E-02 | -1 | ADCY2,ADCY5,CDX1,COL1A2,CRABP1,CREB5,CXCL8,GATA6,HOXB5,HOXD3,IL11,IL17C,IL17D,IL1B,MMP3,TGFB2 |
| Transcriptional Regulatory Network in Embryonic Stem Cells | 6.34E-01 | 4.27E-02 | 0.378 | BMPR1B,FZD6,GATA6,IGF2,INS,ISL1,TGFB2 |
| Role of PKR in Interferon Induction and Antiviral Response | 6.33E-01 | 4.41E-02 | NaN | HSPA1A/HSPA1B,HSPA6,IL1B,NLRP1,NLRP12,SCARA3 |
| Leukocyte Extravasation Signaling | 6.31E-01 | 4.15E-02 | -0.447 | ACTA1,BMX,CLDN16,CLDN4,CLDN5,CXCR4,MMP21,MMP3 |
| Xenobiotic Metabolism PXR Signaling Pathway | 6.31E-01 | 4.15E-02 | 0 | ALDH1L2,CES1,CHST1,GRIP1,MAOB,PPP1R14D,SULT1B1,SULT4A1 |
| PPARα/RXRα Activation | 6.16E-01 | 4.10E-02 | 0.378 | ABCA1,ADCY2,ADCY5,IL1B,IL1R2,INS,NOTUM,TGFB2 |
| Regulation of the Epithelial-Mesenchymal Transition Pathway | 6.16E-01 | 4.10E-02 | NaN | CDH2,FGF1,FGF13,FZD6,HGF,HMGA2,TGFB2,TWIST1 |
| Pathogenesis of Multiple Sclerosis | 6.15E-01 | 1.11E-01 | NaN | CCL4 |
| Role of NFAT in Cardiac Hypertrophy | 6.15E-01 | 4.02E-02 | -0.707 | ADCY2,ADCY5,CACNA1B,CACNA1C,IL11,NKX2-5,NOTUM,PPP3R2,TGFB2 |
| Estrogen-Dependent Breast Cancer Signaling | 6.11E-01 | 4.82E-02 | NaN | AKR1C1/AKR1C2,CREB5,HSD17B13,HSD17B2 |
| HMGB1 Signaling | 6.10E-01 | 4.19E-02 | 1.89 | CXCL8,IL11,IL17C,IL17D,IL1B,NGFR,TGFB2 |
| Dopamine Degradation | 5.96E-01 | 6.25E-02 | NaN | MAOB,SULT1B1 |
| Neuroinflammation Signaling Pathway | 5.88E-01 | 3.79E-02 | 1.89 | CALB2,CREB5,CXCL8,GABRA2,GABRG3,GABRQ,HLA-G,IL1B,MMP3,NTF3,PPP3R2,TGFB2 |
| Germ Cell-Sertoli Cell Junction Signaling | 5.86E-01 | 4.12E-02 | NaN | ACTA1,CDH2,MAP3K15,SORBS1,TGFB2,TUBB2B,TUBB4A |
| FGF Signaling | 5.77E-01 | 4.65E-02 | 0 | CREB5,FGF1,FGF13,HGF |
| Prostanoid Biosynthesis | 5.76E-01 | 1.00E-01 | NaN | PTGES |
| Primary Immunodeficiency Signaling | 5.75E-01 | 5.08E-02 | NaN | CD19,CD79A,ZAP70 |
| ID1 Signaling Pathway | 5.72E-01 | 3.98E-02 | 0 | BHLHA15,BHLHE22,BMPR1B,CCN2,CHRNB2,IGF2,NGFR,TGFB2 |
| Xenobiotic Metabolism Signaling | 5.60E-01 | 3.77E-02 | NaN | ALDH1L2,CES1,CHST1,FMO1,GRIP1,IL1B,MAOB,MAP3K15,PPP2R2C,SULT1B1,SULT4A1 |
| Amyotrophic Lateral Sclerosis Signaling | 5.57E-01 | 4.31E-02 | NaN | CACNA1B,CACNA1C,GRIA2,GRIK3,PRPH |
| Bladder Cancer Signaling | 5.57E-01 | 4.31E-02 | NaN | CXCL8,FGF1,FGF13,MMP21,MMP3 |
| Ketogenesis | 5.41E-01 | 9.09E-02 | NaN | HMGCS1 |
| GPCR-Mediated Nutrient Sensing in Enteroendocrine Cells | 5.39E-01 | 4.24E-02 | -0.447 | ADCY2,ADCY5,CACNA1B,CACNA1C,NOTUM |
| Immunogenic Cell Death Signaling Pathway | 5.34E-01 | 4.44E-02 | 2 | HSPA1A/HSPA1B,HSPA6,IL1B,NGFR |
| Myelination Signaling Pathway | 5.34E-01 | 3.67E-02 | 1.155 | BMP6,BMPR1B,CREB5,ERBB4,FZD6,LAMA5,MAG,NGFR,NKX2-2,NTF3,PLP1,PPP3R2 |
| Hematopoiesis from Multipotent Stem Cells | 5.10E-01 | 8.33E-02 | NaN | THPO |
| PTEN Signaling | 5.08E-01 | 3.97E-02 | -1.342 | BMPR1B,FOXG1,ITGA2B,NGFR,NTRK1,TNFRSF11A |
| Pyroptosis Signaling Pathway | 5.05E-01 | 4.30E-02 | 2 | IL1B,NGFR,NLRP1,NLRP12 |
| AMPK Signaling | 5.01E-01 | 3.72E-02 | 0.447 | ADRA2A,AK7,AK8,CHRNB2,CREB5,FOXG1,INS,PPP2R2C,RAB9B |
| Corticotropin Releasing Hormone Signaling | 5.01E-01 | 3.95E-02 | 0 | ADCY2,ADCY5,CACNA1B,CACNA1C,CREB5,CRHR2 |
| IL-15 Production | 4.97E-01 | 4.07E-02 | 0.447 | BMX,ERBB4,NTRK1,TNK1,ZAP70 |
| Actin Cytoskeleton Signaling | 4.90E-01 | 3.69E-02 | -0.447 | ACTA1,F2R,FGF1,FGF13,INS,ITGA2B,MYH2,MYL11,MYL4 |
| WNT/Ca+ pathway | 4.89E-01 | 4.55E-02 | NaN | CREB5,FZD6,NOTUM |
| NAD biosynthesis II (from tryptophan) | 4.81E-01 | 7.69E-02 | NaN | NMNAT2 |
| Cholesterol Biosynthesis I | 4.81E-01 | 7.69E-02 | NaN | MSMO1 |
| γ-glutamyl Cycle | 4.81E-01 | 7.69E-02 | NaN | CHAC1 |
| Cholesterol Biosynthesis II (via 24,25-dihydrolanosterol) | 4.81E-01 | 7.69E-02 | NaN | MSMO1 |
| Guanosine Nucleotides Degradation III | 4.81E-01 | 7.69E-02 | NaN | NT5E |
| Cholesterol Biosynthesis III (via Desmosterol) | 4.81E-01 | 7.69E-02 | NaN | MSMO1 |
| Pyrimidine Ribonucleotides Interconversion | 4.78E-01 | 5.13E-02 | NaN | AK7,AK8 |
| IL-17A Signaling in Airway Cells | 4.78E-01 | 4.48E-02 | NaN | CCL20,CXCL3,CXCL5 |
| Autophagy | 4.75E-01 | 3.70E-02 | 0.707 | BMP6,CREB5,HGF,INS,NGFR,PPP2R2C,PPP3R2,TGFB2 |
| Role of Pattern Recognition Receptors in Recognition of Bacteria | 4.72E-01 | 3.85E-02 | NaN | CXCL8,IL11,IL17C,IL17D,IL1B,TGFB2 |
| Epithelial Adherens Junction Signaling | 4.65E-01 | 3.82E-02 | 0.816 | CDH2,FGF1,HGF,MYH2,PPP2R2C,TGFB2 |
| Acyl-CoA Hydrolysis | 4.55E-01 | 7.14E-02 | NaN | THEM5 |
| Urate Biosynthesis/Inosine 5'-phosphate Degradation | 4.55E-01 | 7.14E-02 | NaN | NT5E |
| Mevalonate Pathway I | 4.55E-01 | 7.14E-02 | NaN | HMGCS1 |
| Phenylalanine Degradation IV (Mammalian, via Side Chain) | 4.55E-01 | 7.14E-02 | NaN | MAOB |
| Macrophage Alternative Activation Signaling Pathway | 4.55E-01 | 3.70E-02 | -0.378 | ABCA1,CREB5,FCER1A,FCER2,IL1B,IL1R2,TGFB2 |
| IL-6 Signaling | 4.50E-01 | 3.88E-02 | 1.342 | CXCL8,IL1B,IL1R2,NGFR,TNFAIP6 |
| Xenobiotic Metabolism CAR Signaling Pathway | 4.37E-01 | 3.65E-02 | 0.378 | ALDH1L2,CHST1,FMO1,GRIP1,PPP2R2C,SULT1B1,SULT4A1 |
| Intrinsic Prothrombin Activation Pathway | 4.36E-01 | 4.76E-02 | NaN | COL1A2,F13A1 |
| Pyrimidine Ribonucleotides De Novo Biosynthesis | 4.36E-01 | 4.76E-02 | NaN | AK7,AK8 |
| CLEAR Signaling Pathway | 4.36E-01 | 3.51E-02 | -0.632 | BMP6,BMPR1B,CREB5,HGF,NGFR,NTRK1,PPP2R2C,PPP3R2,TGFB2,TNFRSF11A |
| Neuropathic Pain Signaling In Dorsal Horn Neurons | 4.34E-01 | 3.96E-02 | 0 | GRIA2,GRM1,GRM4,NOTUM |
| Protein Kinase A Signaling | 4.33E-01 | 3.41E-02 | -1.155 | ADCY2,ADCY5,CREB5,DUSP21,MYH2,MYL11,MYL4,NGFR,NOTUM,PPP1R14D,PPP3R2,PTPN3,PTPRQ,TGFB2 |
| Thrombin Signaling | 4.25E-01 | 3.56E-02 | 0 | ADCY2,ADCY5,ARHGEF10,F2R,GATA6,MYL11,MYL4,NOTUM |
| P2Y Purigenic Receptor Signaling Pathway | 4.22E-01 | 3.76E-02 | 0 | ADCY2,ADCY5,CREB5,ITGA2B,NOTUM |
| Sperm Motility | 4.21E-01 | 3.50E-02 | NaN | ATP1A4,BMX,ERBB4,NOTUM,NTRK1,PRKG1,TNK1,ZAN,ZAP70 |
| Extrinsic Prothrombin Activation Pathway | 4.09E-01 | 6.25E-02 | NaN | F13A1 |
| Chondroitin Sulfate Degradation (Metazoa) | 4.09E-01 | 6.25E-02 | NaN | HYAL1 |
| Adenosine Nucleotides Degradation II | 4.09E-01 | 6.25E-02 | NaN | NT5E |
| Coronavirus Replication Pathway | 3.99E-01 | 4.44E-02 | NaN | TUBB2B,TUBB4A |
| Pulmonary Healing Signaling Pathway | 3.97E-01 | 3.52E-02 | 1.89 | BMPR1B,CXCR4,FZD6,MMP21,MMP3,NGFR,TGFB2 |
| Adrenomedullin signaling pathway | 3.97E-01 | 3.52E-02 | 0.378 | ADCY2,ADCY5,CFH,IL1B,NOTUM,PRKG1,RAMP2 |
| Dermatan Sulfate Degradation (Metazoa) | 3.89E-01 | 5.88E-02 | NaN | HYAL1 |
| Reelin Signaling in Neurons | 3.88E-01 | 3.62E-02 | 2 | ARHGEF10,CDH2,DNAH3,MAP1B,VLDLR |
| Iron homeostasis signaling pathway | 3.88E-01 | 3.62E-02 | NaN | BMP6,BMPR1B,CP,HEPH,HIF3A |
| IL-23 Signaling Pathway | 3.88E-01 | 4.35E-02 | NaN | IL1B,IL23A |
| PPAR Signaling | 3.88E-01 | 3.74E-02 | -1 | IL1B,IL1R2,INS,NGFR |
| PD-1, PD-L1 cancer immunotherapy pathway | 3.88E-01 | 3.74E-02 | NaN | HLA-G,NGFR,TGFB2,ZAP70 |
| Ephrin A Signaling | 3.77E-01 | 4.26E-02 | NaN | EPHA10,NGFR |
| Retinol Biosynthesis | 3.77E-01 | 4.26E-02 | NaN | CES1,PNPLA5 |
| VDR/RXR Activation | 3.73E-01 | 3.85E-02 | NaN | HR,HSD17B2,TGFB2 |
| Superpathway of Geranylgeranyldiphosphate Biosynthesis I (via | 3.71E-01 | 5.56E-02 | NaN | HMGCS1 |
| Osteoarthritis Pathway | 3.70E-01 | 3.39E-02 | 0.816 | CREB5,CXCL8,FZD6,IL1B,IL1R2,ITGA2B,MMP3,NKX3-2 |
| Circadian Rhythm Signaling | 3.69E-01 | 3.36E-02 | NaN | ADCY2,ADCY5,CACNA1B,CACNA1C,CREB5,GRIA2,NGFR,NOTUM,PRKG1 |
| nNOS Signaling in Skeletal Muscle Cells | 3.66E-01 | 4.17E-02 | NaN | CACNA1B,CACNA1C |
| PFKFB4 Signaling Pathway | 3.66E-01 | 4.17E-02 | NaN | CREB5,TGFB2 |
| Role of JAK family kinases in IL-6-type Cytokine Signaling | 3.65E-01 | 3.80E-02 | NaN | IL11,SERPINA5,TGFB2 |
| Role of Tissue Factor in Cancer | 3.56E-01 | 3.38E-02 | 0.378 | CCN2,CXCL8,F2R,HGF,IL1B,NGFR,TGFB2 |
| Purine Nucleotides Degradation II (Aerobic) | 3.53E-01 | 5.26E-02 | NaN | NT5E |
| Protein Ubiquitination Pathway | 3.48E-01 | 3.30E-02 | NaN | CRYAB,DNAJB13,DNAJC22,DNAJC5G,DNAJC6,HLA-G,HSPA1A/HSPA1B,HSPA6,HSPB6 |
| Erythropoietin Signaling Pathway | 3.46E-01 | 3.39E-02 | -1.633 | CXCL8,IL11,IL17C,IL17D,IL1B,TGFB2 |
| Endocannabinoid Cancer Inhibition Pathway | 3.35E-01 | 3.40E-02 | -0.447 | ADCY2,ADCY5,CREB5,NUPR1,TWIST1 |
| Regulation of Actin-based Motility by Rho | 3.34E-01 | 3.48E-02 | NaN | ACTA1,ITGA2B,MYL11,MYL4 |
| Cardiac β-adrenergic Signaling | 3.31E-01 | 3.33E-02 | -1.342 | ADCY2,ADCY5,CACNA1B,CACNA1C,PPP1R14D,PPP2R2C |
| Estrogen Receptor Signaling | 3.25E-01 | 3.18E-02 | 0.577 | ADCY2,ADCY5,AGT,CACNA1B,CACNA1C,CREB5,FOXG1,IGF2,MMP21,MMP3,MYL11,MYL4,NOTUM |
| Neuregulin Signaling | 3.22E-01 | 3.42E-02 | NaN | AREG,ERBB4,EREG,ITGA2B |
| Opioid Signaling Pathway | 3.20E-01 | 3.21E-02 | -0.378 | ADCY2,ADCY5,CACNA1B,CACNA1C,CREB5,FOSB,PPP3R2,RGS4,SCN7A |
| Putrescine Degradation III | 3.07E-01 | 4.55E-02 | NaN | MAOB |
| Acute Phase Response Signaling | 3.07E-01 | 3.24E-02 | 2 | AGT,C1S,CP,CRABP1,IL1B,NGFR |
| NGF Signaling | 3.04E-01 | 3.33E-02 | 1 | CREB5,MAP3K15,NGFR,NTRK1 |
| Nitric Oxide Signaling in the Cardiovascular System | 3.04E-01 | 3.33E-02 | NaN | BDKRB2,CACNA1B,CACNA1C,PRKG1 |
| p38 MAPK Signaling | 3.04E-01 | 3.33E-02 | 0 | CREB5,IL1B,IL1R2,TGFB2 |
| IL-33 Signaling Pathway | 3.02E-01 | 3.23E-02 | 2.449 | AREG,CREB5,CXCL8,IL1B,PRSS21,PRSS33 |
| Glucocorticoid Receptor Signaling | 2.99E-01 | 3.09E-02 | NaN | AGT,CD3G,CXCL3,CXCL8,DNAH3,HLA-G,HSPA1A/HSPA1B,HSPA6,IL1B,IL1R2,IL22RA2,KRT15,KRT76,MMP3,PPP3R2,SLPI,TGFB2,VIPR1 |
| The Visual Cycle | 2.94E-01 | 4.35E-02 | NaN | RLBP1 |
| Role of NANOG in Mammalian Embryonic Stem Cell Pluripotency | 2.82E-01 | 3.23E-02 | NaN | BMP6,BMPR1B,FZD6,GATA6 |
| IL-22 Signaling | 2.81E-01 | 4.17E-02 | NaN | IL22RA2 |
| Tumoricidal Function of Hepatic Natural Killer Cells | 2.81E-01 | 4.17E-02 | NaN | LYVE1 |
| GNRH Signaling | 2.80E-01 | 3.14E-02 | 0 | ADCY2,ADCY5,CACNA1B,CACNA1C,CREB5,MAP3K15 |
| Crosstalk between Dendritic Cells and Natural Killer Cells | 2.80E-01 | 3.30E-02 | NaN | ACTA1,CCR7,HLA-G |
| RANK Signaling in Osteoclasts | 2.80E-01 | 3.30E-02 | NaN | MAP3K15,PPP3R2,TNFRSF11A |
| BMP signaling pathway | 2.80E-01 | 3.30E-02 | NaN | BMP6,BMPR1B,NKX2-5 |
| ERBB Signaling | 2.68E-01 | 3.23E-02 | NaN | AREG,ERBB4,EREG |
| GADD45 Signaling | 2.61E-01 | 3.33E-02 | NaN | IL1B,TGFB2 |
| TGF-β Signaling | 2.51E-01 | 3.12E-02 | NaN | BMPR1B,NKX2-5,TGFB2 |
| Tryptophan Degradation X (Mammalian, via Tryptamine) | 2.48E-01 | 3.70E-02 | NaN | MAOB |
| Methionine Degradation I (to Homocysteine) | 2.48E-01 | 3.70E-02 | NaN | PRMT8 |
| Role of JAK2 in Hormone-like Cytokine Signaling | 2.48E-01 | 3.23E-02 | NaN | PAEP,THPO |
| Salvage Pathways of Pyrimidine Ribonucleotides | 2.45E-01 | 3.09E-02 | NaN | AK7,AK8,MAK |
| Melanocyte Development and Pigmentation Signaling | 2.40E-01 | 3.06E-02 | NaN | ADCY2,ADCY5,CREB5 |
| Cysteine Biosynthesis III (mammalia) | 2.28E-01 | 3.45E-02 | NaN | PRMT8 |
| PXR/RXR Activation | 2.28E-01 | 3.08E-02 | NaN | FOXA2,INS |
| Induction of Apoptosis by HIV1 | 2.28E-01 | 3.08E-02 | NaN | CXCR4,NGFR |
| Methylglyoxal Degradation III | 2.11E-01 | 3.23E-02 | NaN | AKR1C1/AKR1C2 |

| | | | | |
|---|---|---|---|---|
| CSDE1 Signaling Pathway | 0.00E+00 | 1.79E-02 | NaN | TNC |
| Oxytocin In Brain Signaling Pathway | 0.00E+00 | 3.02E-02 | -0.816 | CACNA1B,CACNA1C,CREB5,IL1B,NLRP1,NLRP12 |
| Oxytocin Signaling Pathway | 0.00E+00 | 2.84E-02 | 0 | CACNA1B,CACNA1C,CREB5,CXCL8,MYH2,MYL4,PPP3R2,PRKG1 |
| IL-13 Signaling Pathway | 0.00E+00 | 2.59E-02 | NaN | CCL26,FOXA2,TGFB2 |
| Oxytocin In Spinal Neurons Signaling Pathway | 0.00E+00 | 2.86E-02 | NaN | PRKG1 |
| MicroRNA Biogenesis Signaling Pathway | 0.00E+00 | 1.60E-02 | NaN | BMP6,HGF,TGFB2 |
| Ribonucleotide Reductase Signaling Pathway | 0.00E+00 | 1.18E-02 | NaN | CREB5,HGF |
| Natural Killer Cell Signaling | 0.00E+00 | 3.03E-02 | 0.816 | COL1A2,HLA-G,HSPA1A/HSPA1B,HSPA6,MAP3K15,ZAP70 |
| IL-10 Signaling | 0.00E+00 | 2.60E-02 | 0 | CREB5,HLA-G,IL1B,IL1R2 |
| Neutrophil Extracellular Trap Signaling Pathway | 0.00E+00 | 2.01E-02 | 0.707 | COL1A2,COL27A1,COL6A5,COL8A1,CXCL8,IL1B,NOTUM,PPP3R2 |
| Fc Epsilon RI Signaling | 0.00E+00 | 1.69E-02 | NaN | FCER1A,GRAP2 |
| Huntington's Disease Signaling | 0.00E+00 | 2.12E-02 | NaN | CACNA1B,CREB5,GRM1,HSPA1A/HSPA1B,HSPA6,NTRK1 |
| Chaperone Mediated Autophagy Signaling Pathway | 0.00E+00 | 1.56E-02 | 0.632 | CD3G,HSPA1A/HSPA1B,HSPA6,IL1B,INS,MMP21,MMP3,MTRNR2L4,PPP3R2,TRBV5-4 |
| DHCR24 Signaling Pathway | 0.00E+00 | 2.19E-02 | NaN | AGT,APOB,HPR |
| WNK Renal Signaling Pathway | 0.00E+00 | 2.83E-02 | NaN | AGT,INS,SLC12A1 |
| NRF2-mediated Oxidative Stress Response | 0.00E+00 | 2.53E-02 | NaN | ACTA1,CYP4Z1,DNAJB13,DNAJC5G,DNAJC6,FMO1 |
| Aryl Hydrocarbon Receptor Signaling | 0.00E+00 | 1.89E-02 | NaN | ALDH1L2,IL1B,TGFB2 |
| Mitochondrial Dysfunction | 0.00E+00 | 2.62E-02 | -1.667 | ATP1A3,ATP1A4,CACNA1B,CACNA1C,CREB5,GPX8,MAOB,PPP3R2,PTGES |
| Ceramide Signaling | 0.00E+00 | 2.20E-02 | NaN | NGFR,PPP2R2C |
| TR/RXR Activation | 0.00E+00 | 1.19E-02 | NaN | AKR1C1/AKR1C2 |
| 14-3-3-mediated Signaling | 0.00E+00 | 2.36E-02 | NaN | NOTUM,TUBB2B,TUBB4A |
| α-Adrenergic Signaling | 0.00E+00 | 2.75E-02 | NaN | ADCY2,ADCY5,ADRA2A |
| Fcγ Receptor-mediated Phagocytosis in Macrophages and Mono | 0.00E+00 | 1.06E-02 | NaN | ACTA1 |
| IL-8 Signaling | 0.00E+00 | 4.76E-03 | NaN | CXCL8 |
| IL-12 Signaling and Production in Macrophages | 0.00E+00 | 2.54E-02 | 0.816 | ADRA2A,APOB,COL1A2,IL23A,TGFB2,TNFRSF11A |
| Role of NFAT in Regulation of the Immune Response | 0.00E+00 | 5.79E-03 | -0.447 | CD3G,CD79A,FCER1A,PPP3R2,TRBV5-4,ZAP70 |
| FcγRIIB Signaling in B Lymphocytes | 0.00E+00 | 5.63E-03 | NaN | CACNA1B,CACNA1C,CD79A |
| CCR5 Signaling in Macrophages | 0.00E+00 | 1.00E-02 | NaN | CACNA1B,CACNA1C,CCL4,CD3G,TRBV5-4 |
| CD40 Signaling | 0.00E+00 | 1.49E-02 | NaN | FCER2 |
| Calcium-induced T Lymphocyte Apoptosis | 0.00E+00 | 8.70E-03 | NaN | CD3G,PPP3R2,TRBV5-4,ZAP70 |
| Cytotoxic T Lymphocyte-mediated Apoptosis of Target Cells | 0.00E+00 | 7.06E-03 | NaN | CD3G,HLA-G,TRBV5-4 |
| CD27 Signaling in Lymphocytes | 0.00E+00 | 1.75E-02 | NaN | MAP3K15 |
| IL-3 Signaling | 0.00E+00 | 1.27E-02 | NaN | PPP3R2 |
| fMLP Signaling in Neutrophils | 0.00E+00 | 7.63E-03 | NaN | PPP3R2 |
| CXCR4 Signaling | 0.00E+00 | 2.98E-02 | 1.342 | ADCY2,ADCY5,CXCR4,MYL11,MYL4 |
| Thrombopoietin Signaling | 0.00E+00 | 1.59E-02 | NaN | THPO |
| CTLA4 Signaling in Cytotoxic T Lymphocytes | 0.00E+00 | 1.15E-02 | -1.134 | CD3G,GRAP2,HLA-G,PPP2R2C,TGFB2,TRBV5-4,ZAP70 |
| T Helper Cell Differentiation | 0.00E+00 | 6.37E-03 | NaN | CD3G,NGFR,TRBV5-4 |
| CCR3 Signaling in Eosinophils | 0.00E+00 | 7.46E-03 | NaN | CCL26 |
| Oncostatin M Signaling | 0.00E+00 | 2.33E-02 | NaN | MMP3 |
| CD28 Signaling in T Helper Cells | 0.00E+00 | 9.67E-03 | NaN | CD3G,GRAP2,PPP3R2,TRBV5-4,ZAP70 |
| IL-15 Signaling | 0.00E+00 | 1.89E-03 | NaN | CXCL8 |
| Virus Entry via Endocytic Pathways | 0.00E+00 | 1.69E-02 | NaN | ACTA1,HLA-G |
| Dendritic Cell Maturation | 0.00E+00 | 2.01E-02 | 1.265 | CCR7,CD1A,CD3G,COL1A2,CREB5,HLA-G,IL1B,IL23A,IL32,NGFR,NOTUM,TRBV5-4 |
| Mechanisms of Viral Exit from Host Cells | 0.00E+00 | 2.44E-02 | NaN | ACTA1 |
| Melatonin Signaling | 0.00E+00 | 1.39E-02 | NaN | NOTUM |
| Endothelin-1 Signaling | 0.00E+00 | 2.06E-02 | 0 | ADCY2,ADCY5,EDN1,NOTUM |
| Relaxin Signaling | 0.00E+00 | 1.29E-02 | NaN | ADCY2,ADCY5 |
| Agrin Interactions at Neuromuscular Junction | 0.00E+00 | 2.90E-02 | NaN | ACTA1,ERBB4 |
| Renin-Angiotensin Signaling | 0.00E+00 | 2.48E-02 | NaN | ADCY2,ADCY5,AGT |
| Docosahexaenoic Acid (DHA) Signaling | 0.00E+00 | 2.63E-02 | NaN | IL1B |
| ICOS-ICOSL Signaling in T Helper Cells | 0.00E+00 | 1.18E-02 | NaN | CD3G,GRAP2,PLEKHA4,PPP3R2,TRBV5-4,ZAP70 |
| Molecular Mechanisms of Cancer | 0.00E+00 | 1.78E-02 | NaN | ADCY2,ADCY5,ARHGEF10,BMP6,BMPR1B,FZD6,ITGA2B,TGFB2 |
| Lipid Antigen Presentation by CD1 | 0.00E+00 | 7.23E-03 | NaN | CD1A,CD3G,TRBV5-4 |
| Cell Cycle Regulation by BTG Family Proteins | 0.00E+00 | 2.63E-02 | NaN | PPP2R2C |
| Mitotic Roles of Polo-Like Kinase | 0.00E+00 | 1.49E-02 | NaN | PPP2R2C |
| HGF Signaling | 0.00E+00 | 2.27E-02 | NaN | HGF,ITGA2B,MAP3K15 |
| Role of CHK Proteins in Cell Cycle Checkpoint Control | 0.00E+00 | 1.72E-02 | NaN | PPP2R2C |
| FLT3 Signaling in Hematopoietic Progenitor Cells | 0.00E+00 | 1.22E-02 | NaN | CREB5 |
| Cholecystokinin/Gastrin-mediated Signaling | 0.00E+00 | 8.40E-03 | NaN | IL1B |
| Human Embryonic Stem Cell Pluripotency | 0.00E+00 | 2.99E-02 | 1.633 | BMP6,BMPR1B,FZD6,NTF3,NTRK1,TGFB2 |
| DNA Methylation and Transcriptional Repression Signaling | 0.00E+00 | 2.04E-02 | NaN | FOXA2,TCF15 |
| ATM Signaling | 0.00E+00 | 2.00E-02 | NaN | CREB5,PPP2R2C |
| Androgen Signaling | 0.00E+00 | 1.18E-02 | NaN | CACNA1B,CACNA1C |
| Role of OCT4 in Mammalian Embryonic Stem Cell Pluripotency | 0.00E+00 | 2.17E-02 | NaN | FOXA2 |
| Growth Hormone Signaling | 0.00E+00 | 1.41E-02 | NaN | IGF2 |
| Prostate Cancer Signaling | 0.00E+00 | 8.77E-03 | NaN | CREB5 |
| Renal Cell Carcinoma Signaling | 0.00E+00 | 1.27E-02 | NaN | HGF |
| Type I Diabetes Mellitus Signaling | 0.00E+00 | 1.18E-02 | NaN | CD3G,HLA-G,IL1B,INS,NGFR,TRBV5-4 |
| Basal Cell Carcinoma Signaling | 0.00E+00 | 2.78E-02 | NaN | BMP6,FZD6 |
| Allograft Rejection Signaling | 0.00E+00 | 6.16E-03 | NaN | CD3G,HLA-G,TRBV5-4 |
| Glioma Signaling | 0.00E+00 | 8.00E-03 | NaN | IGF2 |
| Autoimmune Thyroid Disease Signaling | 0.00E+00 | 8.73E-03 | NaN | CD3G,HLA-G,TRBV5-4,TSHR |
| Graft-versus-Host Disease Signaling | 0.00E+00 | 8.97E-03 | NaN | CD3G,HLA-G,IL1B,TRBV5-4 |
| Type II Diabetes Mellitus Signaling | 0.00E+00 | 2.61E-02 | NaN | CACNA1B,CACNA1C,INS,NGFR |
| Chronic Myeloid Leukemia Signaling | 0.00E+00 | 1.44E-02 | -1 | FZD6,NOTUM,PPP3R2,TGFB2 |
| Production of Nitric Oxide and Reactive Oxygen Species in Macr | 0.00E+00 | 2.62E-02 | -1.342 | APOB,MAP3K15,NGFR,PPP1R14D,PPP2R2C |
| p70S6K Signaling | 0.00E+00 | 1.03E-02 | -1.633 | AGT,CD19,CD79A,F2R,NOTUM,PPP2R2C |
| ERK5 Signaling | 0.00E+00 | 2.70E-02 | NaN | CREB5,NTRK1 |
| Colorectal Cancer Metastasis Signaling | 0.00E+00 | 2.58E-02 | 0 | ADCY2,ADCY5,FZD6,MMP21,MMP3,PTGER1,TGFB2 |
| mTOR Signaling | 0.00E+00 | 9.35E-03 | NaN | INS,PPP2R2C |
| MYC Mediated Apoptosis Signaling | 0.00E+00 | 2.00E-02 | NaN | NGFR |
| Pancreatic Adenocarcinoma Signaling | 0.00E+00 | 7.94E-03 | NaN | TGFB2 |
| G Beta Gamma Signaling | 0.00E+00 | 2.33E-02 | NaN | ADCY2,CACNA1B,CACNA1C |
| G Protein Signaling Mediated by Tubby | 0.00E+00 | 8.55E-03 | 1 | CD3G,GRAP2,INS,TRBV5-4 |
| Communication between Innate and Adaptive Immune Cells | 0.00E+00 | 8.57E-03 | NaN | CCL4,CCR7,CD3G,CD79A,CXCL8,HLA-G,IL1B,TRBV5-4 |
| Sphingosine-1-phosphate Signaling | 0.00E+00 | 2.50E-02 | NaN | ADCY2,ADCY5,NOTUM |
| Systemic Lupus Erythematosus Signaling | 0.00E+00 | 5.62E-03 | NaN | BDKRB2,CD3G,CD79A,HLA-G,IL1B,TRBV5-4 |
| CDC42 Signaling | 0.00E+00 | 1.04E-02 | NaN | CD3G,HLA-G,ITGA2B,MYL11,MYL4,TRBV5-4 |
| ILK Signaling | 0.00E+00 | 2.99E-02 | -0.447 | ACTA1,CREB5,DSP,MYH2,MYL4,PPP2R2C |
| EIF2 Signaling | 0.00E+00 | 8.81E-03 | NaN | ACTA1,INS |
| Retinoic acid Mediated Apoptosis Signaling | 0.00E+00 | 1.67E-02 | NaN | CRABP1 |
| PAK Signaling | 0.00E+00 | 2.56E-02 | NaN | ITGA2B,MYL11,MYL4 |
| RAC Signaling | 0.00E+00 | 7.30E-03 | NaN | ITGA2B |
| RHOA Signaling | 0.00E+00 | 2.42E-02 | NaN | ACTA1,MYL11,MYL4 |
| Phospholipase C Signaling | 0.00E+00 | 1.16E-02 | 0.707 | ADCY2,ADCY5,ARHGEF10,CD3G,CD79A,CREB5,GRAP2,ITGA2B,MYL11,MYL4,PPP3R2,TRBV5-4,ZAP70 |
| Ovarian Cancer Signaling | 0.00E+00 | 1.90E-02 | NaN | EDN1,FZD6,LHCGR |
| HER-2 Signaling in Breast Cancer | 0.00E+00 | 8.81E-03 | NaN | AREG,FCER1A |
| Altered T Cell and B Cell Signaling in Rheumatoid Arthritis | 0.00E+00 | 5.32E-03 | NaN | CD3G,CD79A,IL1B,IL23A,TRBV5-4 |
| Regulation of eIF4 and p70S6K Signaling | 0.00E+00 | 1.10E-02 | NaN | ITGA2B,PPP2R2C |
| Glioma Invasiveness Signaling | 0.00E+00 | 1.37E-02 | NaN | F2R |
| B Cell Development | 0.00E+00 | 4.09E-03 | NaN | CD19,CD79A |
| IL-1 Signaling | 0.00E+00 | 2.08E-02 | NaN | ADCY2,ADCY5 |
| Glioblastoma Multiforme Signaling | 0.00E+00 | 1.75E-02 | NaN | FZD6,IGF2,NOTUM |
| Regulation of IL-2 Expression in Activated and Anergic T Lympho | 0.00E+00 | 1.08E-02 | 1.342 | CD3G,PPP3R2,TGFB2,TRBV5-4,ZAP70 |
| Granzyme A Signaling | 0.00E+00 | 2.67E-02 | NaN | F2R,IL1B |
| Role of WNT/GSK-3β Signaling in the Pathogenesis of Influenza | 0.00E+00 | 1.28E-02 | NaN | FZD6 |
| NUR77 Signaling in T Lymphocytes | 0.00E+00 | 7.80E-03 | NaN | CD3G,HLA-G,PPP3R2,TRBV5-4 |
| PKCθ Signaling in T Lymphocytes | 0.00E+00 | 1.43E-02 | -0.447 | CACNA1B,CACNA1C,CD3G,GRAP2,MAP3K15,PPP3R2,TRBV5-4,ZAP70 |
| Antiproliferative Role of TOB in T Cell Signaling | 0.00E+00 | 7.04E-03 | NaN | CD3G,TGFB2,TRBV5-4 |
| MSP-RON Signaling Pathway | 0.00E+00 | 1.72E-02 | NaN | ACTA1 |
| OX40 Signaling Pathway | 0.00E+00 | 6.28E-03 | NaN | CD3G,HLA-G,TRBV5-4 |
| PI3K Signaling in B Lymphocytes | 0.00E+00 | 8.46E-03 | -2 | CD19,CD79A,NOTUM,PLEKHA4,PPP3R2 |
| Cyclins and Cell Cycle Regulation | 0.00E+00 | 2.35E-02 | NaN | PPP2R2C,TGFB2 |
| IL-17A Signaling in Fibroblasts | 0.00E+00 | 2.63E-02 | NaN | CXCL5 |
| Role of JAK1 and JAK3 in γc Cytokine Signaling | 0.00E+00 | 1.45E-02 | NaN | CRLF2 |
| Actin Nucleation by ARP-WASP Complex | 0.00E+00 | 1.08E-02 | NaN | ITGA2B |
| Paxillin Signaling | 0.00E+00 | 1.87E-02 | NaN | ACTA1,ITGA2B |
| Signaling by Rho Family GTPases | 0.00E+00 | 3.00E-02 | 1.134 | ACTA1,ARHGEF10,CDH17,CDH2,CDH6,ITGA2B,MYL11,MYL4 |
| Telomerase Signaling | 0.00E+00 | 9.26E-03 | NaN | PPP2R2C |
| Mouse Embryonic Stem Cell Pluripotency | 0.00E+00 | 9.62E-03 | NaN | FZD6 |
| Hematopoiesis from Pluripotent Stem Cells | 0.00E+00 | 8.99E-03 | NaN | CD3G,CXCL8,IL11,TRBV5-4 |
| nNOS Signaling in Neurons | 0.00E+00 | 2.13E-02 | NaN | PPP3R2 |
| Ephrin B Signaling | 0.00E+00 | 1.39E-02 | NaN | CXCR4 |
| ERBB4 Signaling | 0.00E+00 | 1.47E-02 | NaN | ERBB4 |
| GDNF Family Ligand-Receptor Interactions | 0.00E+00 | 1.32E-02 | NaN | DOK7 |
| Retinoate Biosynthesis I | 0.00E+00 | 2.38E-02 | NaN | AKR1C1/AKR1C2 |

| | | | | |
|---|---|---|---|---|
| Thyroid Hormone Metabolism II (via Conjugation and/or Degrad | 0.00E+00 | 2.44E-02 | NaN | SULT1B1 |
| D-myo-inositol-5-phosphate Metabolism | 0.00E+00 | 2.04E-02 | NaN | CILP,PPP1R14D,PPP2R2C,PPP3R2 |
| Melatonin Degradation I | 0.00E+00 | 1.59E-02 | NaN | SULT1B1 |
| Pyridoxal 5'-phosphate Salvage Pathway | 0.00E+00 | 1.54E-02 | NaN | MAK |
| Phospholipases | 0.00E+00 | 1.45E-02 | NaN | NOTUM |
| Glutathione-mediated Detoxification | 0.00E+00 | 2.70E-02 | NaN | PTGES |
| D-myo-inositol (1,4,5,6)-Tetrakisphosphate Biosynthesis | 0.00E+00 | 2.22E-02 | NaN | CILP,PPP1R14D,PPP2R2C,PPP3R2 |
| Nicotine Degradation II | 0.00E+00 | 2.99E-02 | NaN | FMO1,INMT |
| Superpathway of Inositol Phosphate Compounds | 0.00E+00 | 1.71E-02 | NaN | CILP,PPP1R14D,PPP2R2C,PPP3R2 |
| Serotonin Degradation | 0.00E+00 | 2.78E-02 | NaN | MAOB,SULT1B1 |
| D-myo-inositol (3,4,5,6)-tetrakisphosphate Biosynthesis | 0.00E+00 | 2.22E-02 | NaN | CILP,PPP1R14D,PPP2R2C,PPP3R2 |
| 3-phosphoinositide Degradation | 0.00E+00 | 2.09E-02 | NaN | CILP,PPP1R14D,PPP2R2C,PPP3R2 |
| 3-phosphoinositide Biosynthesis | 0.00E+00 | 1.94E-02 | NaN | CILP,PPP1R14D,PPP2R2C,PPP3R2 |
| Triacylglycerol Biosynthesis | 0.00E+00 | 1.82E-02 | NaN | DGAT2L6 |
| Superpathway of Melatonin Degradation | 0.00E+00 | 2.94E-02 | NaN | MAOB,SULT1B1 |
| Stearate Biosynthesis I (Animals) | 0.00E+00 | 1.45E-02 | NaN | THEM5 |
| Noradrenaline and Adrenaline Degradation | 0.00E+00 | 2.70E-02 | NaN | MAOB |
| Oxidative Ethanol Degradation III | 0.00E+00 | 1.75E-02 | NaN | CYP2W1 |
| Superpathway of Methionine Degradation | 0.00E+00 | 2.38E-02 | NaN | PRMT8 |
| Antioxidant Action of Vitamin C | 0.00E+00 | 8.77E-03 | NaN | NOTUM |
| Gαq Signaling | 0.00E+00 | 1.76E-02 | NaN | GRM1,PPP3R2,RGS4 |
| Regulation of Cellular Mechanics by Calpain Protease | 0.00E+00 | 1.11E-02 | NaN | ITGA2B |
| TEC Kinase Signaling | 0.00E+00 | 1.04E-02 | NaN | ACTA1,BMX,CD3G,FCER1A,ITGA2B,TRBV5-4 |
| UVA-Induced MAPK Signaling | 0.00E+00 | 1.02E-02 | NaN | NOTUM |
| Adipogenesis pathway | 0.00E+00 | 2.88E-02 | -1 | BMPR1B,FGF1,FZD6,RUNX1T1 |
| HIPPO signaling | 0.00E+00 | 2.33E-02 | NaN | PPP1R14D,PPP2R2C |
| PCP (Planar Cell Polarity) Pathway | 0.00E+00 | 1.67E-02 | NaN | FZD6 |
| Toll-like Receptor Signaling | 0.00E+00 | 1.28E-02 | NaN | IL1B |
| Cell Cycle: G1/S Checkpoint Regulation | 0.00E+00 | 1.47E-02 | NaN | TGFB2 |
| ERK/MAPK Signaling | 0.00E+00 | 1.86E-02 | NaN | CREB5,ITGA2B,PPP1R14D,PPP2R2C |
| SAPK/JNK Signaling | 0.00E+00 | 3.98E-03 | NaN | CD3G,TRBV5-4 |
| PI3K/AKT Signaling | 0.00E+00 | 2.00E-02 | NaN | IL1R2,IL22RA2,ITGA2B,PPP2R2C |
| IL-4 Signaling | 0.00E+00 | 1.56E-02 | 0.333 | CD3G,COL1A2,COL27A1,COL6A5,COL8A1,CREB5,FCER2,TGFB2,TRBV5-4 |
| Antigen Presentation Pathway | 0.00E+00 | 2.56E-02 | NaN | HLA-G |
| B Cell Receptor Signaling | 0.00E+00 | 7.87E-03 | -1.342 | CD19,CD79A,CREB5,MAP3K15,PPP3R2 |
| Insulin Receptor Signaling | 0.00E+00 | 1.43E-02 | NaN | INS,PPP1R14D |
| Chemokine Signaling | 0.00E+00 | 2.47E-02 | NaN | CCL4,CXCR4 |
| Integrin Signaling | 0.00E+00 | 9.43E-03 | NaN | ACTA1,ITGA2B |
| Death Receptor Signaling | 0.00E+00 | 1.04E-02 | NaN | ACTA1 |
| IGF-1 Signaling | 0.00E+00 | 9.52E-03 | NaN | CCN2 |
| Notch Signaling | 0.00E+00 | 2.63E-02 | NaN | MAG |
| NF-κB Signaling | 0.00E+00 | 1.75E-02 | 1.414 | BMPR1B,CD3G,IL1B,IL1R2,INS,NGFR,NTRK1,TNFRSF11A,TRBV5-4,ZAP70 |
| VEGF Signaling | 0.00E+00 | 1.01E-02 | NaN | ACTA1 |
| Hypoxia Signaling in the Cardiovascular System | 0.00E+00 | 2.63E-02 | NaN | CREB5,EDN1 |
| T Cell Receptor Signaling | 0.00E+00 | 9.74E-03 | 0.816 | CD3G,GRAP2,HLA-G,PPP3R2,TRBV5-4,ZAP70 |
| Phagosome Maturation | 0.00E+00 | 1.84E-02 | NaN | HLA-G,TUBB2B,TUBB4A |
| Macropinocytosis Signaling | 0.00E+00 | 2.63E-02 | NaN | HGF,INS |
| Cancer Drug Resistance By Drug Efflux | 0.00E+00 | 1.72E-02 | NaN | FOXG1 |
| Sumoylation Pathway | 0.00E+00 | 9.71E-03 | NaN | ISG20 |
| Th1 and Th2 Activation Pathway | 0.00E+00 | 1.74E-02 | NaN | CD3G,CRLF2,CXCR4 |
| Th1 Pathway | 0.00E+00 | 8.20E-03 | NaN | CD3G |
| Th2 Pathway | 0.00E+00 | 2.19E-02 | NaN | CD3G,CRLF2,CXCR4 |
| IL-7 Signaling Pathway | 0.00E+00 | 2.56E-02 | NaN | FOXG1,HGF |
| Sirtuin Signaling Pathway | 0.00E+00 | 1.37E-02 | NaN | ABCA1,CXCL8,LDHC,OTC |
| Th17 Activation Pathway | 0.00E+00 | 1.03E-02 | NaN | CCL20,CD3G,IL1B,IL23A,TRBV5-4 |
| SPINK1 Pancreatic Cancer Pathway | 0.00E+00 | 1.67E-02 | NaN | CPA4 |
| Endocannabinoid Developing Neuron Pathway | 0.00E+00 | 2.36E-02 | NaN | ADCY2,ADCY5,CREB5 |
| Apelin Pancreas Signaling Pathway | 0.00E+00 | 2.17E-02 | NaN | INS |
| Apelin Cardiomyocyte Signaling Pathway | 0.00E+00 | 3.03E-02 | NaN | MYL11,MYL4,NOTUM |
| Apelin Endothelial Signaling Pathway | 0.00E+00 | 1.42E-02 | NaN | ADCY2,ADCY5 |
| BAG2 Signaling Pathway | 0.00E+00 | 2.38E-02 | NaN | HSPA1A/HSPA1B,HSPA6 |
| T Cell Exhaustion Signaling Pathway | 0.00E+00 | 8.82E-03 | NaN | CD3G,HLA-G,PPP2R2C,TRBV5-4,ZAP70 |
| Systemic Lupus Erythematosus In T Cell Signaling Pathway | 0.00E+00 | 1.09E-02 | 0.447 | CD3G,CREB5,HLA-G,IL23A,PPP2R2C,PPP3R2,TRBV5-4 |
| Systemic Lupus Erythematosus In B Cell Signaling Pathway | 0.00E+00 | 1.52E-02 | 0.302 | CD19,CD79A,CXCL8,FOXG1,IL11,IL17C,IL17D,IL1B,ISG20,PPP3R2,TGFB2 |
| Senescence Pathway | 0.00E+00 | 2.01E-02 | -0.447 | CACNA1B,CACNA1C,CXCL8,PPP2R2C,PPP3R2,TGFB2 |
| Inhibition of ARE-Mediated mRNA Degradation Pathway | 0.00E+00 | 1.23E-02 | NaN | NGFR,PPP2R2C |
| HOTAIR Regulatory Pathway | 0.00E+00 | 2.45E-02 | 0 | COL1A2,MMP21,MMP3,TWIST1 |
| Necroptosis Signaling Pathway | 0.00E+00 | 1.28E-02 | NaN | NGFR,PPP3R2 |
| Xenobiotic Metabolism General Signaling Pathway | 0.00E+00 | 6.99E-03 | NaN | MAP3K15 |
| Xenobiotic Metabolism AHR Signaling Pathway | 0.00E+00 | 2.30E-02 | NaN | ALDH1L2,IL1B |
| Insulin Secretion Signaling Pathway | 0.00E+00 | 2.94E-02 | 0.378 | ADCY2,ADCY5,CACNA1B,CACNA1C,CREB5,INS,NOTUM,PCSK2 |
| Semaphorin Neuronal Repulsive Signaling Pathway | 0.00E+00 | 2.67E-02 | NaN | ITGA2B,MYL11,MYL4,PRKG1 |
| Regulation Of The Epithelial Mesenchymal Transition In Develop | 0.00E+00 | 2.30E-02 | NaN | FZD6,TWIST1 |
| Kinetochore Metaphase Signaling Pathway | 0.00E+00 | 1.80E-02 | NaN | DNAH3,PPP1R14D |
| Coronavirus Pathogenesis Pathway | 0.00E+00 | 1.47E-02 | NaN | AGT,CXCL8,IL1B |
| MSP-RON Signaling In Cancer Cells Pathway | 0.00E+00 | 7.14E-03 | NaN | CREB5 |
| MSP-RON Signaling In Macrophages Pathway | 0.00E+00 | 8.40E-03 | NaN | CREB5 |
| GM-CSF Signaling | 0.00E+00 | 1.43E-02 | NaN | PPP3R2 |
| Eicosanoid Signaling | 0.00E+00 | 2.86E-02 | NaN | PTGER1,PTGES |
| Ephrin Receptor Signaling | 0.00E+00 | 2.97E-02 | NaN | CREB5,CXCR4,EPHA10,FGF1,ITGA2B,SORBS1 |
| Ferroptosis Signaling Pathway | 0.00E+00 | 2.27E-02 | NaN | ABCA1,CHAC1,SLC7A11 |
| Role of MAPK Signaling in Inhibiting the Pathogenesis of Influenz | 0.00E+00 | 2.53E-02 | NaN | CXCL8,IL1B |

IPA Microglia GCSi FC2 adj0.05 920

| | Ingenuity Canonical Pathways | -log(p-value) | Ratio | z-score | Molecules |
|---|---|---|---|---|---|
| 1 | S100 Family Signaling Pathway | 1.11E+01 | 6.48E-02 | 0.283 | ACKR3,ADGRB2,ADGRD2,ADGRE1,ADGRG1,ADGRG6,ADGRL3,ADORA3,AREG,CACNA1G,CACNA2D3,CCR7,EDNRB,EREG,ESR2,F2RL3,FCGR1A,FCGR1BP,FCGR2B,FCGR2C,FFAR4,GLP2R,GPR152,GPR160,GPR37,GPR4,HTR4,IL1B,LHCGR,LPAR3,MAPK15,MMP21,MMP3,MMP8,NOTUM,NPY1R,NTF3,NTSR1,P2RY12,P2RY13,PLA2G1B,RGR,S100Z,S1PR3,SSTR2,TRPV5,WNT1,WNT10B,WNT3,WNT5B |
| 2 | Hepatic Fibrosis / Hepatic Stellate Cell Activatio | 9.52E+00 | 1.13E-01 | NaN | CCR7,COL13A1,COL5A1,COL5A3,COL6A5,COL7A1,CXCL3,EDNRB,FGF2,HGF,IGF1,IGF2,IGFBP3,IL10,IL18RAP,IL1B,MET,MYL4,MYL9,NGFR,PDGFRB,SERPINE1 |
| 3 | CREB Signaling in Neurons | 8.69E+00 | 6.43E-02 | -0.16 | ACKR3,ADCY10,ADGRB2,ADGRD2,ADGRE1,ADGRG1,ADGRG6,ADGRL3,ADORA3,BMP6,CACNA1G,CACNA2D3,CCR7,EDNRB,F2RL3,FFAR4,FGF2,GLP2R,GPR152,GPR160,GPR37,GPR4,GRIA2,GRIN2A,HGF,HTR4,IGF1,LHCGR,LPAR3,NGFR,NOTUM,NPY1R,NTSR1,P2RY12,P2RY13,PDGFRB,RGR,S1PR3,SSTR2 |
| 4 | Phagosome Formation | 8.52E+00 | 6.04E-02 | -2.16 | ACKR3,ADGRB2,ADGRD2,ADGRE1,ADGRG1,ADGRG6,ADGRL3,ADORA3,CCR7,CR1,EDNRB,F2RL3,FCER1A,FCGR1A,FCGR1BP,FCGR2B,FCGR2C,FFAR4,GLP2R,GPR152,GPR160,GPR37,GPR4,HTR4,ITGA2,ITGA9,LHCGR,LPAR3,MAPK15,MYL4,MYL9,NPY1R,NTSR1,P2RY12,P2RY13,PLA2G1B,PLAAT4,PLD4,RGR,S1PR3,SSTR2,TLR10 |
| 5 | Granulocyte Adhesion and Diapedesis | 6.73E+00 | 9.52E-02 | NaN | ACKR3,CCL23,CCL24,CCL7,CCR7,CLDN4,CLDN5,CXCL2,CXCL3,CXCL5,CXCR4,IL18RAP,IL1B,MMP21,MMP3,MMP8,NGFR,SELL |
| 6 | Agranulocyte Adhesion and Diapedesis | 6.72E+00 | 9.05E-02 | NaN | ACKR3,CCL23,CCL24,CCL7,CCR7,CLDN4,CLDN5,CXCL2,CXCL3,CXCL5,CXCR4,IL1B,MMP21,MMP3,MMP8,MYL4,MYL9,PODXL,SELL |
| 7 | Pulmonary Fibrosis Idiopathic Signaling Pathwa | 6.62E+00 | 7.36E-02 | 3.128 | AREG,CAV1,CCN4,COL13A1,COL5A1,COL5A3,COL6A5,COL7A1,EFNB2,FGF2,FOXG1,IL11,IL1B,ITGA2,MMP21,MMP3,MMP8,NOTCH3,PDGFRB,SERPINE1,WNT1,WNT10B,WNT3,WNT5B |
| 8 | Breast Cancer Regulation by Stathmin1 | 5.99E+00 | 5.56E-02 | -0.522 | ACKR3,ADGRB2,ADGRD2,ADGRE1,ADGRG1,ADGRG6,ADGRL3,ADORA3,BMP6,CCR7,EDNRB,F2RL3,FFAR4,FGF2,GLP2R,GPR152,GPR160,GPR37,GPR4,HGF,HTR4,IGF1,LHCGR,LPAR3,NPY1R,NTSR1,P2RY12,P2RY13,PPM1J,RGR,S1PR3,SSTR2,TUBA1A |
| 9 | G-Protein Coupled Receptor Signaling | 5.66E+00 | 5.12E-02 | -1.667 | ACKR3,ADCY10,ADGRB2,ADGRD2,ADGRE1,ADGRG1,ADGRG6,ADGRL3,ADORA3,CARD10,CCR7,EDNRB,F2RL3,FFAR4,FOXG1,GLP2R,GPR152,GPR160,GPR37,GPR4,HTR4,KCNH2,KCNN3,LHCGR,LPAR3,MYL4,MYL9,NPY1R,NTSR1,P2RY12,P2RY13,PDE4C,PDE6B,RGR,S1PR3,SSTR2 |
| 10 | Gustation Pathway | 4.98E+00 | 7.88E-02 | -1.291 | ADCY10,CACNA1G,CACNA2D3,CALHM1,FFAR4,GABRA3,GABRE,GABRQ,GPR37,KCNH2,KCNN3,LRRC8E,NOTUM,P2RY12,P2RY13,SCN7A |
| | Pathogen Induced Cytokine Storm Signaling Pat | 4.61E+00 | 5.93E-02 | 1.706 | CCL23,CCL24,CCL7,CLCF1,COL13A1,COL5A1,COL5A3,COL6A5,COL7A1,CXCL2,CXCL3,CXCL5,CXCR4,EOMES,IL10,IL11,IL18RAP,IL1B,MAPK15,NGFR,TLR10,TNFSF8 |
| | Axonal Guidance Signaling | 4.21E+00 | 5.10E-02 | NaN | ADAMTS20,ADAMTS9,BMP6,CXCR4,EFNB2,EFNB3,EPHA10,IGF1,ITGA2,ITGA9,MET,MMP21,MMP3,MMP8,MYL4,MYL9,NGFR,NOTUM,NTF3,PLXNB1,SHANK2,TUBA1A,WNT1,WNT10B,WNT3,WNT5B |
| | Activin Inhibin Signaling Pathway | 4.05E+00 | 6.91E-02 | 1.291 | COL5A3,CXCR4,FOSB,FOSL2,IL11,IL18RAP,IL1B,INHBA,MAF,MAPK15,NGFR,PMEPA1,SERPINE1,TDGF1,TLR10 |
| | Role of Osteoblasts, Osteoclasts and Chondrocy | 3.81E+00 | 6.58E-02 | NaN | BMP6,DKK3,IGF1,IL10,IL11,IL18RAP,IL1B,MMP3,MMP8,NAIP,NGFR,WNT1,WNT10B,WNT3,WNT5B |
| | Role Of Osteoclasts In Rheumatoid Arthritis Sig | 3.78E+00 | 5.83E-02 | 0 | COL13A1,COL5A1,COL5A3,COL6A5,COL7A1,FCGR1A,FCGR1BP,FCGR2B,FCGR2C,FOXG1,IL10,IL18RAP,IL1B,MMP21,MMP3,MMP8,NAIP,NGFR |
| | Neurovascular Coupling Signaling Pathway | 3.73E+00 | 6.47E-02 | 0.775 | CACNA1G,CACNA2D3,GABRA3,GABRE,GABRQ,GRIA2,GRIN2A,KCNJ15,KCNJ5,KCNMB1,KCNN3,NPR1,PLA2G1B,PLAAT4,SLC1A2 |
| | Myelination Signaling Pathway | 3.48E+00 | 5.50E-02 | 1.886 | ADGRG6,BMP6,FGF2,GPR37,ID4,IGF1,IRS1,ITGA2,LAMA5,MYRF,NGFR,NTF3,PDGFRB,PLP1,WNT1,WNT10B,WNT3,WNT5B |
| | Tumor Microenvironment Pathway | 3.24E+00 | 6.70E-02 | 1.155 | CXCR4,FGF2,FOXG1,HGF,IGF1,IGF2,IL10,IL1B,MMP21,MMP3,MMP8,TNC |
| | GABA Receptor Signaling | 3.19E+00 | 7.58E-02 | NaN | ADCY10,CACNA1G,CACNA2D3,GABRA3,GABRE,GABRQ,GPR37,KCNH2,KCNN3,SLC6A13 |
| | FAK Signaling | 3.12E+00 | 3.75E-02 | 0.16 | ACKR3,ADGRB2,ADGRD2,ADGRE1,ADGRG1,ADGRG6,ADGRL3,ADORA3,CCR7,COL5A3,EDNRB,EFNB2,EFNB3,ELF5,F2RL3,FFAR4,GLP2R,GPR152,GPR160,GPR37,GPR4,HTR4,IL18RAP,IL31RA,ITGA2,ITGA9,LHCGR,LPAR3,MET,NPY1R,NTSR1,P2RY12,P2RY13,PDGFRB,RGR,S1PR3,SSTR2,TRAJ23,TRBV5-4 |
| | Coagulation System | 3.04E+00 | 1.43E-01 | 0.447 | F13A1,F5,SERPIND1,SERPINE1,THBD |
| | HOTAIR Regulatory Pathway | 3.04E+00 | 6.75E-02 | 2.111 | ESR2,JAM2,MET,MMP21,MMP3,MMP8,TWIST1,WNT1,WNT10B,WNT3,WNT5B |
| | Wound Healing Signaling Pathway | 2.87E+00 | 5.56E-02 | 1.069 | CLCF1,COL13A1,COL5A1,COL5A3,COL6A5,COL7A1,FGF2,IL11,IL18RAP,IL1B,LAMA5,MMP8,NGFR,TNFSF8 |
| | TREM1 Signaling | 2.84E+00 | 9.09E-02 | -0.378 | CASP5,CCL7,CXCL3,FCGR2B,IL10,IL1B,TLR10 |
| | Inhibition of Matrix Metalloproteases | 2.82E+00 | 1.28E-01 | -1 | HSPG2,MMP21,MMP3,MMP8,TFPI2 |
| | Serotonin Receptor Signaling | 2.80E+00 | 4.48E-02 | -1.091 | ADCY10,CACNA1G,CACNA2D3,CAV1,CAV2,CCR7,CD38,F13A1,GPR37,HTR4,IL1B,KCNH2,KCNJ5,KCNN3,NOTUM,PLA2G1B,PLAAT4,PLD4,SHANK3,TEAD1,TRPM8 |
| | Cellular Effects of Sildenafil (Viagra) | 2.77E+00 | 6.67E-02 | NaN | ADCY10,CACNA2D3,GPR37,KCNH2,KCNN3,MYL4,MYL9,NOTUM,NPR1,PDE4C |
| | Neuroinflammation Signaling Pathway | 2.77E+00 | 5.05E-02 | 1 | GABRA3,GABRE,GABRQ,GRIN2A,IL10,IL1B,KCNJ5,MAPK15,MMP3,NAIP,NTF3,PLA2G1B,SLC1A2,SLC6A13,TLR10,WNT1 |
| | Cardiac Hypertrophy Signaling (Enhanced) | 2.71E+00 | 4.24E-02 | 1.606 | ADCY10,CACNA2D3,CLCF1,EDNRB,FGF2,IGF1,IL11,IL18RAP,IL1B,IL31RA,ITGA2,ITGA9,NGFR,NKX2-5,NOTUM,NPR1,PDE4C,PDE6B,TNFSF8,WNT1,WNT10B,WNT3,WNT5B |
| | Osteoarthritis Pathway | 2.67E+00 | 5.51E-02 | 0.707 | CASP5,CCN4,FGF2,HTRA1,IL18RAP,IL1B,ITGA2,ITGA9,MMP3,NKX3-2,S1PR3,SLC39A8,SOX9 |
| | STAT3 Pathway | 2.54E+00 | 6.67E-02 | 1.414 | BMP6,FGF2,HGF,IGF1,IL18RAP,IL1B,IL31RA,NGFR,PDGFRB |
| | Transcriptional Regulatory Network in Embryon | 2.49E+00 | 6.10E-02 | 1.265 | EOMES,FGF2,GATA6,IGF1,IGF2,INHBA,WNT1,WNT10B,WNT3,WNT5B |
| | Role of IL-17F in Allergic Inflammatory Airway D | 2.46E+00 | 1.06E-01 | NaN | CCL7,CXCL5,IGF1,IL11,IL1B |
| | Regulation of the Epithelial-Mesenchymal Trans | 2.42E+00 | 5.64E-02 | NaN | FGF2,HGF,HMGA2,MET,NOTCH3,PDGFRB,TWIST1,WNT1,WNT10B,WNT3,WNT5B |
| | Pulmonary Healing Signaling Pathway | 2.36E+00 | 5.53E-02 | 3.317 | ACKR3,CXCR4,MMP21,MMP3,MMP8,NGFR,NOTCH3,WNT1,WNT10B,WNT3,WNT5B |
| | Extrinsic Prothrombin Activation Pathway | 2.33E+00 | 1.88E-01 | NaN | F13A1,F5,THBD |
| | Human Embryonic Stem Cell Pluripotency | 2.32E+00 | 5.47E-02 | 2.714 | BMP6,FGF2,ID4,INHBA,NTF3,PDGFRB,TDGF1,WNT1,WNT10B,WNT3,WNT5B |
| | CDX Gastrointestinal Cancer Signaling Pathway | 2.31E+00 | 5.45E-02 | -0.905 | AXIN2,BMP6,CLCF1,GATA6,IL11,IL1B,TNFSF8,WNT1,WNT10B,WNT3,WNT5B |
| | Role Of Osteoblasts In Rheumatoid Arthritis Sig | 2.12E+00 | 4.92E-02 | 0 | CLCF1,IGF1,IL11,IL1B,MMP21,MMP3,MMP8,TNFSF8,WNT1,WNT10B,WNT3,WNT5B |
| | Atherosclerosis Signaling | 2.06E+00 | 6.02E-02 | NaN | ALB,ALOX15B,COL5A3,CXCR4,IL1B,MMP3,PLA2G1B,PLAAT4 |
| | Inflammasome pathway | 2.05E+00 | 1.50E-01 | NaN | CASP5,IL1B,NAIP |
| | Chondroitin Sulfate Biosynthesis | 2.04E+00 | 8.47E-02 | 2.236 | B3GAT1,CHST1,CHST3,HS3ST3B1,SULT1B1 |
| | SNARE Signaling Pathway | 2.00E+00 | 5.88E-02 | 0.707 | ADCY10,MYL4,MYL9,SNCB,STX1A,SYN1,SYN3,SYT9 |
| | Dermatan Sulfate Biosynthesis | 1.98E+00 | 8.20E-02 | 2.236 | B3GAT1,CHST1,CHST3,HS3ST3B1,SULT1B1 |
| | Heparan Sulfate Biosynthesis | 1.93E+00 | 6.90E-02 | 2.449 | B3GAT1,CHST1,CHST3,HS3ST3B1,NOTUM,SULT1B1 |
| | Regulation Of The Epithelial Mesenchymal Tran | 1.93E+00 | 6.90E-02 | 2.449 | AXIN2,TWIST1,WNT1,WNT10B,WNT3,WNT5B |
| | Histidine Degradation III | 1.92E+00 | 2.50E-01 | NaN | AMDHD1,FTCD |
| | Intrinsic Prothrombin Activation Pathway | 1.89E+00 | 9.52E-02 | NaN | COL5A3,F13A1,F5,THBD |
| | CDK5 Signaling | 1.89E+00 | 6.09E-02 | 0.816 | ADCY10,FOSB,ITGA2,LAMA5,MAPK15,NGFR,PPM1J |
| | Role of Macrophages, Fibroblasts and Endotheli | 1.83E+00 | 4.22E-02 | NaN | DKK3,FCGR1A,FGF2,IL10,IL18RAP,IL1B,MMP3,NGFR,NOTUM,TLR10,WNT1,WNT10B,WNT3,WNT5B |
| | Airway Pathology in Chronic Obstructive Pulmo | 1.83E+00 | 5.93E-02 | NaN | CLCF1,CXCL3,FGF2,IL11,IL1B,MMP8,TNFSF8 |
| | Pyroptosis Signaling Pathway | 1.79E+00 | 6.45E-02 | -0.816 | CASP5,GBP1,IL1B,NAIP,NGFR,TLR10 |
| | HIF1α Signaling | 1.79E+00 | 4.81E-02 | 1.897 | BMP6,FGF2,HGF,IGF1,IGF2,MET,MMP21,MMP3,MMP8,SERPINE1 |
| | Endocannabinoid Neuronal Synapse Pathway | 1.78E+00 | 5.37E-02 | 1.414 | ADCY10,CACNA1G,CACNA2D3,GRIA2,GRIN2A,KCNJ5,MAPK15,NOTUM |
| | Factors Promoting Cardiogenesis in Vertebrates | 1.72E+00 | 5.23E-02 | 2.121 | BMP6,NKX2-5,NOTUM,TDGF1,WNT1,WNT10B,WNT3,WNT5B |
| | Dermatan Sulfate Biosynthesis (Late Stages) | 1.69E+00 | 8.33E-02 | 2 | CHST1,CHST3,HS3ST3B1,SULT1B1 |
| | Basal Cell Carcinoma Signaling | 1.69E+00 | 6.94E-02 | 2 | BMP6,WNT1,WNT10B,WNT3,WNT5B |
| | NAD Salvage Pathway II | 1.69E+00 | 1.11E-01 | NaN | NMNAT2,NT5E,PXYLP1 |
| | Apelin Liver Signaling Pathway | 1.69E+00 | 1.11E-01 | NaN | COL5A3,IRS1,PDGFRB |
| | Synaptogenesis Signaling Pathway | 1.67E+00 | 4.13E-02 | 1.387 | ADCY10,CDH13,EFNB2,EFNB3,EPHA10,GRIA2,GRIN2A,NLGN2,SNCB,STX1A,SYN1,SYN3,SYT9 |
| | Neuropathic Pain Signaling In Dorsal Horn Neur | 1.64E+00 | 5.94E-02 | 2.449 | GPR37,GRIA2,GRIN2A,KCNH2,KCNN3,NOTUM |
| | Caveolar-mediated Endocytosis Signaling | 1.62E+00 | 6.67E-02 | NaN | ALB,CAV1,FLNC,ITGA2,ITGA9 |
| | Hepatic Fibrosis Signaling Pathway | 1.62E+00 | 3.78E-02 | 2.673 | CACNA2D3,COL5A3,FGF2,IL18RAP,IL1B,ITGA2,ITGA9,MYL4,MYL9,NGFR,PDGFRB,SERPINE1,WNT1,WNT10B,WNT3,WNT5B |
| | Chondroitin Sulfate Biosynthesis (Late Stages) | 1.61E+00 | 7.84E-02 | 2 | CHST1,CHST3,HS3ST3B1,SULT1B1 |
| | Leukocyte Extravasation Signaling | 1.59E+00 | 4.66E-02 | 1.633 | CLDN4,CLDN5,CXCR4,EDIL3,ITGA2,JAM2,MMP21,MMP3,MMP8 |
| | Endothelin-1 Signaling | 1.57E+00 | 4.64E-02 | -0.707 | ADCY10,CASP5,EDNRB,MAPK15,NOTUM,NPR1,PLA2G1B,PLAAT4,PLD4 |
| | Synaptic Long Term Depression | 1.52E+00 | 4.55E-02 | -0.333 | CACNA1G,CACNA2D3,GRIA2,IGF1,NOTUM,NPR1,PLA2G1B,PLAAT4,PPM1J |
| | Heparan Sulfate Biosynthesis (Late Stages) | 1.52E+00 | 6.25E-02 | 2.236 | CHST1,CHST3,HS3ST3B1,NOTUM,SULT1B1 |
| | Adrenomedullin signaling pathway | 1.51E+00 | 4.52E-02 | 2.333 | ADCY10,GPR37,IL1B,KCNH2,KCNN3,MAPK15,NOTUM,NPR1,TFAP2C |
| | NAD biosynthesis II (from tryptophan) | 1.50E+00 | 1.54E-01 | NaN | NMNAT2,TDO2 |
| | Gαi Signaling | 1.47E+00 | 5.00E-02 | -0.816 | ADCY10,ADORA3,CAV1,NPY1R,P2RY12,P2RY13,S1PR3 |
| | Glioblastoma Multiforme Signaling | 1.47E+00 | 4.68E-02 | 2.121 | IGF1,IGF2,NOTUM,PDGFRB,WNT1,WNT10B,WNT3,WNT5B |
| | cAMP-mediated signaling | 1.46E+00 | 4.24E-02 | -2.53 | ADCY10,ADORA3,HTR4,LHCGR,NPY1R,P2RY12,P2RY13,PDE4C,PDE6B,S1PR3 |
| | Role Of Chondrocytes In Rheumatoid Arthritis S | 1.46E+00 | 4.96E-02 | 1.89 | CXCR4,IL18RAP,IL1B,MMP21,MMP3,MMP8,NGFR |
| | Role of IL-17A in Psoriasis | 1.44E+00 | 1.43E-01 | NaN | CXCL3,CXCL5 |
| | WNT/β-catenin Signaling | 1.43E+00 | 4.60E-02 | 0.707 | AXIN2,DKK3,PPM1J,SOX9,WNT1,WNT10B,WNT3,WNT5B |
| | Amyotrophic Lateral Sclerosis Signaling | 1.38E+00 | 5.17E-02 | 1.342 | CACNA2D3,GRIA2,GRIN2A,IGF1,NAIP,SLC1A2 |
| | PCP (Planar Cell Polarity) Pathway | 1.38E+00 | 6.67E-02 | 2 | WNT1,WNT10B,WNT3,WNT5B |
| | Tight Junction Signaling | 1.37E+00 | 4.47E-02 | NaN | CLDN4,CLDN5,JAM2,MYL4,MYL9,NGFR,PPM1J,STX1A |
| | NAD Signaling Pathway | 1.32E+00 | 4.64E-02 | 0.378 | BMP6,CD38,FGF2,HGF,IGF1,NMNAT2,NT5E |
| | Isoleucine Degradation I | 1.29E+00 | 1.18E-01 | NaN | ACAT2,SDS |
| | IL-10 Signaling | 1.28E+00 | 4.55E-02 | -1.134 | CCN4,FCGR2B,FCGR2C,IL10,IL18RAP,IL1B,MAF |
| | LXR/RXR Activation | 1.28E+00 | 4.88E-02 | -1.633 | ALB,CCL7,CETP,IL18RAP,IL1B,NGFR |
| | IL-17 Signaling | 1.28E+00 | 4.28E-02 | 1.414 | CLCF1,CXCL3,CXCL5,IL11,IL1B,MMP3,RGS13,TNFSF8 |
| | Role of NANOG in Mammalian Embryonic Stem | 1.27E+00 | 4.84E-02 | NaN | BMP6,GATA6,WNT1,WNT10B,WNT3,WNT5B |
| | eNOS Signaling | 1.26E+00 | 4.49E-02 | -0.447 | ADCY10,AQP2,AQP3,CAV1,CHRNA5,ESR2,LPAR3 |
| | Glutamate Receptor Signaling | 1.25E+00 | 6.06E-02 | NaN | GRIA2,GRIN2A,SLC1A2,SLC1A6 |
| | Differential Regulation of Cytokine Production i | 1.24E+00 | 1.11E-01 | NaN | IL10,IL1B |
| | Hepatic Cholestasis | 1.23E+00 | 4.19E-02 | NaN | ADCY10,CETP,CLCF1,IL11,IL18RAP,IL1B,NGFR,TNFSF8 |
| | GP6 Signaling Pathway | 1.23E+00 | 4.72E-02 | 0.816 | COL13A1,COL5A1,COL5A3,COL6A5,COL7A1,LAMA5 |
| | Regulation Of The Epithelial Mesenchymal Tran | 1.22E+00 | 4.17E-02 | 1.134 | FGF2,HGF,HMGA2,MET,NGFR,PDGFRB,TNFSF8,TWIST1 |
| | Phospholipases | 1.20E+00 | 5.80E-02 | -1 | NOTUM,PLA2G1B,PLAAT4,PLD4 |
| | L-serine Degradation | 1.19E+00 | 3.33E-01 | NaN | SDS |
| | Gap Junction Signaling | 1.16E+00 | 4.04E-02 | NaN | ADCY10,CAV1,GJB6,GJC1,GRIA2,NOTUM,NPR1,TUBA1A |
| | Growth Hormone Signaling | 1.16E+00 | 5.63E-02 | NaN | IGF1,IGF2,IGFBP3,IRS1 |
| | Estrogen Biosynthesis | 1.08E+00 | 6.38E-02 | NaN | CYP4X1,HSD17B13,HSD17B14 |
| | Retinoate Biosynthesis II | 1.07E+00 | 2.50E-01 | NaN | RBP5 |
| | RAR Activation | 1.06E+00 | 3.27E-02 | -0.535 | ADCY10,CLCF1,GATA6,IGFBP3,IL11,IL1B,MMP3,MMP8,PDE4C,PDE6B,RBP5,SOX9,STRA6,TNFSF8 |
| | Opioid Signaling Pathway | 1.06E+00 | 3.57E-02 | 0.707 | ADCY10,CACNA1G,CACNA2D3,FOSB,GRIN2A,KCNJ5,MAPK15,PNOC,RGS13,SCN7A |
| | Differential Regulation of Cytokine Production i | 1.05E+00 | 8.70E-02 | NaN | IL10,IL1B |
| | VDR/RXR Activation | 1.04E+00 | 5.13E-02 | NaN | COL13A1,IGFBP3,THBD,TRPV5 |
| | Role of WNT/GSK-3β Signaling in the Pathogene | 1.04E+00 | 5.13E-02 | 2 | WNT1,WNT10B,WNT3,WNT5B |
| | Maturity Onset Diabetes of Young (MODY) Sign | 1.03E+00 | 5.06E-02 | NaN | ALDOB,CACNA2D3,FOXA2,PKLR |
| | Thyroid Cancer Signaling | 1.03E+00 | 5.06E-02 | 1 | CXCR4,IGF1,IRS1,NTF3 |
| | FAT10 Cancer Signaling Pathway | 1.02E+00 | 6.00E-02 | NaN | ACKR3,CXCR4,NGFR |
| | Neutrophil Extracellular Trap Signaling Pathway | 1.01E+00 | 3.26E-02 | 0.832 | CASP5,COL13A1,COL5A1,COL5A3,COL6A5,COL7A1,IL10,IL1B,ITGA2,KCNN3,NOTUM,PLA2G1B,PLAAT4 |
| | Tryptophan Degradation III (Eukaryotic) | 9.90E-01 | 8.00E-02 | NaN | ACAT2,TDO2 |
| | Glycolysis I | 9.90E-01 | 8.00E-02 | NaN | ALDOB,PKLR |
| | Calcium Signaling | 9.62E-01 | 3.64E-02 | 1.342 | CACNA1G,CACNA2D3,CHRNA5,GRIA2,GRIN2A,MYL4,MYL9,TRPM8 |
| | Semaphorin Neuronal Repulsive Signaling Pathw | 9.58E-01 | 4.00E-02 | 0 | ITGA2,ITGA9,MYL4,MYL9,PDE4C,PLXNB1 |
| | Sperm Motility | 9.57E-01 | 3.50E-02 | -0.816 | ADCY10,CACNA1G,MET,NOTUM,NPR1,PDE4C,PDGFRB,PLA2G1B,PLAAT4 |
| | Neuregulin Signaling | 9.49E-01 | 4.27E-02 | NaN | AREG,EREG,ITGA2,ITGA9,NRG2 |
| | PAK Signaling | 9.49E-01 | 4.27E-02 | 1 | ITGA2,ITGA9,MYL4,MYL9,PDGFRB |
| | PTEN Signaling | 9.48E-01 | 3.97E-02 | -1.342 | FOXG1,ITGA2,ITGA9,NGFR,PDGFRB,SYNJ2 |
| | Dopamine-DARPP32 Feedback in cAMP Signalin | 9.48E-01 | 3.76E-02 | 1 | ADCY10,CACNA2D3,GRIN2A,KCNJ15,KCNJ5,NOTUM,PPM1J |
| | Phototransduction Pathway | 9.46E-01 | 5.56E-02 | NaN | NPR1,PDE6B,RGR |
| | Estrogen Receptor Signaling | 9.46E-01 | 3.18E-02 | 0.302 | ADCY10,CACNA2D3,CAV1,ESR2,FOXG1,IGF1,IGF2,MMP21,MMP3,MMP8,MYL4,MYL9,NOTUM |
| | Multiple Sclerosis Signaling Pathway | 9.46E-01 | 3.60E-02 | 1.414 | CLCF1,GRIN2A,IL10,IL11,IL1B,PLP1,TLR10,TNFSF8 |
| | GPCR-Mediated Nutrient Sensing in Enteroendo | 9.37E-01 | 4.24E-02 | -0.447 | ADCY10,CACNA1G,CACNA2D3,FFAR4,NOTUM |
| | Role of Hypercytokinemia/hyperchemokinemia | 9.27E-01 | 4.65E-02 | 1 | AREG,CXCL3,IL10,IL1B |
| | Sphingosine-1-phosphate Signaling | 9.15E-01 | 4.17E-02 | 2.236 | ADCY10,CASP5,NOTUM,PDGFRB,S1PR3 |
| | Histidine Degradation VI | 9.08E-01 | 7.14E-02 | NaN | AMDHD1,CYP4X1 |
| | UDP-N-acetyl-D-glucosamine Biosynthesis II | 9.07E-01 | 1.67E-01 | NaN | GFPT2 |

| | | | | |
|---|---|---|---|---|
| Tryptophan Degradation to 2-amino-3-carboxyr | 9.07E-01 | 1.67E-01 | NaN | TDO2 |
| NAD Biosynthesis III | 9.07E-01 | 1.67E-01 | NaN | NMNAT2 |
| Neuroprotective Role of THOP1 in Alzheimer's I | 9.04E-01 | 4.13E-02 | 1 | CTRB2,FAP,HTRA1,HTRA3,PNOC |
| Role of IL-17A in Arthritis | 8.95E-01 | 5.26E-02 | NaN | CCL7,CXCL3,CXCL5 |
| RHOA Signaling | 8.71E-01 | 4.03E-02 | 0.447 | CDC42EP5,IGF1,LPAR3,MYL4,MYL9 |
| Colorectal Cancer Metastasis Signaling | 8.56E-01 | 3.32E-02 | 1.667 | ADCY10,MMP21,MMP3,MMP8,TLR10,WNT1,WNT10B,WNT3,WNT5B |
| FXR/RXR Activation | 8.51E-01 | 3.97E-02 | NaN | ALB,CETP,FOXA2,IL1B,PKLR |
| NAD Biosynthesis from 2-amino-3-carboxymucc | 8.45E-01 | 1.43E-01 | NaN | NMNAT2 |
| Ceramide Biosynthesis | 8.45E-01 | 1.43E-01 | NaN | DEGS2 |
| NAD Salvage Pathway III | 8.45E-01 | 1.43E-01 | NaN | NMNAT2 |
| Glycoaminoglycan-protein Linkage Region Biosy | 8.45E-01 | 1.43E-01 | NaN | B3GAT1 |
| IL-12 Signaling and Production in Macrophages | 8.39E-01 | 3.39E-02 | 2.828 | ALB,COL5A3,FCGR1A,FCGR1BP,FCGR2B,FCGR2C,IL10,MAF |
| ID1 Signaling Pathway | 8.23E-01 | 3.48E-02 | 1.134 | CAV1,CHRNA5,FGF2,IGF2,NGFR,PLAC8,PLXNB1 |
| G Beta Gamma Signaling | 8.21E-01 | 3.88E-02 | 0.447 | CACNA1G,CACNA2D3,CAV1,CAV2,KCNJ5 |
| Ephrin Receptor Signaling | 8.15E-01 | 3.47E-02 | NaN | CXCR4,EFNB2,EFNB3,EPHA10,GRIN2A,ITGA2,ITGA9 |
| TGF-β Signaling | 8.03E-01 | 4.17E-02 | 1 | INHBA,NKX2-5,PMEPA1,SERPINE1 |
| Melatonin Degradation I | 8.02E-01 | 4.76E-02 | NaN | CYP4X1,SULT1B1,UGT3A1 |
| HMGB1 Signaling | 8.01E-01 | 3.59E-02 | NaN | CLCF1,IL11,IL1B,NGFR,SERPINE1,TNFSF8 |
| Airway Inflammation in Asthma | 7.93E-01 | 6.06E-02 | NaN | IL10,RNASE2 |
| HGF Signaling | 7.92E-01 | 3.79E-02 | NaN | ELF5,HGF,ITGA2,ITGA9,MET |
| Sertoli Cell-Sertoli Cell Junction Signaling | 7.85E-01 | 3.40E-02 | NaN | ADCY10,CLDN4,CLDN5,ITGA2,ITGA9,JAM2,TUBA1A |
| Gα12/13 Signaling | 7.82E-01 | 3.76E-02 | 0.447 | CDH13,F2RL3,LPAR3,MYL4,MYL9 |
| Role of Tissue Factor in Cancer | 7.77E-01 | 3.38E-02 | 1.134 | FLNC,HGF,IL1B,MAPK15,MET,NGFR,PPM1J |
| Induction of Apoptosis by HIV1 | 7.74E-01 | 4.62E-02 | NaN | CXCR4,NAIP,NGFR |
| Apelin Cardiomyocyte Signaling Pathway | 7.70E-01 | 4.04E-02 | 1 | MAPK15,MYL4,MYL9,NOTUM |
| CLEAR Signaling Pathway | 7.66E-01 | 3.16E-02 | -0.333 | BMP6,FGF2,HGF,IGF1,MAPK15,NGFR,PDGFRB,PPM1J,TLR10 |
| WNT/Ca+ pathway | 7.60E-01 | 4.55E-02 | NaN | NOTUM,PDE6B,WNT5B |
| Th2 Pathway | 7.46E-01 | 3.65E-02 | -0.447 | CD28,CXCR4,IL10,MAF,NOTCH3 |
| Sucrose Degradation V (Mammalian) | 7.45E-01 | 1.11E-01 | NaN | ALDOB |
| White Adipose Tissue Browning Pathway | 7.37E-01 | 3.62E-02 | -0.447 | ADCY10,CACNA1G,CACNA2D3,FCER1A,NPR1 |
| Superpathway of Melatonin Degradation | 7.34E-01 | 4.41E-02 | NaN | CYP4X1,SULT1B1,UGT3A1 |
| Remodeling of Epithelial Adherens Junctions | 7.34E-01 | 4.41E-02 | NaN | HGF,MET,TUBA1A |
| SPINK1 General Cancer Pathway | 7.21E-01 | 4.35E-02 | NaN | MT1E,MT1G,MT1H |
| Autophagy | 7.14E-01 | 3.24E-02 | 0.378 | BMP6,FGF2,HGF,IGF1,IRS1,NGFR,PPM1J |
| LPS/IL-1 Mediated Inhibition of RXR Function | 7.12E-01 | 3.14E-02 | 1 | CETP,CHST1,CHST3,HS3ST3B1,IL18RAP,IL1B,NGFR,SULT1B1 |
| Eicosanoid Signaling | 7.08E-01 | 4.29E-02 | NaN | ALOX15B,PLA2G1B,PLAAT4 |
| IGF-1 Signaling | 7.08E-01 | 3.81E-02 | NaN | IGF1,IGFBP3,IGFBP7,IRS1 |
| Embryonic Stem Cell Differentiation into Cardia | 7.04E-01 | 1.00E-01 | NaN | NKX2-5 |
| Ketolysis | 7.04E-01 | 1.00E-01 | NaN | ACAT2 |
| IL-17A Signaling in Fibroblasts | 6.98E-01 | 5.26E-02 | NaN | CCL7,CXCL5 |
| PPAR Signaling | 6.88E-01 | 3.74E-02 | -1 | IL18RAP,IL1B,NGFR,PDGFRB |
| Ephrin B Signaling | 6.84E-01 | 4.17E-02 | NaN | CXCR4,EFNB2,EFNB3 |
| Cardiac Hypertrophy Signaling | 6.76E-01 | 3.07E-02 | 0 | ADCY10,CACNA2D3,IGF1,IRS1,MYL4,MYL9,NKX2-5,NOTUM |
| Ketogenesis | 6.67E-01 | 9.09E-02 | NaN | ACAT2 |
| Glycine Betaine Degradation | 6.67E-01 | 9.09E-02 | NaN | SDS |
| Acute Phase Response Signaling | 6.64E-01 | 3.24E-02 | 2 | ALB,IL1B,NGFR,RBP5,SERPIND1,SERPINE1 |
| Role of NFAT in Cardiac Hypertrophy | 6.62E-01 | 3.12E-02 | 0 | ADCY10,CACNA1G,CACNA2D3,IGF1,IL11,NKX2-5,NOTUM |
| MicroRNA Biogenesis Signaling Pathway | 6.50E-01 | 3.21E-02 | 0.816 | BMP6,ESR2,FGF2,HGF,HOPX,IGF1 |
| GPCR-Mediated Integration of Enteroendocrine | 6.50E-01 | 4.00E-02 | NaN | ADCY10,GLP2R,NOTUM |
| Thyroid Hormone Metabolism II (via Conjugatio | 6.49E-01 | 4.88E-02 | NaN | SULT1B1,UGT3A1 |
| Dilated Cardiomyopathy Signaling Pathway | 6.39E-01 | 3.33E-02 | NaN | ADCY10,CACNA1G,CACNA2D3,MYL4,MYL9 |
| Macrophage Classical Activation Signaling Path | 6.36E-01 | 3.17E-02 | 1.633 | CLCF1,IL10,IL11,IL1B,MAF,TNFSF8 |
| Macrophage Alternative Activation Signaling Pa | 6.36E-01 | 3.17E-02 | -1.633 | FCER1A,FCGR2B,IL10,IL1B,IRS1,MAF |
| Circadian Rhythm Signaling | 6.36E-01 | 2.99E-02 | NaN | ADCY10,CACNA1G,CACNA2D3,GRIA2,GRIN2A,NGFR,NOTUM,NPR1 |
| Antioxidant Action of Vitamin C | 6.25E-01 | 3.51E-02 | 1 | NOTUM,PLA2G1B,PLAAT4,PLD4 |
| IL-7 Signaling Pathway | 6.17E-01 | 3.85E-02 | NaN | FOXG1,HGF,MET |
| Type II Diabetes Mellitus Signaling | 6.17E-01 | 3.27E-02 | NaN | CACNA1G,CACNA2D3,IRS1,NGFR,PKLR |
| Regulation of Actin-based Motility by Rho | 6.16E-01 | 3.48E-02 | NaN | ITGA2,ITGA9,MYL4,MYL9 |
| IL-13 Signaling Pathway | 6.08E-01 | 3.45E-02 | -1 | ALOX15B,FOSL2,FOXA2,IL10 |
| Bladder Cancer Signaling | 6.08E-01 | 3.45E-02 | NaN | FGF2,MMP21,MMP3,MMP8 |
| Role of JAK family kinases in IL-6-type Cytokine | 6.07E-01 | 3.80E-02 | NaN | CLCF1,IL10,IL11 |
| Role of MAPK Signaling in Inhibiting the Pathog | 6.07E-01 | 3.80E-02 | NaN | IL1B,PLA2G1B,PLAAT4 |
| NAD Phosphorylation and Dephosphorylation | 6.04E-01 | 7.69E-02 | NaN | PXYLP1 |
| γ-glutamyl Cycle | 6.04E-01 | 7.69E-02 | NaN | ANPEP |
| Guanosine Nucleotides Degradation III | 6.04E-01 | 7.69E-02 | NaN | NT5E |
| Role of Pattern Recognition Receptors in Recog | 5.96E-01 | 3.21E-02 | NaN | CLCF1,IL10,IL11,IL1B,TNFSF8 |
| Chemokine Signaling | 5.87E-01 | 3.70E-02 | NaN | CCL24,CCL7,CXCR4 |
| Ovarian Cancer Signaling | 5.82E-01 | 3.16E-02 | NaN | LHCGR,WNT1,WNT10B,WNT3,WNT5B |
| Role of OCT4 in Mammalian Embryonic Stem Ce | 5.77E-01 | 4.35E-02 | NaN | FBXO15,FOXA2 |
| tRNA Splicing | 5.77E-01 | 4.35E-02 | NaN | PDE4C,PDE6B |
| Glycogen Degradation III | 5.76E-01 | 7.14E-02 | NaN | MGAM |
| Urate Biosynthesis/Inosine 5'-phosphate Degrad | 5.76E-01 | 7.14E-02 | NaN | NT5E |
| Mevalonate Pathway I | 5.76E-01 | 7.14E-02 | NaN | ACAT2 |
| Oxytocin In Brain Signaling Pathway | 5.73E-01 | 3.02E-02 | -0.816 | CACNA1G,CACNA2D3,CASP5,IL1B,PLA2G1B,PLAAT4 |
| Estrogen-Dependent Breast Cancer Signaling | 5.68E-01 | 3.61E-02 | NaN | HSD17B13,HSD17B14,IGF1 |
| PI3K/AKT Signaling | 5.67E-01 | 3.00E-02 | NaN | IL18RAP,IL31RA,ITGA2,ITGA9,PPM1J,SYNJ2 |
| Ephrin A Signaling | 5.64E-01 | 4.26E-02 | NaN | EPHA10,NGFR |
| nNOS Signaling in Skeletal Muscle Cells | 5.51E-01 | 4.17E-02 | NaN | CACNA1G,CACNA2D3 |
| Choline Biosynthesis III | 5.51E-01 | 6.67E-02 | NaN | PLD4 |
| Protein Kinase A Signaling | 5.44E-01 | 2.68E-02 | -1.414 | ADCY10,FLNC,MYL4,MYL9,NGFR,NOTUM,PDE4C,PDE6B,PTPN14,PTPRD,PTPRN |
| FGF Signaling | 5.40E-01 | 3.49E-02 | NaN | FGF2,HGF,MET |
| PDGF Signaling | 5.31E-01 | 3.45E-02 | NaN | CAV1,PDGFRB,SYNJ2 |
| D-myo-inositol (1,4,5)-trisphosphate Degradatic | 5.27E-01 | 6.25E-02 | NaN | SYNJ2 |
| Adenosine Nucleotides Degradation II | 5.27E-01 | 6.25E-02 | NaN | NT5E |
| Parkinson's Signaling | 5.27E-01 | 6.25E-02 | NaN | GPR37 |
| 1D-myo-inositol Hexakisphosphate Biosynthesis | 5.05E-01 | 5.88E-02 | NaN | SYNJ2 |
| D-myo-inositol (1,3,4)-trisphosphate Biosynthes | 5.05E-01 | 5.88E-02 | NaN | SYNJ2 |
| Ceramide Signaling | 4.97E-01 | 3.30E-02 | NaN | NGFR,PPM1J,S1PR3 |
| Th1 and Th2 Activation Pathway | 4.94E-01 | 2.91E-02 | NaN | CD28,CXCR4,IL10,MAF,NOTCH3 |
| ABRA Signaling Pathway | 4.89E-01 | 3.26E-02 | NaN | FOSB,IGF1,MYL9 |
| Ferroptosis Signaling Pathway | 4.89E-01 | 3.03E-02 | -1 | ALOX15B,SLC39A8,SLC7A11,TFAP2C |
| Glutaryl-CoA Degradation | 4.85E-01 | 5.56E-02 | NaN | ACAT2 |
| Superpathway of Geranylgeranyldiphosphate Bi | 4.85E-01 | 5.56E-02 | NaN | ACAT2 |
| Role of Cytokines in Mediating Communication | 4.82E-01 | 3.70E-02 | NaN | IL10,IL1B |
| ERBB Signaling | 4.81E-01 | 3.23E-02 | NaN | AREG,EREG,NRG2 |
| Triacylglycerol Biosynthesis | 4.72E-01 | 3.64E-02 | NaN | LPCAT1,PLPP4 |
| Purine Nucleotides Degradation II (Aerobic) | 4.66E-01 | 5.26E-02 | NaN | NT5E |
| Erythropoietin Signaling Pathway | 4.66E-01 | 2.82E-02 | -0.447 | CLCF1,IL11,IL1B,TNFSF8,WNT1 |
| Role of PKR in Interferon Induction and Antivira | 4.63E-01 | 2.94E-02 | NaN | CASP5,FCGR1A,IL1B,PDGFRB |
| RHOGDI Signaling | 4.60E-01 | 2.73E-02 | 0.447 | CDH13,ESR2,ITGA2,ITGA9,MYL4,MYL9 |
| DHCR24 Signaling Pathway | 4.57E-01 | 2.92E-02 | -1 | ALB,CETP,IGF1,SOX9 |
| RAC Signaling | 4.57E-01 | 2.92E-02 | NaN | ANK1,ITGA2,ITGA9,MCF2L |
| D-myo-inositol (1,4,5,6)-Tetrakisphosphate Bios | 4.50E-01 | 2.78E-02 | NaN | ATP1A2,PPM1J,PTPRN,PXYLP1,SYNJ2 |
| D-myo-inositol (3,4,5,6)-tetrakisphosphate Bios | 4.50E-01 | 2.78E-02 | NaN | ATP1A2,PPM1J,PTPRN,PXYLP1,SYNJ2 |
| Cardiac β-adrenergic Signaling | 4.50E-01 | 2.78E-02 | -2 | ADCY10,CACNA2D3,PDE4C,PDE6B,PPM1J |
| Androgen Biosynthesis | 4.48E-01 | 5.00E-02 | NaN | HSD17B14 |
| Valine Degradation I | 4.48E-01 | 5.00E-02 | NaN | SDS |
| Thrombin Signaling | 4.36E-01 | 2.67E-02 | 1 | ADCY10,F2RL3,GATA6,MYL4,MYL9,NOTUM |
| Nicotine Degradation III | 4.33E-01 | 3.39E-02 | NaN | CYP4X1,UGT3A1 |
| Superpathway of D-myo-inositol (1,4,5)-trispho | 4.16E-01 | 4.55E-02 | NaN | SYNJ2 |
| Semaphorin Signaling in Neurons | 4.15E-01 | 3.28E-02 | NaN | MET,PLXNB1 |
| Molecular Mechanisms of Cancer | 4.08E-01 | 2.44E-02 | NaN | ADCY10,BMP6,HIPK2,IRS1,ITGA2,ITGA9,NAIP,WNT1,WNT10B,WNT3,WNT5B |
| Role of JAK2 in Hormone-like Cytokine Signaling | 4.06E-01 | 3.23E-02 | NaN | IGF1,IRS1 |
| NOD1/2 Signaling Pathway | 4.04E-01 | 2.65E-02 | 0.447 | CLCF1,IL11,IL1B,TLR10,TNFSF8 |
| The Visual Cycle | 4.01E-01 | 4.35E-02 | NaN | RBP5 |
| Pyrimidine Deoxyribonucleotides De Novo Biosy | 4.01E-01 | 4.35E-02 | NaN | AK7 |
| Vitamin-C Transport | 4.01E-01 | 4.35E-02 | NaN | LRRC8E |
| Apelin Cardiac Fibroblast Signaling Pathway | 4.01E-01 | 4.35E-02 | NaN | SERPINE1 |
| Chronic Myeloid Leukemia Signaling | 3.95E-01 | 2.52E-02 | 1.89 | MECOM,NOTCH3,NOTUM,WNT1,WNT10B,WNT3,WNT5B |
| 3-phosphoinositide Degradation | 3.95E-01 | 2.62E-02 | NaN | ATP1A2,PPM1J,PTPRN,PXYLP1,SYNJ2 |
| Xenobiotic Metabolism CAR Signaling Pathway | 3.90E-01 | 2.60E-02 | 2.236 | CHST1,CHST3,HS3ST3B1,PPM1J,SULT1B1 |
| WNK Renal Signaling Pathway | 3.90E-01 | 2.83E-02 | NaN | IGF1,IRS1,LRRC8E |
| Tumoricidal Function of Hepatic Natural Killer C | 3.87E-01 | 4.17E-02 | NaN | LYVE1 |
| PPARα/RXRα Activation | 3.77E-01 | 2.56E-02 | -1 | ADCY10,IL18RAP,IL1B,IRS1,NOTUM |
| Corticotropin Releasing Hormone Signaling | 3.74E-01 | 2.63E-02 | 0 | ADCY10,CACNA1G,CACNA2D3,NPR1 |
| Bupropion Degradation | 3.73E-01 | 4.00E-02 | NaN | CYP4X1 |
| Gluconeogenesis I | 3.73E-01 | 4.00E-02 | NaN | ALDOB |

| | | | | |
|---|---|---|---|---|
| D-myo-inositol-5-phosphate Metabolism | 3.73E-01 | 2.55E-02 | NaN | ATP1A2,PPM1J,PTPRN,PXYLP1,SYNJ2 |
| IL-17A Signaling in Airway Cells | 3.66E-01 | 2.99E-02 | NaN | CXCL3,CXCL5 |
| Nicotine Degradation II | 3.66E-01 | 2.99E-02 | NaN | CYP4X1,UGT3A1 |
| Estrogen-mediated S-phase Entry | 3.61E-01 | 3.85E-02 | NaN | ESR2 |
| Relaxin Signaling | 3.59E-01 | 2.58E-02 | NaN | ADCY10,NPR1,PDE4C,PDE6B |
| Agrin Interactions at Neuromuscular Junction | 3.52E-01 | 2.90E-02 | NaN | NRG2,PKLR |
| ILK Signaling | 3.51E-01 | 2.49E-02 | -0.447 | FLNC,IRS1,MYL4,MYL9,PPM1J |
| Epithelial Adherens Junction Signaling | 3.50E-01 | 2.55E-02 | 0 | HGF,MET,NOTCH3,PPM1J |
| Cardiomyocyte Differentiation via BMP Recepto | 3.48E-01 | 3.70E-02 | NaN | NKX2-5 |
| Pregnenolone Biosynthesis | 3.48E-01 | 3.70E-02 | NaN | CYP4X1 |
| Xenobiotic Metabolism Signaling | 3.45E-01 | 2.40E-02 | NaN | CHST1,CHST3,HS3ST3B1,IL1B,MAF,PPM1J,SULT1B1 |
| CDP-diacylglycerol Biosynthesis I | 3.37E-01 | 3.57E-02 | NaN | LPCAT1 |
| 3-phosphoinositide Biosynthesis | 3.31E-01 | 2.43E-02 | NaN | ATP1A2,PPM1J,PTPRN,PXYLP1,SYNJ2 |
| Netrin Signaling | 3.31E-01 | 2.78E-02 | NaN | CACNA1G,CACNA2D3 |
| Serotonin Degradation | 3.31E-01 | 2.78E-02 | NaN | SULT1B1,UGT3A1 |
| Inhibition of ARE-Mediated mRNA Degradation | 3.27E-01 | 2.47E-02 | 0 | MAPK15,NGFR,PPM1J,TNFSF8 |
| Superpathway of Cholesterol Biosynthesis | 3.26E-01 | 3.45E-02 | NaN | ACAT2 |
| Fc Epsilon RI Signaling | 3.23E-01 | 2.54E-02 | NaN | FCER1A,PLA2G1B,SYNJ2 |
| Virus Entry via Endocytic Pathways | 3.23E-01 | 2.54E-02 | NaN | CAV1,FLNC,ITGA2 |
| Phosphatidylglycerol Biosynthesis II (Non-plasti | 3.16E-01 | 3.33E-02 | NaN | LPCAT1 |
| Nitric Oxide Signaling in the Cardiovascular Syst | 3.13E-01 | 2.50E-02 | NaN | CACNA2D3,CAV1,NPR1 |
| p38 MAPK Signaling | 3.13E-01 | 2.50E-02 | NaN | IL18RAP,IL1B,PLA2G1B |
| Ubiquinol-10 Biosynthesis (Eukaryotic) | 3.06E-01 | 3.23E-02 | NaN | CYP4X1 |
| Leptin Signaling in Obesity | 3.05E-01 | 2.63E-02 | NaN | ADCY10,NOTUM |
| Macropinocytosis Signaling | 3.05E-01 | 2.63E-02 | NaN | HGF,MET |
| Th1 Pathway | 3.03E-01 | 2.46E-02 | NaN | CD28,IL10,NOTCH3 |
| CXCR4 Signaling | 3.02E-01 | 2.38E-02 | 1 | ADCY10,CXCR4,MYL4,MYL9 |
| Antiproliferative Role of Somatostatin Receptor | 2.99E-01 | 2.60E-02 | NaN | NPR1,SSTR2 |
| Glucocorticoid Receptor Signaling | 2.99E-01 | 2.23E-02 | NaN | CAV1,CXCL3,FCGR1A,IGF1,IL10,IL18RAP,IL1B,IL31RA,MMP3,MMP8,PLA2G1B,SERPINE1,SLPI |
| ERK/MAPK Signaling | 2.98E-01 | 2.33E-02 | -1 | ELF5,ITGA2,ITGA9,PLA2G1B,PPM1J |
| TNFR2 Signaling | 2.96E-01 | 3.12E-02 | NaN | NAIP |
| Dopamine Degradation | 2.96E-01 | 3.12E-02 | NaN | SULT1B1 |
| Toll-like Receptor Signaling | 2.93E-01 | 2.56E-02 | NaN | IL1B,TLR10 |
| Neurotrophin/TRK Signaling | 2.93E-01 | 2.56E-02 | NaN | NGFR,NTF3 |
| Glioma Signaling | 2.89E-01 | 2.40E-02 | NaN | IGF1,IGF2,PDGFRB |
| Renal Cell Carcinoma Signaling | 2.88E-01 | 2.53E-02 | NaN | HGF,MET |
| Gαs Signaling | 2.84E-01 | 2.38E-02 | NaN | ADCY10,HTR4,LHCGR |
| Dopamine Receptor Signaling | 2.82E-01 | 2.50E-02 | NaN | ADCY10,PPM1J |
| Signaling by Rho Family GTPases | 2.79E-01 | 2.25E-02 | 0.447 | CDC42EP5,CDH13,ITGA2,ITGA9,MYL4,MYL9 |
| Inhibition of Angiogenesis by TSP1 | 2.78E-01 | 2.94E-02 | NaN | HSPG2 |
| Fatty Acid β-oxidation I | 2.78E-01 | 2.94E-02 | NaN | SDS |
| IL-6 Signaling | 2.71E-01 | 2.33E-02 | NaN | IL18RAP,IL1B,NGFR |
| BEX2 Signaling Pathway | 2.71E-01 | 2.44E-02 | NaN | NGFR,PPM1J |
| Dendritic Cell Maturation | 2.70E-01 | 2.18E-02 | 0.302 | CCR7,CD1B,COL5A3,FCGR1A,FCGR1BP,FCGR2B,FCGR2C,IL10,IL1B,NGFR,NOTUM,TRAJ23,TRBV5-4 |
| Oxytocin In Spinal Neurons Signaling Pathway | 2.70E-01 | 2.86E-02 | NaN | NPR1 |
| IL-9 Signaling | 2.70E-01 | 2.86E-02 | NaN | IRS1 |
| MIF-mediated Glucocorticoid Regulation | 2.62E-01 | 2.78E-02 | NaN | PLA2G1B |
| TR/RXR Activation | 2.61E-01 | 2.38E-02 | NaN | RAB3B,SLC16A2 |
| Role of MAPK Signaling in the Pathogenesis of I | 2.61E-01 | 2.38E-02 | NaN | PLA2G1B,PLAAT4 |
| Synaptic Long Term Potentiation | 2.59E-01 | 2.27E-02 | NaN | GRIA2,GRIN2A,NOTUM |
| Cyclins and Cell Cycle Regulation | 2.55E-01 | 2.35E-02 | NaN | MYT1,PPM1J |
| P2Y Purigenic Receptor Signaling Pathway | 2.55E-01 | 2.26E-02 | NaN | ADCY10,NOTUM,P2RY12 |
| TWEAK Signaling | 2.54E-01 | 2.70E-02 | NaN | NAIP |
| Glutathione-mediated Detoxification | 2.54E-01 | 2.70E-02 | NaN | ANPEP |
| Complement System | 2.54E-01 | 2.70E-02 | NaN | CR1 |
| Regulation of eIF4 and p70S6K Signaling | 2.54E-01 | 2.21E-02 | NaN | IRS1,ITGA2,ITGA9,PPM1J |
| HIPPO signaling | 2.51E-01 | 2.33E-02 | NaN | PPM1J,TEAD1 |
| Docosahexaenoic Acid (DHA) Signaling | 2.46E-01 | 2.63E-02 | NaN | IL1B |
| Cell Cycle Regulation by BTG Family Proteins | 2.46E-01 | 2.63E-02 | NaN | PPM1J |
| Notch Signaling | 2.46E-01 | 2.63E-02 | NaN | NOTCH3 |
| Pyrimidine Ribonucleotides Interconversion | 2.39E-01 | 2.56E-02 | NaN | AK7 |
| Immunogenic Cell Death Signaling Pathway | 2.32E-01 | 2.22E-02 | NaN | IL1B,NGFR |
| Regulation of Cellular Mechanics by Calpain Prc | 2.32E-01 | 2.22E-02 | NaN | ITGA2,ITGA9 |
| Crosstalk between Dendritic Cells and Natural K | 2.27E-01 | 2.20E-02 | NaN | CCR7,CD28 |
| BMP signaling pathway | 2.27E-01 | 2.20E-02 | NaN | BMP6,NKX2-5 |
| Apelin Adipocyte Signaling Pathway | 2.27E-01 | 2.20E-02 | NaN | ADCY10,MAPK15 |
| Retinoate Biosynthesis I | 2.19E-01 | 2.38E-02 | NaN | RBP5 |
| Pyrimidine Ribonucleotides De Novo Biosynthes | 2.19E-01 | 2.38E-02 | NaN | AK7 |
| Acetone Degradation I (to Methylglyoxal) | 2.19E-01 | 2.38E-02 | NaN | CYP4X1 |
| Oncostatin M Signaling | 2.13E-01 | 2.33E-02 | NaN | MMP3 |
| MIF Regulation of Innate Immunity | 2.07E-01 | 2.27E-02 | NaN | PLA2G1B |
| Coronavirus Replication Pathway | 2.01E-01 | 2.22E-02 | NaN | TUBA1A |
| CSDE1 Signaling Pathway | 0.00E+00 | 1.79E-02 | NaN | TNC |
| Oxytocin Signaling Pathway | 0.00E+00 | 2.13E-02 | 0 | CACNA2D3,MAPK15,MYL4,MYL9,NPR1,PLA2G1B |
| Ribonucleotide Reductase Signaling Pathway | 0.00E+00 | 1.18E-02 | NaN | HGF,MET |
| Natural Killer Cell Signaling | 0.00E+00 | 2.02E-02 | 0 | COL5A3,IL18RAP,NCR1,ULBP3 |
| Actin Cytoskeleton Signaling | 0.00E+00 | 2.05E-02 | NaN | FGF2,ITGA2,ITGA9,MYL4,MYL9 |
| Huntington's Disease Signaling | 0.00E+00 | 1.06E-02 | NaN | CASP5,IGF1,STX1A |
| Chaperone Mediated Autophagy Signaling Pathv | 0.00E+00 | 1.24E-02 | 0 | EEF1A2,IL1B,MMP21,MMP3,MMP8,PRR5L,TRAJ23,TRBV5-4 |
| IL-33 Signaling Pathway | 0.00E+00 | 2.15E-02 | 2 | AREG,CASP5,CCL24,IL1B |
| NRF2-mediated Oxidative Stress Response | 0.00E+00 | 4.22E-03 | NaN | MAF |
| Aryl Hydrocarbon Receptor Signaling | 0.00E+00 | 1.89E-02 | NaN | ESR2,IL1B,NFIB |
| p53 Signaling | 0.00E+00 | 1.02E-02 | NaN | HIPK2 |
| Mitochondrial Dysfunction | 0.00E+00 | 1.16E-02 | 0 | ADCY10,ATP1A2,CACNA1G,CACNA2D3 |
| PXR/RXR Activation | 0.00E+00 | 1.54E-02 | NaN | FOXA2 |
| 14-3-3-mediated Signaling | 0.00E+00 | 1.57E-02 | NaN | NOTUM,TUBA1A |
| α-Adrenergic Signaling | 0.00E+00 | 9.17E-03 | NaN | ADCY10 |
| Activation of IRF by Cytosolic Pattern Recognitic | 0.00E+00 | 1.54E-02 | NaN | IL10 |
| Clathrin-mediated Endocytosis Signaling | 0.00E+00 | 1.92E-02 | NaN | ALB,FGF2,IGF1,MET |
| Fcγ Receptor-mediated Phagocytosis in Macrop | 0.00E+00 | 2.13E-02 | NaN | FCGR1A,PLD4 |
| IL-8 Signaling | 0.00E+00 | 9.52E-03 | NaN | MYL9,PLD4 |
| Role of NFAT in Regulation of the Immune Resp | 0.00E+00 | 7.71E-03 | -2.449 | CD28,FCER1A,FCGR1A,FCGR1BP,FCGR2B,FCGR2C,TRAJ23,TRBV5-4 |
| FcγRIIB Signaling in B Lymphocytes | 0.00E+00 | 5.63E-03 | NaN | CACNA1G,CACNA2D3,FCGR2B |
| NF-κB Activation by Viruses | 0.00E+00 | 1.28E-02 | NaN | ITGA2 |
| CCR5 Signaling in Macrophages | 0.00E+00 | 8.02E-03 | NaN | CACNA1G,CACNA2D3,TRAJ23,TRBV5-4 |
| Calcium-induced T Lymphocyte Apoptosis | 0.00E+00 | 4.35E-03 | NaN | TRAJ23,TRBV5-4 |
| Cytotoxic T Lymphocyte-mediated Apoptosis of | 0.00E+00 | 4.71E-03 | NaN | TRAJ23,TRBV5-4 |
| CTLA4 Signaling in Cytotoxic T Lymphocytes | 0.00E+00 | 9.87E-03 | 0 | CD28,ITGA2,PLD4,PPM1J,TRAJ23,TRBV5-4 |
| IL-15 Production | 0.00E+00 | 1.63E-02 | NaN | MET,PDGFRB |
| T Helper Cell Differentiation | 0.00E+00 | 1.06E-02 | NaN | CD28,IL10,NGFR,TRAJ23,TRBV5-4 |
| CCR3 Signaling in Eosinophils | 0.00E+00 | 1.49E-02 | NaN | CCL24,PLA2G1B |
| CD28 Signaling in T Helper Cells | 0.00E+00 | 5.80E-03 | NaN | CD28,TRAJ23,TRBV5-4 |
| Reelin Signaling in Neurons | 0.00E+00 | 7.25E-03 | NaN | GRIN2A |
| Melatonin Signaling | 0.00E+00 | 1.39E-02 | NaN | NOTUM |
| Renin-Angiotensin Signaling | 0.00E+00 | 8.26E-03 | NaN | ADCY10 |
| ICOS-ICOSL Signaling in T Helper Cells | 0.00E+00 | 5.92E-03 | NaN | CD28,TRAJ23,TRBV5-4 |
| Lipid Antigen Presentation by CD1 | 0.00E+00 | 7.23E-03 | NaN | CD1B,TRAJ23,TRBV5-4 |
| Mitotic Roles of Polo-Like Kinase | 0.00E+00 | 1.49E-02 | NaN | PPM1J |
| Role of CHK Proteins in Cell Cycle Checkpoint Cc | 0.00E+00 | 1.72E-02 | NaN | PPM1J |
| GNRH Signaling | 0.00E+00 | 1.57E-02 | NaN | ADCY10,CACNA1G,CACNA2D3 |
| Cholecystokinin/Gastrin-mediated Signaling | 0.00E+00 | 8.40E-03 | NaN | IL1B |
| Melanocyte Development and Pigmentation Sig | 0.00E+00 | 1.02E-02 | NaN | ADCY10 |
| DNA Methylation and Transcriptional Repressio | 0.00E+00 | 2.04E-02 | NaN | FOXA2,TEAD1 |
| ATM Signaling | 0.00E+00 | 1.00E-02 | NaN | PPM1J |
| Androgen Signaling | 0.00E+00 | 1.18E-02 | NaN | CACNA1G,CACNA2D3 |
| Germ Cell-Sertoli Cell Junction Signaling | 0.00E+00 | 1.18E-02 | NaN | ITGA2,TUBA1A |
| Aldosterone Signaling in Epithelial Cells | 0.00E+00 | 1.74E-02 | NaN | CRYAB,KCNMB1,NOTUM |
| Prolactin Signaling | 0.00E+00 | 2.11E-02 | NaN | IRS1,KCNMB1 |
| Type I Diabetes Mellitus Signaling | 0.00E+00 | 1.18E-02 | NaN | CD28,IL1B,NGFR,PTPRN,TRAJ23,TRBV5-4 |
| Allograft Rejection Signaling | 0.00E+00 | 8.21E-03 | NaN | CD28,IL10,TRAJ23,TRBV5-4 |
| Autoimmune Thyroid Disease Signaling | 0.00E+00 | 8.73E-03 | NaN | CD28,IL10,TRAJ23,TRBV5-4 |
| Graft-versus-Host Disease Signaling | 0.00E+00 | 8.97E-03 | NaN | CD28,IL1B,TRAJ23,TRBV5-4 |
| Production of Nitric Oxide and Reactive Oxygen | 0.00E+00 | 1.57E-02 | NaN | ALB,NGFR,PPM1J |
| p70S6K Signaling | 0.00E+00 | 6.88E-03 | 0 | F2RL3,IRS1,NOTUM,PPM1J |
| mTOR Signaling | 0.00E+00 | 1.87E-02 | NaN | IRS1,PLD4,PPM1J,PRR5L |
| MYC Mediated Apoptosis Signaling | 0.00E+00 | 2.00E-02 | NaN | NGFR |
| Pancreatic Adenocarcinoma Signaling | 0.00E+00 | 7.94E-03 | NaN | PLD4 |

| | | | | |
|---|---|---|---|---|
| G Protein Signaling Mediated by Tubby | 0.00E+00 | 4.27E-03 | NaN | TRAJ23,TRBV5-4 |
| Communication between Innate and Adaptive Ir | 0.00E+00 | 7.50E-03 | NaN | CCR7,CD28,IL10,IL1B,TLR10,TRAJ23,TRBV5-4 |
| Systemic Lupus Erythematosus Signaling | 0.00E+00 | 8.43E-03 | NaN | CD28,FCGR1A,FCGR1BP,FCGR2B,FCGR2C,IL10,IL1B,TRAJ23,TRBV5-4 |
| CDC42 Signaling | 0.00E+00 | 1.22E-02 | 1 | CDC42EP5,ITGA2,ITGA9,MYL4,MYL9,TRAJ23,TRBV5-4 |
| AMPK Signaling | 0.00E+00 | 2.07E-02 | NaN | AK7,CHRNA5,FOXG1,IRS1,PPM1J |
| Phospholipase C Signaling | 0.00E+00 | 9.81E-03 | -0.816 | ADCY10,FCGR2B,FCGR2C,ITGA2,ITGA9,MYL4,MYL9,PLA2G1B,PLD4,TRAJ23,TRBV5-4 |
| HER-2 Signaling in Breast Cancer | 0.00E+00 | 1.32E-02 | NaN | AREG,ELF5,FCER1A |
| Altered T Cell and B Cell Signaling in Rheumatoi | 0.00E+00 | 6.38E-03 | NaN | CD28,IL10,IL1B,TLR10,TRAJ23,TRBV5-4 |
| IL-1 Signaling | 0.00E+00 | 1.04E-02 | NaN | ADCY10 |
| Regulation of IL-2 Expression in Activated and A | 0.00E+00 | 6.49E-03 | NaN | CD28,TRAJ23,TRBV5-4 |
| Granzyme A Signaling | 0.00E+00 | 1.33E-02 | NaN | IL1B |
| NUR77 Signaling in T Lymphocytes | 0.00E+00 | 5.85E-03 | NaN | CD28,TRAJ23,TRBV5-4 |
| PKCθ Signaling in T Lymphocytes | 0.00E+00 | 8.96E-03 | NaN | CACNA1G,CACNA2D3,CD28,TRAJ23,TRBV5-4 |
| TNFR1 Signaling | 0.00E+00 | 1.96E-02 | NaN | NAIP |
| Role of PI3K/AKT Signaling in the Pathogenesis ( | 0.00E+00 | 1.52E-02 | NaN | PLAC8 |
| Antiproliferative Role of TOB in T Cell Signaling | 0.00E+00 | 7.04E-03 | NaN | CD28,TRAJ23,TRBV5-4 |
| OX40 Signaling Pathway | 0.00E+00 | 4.18E-03 | NaN | TRAJ23,TRBV5-4 |
| PI3K Signaling in B Lymphocytes | 0.00E+00 | 6.77E-03 | NaN | CARD10,FCGR2B,IRS1,NOTUM |
| Role of JAK1 and JAK3 in γc Cytokine Signaling | 0.00E+00 | 1.45E-02 | NaN | IRS1 |
| Actin Nucleation by ARP-WASP Complex | 0.00E+00 | 2.15E-02 | NaN | ITGA2,ITGA9 |
| NGF Signaling | 0.00E+00 | 8.33E-03 | NaN | NGFR |
| Paxillin Signaling | 0.00E+00 | 1.87E-02 | NaN | ITGA2,ITGA9 |
| Telomerase Signaling | 0.00E+00 | 1.85E-02 | NaN | ELF5,PPM1J |
| Mouse Embryonic Stem Cell Pluripotency | 0.00E+00 | 9.62E-03 | NaN | ID4 |
| Hematopoiesis from Pluripotent Stem Cells | 0.00E+00 | 8.99E-03 | NaN | IL10,IL11,TRAJ23,TRBV5-4 |
| nNOS Signaling in Neurons | 0.00E+00 | 2.13E-02 | NaN | GRIN2A |
| VEGF Family Ligand-Receptor Interactions | 0.00E+00 | 1.19E-02 | NaN | PLA2G1B |
| ERB2-ERBB3 Signaling | 0.00E+00 | 1.54E-02 | NaN | NRG2 |
| ERBB4 Signaling | 0.00E+00 | 1.47E-02 | NaN | NRG2 |
| GADD45 Signaling | 0.00E+00 | 1.67E-02 | NaN | IL1B |
| GDNF Family Ligand-Receptor Interactions | 0.00E+00 | 1.32E-02 | NaN | IRS1 |
| Triacylglycerol Degradation | 0.00E+00 | 1.75E-02 | NaN | NOTUM |
| Superpathway of Inositol Phosphate Compound | 0.00E+00 | 2.14E-02 | NaN | ATP1A2,PPM1J,PTPRN,PXYLP1,SYNJ2 |
| Retinol Biosynthesis | 0.00E+00 | 2.13E-02 | NaN | RBP5 |
| Salvage Pathways of Pyrimidine Ribonucleotide: | 0.00E+00 | 1.03E-02 | NaN | AK7 |
| DNA damage-induced 14-3-3σ Signaling | 0.00E+00 | 2.17E-02 | NaN | CASP5 |
| Gαq Signaling | 0.00E+00 | 5.88E-03 | NaN | PLD4 |
| TEC Kinase Signaling | 0.00E+00 | 8.65E-03 | NaN | FCER1A,ITGA2,ITGA9,TRAJ23,TRBV5-4 |
| UVA-Induced MAPK Signaling | 0.00E+00 | 1.02E-02 | NaN | NOTUM |
| Adipogenesis pathway | 0.00E+00 | 2.16E-02 | NaN | FGF2,SOX9,WNT10B |
| SAPK/JNK Signaling | 0.00E+00 | 5.98E-03 | NaN | IRS1,TRAJ23,TRBV5-4 |
| Protein Ubiquitination Pathway | 0.00E+00 | 7.33E-03 | NaN | CRYAB,USP51 |
| Cell Cycle: G2/M DNA Damage Checkpoint Regu | 0.00E+00 | 2.00E-02 | NaN | HIPK2 |
| IL-4 Signaling | 0.00E+00 | 1.91E-02 | 0.302 | ALOX15B,COL13A1,COL5A1,COL5A3,COL6A5,COL7A1,IRS1,MAF,PRR5L,TRAJ23,TRBV5-4 |
| B Cell Receptor Signaling | 0.00E+00 | 6.30E-03 | NaN | CARD10,FCGR2B,FCGR2C,SYNJ2 |
| Insulin Receptor Signaling | 0.00E+00 | 1.43E-02 | NaN | IRS1,SYNJ2 |
| Integrin Signaling | 0.00E+00 | 1.89E-02 | 0 | CAV1,ITGA2,ITGA9,MYL9 |
| Death Receptor Signaling | 0.00E+00 | 1.04E-02 | NaN | NAIP |
| Apoptosis Signaling | 0.00E+00 | 1.92E-02 | NaN | BCL2L10,NAIP |
| NF-κB Signaling | 0.00E+00 | 1.23E-02 | 1.342 | CARD10,IL1B,NGFR,PDGFRB,TLR10,TRAJ23,TRBV5-4 |
| T Cell Receptor Signaling | 0.00E+00 | 6.49E-03 | 1 | CD28,ITGA2,TRAJ23,TRBV5-4 |
| Phagosome Maturation | 0.00E+00 | 1.23E-02 | NaN | STX1A,TUBA1A |
| PD-1, PD-L1 cancer immunotherapy pathway | 0.00E+00 | 1.87E-02 | NaN | CD28,NGFR |
| Cancer Drug Resistance By Drug Efflux | 0.00E+00 | 1.72E-02 | NaN | FOXG1 |
| Sirtuin Signaling Pathway | 0.00E+00 | 6.83E-03 | NaN | MAPK15,TUBA1A |
| Iron homeostasis signaling pathway | 0.00E+00 | 2.17E-02 | NaN | BMP6,PDGFRB,SLC40A1 |
| Th17 Activation Pathway | 0.00E+00 | 8.26E-03 | NaN | IL10,IL1B,TRAJ23,TRBV5-4 |
| SPINK1 Pancreatic Cancer Pathway | 0.00E+00 | 1.67E-02 | NaN | CTRB2 |
| Endocannabinoid Developing Neuron Pathway | 0.00E+00 | 1.57E-02 | NaN | ADCY10,MAPK15 |
| Endocannabinoid Cancer Inhibition Pathway | 0.00E+00 | 2.04E-02 | NaN | ADCY10,CASP5,TWIST1 |
| Apelin Endothelial Signaling Pathway | 0.00E+00 | 1.42E-02 | NaN | ADCY10,AQP2 |
| PFKFB4 Signaling Pathway | 0.00E+00 | 2.08E-02 | NaN | FGF2 |
| T Cell Exhaustion Signaling Pathway | 0.00E+00 | 1.06E-02 | 1 | CD28,EOMES,IL10,PPM1J,TRAJ23,TRBV5-4 |
| IL-23 Signaling Pathway | 0.00E+00 | 2.17E-02 | NaN | IL1B |
| Systemic Lupus Erythematosus In T Cell Signalin | 0.00E+00 | 1.09E-02 | 0.447 | CASP5,CD28,IL10,PPM1J,S1PR3,TRAJ23,TRBV5-4 |
| Systemic Lupus Erythematosus In B Cell Signalin | 0.00E+00 | 1.38E-02 | 0 | CLCF1,FCGR2B,FCGR2C,FOXG1,IL10,IL11,IL1B,PLAAT4,SYNJ2,TNFSF8 |
| Senescence Pathway | 0.00E+00 | 2.01E-02 | 1.342 | CACNA2D3,ELF5,HIPK2,MAPK15,PPM1J,SERPINE1 |
| Necroptosis Signaling Pathway | 0.00E+00 | 1.28E-02 | NaN | NGFR,PLA2G1B |
| Xenobiotic Metabolism General Signaling Pathw | 0.00E+00 | 1.40E-02 | NaN | ABCA4,MAF |
| Xenobiotic Metabolism AHR Signaling Pathway | 0.00E+00 | 1.15E-02 | NaN | IL1B |
| Xenobiotic Metabolism PXR Signaling Pathway | 0.00E+00 | 2.07E-02 | 2 | CHST1,CHST3,HS3ST3B1,SULT1B1 |
| Insulin Secretion Signaling Pathway | 0.00E+00 | 1.84E-02 | NaN | ADCY10,CACNA1G,CACNA2D3,NOTUM,STX1A |
| Coronavirus Pathogenesis Pathway | 0.00E+00 | 1.47E-02 | NaN | EEF1A2,IL1B,SERPINE1 |
| MSP-RON Signaling In Cancer Cells Pathway | 0.00E+00 | 1.43E-02 | NaN | ELF5,MET |
| MSP-RON Signaling In Macrophages Pathway | 0.00E+00 | 8.40E-03 | NaN | IL10 |
| Role of MAPK Signaling in Promoting the Pathog | 0.00E+00 | 1.75E-02 | NaN | PLA2G1B,PLAAT4 |

IPA_Motor_Neurons_GCSi_FC2_adj0.05_576

| | Ingenuity Canonical Pathways | -log(p-value) | Ratio | z-score | Molecules |
|---|---|---|---|---|---|
| 1 | Glutamate Degradation II | 1.84E+00 | 3.33E-01 | NaN | GOT1L1 |
| 2 | Aspartate Biosynthesis | 1.84E+00 | 3.33E-01 | NaN | GOT1L1 |
| 3 | L-cysteine Degradation I | 1.72E+00 | 2.50E-01 | NaN | GOT1L1 |
| 4 | Neuroprotective Role of THOP1 in Alzheimer's Dise | 1.68E+00 | 2.48E-02 | NaN | CTRL,F11,FAP |
| 5 | LXR/RXR Activation | 1.66E+00 | 2.44E-02 | NaN | IL18RAP,IL36RN,S100A8 |
| 6 | FXR/RXR Activation | 1.63E+00 | 2.38E-02 | NaN | FBP1,G6PC1,IL36RN |
| 7 | Osteoarthritis Pathway | 1.55E+00 | 1.69E-02 | NaN | ADIPOQ,H19,IL18RAP,S100A8 |
| 8 | Role of Cytokines in Mediating Communication bet | 1.55E+00 | 3.70E-02 | NaN | IFNL1,IL36RN |
| 9 | Serotonin and Melatonin Biosynthesis | 1.54E+00 | 1.67E-01 | NaN | TPH2 |
| 10 | Aspartate Degradation II | 1.48E+00 | 1.43E-01 | NaN | GOT1L1 |
| | Mineralocorticoid Biosynthesis | 1.25E+00 | 8.33E-02 | NaN | HSD3B2 |
| | Glucocorticoid Biosynthesis | 1.21E+00 | 7.69E-02 | NaN | HSD3B2 |
| | Granulocyte Adhesion and Diapedesis | 1.19E+00 | 1.59E-02 | NaN | CCL15,IL18RAP,IL36RN |
| | Role of Hypercytokinemia/hyperchemokinemia in t | 1.18E+00 | 2.33E-02 | NaN | IFNL1,IL36RN |
| | Role of IL-17A in Psoriasis | 1.18E+00 | 7.14E-02 | NaN | S100A8 |
| | Phenylalanine Degradation IV (Mammalian, via Sid | 1.18E+00 | 7.14E-02 | NaN | GOT1L1 |
| | Hepatic Fibrosis / Hepatic Stellate Cell Activation | 1.17E+00 | 1.55E-02 | NaN | IGF2,IL18RAP,MYL4 |
| | Agranulocyte Adhesion and Diapedesis | 1.09E+00 | 1.43E-02 | NaN | CCL15,IL36RN,MYL4 |
| | Androgen Biosynthesis | 1.03E+00 | 5.00E-02 | NaN | HSD3B2 |
| | PPAR Signaling | 1.02E+00 | 1.87E-02 | NaN | IL18RAP,IL36RN |
| | Pathogen Induced Cytokine Storm Signaling Pathw | 9.74E-01 | 1.08E-02 | 2 | CCL15,IL18RAP,IL36RN,STING1 |
| | Glycolysis I | 9.42E-01 | 4.00E-02 | NaN | FBP1 |
| | Gluconeogenesis I | 9.42E-01 | 4.00E-02 | NaN | FBP1 |
| | p38 MAPK Signaling | 9.39E-01 | 1.67E-02 | NaN | IL18RAP,IL36RN |
| | IL-6 Signaling | 8.88E-01 | 1.55E-02 | NaN | IL18RAP,IL36RN |
| | Atherosclerosis Signaling | 8.66E-01 | 1.50E-02 | NaN | IL36RN,S100A8 |
| | Role Of Chondrocytes In Rheumatoid Arthritis Sign | 8.26E-01 | 1.42E-02 | NaN | IL18RAP,IL36RN |
| | Glucocorticoid Receptor Signaling | 8.17E-01 | 8.59E-03 | NaN | FBP1,G6PC1,IL18RAP,KRT2,KRT34 |
| | Coagulation System | 8.06E-01 | 2.86E-02 | NaN | F11 |
| | Complement System | 7.84E-01 | 2.70E-02 | NaN | CFI |
| | IL-10 Signaling | 7.66E-01 | 1.30E-02 | NaN | IL18RAP,IL36RN |
| | Intrinsic Prothrombin Activation Pathway | 7.34E-01 | 2.38E-02 | NaN | F11 |
| | Superpathway of Methionine Degradation | 7.34E-01 | 2.38E-02 | NaN | GOT1L1 |
| | Serotonin Receptor Signaling | 7.14E-01 | 8.53E-03 | 1 | ADIPOQ,FBP1,G6PC1,TPH2 |
| | PFKFB4 Signaling Pathway | 6.82E-01 | 2.08E-02 | NaN | FBP1 |
| | Tumor Microenvironment Pathway | 6.67E-01 | 1.12E-02 | NaN | CTLA4,IGF2 |
| | Hepatic Cholestasis | 6.26E-01 | 1.05E-02 | NaN | IL18RAP,IL36RN |
| | PPARα/RXRα Activation | 6.13E-01 | 1.03E-02 | NaN | ADIPOQ,IL18RAP |
| | MSP-RON Signaling Pathway | 6.10E-01 | 1.72E-02 | NaN | F11 |
| | SPINK1 Pancreatic Cancer Pathway | 5.97E-01 | 1.67E-02 | NaN | CTRL |
| | PXR/RXR Activation | 5.67E-01 | 1.54E-02 | NaN | G6PC1 |
| | Pyridoxal 5'-phosphate Salvage Pathway | 5.67E-01 | 1.54E-02 | NaN | G6PC1 |
| | Glutamate Receptor Signaling | 5.61E-01 | 1.52E-02 | NaN | SLC1A3 |
| | Activin Inhibin Signaling Pathway | 5.48E-01 | 9.22E-03 | NaN | IL18RAP,IL36RN |
| | Calcium Signaling | 5.40E-01 | 9.09E-03 | NaN | CASQ2,MYL4 |
| | Growth Hormone Signaling | 5.35E-01 | 1.41E-02 | NaN | IGF2 |
| | Netrin Signaling | 5.30E-01 | 1.39E-02 | NaN | NTN1 |
| | Role of Osteoblasts, Osteoclasts and Chondrocytes | 5.19E-01 | 8.77E-03 | NaN | IL18RAP,IL36RN |
| | Toll-like Receptor Signaling | 5.01E-01 | 1.28E-02 | NaN | IL36RN |
| | Maturity Onset Diabetes of Young (MODY) Signalin | 4.96E-01 | 1.27E-02 | NaN | ADIPOQ |
| | S100 Family Signaling Pathway | 4.96E-01 | 6.48E-03 | 2.236 | HCAR1,S100A11,S100A7A,S100A8,S100Z |
| | Lipid Antigen Presentation by CD1 | 4.87E-01 | 7.23E-03 | NaN | CD1B,TRBV4-1,TRBV6-7 |
| | Dendritic Cell Maturation | 4.85E-01 | 6.71E-03 | NaN | CD1B,IL36RN,TRBV4-1,TRBV6-7 |
| | TR/RXR Activation | 4.74E-01 | 1.19E-02 | NaN | G6PC1 |
| | Hepatic Fibrosis Signaling Pathway | 4.73E-01 | 7.09E-03 | NaN | IL18RAP,IL36RN,MYL4 |
| | Wound Healing Signaling Pathway | 4.62E-01 | 7.94E-03 | NaN | IL18RAP,IL36RN |
| | LPS/IL-1 Mediated Inhibition of RXR Function | 4.55E-01 | 7.84E-03 | NaN | IL18RAP,IL36RN |
| | Immunogenic Cell Death Signaling Pathway | 4.50E-01 | 1.11E-02 | NaN | STING1 |
| | Cardiac Hypertrophy Signaling | 4.42E-01 | 7.66E-03 | NaN | ELSPBP1,MYL4 |
| | Graft-versus-Host Disease Signaling | 4.35E-01 | 6.73E-03 | NaN | IL36RN,TRBV4-1,TRBV6-7 |
| | Salvage Pathways of Pyrimidine Ribonucleotides | 4.25E-01 | 1.03E-02 | NaN | G6PC1 |
| | Protein Ubiquitination Pathway | 4.18E-01 | 7.33E-03 | NaN | ODF1,USP29 |
| | Apelin Cardiomyocyte Signaling Pathway | 4.18E-01 | 1.01E-02 | NaN | MYL4 |

| | | | | |
|---|---|---|---|---|
| Mouse Embryonic Stem Cell Pluripotency | 4.01E-01 | 9.62E-03 | NaN | ZFP42 |
| Regulation of Actin-based Motility by Rho | 3.68E-01 | 8.70E-03 | NaN | MYL4 |
| PAK Signaling | 3.62E-01 | 8.55E-03 | NaN | MYL4 |
| Cholecystokinin/Gastrin-mediated Signaling | 3.57E-01 | 8.40E-03 | NaN | IL36RN |
| MSP-RON Signaling In Macrophages Pathway | 3.57E-01 | 8.40E-03 | NaN | F11 |
| Role of NANOG in Mammalian Embryonic Stem Cel | 3.44E-01 | 8.06E-03 | NaN | ZFP42 |
| RHOA Signaling | 3.44E-01 | 8.06E-03 | NaN | MYL4 |
| Glioma Signaling | 3.41E-01 | 8.00E-03 | NaN | IGF2 |
| CD28 Signaling in T Helper Cells | 3.38E-01 | 5.80E-03 | NaN | CTLA4,TRBV4-1,TRBV6-7 |
| Communication between Innate and Adaptive Imm | 3.25E-01 | 5.36E-03 | NaN | CCL15,IGLV1-44,IL36RN,TRBV4-1,TRBV6-7 |
| Gα12/13 Signaling | 3.22E-01 | 7.52E-03 | NaN | MYL4 |
| Role of Macrophages, Fibroblasts and Endothelial C | 3.19E-01 | 6.02E-03 | NaN | IL18RAP,IL36RN |
| STAT3 Pathway | 3.17E-01 | 7.41E-03 | NaN | IL18RAP |
| SNARE Signaling Pathway | 3.15E-01 | 7.35E-03 | NaN | MYL4 |
| MSP-RON Signaling In Cancer Cells Pathway | 3.06E-01 | 7.14E-03 | NaN | F11 |
| Dilated Cardiomyopathy Signaling Pathway | 2.85E-01 | 6.67E-03 | NaN | MYL4 |
| Cellular Effects of Sildenafil (Viagra) | 2.85E-01 | 6.67E-03 | NaN | MYL4 |
| Semaphorin Neuronal Repulsive Signaling Pathway | 2.85E-01 | 6.67E-03 | NaN | MYL4 |
| T Cell Exhaustion Signaling Pathway | 2.84E-01 | 5.29E-03 | NaN | CTLA4,TRBV4-1,TRBV6-7 |
| NF-κB Signaling | 2.80E-01 | 5.25E-03 | NaN | IL36RN,TRBV4-1,TRBV6-7 |
| Type II Diabetes Mellitus Signaling | 2.80E-01 | 6.54E-03 | NaN | ADIPOQ |
| CDC42 Signaling | 2.75E-01 | 5.21E-03 | NaN | MYL4,TRBV4-1,TRBV6-7 |
| Aryl Hydrocarbon Receptor Signaling | 2.68E-01 | 6.29E-03 | NaN | TFF1 |
| Transcriptional Regulatory Network in Embryonic S | 2.59E-01 | 6.10E-03 | NaN | IGF2 |
| CXCR4 Signaling | 2.53E-01 | 5.95E-03 | NaN | MYL4 |
| Glioblastoma Multiforme Signaling | 2.48E-01 | 5.85E-03 | NaN | IGF2 |
| Aldosterone Signaling in Epithelial Cells | 2.46E-01 | 5.81E-03 | NaN | ODF1 |
| CTLA4 Signaling in Cytotoxic T Lymphocytes | 2.46E-01 | 4.93E-03 | NaN | CTLA4,TRBV4-1,TRBV6-7 |
| T Cell Receptor Signaling | 2.39E-01 | 4.87E-03 | NaN | CTLA4,TRBV4-1,TRBV6-7 |
| Tight Junction Signaling | 2.35E-01 | 5.59E-03 | NaN | MYL4 |
| D-myo-inositol (1,4,5,6)-Tetrakisphosphate Biosynt | 2.34E-01 | 5.56E-03 | NaN | G6PC1 |
| D-myo-inositol (3,4,5,6)-tetrakisphosphate Biosyntl | 2.34E-01 | 5.56E-03 | NaN | G6PC1 |
| Estrogen Receptor Signaling | 2.28E-01 | 4.89E-03 | NaN | IGF2,MYL4 |
| Acute Phase Response Signaling | 2.26E-01 | 5.41E-03 | NaN | IL36RN |
| Protein Kinase A Signaling | 2.26E-01 | 4.87E-03 | NaN | MYL4,NTN1 |
| IL-33 Signaling Pathway | 2.25E-01 | 5.38E-03 | NaN | IL36RN |
| Macrophage Classical Activation Signaling Pathway | 2.20E-01 | 5.29E-03 | NaN | CCL15 |
| Macrophage Alternative Activation Signaling Pathw | 2.20E-01 | 5.29E-03 | NaN | IL36RN |
| Production of Nitric Oxide and Reactive Oxygen Spe | 2.18E-01 | 5.24E-03 | NaN | S100A8 |
| 3-phosphoinositide Degradation | 2.18E-01 | 5.24E-03 | NaN | G6PC1 |
| D-myo-inositol-5-phosphate Metabolism | 2.11E-01 | 5.10E-03 | NaN | G6PC1 |
| Natural Killer Cell Signaling | 2.08E-01 | 5.05E-03 | NaN | IL18RAP |
| Pulmonary Healing Signaling Pathway | 2.07E-01 | 5.03E-03 | NaN | mir-183 |
| Adrenomedullin signaling pathway | 2.07E-01 | 5.03E-03 | NaN | IL36RN |
| PI3K/AKT Signaling | 2.06E-01 | 5.00E-03 | NaN | IL18RAP |
| ID1 Signaling Pathway | 2.04E-01 | 4.98E-03 | NaN | IGF2 |
| ILK Signaling | 2.04E-01 | 4.98E-03 | NaN | MYL4 |
| Coronavirus Pathogenesis Pathway | 2.01E-01 | 4.90E-03 | NaN | STING1 |
| 3-phosphoinositide Biosynthesis | 1.98E-01 | 4.85E-03 | NaN | G6PC1 |
| Neurovascular Coupling Signaling Pathway | 0.00E+00 | 4.31E-03 | NaN | SLC1A3 |
| Oxytocin Signaling Pathway | 0.00E+00 | 3.55E-03 | NaN | MYL4 |
| Multiple Sclerosis Signaling Pathway | 0.00E+00 | 4.50E-03 | NaN | CTLA4 |
| Role Of Osteoclasts In Rheumatoid Arthritis Signalii | 0.00E+00 | 3.24E-03 | NaN | IL18RAP |
| Axonal Guidance Signaling | 0.00E+00 | 3.92E-03 | NaN | MYL4,NTN1 |
| Actin Cytoskeleton Signaling | 0.00E+00 | 4.10E-03 | NaN | MYL4 |
| Chaperone Mediated Autophagy Signaling Pathway | 0.00E+00 | 4.67E-03 | NaN | STING1,TRBV4-1,TRBV6-7 |
| Clathrin-mediated Endocytosis Signaling | 0.00E+00 | 4.81E-03 | NaN | S100A8 |
| IL-12 Signaling and Production in Macrophages | 0.00E+00 | 4.24E-03 | NaN | S100A8 |
| Role of NFAT in Regulation of the Immune Respons | 0.00E+00 | 2.89E-03 | NaN | IGLV1-44,TRBV4-1,TRBV6-7 |
| FcγRIIB Signaling in B Lymphocytes | 0.00E+00 | 1.88E-03 | NaN | IGLV1-44 |
| CCR5 Signaling in Macrophages | 0.00E+00 | 4.01E-03 | NaN | TRBV4-1,TRBV6-7 |
| Calcium-induced T Lymphocyte Apoptosis | 0.00E+00 | 4.35E-03 | NaN | TRBV4-1,TRBV6-7 |
| Cytotoxic T Lymphocyte-mediated Apoptosis of Tar | 0.00E+00 | 4.71E-03 | NaN | TRBV4-1,TRBV6-7 |
| T Helper Cell Differentiation | 0.00E+00 | 4.25E-03 | NaN | TRBV4-1,TRBV6-7 |
| IL-15 Signaling | 0.00E+00 | 1.89E-03 | NaN | IGLV1-44 |
| HIF1α Signaling | 0.00E+00 | 4.81E-03 | NaN | IGF2 |
| Thrombin Signaling | 0.00E+00 | 4.44E-03 | NaN | MYL4 |

| | | | | | |
|---|---|---|---|---|---|
| ICOS-ICOSL Signaling in T Helper Cells | 0.00E+00 | 3.94E-03 | NaN | | TRBV4-1,TRBV6-7 |
| CREB Signaling in Neurons | 0.00E+00 | 1.65E-03 | NaN | | HCAR1 |
| Type I Diabetes Mellitus Signaling | 0.00E+00 | 3.93E-03 | NaN | | TRBV4-1,TRBV6-7 |
| Allograft Rejection Signaling | 0.00E+00 | 4.11E-03 | NaN | | TRBV4-1,TRBV6-7 |
| Autoimmune Thyroid Disease Signaling | 0.00E+00 | 4.37E-03 | NaN | | TRBV4-1,TRBV6-7 |
| p70S6K Signaling | 0.00E+00 | 1.72E-03 | NaN | | IGLV1-44 |
| G Protein Signaling Mediated by Tubby | 0.00E+00 | 4.27E-03 | NaN | | TRBV4-1,TRBV6-7 |
| Systemic Lupus Erythematosus Signaling | 0.00E+00 | 3.75E-03 | NaN | | IGLV1-44,IL36RN,TRBV4-1,TRBV6-7 |
| FAK Signaling | 0.00E+00 | 3.84E-03 | | 2 | HCAR1,IL18RAP,TRBV4-1,TRBV6-7 |
| AMPK Signaling | 0.00E+00 | 4.13E-03 | NaN | | ADIPOQ |
| Phospholipase C Signaling | 0.00E+00 | 3.57E-03 | NaN | | IGLV1-44,MYL4,TRBV4-1,TRBV6-7 |
| Altered T Cell and B Cell Signaling in Rheumatoid A | 0.00E+00 | 4.26E-03 | NaN | | IGLV1-44,IL36RN,TRBV4-1,TRBV6-7 |
| B Cell Development | 0.00E+00 | 2.04E-03 | NaN | | IGLV1-44 |
| Breast Cancer Regulation by Stathmin1 | 0.00E+00 | 1.68E-03 | NaN | | HCAR1 |
| Regulation of IL-2 Expression in Activated and Aner | 0.00E+00 | 4.33E-03 | NaN | | TRBV4-1,TRBV6-7 |
| NUR77 Signaling in T Lymphocytes | 0.00E+00 | 3.90E-03 | NaN | | TRBV4-1,TRBV6-7 |
| PKCθ Signaling in T Lymphocytes | 0.00E+00 | 3.58E-03 | NaN | | TRBV4-1,TRBV6-7 |
| Antiproliferative Role of TOB in T Cell Signaling | 0.00E+00 | 4.69E-03 | NaN | | TRBV4-1,TRBV6-7 |
| OX40 Signaling Pathway | 0.00E+00 | 4.18E-03 | NaN | | TRBV4-1,TRBV6-7 |
| PI3K Signaling in B Lymphocytes | 0.00E+00 | 3.38E-03 | NaN | | CD180,IGLV1-44 |
| Signaling by Rho Family GTPases | 0.00E+00 | 3.75E-03 | NaN | | MYL4 |
| RHOGDI Signaling | 0.00E+00 | 4.55E-03 | NaN | | MYL4 |
| Hematopoiesis from Pluripotent Stem Cells | 0.00E+00 | 4.49E-03 | NaN | | TRBV4-1,TRBV6-7 |
| Superpathway of Inositol Phosphate Compounds | 0.00E+00 | 4.27E-03 | NaN | | G6PC1 |
| TEC Kinase Signaling | 0.00E+00 | 3.46E-03 | NaN | | TRBV4-1,TRBV6-7 |
| SAPK/JNK Signaling | 0.00E+00 | 3.98E-03 | NaN | | TRBV4-1,TRBV6-7 |
| IL-4 Signaling | 0.00E+00 | 3.47E-03 | NaN | | TRBV4-1,TRBV6-7 |
| B Cell Receptor Signaling | 0.00E+00 | 1.57E-03 | NaN | | IGLV1-44 |
| G-Protein Coupled Receptor Signaling | 0.00E+00 | 2.84E-03 | NaN | | HCAR1,MYL4 |
| Phagosome Formation | 0.00E+00 | 2.88E-03 | NaN | | HCAR1,MYL4 |
| Neuroinflammation Signaling Pathway | 0.00E+00 | 3.15E-03 | NaN | | SLC1A3 |
| Th17 Activation Pathway | 0.00E+00 | 4.13E-03 | NaN | | TRBV4-1,TRBV6-7 |
| Cardiac Hypertrophy Signaling (Enhanced) | 0.00E+00 | 1.85E-03 | NaN | | IL18RAP |
| Systemic Lupus Erythematosus In T Cell Signaling P | 0.00E+00 | 3.11E-03 | NaN | | TRBV4-1,TRBV6-7 |
| Systemic Lupus Erythematosus In B Cell Signaling P | 0.00E+00 | 2.76E-03 | NaN | | IGLV1-44,STING1 |
| Senescence Pathway | 0.00E+00 | 3.34E-03 | NaN | | STING1 |

IPA_Venn_List_Micro_FB1_FC2_adj0.05_vs._Micro_GCS_FC2_adj0.05_278genes

| | Ingenuity Canonical Pathways | -log(p-valı | Ratio | z-score | Molecules |
|---|---|---|---|---|---|
| 1 | Granulocyte Adhesion and Diapedesis | 6.78E+00 | 5.29E-02 | NaN | CCR7,CLDN4,CLDN5,CXCL3,CXCL5,CXCR4,IL1B,MMP21,MMP3,NGFR |
| 2 | Agranulocyte Adhesion and Diapedesis | 6.36E+00 | 4.76E-02 | NaN | CCR7,CLDN4,CLDN5,CXCL3,CXCL5,CXCR4,IL1B,MMP21,MMP3,MYL4 |
| 3 | Tumor Microenvironment Pathway | 4.99E+00 | 4.47E-02 | NaN | CXCR4,FOXG1,HGF,IGF2,IL1B,MMP21,MMP3,TNC |
| 4 | Hepatic Fibrosis / Hepatic Stellate Cell Activation | 4.73E+00 | 4.12E-02 | NaN | CCR7,COL6A5,CXCL3,HGF,IGF2,IL1B,MYL4,NGFR |
| 5 | S100 Family Signaling Pathway | 4.35E+00 | 1.94E-02 | NaN | ADGRB2,ADGRD2,ADGRL3,AREG,CCR7,EREG,GLP2R,HTR4,IL1B,LHCGR,MMP21,MMP3,NOTUM,NTF3,NTSR1 |
| 6 | CREB Signaling in Neurons | 4.25E+00 | 2.14E-02 | NaN | ADGRB2,ADGRD2,ADGRL3,BMP6,CCR7,GLP2R,GRIA2,HGF,HTR4,LHCGR,NGFR,NOTUM,NTSR1 |
| 7 | Axonal Guidance Signaling | 3.09E+00 | 1.96E-02 | NaN | ADAMTS9,BMP6,CXCR4,EPHA10,MMP21,MMP3,MYL4,NGFR,NOTUM,NTF3 |
| 8 | Role Of Chondrocytes In Rheumatoid Arthritis Signaling Pa | 2.85E+00 | 3.55E-02 | NaN | CXCR4,IL1B,MMP21,MMP3,NGFR |
| 9 | Inhibition of Matrix Metalloproteases | 2.82E+00 | 7.69E-02 | NaN | MMP21,MMP3,TFPI2 |
| 10 | Heparan Sulfate Biosynthesis | 2.78E+00 | 4.60E-02 | NaN | B3GAT1,CHST1,NOTUM,SULT1B1 |
| | Role of Osteoblasts, Osteoclasts and Chondrocytes in Rhe | 2.66E+00 | 2.63E-02 | NaN | BMP6,DKK3,IL11,IL1B,MMP3,NGFR |
| | Breast Cancer Regulation by Stathmin1 | 2.60E+00 | 1.68E-02 | NaN | ADGRB2,ADGRD2,ADGRL3,BMP6,CCR7,GLP2R,HGF,HTR4,LHCGR,NTSR1 |
| | Role of IL-17F in Allergic Inflammatory Airway Diseases | 2.59E+00 | 6.38E-02 | NaN | CXCL5,IL11,IL1B |
| | Role of IL-17A in Psoriasis | 2.54E+00 | 1.43E-01 | NaN | CXCL3,CXCL5 |
| | Pulmonary Fibrosis Idiopathic Signaling Pathway | 2.52E+00 | 2.15E-02 | NaN | AREG,COL6A5,FOXG1,IL11,IL1B,MMP21,MMP3 |
| | IL-17 Signaling | 2.32E+00 | 2.67E-02 | NaN | CXCL3,CXCL5,IL11,IL1B,MMP3 |
| | Chondroitin Sulfate Biosynthesis | 2.31E+00 | 5.08E-02 | NaN | B3GAT1,CHST1,SULT1B1 |
| | Dermatan Sulfate Biosynthesis | 2.27E+00 | 4.92E-02 | NaN | B3GAT1,CHST1,SULT1B1 |
| | Leukocyte Extravasation Signaling | 2.26E+00 | 2.59E-02 | NaN | CLDN4,CLDN5,CXCR4,MMP21,MMP3 |
| | Pathogen Induced Cytokine Storm Signaling Pathway | 2.22E+00 | 1.89E-02 | NaN | COL6A5,CXCL3,CXCL5,CXCR4,IL11,IL1B,NGFR |
| | HIF1α Signaling | 2.13E+00 | 2.40E-02 | NaN | BMP6,HGF,IGF2,MMP21,MMP3 |
| | Phagosome Formation | 2.13E+00 | 1.44E-02 | NaN | ADGRB2,ADGRD2,ADGRL3,CCR7,FCER1A,GLP2R,HTR4,LHCGR,MYL4,NTSR1 |
| | STAT3 Pathway | 2.09E+00 | 2.96E-02 | NaN | BMP6,HGF,IL1B,NGFR |
| | G-Protein Coupled Receptor Signaling | 2.09E+00 | 1.42E-02 | NaN | ADGRB2,ADGRD2,ADGRL3,CCR7,FOXG1,GLP2R,HTR4,LHCGR,MYL4,NTSR1 |
| | Activin Inhibin Signaling Pathway | 2.05E+00 | 2.30E-02 | NaN | CXCR4,FOSB,IL11,IL1B,NGFR |
| | Role Of Osteoclasts In Rheumatoid Arthritis Signaling Path | 2.02E+00 | 1.94E-02 | NaN | COL6A5,FOXG1,IL1B,MMP21,MMP3,NGFR |
| | NAD Salvage Pathway II | 1.97E+00 | 7.41E-02 | NaN | NMNAT2,NT5E |
| | Heparan Sulfate Biosynthesis (Late Stages) | 1.94E+00 | 3.75E-02 | NaN | CHST1,NOTUM,SULT1B1 |
| | NAD Signaling Pathway | 1.93E+00 | 2.65E-02 | NaN | BMP6,HGF,NMNAT2,NT5E |
| | Role of Hypercytokinemia/hyperchemokinemia in the Patl | 1.86E+00 | 3.49E-02 | NaN | AREG,CXCL3,IL1B |
| | Wound Healing Signaling Pathway | 1.79E+00 | 1.98E-02 | NaN | COL6A5,IL11,IL1B,LAMA5,NGFR |
| | Tight Junction Signaling | 1.68E+00 | 2.23E-02 | NaN | CLDN4,CLDN5,MYL4,NGFR |
| | Regulation Of The Epithelial Mesenchymal Transition By G | 1.59E+00 | 2.08E-02 | NaN | HGF,HMGA2,NGFR,TWIST1 |
| | Pulmonary Healing Signaling Pathway | 1.54E+00 | 2.01E-02 | NaN | CXCR4,MMP21,MMP3,NGFR |
| | CDK5 Signaling | 1.53E+00 | 2.61E-02 | NaN | FOSB,LAMA5,NGFR |
| | CDX Gastrointestinal Cancer Signaling Pathway | 1.52E+00 | 1.98E-02 | NaN | BMP6,GATA6,IL11,IL1B |
| | Ephrin A Signaling | 1.51E+00 | 4.26E-02 | NaN | EPHA10,NGFR |
| | Airway Pathology in Chronic Obstructive Pulmonary Disea | 1.50E+00 | 2.54E-02 | NaN | CXCL3,IL11,IL1B |
| | Dermatan Sulfate Biosynthesis (Late Stages) | 1.50E+00 | 4.17E-02 | NaN | CHST1,SULT1B1 |
| | Estrogen Receptor Signaling | 1.49E+00 | 1.47E-02 | NaN | FOXG1,IGF2,MMP21,MMP3,MYL4,NOTUM |
| | FAT10 Cancer Signaling Pathway | 1.47E+00 | 4.00E-02 | NaN | CXCR4,NGFR |
| | UDP-N-acetyl-D-glucosamine Biosynthesis II | 1.46E+00 | 1.67E-01 | NaN | GFPT2 |
| | NAD Biosynthesis III | 1.46E+00 | 1.67E-01 | NaN | NMNAT2 |
| | Chondroitin Sulfate Biosynthesis (Late Stages) | 1.45E+00 | 3.92E-02 | NaN | CHST1,SULT1B1 |
| | NAD Biosynthesis from 2-amino-3-carboxymuconate Semi | 1.40E+00 | 1.43E-01 | NaN | NMNAT2 |
| | Ceramide Biosynthesis | 1.40E+00 | 1.43E-01 | NaN | DEGS2 |
| | NAD Salvage Pathway III | 1.40E+00 | 1.43E-01 | NaN | NMNAT2 |
| | Glycoaminoglycan-protein Linkage Region Biosynthesis | 1.40E+00 | 1.43E-01 | NaN | B3GAT1 |
| | Myelination Signaling Pathway | 1.37E+00 | 1.53E-02 | NaN | BMP6,LAMA5,NGFR,NTF3,PLP1 |
| | Atherosclerosis Signaling | 1.37E+00 | 2.26E-02 | NaN | CXCR4,IL1B,MMP3 |
| | Role of IL-17A in Arthritis | 1.36E+00 | 3.51E-02 | NaN | CXCL3,CXCL5 |
| | Role of Macrophages, Fibroblasts and Endothelial Cells in I | 1.35E+00 | 1.51E-02 | NaN | DKK3,IL1B,MMP3,NGFR,NOTUM |
| | Role Of Osteoblasts In Rheumatoid Arthritis Signaling Path | 1.26E+00 | 1.64E-02 | NaN | IL11,IL1B,MMP21,MMP3 |
| | Induction of Apoptosis by HIV1 | 1.26E+00 | 3.08E-02 | NaN | CXCR4,NGFR |
| | Embryonic Stem Cell Differentiation into Cardiac Lineages | 1.25E+00 | 1.00E-01 | NaN | NKX2-5 |
| | IL-17A Signaling in Airway Cells | 1.24E+00 | 2.99E-02 | NaN | CXCL3,CXCL5 |
| | Factors Promoting Cardiogenesis in Vertebrates | 1.22E+00 | 1.96E-02 | NaN | BMP6,NKX2-5,NOTUM |
| | LPS/IL-1 Mediated Inhibition of RXR Function | 1.21E+00 | 1.57E-02 | NaN | CHST1,IL1B,NGFR,SULT1B1 |
| | HOTAIR Regulatory Pathway | 1.16E+00 | 1.84E-02 | NaN | MMP21,MMP3,TWIST1 |
| | GPCR-Mediated Integration of Enteroendocrine Signaling | 1.15E+00 | 2.67E-02 | NaN | GLP2R,NOTUM |
| | NAD biosynthesis II (from tryptophan) | 1.14E+00 | 7.69E-02 | NaN | NMNAT2 |
| | Guanosine Nucleotides Degradation III | 1.14E+00 | 7.69E-02 | NaN | NT5E |
| | HMGB1 Signaling | 1.13E+00 | 1.80E-02 | NaN | IL11,IL1B,NGFR |
| | TREM1 Signaling | 1.13E+00 | 2.60E-02 | NaN | CXCL3,IL1B |

| | | | | |
|---|---|---|---|---|
| Neurotrophin/TRK Signaling | 1.12E+00 | 2.56E-02 | NaN | NGFR,NTF3 |
| IL-7 Signaling Pathway | 1.12E+00 | 2.56E-02 | NaN | FOXG1,HGF |
| Thyroid Cancer Signaling | 1.11E+00 | 2.53E-02 | NaN | CXCR4,NTF3 |
| Urate Biosynthesis/Inosine 5'-phosphate Degradation | 1.11E+00 | 7.14E-02 | NaN | NT5E |
| Extrinsic Prothrombin Activation Pathway | 1.05E+00 | 6.25E-02 | NaN | F13A1 |
| Adenosine Nucleotides Degradation II | 1.05E+00 | 6.25E-02 | NaN | NT5E |
| Immunogenic Cell Death Signaling Pathway | 1.02E+00 | 2.22E-02 | NaN | IL1B,NGFR |
| BMP signaling pathway | 1.01E+00 | 2.20E-02 | NaN | BMP6,NKX2-5 |
| Differential Regulation of Cytokine Production in Macroph | 1.00E+00 | 5.56E-02 | NaN | IL1B |
| Hepatic Cholestasis | 9.98E-01 | 1.57E-02 | NaN | IL11,IL1B,NGFR |
| Pyroptosis Signaling Pathway | 9.91E-01 | 2.15E-02 | NaN | IL1B,NGFR |
| ERBB Signaling | 9.91E-01 | 2.15E-02 | NaN | AREG,EREG |
| Purine Nucleotides Degradation II (Aerobic) | 9.79E-01 | 5.26E-02 | NaN | NT5E |
| Regulation of the Epithelial-Mesenchymal Transition Path | 9.78E-01 | 1.54E-02 | NaN | HGF,HMGA2,TWIST1 |
| Inflammasome pathway | 9.58E-01 | 5.00E-02 | NaN | IL1B |
| Apelin Cardiomyocyte Signaling Pathway | 9.46E-01 | 2.02E-02 | NaN | MYL4,NOTUM |
| Neuroinflammation Signaling Pathway | 9.44E-01 | 1.26E-02 | NaN | GABRQ,IL1B,MMP3,NTF3 |
| Gustation Pathway | 9.40E-01 | 1.48E-02 | NaN | GABRQ,NOTUM,SCN7A |
| Neuropathic Pain Signaling In Dorsal Horn Neurons | 9.32E-01 | 1.98E-02 | NaN | GRIA2,NOTUM |
| Role of Tissue Factor in Cancer | 9.21E-01 | 1.45E-02 | NaN | HGF,IL1B,NGFR |
| Differential Regulation of Cytokine Production in Intestina | 9.01E-01 | 4.35E-02 | NaN | IL1B |
| Pyrimidine Deoxyribonucleotides De Novo Biosynthesis I | 9.01E-01 | 4.35E-02 | NaN | AK7 |
| PPAR Signaling | 8.91E-01 | 1.87E-02 | NaN | IL1B,NGFR |
| Tumoricidal Function of Hepatic Natural Killer Cells | 8.84E-01 | 4.17E-02 | NaN | LYVE1 |
| Autophagy | 8.81E-01 | 1.39E-02 | NaN | BMP6,HGF,NGFR |
| Serotonin Receptor Signaling | 8.59E-01 | 1.07E-02 | NaN | CCR7,F13A1,HTR4,IL1B,NOTUM |
| Multiple Sclerosis Signaling Pathway | 8.56E-01 | 1.35E-02 | NaN | IL11,IL1B,PLP1 |
| Role of NFAT in Cardiac Hypertrophy | 8.47E-01 | 1.34E-02 | NaN | IL11,NKX2-5,NOTUM |
| Thrombin Signaling | 8.43E-01 | 1.33E-02 | NaN | GATA6,MYL4,NOTUM |
| Cardiomyocyte Differentiation via BMP Receptors | 8.37E-01 | 3.70E-02 | NaN | NKX2-5 |
| Bladder Cancer Signaling | 8.35E-01 | 1.72E-02 | NaN | MMP21,MMP3 |
| Neuregulin Signaling | 8.29E-01 | 1.71E-02 | NaN | AREG,EREG |
| FAK Signaling | 8.16E-01 | 8.65E-03 | NaN | ADGRB2,ADGRD2,ADGRL3,CCR7,GLP2R,HTR4,LHCGR,NTSR1,TRBV5-4 |
| Osteoarthritis Pathway | 8.00E-01 | 1.27E-02 | NaN | IL1B,MMP3,NKX3-2 |
| LXR/RXR Activation | 7.95E-01 | 1.63E-02 | NaN | IL1B,NGFR |
| Role of NANOG in Mammalian Embryonic Stem Cell Plurip | 7.89E-01 | 1.61E-02 | NaN | BMP6,GATA6 |
| FXR/RXR Activation | 7.78E-01 | 1.59E-02 | NaN | FOXA2,IL1B |
| Gαs Signaling | 7.78E-01 | 1.59E-02 | NaN | HTR4,LHCGR |
| GP6 Signaling Pathway | 7.73E-01 | 1.57E-02 | NaN | COL6A5,LAMA5 |
| Dopamine Degradation | 7.69E-01 | 3.12E-02 | NaN | SULT1B1 |
| IL-6 Signaling | 7.63E-01 | 1.55E-02 | NaN | IL1B,NGFR |
| Synaptic Long Term Potentiation | 7.47E-01 | 1.52E-02 | NaN | GRIA2,NOTUM |
| Coagulation System | 7.34E-01 | 2.86E-02 | NaN | F13A1 |
| SNARE Signaling Pathway | 7.27E-01 | 1.47E-02 | NaN | MYL4,SYN3 |
| Cardiac Hypertrophy Signaling | 7.11E-01 | 1.15E-02 | NaN | MYL4,NKX2-5,NOTUM |
| Docosahexaenoic Acid (DHA) Signaling | 7.02E-01 | 2.63E-02 | NaN | IL1B |
| IL-17A Signaling in Fibroblasts | 7.02E-01 | 2.63E-02 | NaN | CXCL5 |
| Pyrimidine Ribonucleotides Interconversion | 6.91E-01 | 2.56E-02 | NaN | AK7 |
| Circadian Rhythm Signaling | 6.89E-01 | 1.12E-02 | NaN | GRIA2,NGFR,NOTUM |
| Cardiac Hypertrophy Signaling (Enhanced) | 6.81E-01 | 9.23E-03 | NaN | IL11,IL1B,NGFR,NKX2-5,NOTUM |
| Thyroid Hormone Metabolism II (via Conjugation and/or D | 6.72E-01 | 2.44E-02 | NaN | SULT1B1 |
| Endocannabinoid Neuronal Synapse Pathway | 6.68E-01 | 1.34E-02 | NaN | GRIA2,NOTUM |
| Cellular Effects of Sildenafil (Viagra) | 6.64E-01 | 1.33E-02 | NaN | MYL4,NOTUM |
| Intrinsic Prothrombin Activation Pathway | 6.63E-01 | 2.38E-02 | NaN | F13A1 |
| Pyrimidine Ribonucleotides De Novo Biosynthesis | 6.63E-01 | 2.38E-02 | NaN | AK7 |
| PTEN Signaling | 6.59E-01 | 1.32E-02 | NaN | FOXG1,NGFR |
| Oncostatin M Signaling | 6.54E-01 | 2.33E-02 | NaN | MMP3 |
| Role of Pattern Recognition Receptors in Recognition of B | 6.39E-01 | 1.28E-02 | NaN | IL11,IL1B |
| CLEAR Signaling Pathway | 6.37E-01 | 1.05E-02 | NaN | BMP6,HGF,NGFR |
| Role of OCT4 in Mammalian Embryonic Stem Cell Pluripot | 6.28E-01 | 2.17E-02 | NaN | FOXA2 |
| IL-23 Signaling Pathway | 6.28E-01 | 2.17E-02 | NaN | IL1B |
| RAR Activation | 6.23E-01 | 9.35E-03 | NaN | GATA6,IL11,IL1B,MMP3 |
| Estrogen Biosynthesis | 6.20E-01 | 2.13E-02 | NaN | HSD17B13 |
| Xenobiotic Metabolism Signaling | 6.17E-01 | 1.03E-02 | NaN | CHST1,IL1B,SULT1B1 |
| Transcriptional Regulatory Network in Embryonic Stem Ce | 6.07E-01 | 1.22E-02 | NaN | GATA6,IGF2 |
| MYC Mediated Apoptosis Signaling | 5.97E-01 | 2.00E-02 | NaN | NGFR |
| CXCR4 Signaling | 5.93E-01 | 1.19E-02 | NaN | CXCR4,MYL4 |
| Glioblastoma Multiforme Signaling | 5.82E-01 | 1.17E-02 | NaN | IGF2,NOTUM |
| Aldosterone Signaling in Epithelial Cells | 5.78E-01 | 1.16E-02 | NaN | CRYAB,NOTUM |
| Dendritic Cell Maturation | 5.75E-01 | 8.39E-03 | NaN | CCR7,IL1B,NGFR,NOTUM,TRBV5-4 |
| Role of Cytokines in Mediating Communication between I | 5.68E-01 | 1.85E-02 | NaN | IL1B |
| Erythropoietin Signaling Pathway | 5.61E-01 | 1.13E-02 | NaN | IL11,IL1B |

| | | | | |
|---|---|---|---|---|
| Synaptogenesis Signaling Pathway | 5.57E-01 | 9.52E-03 | NaN | EPHA10,GRIA2,SYN3 |
| CSDE1 Signaling Pathway | 5.55E-01 | 1.79E-02 | NaN | TNC |
| Triacylglycerol Degradation | 5.48E-01 | 1.75E-02 | NaN | NOTUM |
| Cancer Drug Resistance By Drug Efflux | 5.42E-01 | 1.72E-02 | NaN | FOXG1 |
| Acute Phase Response Signaling | 5.34E-01 | 1.08E-02 | NaN | IL1B,NGFR |
| IL-33 Signaling Pathway | 5.31E-01 | 1.08E-02 | NaN | AREG,IL1B |
| GADD45 Signaling | 5.30E-01 | 1.67E-02 | NaN | IL1B |
| MicroRNA Biogenesis Signaling Pathway | 5.28E-01 | 1.07E-02 | NaN | BMP6,HGF |
| Macrophage Classical Activation Signaling Pathway | 5.22E-01 | 1.06E-02 | NaN | IL11,IL1B |
| Macrophage Alternative Activation Signaling Pathway | 5.22E-01 | 1.06E-02 | NaN | FCER1A,IL1B |
| NOD1/2 Signaling Pathway | 5.22E-01 | 1.06E-02 | NaN | IL11,IL1B |
| Xenobiotic Metabolism CAR Signaling Pathway | 5.13E-01 | 1.04E-02 | NaN | CHST1,SULT1B1 |
| Melatonin Degradation I | 5.12E-01 | 1.59E-02 | NaN | SULT1B1 |
| Xenobiotic Metabolism PXR Signaling Pathway | 5.10E-01 | 1.04E-02 | NaN | CHST1,SULT1B1 |
| PPARα/RXRα Activation | 5.04E-01 | 1.03E-02 | NaN | IL1B,NOTUM |
| PXR/RXR Activation | 5.01E-01 | 1.54E-02 | NaN | FOXA2 |
| WNT/Ca+ pathway | 4.95E-01 | 1.52E-02 | NaN | NOTUM |
| Glutamate Receptor Signaling | 4.95E-01 | 1.52E-02 | NaN | GRIA2 |
| Synaptic Long Term Depression | 4.95E-01 | 1.01E-02 | NaN | GRIA2,NOTUM |
| Gap Junction Signaling | 4.95E-01 | 1.01E-02 | NaN | GRIA2,NOTUM |
| Adrenomedullin signaling pathway | 4.92E-01 | 1.01E-02 | NaN | IL1B,NOTUM |
| ID1 Signaling Pathway | 4.86E-01 | 9.95E-03 | NaN | IGF2,NGFR |
| Human Embryonic Stem Cell Pluripotency | 4.86E-01 | 9.95E-03 | NaN | BMP6,NTF3 |
| Superpathway of Melatonin Degradation | 4.85E-01 | 1.47E-02 | NaN | SULT1B1 |
| Remodeling of Epithelial Adherens Junctions | 4.85E-01 | 1.47E-02 | NaN | HGF |
| Ephrin Receptor Signaling | 4.84E-01 | 9.90E-03 | NaN | CXCR4,EPHA10 |
| Phospholipases | 4.79E-01 | 1.45E-02 | NaN | NOTUM |
| Sertoli Cell-Sertoli Cell Junction Signaling | 4.72E-01 | 9.71E-03 | NaN | CLDN4,CLDN5 |
| Growth Hormone Signaling | 4.69E-01 | 1.41E-02 | NaN | IGF2 |
| Melatonin Signaling | 4.64E-01 | 1.39E-02 | NaN | NOTUM |
| Basal Cell Carcinoma Signaling | 4.64E-01 | 1.39E-02 | NaN | BMP6 |
| Ephrin B Signaling | 4.64E-01 | 1.39E-02 | NaN | CXCR4 |
| Serotonin Degradation | 4.64E-01 | 1.39E-02 | NaN | SULT1B1 |
| Granzyme A Signaling | 4.50E-01 | 1.33E-02 | NaN | IL1B |
| Leptin Signaling in Obesity | 4.46E-01 | 1.32E-02 | NaN | NOTUM |
| Macropinocytosis Signaling | 4.46E-01 | 1.32E-02 | NaN | HGF |
| Toll-like Receptor Signaling | 4.37E-01 | 1.28E-02 | NaN | IL1B |
| Calcium Signaling | 4.36E-01 | 9.09E-03 | NaN | GRIA2,MYL4 |
| Maturity Onset Diabetes of Young (MODY) Signaling | 4.32E-01 | 1.27E-02 | NaN | FOXA2 |
| Renal Cell Carcinoma Signaling | 4.32E-01 | 1.27E-02 | NaN | HGF |
| Role of JAK family kinases in IL-6-type Cytokine Signaling | 4.32E-01 | 1.27E-02 | NaN | IL11 |
| Role of MAPK Signaling in Inhibiting the Pathogenesis of Ir | 4.32E-01 | 1.27E-02 | NaN | IL1B |
| Chemokine Signaling | 4.24E-01 | 1.23E-02 | NaN | CXCR4 |
| BEX2 Signaling Pathway | 4.20E-01 | 1.22E-02 | NaN | NGFR |
| HER-2 Signaling in Breast Cancer | 4.19E-01 | 8.81E-03 | NaN | AREG,FCER1A |
| Estrogen-Dependent Breast Cancer Signaling | 4.16E-01 | 1.20E-02 | NaN | HSD17B13 |
| Neurovascular Coupling Signaling Pathway | 4.08E-01 | 8.62E-03 | NaN | GABRQ,GRIA2 |
| FGF Signaling | 4.04E-01 | 1.16E-02 | NaN | HGF |
| Xenobiotic Metabolism AHR Signaling Pathway | 4.00E-01 | 1.15E-02 | NaN | IL1B |
| Regulation Of The Epithelial Mesenchymal Transition In De | 4.00E-01 | 1.15E-02 | NaN | TWIST1 |
| cAMP-mediated signaling | 3.99E-01 | 8.47E-03 | NaN | HTR4,LHCGR |
| Neutrophil Extracellular Trap Signaling Pathway | 3.87E-01 | 7.52E-03 | NaN | COL6A5,IL1B,NOTUM |
| AMPK Signaling | 3.86E-01 | 8.26E-03 | NaN | AK7,FOXG1 |
| Ceramide Signaling | 3.85E-01 | 1.10E-02 | NaN | NGFR |
| Crosstalk between Dendritic Cells and Natural Killer Cells | 3.85E-01 | 1.10E-02 | NaN | CCR7 |
| ABRA Signaling Pathway | 3.81E-01 | 1.09E-02 | NaN | FOSB |
| Protein Kinase A Signaling | 3.68E-01 | 7.30E-03 | NaN | MYL4,NGFR,NOTUM |
| TGF-β Signaling | 3.67E-01 | 1.04E-02 | NaN | NKX2-5 |
| Salvage Pathways of Pyrimidine Ribonucleotides | 3.64E-01 | 1.03E-02 | NaN | AK7 |
| DNA Methylation and Transcriptional Repression Signaling | 3.61E-01 | 1.02E-02 | NaN | FOXA2 |
| UVA-Induced MAPK Signaling | 3.61E-01 | 1.02E-02 | NaN | NOTUM |
| Glucocorticoid Receptor Signaling | 3.57E-01 | 6.87E-03 | NaN | CXCL3,IL1B,MMP3,SLPI |
| Hepatic Fibrosis Signaling Pathway | 3.50E-01 | 7.09E-03 | NaN | IL1B,MYL4,NGFR |
| PD-1, PD-L1 cancer immunotherapy pathway | 3.33E-01 | 9.35E-03 | NaN | NGFR |
| Colorectal Cancer Metastasis Signaling | 3.29E-01 | 7.38E-03 | NaN | MMP21,MMP3 |
| Opioid Signaling Pathway | 3.14E-01 | 7.14E-03 | NaN | FOSB,SCN7A |
| Antioxidant Action of Vitamin C | 3.13E-01 | 8.77E-03 | NaN | NOTUM |
| Regulation of Actin-based Motility by Rho | 3.10E-01 | 8.70E-03 | NaN | MYL4 |
| IL-13 Signaling Pathway | 3.08E-01 | 8.62E-03 | NaN | FOXA2 |
| Amyotrophic Lateral Sclerosis Signaling | 3.08E-01 | 8.62E-03 | NaN | GRIA2 |
| PAK Signaling | 3.05E-01 | 8.55E-03 | NaN | MYL4 |
| Fc Epsilon RI Signaling | 3.03E-01 | 8.47E-03 | NaN | FCER1A |

| | | | | |
|---|---|---|---|---|
| GPCR-Mediated Nutrient Sensing in Enteroendocrine Cells | 3.03E-01 | 8.47E-03 | NaN | NOTUM |
| Cholecystokinin/Gastrin-mediated Signaling | 3.00E-01 | 8.40E-03 | NaN | IL1B |
| Sphingosine-1-phosphate Signaling | 2.98E-01 | 8.33E-03 | NaN | NOTUM |
| NGF Signaling | 2.98E-01 | 8.33E-03 | NaN | NGFR |
| p38 MAPK Signaling | 2.98E-01 | 8.33E-03 | NaN | IL1B |
| Neuroprotective Role of THOP1 in Alzheimer's Disease | 2.95E-01 | 8.26E-03 | NaN | HTRA3 |
| RHOA Signaling | 2.88E-01 | 8.06E-03 | NaN | MYL4 |
| Chaperone Mediated Autophagy Signaling Pathway | 2.87E-01 | 6.22E-03 | NaN | IL1B,MMP21,MMP3,TRBV5-4 |
| Glioma Signaling | 2.85E-01 | 8.00E-03 | NaN | IGF2 |
| 14-3-3-mediated Signaling | 2.81E-01 | 7.87E-03 | NaN | NOTUM |
| HGF Signaling | 2.69E-01 | 7.58E-03 | NaN | HGF |
| GABA Receptor Signaling | 2.69E-01 | 7.58E-03 | NaN | GABRQ |
| Ferroptosis Signaling Pathway | 2.69E-01 | 7.58E-03 | NaN | SLC7A11 |
| Gα12/13 Signaling | 2.67E-01 | 7.52E-03 | NaN | MYL4 |
| P2Y Purigenic Receptor Signaling Pathway | 2.67E-01 | 7.52E-03 | NaN | NOTUM |
| Role of PKR in Interferon Induction and Antiviral Response | 2.61E-01 | 7.35E-03 | NaN | IL1B |
| Th2 Pathway | 2.59E-01 | 7.30E-03 | NaN | CXCR4 |
| Iron homeostasis signaling pathway | 2.57E-01 | 7.25E-03 | NaN | BMP6 |
| White Adipose Tissue Browning Pathway | 2.57E-01 | 7.25E-03 | NaN | FCER1A |
| Type I Diabetes Mellitus Signaling | 2.44E-01 | 5.89E-03 | NaN | IL1B,NGFR,TRBV5-4 |
| Endocannabinoid Cancer Inhibition Pathway | 2.39E-01 | 6.80E-03 | NaN | TWIST1 |
| Dilated Cardiomyopathy Signaling Pathway | 2.34E-01 | 6.67E-03 | NaN | MYL4 |
| Semaphorin Neuronal Repulsive Signaling Pathway | 2.34E-01 | 6.67E-03 | NaN | MYL4 |
| Type II Diabetes Mellitus Signaling | 2.28E-01 | 6.54E-03 | NaN | NGFR |
| IL-10 Signaling | 2.26E-01 | 6.49E-03 | NaN | IL1B |
| Necroptosis Signaling Pathway | 2.23E-01 | 6.41E-03 | NaN | NGFR |
| Epithelial Adherens Junction Signaling | 2.21E-01 | 6.37E-03 | NaN | HGF |
| Ovarian Cancer Signaling | 2.20E-01 | 6.33E-03 | NaN | LHCGR |
| Aryl Hydrocarbon Receptor Signaling | 2.18E-01 | 6.29E-03 | NaN | IL1B |
| Inhibition of ARE-Mediated mRNA Degradation Pathway | 2.13E-01 | 6.17E-03 | NaN | NGFR |
| Ribonucleotide Reductase Signaling Pathway | 2.01E-01 | 5.88E-03 | NaN | HGF |
| Th1 and Th2 Activation Pathway | 1.98E-01 | 5.81E-03 | NaN | CXCR4 |
| Oxytocin In Brain Signaling Pathway | 0.00E+00 | 5.03E-03 | NaN | IL1B |
| Oxytocin Signaling Pathway | 0.00E+00 | 3.55E-03 | NaN | MYL4 |
| Actin Cytoskeleton Signaling | 0.00E+00 | 4.10E-03 | NaN | MYL4 |
| Role of NFAT in Regulation of the Immune Response | 0.00E+00 | 1.93E-03 | NaN | FCER1A,TRBV5-4 |
| CCR5 Signaling in Macrophages | 0.00E+00 | 2.00E-03 | NaN | TRBV5-4 |
| Calcium-induced T Lymphocyte Apoptosis | 0.00E+00 | 2.17E-03 | NaN | TRBV5-4 |
| Cytotoxic T Lymphocyte-mediated Apoptosis of Target Cel | 0.00E+00 | 2.35E-03 | NaN | TRBV5-4 |
| CTLA4 Signaling in Cytotoxic T Lymphocytes | 0.00E+00 | 1.64E-03 | NaN | TRBV5-4 |
| T Helper Cell Differentiation | 0.00E+00 | 4.25E-03 | NaN | NGFR,TRBV5-4 |
| CD28 Signaling in T Helper Cells | 0.00E+00 | 1.93E-03 | NaN | TRBV5-4 |
| Endothelin-1 Signaling | 0.00E+00 | 5.15E-03 | NaN | NOTUM |
| ICOS-ICOSL Signaling in T Helper Cells | 0.00E+00 | 1.97E-03 | NaN | TRBV5-4 |
| Molecular Mechanisms of Cancer | 0.00E+00 | 2.22E-03 | NaN | BMP6 |
| Lipid Antigen Presentation by CD1 | 0.00E+00 | 2.41E-03 | NaN | TRBV5-4 |
| Allograft Rejection Signaling | 0.00E+00 | 2.05E-03 | NaN | TRBV5-4 |
| Autoimmune Thyroid Disease Signaling | 0.00E+00 | 2.18E-03 | NaN | TRBV5-4 |
| Graft-versus-Host Disease Signaling | 0.00E+00 | 4.48E-03 | NaN | IL1B,TRBV5-4 |
| Chronic Myeloid Leukemia Signaling | 0.00E+00 | 3.60E-03 | NaN | NOTUM |
| Production of Nitric Oxide and Reactive Oxygen Species in | 0.00E+00 | 5.24E-03 | NaN | NGFR |
| p70S6K Signaling | 0.00E+00 | 1.72E-03 | NaN | NOTUM |
| G Protein Signaling Mediated by Tubby | 0.00E+00 | 2.14E-03 | NaN | TRBV5-4 |
| Communication between Innate and Adaptive Immune Ce | 0.00E+00 | 3.22E-03 | NaN | CCR7,IL1B,TRBV5-4 |
| Systemic Lupus Erythematosus Signaling | 0.00E+00 | 1.87E-03 | NaN | IL1B,TRBV5-4 |
| CDC42 Signaling | 0.00E+00 | 3.47E-03 | NaN | MYL4,TRBV5-4 |
| ILK Signaling | 0.00E+00 | 4.98E-03 | NaN | MYL4 |
| Phospholipase C Signaling | 0.00E+00 | 1.78E-03 | NaN | MYL4,TRBV5-4 |
| Altered T Cell and B Cell Signaling in Rheumatoid Arthritis | 0.00E+00 | 2.13E-03 | NaN | IL1B,TRBV5-4 |
| Regulation of IL-2 Expression in Activated and Anergic T Ly | 0.00E+00 | 2.16E-03 | NaN | TRBV5-4 |
| NUR77 Signaling in T Lymphocytes | 0.00E+00 | 1.95E-03 | NaN | TRBV5-4 |
| PKCθ Signaling in T Lymphocytes | 0.00E+00 | 1.79E-03 | NaN | TRBV5-4 |
| Antiproliferative Role of TOB in T Cell Signaling | 0.00E+00 | 2.35E-03 | NaN | TRBV5-4 |
| OX40 Signaling Pathway | 0.00E+00 | 2.09E-03 | NaN | TRBV5-4 |
| PI3K Signaling in B Lymphocytes | 0.00E+00 | 1.69E-03 | NaN | NOTUM |
| Dopamine-DARPP32 Feedback in cAMP Signaling | 0.00E+00 | 5.38E-03 | NaN | NOTUM |
| Signaling by Rho Family GTPases | 0.00E+00 | 3.75E-03 | NaN | MYL4 |
| RHOGDI Signaling | 0.00E+00 | 4.55E-03 | NaN | MYL4 |
| Hematopoiesis from Pluripotent Stem Cells | 0.00E+00 | 4.49E-03 | NaN | IL11,TRBV5-4 |
| Sperm Motility | 0.00E+00 | 3.89E-03 | NaN | NOTUM |
| TEC Kinase Signaling | 0.00E+00 | 3.46E-03 | NaN | FCER1A,TRBV5-4 |
| SAPK/JNK Signaling | 0.00E+00 | 1.99E-03 | NaN | TRBV5-4 |

| | | | | |
|---|---|---|---|---|
| Protein Ubiquitination Pathway | 0.00E+00 | 3.66E-03 | NaN | CRYAB |
| IL-4 Signaling | 0.00E+00 | 3.47E-03 | NaN | COL6A5,TRBV5-4 |
| WNT/β-catenin Signaling | 0.00E+00 | 5.75E-03 | NaN | DKK3 |
| NF-κB Signaling | 0.00E+00 | 5.25E-03 | NaN | IL1B,NGFR,TRBV5-4 |
| T Cell Receptor Signaling | 0.00E+00 | 1.62E-03 | NaN | TRBV5-4 |
| Th17 Activation Pathway | 0.00E+00 | 4.13E-03 | NaN | IL1B,TRBV5-4 |
| T Cell Exhaustion Signaling Pathway | 0.00E+00 | 1.76E-03 | NaN | TRBV5-4 |
| Systemic Lupus Erythematosus In T Cell Signaling Pathway | 0.00E+00 | 1.56E-03 | NaN | TRBV5-4 |
| Systemic Lupus Erythematosus In B Cell Signaling Pathway | 0.00E+00 | 4.14E-03 | NaN | FOXG1,IL11,IL1B |
| Insulin Secretion Signaling Pathway | 0.00E+00 | 3.68E-03 | NaN | NOTUM |
| Coronavirus Pathogenesis Pathway | 0.00E+00 | 4.90E-03 | NaN | IL1B |

IPA_Venn_List_Astro_FB1_FC2_adj0.05_vs._Astro_GCS_FC2_adj0.05_153genes

| | Ingenuity Canonical Pathways | -log(p-value) | Ratio | z-score | Molecules |
|---|---|---|---|---|---|
| 1 | Agranulocyte Adhesion and Diapedesis | 3.01E+00 | 2.86E-02 | NaN | ACTA2,CCL2,CLDN5,ITGB7,MMP10,MMP17 |
| 2 | Inhibition of Matrix Metalloproteases | 2.92E+00 | 7.69E-02 | NaN | MMP10,MMP17,TFPI2 |
| 3 | ABRA Signaling Pathway | 2.81E+00 | 4.35E-02 | NaN | ACTA2,CCN2,EGR1,TAGLN |
| 4 | Pulmonary Fibrosis Idiopathic Signaling Pathway | 2.70E+00 | 2.15E-02 | NaN | ACTA2,CCN2,EGR1,MMP10,MMP17,PMAIP1,THBS1 |
| 5 | Apelin Cardiac Fibroblast Signaling Pathway | 2.17E+00 | 8.70E-02 | NaN | CCN2,SPHK1 |
| 6 | Role Of Chondrocytes In Rheumatoid Arthritis Sign | 2.14E+00 | 2.84E-02 | NaN | CCL2,EGR1,MMP10,MMP17 |
| 7 | Granulocyte Adhesion and Diapedesis | 1.71E+00 | 2.12E-02 | NaN | CCL2,CLDN5,MMP10,MMP17 |
| 8 | WNK Renal Signaling Pathway | 1.70E+00 | 2.83E-02 | NaN | BIRC3,SGK1,WNK4 |
| 9 | Leukocyte Extravasation Signaling | 1.68E+00 | 2.07E-02 | NaN | ACTA2,CLDN5,MMP10,MMP17 |
| 10 | ID1 Signaling Pathway | 1.63E+00 | 1.99E-02 | NaN | ATF3,CCN2,CHRNA9,EGR1 |
| | Bladder Cancer Signaling | 1.60E+00 | 2.59E-02 | NaN | MMP10,MMP17,THBS1 |
| | Osteoarthritis Pathway | 1.41E+00 | 1.69E-02 | NaN | DKK1,ITGB7,MMP10,SPHK1 |
| | MSP-RON Signaling Pathway | 1.41E+00 | 3.45E-02 | NaN | ACTA2,CCL2 |
| | Dilated Cardiomyopathy Signaling Pathway | 1.32E+00 | 2.00E-02 | NaN | ACTA2,PLN,TNNI3 |
| | Caveolar-mediated Endocytosis Signaling | 1.21E+00 | 2.67E-02 | NaN | ACTA2,ITGB7 |
| | Mineralocorticoid Biosynthesis | 1.20E+00 | 8.33E-02 | NaN | GSTA1 |
| | Glucocorticoid Biosynthesis | 1.17E+00 | 7.69E-02 | NaN | GSTA1 |
| | Protein Kinase A Signaling | 1.14E+00 | 1.22E-02 | NaN | DUSP4,MYLK3,PLN,PTPRN,TNNI3 |
| | Tumor Microenvironment Pathway | 1.14E+00 | 1.68E-02 | NaN | CCL2,MMP10,MMP17 |
| | Hepatic Fibrosis Signaling Pathway | 1.10E+00 | 1.18E-02 | NaN | ACTA2,CCL2,CCN2,ITGB7,MYLK3 |
| | NOD1/2 Signaling Pathway | 1.08E+00 | 1.59E-02 | NaN | BIRC3,CCL2,TAB2 |
| | Unfolded protein response | 1.07E+00 | 2.22E-02 | NaN | DNAJB13,SREBF1 |
| | Role Of Osteoclasts In Rheumatoid Arthritis Signali | 1.06E+00 | 1.29E-02 | NaN | BIRC3,MMP10,MMP17,TAB2 |
| | RANK Signaling in Osteoclasts | 1.06E+00 | 2.20E-02 | NaN | BIRC3,TAB2 |
| | Hepatic Fibrosis / Hepatic Stellate Cell Activation | 1.06E+00 | 1.55E-02 | NaN | ACTA2,CCL2,CCN2 |
| | Differential Regulation of Cytokine Production in M | 1.03E+00 | 5.56E-02 | NaN | CCL2 |
| | Pulmonary Healing Signaling Pathway | 1.03E+00 | 1.51E-02 | NaN | MMP10,MMP17,THBS1 |
| | Death Receptor Signaling | 1.02E+00 | 2.08E-02 | NaN | ACTA2,BIRC3 |
| | Salvage Pathways of Pyrimidine Ribonucleotides | 1.02E+00 | 2.06E-02 | NaN | AK7,SGK1 |
| | p53 Signaling | 1.01E+00 | 2.04E-02 | NaN | PMAIP1,THBS1 |
| | Sertoli Cell-Sertoli Cell Junction Signaling | 9.98E-01 | 1.46E-02 | NaN | ACTA2,CLDN5,ITGB7 |
| | Role of Tissue Factor in Cancer | 9.93E-01 | 1.45E-02 | NaN | CCN1,CCN2,EGR1 |
| | Androgen Biosynthesis | 9.89E-01 | 5.00E-02 | NaN | GSTA1 |
| | Integrin Signaling | 9.70E-01 | 1.42E-02 | NaN | ACTA2,ITGB7,MYLK3 |
| | IGF-1 Signaling | 9.58E-01 | 1.90E-02 | NaN | CCN1,CCN2 |
| | Paxillin Signaling | 9.44E-01 | 1.87E-02 | NaN | ACTA2,ITGB7 |
| | Calcium Signaling | 9.35E-01 | 1.36E-02 | NaN | ACTA2,CHRNA9,TNNI3 |
| | Differential Regulation of Cytokine Production in Ir | 9.32E-01 | 4.35E-02 | NaN | CCL2 |
| | Pyrimidine Deoxyribonucleotides De Novo Biosynt | 9.32E-01 | 4.35E-02 | NaN | AK7 |
| | EIF2 Signaling | 9.05E-01 | 1.32E-02 | NaN | ACTA2,ATF3,SREBF1 |
| | Role of Osteoblasts, Osteoclasts and Chondrocytes | 9.01E-01 | 1.32E-02 | NaN | BIRC3,DKK1,TAB2 |
| | Regulation of Actin-based Motility by Rho | 8.93E-01 | 1.74E-02 | NaN | ACTA2,ITGB7 |
| | Virus Entry via Endocytic Pathways | 8.75E-01 | 1.69E-02 | NaN | ACTA2,ITGB7 |
| | NRF2-mediated Oxidative Stress Response | 8.65E-01 | 1.27E-02 | NaN | ACTA2,DNAJB13,GSTA1 |
| | Glutathione Redox Reactions I | 8.52E-01 | 3.57E-02 | NaN | GSTA1 |
| | LXR/RXR Activation | 8.46E-01 | 1.63E-02 | NaN | CCL2,SREBF1 |
| | RHOA Signaling | 8.41E-01 | 1.61E-02 | NaN | ACTA2,MYLK3 |
| | Role Of Osteoblasts In Rheumatoid Arthritis Signal | 8.38E-01 | 1.23E-02 | NaN | DKK1,MMP10,MMP17 |
| | Actin Cytoskeleton Signaling | 8.38E-01 | 1.23E-02 | NaN | ACTA2,ITGB7,MYLK3 |
| | TNFR2 Signaling | 7.99E-01 | 3.12E-02 | NaN | BIRC3 |
| | LPS/IL-1 Mediated Inhibition of RXR Function | 7.98E-01 | 1.18E-02 | NaN | GSTA1,SLCO1A2,SREBF1 |
| | Airway Inflammation in Asthma | 7.87E-01 | 3.03E-02 | NaN | CCL2 |
| | SNARE Signaling Pathway | 7.78E-01 | 1.47E-02 | NaN | STXBP6,SYT6 |
| | Role of PKR in Interferon Induction and Antiviral R | 7.78E-01 | 1.47E-02 | NaN | ATF3,TAB2 |
| | Inhibition of Angiogenesis by TSP1 | 7.75E-01 | 2.94E-02 | NaN | THBS1 |
| | TWEAK Signaling | 7.41E-01 | 2.70E-02 | NaN | BIRC3 |
| | Glutathione-mediated Detoxification | 7.41E-01 | 2.70E-02 | NaN | GSTA1 |

| | | | | |
|---|---|---|---|---|
| Complement System | 7.41E-01 | 2.70E-02 | NaN | MASP1 |
| Protein Ubiquitination Pathway | 7.38E-01 | 1.10E-02 | NaN | BIRC3,DNAJB13,USP43 |
| IL-17A Signaling in Fibroblasts | 7.31E-01 | 2.63E-02 | NaN | CCL2 |
| Pyrimidine Ribonucleotides Interconversion | 7.21E-01 | 2.56E-02 | NaN | AK7 |
| Mechanisms of Viral Exit from Host Cells | 7.01E-01 | 2.44E-02 | NaN | ACTA2 |
| Pyrimidine Ribonucleotides De Novo Biosynthesis | 6.92E-01 | 2.38E-02 | NaN | AK7 |
| eNOS Signaling | 6.87E-01 | 1.28E-02 | NaN | AQP5,CHRNA9 |
| Necroptosis Signaling Pathway | 6.87E-01 | 1.28E-02 | NaN | BIRC3,TAB2 |
| HOTAIR Regulatory Pathway | 6.59E-01 | 1.23E-02 | NaN | MMP10,MMP17 |
| Role of IL-17F in Allergic Inflammatory Airway Dise | 6.49E-01 | 2.13E-02 | NaN | CCL2 |
| Molecular Mechanisms of Cancer | 6.47E-01 | 8.89E-03 | NaN | BIRC3,ITGB7,PMAIP1,TAB2 |
| MYC Mediated Apoptosis Signaling | 6.25E-01 | 2.00E-02 | NaN | PMAIP1 |
| Aldosterone Signaling in Epithelial Cells | 6.25E-01 | 1.16E-02 | NaN | DNAJB13,SGK1 |
| TNFR1 Signaling | 6.18E-01 | 1.96E-02 | NaN | BIRC3 |
| Chondroitin Sulfate Biosynthesis (Late Stages) | 6.18E-01 | 1.96E-02 | NaN | CHST9 |
| Synaptogenesis Signaling Pathway | 6.17E-01 | 9.52E-03 | NaN | STXBP6,SYT6,THBS1 |
| Neuroinflammation Signaling Pathway | 6.12E-01 | 9.46E-03 | NaN | BIRC3,CCL2,GABRG1 |
| Tight Junction Signaling | 6.00E-01 | 1.12E-02 | NaN | ACTA2,CLDN5 |
| D-myo-inositol (1,4,5,6)-Tetrakisphosphate Biosyn | 5.96E-01 | 1.11E-02 | NaN | DUSP4,PTPRN |
| D-myo-inositol (3,4,5,6)-tetrakisphosphate Biosynt | 5.96E-01 | 1.11E-02 | NaN | DUSP4,PTPRN |
| CSDE1 Signaling Pathway | 5.83E-01 | 1.79E-02 | NaN | CCL2 |
| Role of IL-17A in Arthritis | 5.76E-01 | 1.75E-02 | NaN | CCL2 |
| Role of Macrophages, Fibroblasts and Endothelial | 5.75E-01 | 9.04E-03 | NaN | CCL2,DKK1,IL16 |
| IL-17 Signaling | 5.73E-01 | 1.07E-02 | NaN | CCL2,TAB2 |
| Macrophage Alternative Activation Signaling Pathv | 5.67E-01 | 1.06E-02 | NaN | SREBF1,THBS1 |
| Chondroitin Sulfate Biosynthesis | 5.63E-01 | 1.69E-02 | NaN | CHST9 |
| Hepatic Cholestasis | 5.60E-01 | 1.05E-02 | NaN | SLCO1A2,SREBF1 |
| 3-phosphoinositide Degradation | 5.60E-01 | 1.05E-02 | NaN | DUSP4,PTPRN |
| SPINK1 Pancreatic Cancer Pathway | 5.57E-01 | 1.67E-02 | NaN | CPA4 |
| Role of JAK2 in Hormone-like Cytokine Signaling | 5.45E-01 | 1.61E-02 | NaN | BIRC3 |
| D-myo-inositol-5-phosphate Metabolism | 5.45E-01 | 1.02E-02 | NaN | DUSP4,PTPRN |
| ILK Signaling | 5.30E-01 | 9.95E-03 | NaN | ACTA2,ITGB7 |
| PXR/RXR Activation | 5.28E-01 | 1.54E-02 | NaN | GSTA1 |
| Induction of Apoptosis by HIV1 | 5.28E-01 | 1.54E-02 | NaN | BIRC3 |
| Pyridoxal 5'-phosphate Salvage Pathway | 5.28E-01 | 1.54E-02 | NaN | SGK1 |
| 3-phosphoinositide Biosynthesis | 5.16E-01 | 9.71E-03 | NaN | DUSP4,PTPRN |
| Remodeling of Epithelial Adherens Junctions | 5.12E-01 | 1.47E-02 | NaN | ACTA2 |
| Clathrin-mediated Endocytosis Signaling | 5.10E-01 | 9.62E-03 | NaN | ACTA2,ITGB7 |
| HIF1α Signaling | 5.10E-01 | 9.62E-03 | NaN | MMP10,MMP17 |
| Agrin Interactions at Neuromuscular Junction | 5.06E-01 | 1.45E-02 | NaN | ACTA2 |
| ERK/MAPK Signaling | 4.91E-01 | 9.30E-03 | NaN | DUSP4,ITGB7 |
| Activin Inhibin Signaling Pathway | 4.86E-01 | 9.22E-03 | NaN | CCN2,FBXO32 |
| ERK5 Signaling | 4.81E-01 | 1.35E-02 | NaN | SGK1 |
| RHOGDI Signaling | 4.78E-01 | 9.09E-03 | NaN | ACTA2,ITGB7 |
| Macropinocytosis Signaling | 4.72E-01 | 1.32E-02 | NaN | ITGB7 |
| TREM1 Signaling | 4.67E-01 | 1.30E-02 | NaN | CCL2 |
| Toll-like Receptor Signaling | 4.63E-01 | 1.28E-02 | NaN | TAB2 |
| Role of MAPK Signaling in Inhibiting the Pathogene | 4.58E-01 | 1.27E-02 | NaN | CCL2 |
| Chemokine Signaling | 4.50E-01 | 1.23E-02 | NaN | CCL2 |
| Superpathway of Inositol Phosphate Compounds | 4.43E-01 | 8.55E-03 | NaN | DUSP4,PTPRN |
| cAMP-mediated signaling | 4.39E-01 | 8.47E-03 | NaN | DUSP4,RGS4 |
| TR/RXR Activation | 4.37E-01 | 1.19E-02 | NaN | SREBF1 |
| Role of MAPK Signaling in the Pathogenesis of Influ | 4.37E-01 | 1.19E-02 | NaN | CCL2 |
| Role of Hypercytokinemia/hyperchemokinemia in | 4.29E-01 | 1.16E-02 | NaN | CCL2 |
| AMPK Signaling | 4.25E-01 | 8.26E-03 | NaN | AK7,CHRNA9 |
| PDGF Signaling | 4.25E-01 | 1.15E-02 | NaN | SPHK1 |
| Xenobiotic Metabolism AHR Signaling Pathway | 4.25E-01 | 1.15E-02 | NaN | GSTA1 |
| Estrogen Receptor Signaling | 4.21E-01 | 7.33E-03 | NaN | FBXO32,MMP10,MMP17 |
| Regulation of Cellular Mechanics by Calpain Protea | 4.14E-01 | 1.11E-02 | NaN | ITGB7 |
| Ceramide Signaling | 4.10E-01 | 1.10E-02 | NaN | SPHK1 |
| Crosstalk between Dendritic Cells and Natural Kille | 4.10E-01 | 1.10E-02 | NaN | ACTA2 |

| | | | | |
|---|---|---|---|---|
| Apelin Adipocyte Signaling Pathway | 4.10E-01 | 1.10E-02 | NaN | GSTA1 |
| Wound Healing Signaling Pathway | 4.04E-01 | 7.94E-03 | NaN | ACTA2,MMP10 |
| Actin Nucleation by ARP-WASP Complex | 4.03E-01 | 1.08E-02 | NaN | ITGB7 |
| Fcγ Receptor-mediated Phagocytosis in Macropha | 3.99E-01 | 1.06E-02 | NaN | ACTA2 |
| IL-1 Signaling | 3.92E-01 | 1.04E-02 | NaN | TAB2 |
| VEGF Signaling | 3.82E-01 | 1.01E-02 | NaN | ACTA2 |
| Apelin Cardiomyocyte Signaling Pathway | 3.82E-01 | 1.01E-02 | NaN | PLN |
| Signaling by Rho Family GTPases | 3.74E-01 | 7.49E-03 | NaN | ACTA2,ITGB7 |
| Colorectal Cancer Metastasis Signaling | 3.66E-01 | 7.38E-03 | NaN | MMP10,MMP17 |
| Apoptosis Signaling | 3.66E-01 | 9.62E-03 | NaN | BIRC3 |
| Opioid Signaling Pathway | 3.50E-01 | 7.14E-03 | NaN | EGR4,RGS4 |
| CDK5 Signaling | 3.34E-01 | 8.70E-03 | NaN | EGR1 |
| Serotonin Receptor Signaling | 3.32E-01 | 6.40E-03 | NaN | MYLK3,PLN,TNNI3 |
| Amyotrophic Lateral Sclerosis Signaling | 3.31E-01 | 8.62E-03 | NaN | BIRC3 |
| Neuregulin Signaling | 3.28E-01 | 8.55E-03 | NaN | ITGB7 |
| PAK Signaling | 3.28E-01 | 8.55E-03 | NaN | ITGB7 |
| Airway Pathology in Chronic Obstructive Pulmonar | 3.26E-01 | 8.47E-03 | NaN | CCL2 |
| Sphingosine-1-phosphate Signaling | 3.20E-01 | 8.33E-03 | NaN | SPHK1 |
| Nitric Oxide Signaling in the Cardiovascular System | 3.20E-01 | 8.33E-03 | NaN | PLN |
| p38 MAPK Signaling | 3.20E-01 | 8.33E-03 | NaN | TAB2 |
| Senescence Pathway | 3.18E-01 | 6.69E-03 | NaN | ATF3,PDK4 |
| Renin-Angiotensin Signaling | 3.18E-01 | 8.26E-03 | NaN | CCL2 |
| Neuroprotective Role of THOP1 in Alzheimer's Dis | 3.18E-01 | 8.26E-03 | NaN | MASP1 |
| Th1 Pathway | 3.15E-01 | 8.20E-03 | NaN | ICOSLG/LOC102723996 |
| FXR/RXR Activation | 3.06E-01 | 7.94E-03 | NaN | SREBF1 |
| HGF Signaling | 2.92E-01 | 7.58E-03 | NaN | ITGB7 |
| GABA Receptor Signaling | 2.92E-01 | 7.58E-03 | NaN | GABRG1 |
| Ferroptosis Signaling Pathway | 2.92E-01 | 7.58E-03 | NaN | ANGPTL4 |
| Atherosclerosis Signaling | 2.89E-01 | 7.52E-03 | NaN | CCL2 |
| Axonal Guidance Signaling | 2.83E-01 | 5.88E-03 | NaN | ITGB7,MMP10,MMP17 |
| DHCR24 Signaling Pathway | 2.81E-01 | 7.30E-03 | NaN | SREBF1 |
| RAC Signaling | 2.81E-01 | 7.30E-03 | NaN | ITGB7 |
| Th2 Pathway | 2.81E-01 | 7.30E-03 | NaN | ICOSLG/LOC102723996 |
| Reelin Signaling in Neurons | 2.78E-01 | 7.25E-03 | NaN | PDK4 |
| Myelination Signaling Pathway | 2.77E-01 | 6.12E-03 | NaN | ITGB7,SREBF1 |
| Adipogenesis pathway | 2.76E-01 | 7.19E-03 | NaN | SREBF1 |
| Gαi Signaling | 2.74E-01 | 7.14E-03 | NaN | RGS4 |
| Insulin Receptor Signaling | 2.74E-01 | 7.14E-03 | NaN | SGK1 |
| MSP-RON Signaling In Cancer Cells Pathway | 2.74E-01 | 7.14E-03 | NaN | ACTA2 |
| Apelin Endothelial Signaling Pathway | 2.72E-01 | 7.09E-03 | NaN | CCL2 |
| Xenobiotic Metabolism General Signaling Pathway | 2.68E-01 | 6.99E-03 | NaN | GSTA1 |
| Endocannabinoid Cancer Inhibition Pathway | 2.60E-01 | 6.80E-03 | NaN | ATF3 |
| Cellular Effects of Sildenafil (Viagra) | 2.54E-01 | 6.67E-03 | NaN | ACTA2 |
| Semaphorin Neuronal Repulsive Signaling Pathway | 2.54E-01 | 6.67E-03 | NaN | ITGB7 |
| NAD Signaling Pathway | 2.53E-01 | 6.62E-03 | NaN | SREBF1 |
| PTEN Signaling | 2.53E-01 | 6.62E-03 | NaN | ITGB7 |
| Factors Promoting Cardiogenesis in Vertebrates | 2.49E-01 | 6.54E-03 | NaN | DKK1 |
| Aryl Hydrocarbon Receptor Signaling | 2.38E-01 | 6.29E-03 | NaN | GSTA1 |
| HMGB1 Signaling | 2.25E-01 | 5.99E-03 | NaN | CCL2 |
| CXCR4 Signaling | 2.23E-01 | 5.95E-03 | NaN | EGR1 |
| Ribonucleotide Reductase Signaling Pathway | 2.20E-01 | 5.88E-03 | NaN | THBS1 |
| Germ Cell-Sertoli Cell Junction Signaling | 2.20E-01 | 5.88E-03 | NaN | ACTA2 |
| Gαq Signaling | 2.20E-01 | 5.88E-03 | NaN | RGS4 |
| Th1 and Th2 Activation Pathway | 2.17E-01 | 5.81E-03 | NaN | ICOSLG/LOC102723996 |
| WNT/β-catenin Signaling | 2.14E-01 | 5.75E-03 | NaN | DKK1 |
| Erythropoietin Signaling Pathway | 2.10E-01 | 5.65E-03 | NaN | BIRC3 |
| Cardiac β-adrenergic Signaling | 2.05E-01 | 5.56E-03 | NaN | PLN |
| Regulation of eIF4 and p70S6K Signaling | 2.04E-01 | 5.52E-03 | NaN | ITGB7 |
| Neurovascular Coupling Signaling Pathway | 0.00E+00 | 4.31E-03 | NaN | GABRG1 |
| Multiple Sclerosis Signaling Pathway | 0.00E+00 | 4.50E-03 | NaN | MASP1 |
| Pathogen Induced Cytokine Storm Signaling Pathw | 0.00E+00 | 2.70E-03 | NaN | CCL2 |

| | | | | |
|---|---|---|---|---|
| Glucocorticoid Receptor Signaling | 0.00E+00 | 5.15E-03 | NaN | CCL2,PDK4,SGK1 |
| S100 Family Signaling Pathway | 0.00E+00 | 2.59E-03 | NaN | MMP10,MMP17 |
| Huntington's Disease Signaling | 0.00E+00 | 3.53E-03 | NaN | SGK1 |
| Chaperone Mediated Autophagy Signaling Pathwa | 0.00E+00 | 3.11E-03 | NaN | MMP10,MMP17 |
| IL-33 Signaling Pathway | 0.00E+00 | 5.38E-03 | NaN | CCL2 |
| Mitochondrial Dysfunction | 0.00E+00 | 2.91E-03 | NaN | GSTA1 |
| RAR Activation | 0.00E+00 | 2.34E-03 | NaN | DKK1 |
| IL-12 Signaling and Production in Macrophages | 0.00E+00 | 4.24E-03 | NaN | THBS1 |
| T Helper Cell Differentiation | 0.00E+00 | 2.12E-03 | NaN | ICOSLG/LOC102723996 |
| ICOS-ICOSL Signaling in T Helper Cells | 0.00E+00 | 1.97E-03 | NaN | ICOSLG/LOC102723996 |
| GNRH Signaling | 0.00E+00 | 5.24E-03 | NaN | EGR1 |
| Human Embryonic Stem Cell Pluripotency | 0.00E+00 | 4.98E-03 | NaN | SPHK1 |
| Type I Diabetes Mellitus Signaling | 0.00E+00 | 1.96E-03 | NaN | PTPRN |
| CDC42 Signaling | 0.00E+00 | 1.74E-03 | NaN | ITGB7 |
| FAK Signaling | 0.00E+00 | 9.61E-04 | NaN | ITGB7 |
| Phospholipase C Signaling | 0.00E+00 | 8.92E-04 | NaN | ITGB7 |
| HER-2 Signaling in Breast Cancer | 0.00E+00 | 4.41E-03 | NaN | ITGB7 |
| PI3K Signaling in B Lymphocytes | 0.00E+00 | 1.69E-03 | NaN | ATF3 |
| Gap Junction Signaling | 0.00E+00 | 5.05E-03 | NaN | ACTA2 |
| Regulation of the Epithelial-Mesenchymal Transitic | 0.00E+00 | 5.13E-03 | NaN | EGR1 |
| TEC Kinase Signaling | 0.00E+00 | 3.46E-03 | NaN | ACTA2,ITGB7 |
| SAPK/JNK Signaling | 0.00E+00 | 1.99E-03 | NaN | DUSP4 |
| PI3K/AKT Signaling | 0.00E+00 | 5.00E-03 | NaN | ITGB7 |
| Xenobiotic Metabolism Signaling | 0.00E+00 | 3.42E-03 | NaN | GSTA1 |
| B Cell Receptor Signaling | 0.00E+00 | 1.57E-03 | NaN | EGR1 |
| NF-κB Signaling | 0.00E+00 | 1.75E-03 | NaN | TAB2 |
| T Cell Receptor Signaling | 0.00E+00 | 1.62E-03 | NaN | ICOSLG/LOC102723996 |
| G-Protein Coupled Receptor Signaling | 0.00E+00 | 4.27E-03 | NaN | DUSP4,MYLK3,RGS4 |
| Gustation Pathway | 0.00E+00 | 4.93E-03 | NaN | GABRG1 |
| Phagosome Formation | 0.00E+00 | 4.32E-03 | NaN | ITGB7,MYLK3,SPHK1 |
| Sirtuin Signaling Pathway | 0.00E+00 | 3.41E-03 | NaN | SREBF1 |
| Adrenomedullin signaling pathway | 0.00E+00 | 5.03E-03 | NaN | MYLK3 |
| Cardiac Hypertrophy Signaling (Enhanced) | 0.00E+00 | 3.69E-03 | NaN | ITGB7,PLN |
| Systemic Lupus Erythematosus In T Cell Signaling P | 0.00E+00 | 1.56E-03 | NaN | ICOSLG/LOC102723996 |
| Xenobiotic Metabolism CAR Signaling Pathway | 0.00E+00 | 5.21E-03 | NaN | GSTA1 |
| Xenobiotic Metabolism PXR Signaling Pathway | 0.00E+00 | 5.18E-03 | NaN | GSTA1 |
| Regulation Of The Epithelial Mesenchymal Transiti | 0.00E+00 | 5.21E-03 | NaN | EGR1 |
| Coronavirus Pathogenesis Pathway | 0.00E+00 | 4.90E-03 | NaN | CCL2 |
| Ephrin Receptor Signaling | 0.00E+00 | 4.95E-03 | NaN | ITGB7 |

IPA_Venn_List_FB1_FC2_adj0.05_Astro_Micro_143genes

| | Ingenuity Canonical Pathways | -log(p-value) | Ratio | z-score | Molecules |
|---|---|---|---|---|---|
| 1 | Hepatic Fibrosis / Hepatic Stellate Cell Activation | 2.74E+00 | 2.58E-02 | NaN | CCN2,EDN1,IGF2,MYH2,NGFR |
| 2 | Superpathway of Cholesterol Biosynthesis | 2.13E+00 | 6.90E-02 | NaN | HMGCS1,MSMO1 |
| 3 | ABRA Signaling Pathway | 2.09E+00 | 3.26E-02 | NaN | CCN2,FOSB,TPM2 |
| 4 | Granulocyte Adhesion and Diapedesis | 1.99E+00 | 2.12E-02 | NaN | CLDN5,CXCL14,HRH4,NGFR |
| 5 | Melatonin Degradation II | 1.75E+00 | 2.50E-01 | NaN | MAOB |
| 6 | Zymosterol Biosynthesis | 1.58E+00 | 1.67E-01 | NaN | MSMO1 |
| 7 | NAD Biosynthesis III | 1.58E+00 | 1.67E-01 | NaN | NMNAT2 |
| 8 | LPS/IL-1 Mediated Inhibition of RXR Function | 1.56E+00 | 1.57E-02 | NaN | ALDH1L2,HMGCS1,MAOB,NGFR |
| 9 | NAD Biosynthesis from 2-amino-3-carboxymuconate Semialdehyde | 1.51E+00 | 1.43E-01 | NaN | NMNAT2 |
| 10 | NAD Salvage Pathway III | 1.51E+00 | 1.43E-01 | NaN | NMNAT2 |
| | Embryonic Stem Cell Differentiation into Cardiac Lineages | 1.36E+00 | 1.00E-01 | NaN | NKX2-5 |
| | Tight Junction Signaling | 1.33E+00 | 1.68E-02 | NaN | CLDN5,MYH2,NGFR |
| | Ketogenesis | 1.32E+00 | 9.09E-02 | NaN | HMGCS1 |
| | BEX2 Signaling Pathway | 1.28E+00 | 2.44E-02 | NaN | BEX2,NGFR |
| | Regulation Of The Epithelial Mesenchymal Transition By Growth Factors | 1.26E+00 | 1.56E-02 | NaN | FGF13,NGFR,TWIST1 |
| | NAD biosynthesis II (from tryptophan) | 1.25E+00 | 7.69E-02 | NaN | NMNAT2 |
| | Cholesterol Biosynthesis I | 1.25E+00 | 7.69E-02 | NaN | MSMO1 |
| | γ-glutamyl Cycle | 1.25E+00 | 7.69E-02 | NaN | CHAC1 |
| | Cholesterol Biosynthesis II (via 24,25-dihydrolanosterol) | 1.25E+00 | 7.69E-02 | NaN | MSMO1 |
| | Cholesterol Biosynthesis III (via Desmosterol) | 1.25E+00 | 7.69E-02 | NaN | MSMO1 |
| | Role of Hypercytokinemia/hyperchemokinemia in the Pathogenesis of Ir | 1.25E+00 | 2.33E-02 | NaN | AREG,ISG20 |
| | Mevalonate Pathway I | 1.22E+00 | 7.14E-02 | NaN | HMGCS1 |
| | Phenylalanine Degradation IV (Mammalian, via Side Chain) | 1.22E+00 | 7.14E-02 | NaN | MAOB |
| | ID1 Signaling Pathway | 1.21E+00 | 1.49E-02 | NaN | CCN2,IGF2,NGFR |
| | Gustation Pathway | 1.20E+00 | 1.48E-02 | NaN | GABRA2,GABRQ,SCN7A |
| | Agranulocyte Adhesion and Diapedesis | 1.17E+00 | 1.43E-02 | NaN | CLDN5,CXCL14,MYH2 |
| | Salvage Pathways of Pyrimidine Ribonucleotides | 1.16E+00 | 2.06E-02 | NaN | AK7,MAK |
| | Activin Inhibin Signaling Pathway | 1.13E+00 | 1.38E-02 | NaN | CCN2,FOSB,NGFR |
| | Superpathway of Geranylgeranyldiphosphate Biosynthesis I (via Mevalo | 1.11E+00 | 5.56E-02 | NaN | HMGCS1 |
| | Axonal Guidance Signaling | 1.11E+00 | 9.80E-03 | NaN | ADAM19,ADAMTS12,ADAMTS9,NGFR,PAPPA |
| | Neurovascular Coupling Signaling Pathway | 1.07E+00 | 1.29E-02 | NaN | GABRA2,GABRQ,KCNJ4 |
| | CDK5 Signaling | 1.03E+00 | 1.74E-02 | NaN | FOSB,NGFR |
| | Putrescine Degradation III | 1.03E+00 | 4.55E-02 | NaN | MAOB |
| | Cardiac Hypertrophy Signaling (Enhanced) | 1.02E+00 | 9.23E-03 | NaN | EDN1,FGF13,MAP3K15,NGFR,NKX2-5 |
| | Airway Pathology in Chronic Obstructive Pulmonary Disease | 1.01E+00 | 1.69E-02 | NaN | FGF13,LCN9 |
| | Pyrimidine Deoxyribonucleotides De Novo Biosynthesis I | 1.01E+00 | 4.35E-02 | NaN | AK7 |
| | Apelin Cardiac Fibroblast Signaling Pathway | 1.01E+00 | 4.35E-02 | NaN | CCN2 |
| | NGF Signaling | 9.98E-01 | 1.67E-02 | NaN | MAP3K15,NGFR |
| | Cardiomyocyte Differentiation via BMP Receptors | 9.44E-01 | 3.70E-02 | NaN | NKX2-5 |
| | NAD Salvage Pathway II | 9.44E-01 | 3.70E-02 | NaN | NMNAT2 |
| | Tryptophan Degradation X (Mammalian, via Tryptamine) | 9.44E-01 | 3.70E-02 | NaN | MAOB |
| | Apelin Liver Signaling Pathway | 9.44E-01 | 3.70E-02 | NaN | EDN1 |
| | GABA Receptor Signaling | 9.29E-01 | 1.52E-02 | NaN | GABRA2,GABRQ |
| | Ferroptosis Signaling Pathway | 9.29E-01 | 1.52E-02 | NaN | CHAC1,SLC7A11 |
| | Opioid Signaling Pathway | 8.86E-01 | 1.07E-02 | NaN | FOSB,RGS4,SCN7A |
| | Dopamine Degradation | 8.75E-01 | 3.12E-02 | NaN | MAOB |
| | Endocannabinoid Cancer Inhibition Pathway | 8.53E-01 | 1.36E-02 | NaN | NUPR1,TWIST1 |
| | Xenobiotic Metabolism Signaling | 8.47E-01 | 1.03E-02 | NaN | ALDH1L2,MAOB,MAP3K15 |
| | Noradrenaline and Adrenaline Degradation | 8.16E-01 | 2.70E-02 | NaN | MAOB |
| | Pyrimidine Ribonucleotides Interconversion | 7.95E-01 | 2.56E-02 | NaN | AK7 |
| | Inhibition of Matrix Metalloproteases | 7.95E-01 | 2.56E-02 | NaN | TFPI2 |
| | Pyrimidine Ribonucleotides De Novo Biosynthesis | 7.66E-01 | 2.38E-02 | NaN | AK7 |
| | Pulmonary Fibrosis Idiopathic Signaling Pathway | 7.48E-01 | 9.20E-03 | NaN | AREG,CCN2,EDN1 |
| | Aldosterone Signaling in Epithelial Cells | 7.47E-01 | 1.16E-02 | NaN | CRYAB,DNAJB13 |
| | Ephrin A Signaling | 7.22E-01 | 2.13E-02 | NaN | NGFR |
| | Tumor Microenvironment Pathway | 7.20E-01 | 1.12E-02 | NaN | FGF13,IGF2 |
| | Acute Phase Response Signaling | 6.99E-01 | 1.08E-02 | NaN | CP,NGFR |
| | MYC Mediated Apoptosis Signaling | 6.98E-01 | 2.00E-02 | NaN | NGFR |
| | FAT10 Cancer Signaling Pathway | 6.98E-01 | 2.00E-02 | NaN | NGFR |
| | Production of Nitric Oxide and Reactive Oxygen Species in Macrophages | 6.78E-01 | 1.05E-02 | NaN | MAP3K15,NGFR |
| | Xenobiotic Metabolism PXR Signaling Pathway | 6.71E-01 | 1.04E-02 | NaN | ALDH1L2,MAOB |
| | Role of Cytokines in Mediating Communication between Immune Cells | 6.68E-01 | 1.85E-02 | NaN | IL32 |
| | Regulation of the Epithelial-Mesenchymal Transition Pathway | 6.65E-01 | 1.03E-02 | NaN | FGF13,TWIST1 |
| | CD27 Signaling in Lymphocytes | 6.47E-01 | 1.75E-02 | NaN | MAP3K15 |
| | Sertoli Cell-Sertoli Cell Junction Signaling | 6.30E-01 | 9.71E-03 | NaN | CLDN5,MAP3K15 |

| | | | | |
|---|---|---|---|---|
| SPINK1 Pancreatic Cancer Pathway | 6.28E-01 | 1.67E-02 | NaN | CPA4 |
| Role of Tissue Factor in Cancer | 6.27E-01 | 9.66E-03 | NaN | CCN2,NGFR |
| HIF1α Signaling | 6.24E-01 | 9.62E-03 | NaN | EDN1,IGF2 |
| Induction of Apoptosis by HIV1 | 5.97E-01 | 1.54E-02 | NaN | NGFR |
| Pyridoxal 5'-phosphate Salvage Pathway | 5.97E-01 | 1.54E-02 | NaN | MAK |
| Calcium Signaling | 5.89E-01 | 9.09E-03 | NaN | MYH2,TPM2 |
| Nicotine Degradation II | 5.86E-01 | 1.49E-02 | NaN | INMT |
| Superpathway of Melatonin Degradation | 5.81E-01 | 1.47E-02 | NaN | MAOB |
| Growth Hormone Signaling | 5.65E-01 | 1.41E-02 | NaN | IGF2 |
| Serotonin Degradation | 5.59E-01 | 1.39E-02 | NaN | MAOB |
| GDNF Family Ligand-Receptor Interactions | 5.40E-01 | 1.32E-02 | NaN | DOK7 |
| Hypoxia Signaling in the Cardiovascular System | 5.40E-01 | 1.32E-02 | NaN | EDN1 |
| Hepatic Fibrosis Signaling Pathway | 5.33E-01 | 7.09E-03 | NaN | CCN2,EDN1,NGFR |
| Neurotrophin/TRK Signaling | 5.30E-01 | 1.28E-02 | NaN | NGFR |
| Actin Cytoskeleton Signaling | 5.27E-01 | 8.20E-03 | NaN | FGF13,MYH2 |
| Dopamine Receptor Signaling | 5.21E-01 | 1.25E-02 | NaN | MAOB |
| FGF Signaling | 4.95E-01 | 1.16E-02 | NaN | FGF13 |
| Xenobiotic Metabolism AHR Signaling Pathway | 4.91E-01 | 1.15E-02 | NaN | ALDH1L2 |
| Regulation Of The Epithelial Mesenchymal Transition In Development Pa | 4.91E-01 | 1.15E-02 | NaN | TWIST1 |
| Cardiac Hypertrophy Signaling | 4.88E-01 | 7.66E-03 | NaN | MAP3K15,NKX2-5 |
| Immunogenic Cell Death Signaling Pathway | 4.79E-01 | 1.11E-02 | NaN | NGFR |
| Unfolded protein response | 4.79E-01 | 1.11E-02 | NaN | DNAJB13 |
| Ceramide Signaling | 4.75E-01 | 1.10E-02 | NaN | NGFR |
| RANK Signaling in Osteoclasts | 4.75E-01 | 1.10E-02 | NaN | MAP3K15 |
| BMP signaling pathway | 4.75E-01 | 1.10E-02 | NaN | NKX2-5 |
| Pyroptosis Signaling Pathway | 4.67E-01 | 1.08E-02 | NaN | NGFR |
| ERBB Signaling | 4.67E-01 | 1.08E-02 | NaN | AREG |
| Protein Ubiquitination Pathway | 4.62E-01 | 7.33E-03 | NaN | CRYAB,DNAJB13 |
| TGF-β Signaling | 4.56E-01 | 1.04E-02 | NaN | NKX2-5 |
| Sumoylation Pathway | 4.32E-01 | 9.71E-03 | NaN | ISG20 |
| IGF-1 Signaling | 4.25E-01 | 9.52E-03 | NaN | CCN2 |
| Phagosome Formation | 4.25E-01 | 5.76E-03 | NaN | ADGRL3,HRH4,MYH2,SCARA3 |
| PPAR Signaling | 4.19E-01 | 9.35E-03 | NaN | NGFR |
| PD-1, PD-L1 cancer immunotherapy pathway | 4.19E-01 | 9.35E-03 | NaN | NGFR |
| G-Protein Coupled Receptor Signaling | 4.15E-01 | 5.69E-03 | NaN | ADGRL3,HRH4,MAP3K15,RGS4 |
| Role Of Osteoclasts In Rheumatoid Arthritis Signaling Pathway | 3.96E-01 | 6.47E-03 | NaN | ADAM19,NGFR |
| Bladder Cancer Signaling | 3.92E-01 | 8.62E-03 | NaN | FGF13 |
| Neuregulin Signaling | 3.89E-01 | 8.55E-03 | NaN | AREG |
| Neuroinflammation Signaling Pathway | 3.82E-01 | 6.31E-03 | NaN | GABRA2,GABRQ |
| LXR/RXR Activation | 3.73E-01 | 8.13E-03 | NaN | NGFR |
| Glioma Signaling | 3.67E-01 | 8.00E-03 | NaN | IGF2 |
| Myelination Signaling Pathway | 3.66E-01 | 6.12E-03 | NaN | NGFR,PLP1 |
| Role of Macrophages, Fibroblasts and Endothelial Cells in Rheumatoid A | 3.59E-01 | 6.02E-03 | NaN | IL32,NGFR |
| IL-6 Signaling | 3.57E-01 | 7.75E-03 | NaN | NGFR |
| HGF Signaling | 3.50E-01 | 7.58E-03 | NaN | MAP3K15 |
| S100 Family Signaling Pathway | 3.43E-01 | 5.18E-03 | NaN | ADGRL3,AREG,HRH4,TPM2 |
| STAT3 Pathway | 3.43E-01 | 7.41E-03 | NaN | NGFR |
| SNARE Signaling Pathway | 3.40E-01 | 7.35E-03 | NaN | MYH2 |
| Role of PKR in Interferon Induction and Antiviral Response | 3.40E-01 | 7.35E-03 | NaN | SCARA3 |
| Iron homeostasis signaling pathway | 3.36E-01 | 7.25E-03 | NaN | CP |
| Gαi Signaling | 3.31E-01 | 7.14E-03 | NaN | RGS4 |
| Role Of Chondrocytes In Rheumatoid Arthritis Signaling Pathway | 3.29E-01 | 7.09E-03 | NaN | NGFR |
| Xenobiotic Metabolism General Signaling Pathway | 3.25E-01 | 6.99E-03 | NaN | MAP3K15 |
| Dilated Cardiomyopathy Signaling Pathway | 3.10E-01 | 6.67E-03 | NaN | MYH2 |
| Cellular Effects of Sildenafil (Viagra) | 3.10E-01 | 6.67E-03 | NaN | MYH2 |
| NAD Signaling Pathway | 3.08E-01 | 6.62E-03 | NaN | NMNAT2 |
| PTEN Signaling | 3.08E-01 | 6.62E-03 | NaN | NGFR |
| Pathogen Induced Cytokine Storm Signaling Pathway | 3.05E-01 | 5.39E-03 | NaN | CXCL14,NGFR |
| Factors Promoting Cardiogenesis in Vertebrates | 3.04E-01 | 6.54E-03 | NaN | NKX2-5 |
| Type II Diabetes Mellitus Signaling | 3.04E-01 | 6.54E-03 | NaN | NGFR |
| Necroptosis Signaling Pathway | 2.98E-01 | 6.41E-03 | NaN | NGFR |
| Epithelial Adherens Junction Signaling | 2.96E-01 | 6.37E-03 | NaN | MYH2 |
| Ovarian Cancer Signaling | 2.94E-01 | 6.33E-03 | NaN | EDN1 |
| Aryl Hydrocarbon Receptor Signaling | 2.92E-01 | 6.29E-03 | NaN | ALDH1L2 |
| CREB Signaling in Neurons | 2.91E-01 | 4.94E-03 | NaN | ADGRL3,HRH4,NGFR |
| Inhibition of ARE-Mediated mRNA Degradation Pathway | 2.87E-01 | 6.17E-03 | NaN | NGFR |
| HOTAIR Regulatory Pathway | 2.85E-01 | 6.13E-03 | NaN | TWIST1 |
| Transcriptional Regulatory Network in Embryonic Stem Cells | 2.83E-01 | 6.10E-03 | NaN | IGF2 |
| HMGB1 Signaling | 2.78E-01 | 5.99E-03 | NaN | NGFR |
| Germ Cell-Sertoli Cell Junction Signaling | 2.73E-01 | 5.88E-03 | NaN | MAP3K15 |
| Gαq Signaling | 2.73E-01 | 5.88E-03 | NaN | RGS4 |
| Glioblastoma Multiforme Signaling | 2.71E-01 | 5.85E-03 | NaN | IGF2 |

| | | | | |
|---|---|---|---|---|
| Protein Kinase A Signaling | 2.60E-01 | 4.87E-03 | NaN | MYH2,NGFR |
| IL-33 Signaling Pathway | 2.47E-01 | 5.38E-03 | NaN | AREG |
| Dopamine-DARPP32 Feedback in cAMP Signaling | 2.47E-01 | 5.38E-03 | NaN | KCNJ4 |
| Hepatic Cholestasis | 2.40E-01 | 5.24E-03 | NaN | NGFR |
| GNRH Signaling | 2.40E-01 | 5.24E-03 | NaN | MAP3K15 |
| Xenobiotic Metabolism CAR Signaling Pathway | 2.38E-01 | 5.21E-03 | NaN | ALDH1L2 |
| Leukocyte Extravasation Signaling | 2.37E-01 | 5.18E-03 | NaN | CLDN5 |
| Endothelin-1 Signaling | 2.35E-01 | 5.15E-03 | NaN | EDN1 |
| Natural Killer Cell Signaling | 2.30E-01 | 5.05E-03 | NaN | MAP3K15 |
| Pulmonary Healing Signaling Pathway | 2.28E-01 | 5.03E-03 | NaN | NGFR |
| ILK Signaling | 2.26E-01 | 4.98E-03 | NaN | MYH2 |
| Clathrin-mediated Endocytosis Signaling | 2.17E-01 | 4.81E-03 | NaN | FGF13 |
| Autophagy | 2.07E-01 | 4.63E-03 | NaN | NGFR |
| RHOGDI Signaling | 2.02E-01 | 4.55E-03 | NaN | MYH2 |
| Multiple Sclerosis Signaling Pathway | 2.00E-01 | 4.50E-03 | NaN | PLP1 |
| Oxytocin Signaling Pathway | 0.00E+00 | 3.55E-03 | NaN | MYH2 |
| Wound Healing Signaling Pathway | 0.00E+00 | 3.97E-03 | NaN | NGFR |
| CLEAR Signaling Pathway | 0.00E+00 | 3.51E-03 | NaN | NGFR |
| Circadian Rhythm Signaling | 0.00E+00 | 3.73E-03 | NaN | NGFR |
| NRF2-mediated Oxidative Stress Response | 0.00E+00 | 4.22E-03 | NaN | DNAJB13 |
| Mitochondrial Dysfunction | 0.00E+00 | 2.91E-03 | NaN | MAOB |
| T Helper Cell Differentiation | 0.00E+00 | 2.12E-03 | NaN | NGFR |
| Dendritic Cell Maturation | 0.00E+00 | 3.36E-03 | NaN | IL32,NGFR |
| Type I Diabetes Mellitus Signaling | 0.00E+00 | 1.96E-03 | NaN | NGFR |
| FAK Signaling | 0.00E+00 | 1.92E-03 | NaN | ADGRL3,HRH4 |
| AMPK Signaling | 0.00E+00 | 4.13E-03 | NaN | AK7 |
| Role of Osteoblasts, Osteoclasts and Chondrocytes in Rheumatoid Arthri | 0.00E+00 | 4.39E-03 | NaN | NGFR |
| HER-2 Signaling in Breast Cancer | 0.00E+00 | 4.41E-03 | NaN | AREG |
| Role of NFAT in Cardiac Hypertrophy | 0.00E+00 | 4.46E-03 | NaN | NKX2-5 |
| Breast Cancer Regulation by Stathmin1 | 0.00E+00 | 3.37E-03 | NaN | ADGRL3,HRH4 |
| PKCθ Signaling in T Lymphocytes | 0.00E+00 | 1.79E-03 | NaN | MAP3K15 |
| Estrogen Receptor Signaling | 0.00E+00 | 2.44E-03 | NaN | IGF2 |
| B Cell Receptor Signaling | 0.00E+00 | 1.57E-03 | NaN | MAP3K15 |
| Serotonin Receptor Signaling | 0.00E+00 | 4.26E-03 | NaN | EDN1,MAOB |
| cAMP-mediated signaling | 0.00E+00 | 4.24E-03 | NaN | RGS4 |
| NF-κB Signaling | 0.00E+00 | 1.75E-03 | NaN | NGFR |
| Systemic Lupus Erythematosus In B Cell Signaling Pathway | 0.00E+00 | 1.38E-03 | NaN | ISG20 |
