## Supplementary material for "Unravelling neuronal and glial differences in ceramide composition, synthesis, and sensitivity to toxicity": Document1_Venn_List Astrocyte FB1-up GCSi-down

| GeneID | name | | Document1_Venn_List Astrocyte FB1-up GCSi-down |
|---|---|---|---|
| ENSG0000 | PDK4 | | |
| ENSG0000 | KLHDC7B | | |
| ENSG0000 | AQP5 | | |
| ENSG0000 | ANGPTL4 | | |
| ENSG0000 | FIBIN | | |
| ENSG0000 | ERICH2 | | |

| GeneID | Treatment | Treatment | Treatment | Treatment | name | Astrocytes_FB1_FC+2_adj0.05_831 |
|---|---|---|---|---|---|---|
| ENSG0000 | 3.216286 | 9.29391 | 0.002447 | 0.013277 | CASP10 | |
| ENSG0000 | 1.097686 | 2.140111 | 1.73E-05 | 0.000169 | PDK4 | |
| ENSG0000 | 5.177892 | 36.19935 | 8.22E-26 | 6.74E-24 | TAC1 | |
| ENSG0000 | 2.192091 | 4.569672 | 4.36E-220 | 9.59E-217 | IFRD1 | |
| ENSG0000 | 3.322271 | 10.00238 | 0.000827 | 0.005212 | SCN4A | |
| ENSG0000 | 1.118767 | 2.171613 | 3.57E-06 | 4.02E-05 | MAPK8IP2 | |
| ENSG0000 | 1.703892 | 3.257787 | 3.23E-05 | 0.000294 | ETV7 | |
| ENSG0000 | 2.083605 | 4.238649 | 0.0036 | 0.018333 | GPRC5A | |
| ENSG0000 | 1.607582 | 3.047406 | 0.00068 | 0.004385 | ADGRA2 | |
| ENSG0000 | 4.326747 | 20.06692 | 2.89E-117 | 1.91E-114 | KCNG1 | |
| ENSG0000 | 6.215906 | 74.33171 | 0.005365 | 0.025611 | SH2D2A | |
| ENSG0000 | 1.993289 | 3.981437 | 7.42E-161 | 7.25E-158 | SARS1 | |
| ENSG0000 | 1.402133 | 2.64292 | 4.62E-39 | 7.39E-37 | GAB2 | |
| ENSG0000 | 7.27549 | 154.9319 | 3.65E-05 | 0.000329 | TNFRSF9 | |
| ENSG0000 | 2.029585 | 4.082873 | 2.80E-88 | 1.27E-85 | HERPUD1 | |
| ENSG0000 | 1.367377 | 2.58001 | 2.17E-11 | 5.50E-10 | PTPRN | |
| ENSG0000 | 2.777962 | 6.858828 | 0.007455 | 0.033608 | MCOLN3 | |
| ENSG0000 | 1.229975 | 2.345629 | 1.38E-11 | 3.59E-10 | TMCC3 | |
| ENSG0000 | 1.545384 | 2.918818 | 5.60E-06 | 6.06E-05 | TBXAS1 | |
| ENSG0000 | 3.501027 | 11.32176 | 3.09E-197 | 4.29E-194 | BCAT1 | |
| ENSG0000 | 2.708126 | 6.534723 | 0.00465 | 0.02271 | DMRT3 | |
| ENSG0000 | 1.313369 | 2.485212 | 8.64E-32 | 9.71E-30 | UHRF1BP1 | |
| ENSG0000 | 1.541002 | 2.909965 | 3.83E-12 | 1.06E-10 | TLE2 | |
| ENSG0000 | 4.015604 | 16.17399 | 0 | 0 | MTHFD2 | |
| ENSG0000 | 1.022706 | 2.031727 | 3.38E-12 | 9.49E-11 | WIPI1 | |
| ENSG0000 | 4.532646 | 23.14529 | 0 | 0 | ASNS | |
| ENSG0000 | 3.588442 | 12.02898 | 0.009857 | 0.04249 | EPHA8 | |
| ENSG0000 | 2.586952 | 6.00828 | 1.09E-120 | 7.39E-118 | P4HA2 | |
| ENSG0000 | 3.922202 | 15.16004 | 1.65E-10 | 3.75E-09 | MOV10L1 | |
| ENSG0000 | 2.082943 | 4.236707 | 4.98E-08 | 7.83E-07 | PANX2 | |
| ENSG0000 | 1.177696 | 2.262152 | 1.11E-14 | 4.00E-13 | ALPK1 | |
| ENSG0000 | 1.710792 | 3.273404 | 2.39E-52 | 5.44E-50 | TUBE1 | |
| ENSG0000 | 3.108453 | 8.624573 | 4.12E-18 | 2.02E-16 | MOCOS | |
| ENSG0000 | 1.745327 | 3.352708 | 0.011775 | 0.049228 | CAMSAP3 | |
| ENSG0000 | 2.718681 | 6.582705 | 0.004672 | 0.022806 | STXBP2 | |
| ENSG0000 | 1.03823 | 2.053707 | 0.001917 | 0.010762 | EXOSC5 | |
| ENSG0000 | 6.127253 | 69.90159 | 3.30E-201 | 5.80E-198 | LAMP3 | |
| ENSG0000 | 1.060338 | 2.085419 | 4.86E-35 | 6.58E-33 | TULP3 | |
| ENSG0000 | 1.345723 | 2.541575 | 0.000246 | 0.001805 | DCT | |
| ENSG0000 | 2.690197 | 6.454015 | 0.004938 | 0.023858 | EPHA6 | |
| ENSG0000 | 1.141296 | 2.205791 | 6.96E-12 | 1.88E-10 | ARG2 | |
| ENSG0000 | 1.311063 | 2.481242 | 2.33E-06 | 2.73E-05 | GCKR | |
| ENSG0000 | 2.47517 | 5.560328 | 0.001147 | 0.006948 | TTC39A | |
| ENSG0000 | 3.761752 | 13.56439 | 0.001076 | 0.006566 | PPEF1 | |
| ENSG0000 | 1.484289 | 2.797792 | 2.77E-19 | 1.52E-17 | PPP1R15A | |
| ENSG0000 | 1.422043 | 2.679647 | 3.56E-23 | 2.60E-21 | HSD17B14 | |
| ENSG0000 | 1.943584 | 3.846601 | 8.42E-102 | 4.83E-99 | FTL | |
| ENSG0000 | 2.826329 | 7.092669 | 0.000593 | 0.003904 | CASS4 | |
| ENSG0000 | 1.346145 | 2.542319 | 7.66E-22 | 5.13E-20 | SMOX | |

| | | | | | |
|---|---|---|---|---|---|
| ENSG0000 | 2.545048 | 5.836274 | 0.00262 | 0.014085 | FXYD5 |
| ENSG0000 | 1.304372 | 2.469762 | 8.90E-29 | 8.54E-27 | BLVRB |
| ENSG0000 | 1.279219 | 2.427075 | 0.000518 | 0.003468 | TFAP4 |
| ENSG0000 | 3.164362 | 8.965362 | 1.33E-21 | 8.71E-20 | RAPGEF4 |
| ENSG0000 | 1.660092 | 3.160366 | 7.07E-38 | 1.10E-35 | PITPNM3 |
| ENSG0000 | 2.020739 | 4.057916 | 7.58E-05 | 0.000634 | TBL1Y |
| ENSG0000 | 1.816426 | 3.522075 | 1.03E-74 | 3.76E-72 | PHGDH |
| ENSG0000 | 1.704907 | 3.260079 | 1.46E-29 | 1.48E-27 | EPB41L4B |
| ENSG0000 | 5.075796 | 33.72616 | 0.004154 | 0.020652 | CYP26A1 |
| ENSG0000 | 1.885658 | 3.695215 | 6.08E-88 | 2.72E-85 | MKNK2 |
| ENSG0000 | 1.161806 | 2.237374 | 2.90E-36 | 4.18E-34 | XBP1 |
| ENSG0000 | 6.461473 | 88.12461 | 1.02E-06 | 1.28E-05 | MIOX |
| ENSG0000 | 3.062024 | 8.351435 | 2.07E-60 | 5.63E-58 | HMOX1 |
| ENSG0000 | 4.118836 | 17.37374 | 1.72E-06 | 2.06E-05 | IL2RB |
| ENSG0000 | 3.442125 | 10.86883 | 1.36E-06 | 1.67E-05 | KCNK10 |
| ENSG0000 | 5.628153 | 49.45872 | 3.63E-258 | 2.39E-254 | PCK2 |
| ENSG0000 | 1.137119 | 2.199414 | 5.72E-15 | 2.14E-13 | PABPC1L |
| ENSG0000 | 1.135342 | 2.196707 | 1.50E-19 | 8.43E-18 | DOK5 |
| ENSG0000 | 2.081374 | 4.232101 | 5.23E-06 | 5.69E-05 | EEF1A2 |
| ENSG0000 | 4.913376 | 30.13516 | 3.11E-222 | 7.45E-219 | TRIB3 |
| ENSG0000 | 6.619119 | 98.3 | 4.11E-06 | 4.59E-05 | TLDC2 |
| ENSG0000 | 2.142516 | 4.415315 | 0.009919 | 0.042689 | RNF125 |
| ENSG0000 | 1.229888 | 2.345488 | 5.11E-57 | 1.30E-54 | TBL1X |
| ENSG0000 | 5.741331 | 53.49496 | 0.00067 | 0.004337 | RHOXF1 |
| ENSG0000 | 1.0944 | 2.135242 | 4.18E-09 | 7.76E-08 | MOSPD1 |
| ENSG0000 | 1.610873 | 3.054365 | 7.02E-17 | 3.12E-15 | RENBP |
| ENSG0000 | 1.013122 | 2.018274 | 0.003644 | 0.01851 | MTMR8 |
| ENSG0000 | 1.724957 | 3.305702 | 0.001158 | 0.007012 | RUBCNL |
| ENSG0000 | 4.648539 | 25.08128 | 1.28E-41 | 2.17E-39 | SGCG |
| ENSG0000 | 1.813793 | 3.515653 | 8.48E-05 | 0.000701 | FOXF1 |
| ENSG0000 | 1.703362 | 3.25659 | 2.72E-44 | 5.06E-42 | SLC7A5 |
| ENSG0000 | 5.46296 | 44.10774 | 3.87E-05 | 0.000348 | ESRP1 |
| ENSG0000 | 1.435929 | 2.705564 | 4.89E-08 | 7.71E-07 | TNFRSF10A |
| ENSG0000 | 3.830251 | 14.22396 | 0.010772 | 0.045798 | KLC3 |
| ENSG0000 | 5.486988 | 44.84851 | 0.009688 | 0.041884 | RSPH6A |
| ENSG0000 | 1.091628 | 2.131143 | 8.30E-31 | 9.01E-29 | TIMM44 |
| ENSG0000 | 4.419255 | 21.3958 | 0.003606 | 0.018346 | TNNT1 |
| ENSG0000 | 1.000939 | 2.001302 | 3.80E-20 | 2.23E-18 | ZNF419 |
| ENSG0000 | 5.005129 | 32.11397 | 1.15E-241 | 3.78E-238 | SLC1A5 |
| ENSG0000 | 2.093669 | 4.268322 | 8.91E-49 | 1.87E-46 | BBC3 |
| ENSG0000 | 1.528614 | 2.885085 | 2.94E-13 | 9.23E-12 | PLA2G4C |
| ENSG0000 | 4.534343 | 23.17252 | 0.007992 | 0.035676 | PIK3CG |
| ENSG0000 | 5.776674 | 54.82166 | 0.003338 | 0.017216 | ANKRD7 |
| ENSG0000 | 1.140809 | 2.205047 | 1.90E-56 | 4.68E-54 | GRB10 |
| ENSG0000 | 2.269063 | 4.820101 | 1.12E-202 | 2.10E-199 | GARS1 |
| ENSG0000 | 1.063491 | 2.089982 | 2.34E-24 | 1.81E-22 | AIMP2 |
| ENSG0000 | 6.679827 | 102.5247 | 0.010467 | 0.044724 | GALNTL5 |
| ENSG0000 | 1.953827 | 3.874009 | 1.03E-56 | 2.56E-54 | AKNA |
| ENSG0000 | 2.372143 | 5.177094 | 0.009538 | 0.041343 | DOCK8 |
| ENSG0000 | 1.049816 | 2.070266 | 4.18E-14 | 1.43E-12 | PTGDS |

| | | | | | |
|---|---|---|---|---|---|
| ENSG0000 | 3.013646 | 8.076029 | 4.99E-178 | 5.99E-175 | UNC5B |
| ENSG0000 | 1.430163 | 2.694772 | 4.25E-30 | 4.43E-28 | SPOCK2 |
| ENSG0000 | 1.037067 | 2.052052 | 3.18E-28 | 2.93E-26 | SH3PXD2A |
| ENSG0000 | 2.646458 | 6.261281 | 6.65E-24 | 5.03E-22 | DKK1 |
| ENSG0000 | 2.128881 | 4.373781 | 4.51E-09 | 8.29E-08 | SORCS1 |
| ENSG0000 | 1.723262 | 3.301821 | 7.52E-54 | 1.79E-51 | TRIM16L |
| ENSG0000 | 1.220575 | 2.330396 | 0.004712 | 0.022959 | ALDH3A1 |
| ENSG0000 | 2.332851 | 5.037999 | 1.40E-06 | 1.70E-05 | SLC16A6 |
| ENSG0000 | 6.794584 | 111.013 | 0.009193 | 0.040061 | GABRA4 |
| ENSG0000 | 6.363351 | 82.33024 | 0.000993 | 0.006111 | AREG |
| ENSG0000 | 1.415034 | 2.66666 | 1.50E-33 | 1.85E-31 | PPARGC1A |
| ENSG0000 | 1.467087 | 2.764632 | 5.75E-23 | 4.12E-21 | HTATIP2 |
| ENSG0000 | 1.072514 | 2.103095 | 0.003155 | 0.016439 | CTSC |
| ENSG0000 | 1.054397 | 2.07685 | 1.54E-21 | 1.00E-19 | CPT1A |
| ENSG0000 | 6.541455 | 93.14817 | 0.009733 | 0.04203 | FOLR3 |
| ENSG0000 | 1.546561 | 2.921199 | 4.63E-89 | 2.14E-86 | CARS1 |
| ENSG0000 | 6.401251 | 84.52178 | 3.15E-07 | 4.34E-06 | CALCA |
| ENSG0000 | 1.14818 | 2.216341 | 2.62E-32 | 3.10E-30 | ALDH2 |
| ENSG0000 | 2.77899 | 6.863715 | 9.36E-248 | 4.12E-244 | SLC38A1 |
| ENSG0000 | 3.518111 | 11.45663 | 8.89E-11 | 2.08E-09 | TPD52L1 |
| ENSG0000 | 1.847107 | 3.59778 | 9.14E-84 | 3.77E-81 | NCOA7 |
| ENSG0000 | 4.587375 | 24.04017 | 1.99E-133 | 1.46E-130 | ULBP1 |
| ENSG0000 | 1.135901 | 2.197557 | 4.60E-19 | 2.48E-17 | EYA4 |
| ENSG0000 | 1.48183 | 2.793028 | 0.000344 | 0.002419 | TRIM38 |
| ENSG0000 | 3.398914 | 10.54812 | 1.43E-230 | 4.19E-227 | VEGFA |
| ENSG0000 | 1.630105 | 3.095355 | 3.21E-151 | 2.83E-148 | HSPA9 |
| ENSG0000 | 1.28676 | 2.439795 | 0.00089 | 0.005563 | RASGRF2 |
| ENSG0000 | 1.149342 | 2.218127 | 1.09E-40 | 1.80E-38 | CCNG1 |
| ENSG0000 | 1.476107 | 2.78197 | 1.58E-211 | 3.21E-208 | TARS1 |
| ENSG0000 | 4.531501 | 23.12691 | 1.32E-200 | 2.17E-197 | STC2 |
| ENSG0000 | 1.63021 | 3.095581 | 4.18E-06 | 4.65E-05 | EFCC1 |
| ENSG0000 | 2.599238 | 6.059666 | 9.34E-17 | 4.09E-15 | PEX5L |
| ENSG0000 | 1.015414 | 2.021483 | 4.46E-36 | 6.40E-34 | EIF1B |
| ENSG0000 | 3.33947 | 10.12233 | 0.000227 | 0.001677 | IL1A |
| ENSG0000 | 1.35806 | 2.563402 | 1.89E-21 | 1.23E-19 | STEAP3 |
| ENSG0000 | 1.184118 | 2.272245 | 3.34E-32 | 3.88E-30 | MARK1 |
| ENSG0000 | 1.134149 | 2.194891 | 2.85E-28 | 2.63E-26 | ERRFI1 |
| ENSG0000 | 2.526042 | 5.759892 | 1.67E-106 | 1.03E-103 | ARHGEF2 |
| ENSG0000 | 1.037068 | 2.052053 | 0.000665 | 0.004309 | FBXO2 |
| ENSG0000 | 5.510165 | 45.5748 | 0.008005 | 0.035726 | NCF2 |
| ENSG0000 | 1.955819 | 3.879361 | 1.69E-17 | 7.90E-16 | PLA2G4A |
| ENSG0000 | 2.962816 | 7.796444 | 3.79E-75 | 1.41E-72 | CTH |
| ENSG0000 | 1.305315 | 2.471377 | 1.24E-38 | 1.95E-36 | SIPA1L2 |
| ENSG0000 | 2.255509 | 4.775027 | 3.53E-106 | 2.12E-103 | RIMS3 |
| ENSG0000 | 1.602119 | 3.035889 | 0.003735 | 0.018903 | SLC8A2 |
| ENSG0000 | 1.064985 | 2.092149 | 1.95E-11 | 4.97E-10 | RAB32 |
| ENSG0000 | 7.973795 | 251.392 | 0.000119 | 0.000951 | BSPRY |
| ENSG0000 | 2.020076 | 4.056052 | 6.74E-74 | 2.40E-71 | GOT1 |
| ENSG0000 | 1.395646 | 2.631064 | 4.13E-48 | 8.51E-46 | MTHFD1L |
| ENSG0000 | 1.335178 | 2.523067 | 4.59E-10 | 9.77E-09 | ARAP3 |

| | | | | | |
|---|---|---|---|---|---|
| ENSG0000 | 5.933854 | 61.13194 | 0.004044 | 0.0202 | TEX11 |
| ENSG0000 | 1.100618 | 2.144466 | 2.75E-14 | 9.55E-13 | DUSP4 |
| ENSG0000 | 1.397744 | 2.634893 | 1.08E-85 | 4.65E-83 | TNFRSF10B |
| ENSG0000 | 2.600235 | 6.063853 | 0.009217 | 0.040136 | TBX2 |
| ENSG0000 | 6.184548 | 72.73351 | 0.004346 | 0.02147 | TMEM156 |
| ENSG0000 | 1.303695 | 2.468604 | 0.008766 | 0.038485 | RASL11A |
| ENSG0000 | 2.950055 | 7.727786 | 0.001344 | 0.007999 | TWIST1 |
| ENSG0000 | 1.65821 | 3.156247 | 2.25E-30 | 2.40E-28 | KIAA1549 |
| ENSG0000 | 6.191418 | 73.08066 | 2.73E-34 | 3.52E-32 | ARHGAP9 |
| ENSG0000 | 5.392978 | 42.01925 | 0.000122 | 0.000971 | HOXC13 |
| ENSG0000 | 2.103464 | 4.297399 | 1.64E-66 | 5.14E-64 | CDKN1A |
| ENSG0000 | 3.008059 | 8.044816 | 0.008781 | 0.038537 | NRN1 |
| ENSG0000 | 2.892467 | 7.425391 | 5.96E-05 | 0.000511 | TNFSF9 |
| ENSG0000 | 1.023937 | 2.033461 | 5.48E-36 | 7.77E-34 | EML2 |
| ENSG0000 | 1.055064 | 2.077811 | 9.56E-07 | 1.21E-05 | BMP2 |
| ENSG0000 | 6.670854 | 101.889 | 0.001549 | 0.009006 | OVOL2 |
| ENSG0000 | 1.482641 | 2.794599 | 0.003317 | 0.017131 | CHURC1-FNTB |
| ENSG0000 | 1.612648 | 3.058127 | 3.08E-134 | 2.32E-131 | EIF2S2 |
| ENSG0000 | 2.459256 | 5.499332 | 0.000293 | 0.0021 | FLRT1 |
| ENSG0000 | 1.669792 | 3.181687 | 0.002132 | 0.011768 | EVI2A |
| ENSG0000 | 1.299532 | 2.46149 | 0.011568 | 0.048539 | LRRC61 |
| ENSG0000 | 1.10566 | 2.151974 | 0.010056 | 0.043212 | SPINK2 |
| ENSG0000 | 6.34124 | 81.07808 | 5.71E-147 | 4.57E-144 | ADM2 |
| ENSG0000 | 1.611628 | 3.055964 | 1.78E-91 | 8.84E-89 | ATF4 |
| ENSG0000 | 2.079024 | 4.225212 | 0.008251 | 0.036637 | FOXP2 |
| ENSG0000 | 2.172894 | 4.509269 | 5.61E-07 | 7.41E-06 | EIF2S2P4 |
| ENSG0000 | 5.347148 | 40.70539 | 1.52E-158 | 1.43E-155 | CHAC1 |
| ENSG0000 | 1.07143 | 2.101515 | 3.43E-18 | 1.70E-16 | QRICH2 |
| ENSG0000 | 3.935409 | 15.29946 | 7.59E-31 | 8.28E-29 | KCNA5 |
| ENSG0000 | 1.503419 | 2.835138 | 1.82E-05 | 0.000177 | NALF2 |
| ENSG0000 | 7.353912 | 163.5868 | 2.86E-12 | 8.08E-11 | KLHDC7B |
| ENSG0000 | 4.822761 | 28.3006 | 9.91E-173 | 1.09E-169 | GDF15 |
| ENSG0000 | 1.620989 | 3.075858 | 4.62E-08 | 7.33E-07 | NPAS1 |
| ENSG0000 | 2.988189 | 7.934774 | 3.72E-151 | 3.17E-148 | SESN2 |
| ENSG0000 | 1.970782 | 3.919804 | 0.006946 | 0.031737 | CCDC62 |
| ENSG0000 | 3.219607 | 9.31533 | 0.006279 | 0.029299 | DUSP9 |
| ENSG0000 | 1.301663 | 2.465129 | 0.000972 | 0.006001 | RAB11FIP4 |
| ENSG0000 | 1.549705 | 2.927572 | 6.28E-06 | 6.69E-05 | PPARG |
| ENSG0000 | 1.658484 | 3.156846 | 0.000572 | 0.003782 | EMILIN2 |
| ENSG0000 | 1.000359 | 2.000498 | 1.15E-28 | 1.09E-26 | KDM6B |
| ENSG0000 | 1.434911 | 2.703654 | 2.15E-21 | 1.39E-19 | ZSWIM3 |
| ENSG0000 | 3.892161 | 14.84763 | 2.20E-66 | 6.84E-64 | DMGDH |
| ENSG0000 | 2.063795 | 4.180847 | 1.97E-102 | 1.15E-99 | BEX2 |
| ENSG0000 | 1.312632 | 2.483943 | 0.00033 | 0.002331 | AMPD3 |
| ENSG0000 | 1.152936 | 2.22366 | 1.81E-34 | 2.36E-32 | SLC38A2 |
| ENSG0000 | 1.125208 | 2.181331 | 6.34E-59 | 1.69E-56 | NARS1 |
| ENSG0000 | 8.390574 | 335.5942 | 1.22E-06 | 1.51E-05 | HRH4 |
| ENSG0000 | 1.888813 | 3.703304 | 0.004494 | 0.022057 | CABLES1 |
| ENSG0000 | 1.856155 | 3.620414 | 0.001744 | 0.009936 | RAB33A |
| ENSG0000 | 5.156215 | 35.65951 | 0.01116 | 0.047112 | MTNR1B |

| | | | | | |
|---|---|---|---|---|---|
| ENSG0000 | 2.019991 | 4.055813 | 5.98E-199 | 8.76E-196 | YARS1 |
| ENSG0000 | 2.589451 | 6.018698 | 2.05E-162 | 2.16E-159 | PSAT1 |
| ENSG0000 | 2.604711 | 6.082695 | 0.005053 | 0.024345 | TBX3 |
| ENSG0000 | 3.49181 | 11.24967 | 5.46E-64 | 1.57E-61 | HRK |
| ENSG0000 | 2.108996 | 4.313909 | 3.55E-12 | 9.93E-11 | TES |
| ENSG0000 | 1.805097 | 3.494526 | 4.63E-21 | 2.91E-19 | FAIM2 |
| ENSG0000 | 1.334712 | 2.522251 | 0.004145 | 0.020614 | HEY2 |
| ENSG0000 | 1.301757 | 2.46529 | 4.77E-09 | 8.73E-08 | RGS8 |
| ENSG0000 | 3.59393 | 12.07482 | 2.26E-227 | 5.96E-224 | NIBAN1 |
| ENSG0000 | 1.458371 | 2.747979 | 3.16E-31 | 3.51E-29 | SERPINE2 |
| ENSG0000 | 3.318474 | 9.976087 | 5.57E-123 | 3.87E-120 | ALDH1L2 |
| ENSG0000 | 3.365609 | 10.3074 | 7.75E-16 | 3.15E-14 | GPNMB |
| ENSG0000 | 1.640044 | 3.116754 | 0.000386 | 0.002678 | SCN7A |
| ENSG0000 | 2.419906 | 5.351363 | 0.00893 | 0.039081 | GATA4 |
| ENSG0000 | 1.177684 | 2.262133 | 1.59E-70 | 5.23E-68 | EPRS1 |
| ENSG0000 | 1.344611 | 2.539618 | 1.88E-07 | 2.70E-06 | KLF4 |
| ENSG0000 | 1.746203 | 3.354744 | 2.64E-29 | 2.61E-27 | TLR4 |
| ENSG0000 | 1.346782 | 2.543442 | 1.15E-11 | 3.00E-10 | HMGA1 |
| ENSG0000 | 2.861205 | 7.266221 | 7.57E-16 | 3.08E-14 | CASP1 |
| ENSG0000 | 5.088523 | 34.02499 | 3.08E-08 | 5.00E-07 | CASP5 |
| ENSG0000 | 1.933234 | 3.819102 | 2.07E-06 | 2.44E-05 | HERC5 |
| ENSG0000 | 3.236297 | 9.42372 | 4.72E-13 | 1.45E-11 | GPAT3 |
| ENSG0000 | 1.456755 | 2.744903 | 1.47E-06 | 1.78E-05 | CDKL2 |
| ENSG0000 | 5.094502 | 34.1663 | 0.000152 | 0.001183 | C4orf17 |
| ENSG0000 | 2.676572 | 6.39335 | 3.03E-05 | 0.000278 | INHBE |
| ENSG0000 | 1.757348 | 3.38076 | 4.38E-68 | 1.39E-65 | SLC7A1 |
| ENSG0000 | 1.038821 | 2.054548 | 2.10E-18 | 1.06E-16 | WDFY2 |
| ENSG0000 | 3.207671 | 9.238579 | 0.004322 | 0.021373 | ZIC5 |
| ENSG0000 | 1.355064 | 2.558084 | 2.01E-14 | 7.09E-13 | JDP2 |
| ENSG0000 | 2.874877 | 7.335406 | 5.29E-190 | 6.65E-187 | WARS1 |
| ENSG0000 | 6.508305 | 91.0322 | 7.64E-05 | 0.000639 | FGF7 |
| ENSG0000 | 1.279988 | 2.42837 | 0.000227 | 0.001677 | CYP1A1 |
| ENSG0000 | 3.052726 | 8.297782 | 2.39E-62 | 6.72E-60 | OSGIN1 |
| ENSG0000 | 1.764371 | 3.397258 | 0.00644 | 0.029891 | SPATA22 |
| ENSG0000 | 3.045082 | 8.253936 | 0.002406 | 0.013082 | TAF4B |
| ENSG0000 | 1.059203 | 2.08378 | 1.83E-45 | 3.56E-43 | TP53 |
| ENSG0000 | 4.788267 | 27.63198 | 3.53E-16 | 1.49E-14 | PMAIP1 |
| ENSG0000 | 5.193618 | 36.59609 | 0.006698 | 0.030814 | VAV1 |
| ENSG0000 | 7.005351 | 128.4756 | 5.25E-05 | 0.000457 | TRPM2 |
| ENSG0000 | 1.454528 | 2.740668 | 2.39E-14 | 8.35E-13 | SLC47A1 |
| ENSG0000 | 1.516445 | 2.860852 | 6.28E-31 | 6.90E-29 | PRDM16 |
| ENSG0000 | 5.760523 | 54.21136 | 0.004673 | 0.022808 | IL22RA1 |
| ENSG0000 | 3.243119 | 9.468389 | 6.75E-07 | 8.78E-06 | SYTL1 |
| ENSG0000 | 1.217566 | 2.32554 | 3.04E-38 | 4.75E-36 | LMO4 |
| ENSG0000 | 1.707874 | 3.266792 | 6.87E-37 | 1.03E-34 | RGS16 |
| ENSG0000 | 1.313602 | 2.485614 | 4.93E-47 | 9.85E-45 | HAX1 |
| ENSG0000 | 2.108631 | 4.31282 | 7.88E-20 | 4.53E-18 | CNGA3 |
| ENSG0000 | 3.633524 | 12.4108 | 0.001012 | 0.006212 | PTH2R |
| ENSG0000 | 1.166768 | 2.245082 | 2.24E-09 | 4.30E-08 | ACKR3 |
| ENSG0000 | 1.058116 | 2.082211 | 7.05E-05 | 0.000594 | ITGA9 |

| | | | | | |
|---|---|---|---|---|---|
| ENSG0000 | 1.356579 | 2.560773 | 0.001097 | 0.006683 | SLC10A4 |
| ENSG0000 | 1.453621 | 2.738947 | 5.65E-15 | 2.11E-13 | MARCHF1 |
| ENSG0000 | 2.321659 | 4.999066 | 1.08E-05 | 0.00011 | NKD2 |
| ENSG0000 | 2.164667 | 4.48363 | 1.06E-16 | 4.62E-15 | TSLP |
| ENSG0000 | 1.804124 | 3.492172 | 1.65E-58 | 4.32E-56 | PSPH |
| ENSG0000 | 1.440926 | 2.71495 | 8.59E-48 | 1.76E-45 | SH3KBP1 |
| ENSG0000 | 1.288086 | 2.442038 | 3.20E-10 | 6.97E-09 | GCNA |
| ENSG0000 | 1.009225 | 2.012829 | 2.97E-05 | 0.000273 | DOCK11 |
| ENSG0000 | 4.757377 | 27.04663 | 0.005007 | 0.024151 | ATP6V0D2 |
| ENSG0000 | 1.112753 | 2.16258 | 2.86E-11 | 7.13E-10 | MAL2 |
| ENSG0000 | 1.529797 | 2.887452 | 0.00887 | 0.038882 | CDKN2A |
| ENSG0000 | 1.527864 | 2.883586 | 3.35E-33 | 4.09E-31 | FBXO10 |
| ENSG0000 | 1.380316 | 2.603254 | 1.02E-05 | 0.000104 | LCN9 |
| ENSG0000 | 1.548533 | 2.925194 | 1.33E-09 | 2.64E-08 | SLC39A12 |
| ENSG0000 | 3.654205 | 12.58999 | 0.009744 | 0.04206 | HTR7 |
| ENSG0000 | 2.049309 | 4.139076 | 0.000907 | 0.005651 | ADM |
| ENSG0000 | 1.413777 | 2.664338 | 0.004424 | 0.021773 | SLC43A1 |
| ENSG0000 | 1.073172 | 2.104054 | 0.000812 | 0.005124 | MPP7 |
| ENSG0000 | 1.228447 | 2.343146 | 1.37E-09 | 2.71E-08 | MIA2 |
| ENSG0000 | 3.372865 | 10.35937 | 3.59E-162 | 3.64E-159 | SLC7A11 |
| ENSG0000 | 1.209411 | 2.312432 | 9.07E-10 | 1.84E-08 | GABRA2 |
| ENSG0000 | 1.730285 | 3.317933 | 7.97E-26 | 6.55E-24 | KCNE4 |
| ENSG0000 | 1.30059 | 2.463296 | 2.67E-25 | 2.13E-23 | ZNF773 |
| ENSG0000 | 5.479278 | 44.60946 | 0.008715 | 0.038301 | DDX4 |
| ENSG0000 | 3.441969 | 10.86766 | 0.001706 | 0.00976 | CMTM7 |
| ENSG0000 | 1.348714 | 2.546851 | 7.38E-98 | 4.06E-95 | RMND5A |
| ENSG0000 | 2.303835 | 4.937685 | 0 | 0 | CEBPG |
| ENSG0000 | 1.06293 | 2.08917 | 0.003481 | 0.017843 | DGKE |
| ENSG0000 | 2.384051 | 5.220005 | 0.002719 | 0.014514 | LRRK1 |
| ENSG0000 | 1.411203 | 2.659589 | 9.16E-54 | 2.16E-51 | UCHL1 |
| ENSG0000 | 1.470976 | 2.772093 | 0.003822 | 0.019273 | GBP5 |
| ENSG0000 | 1.233145 | 2.350789 | 2.10E-55 | 5.04E-53 | SLC16A1 |
| ENSG0000 | 1.471651 | 2.77339 | 4.87E-43 | 8.86E-41 | RAB39B |
| ENSG0000 | 3.658469 | 12.62725 | 6.62E-13 | 2.00E-11 | FGF18 |
| ENSG0000 | 3.483109 | 11.18202 | 8.29E-99 | 4.66E-96 | PCDH1 |
| ENSG0000 | 1.697271 | 3.24287 | 4.46E-40 | 7.26E-38 | HKDC1 |
| ENSG0000 | 1.108369 | 2.156017 | 5.92E-21 | 3.67E-19 | NMNAT2 |
| ENSG0000 | 1.404618 | 2.647476 | 0.011756 | 0.049172 | STEAP2 |
| ENSG0000 | 1.021025 | 2.02936 | 3.34E-42 | 5.84E-40 | TSC22D3 |
| ENSG0000 | 2.818191 | 7.052776 | 5.24E-49 | 1.11E-46 | CREB3L1 |
| ENSG0000 | 1.388537 | 2.61813 | 1.70E-17 | 7.91E-16 | GAREM2 |
| ENSG0000 | 1.158094 | 2.231625 | 3.17E-05 | 0.000289 | SLC13A3 |
| ENSG0000 | 6.014109 | 64.62895 | 0.009473 | 0.041089 | ALAS2 |
| ENSG0000 | 1.123385 | 2.178575 | 9.06E-34 | 1.13E-31 | RCAN1 |
| ENSG0000 | 1.766583 | 3.402472 | 0.011486 | 0.048282 | SIM2 |
| ENSG0000 | 1.193189 | 2.286576 | 4.79E-57 | 1.23E-54 | HLCS |
| ENSG0000 | 2.44239 | 5.435415 | 4.05E-176 | 4.64E-173 | HK2 |
| ENSG0000 | 3.849076 | 14.41077 | 3.50E-05 | 0.000317 | ISL2 |
| ENSG0000 | 3.820705 | 14.13015 | 5.43E-92 | 2.75E-89 | CBS |
| ENSG0000 | 1.190018 | 2.281556 | 0.003013 | 0.015821 | ICOSLG |

| | | | | | |
|---|---|---|---|---|---|
| ENSG0000 | 1.016831 | 2.02347 | 1.03E-20 | 6.24E-19 | ZBTB7B |
| ENSG0000 | 1.4774 | 2.784466 | 1.87E-80 | 7.38E-78 | SQSTM1 |
| ENSG0000 | 6.812227 | 112.3788 | 0.007861 | 0.03519 | IKZF3 |
| ENSG0000 | 1.234866 | 2.353596 | 4.38E-16 | 1.84E-14 | FDXR |
| ENSG0000 | 1.31959 | 2.495952 | 0.000537 | 0.003585 | AQP5 |
| ENSG0000 | 1.39962 | 2.63832 | 5.26E-05 | 0.000458 | TAMALIN |
| ENSG0000 | 6.920399 | 121.1289 | 0.003649 | 0.018525 | PRR35 |
| ENSG0000 | 3.187707 | 9.111618 | 3.64E-06 | 4.10E-05 | TAL1 |
| ENSG0000 | 1.382682 | 2.607526 | 2.09E-15 | 8.12E-14 | DHRS3 |
| ENSG0000 | 8.223681 | 298.9336 | 0.004638 | 0.022662 | LHX8 |
| ENSG0000 | 1.441623 | 2.716262 | 5.76E-23 | 4.12E-21 | KCNT2 |
| ENSG0000 | 2.049972 | 4.140979 | 6.72E-66 | 2.04E-63 | DDR2 |
| ENSG0000 | 2.781577 | 6.876034 | 1.01E-77 | 3.87E-75 | ATF3 |
| ENSG0000 | 2.749863 | 6.726534 | 5.66E-57 | 1.42E-54 | LRATD1 |
| ENSG0000 | 2.783724 | 6.886278 | 0.000138 | 0.001088 | KCNJ3 |
| ENSG0000 | 1.626063 | 3.086695 | 1.91E-34 | 2.48E-32 | SPATA18 |
| ENSG0000 | 1.243928 | 2.368424 | 4.99E-16 | 2.07E-14 | IGFN1 |
| ENSG0000 | 5.780415 | 54.96399 | 0.011502 | 0.048336 | CRYBA2 |
| ENSG0000 | 1.420024 | 2.6759 | 1.51E-36 | 2.19E-34 | CADPS |
| ENSG0000 | 2.191792 | 4.568727 | 8.45E-45 | 1.60E-42 | RBM47 |
| ENSG0000 | 1.313754 | 2.485876 | 0.00932 | 0.040507 | IL17RE |
| ENSG0000 | 1.276314 | 2.422193 | 7.55E-05 | 0.000632 | UCN |
| ENSG0000 | 6.678484 | 102.4292 | 0.004371 | 0.021555 | SLC6A20 |
| ENSG0000 | 1.11447 | 2.165154 | 4.98E-14 | 1.69E-12 | KLF15 |
| ENSG0000 | 2.204728 | 4.609875 | 0.004155 | 0.020652 | ELOVL7 |
| ENSG0000 | 1.125975 | 2.18249 | 6.14E-15 | 2.28E-13 | NDUFAF2 |
| ENSG0000 | 2.230893 | 4.694245 | 2.99E-08 | 4.87E-07 | F2RL2 |
| ENSG0000 | 1.224091 | 2.336083 | 6.11E-36 | 8.62E-34 | GRPEL2 |
| ENSG0000 | 7.91872 | 241.976 | 7.27E-05 | 0.000611 | FOXQ1 |
| ENSG0000 | 6.316829 | 79.71773 | 0.000147 | 0.001149 | IL31RA |
| ENSG0000 | 3.773774 | 13.67789 | 0.002125 | 0.011737 | FBXL21P |
| ENSG0000 | 1.906361 | 3.748625 | 0.009065 | 0.039561 | CHMP4C |
| ENSG0000 | 1.185012 | 2.273652 | 1.62E-22 | 1.12E-20 | NFIL3 |
| ENSG0000 | 1.962976 | 3.898654 | 7.30E-12 | 1.97E-10 | ZMAT4 |
| ENSG0000 | 2.915478 | 7.544777 | 4.54E-07 | 6.07E-06 | RASEF |
| ENSG0000 | 1.092629 | 2.132623 | 5.78E-33 | 7.03E-31 | MID1IP1 |
| ENSG0000 | 6.603472 | 97.23963 | 0.000101 | 0.000821 | SHOC1 |
| ENSG0000 | 1.562184 | 2.953005 | 1.04E-32 | 1.25E-30 | PCDH19 |
| ENSG0000 | 5.193631 | 36.59644 | 2.80E-65 | 8.29E-63 | SLC7A3 |
| ENSG0000 | 1.021081 | 2.029438 | 2.95E-64 | 8.64E-62 | GHITM |
| ENSG0000 | 1.0177 | 2.024689 | 8.19E-15 | 3.00E-13 | CLMN |
| ENSG0000 | 1.291857 | 2.44843 | 0.006445 | 0.029908 | PTER |
| ENSG0000 | 3.851479 | 14.4348 | 0.004803 | 0.023314 | GLB1L3 |
| ENSG0000 | 2.256513 | 4.77835 | 4.69E-83 | 1.91E-80 | LARP6 |
| ENSG0000 | 2.088462 | 4.252943 | 0.004998 | 0.024116 | CLMP |
| ENSG0000 | 2.623603 | 6.162873 | 0.000233 | 0.001715 | GALR1 |
| ENSG0000 | 1.936419 | 3.827545 | 4.46E-97 | 2.40E-94 | SLFN5 |
| ENSG0000 | 4.59606 | 24.18532 | 0.005684 | 0.026882 | MYO1A |
| ENSG0000 | 1.756949 | 3.379826 | 2.57E-10 | 5.67E-09 | STX3 |
| ENSG0000 | 1.66406 | 3.16907 | 3.32E-73 | 1.15E-70 | MARS1 |

| | | | | | |
|---|---|---|---|---|---|
| ENSG0000 | 1.010009 | 2.013924 | 6.66E-18 | 3.18E-16 | FAM102A |
| ENSG0000 | 2.312237 | 4.966526 | 0.003547 | 0.018133 | ISLR2 |
| ENSG0000 | 1.947099 | 3.855984 | 3.77E-49 | 8.02E-47 | MYO5B |
| ENSG0000 | 2.749288 | 6.72385 | 4.02E-05 | 0.000359 | RHEBL1 |
| ENSG0000 | 3.159205 | 8.933374 | 4.78E-17 | 2.15E-15 | ANGPTL4 |
| ENSG0000 | 1.230044 | 2.345741 | 0.000984 | 0.006065 | TMEM88 |
| ENSG0000 | 1.471201 | 2.772526 | 3.11E-17 | 1.43E-15 | VWCE |
| ENSG0000 | 1.59165 | 3.013939 | 2.09E-29 | 2.09E-27 | RAB3IL1 |
| ENSG0000 | 1.039437 | 2.055425 | 1.67E-32 | 2.00E-30 | FTH1 |
| ENSG0000 | 2.009502 | 4.026431 | 6.39E-131 | 4.56E-128 | SLC3A2 |
| ENSG0000 | 2.174177 | 4.513283 | 2.85E-07 | 3.96E-06 | ENTPD3 |
| ENSG0000 | 1.562888 | 2.954446 | 1.55E-18 | 8.01E-17 | KCNJ4 |
| ENSG0000 | 7.123707 | 139.46 | 0.004677 | 0.022825 | FAM83B |
| ENSG0000 | 1.834467 | 3.566396 | 1.36E-42 | 2.42E-40 | RNF187 |
| ENSG0000 | 3.179225 | 9.058203 | 9.64E-158 | 8.77E-155 | DDIT4 |
| ENSG0000 | 1.01432 | 2.01995 | 1.89E-13 | 6.04E-12 | FAM110B |
| ENSG0000 | 2.880955 | 7.366375 | 1.50E-77 | 5.65E-75 | ATF5 |
| ENSG0000 | 3.315847 | 9.957939 | 2.21E-112 | 1.42E-109 | STK32A |
| ENSG0000 | 9.30765 | 633.6974 | 2.42E-14 | 8.45E-13 | P2RY12 |
| ENSG0000 | 1.746184 | 3.354701 | 0.011577 | 0.048549 | BNC1 |
| ENSG0000 | 5.136546 | 35.17664 | 0.007245 | 0.032849 | SLN |
| ENSG0000 | 1.12654 | 2.183345 | 4.76E-05 | 0.000419 | CEL |
| ENSG0000 | 4.718338 | 26.32457 | 0.008222 | 0.036537 | PLAC1 |
| ENSG0000 | 2.820506 | 7.064103 | 0.001053 | 0.006443 | FUT3 |
| ENSG0000 | 2.764615 | 6.795664 | 4.33E-06 | 4.80E-05 | NMRAL2P |
| ENSG0000 | 1.988854 | 3.969217 | 3.55E-32 | 4.11E-30 | SCG2 |
| ENSG0000 | 1.506136 | 2.840483 | 8.60E-23 | 6.12E-21 | FOXB1 |
| ENSG0000 | 2.048741 | 4.137446 | 1.49E-30 | 1.61E-28 | KLF11 |
| ENSG0000 | 2.093081 | 4.266582 | 2.47E-05 | 0.000232 | ISG20 |
| ENSG0000 | 2.67619 | 6.391656 | 1.59E-19 | 8.87E-18 | CEBPB |
| ENSG0000 | 1.397503 | 2.634452 | 9.73E-37 | 1.44E-34 | TP53RK |
| ENSG0000 | 2.114698 | 4.330993 | 3.09E-91 | 1.48E-88 | GTPBP2 |
| ENSG0000 | 7.162162 | 143.2272 | 2.40E-05 | 0.000226 | THEMIS |
| ENSG0000 | 1.031363 | 2.043954 | 9.79E-08 | 1.47E-06 | ZBTB21 |
| ENSG0000 | 1.708806 | 3.268901 | 3.58E-92 | 1.85E-89 | TNFRSF10D |
| ENSG0000 | 1.214004 | 2.319805 | 1.15E-13 | 3.73E-12 | TNFRSF10C |
| ENSG0000 | 1.969343 | 3.915896 | 3.01E-37 | 4.59E-35 | HAP1 |
| ENSG0000 | 1.448134 | 2.728549 | 4.60E-91 | 2.17E-88 | EIF1 |
| ENSG0000 | 2.408674 | 5.309862 | 0.008141 | 0.036236 | GPR160 |
| ENSG0000 | 5.466427 | 44.21387 | 0.010611 | 0.045238 | SELP |
| ENSG0000 | 2.959626 | 7.779222 | 1.23E-15 | 4.90E-14 | CHRNA9 |
| ENSG0000 | 2.903609 | 7.482959 | 0.000818 | 0.005155 | FUT1 |
| ENSG0000 | 2.021223 | 4.059278 | 3.06E-21 | 1.95E-19 | CADM2 |
| ENSG0000 | 3.607744 | 12.191 | 3.49E-193 | 4.60E-190 | DDIT3 |
| ENSG0000 | 1.229552 | 2.344942 | 1.05E-05 | 0.000107 | PCSK1 |
| ENSG0000 | 5.774038 | 54.72157 | 0.002237 | 0.012284 | LRRC25 |
| ENSG0000 | 2.561579 | 5.903536 | 5.11E-86 | 2.25E-83 | NUPR1 |
| ENSG0000 | 7.461618 | 176.2669 | 0.000652 | 0.004235 | CRYBG2 |
| ENSG0000 | 1.32783 | 2.510248 | 0.001594 | 0.009215 | SPHK1 |
| ENSG0000 | 5.477758 | 44.56248 | 0.007356 | 0.033229 | None |

| | | | | | |
|---|---|---|---|---|---|
| ENSG0000 | 1.983832 | 3.955424 | 1.57E-70 | 5.23E-68 | DIRAS1 |
| ENSG0000 | 1.379237 | 2.601308 | 3.29E-30 | 3.47E-28 | CDK5R1 |
| ENSG0000 | 1.357346 | 2.562133 | 1.71E-05 | 0.000168 | TCERG1L |
| ENSG0000 | 2.061335 | 4.173723 | 0.000205 | 0.00154 | TOB2P1 |
| ENSG0000 | 1.530707 | 2.889274 | 4.34E-31 | 4.80E-29 | FIBIN |
| ENSG0000 | 5.976288 | 62.95668 | 0.000387 | 0.002687 | TRIM72 |
| ENSG0000 | 2.153916 | 4.45034 | 3.30E-30 | 3.47E-28 | LRRN4CL |
| ENSG0000 | 1.291914 | 2.448527 | 1.49E-27 | 1.33E-25 | ZFAS1 |
| ENSG0000 | 1.481021 | 2.791462 | 1.47E-53 | 3.42E-51 | TGIF1 |
| ENSG0000 | 1.059367 | 2.084018 | 5.69E-18 | 2.76E-16 | ERN1 |
| ENSG0000 | 1.108805 | 2.156669 | 2.86E-07 | 3.97E-06 | RPP25 |
| ENSG0000 | 2.181863 | 4.53739 | 0.005054 | 0.024348 | ERFE |
| ENSG0000 | 4.419105 | 21.39357 | 0.004684 | 0.022849 | CPNE7 |
| ENSG0000 | 2.109228 | 4.314604 | 8.92E-35 | 1.18E-32 | OPLAH |
| ENSG0000 | 2.523535 | 5.749894 | 0.000222 | 0.001646 | MSC |
| ENSG0000 | 2.895194 | 7.43944 | 0.008628 | 0.037999 | C14orf39 |
| ENSG0000 | 6.798531 | 111.317 | 0.00275 | 0.014648 | TRIML2 |
| ENSG0000 | 2.051282 | 4.144741 | 0.007888 | 0.035294 | HLA-DQB1 |
| ENSG0000 | 1.125184 | 2.181293 | 0.000407 | 0.002803 | PLD6 |
| ENSG0000 | 1.491177 | 2.811182 | 2.62E-74 | 9.48E-72 | SEPHS2 |
| ENSG0000 | 4.727467 | 26.49168 | 0.002583 | 0.013915 | None |
| ENSG0000 | 2.789651 | 6.914624 | 8.19E-06 | 8.52E-05 | C3orf80 |
| ENSG0000 | 2.034006 | 4.095406 | 0.002466 | 0.01336 | ANKRD18A |
| ENSG0000 | 5.553551 | 46.96621 | 0.001486 | 0.008709 | ALX1 |
| ENSG0000 | 6.101563 | 68.66787 | 0.003554 | 0.018157 | TPRXL |
| ENSG0000 | 2.220202 | 4.659586 | 6.15E-08 | 9.52E-07 | GPR157 |
| ENSG0000 | 2.636516 | 6.21828 | 0.004531 | 0.022207 | MAP3K15 |
| ENSG0000 | 1.367923 | 2.580987 | 0.006703 | 0.030833 | LSMEM1 |
| ENSG0000 | 1.351886 | 2.552456 | 1.80E-19 | 1.00E-17 | AEN |
| ENSG0000 | 1.321499 | 2.499257 | 7.68E-12 | 2.06E-10 | H2AW |
| ENSG0000 | 1.529398 | 2.886654 | 0.003122 | 0.016297 | LINC00471 |
| ENSG0000 | 1.041487 | 2.058349 | 1.20E-07 | 1.77E-06 | FAM89A |
| ENSG0000 | 2.850941 | 7.214708 | 1.31E-199 | 2.04E-196 | SHMT2 |
| ENSG0000 | 4.103758 | 17.19311 | 0.001726 | 0.009852 | BACE2 |
| ENSG0000 | 7.221171 | 149.2069 | 0.003426 | 0.017606 | IZUMO1 |
| ENSG0000 | 2.389358 | 5.239242 | 2.14E-35 | 2.97E-33 | NXPH4 |
| ENSG0000 | 1.85995 | 3.629951 | 6.69E-26 | 5.52E-24 | BGN |
| ENSG0000 | 4.919038 | 30.25366 | 0.002448 | 0.013277 | KCNB2 |
| ENSG0000 | 1.378667 | 2.60028 | 0.004717 | 0.022971 | PAPPA |
| ENSG0000 | 2.293315 | 4.901811 | 1.80E-145 | 1.39E-142 | PYCR1 |
| ENSG0000 | 3.718674 | 13.16535 | 0.002897 | 0.015319 | NKX2-5 |
| ENSG0000 | 2.872828 | 7.324995 | 0.000509 | 0.00342 | CCBE1 |
| ENSG0000 | 1.060337 | 2.085419 | 2.26E-07 | 3.18E-06 | PSMG1 |
| ENSG0000 | 2.268084 | 4.81683 | 0.007919 | 0.035409 | CMKLR2 |
| ENSG0000 | 1.00892 | 2.012405 | 4.48E-22 | 3.05E-20 | UPP1 |
| ENSG0000 | 1.578873 | 2.987365 | 8.17E-10 | 1.67E-08 | KCTD16 |
| ENSG0000 | 6.805374 | 111.8463 | 0.007033 | 0.032073 | DOCK8-AS1 |
| ENSG0000 | 7.16194 | 143.2052 | 2.01E-06 | 2.37E-05 | TREML3P |
| ENSG0000 | 2.279733 | 4.855879 | 0.00842 | 0.03722 | CCSER1 |
| ENSG0000 | 1.47295 | 2.775889 | 0.007092 | 0.032275 | EFNA5 |

| | | | | | |
|---|---|---|---|---|---|
| ENSG0000 | 2.597622 | 6.052883 | 7.59E-254 | 4.01E-250 | XPOT |
| ENSG0000 | 1.132957 | 2.193078 | 3.30E-06 | 3.75E-05 | JAG2 |
| ENSG0000 | 5.304499 | 39.51967 | 2.41E-05 | 0.000227 | IFNLR1 |
| ENSG0000 | 1.205254 | 2.305778 | 1.54E-28 | 1.45E-26 | SH3BGR |
| ENSG0000 | 2.831694 | 7.119094 | 3.39E-05 | 0.000308 | GPRIN3 |
| ENSG0000 | 1.714695 | 3.282273 | 8.44E-42 | 1.46E-39 | TLCD2 |
| ENSG0000 | 4.454039 | 21.91792 | 7.43E-35 | 1.00E-32 | NDUFA4L2 |
| ENSG0000 | 1.425577 | 2.686218 | 0.000553 | 0.003677 | ADARB2 |
| ENSG0000 | 3.848707 | 14.40709 | 0.010435 | 0.04463 | ZNF829 |
| ENSG0000 | 4.732069 | 26.57631 | 0.007709 | 0.034592 | DTX2P1 |
| ENSG0000 | 5.740348 | 53.45851 | 0.001335 | 0.007955 | ANGPTL5 |
| ENSG0000 | 2.18623 | 4.551147 | 0.001556 | 0.009035 | ZNF385C |
| ENSG0000 | 1.617401 | 3.068218 | 0.003161 | 0.016466 | KRT18P59 |
| ENSG0000 | 1.658924 | 3.15781 | 5.80E-96 | 3.06E-93 | TCEA1 |
| ENSG0000 | 8.504265 | 363.1105 | 0.00309 | 0.016156 | HELT |
| ENSG0000 | 3.123041 | 8.712226 | 4.58E-151 | 3.78E-148 | EIF4EBP1 |
| ENSG0000 | 2.036992 | 4.103891 | 0.009356 | 0.040648 | C2orf66 |
| ENSG0000 | 1.238065 | 2.35882 | 0.001801 | 0.010205 | WNT7B |
| ENSG0000 | 6.00493 | 64.21908 | 0.004522 | 0.022174 | OTOG |
| ENSG0000 | 1.220359 | 2.330047 | 1.21E-11 | 3.17E-10 | IER5L |
| ENSG0000 | 2.487233 | 5.607014 | 0.01057 | 0.045095 | INSC |
| ENSG0000 | 1.7159 | 3.285015 | 2.32E-11 | 5.85E-10 | FAM83G |
| ENSG0000 | 2.936304 | 7.654478 | 5.47E-50 | 1.19E-47 | FBLL1 |
| ENSG0000 | 1.388905 | 2.618798 | 4.31E-29 | 4.20E-27 | NANOS1 |
| ENSG0000 | 5.507503 | 45.4908 | 0.005694 | 0.026912 | HMX3 |
| ENSG0000 | 1.83153 | 3.559144 | 0.000526 | 0.003514 | SLC4A5 |
| ENSG0000 | 1.930799 | 3.812664 | 0.001812 | 0.010262 | ZBED6CL |
| ENSG0000 | 1.08952 | 2.128033 | 9.34E-21 | 5.68E-19 | H1-0 |
| ENSG0000 | 1.149398 | 2.218214 | 0.000122 | 0.000968 | RPS2P46 |
| ENSG0000 | 2.124305 | 4.359931 | 6.44E-246 | 2.43E-242 | IARS1 |
| ENSG0000 | 1.141057 | 2.205425 | 6.17E-07 | 8.09E-06 | CD55 |
| ENSG0000 | 1.581716 | 2.993257 | 6.03E-66 | 1.85E-63 | LONP1 |
| ENSG0000 | 1.112269 | 2.161853 | 4.11E-10 | 8.83E-09 | SULT1A1 |
| ENSG0000 | 1.230276 | 2.346118 | 2.27E-25 | 1.82E-23 | SLC6A9 |
| ENSG0000 | 1.931197 | 3.813715 | 0.00322 | 0.016711 | HLA-DQA1 |
| ENSG0000 | 1.068752 | 2.097618 | 0.005912 | 0.027754 | SNHG17 |
| ENSG0000 | 1.652475 | 3.143724 | 0.000178 | 0.001362 | H2BU1 |
| ENSG0000 | 1.753028 | 3.370652 | 0.001566 | 0.009079 | CASP4 |
| ENSG0000 | 1.334984 | 2.522726 | 1.31E-51 | 2.92E-49 | MAFG |
| ENSG0000 | 1.728684 | 3.314253 | 7.94E-11 | 1.88E-09 | BLM |
| ENSG0000 | 6.529486 | 92.37855 | 0.00921 | 0.040119 | GATA3-AS1 |
| ENSG0000 | 1.860495 | 3.631323 | 2.15E-91 | 1.05E-88 | UAP1L1 |
| ENSG0000 | 1.406766 | 2.651422 | 3.83E-35 | 5.21E-33 | SLC2A10 |
| ENSG0000 | 3.029177 | 8.163439 | 0.01012 | 0.043455 | GPX1P1 |
| ENSG0000 | 3.686349 | 12.87364 | 0.008497 | 0.037509 | KLHL14 |
| ENSG0000 | 1.180322 | 2.266274 | 3.42E-35 | 4.67E-33 | SGTB |
| ENSG0000 | 1.874342 | 3.666344 | 1.81E-05 | 0.000176 | PLCG2 |
| ENSG0000 | 5.749661 | 53.80472 | 0.001108 | 0.006741 | SULT1C2 |
| ENSG0000 | 1.491707 | 2.812216 | 0.005028 | 0.024247 | PIM3 |
| ENSG0000 | 1.294489 | 2.452902 | 1.15E-70 | 3.88E-68 | TXNRD1 |

| | | | | | |
|---|---|---|---|---|---|
| ENSG0000 | 1.308808 | 2.477368 | 1.04E-26 | 8.85E-25 | NBR2 |
| ENSG0000 | 4.38371 | 20.87508 | 1.29E-15 | 5.11E-14 | EGFL6 |
| ENSG0000 | 1.238063 | 2.358817 | 3.98E-05 | 0.000356 | BRINP2 |
| ENSG0000 | 6.316856 | 79.71925 | 0.00015 | 0.001172 | PNMA5 |
| ENSG0000 | 5.154441 | 35.6157 | 0.011881 | 0.049563 | U3 |
| ENSG0000 | 4.748697 | 26.88439 | 0.005266 | 0.025201 | IGBP1-AS1 |
| ENSG0000 | 6.155091 | 71.26349 | 0.001514 | 0.008835 | TCEAL6 |
| ENSG0000 | 1.069799 | 2.099141 | 7.43E-31 | 8.14E-29 | INPP5B |
| ENSG0000 | 6.000725 | 64.03217 | 0.001107 | 0.006738 | RPA4 |
| ENSG0000 | 3.366236 | 10.31188 | 1.07E-21 | 7.11E-20 | ERICH2 |
| ENSG0000 | 3.467554 | 11.0621 | 4.34E-07 | 5.84E-06 | CARD16 |
| ENSG0000 | 1.615977 | 3.065191 | 0.005045 | 0.024321 | FAM201A |
| ENSG0000 | 2.034261 | 4.096128 | 2.29E-05 | 0.000217 | None |
| ENSG0000 | 2.602869 | 6.074937 | 0.001855 | 0.010461 | PKHD1L1 |
| ENSG0000 | 6.380821 | 83.33327 | 0.006918 | 0.031617 | KRT17P5 |
| ENSG0000 | 1.352423 | 2.553406 | 0.000698 | 0.004491 | EIF3CL |
| ENSG0000 | 6.673665 | 102.0877 | 0.00934 | 0.040586 | |
| ENSG0000 | 1.30626 | 2.472996 | 0.011394 | 0.047965 | C16orf96 |
| ENSG0000 | 9.124484 | 558.1401 | 0.000222 | 0.001646 | ONECUT3 |
| ENSG0000 | 2.580207 | 5.980253 | 0.000336 | 0.00237 | OLIG2 |
| ENSG0000 | 2.815323 | 7.038767 | 0.011397 | 0.047969 | HNRNPA1P4 |
| ENSG0000 | 6.373131 | 82.89025 | 0.002882 | 0.015258 | Y_RNA |
| ENSG0000 | 2.307321 | 4.949631 | 0.000555 | 0.003686 | MIR616 |
| ENSG0000 | 1.44361 | 2.720006 | 1.22E-12 | 3.60E-11 | MT-TP |
| ENSG0000 | 1.005793 | 2.008047 | 3.76E-07 | 5.12E-06 | GPX3 |
| ENSG0000 | 5.520332 | 45.89714 | 0.010199 | 0.043759 | IGLV1-50 |
| ENSG0000 | 6.008622 | 64.38364 | 0.00162 | 0.009349 | CFL1P4 |
| ENSG0000 | 5.496374 | 45.14125 | 0.002708 | 0.014471 | None |
| ENSG0000 | 7.266538 | 153.9735 | 0.000171 | 0.001314 | YBX1P4 |
| ENSG0000 | 5.732307 | 53.16141 | 0.005522 | 0.026238 | NOC2LP1 |
| ENSG0000 | 2.964151 | 7.803663 | 0.011957 | 0.049827 | EEF1A1P16 |
| ENSG0000 | 1.936479 | 3.827703 | 1.79E-22 | 1.25E-20 | HAUS7 |
| ENSG0000 | 1.518577 | 2.865082 | 0.007422 | 0.033485 | RPLP0P6 |
| ENSG0000 | 6.794261 | 110.9881 | 0.002985 | 0.015702 | KRT18P17 |
| ENSG0000 | 3.264874 | 9.612252 | 0.002865 | 0.015191 | XPOTP1 |
| ENSG0000 | 2.290586 | 4.892549 | 0.00397 | 0.0199 | EEF1A1P12 |
| ENSG0000 | 1.361674 | 2.569832 | 1.35E-07 | 1.98E-06 | LCNL1 |
| ENSG0000 | 1.198657 | 2.29526 | 0.00037 | 0.002578 | PLIN5 |
| ENSG0000 | 1.070075 | 2.099542 | 8.40E-16 | 3.39E-14 | CPEB1 |
| ENSG0000 | 2.971658 | 7.84437 | 0.006669 | 0.030742 | None |
| ENSG0000 | 3.007415 | 8.041222 | 0.008875 | 0.038898 | LINC01588 |
| ENSG0000 | 5.778177 | 54.87881 | 0.008413 | 0.037194 | CHCHD3P3 |
| ENSG0000 | 6.520482 | 91.80381 | 0.001733 | 0.009883 | HSPD1P7 |
| ENSG0000 | 6.971174 | 125.4679 | 4.72E-06 | 5.17E-05 | EEF1A1P29 |
| ENSG0000 | 6.919725 | 121.0723 | 0.002227 | 0.01224 | TOMM40P2 |
| ENSG0000 | 6.157325 | 71.37392 | 0.003254 | 0.016853 | None |
| ENSG0000 | 1.863225 | 3.6382 | 0.006439 | 0.029891 | RPS2P55 |
| ENSG0000 | 2.293224 | 4.901503 | 0.007423 | 0.033485 | SP9 |
| ENSG0000 | 1.629397 | 3.093836 | 0.006861 | 0.031403 | None |
| ENSG0000 | 1.190543 | 2.282386 | 0.005862 | 0.02756 | FTH1P8 |

| | | | | | |
|---|---|---|---|---|---|
| ENSG0000 | 7.371395 | 165.5812 | 7.82E-05 | 0.000652 | RPSAP44 |
| ENSG0000 | 1.18798 | 2.278335 | 0.005248 | 0.025125 | C19orf73 |
| ENSG0000 | 1.041755 | 2.058731 | 7.36E-16 | 3.01E-14 | TRIM16 |
| ENSG0000 | 1.185782 | 2.274867 | 3.92E-17 | 1.79E-15 | APOL6 |
| ENSG0000 | 1.060072 | 2.085035 | 0.000487 | 0.003289 | None |
| ENSG0000 | 5.758098 | 54.1203 | 0.006521 | 0.030187 | MTND6P21 |
| ENSG0000 | 1.113131 | 2.163145 | 2.74E-32 | 3.22E-30 | EXOSC6 |
| ENSG0000 | 1.045887 | 2.064636 | 0.005868 | 0.027577 | LINC00571 |
| ENSG0000 | 5.45122 | 43.75027 | 0.005384 | 0.02569 | None |
| ENSG0000 | 1.029122 | 2.040781 | 0.000296 | 0.00212 | MIR503HG |
| ENSG0000 | 5.308348 | 39.62524 | 7.07E-06 | 7.46E-05 | None |
| ENSG0000 | 5.95804 | 62.16542 | 0.003965 | 0.019879 | MKRN5P |
| ENSG0000 | 1.767221 | 3.403976 | 0.000113 | 0.00091 | None |
| ENSG0000 | 1.839563 | 3.579015 | 0.000378 | 0.002629 | MIR3659HG |
| ENSG0000 | 2.281796 | 4.862828 | 6.28E-11 | 1.50E-09 | HLCS-AS1 |
| ENSG0000 | 6.357724 | 82.00977 | 0.002027 | 0.011273 | RORB-AS1 |
| ENSG0000 | 8.328109 | 321.3739 | 2.24E-10 | 4.98E-09 | LINC00237 |
| ENSG0000 | 1.115434 | 2.166602 | 1.15E-07 | 1.70E-06 | None |
| ENSG0000 | 8.022247 | 259.9783 | 0.000124 | 0.000988 | DUXAP9 |
| ENSG0000 | 1.425574 | 2.686213 | 0.009039 | 0.039474 | LINC01703 |
| ENSG0000 | 6.209152 | 73.98456 | 0.008381 | 0.037083 | PA2G4P5 |
| ENSG0000 | 6.207919 | 73.92135 | 0.005003 | 0.024138 | None |
| ENSG0000 | 6.897439 | 119.2164 | 0.000114 | 0.000919 | RPL36AP54 |
| ENSG0000 | 1.276965 | 2.423286 | 4.20E-10 | 9.01E-09 | SLC16A1-AS1 |
| ENSG0000 | 1.564724 | 2.958209 | 0.003911 | 0.019664 | LINC01748 |
| ENSG0000 | 2.877797 | 7.350271 | 0.006696 | 0.03081 | LINC02549 |
| ENSG0000 | 6.331264 | 80.51938 | 0.000897 | 0.005602 | TRAF6P1 |
| ENSG0000 | 7.800149 | 222.884 | 6.32E-05 | 0.00054 | CYP4F26P |
| ENSG0000 | 1.596915 | 3.024957 | 0.008343 | 0.036964 | FTLP3 |
| ENSG0000 | 6.863273 | 116.4263 | 4.05E-06 | 4.53E-05 | None |
| ENSG0000 | 4.8794 | 29.43376 | 1.81E-05 | 0.000176 | |
| ENSG0000 | 3.848459 | 14.40461 | 3.52E-05 | 0.000319 | None |
| ENSG0000 | 1.120423 | 2.174108 | 0.000551 | 0.003666 | ATE1-AS1 |
| ENSG0000 | 6.185113 | 72.76199 | 0.002907 | 0.015359 | LHFPL3-AS1 |
| ENSG0000 | 1.548852 | 2.925842 | 0.005531 | 0.026269 | BSN-DT |
| ENSG0000 | 6.36629 | 82.49815 | 0.002874 | 0.015231 | NCKAP5-AS2 |
| ENSG0000 | 4.996761 | 31.92824 | 0.000313 | 0.002231 | LINC00629 |
| ENSG0000 | 1.961879 | 3.89569 | 4.28E-09 | 7.92E-08 | None |
| ENSG0000 | 5.155973 | 35.65354 | 0.004357 | 0.02151 | DLG1-AS1 |
| ENSG0000 | 1.30686 | 2.474024 | 0.001787 | 0.010142 | ADGRF5P1 |
| ENSG0000 | 5.963658 | 62.40794 | 0.000404 | 0.00279 | HSD11B1-AS1 |
| ENSG0000 | 6.746975 | 107.4093 | 0.000203 | 0.001528 | None |
| ENSG0000 | 6.539258 | 93.00637 | 0.002961 | 0.015596 | TEX46 |
| ENSG0000 | 1.287435 | 2.440936 | 0.011651 | 0.048842 | SEC63P1 |
| ENSG0000 | 5.740383 | 53.45982 | 0.00034 | 0.002392 | None |
| ENSG0000 | 6.525858 | 92.14655 | 0.00206 | 0.011441 | NACA4P |
| ENSG0000 | 1.193874 | 2.287662 | 0.00023 | 0.001698 | RRAGC-DT |
| ENSG0000 | 1.334122 | 2.521219 | 1.01E-38 | 1.60E-36 | MIR34AHG |
| ENSG0000 | 1.073226 | 2.104133 | 0.007255 | 0.032856 | SNHG26 |
| ENSG0000 | 1.077575 | 2.110486 | 0.009586 | 0.041526 | UBAC2-AS1 |

| | | | | | |
|---|---|---|---|---|---|
| ENSG0000 | 6.6363 | 99.47761 | 0.0002 | 0.001507 | SLC47A1P1 |
| ENSG0000 | 5.136937 | 35.18617 | 0.006406 | 0.029785 | LINC00452 |
| ENSG0000 | 7.540307 | 186.148 | 0.001847 | 0.010423 | SFTA3 |
| ENSG0000 | 4.214443 | 18.56409 | 0.007044 | 0.032098 | LARP4B-DT |
| ENSG0000 | 5.767177 | 54.46196 | 0.005344 | 0.025515 | None |
| ENSG0000 | 6.745495 | 107.2992 | 2.49E-05 | 0.000234 | CT70 |
| ENSG0000 | 6.005739 | 64.2551 | 0.000775 | 0.004916 | HNRNPA1P28 |
| ENSG0000 | 3.912193 | 15.05523 | 6.93E-06 | 7.33E-05 | CNOT6LP1 |
| ENSG0000 | 5.731783 | 53.14209 | 0.010042 | 0.043169 | None |
| ENSG0000 | 1.430186 | 2.694814 | 2.05E-11 | 5.20E-10 | TCEA1P2 |
| ENSG0000 | 3.719082 | 13.16907 | 0.0039 | 0.019623 | PPIAP40 |
| ENSG0000 | 1.440769 | 2.714654 | 1.51E-07 | 2.20E-06 | None |
| ENSG0000 | 6.330843 | 80.4959 | 0.000221 | 0.00164 | KIF5C-AS1 |
| ENSG0000 | 2.042843 | 4.120569 | 0.00715 | 0.032487 | YBX1P2 |
| ENSG0000 | 5.797311 | 55.6115 | 0.007953 | 0.035524 | VDAC1P6 |
| ENSG0000 | 1.031639 | 2.044346 | 0.000475 | 0.003213 | LINC01354 |
| ENSG0000 | 1.364487 | 2.574847 | 0.009646 | 0.04175 | EP300-AS1 |
| ENSG0000 | 6.36427 | 82.3827 | 0.001733 | 0.009883 | PRKD3-DT |
| ENSG0000 | 6.901683 | 119.5676 | 0.003038 | 0.015941 | LINC01873 |
| ENSG0000 | 1.620897 | 3.075663 | 0.002771 | 0.01475 | FTH1P7 |
| ENSG0000 | 7.078947 | 135.1996 | 1.63E-05 | 0.00016 | RPS27AP2 |
| ENSG0000 | 1.372326 | 2.588877 | 0.006877 | 0.031463 | DLGAP4-AS1 |
| ENSG0000 | 1.497784 | 2.824087 | 0.003204 | 0.016638 | SNHG15 |
| ENSG0000 | 6.326755 | 80.26812 | 0.000191 | 0.001446 | HDAC1P2 |
| ENSG0000 | 5.762138 | 54.27208 | 0.011552 | 0.048498 | LNCARSR |
| ENSG0000 | 4.198789 | 18.36375 | 0.003736 | 0.018903 | CNIH3-AS2 |
| ENSG0000 | 6.527691 | 92.26369 | 0.006252 | 0.029183 | None |
| ENSG0000 | 1.766105 | 3.401344 | 2.98E-12 | 8.41E-11 | None |
| ENSG0000 | 1.964726 | 3.903387 | 1.60E-12 | 4.65E-11 | LNCTAM34A |
| ENSG0000 | 1.275638 | 2.421059 | 1.87E-05 | 0.000181 | NPIPB2 |
| ENSG0000 | 1.742986 | 3.347272 | 3.29E-64 | 9.54E-62 | GAS5 |
| ENSG0000 | 1.316668 | 2.490902 | 0.006978 | 0.031855 | ECI2-DT |
| ENSG0000 | 3.853922 | 14.45926 | 0.011729 | 0.049078 | AARSD1P1 |
| ENSG0000 | 1.268474 | 2.409067 | 0.002926 | 0.015442 | FTH1P2 |
| ENSG0000 | 1.128979 | 2.187039 | 0.000249 | 0.001819 | None |
| ENSG0000 | 2.3997 | 5.276932 | 0.000664 | 0.0043 | None |
| ENSG0000 | 6.188112 | 72.91341 | 0.000571 | 0.003775 | TATDN2P3 |
| ENSG0000 | 3.262118 | 9.593902 | 3.92E-28 | 3.56E-26 | L3MBTL2-AS1 |
| ENSG0000 | 2.157799 | 4.462336 | 1.41E-05 | 0.000141 | MSC-AS1 |
| ENSG0000 | 6.207553 | 73.90257 | 0.004066 | 0.020279 | None |
| ENSG0000 | 1.021815 | 2.030471 | 1.92E-06 | 2.28E-05 | PNMA6A |
| ENSG0000 | 2.136678 | 4.397484 | 2.52E-29 | 2.50E-27 | VLDLR-AS1 |
| ENSG0000 | 5.725045 | 52.89448 | 0.003164 | 0.01648 | RPL23AP24 |
| ENSG0000 | 1.0408 | 2.057368 | 0.002711 | 0.01448 | None |
| ENSG0000 | 1.617141 | 3.067665 | 0.003131 | 0.016329 | FTH1P11 |
| ENSG0000 | 2.485056 | 5.598563 | 0.009884 | 0.042566 | SCIRT |
| ENSG0000 | 1.745329 | 3.352713 | 0.01029 | 0.044096 | None |
| ENSG0000 | 4.196677 | 18.33689 | 0.001703 | 0.009745 | IRS4-AS1 |
| ENSG0000 | 3.46142 | 11.01517 | 0.005563 | 0.026393 | CDC26P1 |
| ENSG0000 | 6.499504 | 90.47854 | 0.000461 | 0.003128 | None |

| | | | | | |
|---|---|---|---|---|---|
| ENSG0000 | 6.13305 | 70.18301 | 0.000357 | 0.002502 | SNORA84 |
| ENSG0000 | 3.446724 | 10.90353 | 4.91E-70 | 1.60E-67 | None |
| ENSG0000 | 5.612585 | 48.92787 | 0.000153 | 0.001187 | STRIT1 |
| ENSG0000 | 1.0697 | 2.098997 | 0.000243 | 0.001783 | RPS2P5 |
| ENSG0000 | 3.920337 | 15.14046 | 0.005725 | 0.027031 | ARHGAP8 |
| ENSG0000 | 4.238484 | 18.87604 | 0.005818 | 0.027398 | None |
| ENSG0000 | 1.698033 | 3.244583 | 9.76E-05 | 0.000795 | HLA-DMB |
| ENSG0000 | 5.951547 | 61.88625 | 0.000548 | 0.003651 | None |
| ENSG0000 | 1.910099 | 3.758349 | 5.81E-07 | 7.64E-06 | EIF4EBP3 |
| ENSG0000 | 1.830444 | 3.556464 | 0.004007 | 0.020045 | TWF2-DT |
| ENSG0000 | 5.981886 | 63.20147 | 0.010183 | 0.043708 | MIR1302-2HG |
| ENSG0000 | 2.356458 | 5.121113 | 1.50E-15 | 5.90E-14 | CEBPA |
| ENSG0000 | 1.062568 | 2.088646 | 1.27E-05 | 0.000127 | SBF2-AS1 |
| ENSG0000 | 4.461029 | 22.02438 | 0.006262 | 0.029223 | ZBED9-AS1 |
| ENSG0000 | 4.040591 | 16.45656 | 0.000197 | 0.001489 | TNFRSF10A-DT |
| ENSG0000 | 1.020352 | 2.028413 | 2.54E-06 | 2.95E-05 | RGMB-AS1 |
| ENSG0000 | 1.687835 | 3.221728 | 0.003305 | 0.017083 | MEF2C-AS1 |
| ENSG0000 | 4.056792 | 16.6424 | 0.008707 | 0.038276 | LINC02485 |
| ENSG0000 | 7.72203 | 211.1362 | 3.26E-05 | 0.000297 | LINC01033 |
| ENSG0000 | 1.530711 | 2.889282 | 0.005103 | 0.024524 | EEF1A1P9 |
| ENSG0000 | 4.306318 | 19.78476 | 0.006584 | 0.030439 | HOXC13-AS |
| ENSG0000 | 5.794289 | 55.49511 | 0.003974 | 0.019912 | TNRC18P1 |
| ENSG0000 | 1.451914 | 2.735707 | 8.64E-10 | 1.76E-08 | PVT1 |
| ENSG0000 | 6.461968 | 88.15482 | 0.00032 | 0.002271 | TERB1 |
| ENSG0000 | 1.010414 | 2.01449 | 3.74E-05 | 0.000336 | PURPL |
| ENSG0000 | 1.968001 | 3.912256 | 0.008644 | 0.038049 | STK32A-AS1 |
| ENSG0000 | 5.48939 | 44.92322 | 0.005178 | 0.024827 | RNF138P1 |
| ENSG0000 | 2.454757 | 5.48221 | 0.000784 | 0.00497 | ZFPM2-AS1 |
| ENSG0000 | 5.941333 | 61.44964 | 0.003281 | 0.016975 | LTO1P1 |
| ENSG0000 | 6.611475 | 97.78054 | 0.000153 | 0.00119 | None |
| ENSG0000 | 1.821732 | 3.535054 | 1.92E-14 | 6.77E-13 | LINC01094 |
| ENSG0000 | 1.44596 | 2.724441 | 2.44E-08 | 4.05E-07 | FOXD1 |
| ENSG0000 | 2.449349 | 5.461696 | 0.002899 | 0.015325 | None |
| ENSG0000 | 5.508562 | 45.52421 | 0.005341 | 0.02551 | MIR2116 |
| ENSG0000 | 5.981018 | 63.16346 | 0.005238 | 0.025081 | None |
| ENSG0000 | 5.470174 | 44.32885 | 0.006197 | 0.028957 | None |
| ENSG0000 | 6.679191 | 102.4794 | 0.006828 | 0.031284 | None |
| ENSG0000 | 5.749323 | 53.79211 | 0.000835 | 0.005259 | None |
| ENSG0000 | 1.845911 | 3.5948 | 1.56E-05 | 0.000154 | None |
| ENSG0000 | 1.262497 | 2.399106 | 0.007643 | 0.034357 | LYN |
| ENSG0000 | 6.00664 | 64.29523 | 0.008895 | 0.038967 | None |
| ENSG0000 | 7.02221 | 129.9857 | 0.001573 | 0.009112 | LINC02365 |
| ENSG0000 | 1.03178 | 2.044545 | 0.003291 | 0.017016 | BBOX1-AS1 |
| ENSG0000 | 1.215044 | 2.321479 | 0.011499 | 0.04833 | CSNK2A3 |
| ENSG0000 | 1.488573 | 2.806113 | 1.11E-08 | 1.94E-07 | |
| ENSG0000 | 1.995099 | 3.986434 | 3.25E-05 | 0.000296 | PKNOX2-DT |
| ENSG0000 | 7.468386 | 177.0958 | 0.000549 | 0.003654 | COX5BP4 |
| ENSG0000 | 6.836969 | 114.3228 | 3.67E-05 | 0.000331 | None |
| ENSG0000 | 1.064086 | 2.090844 | 1.98E-08 | 3.34E-07 | SNHG1 |
| ENSG0000 | 1.831884 | 3.560016 | 3.21E-05 | 0.000292 | PRECSIT |

| | | | | | |
|---|---|---|---|---|---|
| ENSG0000 | 5.94688 | 61.68636 | 0.00414 | 0.020597 | None |
| ENSG0000 | 1.519456 | 2.866829 | 0.002632 | 0.014142 | None |
| ENSG0000 | 7.332859 | 161.2169 | 1.12E-06 | 1.40E-05 | MADD-AS1 |
| ENSG0000 | 1.731309 | 3.32029 | 0.008395 | 0.037132 | LBX2-AS1 |
| ENSG0000 | 6.746608 | 107.382 | 0.0008 | 0.005058 | LINC02156 |
| ENSG0000 | 3.233328 | 9.404351 | 0.010967 | 0.046454 | None |
| ENSG0000 | 6.198737 | 73.45235 | 0.00159 | 0.009196 | CHMP4BP1 |
| ENSG0000 | 2.362964 | 5.144262 | 0.001221 | 0.007347 | None |
| ENSG0000 | 5.659402 | 50.5417 | 0.000747 | 0.004767 | None |
| ENSG0000 | 6.014527 | 64.64769 | 0.010522 | 0.044924 | None |
| ENSG0000 | 5.774708 | 54.74699 | 0.00195 | 0.010924 | None |
| ENSG0000 | 4.741193 | 26.74492 | 0.009095 | 0.039664 | None |
| ENSG0000 | 1.438456 | 2.710307 | 3.49E-12 | 9.76E-11 | ITGB3 |
| ENSG0000 | 5.168049 | 35.95323 | 0.009853 | 0.042482 | None |
| ENSG0000 | 1.879081 | 3.678407 | 0.002374 | 0.012922 | CPEB1-AS1 |
| ENSG0000 | 6.883177 | 118.0437 | 4.42E-05 | 0.000392 | None |
| ENSG0000 | 8.174331 | 288.881 | 0.00018 | 0.001372 | PCSK6-AS1 |
| ENSG0000 | 7.439302 | 173.5613 | 1.30E-05 | 0.000131 | SLC22A31 |
| ENSG0000 | 1.596921 | 3.024971 | 1.93E-05 | 0.000186 | None |
| ENSG0000 | 6.750431 | 107.6669 | 0.000225 | 0.001664 | None |
| ENSG0000 | 1.250141 | 2.378646 | 0.005629 | 0.026664 | None |
| ENSG0000 | 7.8691 | 233.795 | 1.89E-08 | 3.19E-07 | None |
| ENSG0000 | 1.099933 | 2.143447 | 0.004765 | 0.02316 | TBC1D22A-DT |
| ENSG0000 | 1.463188 | 2.757169 | 0.002486 | 0.013454 | SLC7A5P1 |
| ENSG0000 | 5.758945 | 54.15209 | 0.002092 | 0.011587 | None |
| ENSG0000 | 1.611536 | 3.055771 | 0.011712 | 0.049028 | PAPOLA-DT |
| ENSG0000 | 6.671455 | 101.9314 | 0.002322 | 0.01268 | None |
| ENSG0000 | 2.418519 | 5.346219 | 0.001695 | 0.009715 | LINC01229 |
| ENSG0000 | 1.55039 | 2.928962 | 3.26E-71 | 1.12E-68 | CCPG1 |
| ENSG0000 | 3.117788 | 8.680557 | 3.64E-06 | 4.10E-05 | None |
| ENSG0000 | 6.157315 | 71.37342 | 0.000914 | 0.005687 | None |
| ENSG0000 | 1.329827 | 2.513725 | 0.00191 | 0.01073 | ZNNT1 |
| ENSG0000 | 6.003593 | 64.15958 | 0.002761 | 0.014706 | LMO7-AS1 |
| ENSG0000 | 2.097808 | 4.280586 | 1.00E-06 | 1.26E-05 | None |
| ENSG0000 | 5.463824 | 44.13416 | 0.005798 | 0.027332 | None |
| ENSG0000 | 1.455136 | 2.741824 | 0.006455 | 0.029944 | ST3GAL1-DT |
| ENSG0000 | 7.713907 | 209.9507 | 2.91E-05 | 0.000268 | ANKRD26P1 |
| ENSG0000 | 1.889316 | 3.704597 | 4.05E-06 | 4.53E-05 | None |
| ENSG0000 | 1.528473 | 2.884803 | 0.004444 | 0.021854 | LINC01686 |
| ENSG0000 | 2.397324 | 5.268251 | 1.00E-58 | 2.64E-56 | SMG1P7 |
| ENSG0000 | 6.473517 | 88.8634 | 0.000683 | 0.004405 | HERC2P11 |
| ENSG0000 | 6.914748 | 120.6553 | 0.002017 | 0.011239 | FOXF2-DT |
| ENSG0000 | 6.347937 | 81.45534 | 0.003625 | 0.018428 | STAG1-DT |
| ENSG0000 | 5.506449 | 45.45759 | 0.00437 | 0.021553 | TUBB8P7 |
| ENSG0000 | 1.912484 | 3.764568 | 2.18E-27 | 1.92E-25 | DLGAP1-AS2 |
| ENSG0000 | 6.172085 | 72.10788 | 0.001928 | 0.010823 | None |
| ENSG0000 | 2.352141 | 5.105815 | 0.008095 | 0.03606 | None |
| ENSG0000 | 1.320043 | 2.496735 | 0.000526 | 0.003514 | ROCK1P1 |
| ENSG0000 | 6.215539 | 74.31281 | 0.004716 | 0.022971 | None |
| ENSG0000 | 1.255514 | 2.387523 | 0.011211 | 0.047271 | None |

| | | | | | |
|---|---|---|---|---|---|
| ENSG0000 | 5.44254 | 43.48783 | 0.003915 | 0.019682 | None |
| ENSG0000 | 1.942436 | 3.84354 | 9.42E-10 | 1.91E-08 | MAFG-DT |
| ENSG0000 | 2.85309 | 7.225464 | 2.38E-13 | 7.53E-12 | ZNF516-AS1 |
| ENSG0000 | 6.358405 | 82.04847 | 0.003181 | 0.016535 | MIR3939 |
| ENSG0000 | 1.323954 | 2.503513 | 0.011574 | 0.048543 | None |
| ENSG0000 | 6.845377 | 114.991 | 0.000814 | 0.005134 | None |
| ENSG0000 | 4.128152 | 17.48628 | 0.0066 | 0.030505 | None |
| ENSG0000 | 2.545988 | 5.84008 | 0.007892 | 0.035305 | None |
| ENSG0000 | 1.578269 | 2.986113 | 1.42E-06 | 1.73E-05 | CEBPA-DT |
| ENSG0000 | 1.520624 | 2.869152 | 1.74E-05 | 0.000169 | None |
| ENSG0000 | 5.940895 | 61.431 | 0.004019 | 0.020099 | None |
| ENSG0000 | 7.917736 | 241.811 | 0.000411 | 0.002828 | FENDRR |
| ENSG0000 | 6.338973 | 80.95078 | 0.000922 | 0.005726 | VN1R80P |
| ENSG0000 | 1.257656 | 2.391069 | 2.52E-07 | 3.51E-06 | PGLS-DT |
| ENSG0000 | 4.732014 | 26.57529 | 0.001377 | 0.008153 | TDGF1P7 |
| ENSG0000 | 1.125111 | 2.181183 | 7.39E-14 | 2.46E-12 | SNHG8 |
| ENSG0000 | 4.464836 | 22.08256 | 0.006325 | 0.029475 | None |
| ENSG0000 | 1.478464 | 2.786519 | 0.000883 | 0.005522 | |
| ENSG0000 | 8.717886 | 421.0612 | 1.94E-10 | 4.34E-09 | |
| ENSG0000 | 1.783562 | 3.442752 | 1.22E-14 | 4.38E-13 | GOLGA4-AS1 |
| ENSG0000 | 5.770698 | 54.59505 | 0.000724 | 0.00464 | SNX2P1 |
| ENSG0000 | 1.085886 | 2.122679 | 6.86E-06 | 7.25E-05 | NAV2-AS6 |
| ENSG0000 | 1.38447 | 2.61076 | 5.58E-27 | 4.85E-25 | RASL10B |
| ENSG0000 | 1.743776 | 3.349105 | 4.43E-18 | 2.17E-16 | SRXN1 |
| ENSG0000 | 6.54148 | 93.14972 | 0.009592 | 0.041546 | None |
| ENSG0000 | 3.968404 | 15.65339 | 0.005464 | 0.025998 | None |
| ENSG0000 | 2.480261 | 5.579984 | 0.000542 | 0.003611 | TP53RK-DT |
| ENSG0000 | 1.182655 | 2.269941 | 0.000693 | 0.00446 | None |
| ENSG0000 | 2.471401 | 5.54582 | 0.003633 | 0.018465 | None |
| ENSG0000 | 2.974981 | 7.862463 | 0.006646 | 0.030668 | None |
| ENSG0000 | 5.463899 | 44.13646 | 0.001145 | 0.00694 | None |
| ENSG0000 | 6.515229 | 91.47016 | 0.003456 | 0.017728 | LINC02084 |
| ENSG0000 | 5.468791 | 44.28636 | 0.005435 | 0.025879 | None |
| ENSG0000 | 6.861332 | 116.2697 | 0.000251 | 0.001833 | None |
| ENSG0000 | 6.358258 | 82.04014 | 0.000763 | 0.004856 | None |
| ENSG0000 | 7.084499 | 135.7209 | 4.88E-06 | 5.33E-05 | None |
| ENSG0000 | 1.018167 | 2.025344 | 0.00325 | 0.016836 | |
| ENSG0000 | 7.80193 | 223.1593 | 3.11E-05 | 0.000285 | CYP4F33P |
| ENSG0000 | 1.575432 | 2.980246 | 0.000509 | 0.003418 | None |
| ENSG0000 | 1.8501 | 3.605252 | 0.00124 | 0.007449 | None |
| ENSG0000 | 5.146143 | 35.41143 | 0.007669 | 0.034449 | None |
| ENSG0000 | 4.003209 | 16.03562 | 0.003335 | 0.017207 | None |
| ENSG0000 | 1.434798 | 2.703443 | 0.000308 | 0.002203 | None |
| ENSG0000 | 1.491219 | 2.811264 | 1.74E-05 | 0.00017 | None |
| ENSG0000 | 1.985636 | 3.960373 | 9.75E-05 | 0.000795 | None |
| ENSG0000 | 1.131219 | 2.190437 | 0.002658 | 0.014249 | FAM27E3 |
| ENSG0000 | 3.776873 | 13.70731 | 1.99E-17 | 9.22E-16 | None |
| ENSG0000 | 3.245462 | 9.483781 | 0.000733 | 0.00469 | H2BC6 |
| ENSG0000 | 3.219677 | 9.315781 | 0.009246 | 0.040239 | LINC02334 |
| ENSG0000 | 6.014094 | 64.62829 | 0.009215 | 0.040136 | None |

| | | | | | |
|---|---|---|---|---|---|
| ENSG0000 | 5.941841 | 61.47131 | 0.00224 | 0.012293 | None |
| ENSG0000 | 5.135211 | 35.14411 | 0.006677 | 0.030767 | None |
| ENSG0000 | 7.604741 | 194.6504 | 0.000732 | 0.004687 | |
| ENSG0000 | 4.163623 | 17.92155 | 0.004881 | 0.023641 | None |
| ENSG0000 | 6.151065 | 71.0649 | 0.001389 | 0.008215 | None |
| ENSG0000 | 3.974659 | 15.72142 | 0.000297 | 0.00213 | None |
| ENSG0000 | 4.255162 | 19.09552 | 0.010185 | 0.043713 | None |
| ENSG0000 | 4.366877 | 20.63293 | 0.003673 | 0.01863 | None |
| ENSG0000 | 1.484749 | 2.798686 | 0.00708 | 0.032234 | None |
| ENSG0000 | 1.853779 | 3.614458 | 0.006832 | 0.031298 | None |
| ENSG0000 | 1.340829 | 2.532969 | 9.63E-05 | 0.000787 | CEBPB-AS1 |
| ENSG0000 | 2.69045 | 6.455149 | 0.00922 | 0.040136 | None |
| ENSG0000 | 1.254752 | 2.386261 | 0.000738 | 0.004713 | None |
| ENSG0000 | 6.626171 | 98.78164 | 0.000134 | 0.001057 | None |
| ENSG0000 | 7.221633 | 149.2548 | 0.005533 | 0.026277 | H4C2 |
| ENSG0000 | 2.165445 | 4.486048 | 0.00414 | 0.020597 | None |
| ENSG0000 | 7.689594 | 206.4422 | 6.78E-07 | 8.81E-06 | None |
| ENSG0000 | 3.418799 | 10.69451 | 0.001874 | 0.010551 | None |
| ENSG0000 | 6.158663 | 71.44012 | 0.006791 | 0.031152 | None |
| ENSG0000 | 5.986476 | 63.40284 | 0.003963 | 0.019875 | None |
| ENSG0000 | 6.673203 | 102.055 | 0.00644 | 0.029891 | |
| ENSG0000 | 1.677657 | 3.199079 | 0.009291 | 0.040398 | None |
| ENSG0000 | 5.468715 | 44.28405 | 0.005826 | 0.027415 | None |
| ENSG0000 | 1.246165 | 2.3721 | 0.010546 | 0.044999 | None |
| ENSG0000 | 6.154912 | 71.25463 | 0.006099 | 0.028573 | None |
| ENSG0000 | 1.011671 | 2.016246 | 0.004367 | 0.021544 | None |
| ENSG0000 | 7.859066 | 232.1745 | 0.011681 | 0.048947 | FOXCUT |
| ENSG0000 | 5.99464 | 63.76267 | 0.002038 | 0.01133 | HERC2P7 |
| ENSG0000 | 5.764868 | 54.37487 | 0.001959 | 0.010961 | RHOXF1P3 |
| ENSG0000 | 5.751242 | 53.86371 | 0.004538 | 0.022233 | None |
| ENSG0000 | 7.302053 | 157.8108 | 0.004893 | 0.023687 | FAM95C |
| ENSG0000 | 8.011799 | 258.1023 | 7.99E-06 | 8.34E-05 | None |

| GeneID | Treatme | Treatment | Treatment | Treatment | name | | Astrocytes_GCSi_FC-2_adj0.05_30 |
|---|---|---|---|---|---|---|---|
| ENSG000 | -1.167 | -2.24471 | 3.21E-05 | 0.000478 | PDK4 | | |
| ENSG000 | -1.245 | -2.36944 | 0.002302 | 0.015921 | DNAH9 | | |
| ENSG000 | -1.161 | -2.23571 | 1.78E-10 | 1.33E-08 | DLEC1 | | |
| ENSG000 | -1.291 | -2.4463 | 0.007281 | 0.038655 | PLEKHG6 | | |
| ENSG000 | -1.08 | -2.1134 | 5.09E-10 | 3.45E-08 | SERPINB1 | | |
| ENSG000 | -1.502 | -2.83199 | 1.07E-09 | 6.59E-08 | TYMP | | |
| ENSG000 | -1.11 | -2.15881 | 3.73E-12 | 3.98E-10 | EFCAB1 | | |
| ENSG000 | -1.134 | -2.19466 | 5.18E-06 | 0.000102 | LMO3 | | |
| ENSG000 | -1.544 | -2.91596 | 3.18E-10 | 2.23E-08 | ENTPD2 | | |
| ENSG000 | -1.03 | -2.0424 | 1.24E-09 | 7.40E-08 | TRAF1 | | |
| ENSG000 | -1.124 | -2.18002 | 0.00955 | 0.047697 | SPAG4 | | |
| ENSG000 | -1.766 | -3.40017 | 4.18E-07 | 1.17E-05 | ZMYND12 | | |
| ENSG000 | -1.301 | -2.46405 | 2.54E-05 | 0.000394 | ADCYAP1R1 | | |
| ENSG000 | -1.293 | -2.4499 | 0.002701 | 0.018019 | KCNK2 | | |
| ENSG000 | -1.282 | -2.43209 | 0.000677 | 0.00602 | SLCO1A2 | | |
| ENSG000 | -1.362 | -2.57043 | 7.39E-06 | 0.000139 | ABCC6 | | |
| ENSG000 | -1.692 | -3.2318 | 1.44E-20 | 9.59E-18 | ADA2 | | |
| ENSG000 | -1.626 | -3.08672 | 5.17E-07 | 1.41E-05 | RASL10A | | |
| ENSG000 | -1.227 | -2.34081 | 1.91E-08 | 8.20E-07 | ISM2 | | |
| ENSG000 | -1.582 | -2.99429 | 4.55E-27 | 1.31E-23 | EFS | | |
| ENSG000 | -1.241 | -2.36422 | 3.87E-09 | 1.99E-07 | NKAIN4 | | |
| ENSG000 | -1.26 | -2.39515 | 0.000974 | 0.00812 | RGCC | | |
| ENSG000 | -1.062 | -2.08834 | 9.19E-07 | 2.33E-05 | PEX11G | | |
| ENSG000 | -1.793 | -3.46582 | 8.78E-10 | 5.52E-08 | ODAD1 | | |
| ENSG000 | -1.015 | -2.02047 | 4.21E-05 | 0.0006 | DNAH11 | | |
| ENSG000 | -1.081 | -2.11496 | 0.002279 | 0.015796 | VSIR | | |
| ENSG000 | -1.038 | -2.05272 | 3.35E-23 | 5.27E-20 | RSPH4A | | |
| ENSG000 | -1.246 | -2.37242 | 1.21E-12 | 1.39E-10 | MDFI | | |
| ENSG000 | -1.03 | -2.04135 | 0.002622 | 0.017588 | SERPINI2 | | |
| ENSG000 | -1.342 | -2.53554 | 0.001101 | 0.008975 | CFAP92 | | |
| ENSG000 | -1.311 | -2.48163 | 0.000106 | 0.001296 | IRAG2 | | |
| ENSG000 | -1.587 | -3.00432 | 0.00506 | 0.029193 | ADGB | | |
| ENSG000 | -1.076 | -2.10776 | 8.03E-08 | 2.81E-06 | ECRG4 | | |
| ENSG000 | -1.138 | -2.20142 | 1.46E-19 | 7.84E-17 | CCDC170 | | |
| ENSG000 | -1.274 | -2.41771 | 1.04E-06 | 2.59E-05 | EPHX2 | | |
| ENSG000 | -1.648 | -3.13452 | 9.81E-21 | 7.06E-18 | DNAI1 | | |
| ENSG000 | -1.136 | -2.19818 | 0.00124 | 0.009825 | NFE2 | | |
| ENSG000 | -1.218 | -2.32634 | 0.00484 | 0.028169 | SLPI | | |
| ENSG000 | -1.403 | -2.64526 | 1.02E-05 | 0.000183 | KCNS1 | | |
| ENSG000 | -1.012 | -2.01691 | 1.85E-05 | 0.000301 | MATN4 | | |
| ENSG000 | -1.08 | -2.11362 | 2.64E-06 | 5.74E-05 | TOX2 | | |
| ENSG000 | -1.08 | -2.11394 | 6.17E-15 | 1.38E-12 | MASP1 | | |
| ENSG000 | -2.001 | -4.00243 | 0.000668 | 0.005968 | TNNI3 | | |
| ENSG000 | -1.021 | -2.02888 | 0.001378 | 0.010674 | PRRG3 | | |
| ENSG000 | -4.175 | -18.0633 | 0.008149 | 0.042113 | KLHDC7B | | |
| ENSG000 | -1.363 | -2.57218 | 0.000163 | 0.001859 | CALY | | |
| ENSG000 | -1.152 | -2.2219 | 1.31E-08 | 5.87E-07 | THEMIS2 | | |
| ENSG000 | -1.304 | -2.46987 | 9.22E-13 | 1.09E-10 | PPP1R1B | | |
| ENSG000 | -1.311 | -2.48175 | 9.64E-11 | 7.86E-09 | RGS22 | | |

| | | | | | |
|---|---|---|---|---|---|
| ENSG000 | -1.093 | -2.13259 | 2.36E-05 | 0.000369 | ANKEF1 |
| ENSG000 | -1.223 | -2.33418 | 0.003241 | 0.020676 | STOML3 |
| ENSG000 | -1.147 | -2.21473 | 0.000435 | 0.004221 | SFTPD |
| ENSG000 | -2.055 | -4.15581 | 1.69E-10 | 1.28E-08 | DYDC2 |
| ENSG000 | -1.437 | -2.70678 | 1.16E-07 | 3.84E-06 | SYT6 |
| ENSG000 | -1.126 | -2.18193 | 3.71E-09 | 1.92E-07 | MYCN |
| ENSG000 | -10.21 | -1184.65 | 0.000251 | 0.002686 | SAA2 |
| ENSG000 | -1.064 | -2.09082 | 5.25E-08 | 1.93E-06 | AGT |
| ENSG000 | -1.036 | -2.05005 | 0.002044 | 0.014497 | PCDH8 |
| ENSG000 | -1.692 | -3.23014 | 7.02E-09 | 3.33E-07 | CNMD |
| ENSG000 | -1.779 | -3.43208 | 5.29E-07 | 1.44E-05 | MYCBPAP |
| ENSG000 | -1.123 | -2.17813 | 0.0037 | 0.022912 | CHAD |
| ENSG000 | -1.038 | -2.05332 | 2.96E-10 | 2.08E-08 | WDR38 |
| ENSG000 | -1.456 | -2.7435 | 1.09E-09 | 6.64E-08 | TTC29 |
| ENSG000 | -1.412 | -2.66116 | 3.36E-07 | 9.62E-06 | SULF1 |
| ENSG000 | -1.015 | -2.02124 | 8.68E-10 | 5.50E-08 | CFAP300 |
| ENSG000 | -1.131 | -2.19041 | 6.69E-08 | 2.41E-06 | ITGB7 |
| ENSG000 | -1.235 | -2.35435 | 1.05E-11 | 1.03E-09 | MORN3 |
| ENSG000 | -1.153 | -2.2239 | 8.83E-20 | 5.09E-17 | AK7 |
| ENSG000 | -1.333 | -2.5187 | 1.04E-05 | 0.000185 | WDR93 |
| ENSG000 | -1.239 | -2.36113 | 1.46E-05 | 0.000248 | MYLK3 |
| ENSG000 | -1.37 | -2.58432 | 0.001952 | 0.014004 | IRF8 |
| ENSG000 | -1.029 | -2.04002 | 6.51E-11 | 5.51E-09 | LRRC46 |
| ENSG000 | -1.389 | -2.61922 | 0.003238 | 0.020668 | SLC14A1 |
| ENSG000 | -1.661 | -3.16195 | 0.007223 | 0.03842 | SLC13A5 |
| ENSG000 | -1.141 | -2.20524 | 0.000215 | 0.002348 | FBXO15 |
| ENSG000 | -1.635 | -3.1055 | 1.90E-13 | 2.76E-11 | CFAP74 |
| ENSG000 | -1.07 | -2.09882 | 1.96E-07 | 6.16E-06 | RXRG |
| ENSG000 | -1.748 | -3.35787 | 1.31E-13 | 1.99E-11 | ANKRD53 |
| ENSG000 | -1.683 | -3.2111 | 6.64E-17 | 1.98E-14 | SCN1A |
| ENSG000 | -1.783 | -3.44223 | 3.97E-07 | 1.11E-05 | DDIT4L |
| ENSG000 | -1.014 | -2.01913 | 0.001659 | 0.012343 | RNF175 |
| ENSG000 | -1.316 | -2.48926 | 5.07E-08 | 1.88E-06 | SLC25A48 |
| ENSG000 | -1.348 | -2.54474 | 0.000206 | 0.002267 | SLC22A3 |
| ENSG000 | -2.115 | -4.33115 | 0.001231 | 0.00978 | NKX6-2 |
| ENSG000 | -1.001 | -2.00115 | 4.77E-08 | 1.78E-06 | ADAM33 |
| ENSG000 | -1.315 | -2.48742 | 8.30E-05 | 0.001061 | KIAA1755 |
| ENSG000 | -1.429 | -2.69308 | 0.009867 | 0.048844 | DOC2A |
| ENSG000 | -1.326 | -2.50749 | 0.000101 | 0.001248 | CAPSL |
| ENSG000 | -1.032 | -2.04443 | 9.21E-17 | 2.70E-14 | DYNLT5 |
| ENSG000 | -1.362 | -2.57105 | 0.002345 | 0.016144 | NRSN1 |
| ENSG000 | -1.345 | -2.53993 | 0.005164 | 0.029667 | BANK1 |
| ENSG000 | -1.202 | -2.30072 | 0.004405 | 0.026216 | FAM81B |
| ENSG000 | -1.469 | -2.76893 | 3.50E-21 | 3.02E-18 | CHST9 |
| ENSG000 | -1.267 | -2.40605 | 8.04E-16 | 2.11E-13 | DNAAF1 |
| ENSG000 | -1.046 | -2.0644 | 2.34E-11 | 2.14E-09 | PITPNC1 |
| ENSG000 | -1.215 | -2.32123 | 2.23E-13 | 3.19E-11 | USP43 |
| ENSG000 | -1.014 | -2.02004 | 0.00401 | 0.024367 | RSPH10B |
| ENSG000 | -1.098 | -2.1401 | 1.50E-11 | 1.44E-09 | PTPRN2 |
| ENSG000 | -1.548 | -2.92476 | 5.55E-10 | 3.75E-08 | SPAG17 |

| | | | | | |
|---|---|---|---|---|---|
| ENSG000 | -1.248 | -2.37543 | 9.05E-10 | 5.63E-08 | CFAP70 |
| ENSG000 | -1.784 | -3.44477 | 5.02E-07 | 1.37E-05 | CFAP161 |
| ENSG000 | -1.049 | -2.06864 | 2.58E-18 | 1.15E-15 | FBXO32 |
| ENSG000 | -1.458 | -2.74674 | 7.51E-10 | 4.82E-08 | GRHL3 |
| ENSG000 | -1.741 | -3.34365 | 2.00E-14 | 3.83E-12 | LRRC43 |
| ENSG000 | -1.036 | -2.05115 | 0.007455 | 0.039315 | XDH |
| ENSG000 | -1.124 | -2.17954 | 0.003534 | 0.022085 | H2BC5 |
| ENSG000 | -1.042 | -2.05956 | 0.000113 | 0.001369 | CATIP |
| ENSG000 | -1.396 | -2.63127 | 3.78E-05 | 0.000549 | VWA5B1 |
| ENSG000 | -1.009 | -2.01217 | 1.02E-11 | 1.01E-09 | None |
| ENSG000 | -1.422 | -2.67984 | 8.54E-23 | 1.13E-19 | DRC7 |
| ENSG000 | -1.373 | -2.58998 | 1.66E-15 | 4.04E-13 | TPPP3 |
| ENSG000 | -1.18 | -2.2661 | 6.70E-22 | 7.24E-19 | CFAP157 |
| ENSG000 | -1.803 | -3.49023 | 2.29E-07 | 7.05E-06 | LRRC71 |
| ENSG000 | -1.046 | -2.065 | 5.41E-05 | 0.00074 | LRRC56 |
| ENSG000 | -1.723 | -3.30108 | 0.000407 | 0.00399 | AQP5 |
| ENSG000 | -1.726 | -3.30902 | 1.90E-08 | 8.19E-07 | UBXN10 |
| ENSG000 | -3.492 | -11.248 | 3.92E-05 | 0.000566 | GABRG1 |
| ENSG000 | -1.017 | -2.02356 | 0.003536 | 0.022091 | DCST2 |
| ENSG000 | -1.208 | -2.3108 | 1.00E-05 | 0.000181 | EFHB |
| ENSG000 | -1.661 | -3.16231 | 1.15E-19 | 6.42E-17 | CFAP100 |
| ENSG000 | -1.099 | -2.14233 | 1.71E-09 | 9.89E-08 | ZNF474 |
| ENSG000 | -1.158 | -2.23104 | 3.03E-05 | 0.000457 | PI16 |
| ENSG000 | -1.108 | -2.15573 | 4.86E-09 | 2.41E-07 | SYTL3 |
| ENSG000 | -1.058 | -2.08236 | 1.34E-08 | 5.95E-07 | HEPACAM |
| ENSG000 | -1.036 | -2.05058 | 0.003318 | 0.021062 | PDZRN4 |
| ENSG000 | -1.097 | -2.13858 | 0.003513 | 0.02199 | TMEM130 |
| ENSG000 | -1.615 | -3.06369 | 6.03E-07 | 1.61E-05 | A2ML1 |
| ENSG000 | -1.195 | -2.2897 | 1.02E-15 | 2.59E-13 | CFAP52 |
| ENSG000 | -1.664 | -3.16936 | 4.07E-07 | 1.14E-05 | SGSM1 |
| ENSG000 | -1.298 | -2.4596 | 2.88E-19 | 1.46E-16 | DNAAF3 |
| ENSG000 | -2.172 | -4.50506 | 3.97E-08 | 1.52E-06 | ANGPTL4 |
| ENSG000 | -1.653 | -3.14511 | 6.51E-18 | 2.62E-15 | TEKT1 |
| ENSG000 | -1.512 | -2.8515 | 0.004623 | 0.027212 | SCNN1B |
| ENSG000 | -1.049 | -2.0695 | 2.05E-05 | 0.000328 | LGI3 |
| ENSG000 | -1.059 | -2.08394 | 1.16E-05 | 0.000205 | VWA3B |
| ENSG000 | -1.407 | -2.65165 | 0.007599 | 0.039902 | RSPH10B2 |
| ENSG000 | -1.494 | -2.816 | 1.51E-06 | 3.54E-05 | FRMPD2 |
| ENSG000 | -1.066 | -2.09378 | 1.19E-07 | 3.92E-06 | DCDC1 |
| ENSG000 | -1.077 | -2.11001 | 5.40E-11 | 4.64E-09 | GRIK1 |
| ENSG000 | -1.703 | -3.25661 | 0.001722 | 0.012704 | DNAI2 |
| ENSG000 | -1.059 | -2.08412 | 3.80E-09 | 1.95E-07 | LRRC34 |
| ENSG000 | -1.793 | -3.46437 | 0.000599 | 0.005454 | IL16 |
| ENSG000 | -1.143 | -2.20905 | 4.04E-24 | 6.98E-21 | EFCAB12 |
| ENSG000 | -1.419 | -2.67484 | 0.000458 | 0.004403 | HPSE2 |
| ENSG000 | -1.19 | -2.28123 | 7.72E-05 | 0.001 | ZNF483 |
| ENSG000 | -6.123 | -69.6968 | 0.00758 | 0.039834 | SAA1 |
| ENSG000 | -1.447 | -2.72674 | 4.86E-06 | 9.64E-05 | FAM166C |
| ENSG000 | -1.328 | -2.50977 | 1.09E-05 | 0.000193 | CCDC13-AS1 |
| ENSG000 | -1.774 | -3.4211 | 1.45E-06 | 3.44E-05 | SNX31 |

| | | | | | |
|---|---|---|---|---|---|
| ENSG000 | -1.551 | -2.92992 | 0.002149 | 0.015096 | DNAH12 |
| ENSG000 | -1.212 | -2.3158 | 2.12E-09 | 1.18E-07 | FAM182B |
| ENSG000 | -1.438 | -2.71032 | 2.88E-20 | 1.78E-17 | VWA3A |
| ENSG000 | -1.005 | -2.00691 | 0.00285 | 0.018767 | GRAMD2A |
| ENSG000 | -1.205 | -2.30517 | 0.000567 | 0.005215 | TEX26 |
| ENSG000 | -1.392 | -2.62421 | 0.001537 | 0.01162 | LINC01106 |
| ENSG000 | -1.061 | -2.08576 | 0.005449 | 0.030996 | PLEKHD1 |
| ENSG000 | -1.377 | -2.59649 | 1.77E-09 | 1.02E-07 | PRR18 |
| ENSG000 | -1.44 | -2.71303 | 4.56E-25 | 8.75E-22 | MAP3K19 |
| ENSG000 | -1.29 | -2.44482 | 8.13E-20 | 4.85E-17 | FIBIN |
| ENSG000 | -1.816 | -3.52068 | 1.87E-14 | 3.63E-12 | ODF3B |
| ENSG000 | -1.39 | -2.6201 | 1.75E-13 | 2.59E-11 | FAR2P2 |
| ENSG000 | -1.081 | -2.11578 | 8.34E-08 | 2.91E-06 | ERICH3 |
| ENSG000 | -1.348 | -2.54479 | 0.00726 | 0.038554 | DAND5 |
| ENSG000 | -1.006 | -2.00818 | 2.88E-14 | 5.24E-12 | SLC17A8 |
| ENSG000 | -1.035 | -2.04944 | 0.008579 | 0.043848 | PCED1B |
| ENSG000 | -1.126 | -2.18276 | 3.87E-06 | 7.97E-05 | KCNE1 |
| ENSG000 | -1.14 | -2.2033 | 1.06E-05 | 0.000189 | TSPYL5 |
| ENSG000 | -1.563 | -2.95469 | 1.41E-14 | 2.87E-12 | FAM83H |
| ENSG000 | -1.376 | -2.59618 | 2.44E-14 | 4.54E-12 | NME9 |
| ENSG000 | -1.353 | -2.55467 | 1.54E-10 | 1.19E-08 | CFAP65 |
| ENSG000 | -2.885 | -7.38917 | 3.43E-06 | 7.18E-05 | ATG9B |
| ENSG000 | -1.021 | -2.02973 | 2.26E-07 | 6.98E-06 | HHIPL1 |
| ENSG000 | -1.2 | -2.2968 | 2.34E-05 | 0.000367 | KIAA2012 |
| ENSG000 | -1.86 | -3.63105 | 0.006969 | 0.037447 | NXPH3 |
| ENSG000 | -1.231 | -2.34749 | 0.002023 | 0.014394 | CSMD1 |
| ENSG000 | -1.071 | -2.10028 | 4.25E-08 | 1.61E-06 | CABCOCO1 |
| ENSG000 | -1.781 | -3.4377 | 2.00E-05 | 0.000321 | HOATZ |
| ENSG000 | -1.366 | -2.57712 | 5.66E-10 | 3.79E-08 | LRRC55 |
| ENSG000 | -1.29 | -2.44487 | 0.000222 | 0.002413 | DNAH2 |
| ENSG000 | -1.291 | -2.4464 | 8.71E-06 | 0.000161 | SDR42E2 |
| ENSG000 | -2.197 | -4.58681 | 1.19E-26 | 2.93E-23 | CLDN5 |
| ENSG000 | -1.05 | -2.06994 | 3.70E-09 | 1.92E-07 | KCNQ3 |
| ENSG000 | -1.151 | -2.22096 | 3.84E-07 | 1.08E-05 | EFCAB10 |
| ENSG000 | -1.118 | -2.17014 | 8.44E-08 | 2.93E-06 | MORN5 |
| ENSG000 | -1.822 | -3.53619 | 3.39E-14 | 5.98E-12 | KLHL32 |
| ENSG000 | -1.756 | -3.37655 | 1.39E-06 | 3.30E-05 | LINC00643 |
| ENSG000 | -1.109 | -2.15727 | 9.30E-06 | 0.00017 | PDE2A |
| ENSG000 | -1.491 | -2.81121 | 5.97E-19 | 2.95E-16 | CFAP73 |
| ENSG000 | -1.455 | -2.7407 | 3.49E-05 | 0.000512 | DNAJB13 |
| ENSG000 | -1.498 | -2.82485 | 1.76E-15 | 4.22E-13 | DIPK1C |
| ENSG000 | -1.311 | -2.4804 | 0.000564 | 0.005191 | LRRC74B |
| ENSG000 | -1.403 | -2.64396 | 0.000141 | 0.001647 | SNHG28 |
| ENSG000 | -1.328 | -2.51099 | 0.000965 | 0.008068 | PRELP |
| ENSG000 | -1.418 | -2.67131 | 1.01E-06 | 2.53E-05 | HACD4 |
| ENSG000 | -2.26 | -4.79132 | 1.11E-08 | 5.05E-07 | KIF19 |
| ENSG000 | -1.054 | -2.07625 | 0.008564 | 0.043787 | SEMA4A |
| ENSG000 | -1.288 | -2.44258 | 1.56E-06 | 3.64E-05 | ARL9 |
| ENSG000 | -1.026 | -2.03672 | 0.000565 | 0.005201 | DTHD1 |
| ENSG000 | -1.062 | -2.0881 | 2.62E-13 | 3.65E-11 | CFAP43 |

| | | | | | |
|---|---|---|---|---|---|
| ENSG000 | -1.514 | -2.85559 | 0.000876 | 0.007477 | CFAP299 |
| ENSG000 | -1.906 | -3.74813 | 0.004889 | 0.028408 | HES5 |
| ENSG000 | -1.336 | -2.52467 | 0.002559 | 0.017233 | LEKR1 |
| ENSG000 | -1.238 | -2.35938 | 2.82E-06 | 6.07E-05 | MMP17 |
| ENSG000 | -1.094 | -2.13466 | 2.18E-11 | 2.03E-09 | DLL1 |
| ENSG000 | -1.395 | -2.62981 | 1.75E-05 | 0.000287 | SMOC1 |
| ENSG000 | -1.808 | -3.50131 | 0.000567 | 0.00521 | Y_RNA |
| ENSG000 | -1.124 | -2.17974 | 0.007054 | 0.037765 | TDRKH-AS1 |
| ENSG000 | -1.406 | -2.64919 | 1.87E-07 | 5.92E-06 | IQANK1 |
| ENSG000 | -1.116 | -2.16721 | 3.19E-07 | 9.19E-06 | PRRT1 |
| ENSG000 | -1.624 | -3.08237 | 0.003372 | 0.021338 | ERICH2 |
| ENSG000 | -1.24 | -2.36261 | 7.03E-12 | 7.19E-10 | CYP21A1P |
| ENSG000 | -1.29 | -2.44445 | 2.91E-06 | 6.24E-05 | BTBD17 |
| ENSG000 | -1.531 | -2.88987 | 8.62E-08 | 2.98E-06 | LRRC10B |
| ENSG000 | -1.6 | -3.03208 | 0.000417 | 0.004069 | MT1M |
| ENSG000 | -1.28 | -2.42806 | 0.000376 | 0.003735 | GMNC |
| ENSG000 | -1.014 | -2.01882 | 3.89E-09 | 1.99E-07 | NYNRIN |
| ENSG000 | -1.378 | -2.59841 | 1.42E-17 | 5.14E-15 | CFAP99 |
| ENSG000 | -1.035 | -2.04896 | 0.00515 | 0.029646 | MIR149 |
| ENSG000 | -1.179 | -2.26376 | 0.008354 | 0.04299 | None |
| ENSG000 | -1.332 | -2.51765 | 1.45E-30 | 2.50E-26 | CFAP45 |
| ENSG000 | -1.558 | -2.94448 | 7.17E-08 | 2.56E-06 | C10orf105 |
| ENSG000 | -1.052 | -2.0729 | 0.000186 | 0.002085 | COL28A1 |
| ENSG000 | -1.09 | -2.12933 | 4.71E-13 | 5.94E-11 | C5orf49 |
| ENSG000 | -1.428 | -2.69128 | 8.64E-06 | 0.00016 | SIAH3 |
| ENSG000 | -1.046 | -2.06479 | 0.000196 | 0.002174 | None |
| ENSG000 | -1.781 | -3.43571 | 0.002356 | 0.016194 | LINC01422 |
| ENSG000 | -1.17 | -2.24998 | 0.00168 | 0.012456 | WARS2-IT1 |
| ENSG000 | -2.133 | -4.38774 | 0.000722 | 0.006344 | LINC00092 |
| ENSG000 | -3.382 | -10.4268 | 0.006056 | 0.033672 | ORM2 |
| ENSG000 | -1.287 | -2.44012 | 1.09E-09 | 6.64E-08 | None |
| ENSG000 | -1.488 | -2.8053 | 1.54E-20 | 9.89E-18 | PROB1 |
| ENSG000 | -1.264 | -2.40184 | 0.000377 | 0.003744 | FRMD3-AS1 |
| ENSG000 | -1.261 | -2.39729 | 0.004763 | 0.027871 | None |
| ENSG000 | -4.483 | -22.3621 | 0.003018 | 0.019614 | ORM1 |
| ENSG000 | -1.512 | -2.85191 | 0.005751 | 0.032381 | None |
| ENSG000 | -1.198 | -2.29395 | 3.46E-08 | 1.36E-06 | CYP21A2 |
| ENSG000 | -1.016 | -2.02296 | 9.03E-07 | 2.29E-05 | TGFB2-AS1 |
| ENSG000 | -1.724 | -3.30387 | 0.006179 | 0.034156 | TMEM229A |
| ENSG000 | -1.77 | -3.41113 | 0.001731 | 0.012759 | OBI1-AS1 |
| ENSG000 | -1.216 | -2.32303 | 3.56E-05 | 0.00052 | DAAM2-AS1 |
| ENSG000 | -1.322 | -2.49995 | 0.007251 | 0.03852 | DNM1P51 |
| ENSG000 | -1.198 | -2.2943 | 0.001529 | 0.011587 | DPY19L2P4 |
| ENSG000 | -1.181 | -2.26801 | 2.89E-07 | 8.46E-06 | DOCK7-DT |
| ENSG000 | -1.147 | -2.21497 | 0.00684 | 0.036917 | None |
| ENSG000 | -2.136 | -4.3953 | 3.05E-05 | 0.000458 | None |
| ENSG000 | -1.705 | -3.26078 | 0.009402 | 0.04712 | None |
| ENSG000 | -1.309 | -2.47807 | 0.000245 | 0.002634 | CT75 |
| ENSG000 | -1.667 | -3.17584 | 0.000121 | 0.001442 | LRIG2-DT |
| ENSG000 | -1.409 | -2.6564 | 0.000929 | 0.007815 | RPS27P25 |

| | | | | | |
|---|---|---|---|---|---|
| ENSG000 | -1.519 | -2.86557 | 0.000112 | 0.001356 | None |
| ENSG000 | -1.099 | -2.14205 | 0.000522 | 0.004881 | SMKR1 |
| ENSG000 | -1.68 | -3.20441 | 0.000465 | 0.004463 | VSIG8 |
| ENSG000 | -1.058 | -2.08134 | 1.07E-14 | 2.21E-12 | CCDC13 |
| ENSG000 | -1.001 | -2.0009 | 1.65E-05 | 0.000273 | USP2-AS1 |
| ENSG000 | -1.087 | -2.12412 | 0.002295 | 0.015887 | None |
| ENSG000 | -1.251 | -2.3796 | 1.86E-07 | 5.92E-06 | TNXA |
| ENSG000 | -1.327 | -2.5092 | 4.17E-06 | 8.46E-05 | None |
| ENSG000 | -1.048 | -2.06801 | 0.004166 | 0.02506 | ALG1L9P |
| ENSG000 | -1.051 | -2.07172 | 3.45E-05 | 0.000505 | C8orf34-AS1 |
| ENSG000 | -1.015 | -2.02082 | 0.00242 | 0.016508 | TICAM2-AS1 |
| ENSG000 | -1.303 | -2.46673 | 5.78E-05 | 0.000784 | None |
| ENSG000 | -1.223 | -2.33355 | 0.004037 | 0.024488 | ZNF474-AS1 |
| ENSG000 | -1.07 | -2.09899 | 1.51E-05 | 0.000254 | None |
| ENSG000 | -1.188 | -2.27808 | 0.004252 | 0.025504 | None |
| ENSG000 | -1.208 | -2.31005 | 0.000455 | 0.004381 | STK19B |
| ENSG000 | -1.259 | -2.39264 | 0.003101 | 0.020034 | KBTBD11-AS1 |
| ENSG000 | -1.93 | -3.80966 | 0.002475 | 0.016799 | RASSF10-DT |
| ENSG000 | -1.397 | -2.63435 | 0.002541 | 0.017139 | B3GAT1-DT |
| ENSG000 | -1.008 | -2.01054 | 4.56E-08 | 1.71E-06 | TRIL |
| ENSG000 | -1.783 | -3.44188 | 3.23E-05 | 0.00048 | None |
| ENSG000 | -2.922 | -7.58144 | 0.007748 | 0.040531 | HP |
| ENSG000 | -1.955 | -3.87653 | 0.000117 | 0.001405 | None |
| ENSG000 | -1.013 | -2.01815 | 8.61E-07 | 2.21E-05 | LINC01579 |
| ENSG000 | -3.114 | -8.65798 | 2.14E-06 | 4.77E-05 | None |
| ENSG000 | -1.366 | -2.57711 | 0.000627 | 0.005662 | LINC02352 |
| ENSG000 | -1.175 | -2.25717 | 0.00154 | 0.011631 | None |
| ENSG000 | -1.082 | -2.11716 | 4.34E-06 | 8.74E-05 | None |
| ENSG000 | -1.148 | -2.21606 | 1.72E-05 | 0.000284 | None |
| ENSG000 | -1.167 | -2.24484 | 0.000538 | 0.004999 | None |
| ENSG000 | -1.003 | -2.00455 | 8.64E-05 | 0.0011 | TSPAN5-DT |
| ENSG000 | -1.823 | -3.53912 | 0.000317 | 0.003252 | |
| ENSG000 | -1.512 | -2.85284 | 0.001573 | 0.011847 | LINC02175 |
| ENSG000 | -1.44 | -2.71283 | 4.32E-07 | 1.20E-05 | SPON1 |
| ENSG000 | -1.317 | -2.4911 | 0.005861 | 0.032853 | None |
| ENSG000 | -1.11 | -2.15795 | 0.002266 | 0.015729 | TMEM220-AS1 |
| ENSG000 | -1.111 | -2.15935 | 0.008058 | 0.041727 | AGAP12P |
| ENSG000 | -1.212 | -2.31597 | 9.15E-21 | 6.88E-18 | MYO15B |
| ENSG000 | -1.493 | -2.81533 | 1.91E-10 | 1.41E-08 | CCDC177 |
| ENSG000 | -1.531 | -2.89076 | 0.00025 | 0.002678 | None |
| ENSG000 | -1.81 | -3.5071 | 0.001445 | 0.011098 | MEI4 |
| ENSG000 | -1.699 | -3.24636 | 0.000211 | 0.00231 | GAS2L2 |
| ENSG000 | -1.327 | -2.50889 | 0.002212 | 0.015428 | None |
| ENSG000 | -1.506 | -2.83978 | 0.002123 | 0.014957 | LINC01607 |
| ENSG000 | -1.484 | -2.79685 | 0.000316 | 0.003249 | None |
| ENSG000 | -1.057 | -2.08057 | 9.61E-07 | 2.43E-05 | MESTIT1 |
| ENSG000 | -1.26 | -2.39526 | 0.009844 | 0.048756 | None |
| ENSG000 | -1.266 | -2.40452 | 6.70E-07 | 1.77E-05 | ADIRF-AS1 |
| ENSG000 | -1.186 | -2.27506 | 0.002359 | 0.016202 | None |
| ENSG000 | -1.48 | -2.78951 | 0.003119 | 0.020112 | LENG9 |

| | | | | | |
|---|---|---|---|---|---|
| ENSG000 | -1.164 | -2.24118 | 1.10E-06 | 2.73E-05 | FLJ16779 |
| ENSG000 | -1.438 | -2.7087 | 3.05E-14 | 5.44E-12 | None |
| ENSG000 | -1.999 | -3.99856 | 1.19E-07 | 3.92E-06 | None |
| ENSG000 | -1.178 | -2.26283 | 0.007941 | 0.041261 | None |
