## Supplementary material for "Unravelling neuronal and glial differences in ceramide composition, synthesis, and sensitivity to toxicity": Document2_Venn_List Astrocyte FB1-down GCSi-up

| GeneID | Treatm | Treatme | Treatment | Treatment | name | | Document2_Venn_List Astrocyte FB1-down GCSi-up |
|---|---|---|---|---|---|---|---|
| ENSG0( | 1.1 | 2.1442 | 3.33E-15 | 7.66E-13 | TNFRSF12A | | |
| ENSG0( | 1.21 | 2.3135 | 7.58E-05 | 0.00099 | BIRC3 | | |
| ENSG0( | 1.09 | 2.1294 | 1.54E-07 | 4.95E-06 | PTPRN | | |
| ENSG0( | -1.03 | -2.042 | 1.24E-09 | 7.40E-08 | TRAF1 | | |
| ENSG0( | 1.506 | 2.8402 | 4.20E-21 | 3.45E-18 | SREBF1 | | |
| ENSG0( | 1.054 | 2.0764 | 0.00506 | 0.02919 | ATP8B1 | | |
| ENSG0( | 1.054 | 2.0769 | 9.68E-06 | 0.00018 | MPP4 | | |
| ENSG0( | 1.513 | 2.8548 | 1.78E-10 | 1.33E-08 | ICAM1 | | |
| ENSG0( | 2.469 | 5.5381 | 1.12E-07 | 3.74E-06 | IL11 | | |
| ENSG0( | 1.578 | 2.9847 | 2.87E-29 | 1.64E-25 | SCD | | |
| ENSG0( | 1.168 | 2.2473 | 6.10E-10 | 4.04E-08 | RELB | | |
| ENSG0( | 1.318 | 2.4923 | 0.00604 | 0.03362 | ICAM5 | | |
| ENSG0( | 1.245 | 2.3699 | 1.15E-17 | 4.31E-15 | TFPI2 | | |
| ENSG0( | -1.015 | -2.02 | 4.21E-05 | 0.0006 | DNAH11 | | |
| ENSG0( | 1.789 | 3.4548 | 5.72E-18 | 2.41E-15 | SERPINE1 | | |
| ENSG0( | 1.151 | 2.2199 | 1.64E-06 | 3.79E-05 | ACTA2 | | |
| ENSG0( | 1.282 | 2.4312 | 2.87E-06 | 6.17E-05 | DKK1 | | |
| ENSG0( | 1.371 | 2.5868 | 6.70E-14 | 1.07E-11 | CCL2 | | |
| ENSG0( | 1.379 | 2.6014 | 0.00147 | 0.01124 | VWF | | |
| ENSG0( | -1.038 | -2.053 | 3.35E-23 | 5.27E-20 | RSPH4A | | |
| ENSG0( | -1.03 | -2.041 | 0.00262 | 0.01759 | SERPINI2 | | |
| ENSG0( | 1.178 | 2.2628 | 2.80E-09 | 1.51E-07 | IL1R1 | | |
| ENSG0( | 1.176 | 2.2594 | 7.21E-22 | 7.33E-19 | GADD45A | | |
| ENSG0( | 2.175 | 4.5156 | 1.47E-18 | 6.85E-16 | RGS4 | | |
| ENSG0( | 1.312 | 2.4823 | 1.24E-20 | 8.60E-18 | TNFAIP3 | | |
| ENSG0( | 1.363 | 2.5727 | 8.93E-21 | 6.88E-18 | SGK1 | | |
| ENSG0( | 1.105 | 2.1516 | 1.58E-05 | 0.00026 | CCN2 | | |
| ENSG0( | 1.18 | 2.2664 | 1.96E-08 | 8.34E-07 | DUSP1 | | |
| ENSG0( | 1.605 | 3.0419 | 2.57E-16 | 7.29E-14 | EGR1 | | |
| ENSG0( | 1.593 | 3.0164 | 3.79E-29 | 1.64E-25 | DUSP4 | | |
| ENSG0( | 1.223 | 2.334 | 2.03E-06 | 4.55E-05 | SV2C | | |
| ENSG0( | 1.904 | 3.7433 | 7.81E-22 | 7.50E-19 | EGR2 | | |
| ENSG0( | -1.012 | -2.017 | 1.85E-05 | 0.0003 | MATN4 | | |
| ENSG0( | 1.027 | 2.0383 | 9.83E-05 | 0.00122 | CHST8 | | |
| ENSG0( | 1.059 | 2.0834 | 0.00013 | 0.0015 | GRPR | | |
| ENSG0( | 1.258 | 2.3917 | 8.05E-11 | 6.63E-09 | WNK4 | | |
| ENSG0( | 1.215 | 2.3209 | 2.44E-14 | 4.54E-12 | LIF | | |
| ENSG0( | 1.791 | 3.4613 | 1.05E-13 | 1.63E-11 | CPA4 | | |
| ENSG0( | 1.557 | 2.9422 | 6.80E-10 | 4.47E-08 | VGF | | |
| ENSG0( | -1.021 | -2.029 | 0.00138 | 0.01067 | PRRG3 | | |
| ENSG0( | 1.098 | 2.1411 | 0.00203 | 0.01443 | NALF2 | | |
| ENSG0( | 1.02 | 2.0276 | 5.49E-14 | 8.95E-12 | GFPT2 | | |
| ENSG0( | 1.493 | 2.8147 | 4.54E-23 | 6.54E-20 | CHI3L1 | | |
| ENSG0( | 2.061 | 4.1729 | 3.71E-18 | 1.60E-15 | SPOCD1 | | |
| ENSG0( | 1.003 | 2.0035 | 1.44E-11 | 1.38E-09 | ETS1 | | |
| ENSG0( | 2.088 | 4.2504 | 4.31E-06 | 8.69E-05 | EGR4 | | |
| ENSG0( | -1.036 | -2.05 | 0.00204 | 0.0145 | PCDH8 | | |
| ENSG0( | 2.089 | 4.2546 | 1.11E-06 | 2.73E-05 | GPNMB | | |
| ENSG0( | -1.038 | -2.053 | 2.96E-10 | 2.08E-08 | WDR38 | | |
| ENSG0( | 1.136 | 2.1971 | 1.33E-15 | 3.27E-13 | IER3 | | |
| ENSG0( | -1.015 | -2.021 | 8.68E-10 | 5.50E-08 | CFAP300 | | |
| ENSG0( | 1.161 | 2.2358 | 1.58E-10 | 1.21E-08 | THBS1 | | |
| ENSG0( | 1.001 | 2.0021 | 4.16E-08 | 1.59E-06 | DUSP6 | | |

| | | | | | |
|---|---|---|---|---|---|
| ENSG0( | -1.029 | -2.04 | 6.51E-11 | 5.51E-09 | LRRC46 |
| ENSG0( | 1.582 | 2.993 | 0.0073 | 0.03872 | PMAIP1 |
| ENSG0( | 1.042 | 2.0584 | 5.87E-09 | 2.82E-07 | CCN1 |
| ENSG0( | 1.034 | 2.0478 | 0.00148 | 0.01133 | KCNH1 |
| ENSG0( | -1.014 | -2.019 | 0.00166 | 0.01234 | RNF175 |
| ENSG0( | 1.029 | 2.0404 | 1.69E-21 | 1.54E-18 | CDKN2B |
| ENSG0( | 1.059 | 2.0837 | 2.19E-13 | 3.15E-11 | ADAM12 |
| ENSG0( | 1.235 | 2.3542 | 0.0001 | 0.00125 | P4HA3 |
| ENSG0( | -1.001 | -2.001 | 4.77E-08 | 1.78E-06 | ADAM33 |
| ENSG0( | 1.113 | 2.1632 | 2.77E-05 | 0.00042 | TAGLN |
| ENSG0( | 1.202 | 2.3011 | 5.71E-07 | 1.54E-05 | IL18 |
| ENSG0( | -1.032 | -2.044 | 9.21E-17 | 2.70E-14 | DYNLT5 |
| ENSG0( | -1.046 | -2.064 | 2.34E-11 | 2.14E-09 | PITPNC1 |
| ENSG0( | -1.014 | -2.02 | 0.00401 | 0.02437 | RSPH10B |
| ENSG0( | -1.049 | -2.069 | 2.58E-18 | 1.15E-15 | FBXO32 |
| ENSG0( | 1.116 | 2.1678 | 4.59E-13 | 5.83E-11 | NRG1 |
| ENSG0( | -1.036 | -2.051 | 0.00746 | 0.03932 | XDH |
| ENSG0( | -1.042 | -2.06 | 0.00011 | 0.00137 | CATIP |
| ENSG0( | -1.009 | -2.012 | 1.02E-11 | 1.01E-09 | None |
| ENSG0( | 1.003 | 2.0039 | 1.76E-09 | 1.01E-07 | ABCG1 |
| ENSG0( | 1.059 | 2.0834 | 0.00147 | 0.01125 | GAB3 |
| ENSG0( | 1.433 | 2.6994 | 0.00031 | 0.00317 | ICOSLG |
| ENSG0( | -1.046 | -2.065 | 5.41E-05 | 0.00074 | LRRC56 |
| ENSG0( | 1.031 | 2.0437 | 8.93E-12 | 8.98E-10 | ATF3 |
| ENSG0( | 1.025 | 2.0345 | 9.53E-06 | 0.00017 | KCNF1 |
| ENSG0( | -1.017 | -2.024 | 0.00354 | 0.02209 | DCST2 |
| ENSG0( | 1.134 | 2.1946 | 8.00E-05 | 0.00103 | CDCP1 |
| ENSG0( | 1.275 | 2.4194 | 1.42E-12 | 1.62E-10 | DLC1 |
| ENSG0( | -1.036 | -2.051 | 0.00332 | 0.02106 | PDZRN4 |
| ENSG0( | 1.2 | 2.2975 | 0.00224 | 0.01557 | MMP10 |
| ENSG0( | -1.049 | -2.07 | 2.05E-05 | 0.00033 | LGI3 |
| ENSG0( | 1.323 | 2.5013 | 1.10E-12 | 1.28E-10 | STXBP6 |
| ENSG0( | 1.181 | 2.2669 | 5.57E-07 | 1.51E-05 | VXN |
| ENSG0( | 1.082 | 2.1171 | 8.33E-12 | 8.42E-10 | EFNA1 |
| ENSG0( | 2.036 | 4.1004 | 0.00227 | 0.01577 | CXCL8 |
| ENSG0( | 1.141 | 2.2058 | 5.84E-13 | 7.21E-11 | FOS |
| ENSG0( | 1.129 | 2.1865 | 1.38E-12 | 1.57E-10 | METTL7B |
| ENSG0( | 1.26 | 2.3955 | 5.18E-17 | 1.63E-14 | JUNB |
| ENSG0( | 1.396 | 2.6322 | 0.00033 | 0.00339 | CHRNA9 |
| ENSG0( | 1.622 | 3.0786 | 2.35E-06 | 5.19E-05 | CMKLR1 |
| ENSG0( | 1.13 | 2.188 | 4.63E-05 | 0.00065 | ASPHD1 |
| ENSG0( | -1.005 | -2.007 | 0.00285 | 0.01877 | GRAMD2A |
| ENSG0( | 1.277 | 2.4235 | 1.03E-28 | 3.55E-25 | LPL |
| ENSG0( | 1.457 | 2.7455 | 5.52E-17 | 1.70E-14 | CLCF1 |
| ENSG0( | 1.893 | 3.7141 | 8.69E-18 | 3.34E-15 | FOSL1 |
| ENSG0( | 1.16 | 2.2345 | 0.00592 | 0.03306 | SPHK1 |
| ENSG0( | 1.041 | 2.0578 | 2.42E-14 | 4.54E-12 | MYO1D |
| ENSG0( | 1.38 | 2.6035 | 1.54E-11 | 1.46E-09 | BDNF |
| ENSG0( | 1.04 | 2.0559 | 0.00013 | 0.00156 | GCNT4 |
| ENSG0( | 1.594 | 3.0193 | 2.64E-25 | 5.69E-22 | EGR3 |
| ENSG0( | -1.006 | -2.008 | 2.88E-14 | 5.24E-12 | SLC17A8 |
| ENSG0( | -1.035 | -2.049 | 0.00858 | 0.04385 | PCED1B |
| ENSG0( | 1.873 | 3.6631 | 0.00347 | 0.02179 | C3orf80 |
| ENSG0( | 1.053 | 2.0742 | 6.02E-13 | 7.38E-11 | ARSJ |

| | | | | | |
|---|---|---|---|---|---|
| ENSG0( | 1.103 | 2.148 | 1.33E-08 | 5.95E-07 | GPR3 |
| ENSG0( | -1.021 | -2.03 | 2.26E-07 | 6.98E-06 | HHIPL1 |
| ENSG0( | 1.126 | 2.1826 | 0.00016 | 0.00184 | TNFAIP8L3 |
| ENSG0( | -1.05 | -2.07 | 3.70E-09 | 1.92E-07 | KCNQ3 |
| ENSG0( | 1.282 | 2.4326 | 4.35E-12 | 4.61E-10 | MAFF |
| ENSG0( | 1.199 | 2.2958 | 9.65E-07 | 2.43E-05 | MYBL1 |
| ENSG0( | 1.342 | 2.5348 | 2.94E-14 | 5.29E-12 | SPRY4 |
| ENSG0( | 1.272 | 2.4152 | 4.05E-16 | 1.09E-13 | SPRED3 |
| ENSG0( | 1.667 | 3.1747 | 7.44E-05 | 0.00097 | SH2D5 |
| ENSG0( | -1.054 | -2.076 | 0.00856 | 0.04379 | SEMA4A |
| ENSG0( | 1.024 | 2.0338 | 4.42E-09 | 2.21E-07 | HRH1 |
| ENSG0( | -1.026 | -2.037 | 0.00057 | 0.0052 | DTHD1 |
| ENSG0( | 2.377 | 5.1959 | 0.00023 | 0.00247 | PLN |
| ENSG0( | 1.969 | 3.9152 | 7.46E-30 | 6.45E-26 | ARC |
| ENSG0( | -1.014 | -2.019 | 3.89E-09 | 1.99E-07 | NYNRIN |
| ENSG0( | -1.035 | -2.049 | 0.00515 | 0.02965 | MIR149 |
| ENSG0( | -1.052 | -2.073 | 0.00019 | 0.00209 | COL28A1 |
| ENSG0( | -1.046 | -2.065 | 0.0002 | 0.00217 | None |
| ENSG0( | 1.587 | 3.0037 | 1.65E-06 | 3.79E-05 | ZNF812P |
| ENSG0( | 1.126 | 2.182 | 0.00812 | 0.042 | None |
| ENSG0( | 1.022 | 2.031 | 0.00425 | 0.0255 | None |
| ENSG0( | 1.721 | 3.2971 | 0.00014 | 0.00167 | MYOSLID |
| ENSG0( | -1.016 | -2.023 | 9.03E-07 | 2.29E-05 | TGFB2-AS1 |
| ENSG0( | 1.489 | 2.8062 | 2.41E-05 | 0.00038 | LINC00707 |
| ENSG0( | 1.391 | 2.6228 | 2.10E-07 | 6.53E-06 | |
| ENSG0( | 1.324 | 2.5034 | 1.61E-06 | 3.72E-05 | GSTA1 |
| ENSG0( | -1.001 | -2.001 | 1.65E-05 | 0.00027 | USP2-AS1 |
| ENSG0( | -1.048 | -2.068 | 0.00417 | 0.02506 | ALG1L9P |
| ENSG0( | -1.051 | -2.072 | 3.45E-05 | 0.00051 | C8orf34-AS1 |
| ENSG0( | -1.015 | -2.021 | 0.00242 | 0.01651 | TICAM2-AS1 |
| ENSG0( | 1.091 | 2.1304 | 0.00102 | 0.0084 | LINC02732 |
| ENSG0( | 1.081 | 2.1149 | 1.22E-06 | 2.96E-05 | None |
| ENSG0( | -1.008 | -2.011 | 4.56E-08 | 1.71E-06 | TRIL |
| ENSG0( | 1.734 | 3.3257 | 0.00354 | 0.02211 | PPP1R14B-AS1 |
| ENSG0( | 1.834 | 3.5644 | 0.00132 | 0.01035 | None |
| ENSG0( | -1.013 | -2.018 | 8.61E-07 | 2.21E-05 | LINC01579 |
| ENSG0( | -1.003 | -2.005 | 8.64E-05 | 0.0011 | TSPAN5-DT |
| ENSG0( | 1.679 | 3.2014 | 3.12E-06 | 6.60E-05 | GJA5 |
| ENSG0( | 1.039 | 2.0548 | 8.40E-06 | 0.00016 | None |
| ENSG0( | 2.27 | 4.8248 | 0.00017 | 0.00194 | None |
| ENSG0( | 1.158 | 2.2316 | 1.36E-09 | 8.06E-08 | None |

| GeneID | Treatment | Treatment | Treatment | Treatment | name | Astrocytes_FB1_FC-2_adj0.05_989 |
|---|---|---|---|---|---|---|
| ENSG0000 | -1.25783 | -2.39135 | 5.02E-05 | 0.000439 | SEMA3F | |
| ENSG0000 | -1.62011 | -3.07398 | 3.35E-30 | 3.51E-28 | ZMYND10 | |
| ENSG0000 | -1.16097 | -2.23607 | 0.000694 | 0.004464 | REXO5 | |
| ENSG0000 | -1.34532 | -2.54086 | 1.61E-22 | 1.12E-20 | PROM1 | |
| ENSG0000 | -1.0153 | -2.02133 | 0.010893 | 0.046196 | DNAH9 | |
| ENSG0000 | -1.04969 | -2.07008 | 1.40E-11 | 3.63E-10 | GAS7 | |
| ENSG0000 | -1.37246 | -2.58911 | 9.81E-14 | 3.21E-12 | DLEC1 | |
| ENSG0000 | -1.73381 | -3.32605 | 7.36E-09 | 1.32E-07 | IL32 | |
| ENSG0000 | -1.68152 | -3.20765 | 9.80E-05 | 0.000798 | ANLN | |
| ENSG0000 | -2.24047 | -4.72551 | 6.28E-46 | 1.23E-43 | GABRA3 | |
| ENSG0000 | -1.13299 | -2.19314 | 0.002042 | 0.011349 | TACC3 | |
| ENSG0000 | -2.04151 | -4.11676 | 6.34E-16 | 2.61E-14 | IGF1 | |
| ENSG0000 | -1.13696 | -2.19917 | 1.22E-17 | 5.75E-16 | PHLDB1 | |
| ENSG0000 | -1.02741 | -2.03837 | 0.002245 | 0.012313 | BIRC3 | |
| ENSG0000 | -1.63451 | -3.10482 | 1.42E-22 | 9.98E-21 | EFCAB1 | |
| ENSG0000 | -1.67932 | -3.20276 | 3.12E-35 | 4.31E-33 | CP | |
| ENSG0000 | -1.09453 | -2.13544 | 0.001975 | 0.01104 | ARHGAP6 | |
| ENSG0000 | -1.08895 | -2.1272 | 1.16E-05 | 0.000118 | LMO3 | |
| ENSG0000 | -1.13446 | -2.19536 | 5.87E-12 | 1.60E-10 | ELN | |
| ENSG0000 | -1.29139 | -2.44763 | 6.34E-26 | 5.25E-24 | MSMO1 | |
| ENSG0000 | -1.00232 | -2.00322 | 0.011504 | 0.048336 | CCDC85A | |
| ENSG0000 | -1.29948 | -2.46141 | 3.60E-12 | 1.00E-10 | NGFR | |
| ENSG0000 | -1.91386 | -3.76815 | 1.66E-31 | 1.86E-29 | CNN2 | |
| ENSG0000 | -1.01185 | -2.01649 | 4.77E-13 | 1.47E-11 | TRAM2 | |
| ENSG0000 | -1.00678 | -2.00942 | 3.41E-19 | 1.85E-17 | IDI1 | |
| ENSG0000 | -1.57323 | -2.9757 | 1.95E-11 | 4.97E-10 | PRR11 | |
| ENSG0000 | -1.70804 | -3.26717 | 1.35E-29 | 1.38E-27 | MAOB | |
| ENSG0000 | -1.01397 | -2.01947 | 1.27E-13 | 4.12E-12 | FSTL3 | |
| ENSG0000 | -1.2089 | -2.31161 | 7.64E-10 | 1.57E-08 | ST6GALNAC2 | |
| ENSG0000 | -1.41204 | -2.66114 | 7.41E-22 | 4.98E-20 | LMCD1 | |
| ENSG0000 | -2.17084 | -4.50285 | 1.47E-29 | 1.48E-27 | LIMS2 | |
| ENSG0000 | -1.18287 | -2.27029 | 2.04E-13 | 6.48E-12 | SREBF1 | |
| ENSG0000 | -1.98364 | -3.9549 | 1.14E-15 | 4.58E-14 | IRAG1 | |
| ENSG0000 | -1.15362 | -2.22471 | 7.28E-17 | 3.23E-15 | NTN4 | |
| ENSG0000 | -1.58652 | -3.00324 | 2.01E-29 | 2.02E-27 | FOSL2 | |
| ENSG0000 | -1.29803 | -2.45894 | 2.42E-11 | 6.08E-10 | CACNG4 | |
| ENSG0000 | -1.04017 | -2.05647 | 6.30E-16 | 2.60E-14 | ACTB | |
| ENSG0000 | -1.49937 | -2.82719 | 0.001343 | 0.007995 | SPAG5 | |
| ENSG0000 | -1.03831 | -2.05382 | 2.67E-16 | 1.13E-14 | MCAM | |
| ENSG0000 | -1.17151 | -2.25248 | 5.81E-11 | 1.39E-09 | PAK3 | |
| ENSG0000 | -5.20918 | -36.9931 | 0.002646 | 0.014205 | CAPN6 | |
| ENSG0000 | -1.39456 | -2.62909 | 3.21E-07 | 4.41E-06 | DCX | |
| ENSG0000 | -1.4979 | -2.82431 | 1.98E-07 | 2.83E-06 | EDN1 | |
| ENSG0000 | -1.34195 | -2.53493 | 1.02E-29 | 1.05E-27 | BRINP1 | |
| ENSG0000 | -1.39243 | -2.62521 | 1.57E-13 | 5.06E-12 | TNS1 | |
| ENSG0000 | -1.05111 | -2.07213 | 3.36E-11 | 8.30E-10 | COL5A3 | |
| ENSG0000 | -1.42777 | -2.6903 | 0.006632 | 0.030612 | CHRNA3 | |
| ENSG0000 | -1.38437 | -2.61058 | 4.32E-16 | 1.82E-14 | IGSF9B | |
| ENSG0000 | -1.18119 | -2.26763 | 1.18E-05 | 0.00012 | MPP4 | |

| | | | | | |
|---|---|---|---|---|---|
| ENSG0000 | -1.09274 | -2.13279 | 2.61E-06 | 3.02E-05 | C1QTNF3 |
| ENSG0000 | -1.10936 | -2.1575 | 2.13E-16 | 9.16E-15 | SEMA5B |
| ENSG0000 | -1.31115 | -2.48139 | 1.67E-10 | 3.77E-09 | TRPM3 |
| ENSG0000 | -1.51702 | -2.862 | 8.78E-05 | 0.000724 | SLCO1A2 |
| ENSG0000 | -1.2413 | -2.36411 | 1.15E-12 | 3.39E-11 | IGSF9 |
| ENSG0000 | -1.19794 | -2.29412 | 0.003598 | 0.018328 | FAT2 |
| ENSG0000 | -1.42447 | -2.68415 | 1.33E-23 | 9.94E-22 | ADAMTS2 |
| ENSG0000 | -1.60337 | -3.03852 | 5.11E-20 | 2.98E-18 | P2RX7 |
| ENSG0000 | -3.22683 | -9.36211 | 4.16E-05 | 0.00037 | LHX5 |
| ENSG0000 | -2.24481 | -4.73976 | 0.002723 | 0.014533 | RPH3A |
| ENSG0000 | -1.08606 | -2.12293 | 1.41E-15 | 5.55E-14 | P3H2 |
| ENSG0000 | -1.07856 | -2.11193 | 4.51E-15 | 1.71E-13 | PLEKHG2 |
| ENSG0000 | -1.87261 | -3.66195 | 0.001649 | 0.009494 | LAMB4 |
| ENSG0000 | -1.58162 | -2.99306 | 7.55E-27 | 6.47E-25 | LAMB1 |
| ENSG0000 | -1.22671 | -2.34033 | 0.000265 | 0.001925 | MYH7 |
| ENSG0000 | -1.00977 | -2.01359 | 7.68E-14 | 2.55E-12 | MYL6 |
| ENSG0000 | -1.25517 | -2.38696 | 3.66E-28 | 3.34E-26 | TEKT2 |
| ENSG0000 | -1.14316 | -2.20865 | 0.005151 | 0.02471 | CDC6 |
| ENSG0000 | -1.91498 | -3.77109 | 4.77E-18 | 2.33E-16 | FMO2 |
| ENSG0000 | -1.10876 | -2.1566 | 2.77E-05 | 0.000256 | HSD3B7 |
| ENSG0000 | -2.24533 | -4.74145 | 4.95E-32 | 5.63E-30 | GADD45B |
| ENSG0000 | -1.97588 | -3.93369 | 0.002497 | 0.013501 | DERL3 |
| ENSG0000 | -1.22175 | -2.33229 | 2.66E-06 | 3.08E-05 | GSTT2 |
| ENSG0000 | -1.07947 | -2.11326 | 6.18E-06 | 6.59E-05 | SUSD2 |
| ENSG0000 | -1.04637 | -2.06532 | 7.19E-07 | 9.27E-06 | GGT5 |
| ENSG0000 | -4.27208 | -19.3208 | 0.000891 | 0.005568 | PLA2G3 |
| ENSG0000 | -1.64458 | -3.12656 | 7.31E-12 | 1.97E-10 | LGALS1 |
| ENSG0000 | -1.20419 | -2.30408 | 6.50E-13 | 1.97E-11 | PIK3IP1 |
| ENSG0000 | -2.2389 | -4.72039 | 4.69E-05 | 0.000413 | SLC5A1 |
| ENSG0000 | -2.18646 | -4.55188 | 2.29E-20 | 1.37E-18 | PDGFB |
| ENSG0000 | -1.21543 | -2.3221 | 7.15E-05 | 0.000602 | PNPLA3 |
| ENSG0000 | -1.16205 | -2.23775 | 5.88E-18 | 2.84E-16 | MYH9 |
| ENSG0000 | -3.87836 | -14.7063 | 0.000785 | 0.004973 | PLEK2 |
| ENSG0000 | -1.95924 | -3.88856 | 0.001488 | 0.008713 | SLC8A3 |
| ENSG0000 | -1.38489 | -2.61153 | 9.27E-26 | 7.58E-24 | REC8 |
| ENSG0000 | -1.24939 | -2.3774 | 7.27E-07 | 9.37E-06 | NFATC2 |
| ENSG0000 | -1.10978 | -2.15812 | 1.18E-07 | 1.74E-06 | NKAIN4 |
| ENSG0000 | -1.30336 | -2.46804 | 4.12E-23 | 3.00E-21 | RASSF2 |
| ENSG0000 | -2.37713 | -5.19504 | 6.01E-40 | 9.73E-38 | MYL9 |
| ENSG0000 | -1.92056 | -3.7857 | 3.67E-16 | 1.55E-14 | E2F1 |
| ENSG0000 | -1.65136 | -3.1413 | 0.000492 | 0.003318 | FAM83D |
| ENSG0000 | -6.53106 | -92.4793 | 7.81E-06 | 8.17E-05 | EPPIN |
| ENSG0000 | -1.87175 | -3.65976 | 1.67E-35 | 2.33E-33 | CHRDL1 |
| ENSG0000 | -1.00662 | -2.00919 | 1.60E-08 | 2.74E-07 | MCF2 |
| ENSG0000 | -2.05652 | -4.15981 | 0.002937 | 0.015488 | RS1 |
| ENSG0000 | -1.02846 | -2.03984 | 2.37E-06 | 2.77E-05 | SRPX2 |
| ENSG0000 | -1.13739 | -2.19982 | 1.64E-27 | 1.46E-25 | SYTL4 |
| ENSG0000 | -1.89508 | -3.71942 | 7.41E-05 | 0.000622 | CENPI |
| ENSG0000 | -1.38184 | -2.60601 | 8.06E-10 | 1.65E-08 | CORO1A |
| ENSG0000 | -1.0521 | -2.07354 | 2.42E-11 | 6.08E-10 | MMP15 |

| | | | | | |
|---|---|---|---|---|---|
| ENSG0000 | -1.98416 | -3.95633 | 8.11E-18 | 3.85E-16 | CRISPLD2 |
| ENSG0000 | -1.53777 | -2.90345 | 9.98E-09 | 1.76E-07 | RASL12 |
| ENSG0000 | -1.38425 | -2.61036 | 0.002068 | 0.011473 | OIP5 |
| ENSG0000 | -1.35023 | -2.54953 | 1.81E-10 | 4.09E-09 | STMN2 |
| ENSG0000 | -1.16446 | -2.2415 | 0.000327 | 0.002318 | MCM4 |
| ENSG0000 | -1.08315 | -2.11866 | 0.000513 | 0.00344 | CD37 |
| ENSG0000 | -1.04321 | -2.0608 | 5.28E-12 | 1.44E-10 | IL27RA |
| ENSG0000 | -1.1005 | -2.14428 | 1.03E-12 | 3.06E-11 | PRX |
| ENSG0000 | -1.02004 | -2.02797 | 0.000126 | 0.001 | ODAD1 |
| ENSG0000 | -1.3727 | -2.58956 | 1.14E-12 | 3.38E-11 | SCN1B |
| ENSG0000 | -1.14781 | -2.21577 | 3.23E-13 | 1.01E-11 | TFPI2 |
| ENSG0000 | -1.01009 | -2.01403 | 8.18E-14 | 2.71E-12 | EPHB6 |
| ENSG0000 | -1.18456 | -2.27294 | 3.41E-12 | 9.54E-11 | NPTX2 |
| ENSG0000 | -1.37066 | -2.58589 | 0.004624 | 0.0226 | TFR2 |
| ENSG0000 | -2.45326 | -5.47654 | 1.36E-52 | 3.12E-50 | SFRP4 |
| ENSG0000 | -1.0459 | -2.06466 | 5.40E-21 | 3.36E-19 | RASSF4 |
| ENSG0000 | -1.25377 | -2.38464 | 1.48E-16 | 6.42E-15 | CXCL12 |
| ENSG0000 | -3.69778 | -12.976 | 2.60E-53 | 6.02E-51 | ACTA2 |
| ENSG0000 | -1.9499 | -3.86347 | 4.62E-25 | 3.65E-23 | CCL2 |
| ENSG0000 | -1.04323 | -2.06083 | 0.004429 | 0.021795 | PTGES3L-AARSD1 |
| ENSG0000 | -1.15071 | -2.22023 | 5.30E-18 | 2.58E-16 | TMEM97 |
| ENSG0000 | -1.174 | -2.25637 | 2.79E-22 | 1.92E-20 | ALDOC |
| ENSG0000 | -1.31721 | -2.49184 | 8.38E-17 | 3.69E-15 | BST1 |
| ENSG0000 | -1.31513 | -2.48826 | 5.74E-05 | 0.000494 | DDX25 |
| ENSG0000 | -1.53245 | -2.89277 | 1.37E-16 | 5.94E-15 | CRYAB |
| ENSG0000 | -1.82174 | -3.53508 | 0.001697 | 0.009722 | P2RX3 |
| ENSG0000 | -1.01022 | -2.01421 | 2.43E-10 | 5.40E-09 | FOLR1 |
| ENSG0000 | -1.01463 | -2.02038 | 0.002006 | 0.011183 | SELPLG |
| ENSG0000 | -1.18638 | -2.2758 | 5.26E-19 | 2.82E-17 | MVK |
| ENSG0000 | -1.52507 | -2.87801 | 0.000675 | 0.004365 | FOXM1 |
| ENSG0000 | -1.86951 | -3.6541 | 4.54E-06 | 5.00E-05 | OAS3 |
| ENSG0000 | -1.70521 | -3.26076 | 0.003064 | 0.016045 | FZD10 |
| ENSG0000 | -1.27568 | -2.42114 | 5.81E-07 | 7.64E-06 | CDCA3 |
| ENSG0000 | -6.37296 | -82.8803 | 0.00016 | 0.001234 | KLRB1 |
| ENSG0000 | -1.53346 | -2.8948 | 4.50E-09 | 8.28E-08 | COL12A1 |
| ENSG0000 | -1.46165 | -2.75423 | 6.83E-43 | 1.23E-40 | RSPH4A |
| ENSG0000 | -1.26349 | -2.40075 | 8.15E-13 | 2.45E-11 | MAK |
| ENSG0000 | -2.2912 | -4.89463 | 0.000135 | 0.001067 | ADTRP |
| ENSG0000 | -2.60885 | -6.10015 | 1.66E-85 | 7.05E-83 | MDGA1 |
| ENSG0000 | -2.7484 | -6.71971 | 1.10E-33 | 1.37E-31 | COL9A1 |
| ENSG0000 | -1.10017 | -2.1438 | 9.31E-21 | 5.68E-19 | C6orf118 |
| ENSG0000 | -2.21416 | -4.64011 | 9.73E-50 | 2.11E-47 | SMOC2 |
| ENSG0000 | -1.00754 | -2.01049 | 5.26E-17 | 2.36E-15 | CCND3 |
| ENSG0000 | -1.20028 | -2.29784 | 1.71E-27 | 1.51E-25 | SRF |
| ENSG0000 | -1.40844 | -2.6545 | 6.49E-28 | 5.85E-26 | GHR |
| ENSG0000 | -1.3606 | -2.56792 | 1.25E-25 | 1.01E-23 | HMGCS1 |
| ENSG0000 | -1.78789 | -3.45309 | 0.000276 | 0.001995 | KIF20A |
| ENSG0000 | -2.48833 | -5.61129 | 3.85E-63 | 1.09E-60 | HBEGF |
| ENSG0000 | -1.10719 | -2.15426 | 7.00E-20 | 4.05E-18 | HMGCR |
| ENSG0000 | -1.12234 | -2.17699 | 0.007246 | 0.03285 | LMNB1 |

| | | | | | |
|---|---|---|---|---|---|
| ENSG0000 | -1.58927 | -3.00896 | 1.92E-28 | 1.80E-26 | ST8SIA4 |
| ENSG0000 | -1.17832 | -2.26313 | 8.26E-12 | 2.21E-10 | PDGFRB |
| ENSG0000 | -3.03606 | -8.20248 | 5.97E-17 | 2.66E-15 | UNC5A |
| ENSG0000 | -1.99341 | -3.98176 | 2.74E-09 | 5.19E-08 | CNTN3 |
| ENSG0000 | -5.19769 | -36.6995 | 0.006913 | 0.031601 | CD86 |
| ENSG0000 | -1.64398 | -3.12528 | 2.67E-28 | 2.48E-26 | AMOTL2 |
| ENSG0000 | -1.1905 | -2.28232 | 0.000573 | 0.003783 | SERPINI2 |
| ENSG0000 | -1.95368 | -3.87362 | 4.50E-06 | 4.97E-05 | TNNC1 |
| ENSG0000 | -1.6698 | -3.1817 | 0.000121 | 0.000967 | POMC |
| ENSG0000 | -2.04591 | -4.12933 | 0.001822 | 0.010311 | OTOF |
| ENSG0000 | -1.47085 | -2.77186 | 1.36E-08 | 2.35E-07 | CENPA |
| ENSG0000 | -1.09631 | -2.13808 | 4.02E-05 | 0.000359 | TACR1 |
| ENSG0000 | -1.20475 | -2.30498 | 6.05E-07 | 7.94E-06 | EVA1A |
| ENSG0000 | -1.12272 | -2.17757 | 1.61E-07 | 2.34E-06 | DNAH6 |
| ENSG0000 | -3.38097 | -10.4177 | 2.75E-52 | 6.20E-50 | KCNJ13 |
| ENSG0000 | -1.20037 | -2.29798 | 4.34E-08 | 6.90E-07 | PASK |
| ENSG0000 | -1.23136 | -2.34789 | 3.38E-09 | 6.33E-08 | RMDN2 |
| ENSG0000 | -1.47135 | -2.77281 | 6.22E-45 | 1.19E-42 | EPHA4 |
| ENSG0000 | -1.65315 | -3.14519 | 3.24E-12 | 9.11E-11 | DHCR24 |
| ENSG0000 | -2.42344 | -5.36448 | 2.53E-39 | 4.07E-37 | RPE65 |
| ENSG0000 | -1.45556 | -2.74264 | 6.48E-11 | 1.54E-09 | KMO |
| ENSG0000 | -1.02975 | -2.04167 | 0.000101 | 0.000822 | RGS4 |
| ENSG0000 | -2.12046 | -4.34833 | 2.27E-29 | 2.26E-27 | GBP1 |
| ENSG0000 | -1.33604 | -2.52457 | 9.06E-49 | 1.88E-46 | PIK3R3 |
| ENSG0000 | -1.12347 | -2.1787 | 0.009179 | 0.040014 | CENPF |
| ENSG0000 | -2.33975 | -5.06215 | 0.009924 | 0.042697 | TREH |
| ENSG0000 | -1.95466 | -3.87624 | 0.001378 | 0.008158 | ZNF541 |
| ENSG0000 | -1.76809 | -3.40603 | 0.000741 | 0.00473 | KIF14 |
| ENSG0000 | -1.07221 | -2.10266 | 2.19E-17 | 1.01E-15 | KLF7 |
| ENSG0000 | -1.01446 | -2.02015 | 1.36E-09 | 2.69E-08 | DNAI7 |
| ENSG0000 | -1.34029 | -2.53202 | 7.43E-05 | 0.000623 | IRAG2 |
| ENSG0000 | -7.52219 | -183.825 | 3.36E-08 | 5.42E-07 | ATP10B |
| ENSG0000 | -1.8275 | -3.54922 | 1.18E-28 | 1.12E-26 | FILIP1 |
| ENSG0000 | -1.37563 | -2.59481 | 4.12E-21 | 2.60E-19 | CNR1 |
| ENSG0000 | -1.61849 | -3.07055 | 5.03E-28 | 4.55E-26 | SGIP1 |
| ENSG0000 | -2.46393 | -5.51719 | 0.000127 | 0.001009 | ADGB |
| ENSG0000 | -1.03995 | -2.05615 | 8.82E-11 | 2.07E-09 | SGK1 |
| ENSG0000 | -3.59701 | -12.1006 | 1.19E-44 | 2.24E-42 | CCN2 |
| ENSG0000 | -1.22427 | -2.33637 | 6.08E-56 | 1.47E-53 | TJP2 |
| ENSG0000 | -2.43423 | -5.40477 | 1.65E-109 | 1.04E-106 | NR4A3 |
| ENSG0000 | -1.61871 | -3.07101 | 1.20E-14 | 4.30E-13 | FLVCR2 |
| ENSG0000 | -1.05144 | -2.07259 | 4.85E-11 | 1.17E-09 | TGFB3 |
| ENSG0000 | -3.53768 | -11.6131 | 7.60E-69 | 2.44E-66 | TEK |
| ENSG0000 | -1.33599 | -2.52449 | 5.82E-26 | 4.83E-24 | CCDC170 |
| ENSG0000 | -1.97187 | -3.92276 | 0.007502 | 0.033785 | TNN |
| ENSG0000 | -1.37618 | -2.5958 | 7.75E-16 | 3.15E-14 | ACAT2 |
| ENSG0000 | -1.3663 | -2.57808 | 3.67E-12 | 1.02E-10 | EGR1 |
| ENSG0000 | -1.46773 | -2.76586 | 1.84E-07 | 2.63E-06 | ADRA1A |
| ENSG0000 | -1.40954 | -2.65653 | 1.61E-21 | 1.05E-19 | GLIPR2 |
| ENSG0000 | -1.52427 | -2.87641 | 1.90E-18 | 9.72E-17 | DNAI1 |

| | | | | | |
|---|---|---|---|---|---|
| ENSG0000 | -1.15122 | -2.22102 | 5.29E-17 | 2.37E-15 | CALD1 |
| ENSG0000 | -1.5583 | -2.94507 | 2.91E-06 | 3.33E-05 | CIT |
| ENSG0000 | -1.64988 | -3.13808 | 2.62E-05 | 0.000244 | CDKN2C |
| ENSG0000 | -1.55623 | -2.94084 | 0.000222 | 0.001647 | CENPK |
| ENSG0000 | -1.48492 | -2.79902 | 2.01E-12 | 5.77E-11 | MMP19 |
| ENSG0000 | -1.18792 | -2.27825 | 1.83E-19 | 1.01E-17 | NR4A1 |
| ENSG0000 | -1.55951 | -2.94753 | 0.000723 | 0.00464 | HJURP |
| ENSG0000 | -1.5751 | -2.97956 | 8.39E-35 | 1.12E-32 | PLP1 |
| ENSG0000 | -1.30009 | -2.46244 | 7.61E-08 | 1.16E-06 | MATN4 |
| ENSG0000 | -1.03469 | -2.04867 | 2.40E-08 | 4.00E-07 | COL21A1 |
| ENSG0000 | -1.0552 | -2.07801 | 7.28E-14 | 2.42E-12 | RUNX2 |
| ENSG0000 | -1.08781 | -2.12551 | 0.000108 | 0.000869 | AHNAK |
| ENSG0000 | -1.24335 | -2.36748 | 0.000107 | 0.000868 | WNT1 |
| ENSG0000 | -1.30052 | -2.46318 | 8.13E-05 | 0.000676 | BMP4 |
| ENSG0000 | -2.83366 | -7.12881 | 1.22E-05 | 0.000123 | MYH2 |
| ENSG0000 | -1.10693 | -2.15387 | 0.004129 | 0.020553 | FOSB |
| ENSG0000 | -1.06544 | -2.09281 | 4.44E-12 | 1.23E-10 | FLRT3 |
| ENSG0000 | -2.11323 | -4.3266 | 2.39E-14 | 8.35E-13 | LAMP5 |
| ENSG0000 | -1.83972 | -3.57941 | 1.36E-08 | 2.35E-07 | FAM110A |
| ENSG0000 | -1.89093 | -3.70874 | 4.33E-24 | 3.30E-22 | MMP24 |
| ENSG0000 | -1.18399 | -2.27204 | 6.27E-08 | 9.70E-07 | WNK4 |
| ENSG0000 | -1.15387 | -2.22511 | 7.71E-11 | 1.83E-09 | HSPA2 |
| ENSG0000 | -1.41545 | -2.66744 | 3.52E-26 | 2.93E-24 | PLEKHG3 |
| ENSG0000 | -1.05158 | -2.0728 | 4.21E-10 | 9.03E-09 | FGD3 |
| ENSG0000 | -1.28732 | -2.44075 | 2.24E-20 | 1.34E-18 | MASP1 |
| ENSG0000 | -1.65399 | -3.14704 | 0.000407 | 0.002803 | PKMYT1 |
| ENSG0000 | -1.12627 | -2.18294 | 1.94E-13 | 6.17E-12 | CHTF18 |
| ENSG0000 | -7.6007 | -194.106 | 4.22E-08 | 6.73E-07 | IL17B |
| ENSG0000 | -1.96818 | -3.91275 | 0.010515 | 0.044903 | YWHAH-AS1 |
| ENSG0000 | -2.88587 | -7.39149 | 2.87E-26 | 2.40E-24 | CPA4 |
| ENSG0000 | -1.26526 | -2.4037 | 2.00E-16 | 8.62E-15 | PODXL |
| ENSG0000 | -1.06844 | -2.09717 | 1.98E-20 | 1.19E-18 | CHN1 |
| ENSG0000 | -1.27158 | -2.41426 | 2.72E-19 | 1.49E-17 | ARHGAP22 |
| ENSG0000 | -1.15357 | -2.22464 | 7.18E-15 | 2.64E-13 | PALLD |
| ENSG0000 | -1.78903 | -3.45583 | 1.47E-28 | 1.39E-26 | KCNC1 |
| ENSG0000 | -2.18259 | -4.53968 | 4.65E-12 | 1.28E-10 | PIMREG |
| ENSG0000 | -1.3523 | -2.55319 | 6.77E-07 | 8.80E-06 | RNASE1 |
| ENSG0000 | -1.13285 | -2.19292 | 2.21E-23 | 1.65E-21 | FGF13 |
| ENSG0000 | -1.85383 | -3.61458 | 0.00127 | 0.007612 | TNNI3 |
| ENSG0000 | -1.5356 | -2.89909 | 4.99E-09 | 9.11E-08 | STARD8 |
| ENSG0000 | -2.49843 | -5.65069 | 5.46E-44 | 1.01E-41 | LDLR |
| ENSG0000 | -3.2975 | -9.83213 | 7.25E-58 | 1.88E-55 | CNN1 |
| ENSG0000 | -6.66972 | -101.809 | 0.000475 | 0.003214 | TNNT3 |
| ENSG0000 | -1.10944 | -2.15762 | 1.23E-13 | 3.98E-12 | COL5A1 |
| ENSG0000 | -1.24776 | -2.37473 | 1.34E-16 | 5.83E-15 | RFTN1 |
| ENSG0000 | -1.6668 | -3.1751 | 0.001053 | 0.006443 | TOP2A |
| ENSG0000 | -1.14165 | -2.20634 | 0.001914 | 0.010749 | MATN3 |
| ENSG0000 | -1.68704 | -3.21996 | 2.94E-34 | 3.76E-32 | MATN2 |
| ENSG0000 | -2.29908 | -4.92142 | 7.46E-06 | 7.84E-05 | KANK4 |
| ENSG0000 | -1.28011 | -2.42857 | 5.94E-18 | 2.86E-16 | MYH10 |

| | | | | | |
|---|---|---|---|---|---|
| ENSG0000 | -1.00627 | -2.00871 | 0.000521 | 0.003487 | EPSTI1 |
| ENSG0000 | -1.42174 | -2.67909 | 4.65E-11 | 1.13E-09 | TRPC4 |
| ENSG0000 | -1.97425 | -3.92924 | 1.95E-05 | 0.000188 | STOML3 |
| ENSG0000 | -1.30974 | -2.47896 | 2.15E-19 | 1.18E-17 | STARD13 |
| ENSG0000 | -2.32577 | -5.01334 | 3.29E-14 | 1.13E-12 | MYH11 |
| ENSG0000 | -2.02401 | -4.06712 | 2.18E-10 | 4.85E-09 | DYDC2 |
| ENSG0000 | -1.63285 | -3.10125 | 5.65E-25 | 4.44E-23 | LOXL2 |
| ENSG0000 | -1.23174 | -2.3485 | 0.000259 | 0.001886 | CCNB1 |
| ENSG0000 | -1.70811 | -3.26732 | 3.12E-15 | 1.20E-13 | TSPAN2 |
| ENSG0000 | -1.12639 | -2.18312 | 2.21E-05 | 0.00021 | SYT6 |
| ENSG0000 | -1.05312 | -2.07501 | 1.32E-22 | 9.31E-21 | PSRC1 |
| ENSG0000 | -1.36906 | -2.58303 | 7.06E-15 | 2.61E-13 | WNT2B |
| ENSG0000 | -1.76561 | -3.40017 | 8.77E-11 | 2.06E-09 | APLNR |
| ENSG0000 | -1.5749 | -2.97914 | 9.30E-21 | 5.68E-19 | FADS2 |
| ENSG0000 | -1.56259 | -2.95384 | 6.75E-21 | 4.17E-19 | PDGFRA |
| ENSG0000 | -2.0623 | -4.17652 | 3.92E-14 | 1.34E-12 | ADAMTS8 |
| ENSG0000 | -1.28754 | -2.44111 | 1.75E-30 | 1.88E-28 | NREP |
| ENSG0000 | -1.16975 | -2.24972 | 8.07E-17 | 3.56E-15 | ADAM19 |
| ENSG0000 | -1.68112 | -3.20677 | 0.007037 | 0.032081 | CD36 |
| ENSG0000 | -1.74231 | -3.34569 | 1.01E-12 | 3.00E-11 | STX11 |
| ENSG0000 | -2.45421 | -5.48012 | 0.000448 | 0.003048 | EGR4 |
| ENSG0000 | -1.24544 | -2.37091 | 2.84E-18 | 1.42E-16 | KCNK1 |
| ENSG0000 | -1.214 | -2.3198 | 3.22E-11 | 7.97E-10 | CKAP2 |
| ENSG0000 | -1.69475 | -3.23721 | 0.00164 | 0.009447 | BRIP1 |
| ENSG0000 | -1.73754 | -3.33465 | 2.26E-18 | 1.14E-16 | GALNT5 |
| ENSG0000 | -1.49356 | -2.81583 | 1.57E-18 | 8.13E-17 | WDR38 |
| ENSG0000 | -1.07327 | -2.1042 | 0.000342 | 0.002409 | CTSV |
| ENSG0000 | -1.42214 | -2.67984 | 3.19E-07 | 4.39E-06 | IL33 |
| ENSG0000 | -1.19345 | -2.28699 | 2.17E-21 | 1.39E-19 | SPAG8 |
| ENSG0000 | -1.89058 | -3.70785 | 5.37E-10 | 1.13E-08 | TCF19 |
| ENSG0000 | -1.20196 | -2.30052 | 2.42E-07 | 3.39E-06 | TTC29 |
| ENSG0000 | -1.42491 | -2.68497 | 1.08E-19 | 6.16E-18 | PI15 |
| ENSG0000 | -1.93884 | -3.83396 | 1.79E-26 | 1.51E-24 | THBS1 |
| ENSG0000 | -1.73837 | -3.33657 | 0.000611 | 0.004005 | NUSAP1 |
| ENSG0000 | -1.43997 | -2.71314 | 0.001723 | 0.009842 | KIF23 |
| ENSG0000 | -1.51404 | -2.85609 | 0.000197 | 0.001488 | KNL1 |
| ENSG0000 | -1.00698 | -2.0097 | 1.24E-15 | 4.94E-14 | CENPO |
| ENSG0000 | -1.39333 | -2.62684 | 5.95E-18 | 2.86E-16 | LOXL4 |
| ENSG0000 | -1.40704 | -2.65193 | 9.37E-05 | 0.000767 | KIF11 |
| ENSG0000 | -1.36219 | -2.57076 | 7.87E-17 | 3.47E-15 | DUSP5 |
| ENSG0000 | -1.04616 | -2.06503 | 7.41E-16 | 3.02E-14 | PLCE1 |
| ENSG0000 | -1.05159 | -2.07281 | 1.06E-21 | 7.04E-20 | BARD1 |
| ENSG0000 | -1.08167 | -2.11649 | 1.08E-21 | 7.14E-20 | MNS1 |
| ENSG0000 | -1.22282 | -2.33402 | 6.59E-13 | 2.00E-11 | SHROOM3 |
| ENSG0000 | -1.04985 | -2.07031 | 7.75E-09 | 1.39E-07 | PPFIA2 |
| ENSG0000 | -2.01988 | -4.0555 | 1.54E-34 | 2.03E-32 | GLIPR1 |
| ENSG0000 | -1.50926 | -2.84663 | 3.00E-17 | 1.38E-15 | LUM |
| ENSG0000 | -1.78893 | -3.45557 | 1.51E-15 | 5.90E-14 | ITGB7 |
| ENSG0000 | -1.17078 | -2.25134 | 0.002291 | 0.012527 | DIAPH3 |
| ENSG0000 | -1.61273 | -3.05829 | 0.000189 | 0.001433 | SRRM4 |

| | | | | | |
|---|---|---|---|---|---|
| ENSG0000 | -1.17437 | -2.25695 | 4.20E-15 | 1.60E-13 | ARMH4 |
| ENSG0000 | -1.23051 | -2.34649 | 3.21E-22 | 2.19E-20 | AK7 |
| ENSG0000 | -1.61899 | -3.07159 | 2.89E-22 | 1.98E-20 | FBLN5 |
| ENSG0000 | -1.68507 | -3.21556 | 7.25E-36 | 1.02E-33 | TPM1 |
| ENSG0000 | -2.10665 | -4.30691 | 4.33E-11 | 1.05E-09 | PIF1 |
| ENSG0000 | -1.0383 | -2.0538 | 4.83E-11 | 1.17E-09 | NTRK3 |
| ENSG0000 | -1.34869 | -2.54682 | 1.77E-18 | 9.08E-17 | ST8SIA2 |
| ENSG0000 | -1.52631 | -2.88048 | 0.000241 | 0.001773 | MCTP2 |
| ENSG0000 | -1.39038 | -2.62148 | 1.08E-18 | 5.67E-17 | TGFB1I1 |
| ENSG0000 | -1.30119 | -2.46433 | 5.77E-06 | 6.20E-05 | MYLK3 |
| ENSG0000 | -1.3019 | -2.46553 | 6.78E-21 | 4.18E-19 | KIFC3 |
| ENSG0000 | -1.38244 | -2.6071 | 1.76E-17 | 8.18E-16 | LRRC46 |
| ENSG0000 | -1.21879 | -2.32751 | 4.59E-17 | 2.08E-15 | ASXL3 |
| ENSG0000 | -3.89876 | -14.9157 | 2.30E-10 | 5.10E-09 | SLC14A1 |
| ENSG0000 | -1.38732 | -2.61592 | 7.48E-21 | 4.60E-19 | MAPK4 |
| ENSG0000 | -2.335 | -5.04552 | 1.73E-05 | 0.000169 | CBLN2 |
| ENSG0000 | -1.71676 | -3.28698 | 0.000309 | 0.002207 | STAC2 |
| ENSG0000 | -2.77816 | -6.85979 | 1.31E-05 | 0.000131 | FBN3 |
| ENSG0000 | -2.17605 | -4.51915 | 0.007492 | 0.033749 | SIGLEC10 |
| ENSG0000 | -1.79796 | -3.47728 | 1.33E-15 | 5.28E-14 | CFAP74 |
| ENSG0000 | -1.12005 | -2.17355 | 2.03E-15 | 7.91E-14 | FHAD1 |
| ENSG0000 | -2.96714 | -7.81986 | 1.91E-61 | 5.24E-59 | CCN1 |
| ENSG0000 | -1.00253 | -2.00351 | 0.000699 | 0.004495 | BEST4 |
| ENSG0000 | -1.41365 | -2.6641 | 3.20E-10 | 6.97E-09 | MAEL |
| ENSG0000 | -1.28911 | -2.44377 | 1.53E-14 | 5.47E-13 | ILDR2 |
| ENSG0000 | -2.19894 | -4.59143 | 1.28E-65 | 3.85E-63 | RGS5 |
| ENSG0000 | -1.40774 | -2.65321 | 6.80E-25 | 5.32E-23 | TUFT1 |
| ENSG0000 | -1.02065 | -2.02884 | 8.22E-07 | 1.05E-05 | VASH2 |
| ENSG0000 | -1.43512 | -2.70405 | 6.29E-15 | 2.34E-13 | FBLN7 |
| ENSG0000 | -3.19816 | -9.17784 | 2.80E-08 | 4.59E-07 | GPR17 |
| ENSG0000 | -6.78893 | -110.579 | 5.26E-06 | 5.71E-05 | NYAP2 |
| ENSG0000 | -1.1078 | -2.15516 | 1.78E-32 | 2.11E-30 | TMEM108 |
| ENSG0000 | -1.47119 | -2.7725 | 2.40E-16 | 1.03E-14 | ROPN1L |
| ENSG0000 | -1.3717 | -2.58775 | 2.53E-16 | 1.08E-14 | ADAMTS16 |
| ENSG0000 | -1.31731 | -2.49201 | 7.72E-22 | 5.16E-20 | LIX1 |
| ENSG0000 | -2.56819 | -5.93064 | 2.26E-42 | 3.98E-40 | CXCL14 |
| ENSG0000 | -2.21616 | -4.64656 | 2.96E-08 | 4.83E-07 | C1QTNF2 |
| ENSG0000 | -2.30341 | -4.93622 | 2.23E-14 | 7.82E-13 | GABRB2 |
| ENSG0000 | -2.57606 | -5.96307 | 2.75E-22 | 1.89E-20 | KCNMB1 |
| ENSG0000 | -1.13576 | -2.19734 | 4.65E-09 | 8.53E-08 | DCDC2 |
| ENSG0000 | -1.65622 | -3.1519 | 1.56E-05 | 0.000154 | MLIP |
| ENSG0000 | -1.44168 | -2.71638 | 4.06E-15 | 1.55E-13 | SCUBE3 |
| ENSG0000 | -1.52435 | -2.87656 | 0.002804 | 0.014912 | CRIP3 |
| ENSG0000 | -1.02073 | -2.02894 | 1.21E-15 | 4.83E-14 | PRSS35 |
| ENSG0000 | -1.32162 | -2.49947 | 0.000661 | 0.004285 | RSPO3 |
| ENSG0000 | -2.10023 | -4.28779 | 2.19E-34 | 2.84E-32 | IGFBP3 |
| ENSG0000 | -1.81859 | -3.52736 | 1.54E-09 | 3.02E-08 | EPHA1 |
| ENSG0000 | -1.02813 | -2.03938 | 5.22E-16 | 2.16E-14 | NLGN4X |
| ENSG0000 | -2.0156 | -4.04347 | 3.53E-16 | 1.49E-14 | RIPPLY1 |
| ENSG0000 | -2.47288 | -5.55151 | 0.000519 | 0.003474 | ST18 |

| | | | | | |
|---|---|---|---|---|---|
| ENSG0000 | -1.4519 | -2.73568 | 2.12E-12 | 6.08E-11 | GINS4 |
| ENSG0000 | -1.30638 | -2.4732 | 3.23E-23 | 2.39E-21 | AOPEP |
| ENSG0000 | -1.79102 | -3.46059 | 0.00145 | 0.008522 | MKI67 |
| ENSG0000 | -1.43692 | -2.70743 | 2.05E-11 | 5.20E-10 | INA |
| ENSG0000 | -1.1488 | -2.2173 | 6.82E-29 | 6.59E-27 | CNNM2 |
| ENSG0000 | -1.31571 | -2.48925 | 4.50E-06 | 4.96E-05 | KLHL35 |
| ENSG0000 | -3.74193 | -13.3793 | 5.69E-45 | 1.10E-42 | TAGLN |
| ENSG0000 | -2.71013 | -6.54382 | 1.07E-41 | 1.82E-39 | JPH2 |
| ENSG0000 | -1.55643 | -2.94124 | 4.85E-06 | 5.31E-05 | KIAA1755 |
| ENSG0000 | -3.20744 | -9.23709 | 0.002735 | 0.014581 | OCSTAMP |
| ENSG0000 | -1.18378 | -2.27172 | 1.22E-18 | 6.40E-17 | ADGRL3 |
| ENSG0000 | -1.24327 | -2.36735 | 3.60E-23 | 2.63E-21 | FAM124A |
| ENSG0000 | -1.09691 | -2.13896 | 3.72E-20 | 2.19E-18 | LYPD1 |
| ENSG0000 | -1.62973 | -3.09456 | 1.15E-19 | 6.51E-18 | C2orf50 |
| ENSG0000 | -1.43798 | -2.70941 | 1.73E-13 | 5.54E-12 | ADAMTS12 |
| ENSG0000 | -1.51202 | -2.8521 | 2.65E-22 | 1.83E-20 | TMEM163 |
| ENSG0000 | -1.35134 | -2.55149 | 7.42E-05 | 0.000622 | CAPSL |
| ENSG0000 | -1.47295 | -2.77589 | 4.81E-32 | 5.49E-30 | DYNLT5 |
| ENSG0000 | -1.71021 | -3.2721 | 0.000415 | 0.002849 | LMNTD1 |
| ENSG0000 | -1.52023 | -2.86837 | 0.000792 | 0.005012 | NRSN1 |
| ENSG0000 | -1.56269 | -2.95403 | 1.34E-10 | 3.05E-09 | JAKMIP1 |
| ENSG0000 | -1.15942 | -2.23368 | 1.35E-11 | 3.52E-10 | RASSF3 |
| ENSG0000 | -1.0398 | -2.05594 | 0.004931 | 0.02384 | CIBAR2 |
| ENSG0000 | -1.09995 | -2.14347 | 2.63E-17 | 1.21E-15 | DDAH1 |
| ENSG0000 | -1.38929 | -2.6195 | 2.87E-19 | 1.57E-17 | CHST9 |
| ENSG0000 | -1.53066 | -2.88918 | 5.50E-34 | 6.91E-32 | THY1 |
| ENSG0000 | -1.43817 | -2.70978 | 1.08E-19 | 6.14E-18 | DNAAF1 |
| ENSG0000 | -1.67372 | -3.19035 | 1.10E-13 | 3.57E-12 | NRGN |
| ENSG0000 | -1.18636 | -2.27577 | 1.21E-08 | 2.11E-07 | ABI3BP |
| ENSG0000 | -1.43068 | -2.69574 | 6.72E-23 | 4.79E-21 | SORBS2 |
| ENSG0000 | -1.11353 | -2.16374 | 0.011794 | 0.049295 | CEP170P1 |
| ENSG0000 | -2.31289 | -4.96879 | 8.53E-44 | 1.56E-41 | CHODL |
| ENSG0000 | -1.92759 | -3.80419 | 0.000213 | 0.001591 | TMPRSS15 |
| ENSG0000 | -1.21808 | -2.32636 | 2.47E-09 | 4.71E-08 | SLFN13 |
| ENSG0000 | -1.45213 | -2.73612 | 7.65E-18 | 3.64E-16 | USP43 |
| ENSG0000 | -1.60728 | -3.04678 | 1.00E-12 | 2.99E-11 | DKK2 |
| ENSG0000 | -1.03123 | -2.04376 | 1.85E-10 | 4.18E-09 | PTPRN2 |
| ENSG0000 | -1.41404 | -2.66482 | 4.84E-24 | 3.68E-22 | KCNMA1 |
| ENSG0000 | -1.71378 | -3.28018 | 1.07E-06 | 1.33E-05 | CFAP161 |
| ENSG0000 | -1.42723 | -2.6893 | 1.35E-11 | 3.52E-10 | MAP3K7CL |
| ENSG0000 | -1.9012 | -3.73524 | 1.87E-27 | 1.65E-25 | GDF6 |
| ENSG0000 | -1.5521 | -2.93244 | 0.000348 | 0.002442 | FBXO43 |
| ENSG0000 | -1.66812 | -3.178 | 0.000222 | 0.001644 | GLYATL2 |
| ENSG0000 | -2.32159 | -4.99883 | 3.61E-79 | 1.40E-76 | FBXO32 |
| ENSG0000 | -1.36672 | -2.57883 | 0.008986 | 0.039289 | BUB1B |
| ENSG0000 | -2.36951 | -5.16765 | 2.97E-10 | 6.49E-09 | SST |
| ENSG0000 | -1.49428 | -2.81723 | 0.0006 | 0.003941 | CLEC18C |
| ENSG0000 | -1.36347 | -2.57304 | 1.99E-50 | 4.41E-48 | KIT |
| ENSG0000 | -1.29564 | -2.45486 | 0.004694 | 0.022887 | CCNB2 |
| ENSG0000 | -1.71503 | -3.28303 | 4.73E-10 | 1.00E-08 | KCNJ6 |

| | | | | | |
|---|---|---|---|---|---|
| ENSG0000 | -1.9452 | -3.8509 | 0.006114 | 0.028624 | ACAN |
| ENSG0000 | -1.28354 | -2.43436 | 1.32E-20 | 7.99E-19 | DZIP1L |
| ENSG0000 | -1.50926 | -2.84663 | 1.12E-13 | 3.65E-12 | TMSB15A |
| ENSG0000 | -2.65359 | -6.29231 | 1.25E-46 | 2.49E-44 | TENT5B |
| ENSG0000 | -1.22004 | -2.32953 | 5.00E-14 | 1.69E-12 | CLSTN2 |
| ENSG0000 | -1.63873 | -3.11392 | 1.42E-36 | 2.07E-34 | COLEC12 |
| ENSG0000 | -1.22592 | -2.33904 | 1.99E-18 | 1.01E-16 | RIBC1 |
| ENSG0000 | -2.06712 | -4.19051 | 4.50E-37 | 6.82E-35 | GDPD5 |
| ENSG0000 | -1.05855 | -2.08284 | 5.55E-10 | 1.16E-08 | DYNC1I1 |
| ENSG0000 | -1.97789 | -3.93917 | 4.27E-08 | 6.79E-07 | VWA5B1 |
| ENSG0000 | -1.14273 | -2.20799 | 0.005906 | 0.027732 | ADAMTS4 |
| ENSG0000 | -4.66193 | -25.3151 | 0.000522 | 0.003491 | TNNI1 |
| ENSG0000 | -1.4317 | -2.69765 | 1.20E-21 | 7.91E-20 | CSRP1 |
| ENSG0000 | -4.05984 | -16.6775 | 8.48E-33 | 1.03E-30 | ACTC1 |
| ENSG0000 | -1.39674 | -2.63306 | 2.80E-06 | 3.22E-05 | CELF3 |
| ENSG0000 | -1.45791 | -2.74711 | 3.33E-28 | 3.05E-26 | ALDH4A1 |
| ENSG0000 | -1.43382 | -2.70162 | 3.37E-23 | 2.47E-21 | DRC7 |
| ENSG0000 | -1.52581 | -2.87948 | 9.52E-19 | 5.02E-17 | TPPP3 |
| ENSG0000 | -1.01603 | -2.02234 | 8.38E-14 | 2.76E-12 | ZYX |
| ENSG0000 | -1.26522 | -2.40363 | 0.001694 | 0.009714 | GAB3 |
| ENSG0000 | -1.10841 | -2.15609 | 2.87E-07 | 3.98E-06 | C21orf58 |
| ENSG0000 | -1.09645 | -2.13828 | 1.11E-14 | 4.00E-13 | S100B |
| ENSG0000 | -1.19055 | -2.28239 | 5.12E-12 | 1.40E-10 | SLC2A6 |
| ENSG0000 | -1.53573 | -2.89936 | 8.22E-35 | 1.10E-32 | CFAP157 |
| ENSG0000 | -1.58759 | -3.00548 | 2.71E-28 | 2.51E-26 | PCSK7 |
| ENSG0000 | -1.91449 | -3.76981 | 5.04E-08 | 7.93E-07 | LRRC71 |
| ENSG0000 | -1.07871 | -2.11215 | 0.00107 | 0.006536 | RECQL4 |
| ENSG0000 | -2.97469 | -7.86088 | 1.26E-61 | 3.49E-59 | COL26A1 |
| ENSG0000 | -1.34051 | -2.53241 | 0.000561 | 0.003719 | RACGAP1 |
| ENSG0000 | -1.63058 | -3.09638 | 0.000558 | 0.003707 | SPC24 |
| ENSG0000 | -1.16161 | -2.23708 | 6.16E-39 | 9.79E-37 | PAQR4 |
| ENSG0000 | -1.98871 | -3.96883 | 1.36E-11 | 3.55E-10 | SHANK2 |
| ENSG0000 | -1.83335 | -3.56363 | 0.000629 | 0.004108 | ELAVL4 |
| ENSG0000 | -3.93223 | -15.2657 | 0.005582 | 0.026477 | KNCN |
| ENSG0000 | -1.17866 | -2.26366 | 9.58E-12 | 2.53E-10 | SYNC |
| ENSG0000 | -1.26942 | -2.41065 | 1.18E-05 | 0.000119 | UBXN10 |
| ENSG0000 | -1.24188 | -2.36507 | 4.56E-05 | 0.000403 | ALPL |
| ENSG0000 | -1.33319 | -2.51959 | 5.79E-06 | 6.22E-05 | MEGF6 |
| ENSG0000 | -2.18807 | -4.55695 | 6.80E-43 | 1.23E-40 | NEXN |
| ENSG0000 | -1.86413 | -3.64047 | 2.41E-05 | 0.000227 | B3GALT2 |
| ENSG0000 | -1.10781 | -2.15519 | 0.000335 | 0.002365 | GBP2 |
| ENSG0000 | -1.8754 | -3.66904 | 5.11E-25 | 4.02E-23 | OLFML2B |
| ENSG0000 | -5.52958 | -46.1924 | 0.003978 | 0.019923 | LMX1A |
| ENSG0000 | -1.53715 | -2.90221 | 5.28E-22 | 3.57E-20 | KLHDC8A |
| ENSG0000 | -1.16539 | -2.24295 | 7.51E-06 | 7.89E-05 | KCNF1 |
| ENSG0000 | -1.90399 | -3.74246 | 0.003647 | 0.018517 | ACTG2 |
| ENSG0000 | -3.74012 | -13.3625 | 0.007879 | 0.035258 | TEKT4 |
| ENSG0000 | -1.12231 | -2.17696 | 0.005268 | 0.025206 | CFAP141 |
| ENSG0000 | -1.61194 | -3.05662 | 0.005561 | 0.026393 | GABRG1 |
| ENSG0000 | -1.02535 | -2.03545 | 1.17E-09 | 2.34E-08 | INAVA |

| | | | | | |
|---|---|---|---|---|---|
| ENSG0000 | -3.33951 | -10.1226 | 1.11E-73 | 3.91E-71 | LMOD1 |
| ENSG0000 | -1.04471 | -2.06295 | 1.60E-19 | 8.93E-18 | TGFBR2 |
| ENSG0000 | -1.21679 | -2.32428 | 6.32E-05 | 0.000539 | SGO2 |
| ENSG0000 | -1.30182 | -2.4654 | 2.14E-06 | 2.52E-05 | EFHB |
| ENSG0000 | -1.42662 | -2.68815 | 3.26E-14 | 1.12E-12 | CDS1 |
| ENSG0000 | -1.15845 | -2.23218 | 2.08E-21 | 1.35E-19 | ADAMTS9 |
| ENSG0000 | -1.52613 | -2.88012 | 3.46E-17 | 1.58E-15 | CFAP100 |
| ENSG0000 | -2.2075 | -4.61875 | 1.39E-05 | 0.000139 | CLDN19 |
| ENSG0000 | -1.58548 | -3.00108 | 0.004332 | 0.021413 | GRM2 |
| ENSG0000 | -1.74374 | -3.34902 | 0.011024 | 0.046638 | NPY1R |
| ENSG0000 | -1.09469 | -2.13567 | 1.78E-09 | 3.47E-08 | ZNF474 |
| ENSG0000 | -1.46023 | -2.75153 | 3.42E-17 | 1.56E-15 | STARD4 |
| ENSG0000 | -1.57377 | -2.97682 | 2.97E-30 | 3.15E-28 | ANKRD33B |
| ENSG0000 | -1.26194 | -2.39818 | 6.43E-05 | 0.000549 | CMYA5 |
| ENSG0000 | -3.64302 | -12.4927 | 0.001267 | 0.007599 | DACT2 |
| ENSG0000 | -1.27103 | -2.41334 | 5.69E-06 | 6.13E-05 | PI16 |
| ENSG0000 | -1.15767 | -2.23097 | 1.07E-09 | 2.14E-08 | SYTL3 |
| ENSG0000 | -1.84091 | -3.58235 | 1.98E-18 | 1.01E-16 | FNDC1 |
| ENSG0000 | -1.6576 | -3.15491 | 9.18E-41 | 1.52E-38 | C7orf57 |
| ENSG0000 | -1.53196 | -2.89178 | 7.05E-14 | 2.36E-12 | PHKG1 |
| ENSG0000 | -1.16791 | -2.24686 | 0.00152 | 0.008864 | CSMD3 |
| ENSG0000 | -3.44167 | -10.8654 | 0.000737 | 0.004709 | UNCX |
| ENSG0000 | -1.48412 | -2.79747 | 1.18E-17 | 5.56E-16 | CA3 |
| ENSG0000 | -1.53105 | -2.88996 | 3.64E-07 | 4.97E-06 | OSR2 |
| ENSG0000 | -1.12886 | -2.18686 | 0.000454 | 0.003087 | C9orf24 |
| ENSG0000 | -3.43575 | -10.8209 | 1.66E-44 | 3.11E-42 | MAMDC2 |
| ENSG0000 | -1.66341 | -3.16765 | 4.71E-18 | 2.30E-16 | SVEP1 |
| ENSG0000 | -1.75681 | -3.3795 | 1.76E-11 | 4.53E-10 | CFAP47 |
| ENSG0000 | -1.91992 | -3.78402 | 2.10E-05 | 0.000201 | ZNF367 |
| ENSG0000 | -2.2864 | -4.87838 | 6.08E-27 | 5.24E-25 | AQP3 |
| ENSG0000 | -1.53317 | -2.89422 | 0.003426 | 0.017606 | DEUP1 |
| ENSG0000 | -1.26162 | -2.39765 | 4.56E-08 | 7.24E-07 | LRFN5 |
| ENSG0000 | -1.50533 | -2.8389 | 4.23E-20 | 2.47E-18 | GJB2 |
| ENSG0000 | -1.04654 | -2.06557 | 4.28E-12 | 1.18E-10 | AMER2 |
| ENSG0000 | -1.03257 | -2.04566 | 3.68E-18 | 1.82E-16 | STOX1 |
| ENSG0000 | -1.11328 | -2.16337 | 6.67E-18 | 3.18E-16 | NDRG2 |
| ENSG0000 | -1.06635 | -2.09412 | 1.40E-15 | 5.50E-14 | HSPA12A |
| ENSG0000 | -1.32758 | -2.50981 | 0.00025 | 0.00183 | PDZRN4 |
| ENSG0000 | -1.11819 | -2.17075 | 7.82E-17 | 3.46E-15 | HACD1 |
| ENSG0000 | -1.70475 | -3.25973 | 2.58E-09 | 4.91E-08 | RRAD |
| ENSG0000 | -1.47278 | -2.77557 | 1.29E-22 | 9.16E-21 | CFAP52 |
| ENSG0000 | -1.67944 | -3.20303 | 0.000602 | 0.003952 | MMP10 |
| ENSG0000 | -2.70514 | -6.52122 | 6.02E-06 | 6.45E-05 | PCLAF |
| ENSG0000 | -1.10264 | -2.14747 | 0.008502 | 0.037519 | GLYATL1 |
| ENSG0000 | -1.20205 | -2.30066 | 3.59E-09 | 6.70E-08 | C18orf54 |
| ENSG0000 | -2.14701 | -4.42908 | 1.36E-06 | 1.67E-05 | GREM1 |
| ENSG0000 | -1.33316 | -2.51954 | 8.02E-08 | 1.22E-06 | IGF2 |
| ENSG0000 | -1.05003 | -2.07057 | 3.72E-20 | 2.19E-18 | PRRT2 |
| ENSG0000 | -1.15779 | -2.23115 | 2.86E-21 | 1.83E-19 | TPM4 |
| ENSG0000 | -1.04812 | -2.06784 | 7.93E-19 | 4.22E-17 | MVD |

| | | | | | |
|---|---|---|---|---|---|
| ENSG0000 | -1.83848 | -3.57634 | 0.007448 | 0.033588 | PPP1R14A |
| ENSG0000 | -1.01827 | -2.02549 | 7.93E-13 | 2.39E-11 | DNAAF3 |
| ENSG0000 | -1.69286 | -3.23298 | 1.10E-07 | 1.64E-06 | ATCAY |
| ENSG0000 | -1.12856 | -2.18641 | 3.76E-10 | 8.10E-09 | SLC43A2 |
| ENSG0000 | -2.58479 | -5.9993 | 2.11E-05 | 0.000202 | KLK5 |
| ENSG0000 | -1.33858 | -2.52902 | 0.000506 | 0.003402 | KRT80 |
| ENSG0000 | -1.73782 | -3.3353 | 1.54E-19 | 8.59E-18 | TEKT1 |
| ENSG0000 | -1.29868 | -2.46003 | 2.85E-14 | 9.86E-13 | SCARA3 |
| ENSG0000 | -1.62822 | -3.09132 | 2.70E-13 | 8.49E-12 | FILIP1L |
| ENSG0000 | -1.78712 | -3.45126 | 8.09E-28 | 7.26E-26 | MFSD2A |
| ENSG0000 | -6.3764 | -83.0784 | 0.000153 | 0.00119 | MYL1 |
| ENSG0000 | -1.52925 | -2.88636 | 0.000111 | 0.000892 | FAM178B |
| ENSG0000 | -1.02904 | -2.04067 | 1.84E-10 | 4.16E-09 | NSG1 |
| ENSG0000 | -1.05753 | -2.08136 | 6.72E-07 | 8.75E-06 | STXBP6 |
| ENSG0000 | -1.07746 | -2.11032 | 2.73E-15 | 1.05E-13 | IRS1 |
| ENSG0000 | -3.38226 | -10.4271 | 1.69E-08 | 2.88E-07 | PCSK9 |
| ENSG0000 | -1.20207 | -2.3007 | 9.50E-05 | 0.000777 | RSPO1 |
| ENSG0000 | -3.42434 | -10.7357 | 0.004063 | 0.020277 | CXCL10 |
| ENSG0000 | -2.00408 | -4.01134 | 3.23E-10 | 7.02E-09 | SH3TC2 |
| ENSG0000 | -1.37456 | -2.59289 | 0.001585 | 0.009167 | ADRB2 |
| ENSG0000 | -1.37238 | -2.58898 | 0.00874 | 0.038383 | RSPH10B2 |
| ENSG0000 | -1.27819 | -2.42535 | 3.45E-07 | 4.72E-06 | KCNK9 |
| ENSG0000 | -3.27169 | -9.65776 | 5.11E-06 | 5.58E-05 | GPR183 |
| ENSG0000 | -1.03017 | -2.04227 | 1.18E-25 | 9.55E-24 | LUZP1 |
| ENSG0000 | -1.4443 | -2.72131 | 3.49E-27 | 3.05E-25 | IFFO2 |
| ENSG0000 | -1.0922 | -2.13199 | 3.46E-12 | 9.68E-11 | MYRIP |
| ENSG0000 | -1.09124 | -2.13057 | 8.65E-10 | 1.76E-08 | GPR37L1 |
| ENSG0000 | -1.34433 | -2.53913 | 0.006847 | 0.031347 | CDK1 |
| ENSG0000 | -2.01725 | -4.04812 | 8.52E-10 | 1.74E-08 | FRMPD2 |
| ENSG0000 | -1.5456 | -2.91926 | 0.000156 | 0.001209 | TCAF2 |
| ENSG0000 | -5.80109 | -55.7574 | 0.00031 | 0.00221 | DYDC1 |
| ENSG0000 | -1.11403 | -2.16449 | 7.77E-06 | 8.13E-05 | CYTL1 |
| ENSG0000 | -1.33061 | -2.51508 | 1.16E-18 | 6.07E-17 | PDGFD |
| ENSG0000 | -1.0639 | -2.09057 | 1.53E-12 | 4.47E-11 | S1PR1 |
| ENSG0000 | -1.78454 | -3.44509 | 0.000133 | 0.001056 | SHCBP1 |
| ENSG0000 | -1.5892 | -3.00882 | 2.09E-09 | 4.04E-08 | KCNK3 |
| ENSG0000 | -2.05875 | -4.16627 | 0.00022 | 0.001635 | ESCO2 |
| ENSG0000 | -1.82911 | -3.55317 | 3.88E-07 | 5.26E-06 | PTGER4 |
| ENSG0000 | -2.1794 | -4.52966 | 0.000829 | 0.005219 | NEUROD2 |
| ENSG0000 | -2.3617 | -5.13975 | 0.003183 | 0.016541 | FGB |
| ENSG0000 | -1.42857 | -2.69179 | 1.42E-24 | 1.11E-22 | ENC1 |
| ENSG0000 | -1.06188 | -2.08765 | 6.61E-18 | 3.16E-16 | PLEKHG5 |
| ENSG0000 | -1.57049 | -2.97006 | 3.63E-29 | 3.54E-27 | VAT1L |
| ENSG0000 | -1.0805 | -2.11477 | 9.23E-20 | 5.28E-18 | CFAP46 |
| ENSG0000 | -2.49648 | -5.64307 | 2.41E-08 | 4.01E-07 | RRM2 |
| ENSG0000 | -2.09055 | -4.25911 | 0.000627 | 0.004097 | LRRC15 |
| ENSG0000 | -1.09688 | -2.13892 | 1.31E-15 | 5.20E-14 | SLFN12 |
| ENSG0000 | -1.32229 | -2.50062 | 1.03E-15 | 4.14E-14 | FRMD3 |
| ENSG0000 | -1.10968 | -2.15797 | 1.25E-10 | 2.87E-09 | NEGR1 |
| ENSG0000 | -1.07498 | -2.10669 | 9.50E-10 | 1.92E-08 | SPTLC3 |

| | | | | | |
|---|---|---|---|---|---|
| ENSG0000 | -1.9699 | -3.91742 | 0.000223 | 0.001651 | IL16 |
| ENSG0000 | -1.39583 | -2.63139 | 3.79E-18 | 1.88E-16 | SYNPO2 |
| ENSG0000 | -1.4958 | -2.82021 | 9.68E-38 | 1.49E-35 | EFCAB12 |
| ENSG0000 | -1.11264 | -2.16241 | 1.28E-10 | 2.93E-09 | DHCR7 |
| ENSG0000 | -1.97399 | -3.92854 | 4.72E-13 | 1.45E-11 | MRGPRF |
| ENSG0000 | -1.56917 | -2.96734 | 0.000135 | 0.001069 | HPSE2 |
| ENSG0000 | -1.2597 | -2.39445 | 2.49E-26 | 2.09E-24 | ABCD2 |
| ENSG0000 | -1.33144 | -2.51653 | 1.88E-10 | 4.22E-09 | SYT12 |
| ENSG0000 | -1.11972 | -2.17304 | 8.31E-14 | 2.74E-12 | HEG1 |
| ENSG0000 | -1.54005 | -2.90804 | 5.25E-07 | 6.98E-06 | CCDC13-AS1 |
| ENSG0000 | -1.32044 | -2.49743 | 9.20E-14 | 3.02E-12 | C1QTNF1 |
| ENSG0000 | -1.13317 | -2.1934 | 7.38E-22 | 4.97E-20 | MSRB3 |
| ENSG0000 | -2.18637 | -4.55158 | 8.87E-09 | 1.58E-07 | MIR1-1HG-AS1 |
| ENSG0000 | -2.95446 | -7.75144 | 1.67E-10 | 3.78E-09 | NPAS4 |
| ENSG0000 | -2.83546 | -7.13772 | 2.49E-23 | 1.85E-21 | SLCO2A1 |
| ENSG0000 | -1.16219 | -2.23796 | 0.003053 | 0.016002 | CD248 |
| ENSG0000 | -1.17283 | -2.25454 | 9.25E-12 | 2.45E-10 | CNIH2 |
| ENSG0000 | -1.75042 | -3.36458 | 5.17E-05 | 0.000451 | UBE2C |
| ENSG0000 | -1.03494 | -2.04903 | 5.37E-21 | 3.35E-19 | CSRP2 |
| ENSG0000 | -1.64229 | -3.12161 | 3.28E-23 | 2.42E-21 | GAL3ST3 |
| ENSG0000 | -1.42097 | -2.67766 | 7.55E-20 | 4.36E-18 | VWA3A |
| ENSG0000 | -1.67551 | -3.19431 | 1.30E-09 | 2.59E-08 | CCNE2 |
| ENSG0000 | -1.77039 | -3.41147 | 2.63E-20 | 1.56E-18 | A2M |
| ENSG0000 | -1.28852 | -2.44277 | 4.51E-07 | 6.04E-06 | DOK7 |
| ENSG0000 | -1.11997 | -2.17342 | 0.003452 | 0.017719 | PLEKHD1 |
| ENSG0000 | -1.31478 | -2.48764 | 2.60E-25 | 2.08E-23 | TUBB6 |
| ENSG0000 | -3.17758 | -9.0479 | 6.63E-13 | 2.01E-11 | TMPRSS7 |
| ENSG0000 | -3.72107 | -13.1872 | 0.008318 | 0.036888 | CIDEA |
| ENSG0000 | -1.76087 | -3.38903 | 1.70E-29 | 1.72E-27 | ACBD7 |
| ENSG0000 | -2.31365 | -4.97141 | 2.23E-56 | 5.45E-54 | MAP3K19 |
| ENSG0000 | -5.98777 | -63.4599 | 0.003737 | 0.018903 | LINC02910 |
| ENSG0000 | -1.34336 | -2.53742 | 2.24E-11 | 5.66E-10 | BOK |
| ENSG0000 | -1.68762 | -3.22125 | 1.10E-07 | 1.64E-06 | GCNT4 |
| ENSG0000 | -1.06627 | -2.09401 | 5.78E-07 | 7.61E-06 | RIMKLA |
| ENSG0000 | -2.1482 | -4.43273 | 1.96E-09 | 3.80E-08 | CLVS1 |
| ENSG0000 | -1.26067 | -2.39607 | 0.00181 | 0.010253 | KCNA3 |
| ENSG0000 | -1.23629 | -2.35591 | 2.42E-17 | 1.11E-15 | CAVIN1 |
| ENSG0000 | -1.62628 | -3.08716 | 2.47E-05 | 0.000232 | ST8SIA3 |
| ENSG0000 | -1.52425 | -2.87637 | 3.91E-11 | 9.58E-10 | ODF3B |
| ENSG0000 | -5.49887 | -45.2195 | 0.011166 | 0.047126 | H1-8 |
| ENSG0000 | -1.42278 | -2.68101 | 0.007684 | 0.034507 | TMEM139 |
| ENSG0000 | -2.3136 | -4.97123 | 8.19E-24 | 6.17E-22 | ERICH3 |
| ENSG0000 | -2.04964 | -4.14003 | 3.14E-34 | 4.00E-32 | EGR3 |
| ENSG0000 | -1.39852 | -2.63631 | 6.89E-14 | 2.31E-12 | VWA1 |
| ENSG0000 | -1.13542 | -2.19682 | 2.08E-12 | 5.96E-11 | FJX1 |
| ENSG0000 | -1.30742 | -2.47498 | 0.00734 | 0.033191 | RPRML |
| ENSG0000 | -2.53179 | -5.78289 | 5.81E-12 | 1.58E-10 | APOBEC3B |
| ENSG0000 | -1.33432 | -2.52157 | 1.01E-13 | 3.29E-12 | LRRC3B |
| ENSG0000 | -1.67285 | -3.18843 | 6.82E-29 | 6.59E-27 | AKAP5 |
| ENSG0000 | -1.05424 | -2.07662 | 2.86E-05 | 0.000264 | CFAP276 |

| | | | | | |
|---|---|---|---|---|---|
| ENSG0000 | -1.6142 | -3.06143 | 1.09E-06 | 1.36E-05 | NRXN1 |
| ENSG0000 | -3.49691 | -11.2895 | 2.54E-46 | 4.99E-44 | ACTA2-AS1 |
| ENSG0000 | -1.7535 | -3.37176 | 3.55E-19 | 1.92E-17 | PLD5 |
| ENSG0000 | -1.2511 | -2.38022 | 1.59E-18 | 8.19E-17 | GPR137C |
| ENSG0000 | -1.39724 | -2.63398 | 3.58E-34 | 4.53E-32 | MAPK15 |
| ENSG0000 | -1.41759 | -2.67139 | 4.43E-15 | 1.68E-13 | NME9 |
| ENSG0000 | -1.29899 | -2.46056 | 6.45E-10 | 1.34E-08 | CFAP65 |
| ENSG0000 | -7.00542 | -128.482 | 3.71E-05 | 0.000334 | TIGIT |
| ENSG0000 | -1.50866 | -2.84545 | 2.91E-07 | 4.03E-06 | KIAA2012 |
| ENSG0000 | -3.82475 | -14.1698 | 0.008608 | 0.037931 | KCNK4 |
| ENSG0000 | -1.13968 | -2.20332 | 0.00435 | 0.021482 | MAFA |
| ENSG0000 | -1.76268 | -3.39328 | 2.25E-18 | 1.14E-16 | PLCXD3 |
| ENSG0000 | -1.76846 | -3.40691 | 0.011195 | 0.047219 | CALN1 |
| ENSG0000 | -1.50196 | -2.83227 | 2.04E-26 | 1.72E-24 | CHST6 |
| ENSG0000 | -2.05908 | -4.16721 | 2.35E-06 | 2.75E-05 | HOATZ |
| ENSG0000 | -1.14114 | -2.20556 | 0.005223 | 0.02503 | OPCML |
| ENSG0000 | -2.00333 | -4.00924 | 3.76E-12 | 1.04E-10 | SLC35F3 |
| ENSG0000 | -1.98644 | -3.96259 | 1.35E-10 | 3.07E-09 | OLFML1 |
| ENSG0000 | -1.38691 | -2.61518 | 0.000442 | 0.003015 | ZNF730 |
| ENSG0000 | -1.15424 | -2.22566 | 3.98E-05 | 0.000356 | ARSI |
| ENSG0000 | -1.19625 | -2.29143 | 0.000558 | 0.003707 | DNAH2 |
| ENSG0000 | -1.45295 | -2.73768 | 8.13E-07 | 1.04E-05 | SDR42E2 |
| ENSG0000 | -1.17714 | -2.26128 | 4.99E-12 | 1.37E-10 | SMTN |
| ENSG0000 | -1.20821 | -2.31051 | 2.67E-29 | 2.63E-27 | ST6GALNAC3 |
| ENSG0000 | -1.10647 | -2.15318 | 3.35E-08 | 5.41E-07 | CLDN5 |
| ENSG0000 | -2.13699 | -4.39845 | 1.80E-16 | 7.77E-15 | CNTN2 |
| ENSG0000 | -1.89571 | -3.72104 | 0.00559 | 0.026508 | NCMAP |
| ENSG0000 | -1.12137 | -2.17554 | 3.23E-35 | 4.44E-33 | NELL2 |
| ENSG0000 | -2.08619 | -4.24625 | 4.77E-36 | 6.81E-34 | ZNF93 |
| ENSG0000 | -1.27458 | -2.41929 | 0.006872 | 0.031448 | CDCA2 |
| ENSG0000 | -1.09066 | -2.12971 | 9.08E-15 | 3.31E-13 | TNFAIP8L1 |
| ENSG0000 | -1.43443 | -2.70275 | 1.42E-11 | 3.69E-10 | MORN5 |
| ENSG0000 | -1.29383 | -2.45178 | 4.22E-07 | 5.69E-06 | MYBL1 |
| ENSG0000 | -1.88491 | -3.6933 | 0.005142 | 0.024679 | IKZF1 |
| ENSG0000 | -3.36573 | -10.3082 | 2.40E-06 | 2.80E-05 | CCIN |
| ENSG0000 | -3.83611 | -14.2819 | 0.009385 | 0.040744 | AGBL4 |
| ENSG0000 | -1.27497 | -2.41993 | 8.99E-09 | 1.60E-07 | KLHL32 |
| ENSG0000 | -1.62034 | -3.07447 | 1.20E-19 | 6.79E-18 | |
| ENSG0000 | -1.03716 | -2.05219 | 4.36E-10 | 9.33E-09 | GLDN |
| ENSG0000 | -1.31638 | -2.49041 | 3.28E-17 | 1.50E-15 | KIF24 |
| ENSG0000 | -2.07849 | -4.22364 | 3.61E-32 | 4.16E-30 | CFAP73 |
| ENSG0000 | -1.32886 | -2.51204 | 0.000899 | 0.005608 | FSCN2 |
| ENSG0000 | -1.65985 | -3.15984 | 6.87E-32 | 7.75E-30 | SPIN4 |
| ENSG0000 | -4.26813 | -19.2679 | 0.001431 | 0.008429 | ZNF732 |
| ENSG0000 | -1.034 | -2.0477 | 7.51E-08 | 1.15E-06 | FAM183A |
| ENSG0000 | -1.1821 | -2.26907 | 2.55E-16 | 1.09E-14 | C3orf70 |
| ENSG0000 | -1.55151 | -2.93125 | 3.47E-34 | 4.40E-32 | LYPD6 |
| ENSG0000 | -5.53053 | -46.2227 | 0.000732 | 0.004687 | None |
| ENSG0000 | -1.67833 | -3.20058 | 3.45E-13 | 1.08E-11 | LUZP2 |
| ENSG0000 | -1.13064 | -2.18957 | 1.49E-08 | 2.55E-07 | LHFPL3 |

| | | | | | |
|---|---|---|---|---|---|
| ENSG0000 | -1.11381 | -2.16417 | 0.010904 | 0.046217 | TLR5 |
| ENSG0000 | -1.37646 | -2.59631 | 7.85E-05 | 0.000654 | DNAJB13 |
| ENSG0000 | -1.57473 | -2.9788 | 5.12E-05 | 0.000447 | LRRC74B |
| ENSG0000 | -1.64475 | -3.12693 | 2.06E-06 | 2.43E-05 | LDLRAD2 |
| ENSG0000 | -1.01667 | -2.02324 | 4.84E-47 | 9.76E-45 | TUBB4B |
| ENSG0000 | -1.14168 | -2.20638 | 1.65E-20 | 9.93E-19 | PLSCR1 |
| ENSG0000 | -1.03154 | -2.04421 | 2.63E-09 | 5.01E-08 | ENO4 |
| ENSG0000 | -1.58575 | -3.00165 | 3.69E-08 | 5.92E-07 | |
| ENSG0000 | -1.13952 | -2.20308 | 6.03E-10 | 1.26E-08 | CFAP77 |
| ENSG0000 | -1.05904 | -2.08354 | 5.17E-11 | 1.25E-09 | CFAP54 |
| ENSG0000 | -1.18395 | -2.27198 | 0.008205 | 0.036472 | C2orf80 |
| ENSG0000 | -1.59086 | -3.01228 | 0.000109 | 0.000876 | PRELP |
| ENSG0000 | -5.78285 | -55.0568 | 0.005342 | 0.025513 | SLC15A5 |
| ENSG0000 | -4.30044 | -19.7043 | 2.55E-81 | 1.02E-78 | FAM111B |
| ENSG0000 | -1.43103 | -2.6964 | 0.007091 | 0.032275 | VSTM1 |
| ENSG0000 | -7.01984 | -129.772 | 8.52E-06 | 8.83E-05 | KRT77 |
| ENSG0000 | -2.53252 | -5.78581 | 6.88E-10 | 1.42E-08 | KIF19 |
| ENSG0000 | -1.63514 | -3.10617 | 4.96E-13 | 1.52E-11 | HRCT1 |
| ENSG0000 | -1.16386 | -2.24056 | 5.46E-18 | 2.65E-16 | CPNE4 |
| ENSG0000 | -2.18883 | -4.55936 | 3.55E-13 | 1.11E-11 | AJAP1 |
| ENSG0000 | -1.76223 | -3.39222 | 2.74E-24 | 2.11E-22 | PDLIM7 |
| ENSG0000 | -1.14216 | -2.20711 | 3.29E-13 | 1.03E-11 | FLNA |
| ENSG0000 | -1.50287 | -2.83406 | 1.71E-23 | 1.28E-21 | NEK5 |
| ENSG0000 | -1.90372 | -3.74178 | 6.02E-10 | 1.26E-08 | FBXL22 |
| ENSG0000 | -1.16165 | -2.23713 | 2.69E-05 | 0.00025 | VEPH1 |
| ENSG0000 | -6.01112 | -64.4953 | 0.002885 | 0.015264 | GGT3P |
| ENSG0000 | -2.07993 | -4.22786 | 1.18E-40 | 1.94E-38 | MFAP5 |
| ENSG0000 | -1.63983 | -3.11629 | 0.000371 | 0.002585 | CFAP299 |
| ENSG0000 | -1.323 | -2.50186 | 8.30E-20 | 4.76E-18 | TMEM229B |
| ENSG0000 | -1.60392 | -3.03969 | 0.007216 | 0.032741 | |
| ENSG0000 | -1.06976 | -2.09909 | 1.01E-08 | 1.78E-07 | ZNF69 |
| ENSG0000 | -1.65956 | -3.15921 | 1.03E-21 | 6.84E-20 | TPM2 |
| ENSG0000 | -7.10595 | -137.754 | 9.77E-08 | 1.47E-06 | PLN |
| ENSG0000 | -1.32049 | -2.4975 | 6.56E-07 | 8.56E-06 | MMP17 |
| ENSG0000 | -2.0394 | -4.11075 | 1.17E-49 | 2.52E-47 | LRRTM3 |
| ENSG0000 | -1.71668 | -3.28678 | 3.37E-06 | 3.82E-05 | RCSD1 |
| ENSG0000 | -1.39472 | -2.62937 | 1.66E-42 | 2.94E-40 | MT-CO1 |
| ENSG0000 | -1.77226 | -3.41588 | 1.21E-08 | 2.11E-07 | ARHGAP11A |
| ENSG0000 | -5.52955 | -46.1912 | 0.009188 | 0.040044 | DCAF12L1 |
| ENSG0000 | -1.08272 | -2.11803 | 7.91E-07 | 1.01E-05 | L1CAM |
| ENSG0000 | -1.21704 | -2.32469 | 1.63E-32 | 1.95E-30 | MT-CO3 |
| ENSG0000 | -5.79862 | -55.662 | 0.007123 | 0.032393 | RNU1-72P |
| ENSG0000 | -3.00337 | -8.01872 | 3.55E-06 | 4.01E-05 | Y_RNA |
| ENSG0000 | -1.69036 | -3.22737 | 6.72E-10 | 1.40E-08 | IQANK1 |
| ENSG0000 | -1.1761 | -2.25964 | 1.58E-12 | 4.60E-11 | SERTAD4-AS1 |
| ENSG0000 | -1.20627 | -2.3074 | 9.88E-06 | 0.000101 | SAMD5 |
| ENSG0000 | -1.73892 | -3.33786 | 3.37E-16 | 1.43E-14 | SMIM5 |
| ENSG0000 | -1.22262 | -2.3337 | 8.06E-06 | 8.41E-05 | BTBD17 |
| ENSG0000 | -1.08962 | -2.12818 | 5.20E-24 | 3.94E-22 | LAYN |
| ENSG0000 | -6.19649 | -73.3381 | 0.000888 | 0.005551 | PSORS1C3 |

| | | | | | |
|---|---|---|---|---|---|
| ENSG0000 | -2.3753 | -5.18843 | 1.19E-10 | 2.74E-09 | C6orf15 |
| ENSG0000 | -4.79865 | -27.8315 | 0.011886 | 0.049578 | HSP90AA4P |
| ENSG0000 | -2.23428 | -4.70527 | 0.006369 | 0.029638 | CCDC144NL |
| ENSG0000 | -2.05031 | -4.14194 | 1.66E-07 | 2.40E-06 | GMNC |
| ENSG0000 | -5.79914 | -55.6822 | 0.008113 | 0.036128 | FAM99A |
| ENSG0000 | -2.40598 | -5.29997 | 1.92E-41 | 3.23E-39 | C21orf62 |
| ENSG0000 | -1.54811 | -2.92435 | 2.16E-21 | 1.39E-19 | CFAP99 |
| ENSG0000 | -1.34717 | -2.54413 | 2.12E-06 | 2.49E-05 | COL6A6 |
| ENSG0000 | -1.0078 | -2.01084 | 8.06E-09 | 1.44E-07 | H1-10-AS1 |
| ENSG0000 | -1.2397 | -2.3615 | 3.99E-17 | 1.81E-15 | |
| ENSG0000 | -1.84115 | -3.58295 | 3.20E-11 | 7.92E-10 | VGLL3 |
| ENSG0000 | -3.18868 | -9.11778 | 0.008609 | 0.037931 | SNORD116-8 |
| ENSG0000 | -5.51166 | -45.622 | 0.008008 | 0.035726 | Y_RNA |
| ENSG0000 | -1.64118 | -3.11922 | 0.011626 | 0.048748 | MIR23B |
| ENSG0000 | -1.65448 | -3.14809 | 0.002715 | 0.014498 | None |
| ENSG0000 | -4.78214 | -27.5149 | 0.006954 | 0.031766 | MIR33B |
| ENSG0000 | -1.5516 | -2.93142 | 0.009862 | 0.042507 | MIR27B |
| ENSG0000 | -1.32526 | -2.50578 | 0.002728 | 0.014556 | MT-TA |
| ENSG0000 | -5.80166 | -55.7795 | 0.005823 | 0.027413 | MIR3065 |
| ENSG0000 | -1.59521 | -3.02138 | 2.87E-41 | 4.79E-39 | CFAP45 |
| ENSG0000 | -2.14838 | -4.43331 | 7.23E-27 | 6.22E-25 | MXD3 |
| ENSG0000 | -5.76861 | -54.5162 | 0.011036 | 0.046679 | YBX2P1 |
| ENSG0000 | -1.93958 | -3.83594 | 9.82E-42 | 1.68E-39 | LBH |
| ENSG0000 | -1.20715 | -2.3088 | 1.29E-17 | 6.04E-16 | S1PR3 |
| ENSG0000 | -1.39924 | -2.63763 | 7.87E-60 | 2.12E-57 | EMP2 |
| ENSG0000 | -1.29662 | -2.45652 | 4.01E-08 | 6.40E-07 | None |
| ENSG0000 | -6.00212 | -64.0941 | 0.001557 | 0.009043 | EFCAB9 |
| ENSG0000 | -1.38036 | -2.60334 | 1.02E-06 | 1.28E-05 | C10orf105 |
| ENSG0000 | -1.61273 | -3.0583 | 6.22E-06 | 6.63E-05 | TUBA5P |
| ENSG0000 | -3.13742 | -8.79952 | 0.000858 | 0.005381 | FAM166B |
| ENSG0000 | -2.36138 | -5.13862 | 2.93E-06 | 3.36E-05 | SCRT2 |
| ENSG0000 | -2.10785 | -4.31048 | 2.00E-09 | 3.87E-08 | SIAH3 |
| ENSG0000 | -1.00981 | -2.01365 | 1.34E-11 | 3.49E-10 | TNFRSF25 |
| ENSG0000 | -5.78564 | -55.1632 | 0.001111 | 0.006753 | UQCRFS1P3 |
| ENSG0000 | -1.24152 | -2.36447 | 0.009816 | 0.042327 | TATDN2P2 |
| ENSG0000 | -2.02073 | -4.05789 | 0.006553 | 0.03032 | None |
| ENSG0000 | -5.50508 | -45.4144 | 0.01065 | 0.045358 | TUBB8P2 |
| ENSG0000 | -1.02194 | -2.03065 | 0.008674 | 0.038153 | None |
| ENSG0000 | -1.19661 | -2.292 | 4.78E-12 | 1.31E-10 | PLXNA4 |
| ENSG0000 | -1.0218 | -2.03045 | 0.002608 | 0.014029 | HMSD |
| ENSG0000 | -6.03064 | -65.374 | 0.005393 | 0.025716 | RNU2-37P |
| ENSG0000 | -2.217 | -4.64925 | 0.003454 | 0.017723 | None |
| ENSG0000 | -1.29313 | -2.45059 | 0.006378 | 0.029671 | None |
| ENSG0000 | -5.52371 | -46.0048 | 0.002905 | 0.015349 | LINC01449 |
| ENSG0000 | -1.11816 | -2.1707 | 0.002455 | 0.013313 | WARS2-IT1 |
| ENSG0000 | -6.53589 | -92.7896 | 0.000108 | 0.000874 | OLFM5P |
| ENSG0000 | -1.28138 | -2.43071 | 5.01E-14 | 1.70E-12 | PRR29 |
| ENSG0000 | -4.19488 | -18.314 | 5.68E-09 | 1.03E-07 | ZNF812P |
| ENSG0000 | -6.89116 | -118.698 | 1.10E-05 | 0.000112 | None |
| ENSG0000 | -1.58697 | -3.00418 | 0.011207 | 0.047261 | PGM5-AS1 |

| | | | | | |
|---|---|---|---|---|---|
| ENSG0000 | -2.77693 | -6.85394 | 0.00194 | 0.010872 | ACKR4P1 |
| ENSG0000 | -5.52163 | -45.9385 | 0.001648 | 0.009487 | MPRIP-AS1 |
| ENSG0000 | -1.10612 | -2.15266 | 1.08E-07 | 1.61E-06 | ZNF469 |
| ENSG0000 | -4.79168 | -27.6974 | 0.01176 | 0.049184 | LINC01831 |
| ENSG0000 | -1.5125 | -2.85304 | 0.001193 | 0.007197 | S1PR1-DT |
| ENSG0000 | -1.01472 | -2.0205 | 0.000357 | 0.002499 | FGF13-AS1 |
| ENSG0000 | -1.1258 | -2.18223 | 0.01046 | 0.044708 | AP4B1-AS1 |
| ENSG0000 | -1.35334 | -2.55502 | 0.000766 | 0.004873 | GAS1RR |
| ENSG0000 | -5.80472 | -55.8976 | 0.009663 | 0.041809 | |
| ENSG0000 | -1.4739 | -2.77771 | 0.006787 | 0.031138 | None |
| ENSG0000 | -1.01953 | -2.02726 | 2.52E-05 | 0.000236 | None |
| ENSG0000 | -1.12914 | -2.18729 | 1.59E-05 | 0.000156 | None |
| ENSG0000 | -1.57408 | -2.97746 | 0.008278 | 0.03673 | SUCLA2-AS1 |
| ENSG0000 | -1.27979 | -2.42803 | 0.00103 | 0.006311 | None |
| ENSG0000 | -1.64349 | -3.12421 | 2.77E-14 | 9.61E-13 | None |
| ENSG0000 | -2.21496 | -4.6427 | 3.13E-11 | 7.76E-10 | None |
| ENSG0000 | -6.19821 | -73.4257 | 0.003549 | 0.018136 | GRK5-IT1 |
| ENSG0000 | -5.78753 | -55.2357 | 0.007898 | 0.035322 | None |
| ENSG0000 | -2.37258 | -5.17866 | 0.000178 | 0.001365 | HCG22 |
| ENSG0000 | -4.20158 | -18.3993 | 0.006887 | 0.031506 | None |
| ENSG0000 | -1.25798 | -2.39161 | 8.81E-17 | 3.87E-15 | MBNL1-AS1 |
| ENSG0000 | -2.25296 | -4.7666 | 0.000526 | 0.003515 | MYOSLID |
| ENSG0000 | -2.02457 | -4.06872 | 0.000157 | 0.001217 | None |
| ENSG0000 | -1.01235 | -2.0172 | 0.003498 | 0.017922 | PPIAP39 |
| ENSG0000 | -2.51242 | -5.70577 | 2.09E-19 | 1.15E-17 | ANKRD66 |
| ENSG0000 | -2.24528 | -4.74129 | 0.001417 | 0.008356 | ARMC2-AS1 |
| ENSG0000 | -6.52633 | -92.1767 | 6.06E-06 | 6.48E-05 | GGTLC4P |
| ENSG0000 | -1.32811 | -2.51074 | 0.010345 | 0.044299 | MEIS1-AS2 |
| ENSG0000 | -5.79694 | -55.5974 | 0.000913 | 0.005678 | |
| ENSG0000 | -2.18484 | -4.54676 | 0.007382 | 0.033336 | NEK2-DT |
| ENSG0000 | -5.77501 | -54.7584 | 0.000775 | 0.004916 | |
| ENSG0000 | -1.10624 | -2.15284 | 0.002674 | 0.014322 | TSPAN19 |
| ENSG0000 | -4.78786 | -27.6241 | 0.009725 | 0.042015 | None |
| ENSG0000 | -1.32692 | -2.50866 | 4.94E-15 | 1.86E-13 | GNG12-AS1 |
| ENSG0000 | -1.16648 | -2.24463 | 1.84E-08 | 3.11E-07 | TGFB2-AS1 |
| ENSG0000 | -2.47139 | -5.54579 | 3.19E-08 | 5.16E-07 | None |
| ENSG0000 | -1.10778 | -2.15514 | 0.010991 | 0.046525 | None |
| ENSG0000 | -2.01645 | -4.04588 | 0.002855 | 0.015149 | None |
| ENSG0000 | -1.58542 | -3.00094 | 0.003665 | 0.018603 | HMGA1P8 |
| ENSG0000 | -1.04743 | -2.06685 | 6.99E-10 | 1.45E-08 | LINC00472 |
| ENSG0000 | -1.60865 | -3.04965 | 0.000397 | 0.002746 | FNDC1-AS1 |
| ENSG0000 | -1.10059 | -2.14442 | 0.008179 | 0.036379 | MYCNOS |
| ENSG0000 | -2.29388 | -4.90373 | 4.34E-06 | 4.80E-05 | |
| ENSG0000 | -4.38987 | -20.9644 | 0.000608 | 0.003988 | None |
| ENSG0000 | -1.65356 | -3.14609 | 5.63E-06 | 6.08E-05 | ACTG1P25 |
| ENSG0000 | -5.51193 | -45.6305 | 0.006775 | 0.031095 | None |
| ENSG0000 | -1.84889 | -3.60223 | 0.000827 | 0.005209 | LINC01102 |
| ENSG0000 | -1.11999 | -2.17345 | 0.00061 | 0.003998 | BHLHE40-AS1 |
| ENSG0000 | -1.29062 | -2.44633 | 5.93E-05 | 0.000509 | None |
| ENSG0000 | -1.30221 | -2.46606 | 3.03E-06 | 3.46E-05 | None |

| | | | | | |
|---|---|---|---|---|---|
| ENSG0000 | -3.74505 | -13.4082 | 3.39E-08 | 5.46E-07 | None |
| ENSG0000 | -4.77759 | -27.4283 | 0.011012 | 0.046596 | KCNMA1-AS1 |
| ENSG0000 | -5.5068 | -45.4688 | 0.000957 | 0.005919 | None |
| ENSG0000 | -2.41029 | -5.31582 | 0.011125 | 0.046988 | LINC01186 |
| ENSG0000 | -2.38601 | -5.22708 | 0.000292 | 0.002095 | COX5BP6 |
| ENSG0000 | -1.83398 | -3.56519 | 0.000245 | 0.0018 | KIFC1 |
| ENSG0000 | -1.19798 | -2.29418 | 6.38E-14 | 2.15E-12 | MTCO1P12 |
| ENSG0000 | -1.7748 | -3.42191 | 0.003505 | 0.017949 | LINC01625 |
| ENSG0000 | -2.3276 | -5.01971 | 1.32E-06 | 1.62E-05 | LINC00707 |
| ENSG0000 | -5.53233 | -46.2805 | 0.003334 | 0.017203 | RNU2-11P |
| ENSG0000 | -5.77757 | -54.8557 | 0.005715 | 0.026996 | RSPH1-DT |
| ENSG0000 | -1.40941 | -2.65629 | 2.71E-20 | 1.61E-18 | AQP1 |
| ENSG0000 | -2.33486 | -5.04502 | 0.004593 | 0.022485 | None |
| ENSG0000 | -5.21265 | -37.082 | 0.004226 | 0.020953 | None |
| ENSG0000 | -5.52953 | -46.1908 | 0.008634 | 0.038019 | TDGF1 |
| ENSG0000 | -1.5965 | -3.02409 | 5.91E-05 | 0.000508 | INMT |
| ENSG0000 | -6.78779 | -110.491 | 0.000135 | 0.001068 | RPL7AP2 |
| ENSG0000 | -1.36492 | -2.57562 | 1.64E-19 | 9.10E-18 | NECTIN3-AS1 |
| ENSG0000 | -1.24728 | -2.37393 | 5.31E-11 | 1.28E-09 | CNTF |
| ENSG0000 | -2.32229 | -5.00125 | 6.51E-06 | 6.91E-05 | VSIG8 |
| ENSG0000 | -1.01671 | -2.0233 | 0.004696 | 0.022895 | TICAM2 |
| ENSG0000 | -1.35625 | -2.56019 | 1.61E-13 | 5.18E-12 | CFAP57 |
| ENSG0000 | -1.23271 | -2.35008 | 0.000115 | 0.000924 | GSTA1 |
| ENSG0000 | -2.89929 | -7.46057 | 0.006297 | 0.029367 | None |
| ENSG0000 | -5.5362 | -46.4049 | 0.000603 | 0.003959 | CYP2U1-AS1 |
| ENSG0000 | -1.73876 | -3.33749 | 8.39E-29 | 8.08E-27 | STARD4-AS1 |
| ENSG0000 | -1.91063 | -3.75974 | 7.19E-07 | 9.27E-06 | None |
| ENSG0000 | -1.52132 | -2.87053 | 2.36E-07 | 3.31E-06 | None |
| ENSG0000 | -2.12522 | -4.36268 | 0.002311 | 0.012624 | PCP4L1 |
| ENSG0000 | -1.02334 | -2.03262 | 5.13E-05 | 0.000448 | C8orf34-AS1 |
| ENSG0000 | -1.27401 | -2.41833 | 0.002793 | 0.014855 | ZNF474-AS1 |
| ENSG0000 | -2.40546 | -5.29803 | 0.00165 | 0.009496 | None |
| ENSG0000 | -3.33718 | -10.1063 | 0.000645 | 0.004202 | CARMN |
| ENSG0000 | -1.84185 | -3.58469 | 0.00538 | 0.025675 | None |
| ENSG0000 | -2.06934 | -4.19696 | 3.06E-05 | 0.000281 | HS3ST5 |
| ENSG0000 | -5.53248 | -46.2851 | 0.005816 | 0.027391 | None |
| ENSG0000 | -1.08504 | -2.12144 | 1.36E-09 | 2.68E-08 | LINC02200 |
| ENSG0000 | -3.44316 | -10.8766 | 4.90E-05 | 0.000429 | None |
| ENSG0000 | -1.01886 | -2.02632 | 0.002546 | 0.013736 | STK19B |
| ENSG0000 | -1.39874 | -2.63672 | 1.86E-06 | 2.21E-05 | ADAMTS16-DT |
| ENSG0000 | -1.15218 | -2.2225 | 2.37E-18 | 1.19E-16 | YPEL3-DT |
| ENSG0000 | -3.83677 | -14.2883 | 0.001443 | 0.008491 | None |
| ENSG0000 | -5.78293 | -55.0597 | 0.003904 | 0.019635 | RNU7-71P |
| ENSG0000 | -6.01975 | -64.882 | 0.001125 | 0.006826 | RNU7-160P |
| ENSG0000 | -5.51879 | -45.8482 | 0.003184 | 0.016541 | SCARNA21 |
| ENSG0000 | -5.52966 | -46.195 | 0.004842 | 0.023461 | SCARNA1 |
| ENSG0000 | -4.79093 | -27.683 | 0.006891 | 0.031517 | None |
| ENSG0000 | -1.68352 | -3.2121 | 1.33E-07 | 1.95E-06 | MIR124-1HG |
| ENSG0000 | -1.24542 | -2.37087 | 8.62E-12 | 2.30E-10 | TRNP1 |
| ENSG0000 | -5.99232 | -63.6603 | 0.000918 | 0.005705 | KCNIP1-AS1 |

| | | | | | |
|---|---|---|---|---|---|
| ENSG0000 | -5.19809 | -36.7098 | 0.004765 | 0.02316 | KCNIP1-OT1 |
| ENSG0000 | -1.75628 | -3.37827 | 0.009666 | 0.041809 | None |
| ENSG0000 | -5.79639 | -55.5759 | 0.000602 | 0.003952 | None |
| ENSG0000 | -1.05578 | -2.07885 | 0.001394 | 0.008236 | PCDHGB1 |
| ENSG0000 | -1.76336 | -3.39487 | 0.007266 | 0.032903 | None |
| ENSG0000 | -3.13484 | -8.78378 | 1.81E-30 | 1.93E-28 | None |
| ENSG0000 | -1.26013 | -2.39517 | 0.000725 | 0.00465 | None |
| ENSG0000 | -6.64505 | -100.083 | 2.87E-05 | 0.000265 | GLYATL1P2 |
| ENSG0000 | -3.79666 | -13.8966 | 3.56E-47 | 7.23E-45 | None |
| ENSG0000 | -1.55341 | -2.9351 | 0.004206 | 0.020862 | None |
| ENSG0000 | -5.19412 | -36.6088 | 0.002185 | 0.012033 | None |
| ENSG0000 | -1.26135 | -2.3972 | 0.000801 | 0.005064 | None |
| ENSG0000 | -6.02519 | -65.1275 | 0.002421 | 0.013151 | None |
| ENSG0000 | -1.34502 | -2.54034 | 1.15E-36 | 1.68E-34 | RMST |
| ENSG0000 | -1.80912 | -3.5043 | 2.46E-05 | 0.000232 | None |
| ENSG0000 | -1.46801 | -2.7664 | 0.009436 | 0.040942 | None |
| ENSG0000 | -5.20112 | -36.787 | 0.005453 | 0.025957 | None |
| ENSG0000 | -5.52407 | -46.0162 | 0.0051 | 0.024515 | None |
| ENSG0000 | -6.77111 | -109.221 | 0.000179 | 0.001371 | None |
| ENSG0000 | -1.67803 | -3.1999 | 1.11E-06 | 1.39E-05 | PAFAH1B2P2 |
| ENSG0000 | -5.52191 | -45.9473 | 0.000941 | 0.005829 | None |
| ENSG0000 | -1.00095 | -2.00132 | 0.000333 | 0.002352 | AJUBA-DT |
| ENSG0000 | -5.21019 | -37.0189 | 0.006805 | 0.031203 | RRAGAP1-AS1 |
| ENSG0000 | -1.98324 | -3.95379 | 0.007306 | 0.033063 | TRIM34 |
| ENSG0000 | -5.80113 | -55.7587 | 0.000311 | 0.002218 | None |
| ENSG0000 | -3.70625 | -13.0525 | 5.58E-08 | 8.70E-07 | None |
| ENSG0000 | -5.7835 | -55.0818 | 0.001136 | 0.00689 | CKS1BP1 |
| ENSG0000 | -6.00553 | -64.2459 | 0.000215 | 0.001606 | None |
| ENSG0000 | -1.21603 | -2.32306 | 2.71E-13 | 8.52E-12 | TUBB3 |
| ENSG0000 | -2.96369 | -7.80115 | 1.17E-25 | 9.48E-24 | TMEM179 |
| ENSG0000 | -6.37819 | -83.1813 | 0.000417 | 0.002859 | None |
| ENSG0000 | -1.13632 | -2.1982 | 0.00128 | 0.007663 | LINC00639 |
| ENSG0000 | -5.78537 | -55.153 | 0.002458 | 0.013321 | GNRHR2P1 |
| ENSG0000 | -2.27177 | -4.82915 | 7.60E-11 | 1.80E-09 | TPM1-AS |
| ENSG0000 | -5.19402 | -36.6064 | 0.008431 | 0.037249 | HMGB1P8 |
| ENSG0000 | -5.51759 | -45.8099 | 0.00825 | 0.036637 | |
| ENSG0000 | -1.85387 | -3.61469 | 0.006224 | 0.029065 | None |
| ENSG0000 | -1.15422 | -2.22564 | 1.38E-08 | 2.38E-07 | None |
| ENSG0000 | -1.57567 | -2.98074 | 0.001477 | 0.008658 | None |
| ENSG0000 | -1.217 | -2.32462 | 0.002228 | 0.012247 | AQP4-AS1 |
| ENSG0000 | -5.99946 | -63.9763 | 0.003453 | 0.017719 | CES1P2 |
| ENSG0000 | -1.21206 | -2.31668 | 1.45E-09 | 2.86E-08 | None |
| ENSG0000 | -7.2073 | -147.779 | 4.80E-05 | 0.000421 | None |
| ENSG0000 | -1.75794 | -3.38214 | 0.000177 | 0.001354 | CLEC19A |
| ENSG0000 | -2.48754 | -5.60822 | 5.48E-05 | 0.000475 | |
| ENSG0000 | -1.2256 | -2.33852 | 1.89E-10 | 4.25E-09 | None |
| ENSG0000 | -2.14407 | -4.42007 | 2.15E-07 | 3.05E-06 | None |
| ENSG0000 | -5.52699 | -46.1093 | 0.003114 | 0.016259 | GREP1 |
| ENSG0000 | -1.71889 | -3.29183 | 0.00178 | 0.010111 | |
| ENSG0000 | -7.10136 | -137.316 | 7.72E-06 | 8.08E-05 | None |

| | | | | | |
|---|---|---|---|---|---|
| ENSG0000 | -4.55314 | -23.4764 | 0.002969 | 0.015634 | None |
| ENSG0000 | -5.53238 | -46.2821 | 0.000732 | 0.004687 | MIR3138 |
| ENSG0000 | -1.39808 | -2.6355 | 0.001228 | 0.007386 | AGAP12P |
| ENSG0000 | -1.20858 | -2.3111 | 1.09E-05 | 0.000111 | TSPOAP1-AS1 |
| ENSG0000 | -1.27177 | -2.41457 | 1.71E-05 | 0.000167 | None |
| ENSG0000 | -6.00114 | -64.0505 | 0.000501 | 0.003369 | None |
| ENSG0000 | -3.1431 | -8.83421 | 0.003583 | 0.018292 | None |
| ENSG0000 | -1.13307 | -2.19325 | 0.00026 | 0.00189 | PTGES3L |
| ENSG0000 | -5.15039 | -35.5157 | 1.29E-08 | 2.24E-07 | None |
| ENSG0000 | -1.30708 | -2.4744 | 1.96E-07 | 2.80E-06 | None |
| ENSG0000 | -1.4666 | -2.76369 | 7.49E-08 | 1.15E-06 | None |
| ENSG0000 | -1.06729 | -2.0955 | 0.000405 | 0.00279 | MIR924HG |
| ENSG0000 | -1.76393 | -3.39622 | 0.00549 | 0.026103 | ZNF887P |
| ENSG0000 | -2.31095 | -4.9621 | 8.66E-11 | 2.04E-09 | LINC01836 |
| ENSG0000 | -6.9061 | -119.934 | 1.39E-05 | 0.000138 | None |
| ENSG0000 | -1.62489 | -3.08419 | 0.001951 | 0.010924 | ASXL3-DT |
| ENSG0000 | -1.29967 | -2.46173 | 0.000188 | 0.001432 | None |
| ENSG0000 | -5.52898 | -46.1732 | 0.001786 | 0.010136 | None |
| ENSG0000 | -6.19645 | -73.3362 | 0.00117 | 0.007073 | LINC01999 |
| ENSG0000 | -1.41983 | -2.67554 | 8.21E-05 | 0.000681 | None |
| ENSG0000 | -1.14575 | -2.2126 | 3.98E-07 | 5.38E-06 | CCDC177 |
| ENSG0000 | -5.53883 | -46.4893 | 0.001562 | 0.00907 | None |
| ENSG0000 | -5.52647 | -46.0929 | 0.002446 | 0.013276 | None |
| ENSG0000 | -1.54196 | -2.91189 | 1.83E-10 | 4.13E-09 | GABRQ |
| ENSG0000 | -1.60028 | -3.03203 | 0.004202 | 0.020844 | PLCE1-AS1 |
| ENSG0000 | -2.6921 | -6.46254 | 1.35E-08 | 2.33E-07 | LINC01224 |
| ENSG0000 | -4.79162 | -27.6962 | 0.011863 | 0.04952 | None |
| ENSG0000 | -3.0101 | -8.05622 | 0.008783 | 0.038542 | None |
| ENSG0000 | -2.27205 | -4.8301 | 0.003065 | 0.01605 | None |
| ENSG0000 | -1.7568 | -3.37947 | 8.23E-14 | 2.72E-12 | |
| ENSG0000 | -5.78756 | -55.2371 | 0.005917 | 0.027772 | None |
| ENSG0000 | -1.46632 | -2.76316 | 2.63E-06 | 3.05E-05 | LINC01235 |
| ENSG0000 | -1.11215 | -2.16168 | 0.007472 | 0.033678 | |
| ENSG0000 | -2.36942 | -5.16734 | 2.34E-06 | 2.73E-05 | GAS2L2 |
| ENSG0000 | -1.03702 | -2.05198 | 0.001905 | 0.010709 | None |
| ENSG0000 | -6.01835 | -64.819 | 0.000329 | 0.002325 | None |
| ENSG0000 | -6.02265 | -65.0126 | 0.002162 | 0.011919 | None |
| ENSG0000 | -2.11832 | -4.34189 | 0.006495 | 0.030094 | None |
| ENSG0000 | -6.6618 | -101.251 | 9.78E-05 | 0.000797 | |
| ENSG0000 | -5.77935 | -54.9233 | 0.000675 | 0.004365 | None |
| ENSG0000 | -1.55623 | -2.94085 | 1.93E-06 | 2.29E-05 | C2orf15 |
| ENSG0000 | -6.21327 | -74.196 | 0.000283 | 0.002041 | None |
| ENSG0000 | -3.94353 | -15.3859 | 0.008427 | 0.037238 | |
| ENSG0000 | -2.02645 | -4.07402 | 0.00827 | 0.036702 | None |
| ENSG0000 | -6.796 | -111.122 | 5.96E-05 | 0.000512 | None |
| ENSG0000 | -3.79521 | -13.8827 | 2.45E-10 | 5.44E-09 | DGKK |
| ENSG0000 | -5.80283 | -55.8246 | 0.002372 | 0.012914 | None |
| ENSG0000 | -1.79146 | -3.46165 | 0.000297 | 0.00213 | None |
| ENSG0000 | -6.52713 | -92.2279 | 0.000263 | 0.001909 | None |
| ENSG0000 | -5.5241 | -46.0172 | 0.00416 | 0.020674 | None |

| | | | | | |
|---|---|---|---|---|---|
| ENSG0000 | -1.54149 | -2.91095 | 2.92E-10 | 6.42E-09 | FLJ16779 |
| ENSG0000 | -4.53083 | -23.1161 | 0.003275 | 0.016944 | MIR6505 |
| ENSG0000 | -1.16538 | -2.24293 | 0.005317 | 0.025404 | UHRF1 |
| ENSG0000 | -2.75611 | -6.75573 | 1.52E-06 | 1.84E-05 | THBS1-IT1 |
| ENSG0000 | -1.48798 | -2.80496 | 1.02E-05 | 0.000104 | None |
| ENSG0000 | -5.7876 | -55.2386 | 0.001488 | 0.008713 | LINC01632 |
| ENSG0000 | -1.31078 | -2.48075 | 0.002695 | 0.014409 | None |
| ENSG0000 | -5.77105 | -54.6085 | 0.010716 | 0.04559 | None |
| ENSG0000 | -5.1983 | -36.715 | 0.003878 | 0.019525 | PTPN20CP |
| ENSG0000 | -2.92731 | -7.60691 | 0.000223 | 0.001649 | THBS1-AS1 |
| ENSG0000 | -3.00025 | -8.00137 | 0.001306 | 0.007802 | None |
| ENSG0000 | -5.78029 | -54.9591 | 0.000142 | 0.001113 | BANCR |
| ENSG0000 | -1.20673 | -2.30813 | 0.004341 | 0.021449 | |
| ENSG0000 | -1.22187 | -2.33248 | 0.001366 | 0.008109 | None |
| ENSG0000 | -1.58921 | -3.00884 | 2.85E-10 | 6.28E-09 | None |
| ENSG0000 | -2.00609 | -4.01692 | 1.02E-07 | 1.52E-06 | None |
| ENSG0000 | -4.54513 | -23.3464 | 0.000429 | 0.002934 | None |
| ENSG0000 | -6.35917 | -82.0921 | 7.77E-05 | 0.000649 | None |
| ENSG0000 | -5.78793 | -55.251 | 0.000661 | 0.004283 | None |
| ENSG0000 | -1.07409 | -2.1054 | 0.007695 | 0.034542 | None |
| ENSG0000 | -2.52731 | -5.76497 | 4.75E-30 | 4.94E-28 | None |
| ENSG0000 | -1.19052 | -2.28236 | 7.55E-06 | 7.92E-05 | None |
| ENSG0000 | -1.98855 | -3.96837 | 2.20E-28 | 2.05E-26 | None |
| ENSG0000 | -2.19988 | -4.59442 | 4.69E-08 | 7.42E-07 | None |
| ENSG0000 | -1.5119 | -2.85185 | 0.005304 | 0.025361 | None |
| ENSG0000 | -4.05863 | -16.6637 | 4.44E-85 | 1.86E-82 | None |
| ENSG0000 | -1.32489 | -2.50513 | 5.21E-15 | 1.96E-13 | |
| ENSG0000 | -5.78216 | -55.0305 | 0.010404 | 0.044536 | None |
| ENSG0000 | -1.03148 | -2.04412 | 0.000344 | 0.002419 | |
| ENSG0000 | -5.52896 | -46.1726 | 0.001699 | 0.009728 | None |
| ENSG0000 | -5.52648 | -46.0932 | 0.003221 | 0.016711 | None |
| ENSG0000 | -4.78782 | -27.6234 | 0.004727 | 0.023013 | |
| ENSG0000 | -1.93428 | -3.82188 | 5.51E-08 | 8.60E-07 | None |
| ENSG0000 | -1.20576 | -2.30658 | 1.78E-06 | 2.13E-05 | PEG13 |
| ENSG0000 | -1.06701 | -2.09509 | 0.001552 | 0.009019 | C1orf232 |
| ENSG0000 | -1.72733 | -3.31115 | 0.002238 | 0.01229 | None |
| ENSG0000 | -6.00318 | -64.1414 | 0.00051 | 0.003426 | U6 |
| ENSG0000 | -2.45821 | -5.49536 | 0.001459 | 0.008562 | Metazoa_SRP |
| ENSG0000 | -3.31237 | -9.93399 | 0.00136 | 0.008077 | LINC01902 |
| ENSG0000 | -5.77808 | -54.8752 | 0.003336 | 0.017207 | TCAF2C |

| GeneID | Treatment | Treatment | Treatment | Treatment | name | Astrocytes_GCSi_FC+2_adj0.05_14 |
|---|---|---|---|---|---|---|
| ENSG0000 | 1.100447 | 2.144212 | 3.33E-15 | 7.66E-13 | TNFRSF12A | |
| ENSG0000 | 1.21005 | 2.313457 | 7.58E-05 | 0.000986 | BIRC3 | |
| ENSG0000 | 1.090425 | 2.129368 | 1.54E-07 | 4.95E-06 | PTPRN | |
| ENSG0000 | -1.03027 | -2.0424 | 1.24E-09 | 7.40E-08 | TRAF1 | |
| ENSG0000 | 1.506003 | 2.840221 | 4.20E-21 | 3.45E-18 | SREBF1 | |
| ENSG0000 | 1.054077 | 2.076389 | 0.005059 | 0.029193 | ATP8B1 | |
| ENSG0000 | 1.054422 | 2.076886 | 9.68E-06 | 0.000175 | MPP4 | |
| ENSG0000 | 1.513387 | 2.854795 | 1.78E-10 | 1.33E-08 | ICAM1 | |
| ENSG0000 | 2.469383 | 5.538069 | 1.12E-07 | 3.74E-06 | IL11 | |
| ENSG0000 | 1.577564 | 2.984654 | 2.87E-29 | 1.64E-25 | SCD | |
| ENSG0000 | 1.168175 | 2.247272 | 6.10E-10 | 4.04E-08 | RELB | |
| ENSG0000 | 1.317504 | 2.492345 | 0.006039 | 0.033622 | ICAM5 | |
| ENSG0000 | 1.244811 | 2.369875 | 1.15E-17 | 4.31E-15 | TFPI2 | |
| ENSG0000 | -1.01469 | -2.02047 | 4.21E-05 | 0.0006 | DNAH11 | |
| ENSG0000 | 1.788596 | 3.454786 | 5.72E-18 | 2.41E-15 | SERPINE1 | |
| ENSG0000 | 1.150505 | 2.219916 | 1.64E-06 | 3.79E-05 | ACTA2 | |
| ENSG0000 | 1.281685 | 2.431228 | 2.87E-06 | 6.17E-05 | DKK1 | |
| ENSG0000 | 1.37118 | 2.586821 | 6.70E-14 | 1.07E-11 | CCL2 | |
| ENSG0000 | 1.379297 | 2.601416 | 0.001467 | 0.011244 | VWF | |
| ENSG0000 | -1.03753 | -2.05272 | 3.35E-23 | 5.27E-20 | RSPH4A | |
| ENSG0000 | -1.02953 | -2.04135 | 0.002622 | 0.017588 | SERPINI2 | |
| ENSG0000 | 1.178138 | 2.262845 | 2.80E-09 | 1.51E-07 | IL1R1 | |
| ENSG0000 | 1.175926 | 2.259379 | 7.21E-22 | 7.33E-19 | GADD45A | |
| ENSG0000 | 2.174913 | 4.515585 | 1.47E-18 | 6.85E-16 | RGS4 | |
| ENSG0000 | 1.311658 | 2.482267 | 1.24E-20 | 8.60E-18 | TNFAIP3 | |
| ENSG0000 | 1.363292 | 2.572716 | 8.93E-21 | 6.88E-18 | SGK1 | |
| ENSG0000 | 1.105419 | 2.151613 | 1.58E-05 | 0.000264 | CCN2 | |
| ENSG0000 | 1.18038 | 2.266365 | 1.96E-08 | 8.34E-07 | DUSP1 | |
| ENSG0000 | 1.604952 | 3.041856 | 2.57E-16 | 7.29E-14 | EGR1 | |
| ENSG0000 | 1.592829 | 3.016404 | 3.79E-29 | 1.64E-25 | DUSP4 | |
| ENSG0000 | 1.222835 | 2.334049 | 2.03E-06 | 4.55E-05 | SV2C | |
| ENSG0000 | 1.904321 | 3.743327 | 7.81E-22 | 7.50E-19 | EGR2 | |
| ENSG0000 | -1.01215 | -2.01691 | 1.85E-05 | 0.000301 | MATN4 | |
| ENSG0000 | 1.027337 | 2.038259 | 9.83E-05 | 0.001219 | CHST8 | |
| ENSG0000 | 1.05892 | 2.083371 | 0.000127 | 0.001505 | GRPR | |
| ENSG0000 | 1.258038 | 2.391703 | 8.05E-11 | 6.63E-09 | WNK4 | |
| ENSG0000 | 1.214656 | 2.320854 | 2.44E-14 | 4.54E-12 | LIF | |
| ENSG0000 | 1.791326 | 3.461329 | 1.05E-13 | 1.63E-11 | CPA4 | |
| ENSG0000 | 1.556878 | 2.942165 | 6.80E-10 | 4.47E-08 | VGF | |
| ENSG0000 | -1.02068 | -2.02888 | 0.001378 | 0.010674 | PRRG3 | |
| ENSG0000 | 1.098386 | 2.14115 | 0.002031 | 0.014429 | NALF2 | |
| ENSG0000 | 1.019768 | 2.027593 | 5.49E-14 | 8.95E-12 | GFPT2 | |
| ENSG0000 | 1.492986 | 2.81471 | 4.54E-23 | 6.54E-20 | CHI3L1 | |
| ENSG0000 | 2.061046 | 4.172888 | 3.71E-18 | 1.60E-15 | SPOCD1 | |
| ENSG0000 | 1.002531 | 2.003512 | 1.44E-11 | 1.38E-09 | ETS1 | |
| ENSG0000 | 2.087594 | 4.250386 | 4.31E-06 | 8.69E-05 | EGR4 | |
| ENSG0000 | -1.03566 | -2.05005 | 0.002044 | 0.014497 | PCDH8 | |
| ENSG0000 | 2.089025 | 4.254603 | 1.11E-06 | 2.73E-05 | GPNMB | |
| ENSG0000 | -1.03796 | -2.05332 | 2.96E-10 | 2.08E-08 | WDR38 | |

| | | | | | |
|---|---|---|---|---|---|
| ENSG0000 | 1.135619 | 2.197128 | 1.33E-15 | 3.27E-13 | IER3 |
| ENSG0000 | -1.01524 | -2.02124 | 8.68E-10 | 5.50E-08 | CFAP300 |
| ENSG0000 | 1.160816 | 2.235839 | 1.58E-10 | 1.21E-08 | THBS1 |
| ENSG0000 | 1.001483 | 2.002057 | 4.16E-08 | 1.59E-06 | DUSP6 |
| ENSG0000 | -1.02859 | -2.04002 | 6.51E-11 | 5.51E-09 | LRRC46 |
| ENSG0000 | 1.581609 | 2.993036 | 0.007304 | 0.038723 | PMAIP1 |
| ENSG0000 | 1.041503 | 2.058371 | 5.87E-09 | 2.82E-07 | CCN1 |
| ENSG0000 | 1.034074 | 2.047799 | 0.001483 | 0.011334 | KCNH1 |
| ENSG0000 | -1.01374 | -2.01913 | 0.001659 | 0.012343 | RNF175 |
| ENSG0000 | 1.028834 | 2.040374 | 1.69E-21 | 1.54E-18 | CDKN2B |
| ENSG0000 | 1.059155 | 2.08371 | 2.19E-13 | 3.15E-11 | ADAM12 |
| ENSG0000 | 1.235243 | 2.35421 | 0.000102 | 0.001254 | P4HA3 |
| ENSG0000 | -1.00083 | -2.00115 | 4.77E-08 | 1.78E-06 | ADAM33 |
| ENSG0000 | 1.113192 | 2.163237 | 2.77E-05 | 0.000424 | TAGLN |
| ENSG0000 | 1.202326 | 2.301103 | 5.71E-07 | 1.54E-05 | IL18 |
| ENSG0000 | -1.0317 | -2.04443 | 9.21E-17 | 2.70E-14 | DYNLT5 |
| ENSG0000 | -1.04572 | -2.0644 | 2.34E-11 | 2.14E-09 | PITPNC1 |
| ENSG0000 | -1.01438 | -2.02004 | 0.00401 | 0.024367 | RSPH10B |
| ENSG0000 | -1.04869 | -2.06864 | 2.58E-18 | 1.15E-15 | FBXO32 |
| ENSG0000 | 1.116232 | 2.167801 | 4.59E-13 | 5.83E-11 | NRG1 |
| ENSG0000 | -1.03643 | -2.05115 | 0.007455 | 0.039315 | XDH |
| ENSG0000 | -1.04234 | -2.05956 | 0.000113 | 0.001369 | CATIP |
| ENSG0000 | -1.00875 | -2.01217 | 1.02E-11 | 1.01E-09 | None |
| ENSG0000 | 1.002789 | 2.003871 | 1.76E-09 | 1.01E-07 | ABCG1 |
| ENSG0000 | 1.058909 | 2.083355 | 0.001467 | 0.011246 | GAB3 |
| ENSG0000 | 1.432644 | 2.699409 | 0.000307 | 0.003174 | ICOSLG |
| ENSG0000 | -1.04614 | -2.065 | 5.41E-05 | 0.00074 | LRRC56 |
| ENSG0000 | 1.031216 | 2.043746 | 8.93E-12 | 8.98E-10 | ATF3 |
| ENSG0000 | 1.024646 | 2.03446 | 9.53E-06 | 0.000173 | KCNF1 |
| ENSG0000 | -1.01689 | -2.02356 | 0.003536 | 0.022091 | DCST2 |
| ENSG0000 | 1.133951 | 2.19459 | 8.00E-05 | 0.00103 | CDCP1 |
| ENSG0000 | 1.274639 | 2.419383 | 1.42E-12 | 1.62E-10 | DLC1 |
| ENSG0000 | -1.03603 | -2.05058 | 0.003318 | 0.021062 | PDZRN4 |
| ENSG0000 | 1.200072 | 2.297511 | 0.002238 | 0.015574 | MMP10 |
| ENSG0000 | -1.04928 | -2.0695 | 2.05E-05 | 0.000328 | LGI3 |
| ENSG0000 | 1.322673 | 2.501292 | 1.10E-12 | 1.28E-10 | STXBP6 |
| ENSG0000 | 1.180722 | 2.266902 | 5.57E-07 | 1.51E-05 | VXN |
| ENSG0000 | 1.082118 | 2.117142 | 8.33E-12 | 8.42E-10 | EFNA1 |
| ENSG0000 | 2.035768 | 4.10041 | 0.002274 | 0.015767 | CXCL8 |
| ENSG0000 | 1.141278 | 2.205763 | 5.84E-13 | 7.21E-11 | FOS |
| ENSG0000 | 1.128634 | 2.186516 | 1.38E-12 | 1.57E-10 | METTL7B |
| ENSG0000 | 1.260301 | 2.395457 | 5.18E-17 | 1.63E-14 | JUNB |
| ENSG0000 | 1.396293 | 2.632244 | 0.000335 | 0.003389 | CHRNA9 |
| ENSG0000 | 1.622283 | 3.078618 | 2.35E-06 | 5.19E-05 | CMKLR1 |
| ENSG0000 | 1.129631 | 2.188027 | 4.63E-05 | 0.00065 | ASPHD1 |
| ENSG0000 | -1.00497 | -2.00691 | 0.00285 | 0.018767 | GRAMD2A |
| ENSG0000 | 1.277117 | 2.423542 | 1.03E-28 | 3.55E-25 | LPL |
| ENSG0000 | 1.457078 | 2.745517 | 5.52E-17 | 1.70E-14 | CLCF1 |
| ENSG0000 | 1.893023 | 3.714125 | 8.69E-18 | 3.34E-15 | FOSL1 |
| ENSG0000 | 1.159949 | 2.234495 | 0.005915 | 0.033064 | SPHK1 |

| | | | | | |
|---|---|---|---|---|---|
| ENSG0000 | 1.04107 | 2.057753 | 2.42E-14 | 4.54E-12 | MYO1D |
| ENSG0000 | 1.380439 | 2.603476 | 1.54E-11 | 1.46E-09 | BDNF |
| ENSG0000 | 1.039736 | 2.055851 | 0.000131 | 0.001559 | GCNT4 |
| ENSG0000 | 1.594231 | 3.019334 | 2.64E-25 | 5.69E-22 | EGR3 |
| ENSG0000 | -1.00589 | -2.00818 | 2.88E-14 | 5.24E-12 | SLC17A8 |
| ENSG0000 | -1.03523 | -2.04944 | 0.008579 | 0.043848 | PCED1B |
| ENSG0000 | 1.873076 | 3.663128 | 0.003472 | 0.021793 | C3orf80 |
| ENSG0000 | 1.052581 | 2.074238 | 6.02E-13 | 7.38E-11 | ARSJ |
| ENSG0000 | 1.10302 | 2.148038 | 1.33E-08 | 5.95E-07 | GPR3 |
| ENSG0000 | -1.02129 | -2.02973 | 2.26E-07 | 6.98E-06 | HHIPL1 |
| ENSG0000 | 1.12602 | 2.182559 | 0.000161 | 0.001838 | TNFAIP8L3 |
| ENSG0000 | -1.04959 | -2.06994 | 3.70E-09 | 1.92E-07 | KCNQ3 |
| ENSG0000 | 1.282488 | 2.432581 | 4.35E-12 | 4.61E-10 | MAFF |
| ENSG0000 | 1.198987 | 2.295784 | 9.65E-07 | 2.43E-05 | MYBL1 |
| ENSG0000 | 1.341858 | 2.534776 | 2.94E-14 | 5.29E-12 | SPRY4 |
| ENSG0000 | 1.272116 | 2.415156 | 4.05E-16 | 1.09E-13 | SPRED3 |
| ENSG0000 | 1.666634 | 3.174731 | 7.44E-05 | 0.000971 | SH2D5 |
| ENSG0000 | -1.05398 | -2.07625 | 0.008564 | 0.043787 | SEMA4A |
| ENSG0000 | 1.024183 | 2.033807 | 4.42E-09 | 2.21E-07 | HRH1 |
| ENSG0000 | -1.02625 | -2.03672 | 0.000565 | 0.005201 | DTHD1 |
| ENSG0000 | 2.377378 | 5.195917 | 0.000228 | 0.002472 | PLN |
| ENSG0000 | 1.9691 | 3.915238 | 7.46E-30 | 6.45E-26 | ARC |
| ENSG0000 | -1.01351 | -2.01882 | 3.89E-09 | 1.99E-07 | NYNRIN |
| ENSG0000 | -1.03489 | -2.04896 | 0.00515 | 0.029646 | MIR149 |
| ENSG0000 | -1.05165 | -2.0729 | 0.000186 | 0.002085 | COL28A1 |
| ENSG0000 | -1.046 | -2.06479 | 0.000196 | 0.002174 | None |
| ENSG0000 | 1.586721 | 3.003658 | 1.65E-06 | 3.79E-05 | ZNF812P |
| ENSG0000 | 1.125619 | 2.181951 | 0.008118 | 0.042 | None |
| ENSG0000 | 1.022212 | 2.031031 | 0.004251 | 0.025504 | None |
| ENSG0000 | 1.721197 | 3.297098 | 0.000144 | 0.001674 | MYOSLID |
| ENSG0000 | -1.01647 | -2.02296 | 9.03E-07 | 2.29E-05 | TGFB2-AS1 |
| ENSG0000 | 1.488611 | 2.806186 | 2.41E-05 | 0.000377 | LINC00707 |
| ENSG0000 | 1.391119 | 2.62282 | 2.10E-07 | 6.53E-06 | |
| ENSG0000 | 1.323907 | 2.503432 | 1.61E-06 | 3.72E-05 | GSTA1 |
| ENSG0000 | -1.00065 | -2.0009 | 1.65E-05 | 0.000273 | USP2-AS1 |
| ENSG0000 | -1.04824 | -2.06801 | 0.004166 | 0.02506 | ALG1L9P |
| ENSG0000 | -1.05083 | -2.07172 | 3.45E-05 | 0.000505 | C8orf34-AS1 |
| ENSG0000 | -1.01494 | -2.02082 | 0.00242 | 0.016508 | TICAM2-AS1 |
| ENSG0000 | 1.091105 | 2.130371 | 0.001016 | 0.008396 | LINC02732 |
| ENSG0000 | 1.080583 | 2.11489 | 1.22E-06 | 2.96E-05 | None |
| ENSG0000 | -1.00758 | -2.01054 | 4.56E-08 | 1.71E-06 | TRIL |
| ENSG0000 | 1.733677 | 3.325743 | 0.003541 | 0.022109 | PPP1R14B-AS1 |
| ENSG0000 | 1.833678 | 3.564447 | 0.001321 | 0.010354 | None |
| ENSG0000 | -1.01303 | -2.01815 | 8.61E-07 | 2.21E-05 | LINC01579 |
| ENSG0000 | -1.00328 | -2.00455 | 8.64E-05 | 0.0011 | TSPAN5-DT |
| ENSG0000 | 1.678717 | 3.201432 | 3.12E-06 | 6.60E-05 | GJA5 |
| ENSG0000 | 1.039005 | 2.05481 | 8.40E-06 | 0.000156 | None |
| ENSG0000 | 2.270464 | 4.824784 | 0.000171 | 0.001936 | None |
| ENSG0000 | 1.158074 | 2.231594 | 1.36E-09 | 8.06E-08 | None |
