## Supplementary material for "Unravelling neuronal and glial differences in ceramide composition, synthesis, and sensitivity to toxicity": Venn_List_FB1 Astrocyte Microglia common gene changes

| GeneID | name |
|---|---|
| ENSG0000 | IL32 |
| ENSG0000 | CP |
| ENSG0000 | MSMO1 |
| ENSG0000 | CCDC85A |
| ENSG0000 | DMRT3 |
| ENSG0000 | NGFR |
| ENSG0000 | MAOB |
| ENSG0000 | PANX2 |
| ENSG0000 | EDN1 |
| ENSG0000 | BRINP1 |
| ENSG0000 | SEMA5B |
| ENSG0000 | GCKR |
| ENSG0000 | NKAIN4 |
| ENSG0000 | RS1 |
| ENSG0000 | KLC3 |
| ENSG0000 | TFPI2 |
| ENSG0000 | NPTX2 |
| ENSG0000 | AREG |
| ENSG0000 | CRYAB |
| ENSG0000 | MAK |
| ENSG0000 | HMGCS1 |
| ENSG0000 | STC2 |
| ENSG0000 | DNAH6 |
| ENSG0000 | RGS4 |
| ENSG0000 | CCN2 |
| ENSG0000 | TEX11 |
| ENSG0000 | TWIST1 |
| ENSG0000 | PLP1 |
| ENSG0000 | MYH2 |
| ENSG0000 | FOSB |
| ENSG0000 | ADM2 |
| ENSG0000 | CPA4 |
| ENSG0000 | FOXP2 |
| ENSG0000 | CHAC1 |
| ENSG0000 | RNASE1 |
| ENSG0000 | FGF13 |
| ENSG0000 | NALF2 |
| ENSG0000 | KLHDC7B |
| ENSG0000 | BEX2 |
| ENSG0000 | HRH4 |
| ENSG0000 | ADAM19 |
| ENSG0000 | ALDH1L2 |
| ENSG0000 | GALNT5 |
| ENSG0000 | SCN7A |
| ENSG0000 | AK7 |
| ENSG0000 | CBLN2 |

ENSG0000 BEST4
ENSG0000 NYAP2
ENSG0000 TMEM108
ENSG0000 LIX1
ENSG0000 CXCL14
ENSG0000 LCN9
ENSG0000 SLC39A12
ENSG0000 ADGRL3
ENSG0000 SLC7A11
ENSG0000 ADAMTS12
ENSG0000 GABRA2
ENSG0000 DNAAF1
ENSG0000 NMNAT2
ENSG0000 STEAP2
ENSG0000 CELF3
ENSG0000 ISL2
ENSG0000 ADAMTS9
ENSG0000 CFAP100
ENSG0000 DACT2
ENSG0000 CSMD3
ENSG0000 UNCX
ENSG0000 CFAP47
ENSG0000 PCDH19
ENSG0000 IGF2
ENSG0000 SCARA3
ENSG0000 KCNJ4
ENSG0000 NMRAL2P
ENSG0000 ISG20
ENSG0000 MRGPRF
ENSG0000 DOK7
ENSG0000 NUPR1
ENSG0000 CRYBG2
ENSG0000 MAP3K15
ENSG0000 NME9
ENSG0000 CFAP65
ENSG0000 PAPPA
ENSG0000 NKX2-5
ENSG0000 CHST6
ENSG0000 SLC35F3
ENSG0000 ARSI
ENSG0000 TREML3P
ENSG0000 CLDN5
ENSG0000 CNTN2
ENSG0000 CCIN
ENSG00000186354
ENSG0000 None
ENSG0000 LHFPL3

ENSG0000 KRT18P59
ENSG0000 DNAJB13
ENSG0000 INSC
ENSG0000 AJAP1
ENSG0000 TPM2
ENSG0000 RNU1-72P
ENSG0000 IGBP1-AS1
ENSG0000 MIR616
ENSG0000 MT-TA
ENSG0000 None
ENSG0000 EEF1A1P29
ENSG0000 HMSD
ENSG0000 MTND6P21
ENSG0000 LINC01449
ENSG0000 OLFM5P
ENSG0000 PRR29
ENSG0000 None
ENSG0000 HSD11B1-AS1
ENSG0000 TEX46
ENSG0000 None
ENSG0000 NEK2-DT
ENSG0000 TGFB2-AS1
ENSG0000 None
ENSG0000 None
ENSG0000 INMT
ENSG0000 CFAP57
ENSG0000 CYP2U1-AS1
ENSG0000 RGMB-AS1
ENSG0000 LINC02485
ENSG0000 FOXD1
ENSG0000 None
ENSG0000 LINC02365
ENSG0000 MADD-AS1
ENSG0000 CHMP4BP1
ENSG0000 None
ENSG0000 None
ENSG0000 None
ENSG0000 PTGES3L
ENSG0000 GABRQ
ENSG00000270246
ENSG0000 NAV2-AS6
ENSG0000 RASL10B
ENSG0000 TP53RK-DT
ENSG0000 None
ENSG0000 None
ENSG00000273301
ENSG0000 None

ENSG0000 None
ENSG00000280237
ENSG0000 LINC01902

| GeneID | name | Astrocytes and microglia common decreased genes with FB1 |
|---|---|---|
| ENSG0000 | MSMO1 | |
| ENSG0000 | MAOB | |
| ENSG0000 | MAK | |
| ENSG0000 | HMGCS1 | |
| ENSG0000 | DNAH6 | |
| ENSG0000 | RGS4 | |
| ENSG0000 | MYH2 | |
| ENSG0000 | RNASE1 | |
| ENSG0000 | ADAM19 | |
| ENSG0000 | GALNT5 | |
| ENSG0000 | AK7 | |
| ENSG0000 | CBLN2 | |
| ENSG0000 | NYAP2 | |
| ENSG0000 | TMEM108 | |
| ENSG0000 | CXCL14 | |
| ENSG0000 | ADAMTS12 | |
| ENSG0000 | CFAP47 | |
| ENSG0000 | MRGPRF | |
| ENSG0000 | CFAP65 | |
| ENSG0000 | SLC35F3 | |
| ENSG0000 | ARSI | |
| ENSG0000 | CLDN5 | |
| ENSG0000 | CNTN2 | |
| ENSG0000 | CCIN | |
| ENSG0000 | None | |
| ENSG0000 | LHFPL3 | |
| ENSG0000 | DNAJB13 | |
| ENSG0000 | TPM2 | |
| ENSG0000 | HMSD | |
| ENSG0000 | LINC01449 | |
| ENSG0000 | PRR29 | |
| ENSG0000 | None | |
| ENSG0000 | NEK2-DT | |
| ENSG0000 | TGFB2-AS1 | |
| ENSG0000 | INMT | |
| ENSG0000 | CYP2U1-AS1 | |
| ENSG0000 | None | |
| ENSG0000 | PTGES3L | |
| ENSG00000270246 | | |
| ENSG00000273301 | | |
| ENSG0000 | LINC01902 | |

| GeneID | name | Astrocytes and microglia common increased genes with FB1 |
|---|---|---|
| ENSG0000 | DMRT3 | |
| ENSG0000 | PANX2 | |
| ENSG0000 | GCKR | |
| ENSG0000 | AREG | |
| ENSG0000 | STC2 | |
| ENSG0000 | TEX11 | |
| ENSG0000 | TWIST1 | |
| ENSG0000 | ADM2 | |
| ENSG0000 | FOXP2 | |
| ENSG0000 | CHAC1 | |
| ENSG0000 | NALF2 | |
| ENSG0000 | KLHDC7B | |
| ENSG0000 | BEX2 | |
| ENSG0000 | ALDH1L2 | |
| ENSG0000 | SLC39A12 | |
| ENSG0000 | SLC7A11 | |
| ENSG0000 | GABRA2 | |
| ENSG0000 | NMNAT2 | |
| ENSG0000 | ISL2 | |
| ENSG0000 | PCDH19 | |
| ENSG0000 | KCNJ4 | |
| ENSG0000 | NMRAL2P | |
| ENSG0000 | ISG20 | |
| ENSG0000 | NUPR1 | |
| ENSG0000 | CRYBG2 | |
| ENSG0000 | PAPPA | |
| ENSG0000 | NKX2-5 | |
| ENSG0000 | TREML3P | |
| ENSG0000 | MIR616 | |
| ENSG0000 | MTND6P21 | |
| ENSG0000 | None | |
| ENSG0000 | FOXD1 | |
| ENSG0000 | MADD-AS1 | |
| ENSG0000 | None | |
| ENSG0000 | NAV2-AS6 | |
| ENSG0000 | RASL10B | |
| ENSG0000 | TP53RK-DT | |
| ENSG0000 | None | |
