## Supplementary material for "Unravelling neuronal and glial differences in ceramide composition, synthesis, and sensitivity to toxicity": Microglia FB1 and GCSi gene changes

| GeneID | name | Microglia FB1 gene changes |
| --- | --- | --- |
| ENSG0000 | MYH16 | |
| ENSG0000 | HGF | |
| ENSG0000 | INSRR | |
| ENSG0000 | TNC | |
| ENSG0000 | DKK3 | |
| ENSG0000 | NGFR | |
| ENSG0000 | GLP2R | |
| ENSG0000 | SCT | |
| ENSG0000 | FNDC8 | |
| ENSG0000 | CHGB | |
| ENSG0000 | IL11 | |
| ENSG0000 | NTSR1 | |
| ENSG0000 | WFDC2 | |
| ENSG0000 | RS1 | |
| ENSG0000 | CORO2B | |
| ENSG0000 | TFPI2 | |
| ENSG0000 | CHCHD2 | |
| ENSG0000 | KCNT1 | |
| ENSG0000 | AREG | |
| ENSG0000 | NKX3-2 | |
| ENSG0000 | CRYAB | |
| ENSG0000 | B3GAT1 | |
| ENSG0000 | C3orf52 | |
| ENSG0000 | ADGRL2 | |
| ENSG0000 | GRIA2 | |
| ENSG0000 | ADGRB2 | |
| ENSG0000 | TMEM54 | |
| ENSG0000 | CXCR4 | |
| ENSG0000 | TWIST1 | |
| ENSG0000 | PLP1 | |
| ENSG0000 | PI3 | |
| ENSG0000 | SLPI | |
| ENSG0000 | F13A1 | |
| ENSG0000 | EREG | |
| ENSG0000 | IL1B | |
| ENSG0000 | FOSB | |
| ENSG0000 | FOXA2 | |
| ENSG0000 | CCR7 | |
| ENSG0000 | ISLR | |
| ENSG0000 | LAMA5 | |
| ENSG0000 | TEX101 | |
| ENSG0000 | GFPT2 | |
| ENSG0000 | LYVE1 | |
| ENSG0000 | NT5E | |
| ENSG0000 | SCN7A | |
| ENSG0000 | SLC28A2 | |

ENSG0000 ARHGAP29
ENSG0000 LHCGR
ENSG0000 AK7
ENSG0000 ABCC12
ENSG0000 GATA6
ENSG0000 CBLN2
ENSG0000 TINAGL1
ENSG0000 ANKRD1
ENSG0000 HMGA2
ENSG0000 MMP3
ENSG0000 ADGRL3
ENSG0000 FREM2
ENSG0000 SLC7A11
ENSG0000 ENKUR
ENSG0000 BMP6
ENSG0000 MMP21
ENSG0000 KCNS2
ENSG0000 NMNAT2
ENSG0000 SLC34A2
ENSG0000 LAD1
ENSG0000 STC1
ENSG0000 LRRTM1
ENSG0000 ADAMTS9
ENSG0000 CXCL3
ENSG0000 CXCL5
ENSG0000 HTR4
ENSG0000 DACT2
ENSG0000 SMAD5-AS1
ENSG0000 STEAP1
ENSG0000 IGF2
ENSG0000 DEGS2
ENSG0000 TM4SF1
ENSG0000 HSD17B13
ENSG0000 HTRA3
ENSG0000 PURG
ENSG0000 COL6A5
ENSG0000 SULT1B1
ENSG0000 SLCO4C1
ENSG0000 C9orf131
ENSG0000 CHST1
ENSG0000 None
ENSG0000 CRYBG2
ENSG0000 FOXG1
ENSG0000 IRX5
ENSG0000 FCER1A
ENSG0000 ADGRD2
ENSG0000 CRIP2

ENSG0000 NKX2-5
ENSG0000 EPHA10
ENSG0000 CABCOCO1
ENSG0000 COL18A1-AS1
ENSG0000 RFLNB
ENSG0000 KIRREL1
ENSG0000 ARSI
ENSG0000 CLDN5
ENSG0000 LINC00313
ENSG0000 NOTUM
ENSG0000 NTF3
ENSG0000 SYN3
ENSG0000 MATN1-AS1
ENSG00000186354
ENSG0000 None
ENSG0000 C11orf96
ENSG0000 DPPA3
ENSG0000 C5orf52
ENSG0000 C9orf153
ENSG0000 NCCRP1
ENSG0000 RPSAP47
ENSG0000 SP6
ENSG0000 CLDN4
ENSG0000 SH2D5
ENSG0000 H4C3
ENSG0000 FAM177B
ENSG0000 ZNF560
ENSG0000 MYL4
ENSG0000 CSAG1
ENSG0000 SNORD104
ENSG0000 Y_RNA
ENSG0000 SNORA63D
ENSG0000 Y_RNA
ENSG0000 Y_RNA
ENSG0000 Y_RNA
ENSG0000 MIR645
ENSG0000 MT-TY
ENSG0000 MT-TS1
ENSG0000 MT-TG
ENSG0000 MT-TH
ENSG0000 MT-TL2
ENSG0000 TRGV1
ENSG0000 SNORA3B
ENSG0000 PDCL3P5
ENSG0000 ARHGEF35
ENSG0000 MAGEA12
ENSG0000 None

| | |
|---|---|
| ENSG0000 | MARCKSL1P2 |
| ENSG0000 | RPL31P52 |
| ENSG0000 | CATSPERZ |
| ENSG0000 | TAFA5 |
| ENSG0000 | None |
| ENSG0000 | None |
| ENSG0000 | LINC02860 |
| ENSG0000 | RNU4-86P |
| ENSG0000 | RN7SKP150 |
| ENSG0000 | None |
| ENSG0000 | None |
| ENSG0000 | ARHGEF2-AS1 |
| ENSG0000 | OLFM5P |
| ENSG0000 | ARL14EPP1 |
| ENSG0000 | LINC01733 |
| ENSG0000 | None |
| ENSG0000 | RPL22P3 |
| ENSG0000 | USF1P1 |
| ENSG0000 | SMCR5 |
| ENSG0000 | None |
| ENSG0000 | MTND5P1 |
| ENSG0000 | None |
| ENSG0000 | None |
| ENSG0000 | None |
| ENSG0000 | HLA-DRB6 |
| ENSG0000 | None |
| ENSG0000 | None |
| ENSG0000 | None |
| ENSG0000 | STK24P1 |
| ENSG0000 | TRBV5-4 |
| ENSG0000 | None |
| ENSG0000 | SRGAP2-AS1 |
| ENSG0000 | TMEM114 |
| ENSG0000 | None |
| ENSG0000 | CRYZP1 |
| ENSG0000 | MROH3P |
| ENSG0000 | CCNQP1 |
| ENSG0000 | None |
| ENSG0000 | CSNK1G2P1 |
| ENSG0000 | None |
| ENSG0000 | None |
| ENSG0000 | NSRP1P1 |
| ENSG0000 | SNRPFP4 |
| ENSG0000 | None |
| ENSG0000 | COX7CP1 |
| ENSG0000 | EHMT2-AS1 |
| ENSG0000 | VDAC1P11 |

ENSG0000 RNU7-38P
ENSG0000 None
ENSG0000 None
ENSG0000 None
ENSG0000 RPSAP52
ENSG0000 None
ENSG0000 RPL12P21
ENSG0000 PDE6B-AS1
ENSG0000 None
ENSG0000 GATA2-AS1
ENSG0000 DUXAP10
ENSG0000 None
ENSG0000 RPL12P33
ENSG0000 None
ENSG0000 None
ENSG0000 RNU7-18P
ENSG0000 RNU4ATAC12P
ENSG0000 RNU6-703P
ENSG0000 USP12P1
ENSG0000 None
ENSG0000 None
ENSG0000 LINC02749
ENSG0000 None
ENSG0000 LINC02551
ENSG0000 DDX18P5
ENSG0000 None
ENSG0000 ELOCP31
ENSG0000 RPL7AP3
ENSG0000 LINC02454
ENSG0000 None
ENSG0000 None
ENSG0000 None
ENSG0000 None
ENSG0000 None
ENSG0000 None
ENSG0000 None
ENSG0000 GCSHP2
ENSG0000 None
ENSG0000 None
ENSG0000 ATP5MFP6
ENSG0000 TNRC6B-DT
ENSG0000 None
ENSG0000 GOLGA8T
ENSG0000 None
ENSG0000 None
ENSG0000 None
ENSG0000 SPON1

| | |
|---|---|
| ENSG0000 | MRPS21P9 |
| ENSG0000 | RYKP1 |
| ENSG0000 | ABHD17AP5 |
| ENSG0000 | MIR5188 |
| ENSG0000 | None |
| ENSG0000 | None |
| ENSG0000 | None |
| ENSG0000 | PTGES3L |
| ENSG0000 | KCNJ2-AS1 |
| ENSG0000 | None |
| ENSG0000 | LINC02073 |
| ENSG0000 | None |
| ENSG0000 | GABRQ |
| ENSG0000 | AIRN |
| ENSG0000 | None |
| ENSG0000 | None |
| ENSG0000 | None |
| ENSG0000 | None |
| ENSG0000 | None |
| ENSG0000 | None |
| ENSG0000 | MIR4787 |
| ENSG0000 | None |
| ENSG0000 | None |
| ENSG0000 | None |
| ENSG0000 | None |
| ENSG0000 | MIR7152 |
| ENSG0000 | None |
| ENSG0000 | H3C11 |
| ENSG0000 | OR7G15P |
| ENSG0000 | None |
| ENSG0000 | DACH1 |
| ENSG0000 | None |
| ENSG0000 | None |
| ENSG0000 | H3C7 |
| ENSG0000 | None |
| ENSG0000 | RN7SL113P |
| ENSG0000 | H4C1 |
| ENSG0000 | None |
| ENSG0000 | None |
| ENSG0000 | None |
| ENSG0000 | None |
| ENSG00000280237 | |
| ENSG0000 | None |
| ENSG0000 | LINC02452 |
